# Gasdermin E is Dispensable for H1N1 Influenza Virus Pathogenesis in Mice

**DOI:** 10.1101/2025.07.29.667514

**Authors:** Samuel Speaks, Jonathan Papa, Matthew McFadden, Jack E. Roettger, Benjamin D. Liu, Shreenath Mohan, Brendan M. Reznik, Steve Leumi, Jana M. Cable, Adriana Forero, Jacob S. Yount

## Abstract

Targeting cell death pathways, including pyroptosis and necroptosis, has been shown to mitigate influenza virus infection severity. Here, we examined whether pyroptosis specifically driven by the pore-forming protein gasdermin E (GSDME) is involved in regulating influenza virus infection outcomes. We found that *Gsdme^−/−^* mice showed similar weight loss and survival in severe A/PR/8/34 (H1N1) virus infections compared to WT counterparts. Likewise, lung dysfunction, histopathological damage, viral titers, and inflammatory cytokine levels were similar in the two groups. Global transcriptomic analysis also revealed similar inflammatory and antiviral gene expression programs in WT versus *Gsdme^−/−^* mouse lungs at baseline and in response to infection. To confirm the generality of these findings, we infected mice with 2009 pandemic H1N1 virus and again observed similar weight loss, mortality, and lung dysfunction in WT and *Gsdme^−/−^* mice. Our results overall demonstrate that GSDME contributes negligibly to the host response against H1N1 influenza virus, refining our understanding of cell death pathways in influenza pathogenesis.

## INTRODUCTION

Influenza virus remains a major global cause of hospitalization and death, causing severe illness in 3-5 million people and over 500,000 deaths annually^1^. While many influenza virus strains circulate in animal reservoirs, the primary culprits of severe human disease are influenza A virus (IAV) subtypes H1N1 and H3N2, and influenza B virus^2^. Severe influenza pathogenesis is driven in part by hyperactive host immune responses, which trigger excessive inflammation, tissue damage, and ultimately respiratory failure^3^. A better understanding of the host factors that contribute to this damaging inflammatory response is critical to improving outcomes in severe influenza.

Multiple forms of programmed cell death contribute to lung pathology during severe influenza virus infections, including apoptosis^4–6^, necroptosis^7^, and pyroptosis^8–11^. While each pathway can influence the balance between pathogen clearance and host-mediated immunopathology, pyroptosis has emerged as a particularly important determinant within this axis^12, 13^. This lytic form of cell death is mediated by members of the Gasdermin family of proteins, a group of pore-forming molecules that execute inflammatory cell death via the release of inflammatory mediators from the cytoplasm, ion gradient disruptions, and osmotic cell lysis^14–17^. Gasdermin D (GSDMD) and gasdermin E (GSDME) are among the best characterized of this pore-forming family of proteins, with both overlapping and unique functionality.

GSDMD and GSDME share a conserved structure including an N-terminal pore-forming domain and a C-terminal repressor domain^16, 18^. Upon proteolytic cleavage, the N-terminal fragment oligomerizes in the plasma membrane to form pores, leading to cell lysis and the release of inflammatory molecules such as IL-1β and IL-18^19–23^. Their upstream activators, however, are distinct. GSDMD is primarily cleaved by inflammatory caspases (1 and 11 in mice and 1, 4, and 5 in humans), whereas GSDME is activated by apoptotic caspase-3 and can redirect apoptotic cells toward lytic pyroptotic death^16^. Notably, we and others demonstrated that GSDMD exacerbates IAV-induced lung pathology by amplifying neutrophil-driven inflammation^24,25^. These findings underscore the pathological function of gasdermin-mediated pyroptosis in IAV infection.

In addition to the established role of GSDMD in driving IAV pathogenesis, recent studies have investigated whether GSDME contributes similarly to influenza severity. GSDME was found to promote caspase-3 mediated pyroptosis in human alveolar epithelial cells infected with highly pathogenic H7N9 IAV, which was consistent with decreased lung inflammation and lethality in *Gsdme^−/−^* mice infected with this virus^26^. Likewise, in an H3N2 IAV infection model, GSDME-deficient mice showed improved survival and reduced cytokine responses, suggesting that GSDME exacerbates inflammation and tissue damage during H3N2 IAV infection^27^.

Despite growing evidence for a role of GSDME in IAV pathogenesis, its contribution to H1N1 IAV pathology remains unexplored. H1N1 IAV has been responsible for a substantial portion of severe influenza cases in humans worldwide, highlighting the importance of understanding its disease pathogenesis^28^. Interestingly, *in vitro* studies have shown that H1N1 infection can induce GSDME cleavage in human epithelial cells^27, 29–31^, but whether GSDME contributes to morbidity or mortality *in vivo* has not been established.

Here, we investigated the role of GSDME in two distinct models of severe H1N1 influenza using *Gsdme^−/−^* mice. We comprehensively assessed disease through measurements of weight loss, lung function, lung histopathology, viral burden, inflammatory cytokines, and global lung transcriptomic changes. We found that GSDME deficiency resulted in negligible changes in inflammatory pathways and did not alter lung dysfunction, tissue pathology, weight loss, viral replication, or mortality. These findings stand in contrast to prior studies of H3N2 and H7N9 IAV and suggest that the contribution of GSDME to influenza pathogenesis is strain-specific and context-dependent.

## RESULTS

### GSDME does not impact morbidity and mortality in H1N1 IAV infection

To examine effects of GSDME on H1N1 IAV infection, we obtained *Gsdme^−/−^* mice and first confirmed loss of this protein in lung homogenates from these animals (**Fig 1A**). We infected WT and *Gsdme^−/−^* mice with 50 TCID50 of the influenza virus strain A/PR/8/34 (H1N1, known as PR8). This viral dose was chosen as it results in severe infection near the LD_50_ for C57BL/6 mice, allowing examination of whether loss of GSDME influences outcomes near this critical disease threshold. We then assessed lung viral titers at day 7 post-infection and observed that viral burdens were similar in WT and KO animals (**Fig 1B**). We also measured body weight loss throughout a time course of infection as an indicator of overall disease severity and observed no significant differences between genotypes (**Fig 1C**). In terms of survival, *Gsdme^−/−^* mice fared slightly better than WT, but the difference was not statistically significant (**Fig 1D**). Lastly, given the known role of GSDME in inflammatory cell death and release of inflammatory mediators, we measured levels of IL-1β, IL-18, TNF, IL-6, and IFNβ in lung homogenates at day 7 post-infection and found no significant differences (**Fig 1E)**. These data indicate that GSDME does not significantly influence disease severity or inflammatory responses in PR8 IAV infection.

**Figure 1:**
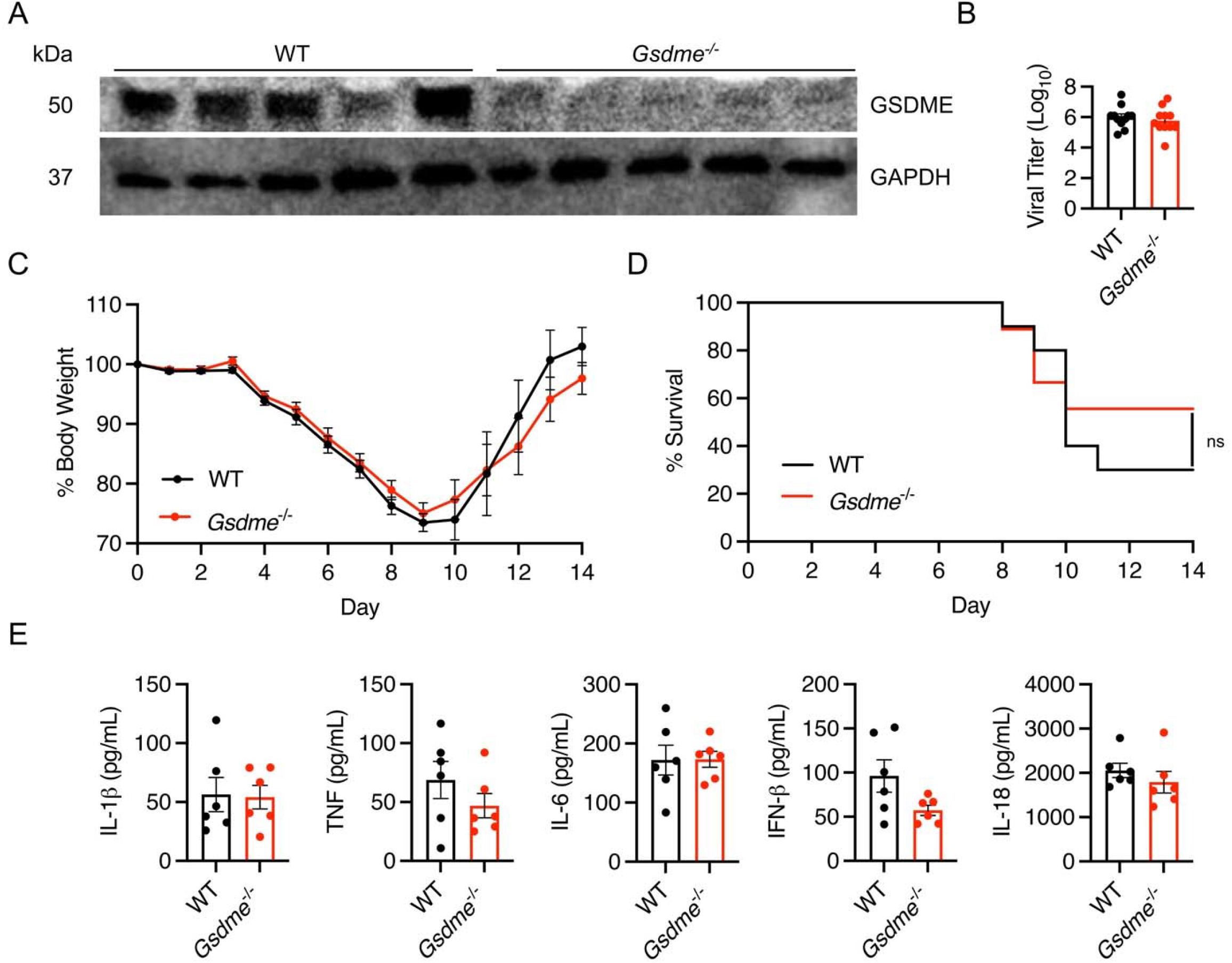
PR8 H1N1 IAV pathogenesis is similar in WT and *Gsdme*^−/−^ mice. A-E. WT and *Gsdme^−/−^* mice were infected with 50 TCID50 of PR8 intranasally. **A** Western blot of lung lysates. Each lane represents one individual mouse. **B** Viral titers of lung homogenates taken on day 7 post infection (WT n = 11, *Gsdme^−/−^* n = 11, error bars represent SEM, not significant by unpaired t-test). **C** Weight loss measurements (WT n = 20 and *Gsdme^−/−^* n = 17 for days 0-7, WT n = 10 and *Gsdme^−/−^* n = 9 for days 8-14, each dot is an average of individual mouse weights normalized to 100% relative to day 0, error bars indicate SEM, no significant differences between genotypes at any timepoint by two-way ANOVA followed by Bonferroni’s multiple comparisons test). **D** Survival analysis (WT n = 10, *Gsdme^−/−^* n = 9, non-significant by Log-rank Mantel-Cox test). **E** Cytokine quantifications on lung homogenates taken at day 7 post infection (WT n = 6, *Gsdme^−/−^* n = 6, error bars indicate SEM, non-significant by unpaired t-test).

### GSDME does not contribute to lung pathology or dysfunction during H1N1 IAV infection

Given the lack of overt effects of GSDME on H1N1 IAV infection outcomes (**Fig 1**), we examined whether more subtle effects could be observed in the lungs of mice lacking GSDME.

We first examined WT and KO lungs collected at day 7 post-infection with hematoxylin and eosin staining. We observed areas of inflammation, immune cell infiltration, alveolar thickening, and tissue damage in both WT and *Gsdme^−/−^* mice (**Fig 2A**), with unbiased software-based quantification of tissue consolidation showing no significant differences between groups (**Fig 2B**). Likewise, we saw similar levels of anti-CD45 staining, indicating similar immune cell infiltration in WT and KO lungs (**Fig 2C, D**). Similarly, staining for smooth muscle actin revealed comparable peribronchiolar and perivascular patterns in WT and *Gsdme^−/−^* lungs, suggesting no genotype-specific differences in smooth muscle activation or fibroproliferative remodeling (**Fig 2E, F**). As a highly sensitive measure of lung dysfunction, we used whole body plethysmography to examine potential differences in WT versus *Gsdme^−/−^* lungs during infection. Enhanced pause (Penh), a surrogate indicator of airway obstruction, increased similarly in both genotypes during infection, with a trend toward delayed resolution in WT animals, though differences were not statistically significant (**Fig 2G**). Likewise, the ratio of time to peak expiratory flow over total expiratory time (Rpef), considered an additional sensitive indicator of functional airflow limitation, showed similar values for WT and *Gsdme^−/−^* animal cohorts (**Fig 2H**). Collectively, these results further indicate that GSDME is dispensable for lung pathology and dysfunction in H1N1 IAV infection.

**Figure 2:**
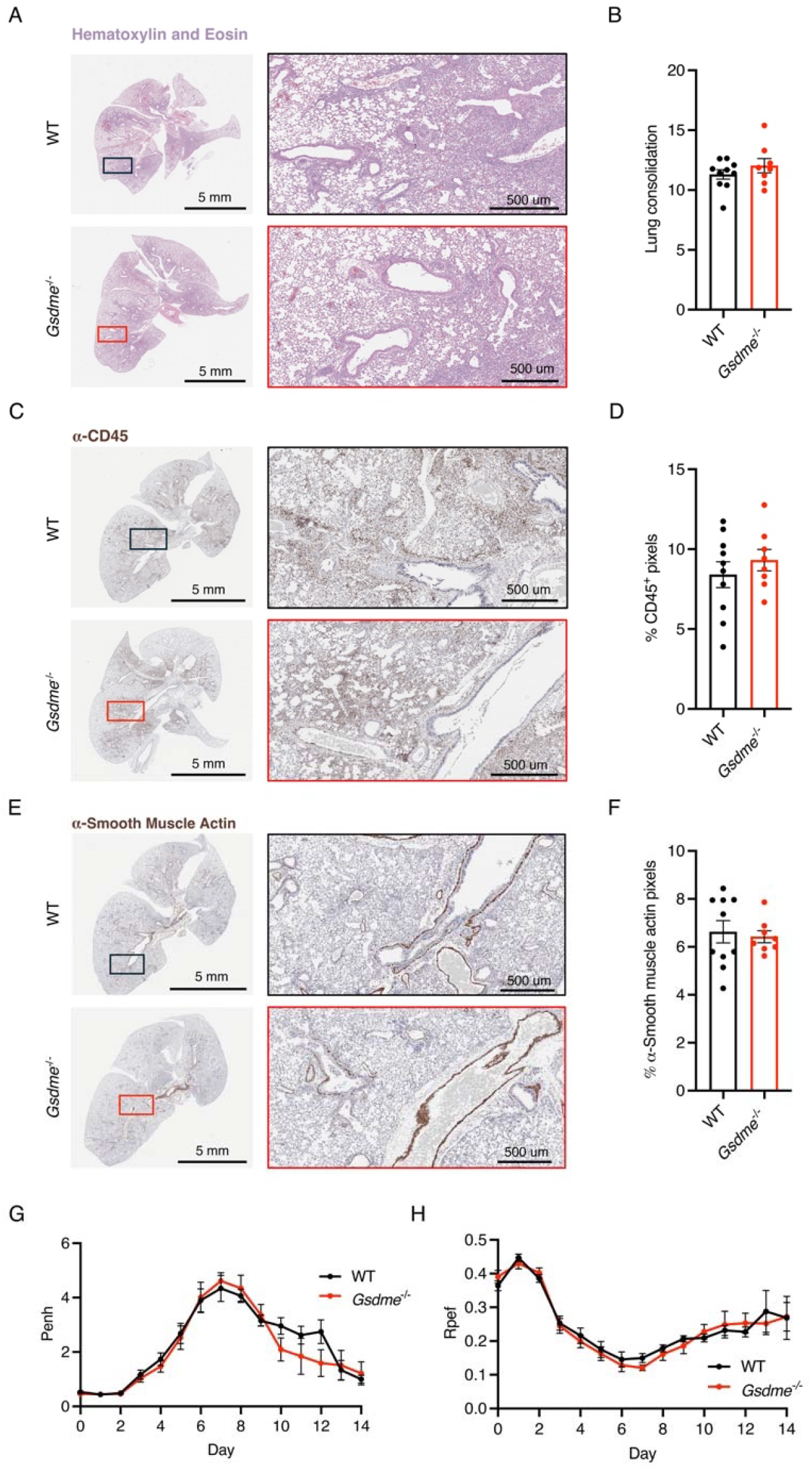
Lung function and histopathology is unchanged in *Gsdme^−/−^* mice in severe PR8 (H1N1) infection. A-E. WT and *Gsdme^−/−^* mice were infected with 50 TCID50 of PR8 intranasally. **A, C, E** Representative images from H&E, α-CD45, and α-smooth muscle actin staining, respectively, from lungs collected on day 7 post infection. Boxes show placement of magnified images. **B, D, F** Quantification of lung images from day 7 post infection of H&E, α-CD45, and α-smooth muscle actin staining, respectively (WT n = 10, *Gsdme^−/−^* n = 8, each dot represents an individual mouse, error bars indicate SEM, not significant by unpaired t-test). **G** Enhanced pause (Penh) measurements from daily whole body plethysmography (WT n = 20 and *Gsdme^−/−^* n = 17 for days 0-7, WT n = 10 and *Gsdme^−/−^* n = 9 for days 8-14, each dot is an average of individual mouse Penh values for that day, error bars represent SEM, no significant differences between genotypes at any timepoint by two-way ANOVA followed by Bonferroni’s multiple comparisons test). **H** Ratio of time to peak expiratory flow (Rpef) measurements from daily whole-body plethysmography (WT n = 20 and *Gsdme^−/−^* n = 17 for days 0-7, WT n = 10 and *Gsdme^−/−^* n = 9 for days 8-14, each dot is an average of individual mouse Rpef values for that day, error bars represent SEM, no significant differences between genotypes at any timepoint by two-way ANOVA followed by Bonferroni’s multiple comparisons test).

### GSDME has minimal effects on global lung transcription programs with or without infection

To investigate whether GSDME influences global gene expression in the lungs during IAV infection, we performed bulk RNA sequencing on whole lung homogenates from WT and *Gsdme^−/−^* mice at day 7 post-infection and genotype-matched mock controls. Principal component analysis revealed a strong separation between infected and mock samples, with minimal separation between WT and *Gsdme^−/−^* animals within either mock or infected conditions (**Fig 3A**). This was further confirmed by differential gene expression analysis (log2fc |0.58|; p-value adj 0.01) where minimal changes in baseline gene expression were observed between mock control WT and *Gsdme^−/−^* (KO) mice (**Fig 3B, left**) or PR8 infected WT and *Gsdme^−/−^* mice (**Fig 3B, right**). We then contrasted the lung transcriptional responses of infected mice to their genotype-matched uninfected controls (**Fig 3C**). Analysis of the overlap of differentially expressed genes (DEG) revealed an overlap of around 66% of transcripts modulated by viral infection across genotypes. Evaluation of the relative gene expression after infection demonstrated a high concordance in the directionality of change of DEG (**Fig 3D, Supplementary Table 1**), demonstrating that *Gsdme* does not significantly contribute to the direction of transcriptional changes. Consistent with prior results (**Fig 1E**), we observed that *Gsdme* expression had minimal impact on the magnitude of induction of antiviral effectors encoded by interferon-stimulated genes^32^ (ISGs) (**Fig 3E**). Gene ontology (GO) analysis of the overlapping DEG across genotypes indicate involvement in the ‘regulation of innate immune responses’ and the ‘regulation of inflammatory responses’ (**Fig 3F**). This was consistent with results from gene set enrichment analysis (GSEA), which indicated that the top 10 induced pathways, which were highly concordant and equivalently enriched in both WT and *Gsdme^−/−^* mice, corresponded to processes involved in antiviral and proinflammatory responses (**Fig 3G**). To better understand host response pathways shaped by GSDME, we conducted GO enrichment analysis in the subset of genes differentially expressed only in WT or *Gsdme^−/−^* mice (**Fig 3C**). This analysis revealed that WT mice had a significant and unique enrichment of pathways involved in ‘lung development’ and ‘muscle cell differentiation’ (**Fig 3F**). In contrast, PR8 infection uniquely elicited changes in genes involved in the ‘negative regulation of transmembrane transport’ and ‘regulation of muscle system processing in mice’ in *Gsdme^−/−^* mice (**Fig 3F**). Together, these data indicate that GSDME is dispensable for mounting a general antiviral and inflammatory response during acute infection. However, GSDME may contribute to non-immunological responses that coordinate tissue remodeling later during infection.

**Figure 3:**
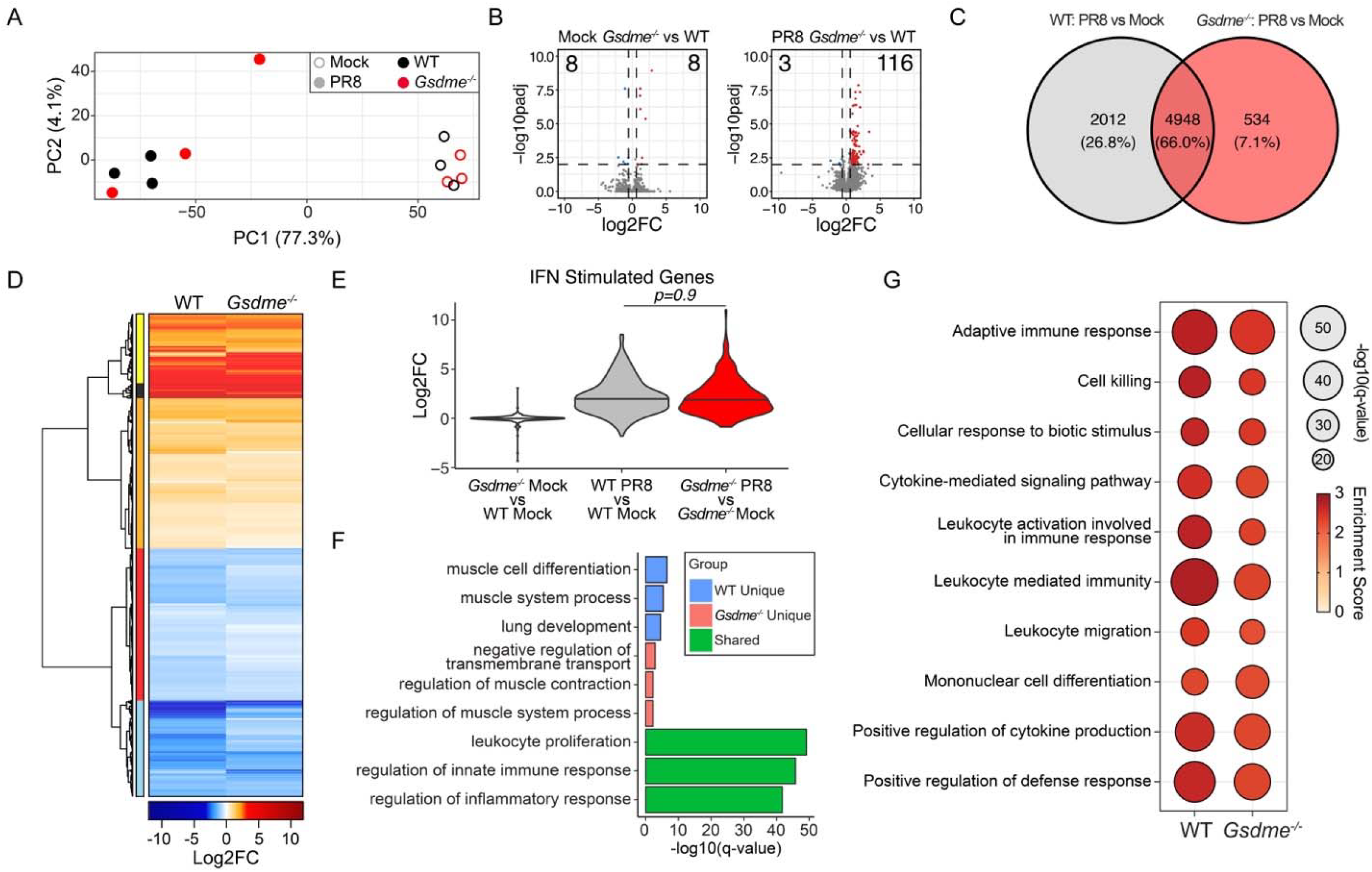
Global transcriptomic analysis reveals minor impacts of GSDME during H1N1 IAV infection. A-G. WT and *Gsdme^−/−^* mice were mock-infected or infected with 50 TCID50 of PR8 intranasally, and bulk lung RNA was isolated at day 7 post-infection for sequencing. **A** Principal component analysis of bulk lung transcriptomes. Open circles indicate mock-infected mice, and closed circles indicate PR8-infected mice. Color indicates genotype. **B** Volcano plots of differential gene expression results comparing (left) mock-infected *Gsdme^−/−^* to mock-infected WT lungs and (right) PR8-infected *Gsdme^−/−^* to PR8-infected WT lungs. Significant DEG are indicated by color, with blue indicating downregulation and red indicating upregulation (log2fc |0.58|; p-value adj 0.01). For the mock comparison (left), there are 8 downregulated and 8 upregulated DEG. For the infected comparison (right), there are 3 downregulated DEG and 116 upregulated DEG. **C-D** Differential gene expression results comparing PR8-infected lungs with mock-infected lungs for each genotype. 7494 DEG were identified (log2fc |0.58|; p-value adj 0.01). **C** Venn diagram comparing DEG overlap. **D** Differential expression heatmap of the 7494 DEG arranged by hierarchical clustering using Euclidean distance. Blue indicates downregulation during infection, and red indicates upregulation during infection. Colored bar (left) indicates gene clusters. **E** Violin plot showing log2fc values of interferon (IFN) stimulated genes for the indicated comparisons. Significance determined by one-way ANOVA with Tukey’s HSD test. **F** Bar graph of top GO term enrichment results for the gene sets from the Venn diagram in panel C. Blue indicates pathways enriched for the 2012 DEG unique to WT lungs; pink for the 534 DEG unique to *Gsdme^−/−^* lungs; green for the 4948 DEG shared between genotypes. X-axis indicates significance of enrichment. **G** Selected antiviral and inflammatory gene set enrichment analysis (GSEA) results for each genotype. Bubble size indicates significance, and color indicates pathway enrichment score.

### GSDME does not impact outcomes of pandemic 2009 H1N1 influenza virus infection

To rigorously test the generalizability of our findings from experiments with the PR8 H1N1 IAV strain, we infected WT and *Gsdme^−/−^* mice with an additional H1N1 strain, specifically the A/California/04/09 (H1N1) strain with a single change in its PB2 protein (E158A) that enhances replication in mouse cells^33^. A dose of 100 TCID50 of this strain was chosen to model a severe, partially lethal dose in which a significant portion of mice succumb to the infection. We observed similar patterns of weight loss between WT and KO animals (**Fig 4A**), nearly identical rates of survival (**Fig 4B**), and similar patterns of lung dysfunction as indicated by Penh (**Fig 4C**) and Rpef (**Fig 4D**) plethysmography measurements. These results further demonstrate that GSDME is not a major contributor to morbidity or mortality in severe H1N1 IAV infections and that these findings apply across divergent H1N1 strains.

**Figure 4:**
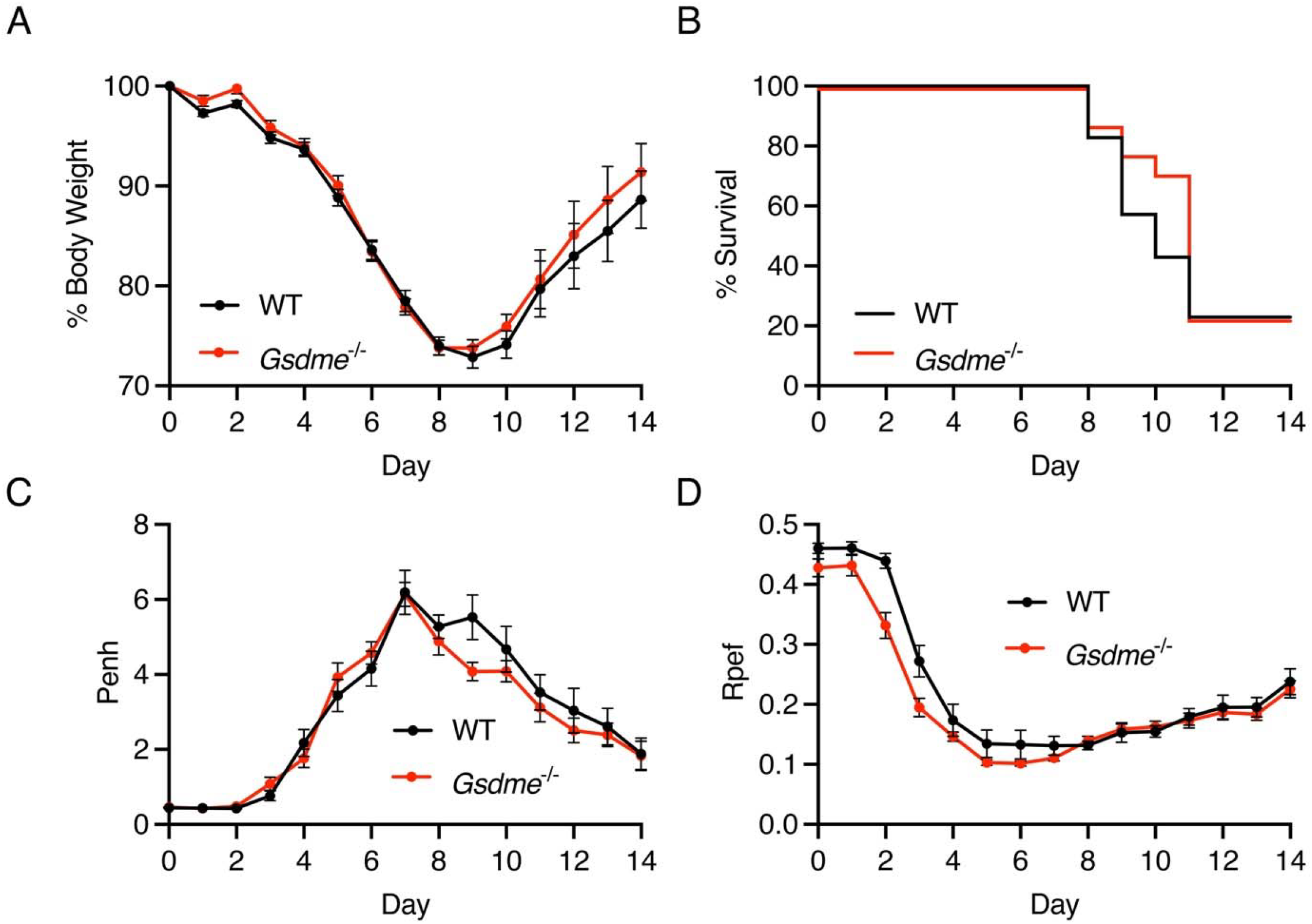
Loss of GSDME does not affect lung function or viral pathogenesis in a severe pandemic 2009 H1N1 IAV infection. A-D. WT and Gsdme^−/−^ mice were infected with 100 TCID50 of 2009 H1N1 intranasally. **A** Weight loss measurement (WT n = 35 and *Gsdme^−/−^* n = 31 for days 0-10, WT n = 20 and *Gsdme^−/−^* n = 17 for days 11-14, each dot is an average of individual mouse weights normalized to 100% relative to day 0, error bars indicate SEM, no significant differences between genotypes at any timepoint by two-way ANOVA followed by Bonferroni’s multiple comparisons test). **B** Survival analysis (WT n = 20, *Gsdme^−/−^* n = 17, not significant by Log-rank Mantel-Cox test. **C** Enhanced pause (Penh) measurements from daily whole body plethysmography (WT n = 35 and *Gsdme^−/−^* n = 31 for days 0-10, WT n = 20 and *Gsdme^−/−^* n = 17 for days 11-14, each dot is an average of individual mouse Penh values for that day, error bars represent SEM, no significant differences between genotypes at any timepoint by two-way ANOVA followed by Bonferroni’s multiple comparisons test). Ratio of time to peak experiatory flow (Rpef) measurements from daily whole-body plethysmography (WT n = 35 and *Gsdme^−/−^* n = 31 for days 0-10, WT n = 20 and *Gsdme^−/−^* n = 17 for days 11-14, each dot is an average of individual mouse Rpef values for that day, error bars represent SEM, no significant differences between genotypes at any timepoint by two-way ANOVA followed by Bonferroni’s multiple comparisons test).

## DISCUSSION

Prior studies have implicated pyroptosis, particularly GSDMD-dependent pyroptosis, in driving lung inflammation and pathology during IAV infections. This was demonstrated by our group examining H1N1 PR8 infection^24^ and by a second group using an IAV H3N2 strain^25^. In both cases, morbidity and mortality were decreased in *Gsdmd^−/−^* mice. Upon mining expression data for other gasdermins in published lung single-cell RNA sequencing data, we noted expression of GSDME in a variety of human and mouse lung cells, including macrophages, neutrophils, and epithelial cells^34, 35^, which are known targets of IAV infection and play roles in influenza pathogenesis^36^. Interestingly, *Gsdme^−/−^* mice were shown to be protected from death during a lethal dose infection with highly pathogenic avian H7N9 virus isolated from a fatal human case^26^. An additional recent study reported modest improvements in disease scores and survival for *Gsdme^−/−^* mice during H3N2 IAV infection, although the statistical approaches and endpoint criteria used in the analysis may limit the strength of the conclusions^27^. Nonetheless, whether lung pathogenesis caused by H1N1 IAV, which significantly contributes to the burden of influenza cases and hospitalizations worldwide^28, 37, 38^, is similarly impacted by GSDME was not previously examined.

Several studies have suggested that H1N1 IAV infections trigger caspase-3 and subsequent GSDME cleavage in human lung epithelial cell lines, human primary bronchial epithelial cell lines, and mouse lung tissue^27, 29–31^. However, we are the first to investigate the contribution of GSDME to H1N1 IAV morbidity and mortality *in vivo*. Here, we found that unlike H7N9 and H3N2 infections, H1N1 IAV pathogenesis is largely unaffected by loss of GSDME. While global transcriptomic analysis suggested a subtle decrease in some lung inflammatory gene programs in *Gsdme^−/−^* mice, this did not affect overall outcomes in terms of weight loss, survival, lung function, or histopathology. Importantly, we observed minimal effects of GSDME loss in two distinct models of severe H1N1 infection, namely in infections with the classic and well-studied PR8 strain from 1934 and with the 2009 H1N1 pandemic virus, which is the precursor of H1N1 strains currently circulating in humans. Given that our previously published results with *Gsdmd^−/−^* mice showed major benefits in PR8 infections when using the same viral stock and dose used in the present study, this suggests that inflammasome/GSDMD-dependent pyroptosis is more relevant in H1N1 infections than apoptosis-linked, GSDME-dependent pyroptosis.

Our results on H1N1 IAV infections in the context of the literature also suggest that distinct IAV subtypes may activate unique cell death and pathogenic programs. While all IAV subtypes can likely induce host cell death^4–11^, the relative contribution of specific pathways such as pyroptosis, apoptosis, and necroptosis may vary based on viral strain, replication kinetics, and host immune engagement and antagonism. H1N1 viruses, particularly PR8 and the 2009 pandemic strain, have been shown to strongly activate inflammasome signaling and drive caspase-1-dependent pyroptosis, consistent with the critical role of GSDMD in these models^24, 39, 40^. H3N2 and H7N9 strains may potentially evade this GSDMD-dependent inflammasome pathway more effectively^41^ or more strongly activate apoptotic pathways, resulting in increased cleavage of caspase-3 and GSDME. These differences may reflect underlying variation in how each subtype interacts with or evades innate immune sensors, induces stress responses, or infects distinct target cell populations within the lung. Overall, our findings reveal the importance of considering IAV subtype when evaluating the contribution of specific cell death pathways to disease severity.

GSDME has been implicated in tumor suppression^42^ and chemotherapy-induced cell death^43, 44^. It remains to be determined whether GSDME may play a more significant role in the context of secondary bacterial infections, which are common complications of severe influenza, including H1N1 infections. Co-infection could increase epithelial or immune cell apoptosis, leading to enhanced GSDME cleavage and potentially shifting the balance of inflammatory cell death pathways. Additionally, although our transcriptomic analysis revealed only minimal changes in gene expression of antiviral and inflammatory genes, it remains to be determined whether GSDME may impact lung repair processes. Further studies incorporating co-infection models, high-resolution transcriptomic approaches, or long-term follow-up of infected mice may be necessary to fully define subtle or context-dependent roles of GSDME in H1N1 IAV infections. Lastly, our finding that *Gsdme^−/−^* mice show minimal differences in disease outcome during H1N1 IAV infection suggests that therapeutic targeting of GSDME in other clinical settings may not carry unintended consequences for H1N1 IAV susceptibility or pathogenesis.

In summary, our findings demonstrate that GSDME plays a minimal role in the pathogenesis of H1N1 IAV infection, distinguishing it from other subtypes where GSDME-dependent pyroptosis may be more relevant. These results refine our understanding of gasdermin family members in viral disease and emphasize the need to consider viral strain-specific differences in host response and disease pathogenesis.

## Methods

### Virus Stocks

Influenza viruses used in this study include A/PR/8/34 (H1N1) (PR8, provided by Dr. Thomas Moran of the Icahn School of Medicine at Mt. Siani) and A/California/04/2009 (H1N1) containing the PB2 E158A mutation (originally from BEI Resources, NR-13659, and mouse adapted as previously described^33^). PR8 was propagated in 10-day embryonated specific pathogen free chicken eggs (AVSbio, 10100331) at 37 degrees Celsius for 2 days. 2009 H1N1 virus was propagated for 2 days in MDCK cells (BEI Resources, NR-26-28) in the presence of TPCK-treated trypsin (Worthington Biochemical). Viruses were flash frozen for storage at −80 degrees Celsius for one-time usage.

### Mouse studies

Male and female *Gsdme^−/−^* mice (Jackson Laboratory strain #032411)^45^ and C57BL/6NJ WT control mice (Jackson Laboratory strain #005304) ages 7-10 weeks were used for all experiments. For *in vivo* infections, mice were anesthetized using 4% isoflurane, and 50 μL of diluted virus (50 TCID50 of PR8 or 100 TCID50 of 2009 H1N1) in sterile saline was delivered intranasally. Mice were weighed each day and euthanized via CO_2_ gas and subsequent cervical dislocation based on the humane endpoint criteria of weight loss greater than 30% of their baseline and lack of mobility. Throughout infection, lung function was measured via whole body plethysmography (Buxco Small Animal Whole Body Plethysmography). Mice were acclimated to the plethysmography chamber for 3 days prior to infection for 10 min/day. During recordings, mice were put in the plethysmography chamber for a 5 min acclimation period followed by a 5 min reading. All plethysmography variables were recorded every 10 seconds during the 5 min reading period and averaged across the 5 min reading for each individual day. All procedures were approved by the OSU IACUC.

### Viral titers and cytokine quantification

For determining viral titer and cytokine levels, mice were sacrificed at day 7 post-infection using CO_2_ gas and subsequent cervical dislocation. Lungs were harvested and homogenized in 1 mL PBS using a tissue bead-beating system (Precellys Evolution Touch Homogenizer). After homogenization, tubes were centrifuged at 3000 x g for 3 min to pellet tissue debris. The supernatant was aliquoted, flash frozen, and stored at −80 degrees Celsius for one-time use in titer or cytokine assays. Viral titer determinations for stock viruses and tissue samples were performed as previously described^46–48^. In brief, MDCK cells were grown in Dulbecco’s Modified Eagle’s Medium (DMEM) with 10% fetal bovine serum (Atlas Biologicals EquaFetal serum) and 1% penicillin-streptomycin at 37 degrees Celsius with 5% CO_2_ in a humidified incubator. For titer assays, samples at 1:10 dilutions were incubated for 3 days on cells in DMEM with 2% BSA and TPCK-treated trypsin (Worthington Biochemical Corporation). After incubation, cells were fixed with 4% PFA, permeabilized in 0.1% Triton X 100 in PBS, and blocked with 2% FBS in PBS for 10 min each. Virus-positive cells were visualized using IAV nucleoprotein primary antibody (BEI Resources, NR-19868) and AlexaFlour 488-conjugated secondary antibody (ThermoFisher Scientific, A11029) via fluorescent microscopy (EVOS Cell Imaging System). The Reed-Muench method was used in determining the final titers of samples. For cytokine quantification, R&D Systems DuoSet ELISA kits for IL-1β (DY-401), TNF (DY410), IL-6 (DY406), IFN-β (DY8234-05), and IL-18 (DY7625-05) were used as specified with lung homogenates. Resulting samples were analyzed on a spectrophotometer plate reader at 450 nm.

### Histology

For histology, mice were sacrificed at day 7 post-infection as described above, and lung tissue was carefully dissected out of the thoracic cavity to preserve tissue architecture. Lung tissue was then fixed in 10% neutral buffered formalin at 4 degrees Celsius for 24 h and then transferred to 70% ethanol. Lungs were embedded in paraffin, sectioned, stained (H&E, α-CD45, and α-smooth muscle actin), and imaged by Histowiz (Histowiz.com, Brooklyn, NY, USA). Unbiased electronic quantification of the resultant images was performed using ImageJ software and the color deconvolution method as described previously^24, 47, 49–51^.

### Western Blotting

Detection of GSDME expression and cleavage in the lung was performed via western blot. Mice were sacrificed as above, and lungs were collected. Lung tissue was homogenized and lysed in 1% SDS buffer (1% SDS, 50mM triethanolamine pH 7.4, 150mM NaCl) containing protease inhibitors (Thermo Scientific, A32965). Cell debris was removed by centrifugation for 10 min at 20,000 x g, and clarified supernatants were used for Western blotting. Total protein per sample was quantified via BSA assay, and 30 μg protein was loaded for separation by SDS-PAGE and subsequently transferred onto membranes. Membranes were blocked with 10% non-fat milk in PBS with 0.1% Tween-20 (PBST) and probed with primary antibodies against GAPDH (Invitrogen, 39-8600) and GSDME (Abcam, EPR19859). HRP-conjugated anti-rabbit IgG secondary antibody (1:10,000, Cell Signaling Technologies #7074) was subsequently used, and chemiluminescent images were acquired using a ChemiDoc Touch (BioRad).

### RNA isolation, sequencing, and analysis

Mice were sacrificed as described above on day 7 post-infection, and lung tissue was homogenized in TRIzol reagent as described above. Following separation with chloroform, the aqueous phase was collected and RNA was precipitated with ethanol. RNA library preparation and sequencing was performed by GENEWIZ from Azenta Life Sciences (South Plainfield, NJ, USA). Sequencing data were analyzed as previously described^24, 51^. Principal components analysis and data visualization was done using R (v4.4.2) with packages ggplot2 (v3.5.1) and ggvenn (v0.1.10). Sample count normalization and differential gene expression were performed using DESeq2 (v1.46.0). Gene Ontology analysis was performed using the clusterProfiler R package (v4.14.6). Gene set enrichment analysis (GSEA) was performed using published methods^52^.

### Quantification and statistical analysis

Statistical analysis was performed using t-tests, ANOVA, and Log-rank (Mantel-Cox) tests, as appropriate. Parametric tests assumed (i) normally distributed residuals within groups, (ii) equal variances across groups, and (iii) independent observations. The Log-rank test was applied to fully observed survival data, assuming independent survival times. All analyses were performed using GraphPad Prism v10.5.0.

## Data Availability

RNA sequencing data files have been deposited in NCBI’s Gene Expression Omnibus and will be released upon publication.

