## Supplemental Table 1 for "Gasdermin E is Dispensable for H1N1 Influenza Virus Pathogenesis in Mice"

| gene | cluster | WT_InfvsMoc | WT_InfvsMoc | KO_InfvsMoc | KO_InfvsMock_padj |
|---|---|---|---|---|---|
| Cdc45 | orange | 1.92884334 | 2.09E-19 | 1.88534913 | 8.60E-18 |
| H19 | skyblue | -1.6424284 | 7.03E-06 | -0.851421 | 0.02878107 |
| Cav2 | red | -0.9335846 | 1.34E-05 | -0.8042964 | 0.00027357 |
| Klf6 | orange | 0.8728701 | 1.30E-12 | 0.55555351 | 1.43E-05 |
| Tbx2 | red | -0.733841 | 7.83E-07 | -0.5594865 | 0.0002772 |
| Wnt9a | red | -0.8498308 | 0.00240453 | -0.2438622 | 0.44456332 |
| Xpo6 | orange | 0.82317906 | 2.51E-10 | 0.77908927 | 3.67E-09 |
| Axin2 | red | -0.7959924 | 5.56E-05 | -0.7731261 | 0.00013123 |
| Slc22a18 | red | -1.0469053 | 2.56E-06 | -0.6448993 | 0.00573982 |
| Pih1d2 | red | -1.0308542 | 0.00050357 | -0.0922554 | 0.79537579 |
| Fgf23 | yellow | 3.82814158 | 0.00040528 | 3.5248553 | 3.58E-05 |
| Nalcn | skyblue | -2.150931 | 0.00332943 | -0.756844 | 0.32653805 |
| Slfn4 | black | 8.52797878 | 3.06E-111 | 7.48233918 | 7.16E-97 |
| Tspan32 | orange | 0.73564681 | 0.00030458 | 1.25944637 | 1.24E-09 |
| Lhx2 | black | 4.68599788 | 1.61E-09 | 5.97836216 | 9.32E-09 |
| Gmpr | skyblue | -1.5000948 | 6.54E-09 | -1.3323724 | 5.43E-07 |
| Mid2 | red | -0.9436883 | 4.96E-06 | -0.6942044 | 0.00121025 |
| Trim25 | orange | 1.43192406 | 4.44E-25 | 1.21474386 | 3.78E-18 |
| Scpep1 | orange | 0.71017073 | 0.00116841 | 0.58002007 | 0.01075027 |
| Itgb2 | yellow | 2.59852941 | 2.42E-15 | 2.36860263 | 9.69E-13 |
| Hddc2 | orange | 0.55529533 | 0.01094454 | 0.63921507 | 0.0040822 |
| Tpd52l1 | red | -1.5054475 | 0.00012265 | -0.7632342 | 0.07868237 |
| Cdh4 | red | -1.0897592 | 0.00775348 | -0.8696099 | 0.04073213 |
| Bcl6b | orange | 1.18826353 | 0.00089035 | 1.15240166 | 0.00168806 |
| Clec10a | orange | 1.03561928 | 0.00023132 | 1.32690684 | 3.22E-06 |
| Arvcf | red | -1.502843 | 1.33E-07 | -1.02996 | 0.00049638 |
| Rtca | orange | 0.63360097 | 4.52E-06 | 0.7487309 | 9.40E-08 |
| Rem1 | red | -1.0481287 | 8.64E-05 | -0.9056515 | 0.00099752 |
| Tbrg4 | orange | 0.93475539 | 4.64E-09 | 0.82561347 | 4.15E-07 |
| Tmprss2 | red | -1.4942138 | 1.06E-13 | -1.0025063 | 1.41E-06 |
| Mx1 | black | 6.21935933 | 4.11E-79 | 6.35017095 | 4.89E-81 |
| Egfl6 | red | -1.0093457 | 0.00066741 | -0.4634148 | 0.15191645 |
| Lck | yellow | 2.64914809 | 3.81E-07 | 3.59907634 | 4.99E-12 |
| Cttnbp2 | red | -1.4415748 | 1.88E-07 | -0.7076606 | 0.01556756 |
| Galnt1 | orange | 0.72093665 | 5.66E-06 | 0.63875725 | 9.29E-05 |
| Sept1 | yellow | 2.39398302 | 1.13E-33 | 2.59264637 | 8.11E-39 |
| Acvrl1 | red | -0.9657486 | 4.83E-11 | -0.9096154 | 1.03E-09 |
| Tom1l2 | red | -0.7830779 | 2.09E-08 | -0.6748216 | 2.47E-06 |
| Zfp385a | orange | 0.88091893 | 0.00128741 | 0.6676618 | 0.01971043 |
| Itga5 | orange | 1.39715245 | 5.31E-06 | 1.35115856 | 1.70E-05 |
| Adora3 | yellow | 2.52034982 | 2.59E-09 | 2.75479694 | 1.20E-09 |
| Hnrnpd | orange | 0.85005339 | 1.85E-13 | 0.79117525 | 1.50E-11 |

| | | | | | |
|---|---|---|---|---|---|
| Gm2a | orange | 0.58957536 | 0.00141192 | 0.27320184 | 0.17546667 |
| Sema4f | red | -1.3794762 | 3.59E-09 | -0.9922996 | 4.33E-05 |
| Haao | orange | 1.22233445 | 2.15E-05 | 1.03567733 | 0.00050859 |
| Cd52 | yellow | 3.44558914 | 1.12E-35 | 3.08200614 | 1.67E-28 |
| Loxl3 | orange | 0.74709353 | 8.44E-06 | 0.64303449 | 0.00019674 |
| Rab5b | red | -0.607886 | 4.49E-07 | -0.5249424 | 2.26E-05 |
| Serpinf1 | skyblue | -1.8240437 | 4.06E-13 | -0.9641605 | 0.00024156 |
| Tcf7 | orange | -0.0959541 | 0.84627805 | 1.12034345 | 0.00887383 |
| Il12rb1 | black | 5.40474755 | 4.77E-42 | 5.4378242 | 1.91E-41 |
| Slc5a5 | skyblue | -0.7727786 | 0.30827757 | -2.1009179 | 0.0071946 |
| Kcnn3 | red | -1.0553365 | 0.0010009 | -0.5070087 | 0.14083378 |
| Car4 | red | -0.7576541 | 0.00108077 | -0.6995335 | 0.00336138 |
| Txnrd3 | red | -0.9406323 | 1.52E-05 | -0.8264709 | 0.00023557 |
| Fasl | yellow | 3.77810147 | 3.80E-11 | 3.43639052 | 2.32E-08 |
| Il4 | skyblue | -1.566781 | 0.00327445 | -1.3620086 | 0.01234153 |
| Pxmp4 | red | -0.7450768 | 7.52E-06 | -0.654841 | 0.00013237 |
| Dlg3 | red | -0.6819356 | 1.06E-06 | -0.5668882 | 7.73E-05 |
| Mmp11 | red | -1.2277788 | 8.08E-09 | -0.8591474 | 0.00010278 |
| Vpreb3 | skyblue | -3.6825086 | 3.15E-15 | -2.5155342 | 9.88E-08 |
| Hoxa4 | red | -0.6466697 | 0.00486645 | -0.3899176 | 0.12040627 |
| Mmp14 | orange | 1.93987109 | 2.32E-10 | 1.70249385 | 5.19E-08 |
| Ccl3 | black | 5.76593325 | 6.07E-65 | 5.11484331 | 6.60E-57 |
| S100a4 | yellow | 1.91923593 | 0.00743203 | 2.56887421 | 0.00783685 |
| S100a6 | orange | 1.33819662 | 5.35E-08 | 1.27028576 | 4.16E-07 |
| Scn4a | red | -0.9571357 | 0.00032045 | -0.762562 | 0.00641423 |
| Epn2 | red | -0.7852815 | 1.69E-06 | -0.6338744 | 0.00018271 |
| N4bp3 | red | -0.6537136 | 2.65E-09 | -0.6480793 | 5.92E-09 |
| Nhp2 | orange | 1.05138993 | 5.49E-07 | 1.02570167 | 1.85E-06 |
| Mfsd10 | orange | 0.71755246 | 5.01E-05 | 0.57258917 | 0.00179679 |
| Lgals9 | orange | 1.88460139 | 5.31E-24 | 1.85540893 | 3.16E-23 |
| Cfp | orange | 0.70706001 | 0.00027417 | 0.64638754 | 0.00130395 |
| Timp1 | black | 6.15335123 | 6.03E-35 | 6.01825166 | 2.76E-33 |
| Uxt | orange | 0.73976994 | 0.00148705 | 0.37250742 | 0.14075643 |
| Cnnm3 | red | -0.6698148 | 0.00086621 | -0.5692173 | 0.00620659 |
| Mxd1 | orange | 1.72628126 | 1.27E-18 | 1.53686454 | 7.98E-15 |
| Oas1c | orange | 1.17037135 | 7.80E-08 | 1.04374748 | 4.06E-06 |
| Oas1h | black | 6.16179678 | 6.32E-06 | 4.87204506 | 4.38E-06 |
| Sema6b | orange | 1.57327672 | 7.49E-08 | 1.07433844 | 0.00041434 |
| Uhrf1 | yellow | 2.64224952 | 1.01E-23 | 2.92551609 | 1.99E-28 |
| Ramp2 | red | -0.7079532 | 0.00217021 | -0.6750532 | 0.00445001 |
| Gramd1a | orange | 1.01362849 | 2.72E-29 | 0.73175161 | 1.64E-15 |
| Hpn | red | -0.6247772 | 6.90E-05 | -0.1933128 | 0.26862157 |
| Itgb7 | yellow | 2.30773289 | 5.04E-27 | 2.12325202 | 7.12E-23 |

| | | | | | |
|---|---|---|---|---|---|
| Rarg | red | -0.9489458 | 2.85E-12 | -0.7143145 | 3.29E-07 |
| Efnb2 | skyblue | -1.8416586 | 2.19E-15 | -1.4259874 | 1.95E-09 |
| Rrp15 | orange | 0.66166073 | 0.00922179 | 0.84578863 | 0.00105077 |
| Rnd2 | red | -0.6653604 | 0.00219818 | -0.6435987 | 0.00395558 |
| Srr | red | -0.6658932 | 1.88E-06 | -0.4772369 | 0.00106378 |
| Sync | skyblue | -1.687352 | 2.42E-11 | -0.9097688 | 0.00060339 |
| Fndc5 | skyblue | -1.7476784 | 0.01064364 | -2.7448222 | 6.25E-05 |
| Acp5 | orange | 1.64557654 | 1.43E-08 | 1.25558707 | 2.87E-05 |
| Cnn1 | skyblue | -1.6464721 | 1.15E-11 | -0.6985339 | 0.00669709 |
| Ube2c | yellow | 3.11661452 | 2.02E-24 | 3.33605824 | 1.14E-26 |
| Cct3 | orange | 0.70616329 | 2.41E-05 | 0.56914937 | 0.00101282 |
| Col18a1 | orange | 0.64096284 | 0.01268739 | 0.7711614 | 0.00305708 |
| Kpnb1 | orange | 0.59676614 | 1.77E-06 | 0.59244166 | 3.36E-06 |
| Tbx21 | yellow | 3.54029342 | 2.76E-68 | 3.77578234 | 5.17E-71 |
| Tubb6 | orange | 1.76094442 | 1.56E-06 | 1.91832281 | 2.49E-07 |
| Meox1 | red | -1.3013653 | 1.73E-08 | -0.8322526 | 0.00052788 |
| Sost | skyblue | -1.856129 | 2.74E-06 | -1.577481 | 0.00015844 |
| Nkx2-1 | red | -1.2622317 | 3.87E-08 | -0.8027626 | 0.00081491 |
| Irx2 | red | -0.8042901 | 0.00028152 | -0.4562103 | 0.05352411 |
| Col1a1 | orange | 0.72398668 | 0.0280677 | 0.92436992 | 0.00542937 |
| Itga3 | orange | 0.50194573 | 0.00365738 | 0.73659005 | 2.12E-05 |
| Sgca | red | -0.8429488 | 0.00066005 | -1.1524014 | 6.98E-06 |
| Foxm1 | yellow | 2.46961986 | 5.50E-34 | 2.62130763 | 2.02E-37 |
| Tubb5 | orange | 1.15071771 | 1.58E-09 | 1.01018544 | 2.17E-07 |
| Ell2 | orange | 0.78350654 | 4.69E-05 | 0.500292 | 0.01332754 |
| Jup | red | -0.9852849 | 6.94E-15 | -0.8414938 | 6.33E-11 |
| Acap1 | yellow | 1.81769927 | 1.49E-25 | 2.04351233 | 1.48E-31 |
| Tcea3 | red | -1.4005625 | 8.60E-11 | -1.0354566 | 3.53E-06 |
| Ifrd1 | orange | 0.801628 | 4.73E-12 | 0.52808808 | 1.14E-05 |
| Stk38l | orange | 0.41206265 | 0.00479216 | 0.70919302 | 1.01E-06 |
| Brpf1 | orange | 0.81213617 | 2.56E-10 | 0.57386701 | 1.52E-05 |
| Gstt1 | red | -1.2535379 | 3.14E-06 | -0.9772404 | 0.00044831 |
| Gstt3 | red | -0.8828657 | 1.39E-05 | -0.6514478 | 0.00216128 |
| Tat | orange | 1.28316248 | 0.00311832 | -1.1154195 | 0.01455459 |
| Gramd3 | orange | 0.83978592 | 2.03E-10 | 0.85787475 | 1.68E-10 |
| Eef1e1 | orange | 0.99229352 | 0.00081065 | 0.92806951 | 0.00239615 |
| Cldn15 | red | -1.0307108 | 0.0052167 | -0.6655131 | 0.09015648 |
| Il16 | orange | 0.50492568 | 0.00139825 | 0.63342781 | 7.74E-05 |
| Tcirg1 | orange | 1.89659502 | 4.32E-19 | 1.83803358 | 7.26E-18 |
| Smo | red | -0.8643607 | 7.41E-13 | -0.5935533 | 1.87E-06 |
| Tspan33 | orange | 0.93319028 | 6.18E-06 | 0.39771765 | 0.0827103 |
| Folh1 | orange | 2.32582638 | 0.00189309 | 1.36974162 | 0.06721875 |
| Lrp3 | skyblue | -1.4781644 | 1.43E-14 | -1.6114027 | 5.01E-17 |

| | | | | | |
|---|---|---|---|---|---|
| Cpa3 | skyblue | -2.3283458 | 3.62E-08 | -1.4657094 | 0.00056798 |
| Nfix | red | -0.7569254 | 9.10E-13 | -0.5521654 | 4.37E-07 |
| Uba1 | orange | 0.65162936 | 6.77E-06 | 0.61505265 | 3.37E-05 |
| Siae | red | -0.3003766 | 0.15205863 | -0.5966088 | 0.00367343 |
| Spa17 | red | -1.2350424 | 3.63E-09 | -1.1703319 | 5.87E-08 |
| Grik3 | skyblue | -1.9863301 | 0.00022724 | -0.8584816 | 0.18679042 |
| Sipa1l2 | red | -0.8206547 | 3.97E-08 | -0.6782076 | 1.01E-05 |
| Blvra | orange | 1.10772303 | 5.11E-07 | 0.88132782 | 0.00010904 |
| Pdzd4 | red | -0.6863953 | 0.00028428 | -0.5287061 | 0.00753812 |
| Srpk3 | red | -1.1719466 | 0.00021365 | -1.1296324 | 0.00060667 |
| Pnck | skyblue | -1.4729154 | 4.32E-09 | -1.1707972 | 6.17E-06 |
| Ssr4 | orange | 0.65529103 | 0.00038707 | 0.69910083 | 0.00020895 |
| Ltbp2 | red | -0.704733 | 0.00300127 | -0.6454668 | 0.00837543 |
| Ift46 | red | -0.7159186 | 1.88E-05 | -0.4463794 | 0.01104735 |
| Tmem25 | red | -1.4282462 | 4.50E-06 | -0.8000323 | 0.01714846 |
| Cd3g | yellow | 3.56368443 | 2.97E-25 | 3.43154567 | 5.30E-23 |
| Spag5 | yellow | 2.99686472 | 1.62E-34 | 3.5274723 | 5.87E-43 |
| Unc119 | orange | 0.59094572 | 0.00615248 | 0.53015259 | 0.0178705 |
| Ccne1 | yellow | 3.29856608 | 1.11E-25 | 2.83914099 | 2.34E-19 |
| Psmc3 | orange | 0.67857414 | 5.41E-05 | 0.58050881 | 0.00082652 |
| Acp2 | orange | 0.7722992 | 0.00056369 | 0.50293389 | 0.03349515 |
| Rapsn | red | -0.928292 | 0.00457567 | -0.6608537 | 0.05925388 |
| Celf2 | orange | 0.67924549 | 4.65E-06 | 0.30142218 | 0.06008948 |
| Spi1 | yellow | 2.90859969 | 1.41E-23 | 2.67283472 | 6.70E-20 |
| Smg9 | orange | 0.82386455 | 1.52E-09 | 0.80772869 | 5.56E-09 |
| Rmnd5a | red | -0.8881111 | 0.00072556 | -0.4484255 | 0.11418127 |
| Mov10 | orange | 1.50543222 | 2.66E-28 | 1.20035152 | 3.03E-18 |
| Ppm1j | orange | 1.11875321 | 1.06E-05 | 1.38595455 | 8.22E-08 |
| Rhoc | orange | 1.49398877 | 1.42E-07 | 1.41656305 | 1.02E-06 |
| Def6 | orange | 1.86452293 | 3.17E-33 | 1.88353237 | 1.29E-33 |
| Zim1 | red | -0.4287768 | 0.40496438 | -1.5054131 | 0.00332419 |
| Angptl4 | orange | 1.39108031 | 8.17E-17 | 1.10665261 | 6.99E-11 |
| Dbf4 | orange | 2.05035705 | 3.77E-16 | 1.65824375 | 7.46E-11 |
| Daxx | yellow | 2.46876625 | 7.02E-35 | 2.26865601 | 1.51E-29 |
| Ipo4 | orange | 0.46981074 | 0.02415679 | 0.63160045 | 0.00259339 |
| Rec8 | red | -1.3100133 | 2.45E-05 | -0.5741791 | 0.08529432 |
| Irf9 | orange | 1.21852812 | 1.22E-23 | 1.21109055 | 2.71E-23 |
| Slc25a42 | red | -1.157098 | 2.38E-05 | -1.1069896 | 8.20E-05 |
| Dyrk1b | red | -0.8427883 | 1.81E-05 | -0.675155 | 0.00090016 |
| Ndufb2 | red | -0.8352005 | 0.00109045 | -0.7731059 | 0.00364483 |
| Rgs19 | orange | 1.18723118 | 2.55E-07 | 1.09356304 | 3.63E-06 |
| Abhd3 | skyblue | -2.6216739 | 1.94E-12 | -1.5510852 | 3.59E-05 |
| Snrpd1 | orange | 1.14990523 | 2.39E-09 | 0.96677301 | 1.16E-06 |

| | | | | | |
|---|---|---|---|---|---|
| Tsc2 | red | -0.792668 | 3.95E-07 | -0.6177061 | 0.00013121 |
| Rpl3l | skyblue | -1.6153837 | 8.03E-06 | -1.3263579 | 0.00042005 |
| Slc9a3r2 | red | -1.0904485 | 1.40E-08 | -0.9704483 | 7.96E-07 |
| Ikzf4 | orange | 1.66790866 | 7.42E-10 | 1.51139606 | 4.61E-08 |
| Pon1 | skyblue | -2.6564591 | 0.00013546 | -1.2908381 | 0.0844935 |
| Tgfb1 | orange | 0.58581467 | 0.00011203 | 0.48981696 | 0.00178158 |
| Shh | skyblue | -1.6468214 | 3.64E-08 | -1.1956509 | 0.00012152 |
| Gtf2f1 | orange | 1.21265107 | 7.46E-11 | 1.11695019 | 3.86E-09 |
| Dennd1c | orange | 1.86546049 | 1.12E-25 | 1.94943437 | 1.34E-27 |
| Prkd1 | red | -0.8390857 | 2.16E-09 | -0.7501332 | 1.60E-07 |
| Lcp2 | yellow | 3.25944095 | 7.23E-38 | 3.28626334 | 2.87E-38 |
| Naa20 | orange | 0.92228253 | 1.03E-07 | 0.687372 | 0.00013063 |
| Pex6 | red | -0.8818601 | 3.19E-10 | -0.8411738 | 3.71E-09 |
| Tmem143 | red | -1.0112801 | 9.54E-09 | -0.6991277 | 0.000149 |
| Ggct | orange | 1.79041787 | 1.88E-19 | 1.67363451 | 6.22E-17 |
| Jag2 | red | -0.5735144 | 1.27E-05 | -0.6072663 | 5.54E-06 |
| Epdr1 | skyblue | -1.9665204 | 9.72E-40 | -1.2128496 | 5.20E-16 |
| Top3a | orange | 0.61518489 | 0.0001032 | 0.46214495 | 0.00492803 |
| Qtrt1 | orange | 0.52857175 | 0.00833918 | 0.61232402 | 0.00271527 |
| Chaf1a | orange | 1.86517562 | 2.82E-13 | 1.82592047 | 1.73E-12 |
| Pla1a | yellow | 3.92611013 | 2.01E-25 | 3.91562714 | 6.03E-25 |
| Mcm2 | orange | 1.55727907 | 3.22E-14 | 1.47990645 | 9.98E-13 |
| Tpra1 | red | -0.7738334 | 1.52E-07 | -0.5932909 | 1.00E-04 |
| Adgre5 | skyblue | -1.7184481 | 7.91E-12 | -1.3569631 | 1.40E-07 |
| Il17ra | yellow | 1.96390946 | 6.08E-19 | 2.04864795 | 2.05E-20 |
| Kcnn1 | red | -1.0858328 | 0.00340955 | -1.1212048 | 0.00571437 |
| Arrdc2 | orange | 0.66185281 | 0.00011469 | -0.0875434 | 0.67342967 |
| Cd36 | red | -1.2181073 | 2.11E-09 | -1.0567978 | 3.78E-07 |
| Bcam | red | -0.622308 | 7.20E-05 | -0.4464289 | 0.00635042 |
| Relb | orange | 1.5688286 | 2.38E-11 | 1.5217 | 1.59E-10 |
| Tomm40 | orange | 0.7990841 | 4.28E-08 | 0.62216767 | 3.71E-05 |
| Hbp1 | red | -0.7161276 | 0.00032405 | -0.5603927 | 0.00670953 |
| Prkar2b | skyblue | -1.7138995 | 4.59E-09 | -1.4515505 | 1.30E-06 |
| Cdkn1b | red | -0.622768 | 0.00063239 | -0.5522104 | 0.0033082 |
| Rab8a | orange | 0.94531292 | 5.92E-10 | 0.74784562 | 1.95E-06 |
| Hmgn2 | orange | 0.74023807 | 3.66E-08 | 0.57438705 | 3.50E-05 |
| Cnot11 | orange | 0.59281099 | 4.64E-06 | 0.43393269 | 0.00125592 |
| Slc2a3 | orange | 1.04849294 | 2.08E-14 | 0.500045 | 0.0005753 |
| Ebi3 | yellow | 1.76095866 | 9.24E-05 | 2.06487827 | 2.27E-05 |
| Edar | red | -1.1660795 | 0.00584514 | -0.0038965 | 0.99385519 |
| Car11 | skyblue | -2.1609064 | 2.62E-14 | -1.9430145 | 3.45E-12 |
| Hck | yellow | 2.12558027 | 1.77E-12 | 1.93139016 | 2.86E-10 |
| Ap1m2 | red | -0.6158221 | 0.00099318 | -0.2764812 | 0.17526073 |

| | | | | | |
|---|---|---|---|---|---|
| Mob3a | orange | 1.15943103 | 4.89E-06 | 1.05951798 | 4.61E-05 |
| Cacnb3 | orange | 0.77620198 | 4.08E-05 | 0.71955597 | 0.00021896 |
| Ccdc65 | skyblue | -1.6275461 | 4.38E-09 | -1.1696736 | 3.14E-05 |
| Pld3 | orange | 0.68326351 | 2.46E-05 | 0.48410832 | 0.00417317 |
| St8sia6 | skyblue | -1.6640615 | 2.63E-05 | -0.7094797 | 0.09181042 |
| Fcgrt | red | -0.8466156 | 4.23E-07 | -0.7657637 | 8.15E-06 |
| Rps11 | orange | 0.67812667 | 0.00047424 | 0.4927472 | 0.01519356 |
| Phyhip | skyblue | -1.2263351 | 0.00012632 | -1.2709809 | 0.00012635 |
| Crhr2 | skyblue | -1.4637578 | 1.05E-05 | -1.2964534 | 0.0001901 |
| Inmt | skyblue | -2.2984706 | 1.03E-05 | -1.8909459 | 0.00044595 |
| Cyp4f18 | yellow | 4.02440329 | 4.07E-21 | 3.40740941 | 2.91E-15 |
| Impdh1 | red | -0.6748478 | 9.31E-09 | -0.6475494 | 6.41E-08 |
| Dusp3 | red | -0.7849959 | 1.74E-06 | -0.6568598 | 0.00010374 |
| Prodh | red | -1.0738231 | 0.00276502 | -0.760746 | 0.04316857 |
| Slc25a1 | red | -0.3546491 | 0.12890946 | -0.62025 | 0.00697151 |
| Homer3 | red | -0.7244573 | 9.33E-14 | -0.6207599 | 4.53E-10 |
| Rnf215 | red | -0.7826216 | 3.62E-07 | -0.5356003 | 0.00081505 |
| Sec14l2 | red | -1.2779483 | 6.91E-16 | -0.9830983 | 7.04E-10 |
| Cp | orange | 0.51636982 | 0.00048944 | 0.61231196 | 4.56E-05 |
| Rps6ka1 | orange | 0.67490046 | 1.25E-11 | 0.27665 | 0.00931167 |
| Has1 | orange | 1.3886948 | 0.00435064 | 2.48411867 | 4.74E-07 |
| Itpkc | orange | 0.77237287 | 0.00019048 | 0.6277805 | 0.00342738 |
| Kif20a | yellow | 2.74955221 | 8.86E-25 | 2.71693615 | 1.24E-23 |
| Farsa | orange | 0.80461085 | 1.65E-10 | 0.6988219 | 5.73E-08 |
| Dnase2a | orange | 1.1481101 | 1.10E-07 | 0.97196433 | 1.40E-05 |
| Syce2 | orange | 1.36403542 | 1.49E-07 | 0.81149792 | 0.00290512 |
| Nqo1 | red | -1.060672 | 1.29E-05 | -0.4887044 | 0.0600515 |
| Gys1 | red | -0.912748 | 1.78E-08 | -0.5425176 | 0.00142347 |
| Bax | orange | 0.8784917 | 1.49E-05 | 0.67253089 | 0.00140521 |
| Il7r | yellow | 2.24201949 | 3.01E-14 | 1.61902291 | 8.56E-08 |
| Zfp81 | red | -0.4010569 | 0.0651282 | -0.5830466 | 0.00717665 |
| Efnb3 | red | -0.2446738 | 0.59777036 | -1.1441982 | 0.00799838 |
| Mmd | red | -0.3597896 | 0.03586152 | -0.6540355 | 0.00011999 |
| Hlf | skyblue | -2.3617798 | 5.09E-13 | -2.1068349 | 2.16E-10 |
| Fam162a | red | -0.6446793 | 7.73E-07 | -0.5204247 | 0.00013172 |
| Ssbp2 | red | -0.5805315 | 0.0060504 | -0.3023456 | 0.1874343 |
| Gstm5 | red | -0.5478984 | 0.03212475 | -0.9299228 | 0.00029859 |
| Gstm7 | red | -1.3581931 | 0.00049254 | -0.9304428 | 0.02373254 |
| Stat3 | orange | 0.93703135 | 3.62E-14 | 0.96129243 | 1.00E-14 |
| Stat5a | orange | 0.6398536 | 3.29E-07 | 0.76192547 | 1.73E-09 |
| Ptrf | red | -0.8918728 | 1.30E-11 | -0.716475 | 1.15E-07 |
| Zak | red | -0.9246649 | 9.95E-08 | -0.6860956 | 0.00013724 |
| Col5a3 | red | -0.9161141 | 3.22E-05 | -0.7685358 | 0.00081829 |

| | | | | | |
|---|---|---|---|---|---|
| Dnmt1 | orange | 1.09903747 | 3.41E-14 | 0.95405828 | 9.46E-11 |
| Ppan | orange | 0.8471781 | 0.000139 | 0.87316 | 0.00012733 |
| Angptl2 | skyblue | -2.0078688 | 6.30E-21 | -1.6607001 | 1.78E-14 |
| Etv1 | skyblue | -1.2352793 | 0.00015864 | -1.5244295 | 4.13E-06 |
| Kifc2 | skyblue | -1.5436899 | 4.01E-06 | -1.2304582 | 0.00036749 |
| Psap | orange | 0.73838035 | 0.00032475 | 0.53446498 | 0.01276852 |
| Phb2 | orange | 0.59789203 | 2.32E-05 | 0.45201668 | 0.0021007 |
| Ptpn6 | yellow | 2.28417502 | 6.87E-28 | 2.20273981 | 7.57E-26 |
| Il12b | yellow | 2.59677255 | 0.000228 | 1.31686307 | 0.07055532 |
| Clcn5 | orange | 0.9175304 | 7.55E-06 | 1.09968566 | 1.19E-07 |
| Hif3a | orange | 1.92741888 | 8.25E-24 | 0.5873196 | 0.00335275 |
| Pde1c | skyblue | -0.7831226 | 0.25009345 | -2.6270314 | 0.00011257 |
| Spic | yellow | 1.84308523 | 0.00545182 | 2.19102391 | 0.00098898 |
| 9330159F19I | skyblue | -1.8753224 | 2.25E-06 | -1.4762713 | 0.00037741 |
| Il11 | orange | 1.46403802 | 0.00064088 | 1.47824737 | 0.00077926 |
| Bid | orange | 1.87895197 | 1.89E-18 | 1.94544358 | 7.92E-20 |
| Dnajb11 | orange | 0.83365619 | 1.15E-06 | 0.61461759 | 0.00056086 |
| Gab2 | orange | 0.61166882 | 4.36E-08 | 0.44990851 | 9.54E-05 |
| Ctse | yellow | 2.45888567 | 1.48E-09 | 2.53291721 | 5.53E-10 |
| Ndrg2 | red | -1.1525215 | 0.00011305 | -1.0636781 | 0.00053438 |
| Arhgef40 | red | -0.590778 | 4.61E-07 | -0.4219159 | 0.0005295 |
| Cd33 | orange | 1.32635508 | 0.00011303 | 1.01516142 | 0.00444018 |
| Nkg7 | black | 4.59832707 | 5.79E-72 | 4.33235126 | 1.98E-62 |
| Stxbp2 | orange | 0.72343289 | 1.23E-06 | 0.66003194 | 1.62E-05 |
| Sgce | red | -0.8639552 | 6.74E-09 | -0.750252 | 9.10E-07 |
| Slbp | orange | 0.9752449 | 7.36E-13 | 0.71526745 | 3.60E-07 |
| Aqp1 | red | -1.2413124 | 1.76E-16 | -1.0213003 | 2.84E-11 |
| Myo9b | orange | 0.71944532 | 2.45E-06 | 0.62191876 | 7.57E-05 |
| Ly9 | yellow | 2.71556393 | 1.94E-23 | 2.42410262 | 1.53E-18 |
| Cd244 | yellow | 2.96674007 | 1.39E-13 | 2.805945 | 5.06E-12 |
| Adgre1 | orange | 1.77178329 | 0.00210365 | 1.464855 | 0.01443111 |
| Mtfp1 | skyblue | -1.3636188 | 3.15E-07 | -1.5375534 | 1.72E-08 |
| Pgf | orange | 1.63017753 | 2.15E-14 | 1.80461 | 4.44E-17 |
| Ulk2 | red | -0.9419357 | 6.20E-06 | -0.7372769 | 0.00064537 |
| Ccl24 | yellow | 2.8468525 | 1.79E-09 | 2.45850137 | 2.12E-07 |
| Plod3 | orange | 0.71101203 | 7.66E-07 | 0.62108409 | 2.66E-05 |
| Ap1s1 | orange | 0.72276504 | 0.00062755 | 0.55126025 | 0.01246687 |
| Lbr | orange | 0.60803959 | 7.56E-08 | 0.61980575 | 6.74E-08 |
| Bcan | skyblue | -2.4552676 | 1.81E-14 | -1.2863466 | 8.45E-05 |
| Matk | yellow | 2.38607302 | 4.56E-14 | 2.09591249 | 7.03E-11 |
| Nmrk2 | skyblue | -1.6621408 | 0.00152962 | -0.8705517 | 0.15470178 |
| Rasa4 | orange | 1.77786201 | 2.73E-22 | 1.46898603 | 2.45E-15 |
| Syt5 | skyblue | -1.5205772 | 3.97E-10 | -1.218616 | 1.17E-06 |

| | | | | | |
|---|---|---|---|---|---|
| Sgsh | red | -0.8450335 | 1.73E-08 | -0.8122331 | 9.59E-08 |
| Chd5 | red | -1.3711889 | 0.00042856 | -0.9783777 | 0.01670933 |
| Cstb | orange | 1.67320644 | 4.19E-11 | 1.55130251 | 1.80E-09 |
| Sh2b2 | orange | 1.17682853 | 0.00180716 | 0.7697428 | 0.05316345 |
| Cd44 | orange | 1.06189785 | 1.08E-06 | 1.0515958 | 2.21E-06 |
| Wdr1 | orange | 0.92048328 | 1.67E-07 | 0.89985465 | 5.21E-07 |
| Slc2a9 | orange | 0.62817989 | 3.34E-05 | 0.4687786 | 0.00295515 |
| 4930550C14 | red | -1.3542168 | 0.00082246 | -1.1024744 | 0.00871465 |
| Man2b1 | orange | 0.867267 | 4.26E-05 | 0.52918598 | 0.01789837 |
| Spc25 | orange | 1.70561522 | 1.02E-13 | 1.59805675 | 8.72E-12 |
| Dnah2 | red | -1.1158472 | 2.87E-10 | -0.7469533 | 4.46E-05 |
| Ripk4 | red | -0.7308372 | 9.64E-07 | -0.3428302 | 0.03060396 |
| Fgfr4 | skyblue | -2.0907495 | 3.96E-15 | -1.6996996 | 3.68E-10 |
| Cadm3 | red | -1.4653268 | 1.37E-13 | -0.3280849 | 0.11981253 |
| Slc1a3 | red | -1.135299 | 1.39E-05 | -0.938647 | 0.00050291 |
| Mlxipl | skyblue | -2.3399896 | 5.92E-08 | -1.7278693 | 0.00010496 |
| Mcm5 | yellow | 2.46325542 | 2.46E-26 | 2.28998628 | 1.17E-22 |
| Hmox1 | orange | 1.51821007 | 3.86E-08 | 1.77447803 | 1.64E-10 |
| Il27ra | orange | 1.57040112 | 7.43E-19 | 1.56332358 | 2.94E-18 |
| Asf1b | yellow | 2.97210021 | 1.50E-32 | 2.48102502 | 1.06E-22 |
| Myl10 | skyblue | -2.8772678 | 0.0014569 | -0.9703596 | 0.2970046 |
| Ddx39 | orange | 1.15376848 | 3.45E-19 | 0.74574895 | 2.20E-08 |
| Por | red | -0.9181607 | 3.50E-08 | -0.7276221 | 2.24E-05 |
| Igf1r | red | -0.7537563 | 6.25E-07 | -0.6078426 | 9.91E-05 |
| Fcer2a | red | -0.9014767 | 0.00466815 | -0.2597894 | 0.48094338 |
| Cyp2a5 | skyblue | -2.6857159 | 2.67E-07 | -2.8131827 | 1.09E-07 |
| Ube2m | orange | 0.59522965 | 0.00011523 | 0.53891188 | 0.00070956 |
| Adcy9 | red | -0.9958386 | 0.00098155 | -0.6767427 | 0.03300864 |
| Mef2c | red | -1.0813746 | 3.17E-08 | -0.74272 | 0.00027832 |
| Mrvi1 | red | -1.0150802 | 1.72E-08 | -0.8867756 | 1.20E-06 |
| Psmd4 | orange | 0.81478838 | 1.27E-08 | 0.59410018 | 6.27E-05 |
| Tmod4 | red | -1.1892694 | 0.000345 | -0.2912889 | 0.45388119 |
| Insrr | yellow | 3.19658242 | 3.25E-13 | 2.45435058 | 1.80E-09 |
| Mthfd2 | yellow | 2.99490762 | 1.48E-28 | 3.10162473 | 4.70E-30 |
| Kit | red | -1.3499063 | 3.21E-09 | -1.1289714 | 1.39E-06 |
| Ampd3 | orange | 0.92261454 | 1.59E-06 | 0.67326194 | 0.00073068 |
| Sh2d1a | yellow | 2.97259694 | 8.23E-07 | 4.02164079 | 3.18E-10 |
| Pvalb | skyblue | -2.0922892 | 3.32E-05 | -1.3579152 | 0.01061247 |
| Ranbp1 | orange | 1.10076226 | 2.88E-06 | 0.85089741 | 0.00050977 |
| Cd247 | yellow | 2.20108056 | 7.58E-21 | 2.70906667 | 4.73E-30 |
| Psmb4 | orange | 1.03756162 | 1.22E-08 | 0.86719129 | 3.60E-06 |
| Mmp8 | black | 4.84208227 | 2.21E-09 | 3.60315342 | 1.28E-05 |
| Tnfsf14 | yellow | 2.9141841 | 6.16E-17 | 2.69931135 | 4.12E-15 |

| | | | | | |
|---|---|---|---|---|---|
| Gata6 | red | -0.7860224 | 1.37E-08 | -0.5740818 | 6.10E-05 |
| Rsl1d1 | orange | 0.6494215 | 2.70E-05 | 0.43501096 | 0.00737904 |
| Spo11 | yellow | -0.0248985 | 0.98753802 | 5.72613793 | 5.23E-05 |
| Nr2c1 | red | -0.5966702 | 3.32E-06 | -0.3430085 | 0.0111214 |
| Otx1 | red | -1.2114869 | 0.00937356 | -0.8129464 | 0.08921787 |
| Itgae | orange | 1.32784524 | 7.50E-07 | 0.70053404 | 0.01493997 |
| P2rx5 | red | -1.3380665 | 1.69E-06 | -0.7789193 | 0.00836291 |
| Ephb3 | red | -0.8977713 | 8.22E-10 | -0.4440478 | 0.00400605 |
| Calcoco2 | skyblue | -3.9296267 | 0.00041713 | -4.6636082 | 0.00035256 |
| Snf8 | orange | 0.60121219 | 6.44E-05 | 0.37265532 | 0.01947137 |
| Tbcb | orange | 0.79018077 | 9.26E-08 | 0.49154727 | 0.0015166 |
| Inpp5k | red | -0.5646891 | 0.00298822 | -0.6037068 | 0.0018673 |
| Eps8l1 | red | -1.1007334 | 0.00042959 | -0.7501095 | 0.0222085 |
| Htra1 | red | -1.1645399 | 7.32E-05 | -0.57749 | 0.0671227 |
| Epor | skyblue | -2.60582 | 3.07E-10 | -1.6777218 | 7.79E-05 |
| Ccdc159 | red | -1.1669313 | 0.00193254 | -0.8463609 | 0.02824091 |
| Atp6v1b1 | skyblue | -4.4687611 | 5.06E-12 | -2.3459606 | 0.00017431 |
| Tep1 | orange | 0.98838693 | 6.47E-08 | 0.81591099 | 1.49E-05 |
| Arpc2 | orange | 1.07367644 | 6.45E-08 | 0.89816434 | 1.10E-05 |
| Zbtb32 | yellow | 2.70124508 | 8.53E-28 | 2.65519293 | 8.55E-27 |
| Susd2 | red | -1.2948322 | 6.23E-14 | -0.8021664 | 6.53E-06 |
| Ggt5 | orange | 0.20000756 | 0.25384648 | 0.5884657 | 0.00042144 |
| Ggt1 | orange | 1.83070369 | 3.53E-09 | 1.7918809 | 8.31E-09 |
| Crip2 | red | -0.910323 | 3.82E-06 | -0.7554208 | 0.00020317 |
| Cbfa2t3 | red | -1.2409841 | 5.64E-11 | -1.0789503 | 2.32E-08 |
| Fbln1 | skyblue | -1.4827617 | 2.67E-10 | -1.1342239 | 2.75E-06 |
| Pgrmc1 | red | -0.9177827 | 8.08E-13 | -0.7767864 | 2.92E-09 |
| Tek | red | -1.2052115 | 1.77E-15 | -1.0964068 | 8.34E-13 |
| Cdc20 | yellow | 2.52189527 | 4.43E-36 | 2.45734524 | 1.21E-33 |
| Adamts4 | black | 5.07188798 | 5.84E-26 | 5.6621652 | 1.88E-30 |
| Rnf114 | orange | 1.36990814 | 1.28E-34 | 1.27195396 | 8.14E-30 |
| Neurl1a | red | -1.3470689 | 1.37E-06 | -0.961757 | 0.00097534 |
| Srm | orange | 1.05635421 | 0.0001125 | 1.01606733 | 0.00030572 |
| Epha2 | orange | 1.16399931 | 5.11E-07 | 1.24769309 | 1.09E-07 |
| Bbs1 | red | -1.0790212 | 1.14E-08 | -0.8259575 | 1.70E-05 |
| Ptbp1 | orange | 0.58529366 | 1.26E-05 | 0.61924677 | 5.62E-06 |
| Mvd | orange | 0.88845664 | 0.00023248 | 0.96852611 | 8.38E-05 |
| Cyba | yellow | 2.15677498 | 4.58E-14 | 1.66436689 | 1.39E-08 |
| Prkag3 | red | -1.0437997 | 0.0048907 | -0.2676541 | 0.53428635 |
| Atp7b | skyblue | -1.7824392 | 0.00171949 | -1.2366565 | 0.03860406 |
| Slc4a3 | red | -1.004569 | 0.00012369 | -0.7753388 | 0.00427972 |
| Cdt1 | yellow | 2.50215479 | 1.98E-24 | 2.00040469 | 7.92E-16 |
| Runx1t1 | red | -1.017632 | 3.05E-05 | -0.8231256 | 0.00119246 |

| | | | | | |
|---|---|---|---|---|---|
| Aprt | orange | 1.44917985 | 1.70E-10 | 1.17770245 | 4.45E-07 |
| Slc5a6 | red | -0.9970808 | 3.23E-05 | -0.9663198 | 8.12E-05 |
| Tcf23 | orange | 1.75393994 | 2.31E-25 | 0.8229949 | 2.09E-06 |
| P4htm | red | -0.8938257 | 2.68E-05 | -0.9009045 | 3.10E-05 |
| Pola1 | orange | 0.89781891 | 6.00E-06 | 0.96264735 | 2.66E-06 |
| D130043K22 | skyblue | -1.7321631 | 1.70E-07 | -0.9655028 | 0.00159845 |
| Gmnn | orange | 1.35365302 | 4.82E-06 | 0.92632817 | 0.00284447 |
| Acot13 | red | -0.5959074 | 0.00010192 | -0.8127127 | 2.20E-07 |
| Zfp184 | red | -1.3803182 | 0.00591516 | -0.9199383 | 0.05645312 |
| B4galnt1 | orange | 1.51737645 | 2.17E-11 | 1.2499184 | 7.37E-08 |
| Krt23 | red | -0.7986004 | 0.00353942 | -0.5206853 | 0.07359249 |
| Cnp | orange | 0.76557252 | 2.36E-07 | 0.76121369 | 4.63E-07 |
| Ttc25 | skyblue | -1.6739376 | 1.20E-06 | -0.9081658 | 0.01130202 |
| Ezh1 | red | -0.6554894 | 3.64E-07 | -0.576293 | 1.34E-05 |
| Hap1 | orange | 1.49315021 | 1.38E-25 | 1.668376 | 1.35E-31 |
| P3h4 | red | -0.7123161 | 4.38E-07 | -0.5269142 | 0.00033319 |
| Ctnnb1 | red | -0.6905331 | 8.46E-10 | -0.4549955 | 0.0001016 |
| Psmd2 | orange | 0.58151303 | 1.06E-05 | 0.56293802 | 3.14E-05 |
| Vars | orange | 1.14215818 | 1.24E-14 | 1.0547309 | 1.93E-12 |
| Vwa7 | red | -0.7602383 | 0.00056823 | -0.4078063 | 0.09360576 |
| Slc44a4 | red | -1.2796068 | 4.02E-08 | -0.6644165 | 0.0065168 |
| Ddah2 | red | -0.7725031 | 3.57E-05 | -0.6205622 | 0.0013622 |
| Clic1 | orange | 1.51825831 | 7.99E-13 | 1.30998255 | 1.33E-09 |
| Lsm2 | orange | 0.83723247 | 0.0002264 | 0.94585268 | 4.77E-05 |
| Pole | yellow | 2.79877326 | 6.25E-24 | 2.75945935 | 5.79E-23 |
| Atp1a2 | red | -1.0811511 | 8.78E-07 | -0.9804665 | 1.39E-05 |
| Casq1 | skyblue | -1.9220242 | 3.70E-08 | -1.1782183 | 0.00129991 |
| Zfp775 | red | -0.509405 | 0.00337113 | -0.5839711 | 0.00098223 |
| Scube2 | skyblue | -2.5750721 | 2.22E-06 | -1.8108521 | 0.00136111 |
| M6pr | orange | 0.60188858 | 3.95E-05 | 0.47479524 | 0.0017526 |
| Tinf2 | orange | 0.58451867 | 3.59E-10 | 0.61650698 | 7.05E-11 |
| Cav1 | red | -1.0890916 | 7.42E-06 | -1.0019667 | 5.89E-05 |
| Bcl2l1 | orange | 1.455173 | 1.73E-18 | 0.93269567 | 5.51E-08 |
| Dio2 | orange | 1.2319428 | 0.00988051 | 2.52731956 | 1.18E-07 |
| Hnrnpa0 | orange | 0.47694278 | 0.00199301 | 0.64068205 | 3.85E-05 |
| Ift43 | red | -0.5983049 | 1.41E-08 | -0.4665886 | 2.11E-05 |
| Tcap | skyblue | -1.7428325 | 8.00E-06 | -1.4496852 | 0.00032394 |
| Crlf1 | red | -1.4227223 | 1.25E-09 | -0.6818943 | 0.00571612 |
| Hmgcll1 | red | -1.1165634 | 1.17E-10 | -0.8641596 | 1.08E-06 |
| Fzd3 | skyblue | -1.358928 | 1.15E-06 | -1.2594392 | 1.02E-05 |
| Mid1ip1 | red | -0.8358137 | 2.44E-09 | -0.7407091 | 2.38E-07 |
| Ap2s1 | orange | 1.00088566 | 2.04E-14 | 0.68022074 | 5.42E-07 |
| Fgfrl1 | red | -0.9147558 | 0.00030532 | -0.7778949 | 0.00297336 |

| | | | | | |
|---|---|---|---|---|---|
| Fhl2 | red | -0.3946902 | 0.27020049 | -0.9209074 | 0.00756585 |
| Clstn3 | skyblue | -3.4486704 | 7.73E-11 | -2.7063528 | 7.83E-07 |
| Fnbp4 | orange | 1.00111597 | 2.23E-16 | 0.91904902 | 9.10E-14 |
| Cers4 | red | -0.9438649 | 2.87E-10 | -0.8010113 | 1.72E-07 |
| Scrn3 | skyblue | -1.4215574 | 3.08E-09 | -1.1729368 | 1.83E-06 |
| 1700109H08 | orange | 0.91749124 | 0.00530109 | 0.9813464 | 0.0037562 |
| Relt | yellow | 1.92751497 | 2.04E-17 | 2.20133602 | 8.45E-22 |
| Snrpb2 | orange | 0.92938826 | 1.62E-06 | 0.70273393 | 0.0005131 |
| Ubc | orange | 0.60177448 | 1.33E-06 | 0.40689006 | 0.00171004 |
| Prpf31 | orange | 0.90844616 | 4.59E-08 | 0.79738607 | 3.01E-06 |
| Sertad1 | orange | 1.17338263 | 1.52E-10 | 0.90134311 | 1.80E-06 |
| Arpc5 | orange | 1.03582862 | 2.38E-08 | 0.82563648 | 1.60E-05 |
| Pou2f2 | yellow | 2.21101839 | 2.41E-12 | 2.07558204 | 1.00E-10 |
| Mgst1 | red | -1.0280003 | 1.90E-06 | -0.697274 | 0.00194532 |
| Nfib | red | -0.6465503 | 1.89E-05 | -0.7116881 | 3.68E-06 |
| Rab25 | red | -0.6821699 | 0.00062742 | -0.4336972 | 0.04043511 |
| Rbfox1 | skyblue | -1.3682055 | 0.00104775 | -1.4405306 | 0.00089762 |
| Rps18 | orange | 0.69062166 | 2.61E-08 | 0.6273019 | 7.68E-07 |
| Rps15a | orange | 0.69230017 | 3.55E-08 | 0.638788 | 6.51E-07 |
| Acyp1 | red | -0.7894536 | 0.00122065 | -0.7890825 | 0.00173272 |
| Hdac5 | red | -0.6858916 | 5.36E-08 | -0.6056216 | 2.75E-06 |
| Bmp7 | red | -0.8796454 | 0.00298908 | -0.0193217 | 0.96091109 |
| Rsph14 | skyblue | -1.6761077 | 8.69E-06 | -1.0352976 | 0.00667913 |
| Tbx1 | red | -1.0196139 | 3.89E-05 | -0.7149712 | 0.00575599 |
| 2610028H24 | skyblue | -1.5800232 | 3.91E-09 | -0.7361707 | 0.00924512 |
| Dqx1 | red | -0.8388125 | 0.00073289 | -0.7370762 | 0.00376711 |
| Ccl8 | black | 5.57912962 | 2.08E-62 | 5.90746372 | 6.68E-58 |
| Tmem8c | yellow | 2.92294863 | 0.00391944 | 3.34885549 | 0.00075327 |
| Trpm2 | yellow | 4.40778166 | 1.33E-23 | 3.66920615 | 8.58E-17 |
| Met | red | -1.0588691 | 5.84E-09 | -0.5958704 | 0.00179416 |
| Slc16a12 | skyblue | -1.6875234 | 6.37E-08 | -1.1023941 | 0.00052038 |
| Syn2 | red | -0.7433081 | 0.0050472 | -0.7714755 | 0.00491328 |
| Myod1 | skyblue | -1.6050856 | 0.00910399 | -1.1634216 | 0.08677743 |
| Kcnq1 | skyblue | -1.3942052 | 2.15E-11 | -1.1278739 | 1.15E-07 |
| Tor2a | orange | 0.82401164 | 1.28E-07 | 0.6174729 | 0.00013137 |
| Apobec3 | yellow | 2.18573663 | 1.14E-108 | 1.92112062 | 6.16E-84 |
| Tex15 | red | -1.3745431 | 3.38E-05 | -0.6147954 | 0.08939315 |
| G0s2 | red | -1.2119558 | 3.08E-11 | -0.802239 | 2.10E-05 |
| Tex11 | skyblue | -2.6067096 | 7.73E-06 | -1.8980245 | 0.0011926 |
| Fxyd5 | yellow | 2.46144638 | 5.38E-27 | 2.11831271 | 4.34E-20 |
| Tfcp2 | red | -0.6629079 | 3.35E-05 | -0.4856234 | 0.00343852 |
| Nuak2 | yellow | 2.37505134 | 2.81E-36 | 2.02603266 | 1.19E-26 |
| Cox4i2 | red | -0.7929839 | 9.45E-06 | -0.8551324 | 2.71E-06 |

| | | | | | |
|---|---|---|---|---|---|
| Snap47 | red | -0.6229169 | 2.31E-06 | -0.731541 | 4.04E-08 |
| Wnt3a | skyblue | -2.2469952 | 4.12E-09 | -1.5634205 | 7.65E-05 |
| Kif19a | red | -1.4346221 | 2.38E-13 | -0.6371774 | 0.00159781 |
| Zmynd10 | skyblue | -1.6765708 | 2.92E-10 | -0.7617189 | 0.00599553 |
| Slc38a3 | skyblue | -1.8649718 | 7.88E-09 | -1.567314 | 4.24E-06 |
| Rassf1 | orange | 0.86091077 | 1.21E-06 | 0.80852615 | 8.21E-06 |
| Rnf112 | skyblue | -3.2195234 | 0.00077501 | -1.6242879 | 0.09546606 |
| Slc47a1 | skyblue | -1.7470846 | 0.00044315 | -1.4957422 | 0.00353992 |
| Pifo | skyblue | -1.9568023 | 1.07E-14 | -1.5343636 | 1.80E-09 |
| Tnfrsf13b | orange | 1.20963595 | 2.66E-06 | 1.03418169 | 0.00010311 |
| AI597479 | red | -0.6226528 | 4.07E-08 | -0.4407698 | 0.00019387 |
| Tmem86a | orange | 1.00635426 | 0.00382504 | 0.60299044 | 0.10559857 |
| Optc | skyblue | -3.010936 | 1.08E-07 | -2.0867077 | 0.00019153 |
| Ifi35 | orange | 1.62680867 | 6.43E-33 | 1.45832843 | 2.37E-26 |
| Rdm1 | skyblue | -1.2738675 | 1.64E-07 | -1.2664578 | 3.33E-07 |
| Mrpl52 | orange | 0.86551021 | 4.26E-06 | 0.88404836 | 4.55E-06 |
| Ebf3 | skyblue | -1.7009655 | 0.00361302 | -0.8099884 | 0.20013377 |
| Uckl1os | red | -1.4054416 | 2.45E-06 | -0.9218415 | 0.00235985 |
| Apol7a | yellow | 2.91116898 | 1.37E-16 | 3.14641989 | 1.47E-19 |
| Psen2 | orange | 1.22817786 | 4.80E-08 | 0.69343901 | 0.00335341 |
| Acaa1b | skyblue | -1.8789505 | 5.01E-07 | -1.5657628 | 4.18E-05 |
| Fads1 | red | -0.6837731 | 6.28E-12 | -0.4835319 | 2.56E-06 |
| Lmbr1 | red | -0.745343 | 1.35E-06 | -0.4233818 | 0.00874184 |
| Tnfrsf22 | red | -0.7921418 | 4.05E-05 | -0.5990878 | 0.00279067 |
| Cars | orange | 0.59284075 | 4.82E-09 | 0.64201922 | 3.74E-10 |
| Wnt2 | red | -1.3462486 | 4.26E-09 | -1.1069121 | 2.76E-06 |
| Grid2ip | skyblue | -1.3823918 | 0.00993075 | -1.2412934 | 0.02417584 |
| Vac14 | orange | 0.6747288 | 1.18E-05 | 0.56565731 | 0.00038508 |
| Mcoln2 | orange | 1.98769458 | 5.81E-07 | 1.94115935 | 1.39E-06 |
| Tbrg1 | orange | 0.75187742 | 9.59E-10 | 0.71611154 | 9.73E-09 |
| 1700026D08 | skyblue | -2.0387589 | 1.16E-11 | -1.4696129 | 1.50E-06 |
| Odc1 | orange | 0.89902955 | 3.65E-07 | 0.91558294 | 3.57E-07 |
| Thg1l | orange | 0.69014637 | 0.00163922 | 0.61243359 | 0.00711832 |
| Adam19 | orange | 1.1094896 | 5.14E-06 | 0.99659052 | 6.82E-05 |
| Exoc3l2 | orange | 1.15239809 | 0.00289482 | 0.88590964 | 0.02957005 |
| Zfp296 | orange | 0.84876723 | 0.00321835 | 0.46641026 | 0.14455211 |
| Plin5 | skyblue | -1.1577322 | 0.00013314 | -1.3713162 | 1.14E-05 |
| Dhdh | red | -1.1788401 | 2.08E-13 | -1.0772609 | 3.85E-11 |
| Zfp790 | red | -0.6571795 | 6.49E-06 | -0.5879203 | 8.46E-05 |
| Cpb1 | skyblue | -2.1871554 | 0.00313996 | 0.13258702 | 0.88354444 |
| Pinlyp | red | -1.0155093 | 0.00708528 | -0.5889527 | 0.14820569 |
| Fuz | red | -0.5631705 | 0.00043705 | -0.5999398 | 0.00025281 |
| Evi5 | red | -0.6539109 | 3.37E-06 | -0.6089497 | 2.44E-05 |

| | | | | | |
|---|---|---|---|---|---|
| Git1 | orange | 0.67436995 | 7.80E-05 | 0.57790079 | 0.00103415 |
| Scarf2 | red | -0.6794961 | 0.00081956 | -0.3967077 | 0.06711942 |
| Med15 | orange | 0.70371577 | 1.45E-05 | 0.57038199 | 0.00067517 |
| Aim1l | red | -0.8873727 | 8.08E-05 | -0.8896597 | 0.0001121 |
| Ehf | red | -0.7088015 | 0.00343133 | -0.4408224 | 0.08756236 |
| Tmem167 | orange | 0.81702937 | 8.93E-08 | 0.69013805 | 1.20E-05 |
| Kif11 | yellow | 3.07724319 | 1.89E-18 | 3.60431246 | 1.45E-23 |
| Mlkl | yellow | 3.29323542 | 2.92E-31 | 3.2343938 | 4.30E-30 |
| Ttll5 | red | -0.7520154 | 4.01E-05 | -0.5533626 | 0.00344698 |
| Retn | skyblue | -3.4333667 | 5.44E-08 | -1.4602609 | 0.02603587 |
| Cdh23 | orange | 1.12963872 | 0.00610173 | 1.35556708 | 0.00157139 |
| Podnl1 | yellow | 3.00353596 | 2.88E-18 | 2.49671608 | 6.06E-13 |
| Adgrl1 | red | -0.7937719 | 2.39E-13 | -0.7402085 | 1.47E-11 |
| Etv5 | red | -1.3809244 | 4.59E-09 | -0.6965656 | 0.00512895 |
| Ptprs | red | -0.7177196 | 1.71E-06 | -0.663625 | 1.58E-05 |
| Fer1l4 | red | -1.3818716 | 0.00139231 | -0.7670784 | 0.04975381 |
| Zfp651 | red | -0.900173 | 2.59E-07 | -0.6827726 | 0.00016175 |
| Bcas1 | skyblue | -1.5208064 | 2.39E-08 | -1.2429834 | 6.47E-06 |
| Snx31 | skyblue | -2.3460407 | 0.00983461 | -1.0230194 | 0.22021588 |
| Cad | orange | 0.65458009 | 0.00081643 | 0.53642669 | 0.00823969 |
| Lypd8 | black | 6.00009854 | 4.04E-05 | 3.82666514 | 1.71E-05 |
| Timm23 | orange | 0.65091634 | 0.00280186 | 0.5605344 | 0.01316017 |
| Tnfaip8l2 | yellow | 2.50548451 | 1.75E-20 | 2.27549733 | 6.84E-17 |
| Ly6g6e | red | -1.1362436 | 0.00866942 | -0.8277164 | 0.07324777 |
| St3gal1 | orange | 0.62483161 | 6.32E-05 | 0.41439059 | 0.01137191 |
| Clip3 | red | -0.933386 | 2.40E-14 | -0.8681761 | 1.35E-12 |
| Tctex1d2 | orange | 0.66660623 | 0.00167705 | 0.45284891 | 0.04538179 |
| Zfp385c | skyblue | -1.631575 | 2.49E-05 | -1.3237126 | 0.00138941 |
| Glis2 | red | -1.0092706 | 1.45E-19 | -0.5729069 | 8.01E-07 |
| Bicc1 | red | -0.7766252 | 7.88E-10 | -0.6439402 | 7.18E-07 |
| Ankrd28 | orange | 0.58169017 | 0.00039129 | 0.66852835 | 5.94E-05 |
| Klra17 | black | 5.66150287 | 0.00018179 | 1.44371556 | 0.16566181 |
| Csf1 | orange | 1.05617862 | 1.54E-12 | 1.16291223 | 7.07E-15 |
| Chrne | orange | 1.97958135 | 0.00343671 | 1.39585045 | 0.04878127 |
| Chfr | orange | 0.71424315 | 9.97E-16 | 0.70526734 | 4.83E-15 |
| Ceacam16 | black | 5.42255002 | 7.51E-13 | 3.98731917 | 3.29E-09 |
| Adam28 | skyblue | -1.6953806 | 1.16E-07 | -0.7335222 | 0.0302225 |
| Psmb1 | orange | 0.90969863 | 3.44E-13 | 0.63600105 | 9.14E-07 |
| Nol3 | red | -0.6266252 | 0.00139319 | -0.2895356 | 0.17928083 |
| Plekhg4 | yellow | 2.77415633 | 8.40E-09 | 1.80743483 | 0.00013507 |
| Tppp3 | skyblue | -2.3665172 | 2.82E-21 | -1.7903636 | 2.12E-12 |
| E2f4 | orange | 0.72337979 | 2.01E-09 | 0.55388898 | 8.41E-06 |
| Surf4 | orange | 0.67488739 | 3.18E-05 | 0.5810194 | 0.00052044 |

| | | | | | |
|---|---|---|---|---|---|
| Yes1 | red | -0.8736163 | 3.32E-07 | -0.7165477 | 5.08E-05 |
| Galns | orange | 1.31623846 | 1.43E-07 | 1.23505714 | 1.38E-06 |
| Entpd2 | red | -1.0364407 | 5.76E-07 | -0.8389177 | 9.51E-05 |
| Aldh1a3 | yellow | 1.98817791 | 4.19E-11 | 2.2259749 | 1.00E-13 |
| Actn1 | orange | 0.73192261 | 0.00034571 | 0.82366724 | 7.54E-05 |
| Nolc1 | orange | 0.69384526 | 1.97E-05 | 0.53294573 | 0.00156425 |
| Casd1 | red | -0.7656731 | 6.05E-06 | -0.7271784 | 2.86E-05 |
| Cyp2j9 | red | -1.0030785 | 0.00178843 | -0.7680002 | 0.02299474 |
| Nipsnap3b | orange | 0.76108367 | 2.48E-05 | 0.51877025 | 0.00621304 |
| Gadd45b | orange | 0.6111845 | 4.71E-08 | 0.50460979 | 1.20E-05 |
| Slamf6 | yellow | 2.5560734 | 5.11E-19 | 3.07067519 | 9.10E-26 |
| Slamf1 | yellow | 2.24052497 | 7.12E-13 | 1.99414988 | 4.24E-10 |
| Zdhhc12 | orange | 0.62964939 | 8.13E-05 | 0.53929789 | 0.00107709 |
| Endog | red | -1.0674517 | 1.12E-07 | -0.8464583 | 4.77E-05 |
| Cybb | yellow | 2.18943997 | 9.60E-10 | 1.72777606 | 2.74E-06 |
| Pcolce2 | skyblue | -2.272204 | 8.66E-10 | -1.756073 | 4.18E-06 |
| Cd48 | yellow | 2.52281542 | 1.15E-24 | 2.60540486 | 1.73E-25 |
| Dennd6b | orange | 0.81294959 | 0.00014683 | 0.70352364 | 0.00142347 |
| Ace2 | red | -1.4183695 | 0.00022229 | -1.0017102 | 0.0126004 |
| Gzmb | black | 8.11729481 | 2.29E-174 | 7.34978131 | 1.38E-145 |
| C4a | orange | 1.32007407 | 2.77E-06 | 1.01043117 | 0.00052583 |
| Notch4 | red | -0.6228404 | 0.00116589 | -0.6095978 | 0.00194153 |
| Lpl | red | -0.394915 | 0.09548551 | -0.7420938 | 0.00140272 |
| Atp6v0e | orange | 0.5949828 | 8.30E-06 | 0.35995666 | 0.01059648 |
| Gata5 | red | -0.7852369 | 0.0014092 | -0.5123834 | 0.05135471 |
| Lama5 | orange | 0.47036508 | 0.01407994 | 0.61015625 | 0.00157427 |
| Steap1 | orange | 1.63454995 | 0.00056993 | 1.37105311 | 0.00509373 |
| Steap2 | orange | 1.08977561 | 1.22E-09 | 0.76022861 | 4.28E-05 |
| Psma2 | orange | 1.03241341 | 8.55E-12 | 0.86672441 | 2.17E-08 |
| Capza2 | orange | 0.9107194 | 2.36E-06 | 0.75770691 | 0.00014253 |
| Plekho1 | orange | 0.80375069 | 3.46E-05 | 0.45267179 | 0.02804612 |
| Rxrg | skyblue | -1.7039428 | 0.00033694 | -0.9464511 | 0.06418404 |
| Rxra | red | -0.7002944 | 3.37E-07 | -0.5947996 | 2.49E-05 |
| Fcrls | skyblue | -3.5507154 | 1.09E-46 | -2.3969244 | 1.98E-22 |
| Ncapg | yellow | 3.84103152 | 3.10E-33 | 4.13989861 | 3.12E-32 |
| H2afy | orange | 0.94864351 | 2.35E-09 | 0.87339114 | 7.28E-08 |
| Gatsl2 | red | -0.6135401 | 2.96E-07 | -0.3162066 | 0.01293568 |
| Fcgr1 | black | 5.99585467 | 9.66E-27 | 5.76012717 | 1.38E-24 |
| Ncf1 | orange | 2.00683103 | 1.49E-11 | 1.63090328 | 9.06E-08 |
| Wnt11 | red | -0.9935327 | 2.71E-12 | -0.6010059 | 4.07E-05 |
| Chdh | red | -1.6615729 | 0.00044376 | -0.2949519 | 0.59493584 |
| Stk32c | orange | 1.74405811 | 4.08E-08 | 1.43955363 | 1.25E-05 |
| Fli1 | orange | 0.58220772 | 0.00054146 | 0.6189358 | 0.00033184 |

| | | | | | |
|---|---|---|---|---|---|
| Stard13 | red | -0.7599813 | 3.21E-06 | -0.5001705 | 0.00335334 |
| H2-M3 | orange | 1.88747941 | 8.66E-30 | 1.69865888 | 4.74E-24 |
| Ctsz | orange | 1.79505659 | 1.46E-10 | 1.50398396 | 1.57E-07 |
| Slmo2 | orange | 0.92532686 | 6.26E-09 | 0.79955711 | 1.05E-06 |
| Sertad4 | red | -1.3089826 | 1.65E-06 | -1.1392521 | 5.92E-05 |
| Slc25a5 | orange | 0.92815871 | 1.55E-11 | 0.76219622 | 6.41E-08 |
| Eef1a2 | red | -1.2095276 | 4.81E-06 | -1.1317175 | 3.07E-05 |
| Col20a1 | red | -1.0371445 | 0.0002145 | -0.5798842 | 0.05086006 |
| Mpped2 | red | -1.3011258 | 0.00023068 | -1.0539618 | 0.00421747 |
| E2f3 | orange | 0.75872246 | 2.28E-06 | 0.74121752 | 8.62E-06 |
| Cd274 | black | 4.5924278 | 5.06E-210 | 4.60492984 | 7.21E-214 |
| Pdcd1lg2 | black | 4.37182205 | 5.30E-24 | 5.3584594 | 7.95E-21 |
| Il10 | black | 6.19473926 | 6.66E-22 | 5.74410153 | 1.23E-22 |
| Foxred2 | orange | 1.63429004 | 3.59E-10 | 0.79581767 | 0.0027981 |
| Eif3d | orange | 0.78491334 | 6.98E-11 | 0.6519232 | 1.36E-07 |
| H3f3b | orange | 0.76095866 | 6.39E-09 | 0.69285622 | 2.23E-07 |
| Nup50 | orange | 0.72028173 | 2.92E-06 | 0.46052328 | 0.00431352 |
| Cmah | orange | 0.29564328 | 0.06533692 | 0.76129971 | 1.36E-06 |
| Scube1 | red | -0.6436864 | 0.00025862 | -1.0014463 | 1.24E-08 |
| Sulf1 | red | -1.2031751 | 7.53E-12 | -0.8414156 | 3.53E-06 |
| Matn4 | skyblue | -1.7845442 | 4.28E-08 | -1.1493315 | 0.0007174 |
| Rbpjl | skyblue | -1.682425 | 8.35E-08 | -1.0152345 | 0.00198396 |
| Sdc4 | orange | 0.9363847 | 0.00012456 | 0.93841403 | 0.00017376 |
| Il13ra1 | orange | 0.90710589 | 0.0006858 | 0.64296591 | 0.02172892 |
| Nbr1 | red | -0.6329527 | 8.60E-05 | -0.50471 | 0.00251416 |
| Cyth1 | orange | 0.73065663 | 1.65E-07 | 0.48863011 | 0.00078501 |
| Rnd3 | orange | 0.96476764 | 0.0004105 | 0.95474303 | 0.00065789 |
| Brca1 | yellow | 3.1407759 | 1.22E-24 | 3.00243148 | 7.30E-22 |
| Cntnap1 | skyblue | -1.7213185 | 2.57E-08 | -2.0579009 | 5.29E-11 |
| Psmd3 | orange | 0.87281782 | 2.50E-10 | 0.75104061 | 1.02E-07 |
| Cd300lg | skyblue | -2.0675294 | 0.00082409 | -1.9105273 | 0.00267105 |
| Mpp2 | red | -0.5691309 | 0.00627263 | -0.7260569 | 0.00056954 |
| Vtn | skyblue | -2.2350371 | 3.41E-09 | -1.5370072 | 8.61E-05 |
| Rpl19 | orange | 0.60573746 | 1.10E-06 | 0.46661771 | 0.00028922 |
| Timp2 | red | -0.9083791 | 1.77E-14 | -0.6789521 | 2.46E-08 |
| Rarb | skyblue | -1.6286961 | 5.09E-09 | -1.0371174 | 0.00022212 |
| Cdc6 | yellow | 2.3717837 | 6.67E-21 | 1.95526486 | 2.03E-14 |
| Atad5 | orange | 1.70391527 | 1.88E-14 | 1.40554764 | 6.08E-10 |
| Abr | orange | 0.66684284 | 8.03E-06 | 0.55025494 | 0.00035979 |
| Rab11fip4 | orange | 1.20496166 | 8.50E-15 | 1.10731524 | 1.94E-12 |
| Cd40 | orange | 1.5454023 | 2.65E-19 | 1.48387885 | 1.11E-17 |
| Slc35c2 | orange | 0.65446023 | 0.00212733 | 0.58204061 | 0.00835184 |
| Zfp334 | red | -0.4424973 | 0.01435791 | -0.6038454 | 0.00086404 |

| | | | | | |
|---|---|---|---|---|---|
| Wsb1 | orange | 0.90145649 | 2.39E-06 | 1.00670806 | 2.04E-07 |
| Ttpal | orange | 0.70886676 | 4.88E-15 | 0.23604657 | 0.01527019 |
| Serinc3 | orange | 0.80908392 | 4.32E-13 | 0.42183415 | 0.00033098 |
| Pgs1 | orange | 1.02416659 | 2.27E-08 | 0.9920343 | 1.05E-07 |
| Birc5 | yellow | 3.1203825 | 3.58E-15 | 3.3580936 | 9.33E-17 |
| Afmid | orange | 1.14387655 | 0.00142236 | 0.53324341 | 0.17098543 |
| Wfdc2 | red | -0.936165 | 2.63E-08 | -0.5288274 | 0.00277669 |
| Etv4 | orange | 1.48081524 | 0.00070093 | 1.8980032 | 2.58E-05 |
| Mmp9 | orange | 1.02502219 | 0.0178753 | 1.23692519 | 0.00451485 |
| Slc12a5 | skyblue | -1.4009239 | 3.28E-07 | -1.4050658 | 5.23E-07 |
| Pltp | skyblue | -1.7036657 | 1.04E-15 | -1.3379377 | 7.18E-10 |
| Ctsa | orange | 0.65590005 | 1.05E-05 | 0.4252861 | 0.00639483 |
| Pitpna | orange | 0.58598573 | 0.00071498 | 0.54658181 | 0.00216128 |
| Jph2 | red | -1.4012783 | 7.18E-12 | -0.9483671 | 7.80E-06 |
| Dhx58 | yellow | 3.80595916 | 8.23E-49 | 3.75034241 | 2.41E-47 |
| Hspb9 | yellow | 2.59668487 | 1.82E-07 | 2.67763788 | 5.46E-07 |
| Mybl2 | yellow | 2.44576143 | 8.66E-17 | 2.36374708 | 1.37E-15 |
| B4galt5 | orange | 1.51061778 | 1.44E-12 | 1.28140098 | 3.89E-09 |
| Cadps2 | red | -1.3044553 | 2.25E-23 | -0.8504178 | 2.11E-10 |
| Ddx27 | orange | 0.66239318 | 9.13E-06 | 0.58736165 | 0.00013184 |
| Cyth3 | red | -0.8196607 | 9.33E-07 | -0.6658528 | 0.0001123 |
| Cyth4 | yellow | 2.63076149 | 9.80E-19 | 2.60811421 | 2.73E-18 |
| Rac3 | skyblue | -1.8568126 | 4.46E-08 | -1.2865923 | 0.00025353 |
| Hist1h2bc | red | -0.9803651 | 6.70E-08 | -0.8055581 | 1.82E-05 |
| Baiap2l2 | skyblue | -1.7014608 | 0.0019524 | -1.9745993 | 0.00071136 |
| Mafk | orange | 0.89395458 | 1.21E-07 | 0.84802635 | 8.65E-07 |
| Erbb3 | red | -0.8122366 | 0.00028872 | -0.4798368 | 0.04363821 |
| Ikzf3 | yellow | 3.05209836 | 5.19E-35 | 3.26074409 | 4.10E-39 |
| Vmp1 | orange | 1.68090537 | 2.02E-13 | 1.52107858 | 5.54E-11 |
| Uchl5 | orange | 0.90192128 | 1.51E-05 | 0.52167037 | 0.01831381 |
| Trove2 | red | -0.779132 | 1.68E-05 | -0.6422261 | 0.00060806 |
| Pmp22 | red | -0.9389685 | 5.29E-07 | -0.6155278 | 0.00162921 |
| Psmb6 | orange | 0.89322012 | 5.90E-08 | 0.6726801 | 8.09E-05 |
| Pfn1 | orange | 1.62227837 | 6.00E-18 | 1.36920418 | 7.17E-13 |
| Gpx3 | orange | 0.58791733 | 0.00106595 | 0.36356565 | 0.05683334 |
| Il12rb2 | yellow | 2.45545644 | 6.77E-08 | 3.69875789 | 5.84E-13 |
| Kpna2 | orange | 1.25248622 | 5.02E-18 | 1.03584937 | 2.12E-12 |
| Smurf2 | red | -0.944056 | 5.15E-16 | -0.6543856 | 4.90E-08 |
| Cuedc1 | red | -0.6745216 | 4.33E-07 | -0.4231659 | 0.00247729 |
| Abi3 | orange | 0.63341695 | 1.57E-05 | 0.4956419 | 0.00114952 |
| Shroom1 | red | -1.4204712 | 1.16E-05 | -0.801272 | 0.01641569 |
| Kif3a | red | -0.6807974 | 5.86E-05 | -0.7315366 | 2.53E-05 |
| Sept8 | red | -0.5895426 | 1.41E-07 | -0.4789838 | 3.29E-05 |

| | | | | | |
|---|---|---|---|---|---|
| Mapt | skyblue | -1.3939292 | 1.32E-11 | -1.2219331 | 5.89E-09 |
| Gid4 | red | -1.3710655 | 7.84E-13 | -1.1049884 | 1.70E-08 |
| Myo1b | red | -1.3150891 | 4.29E-13 | -1.0760647 | 6.64E-09 |
| Ypel2 | red | -1.0381106 | 3.84E-06 | -0.6522758 | 0.00564123 |
| Akap1 | red | -0.810944 | 1.35E-14 | -0.4394416 | 6.63E-05 |
| C1qbp | orange | 0.64231807 | 0.00067901 | 0.65054174 | 0.00081826 |
| 6330403K07I | skyblue | -1.3775818 | 2.14E-06 | -1.1511719 | 0.00019313 |
| Slc13a3 | orange | 1.05584523 | 0.00918925 | 0.56241021 | 0.2038109 |
| Chd3 | red | -0.7175084 | 3.86E-12 | -0.4394002 | 4.55E-05 |
| Wnt9b | orange | 1.76415764 | 0.00533573 | 1.09194777 | 0.10377541 |
| Adora2b | orange | 0.52046402 | 0.02079103 | 0.65456916 | 0.00437556 |
| Trpv2 | orange | 1.87614533 | 1.29E-19 | 1.64987643 | 3.67E-15 |
| Cenpv | red | -0.5060084 | 0.0235158 | -0.6171591 | 0.00549536 |
| Trim37 | red | -0.5943062 | 5.43E-09 | -0.6094679 | 2.97E-09 |
| Ybx2 | skyblue | -1.9423955 | 5.21E-08 | -1.3025616 | 0.0003713 |
| Slc2a4 | skyblue | -1.74977 | 2.53E-07 | -1.4744361 | 2.45E-05 |
| Dnah11 | skyblue | -2.425973 | 2.49E-16 | -1.2881938 | 2.00E-05 |
| Atox1 | orange | 0.63692511 | 1.98E-06 | 0.5793465 | 2.61E-05 |
| Tbx3 | red | -1.1070259 | 3.57E-07 | -1.0278853 | 3.82E-06 |
| Dusp14 | red | -1.1054092 | 8.26E-08 | -0.8185198 | 0.0001279 |
| Ikzf1 | yellow | 2.3551074 | 2.49E-33 | 2.54435674 | 1.52E-38 |
| Copz2 | red | -0.685888 | 4.65E-05 | -0.4037183 | 0.02387607 |
| Aatf | orange | 0.61064236 | 0.00161326 | 0.78849492 | 6.09E-05 |
| Slc25a35 | red | -1.125318 | 3.43E-07 | -0.8209096 | 0.00030377 |
| Zbtb4 | red | -1.0932783 | 3.92E-12 | -0.8220419 | 3.96E-07 |
| Mpdu1 | orange | 0.63547025 | 0.00029971 | 0.43664392 | 0.01819879 |
| Cd68 | yellow | 2.72362212 | 4.17E-12 | 2.4035643 | 1.87E-09 |
| Slc35g3 | skyblue | -2.7462638 | 1.52E-06 | -2.5393026 | 6.99E-06 |
| Acsl1 | skyblue | -1.5718329 | 1.88E-09 | -1.3543398 | 4.18E-07 |
| Abca5 | red | -1.1418715 | 0.00065823 | -0.7309645 | 0.03905287 |
| Lsp1 | orange | 1.48626366 | 1.44E-15 | 1.28762458 | 1.03E-11 |
| Sfrp5 | skyblue | -1.1852715 | 0.02597384 | -1.4548168 | 0.00688925 |
| Myh11 | red | -1.4513404 | 7.01E-16 | -0.6395293 | 0.00078964 |
| Unc45b | red | -0.8571411 | 1.47E-05 | -0.7525467 | 0.00022863 |
| Rars | orange | 0.947443 | 1.03E-09 | 0.85544 | 6.90E-08 |
| Wwc1 | red | -1.1632059 | 3.72E-12 | -0.6699094 | 0.00013122 |
| Pnpla5 | skyblue | -4.0106798 | 0.00277516 | -0.5502616 | 0.67847563 |
| Mrpl45 | orange | 0.82104864 | 1.20E-07 | 0.42236695 | 0.01086707 |
| Mb | skyblue | -1.1127183 | 7.33E-06 | -1.457134 | 5.07E-09 |
| Irf1 | yellow | 2.2716982 | 3.68E-102 | 2.25454949 | 1.23E-100 |
| Slc22a5 | red | -0.5489639 | 0.00946126 | -0.591044 | 0.00611305 |
| Alox12e | yellow | 4.11110519 | 2.16E-06 | 4.19568534 | 1.39E-06 |
| Cxcl16 | yellow | 2.15630656 | 4.35E-15 | 1.83679573 | 5.17E-11 |

| | | | | | |
|---|---|---|---|---|---|
| Med11 | orange | 0.71543303 | 1.75E-05 | 0.52903753 | 0.00244874 |
| Alox15 | orange | 0.61299939 | 0.19409486 | 1.73432596 | 0.00012221 |
| Heatr9 | yellow | 2.26529708 | 1.06E-05 | 2.40057662 | 6.06E-06 |
| Ccl4 | black | 7.72705814 | 1.09E-57 | 6.48000458 | 3.67E-68 |
| Natd1 | red | -0.6346772 | 0.0012432 | -0.4637645 | 0.02403674 |
| E2f2 | orange | 0.53382778 | 4.63E-05 | 0.58035511 | 1.47E-05 |
| Slfn3 | yellow | 2.76005863 | 2.56E-12 | 2.92371158 | 1.76E-13 |
| Dnah1 | skyblue | -2.133496 | 8.65E-09 | -1.2112408 | 0.00148407 |
| Dalrd3 | red | -0.7818852 | 3.44E-06 | -0.5278889 | 0.00262399 |
| Rab3d | red | -0.590746 | 0.00016365 | -0.4524414 | 0.00550002 |
| Slc25a22 | orange | 1.31804984 | 4.79E-19 | 1.21346897 | 3.60E-16 |
| Ccl9 | orange | 1.68963306 | 7.02E-06 | 1.61135356 | 2.83E-05 |
| Scrn1 | red | -1.3328928 | 2.89E-07 | -0.9597912 | 0.00035935 |
| Tmem160 | orange | 0.87434566 | 0.00024958 | 0.74921613 | 0.00258872 |
| Rab5c | orange | 1.07618241 | 2.53E-09 | 0.99261018 | 6.90E-08 |
| Styxl1 | skyblue | -1.6242927 | 0.00098498 | -0.9727149 | 0.05348257 |
| H13 | orange | 0.74748618 | 1.79E-08 | 0.80137006 | 2.41E-09 |
| Scn1b | red | -1.3808935 | 7.83E-10 | -0.7976477 | 0.00072088 |
| Chtf18 | orange | 1.73875049 | 3.26E-10 | 1.88055674 | 2.55E-11 |
| Ahr | red | -1.0049731 | 1.94E-07 | -0.7185292 | 0.00033229 |
| Nop9 | orange | 0.90180386 | 4.87E-16 | 0.73192165 | 1.21E-10 |
| Grb7 | red | -1.4282341 | 1.30E-08 | -1.0644299 | 4.17E-05 |
| Aoc3 | skyblue | -2.7348818 | 8.02E-17 | -1.9759014 | 2.99E-09 |
| Gdpd2 | orange | 0.4257243 | 0.24637488 | 0.94520076 | 0.00653342 |
| Tlcd1 | red | -0.994991 | 6.55E-06 | -0.7330658 | 0.00132991 |
| Arhgef25 | red | -0.9107072 | 1.73E-08 | -0.7978057 | 1.39E-06 |
| Arid3a | orange | 1.09834781 | 4.04E-06 | 0.89560866 | 0.00027277 |
| Mydgf | orange | 0.62683213 | 0.0018676 | 0.52535422 | 0.01210618 |
| Sema6a | skyblue | -1.4543342 | 2.00E-06 | -1.1579286 | 0.00025151 |
| Ccdc12 | orange | 0.64822648 | 7.39E-07 | 0.50045395 | 0.00024087 |
| L3hypdh | red | -0.8887136 | 7.29E-05 | -0.8504343 | 0.00026982 |
| Lyst | orange | 0.42386716 | 0.0010497 | 0.58048278 | 8.37E-06 |
| Calr3 | red | -1.2232886 | 0.00099279 | -0.9112033 | 0.01985878 |
| Iyd | red | -1.7247007 | 3.56E-05 | -0.4972598 | 0.2603872 |
| Ccdc170 | skyblue | -2.3323032 | 1.82E-19 | -1.008549 | 4.16E-05 |
| Fbxo5 | orange | 1.45792261 | 1.85E-24 | 1.23909555 | 4.23E-17 |
| Trdn | red | -0.9487697 | 0.00558777 | -1.2441814 | 0.00036691 |
| Stxbp5 | orange | 0.97091766 | 3.49E-11 | 0.89687573 | 1.56E-09 |
| Katna1 | orange | 0.99033226 | 5.98E-12 | 0.84611932 | 1.00E-08 |
| Adat2 | orange | 0.90088172 | 7.28E-05 | 0.91429052 | 8.42E-05 |
| Ltv1 | orange | 0.83976758 | 2.49E-05 | 0.70622529 | 0.00061173 |
| Plagl1 | red | -1.2024027 | 3.20E-09 | -0.9293657 | 9.04E-06 |
| Utrn | red | -0.8097419 | 3.10E-09 | -0.6475591 | 4.07E-06 |

| | | | | | |
|---|---|---|---|---|---|
| Mical1 | orange | 1.15614838 | 3.26E-10 | 1.0442666 | 2.51E-08 |
| Wasf1 | skyblue | -1.1070773 | 0.00376979 | -1.5234239 | 7.68E-05 |
| Rab32 | orange | 1.32761266 | 6.07E-06 | 1.40266999 | 2.84E-06 |
| Fyn | orange | 0.69680544 | 4.14E-07 | 0.69660938 | 6.96E-07 |
| Tube1 | orange | 1.05764778 | 0.05616682 | 2.09378047 | 0.00013593 |
| Popdc3 | red | -0.5995476 | 1.13E-05 | -0.4026661 | 0.00549357 |
| Tnfaip3 | yellow | 2.31649424 | 7.40E-30 | 2.085308 | 2.54E-24 |
| Perp | red | -0.5858062 | 0.05067841 | -0.802255 | 0.0076288 |
| Hebp2 | red | -0.4285114 | 0.15565868 | -0.8136998 | 0.0064885 |
| Fam184a | red | -1.1501873 | 1.06E-05 | -1.028611 | 0.00011759 |
| Qrsl1 | orange | 0.52658964 | 0.00489547 | 0.63080193 | 0.00109536 |
| Aim1 | orange | 1.46894173 | 2.39E-14 | 1.26756707 | 9.33E-11 |
| Vta1 | orange | 0.75929297 | 0.00068168 | 0.5048332 | 0.03246 |
| Smpdl3a | red | -0.5971164 | 2.47E-05 | -0.5895691 | 4.68E-05 |
| Pkib | yellow | 2.52770596 | 7.28E-11 | 1.90157997 | 1.58E-06 |
| Rspo3 | orange | 1.57311856 | 0.00447976 | 1.81049445 | 0.00182277 |
| Echdc1 | red | -0.882677 | 0.00266565 | -0.7896706 | 0.00902243 |
| Slc6a15 | skyblue | -2.6813458 | 8.97E-09 | -2.2808888 | 1.26E-06 |
| Lama2 | red | -0.9254909 | 0.00016111 | -0.5222205 | 0.04559019 |
| Gprc6a | skyblue | -1.6141348 | 0.00073607 | -0.9648976 | 0.05562189 |
| Fam162b | skyblue | -2.1464051 | 1.01E-09 | -1.618549 | 7.76E-06 |
| P4ha1 | red | -0.5930819 | 5.11E-11 | -0.6180106 | 1.54E-11 |
| Zwint | orange | 0.69470072 | 0.00014368 | 0.53515463 | 0.00486039 |
| Dcn | red | -1.1279534 | 3.12E-07 | -0.7932571 | 0.00054434 |
| Mrln | red | -1.4217091 | 0.00317859 | -0.0814267 | 0.89505588 |
| Cdk1 | yellow | 2.80820711 | 2.67E-24 | 3.75828674 | 4.13E-38 |
| Rhobtb1 | red | -0.9161282 | 1.04E-05 | -1.1275857 | 7.27E-08 |
| 1700040L02F | skyblue | -2.0215174 | 1.01E-07 | -1.4217923 | 0.00024387 |
| Dusp6 | orange | 0.8438277 | 0.00017 | 0.76839775 | 0.00088164 |
| Kitl | red | -0.9814712 | 8.51E-06 | -0.7607119 | 0.00086573 |
| Sgk1 | orange | 0.64238259 | 2.22E-07 | 0.2749987 | 0.03897522 |
| Apaf1 | orange | 1.06602878 | 3.34E-15 | 0.91472063 | 2.89E-11 |
| Arg1 | black | 5.02439117 | 6.73E-17 | 4.54693112 | 7.13E-14 |
| Enpp3 | skyblue | -1.6266545 | 9.46E-07 | -1.2045981 | 0.00034298 |
| Pde7b | red | -1.0963651 | 2.00E-10 | -0.5239602 | 0.00432443 |
| Mtfr2 | yellow | 3.14657234 | 2.37E-20 | 3.23486753 | 3.43E-19 |
| Map7 | red | -0.7500581 | 0.00014944 | -0.4695022 | 0.02407771 |
| Ctgf | orange | 0.64591615 | 0.00355766 | 0.6838916 | 0.00250534 |
| Vnn3 | red | -1.4879395 | 3.20E-05 | -0.4266107 | 0.27923497 |
| Cfap54 | skyblue | -2.243425 | 6.43E-15 | -1.2951595 | 6.53E-06 |
| Ntn4 | skyblue | -1.6136872 | 2.66E-11 | -1.1605349 | 3.48E-06 |
| Usp44 | skyblue | -1.6268028 | 0.00500155 | -0.7905836 | 0.23211058 |
| Nudt4 | red | -0.6691216 | 1.73E-05 | -0.8972257 | 9.01E-09 |

| | | | | | |
|---|---|---|---|---|---|
| Nuak1 | skyblue | -1.7283947 | 2.71E-10 | -1.4702389 | 1.50E-07 |
| Tcp11l2 | red | -1.1778135 | 1.34E-05 | -0.9847819 | 0.00042261 |
| Timp3 | orange | 0.77105093 | 1.11E-08 | 0.54502308 | 9.87E-05 |
| Ascl1 | skyblue | -3.7491986 | 2.59E-05 | -1.1719409 | 0.11043007 |
| Igf1 | orange | 1.11532703 | 0.00044539 | 1.18271041 | 0.00028216 |
| Slc5a8 | skyblue | -6.0745439 | 9.82E-07 | -3.2746029 | 1.15E-06 |
| Mypn | red | -1.0929812 | 2.28E-06 | -0.8424961 | 0.00047762 |
| Hnrnph3 | red | -0.7151779 | 6.14E-07 | -0.4965978 | 0.00091978 |
| Ddx21 | orange | 1.03717246 | 7.56E-08 | 0.96628425 | 9.47E-07 |
| Srgn | orange | 1.90117932 | 5.91E-16 | 1.62713597 | 9.44E-12 |
| Ppa1 | orange | 1.91666722 | 4.52E-23 | 1.8636385 | 1.03E-21 |
| Eif4ebp2 | red | -0.7506011 | 1.53E-07 | -0.6867679 | 2.69E-06 |
| Sgpl1 | orange | 0.64823377 | 0.0003218 | 0.65846332 | 0.00036073 |
| Pcbd1 | red | -1.0479388 | 4.72E-05 | -0.6844979 | 0.01268095 |
| Slc29a3 | orange | 1.53055348 | 1.15E-06 | 1.31061392 | 5.13E-05 |
| Slc16a7 | skyblue | -2.2369173 | 3.10E-11 | -1.4079386 | 4.97E-05 |
| Lrig3 | red | -0.8753102 | 5.52E-06 | -0.4195036 | 0.04051185 |
| Ddit4 | orange | 1.69907365 | 3.39E-07 | 0.72845599 | 0.04230973 |
| Tbk1 | orange | 0.97128039 | 2.02E-14 | 0.80060604 | 6.49E-10 |
| Pno1 | orange | 0.89015407 | 0.00018749 | 0.84191697 | 0.00061088 |
| Plek | yellow | 2.41260975 | 5.95E-16 | 2.01823581 | 2.93E-11 |
| Avpr1a | skyblue | -1.6327662 | 0.00397082 | -1.1534875 | 0.05231524 |
| Vps54 | orange | 0.76481719 | 1.41E-09 | 0.5998642 | 4.00E-06 |
| Pcsk4 | red | -0.9348702 | 0.0004496 | -0.4879623 | 0.08245111 |
| Peli1 | orange | 0.65686028 | 2.57E-07 | 0.52797167 | 5.98E-05 |
| Apc2 | skyblue | -1.3556684 | 2.14E-07 | -1.2882669 | 1.29E-06 |
| Thap2 | red | -0.8915996 | 1.95E-07 | -0.8704981 | 5.85E-07 |
| Slc1a4 | red | -0.9335371 | 2.66E-09 | -0.534829 | 0.00105441 |
| Dock2 | yellow | 2.47238226 | 1.28E-22 | 2.37517596 | 8.41E-21 |
| Gamt | skyblue | -1.3627903 | 6.07E-07 | -1.5234026 | 7.70E-08 |
| Actr2 | orange | 0.87426759 | 5.69E-07 | 0.70429035 | 9.58E-05 |
| Ptprb | red | -1.158779 | 5.88E-07 | -0.9515309 | 6.97E-05 |
| Kcnmb1 | red | -1.5081411 | 1.69E-10 | -0.9539896 | 8.71E-05 |
| Gabrp | skyblue | -1.9070565 | 5.80E-15 | -0.9368197 | 0.00025379 |
| Meis1 | red | -1.2033903 | 7.79E-09 | -0.8249469 | 0.00013786 |
| Best3 | skyblue | -2.0137226 | 1.24E-05 | -0.4229407 | 0.41256379 |
| Cobl | red | -0.8264181 | 2.92E-05 | -0.6592907 | 0.00126808 |
| Rab36 | red | -0.7807876 | 1.83E-06 | -0.481748 | 0.00454784 |
| Adora2a | orange | 1.7916193 | 6.33E-26 | 1.7018122 | 4.14E-23 |
| Snrpd3 | orange | 0.63023363 | 0.00260372 | 0.49959846 | 0.02241468 |
| Ddc | skyblue | -1.5757035 | 0.00018185 | -1.7188603 | 6.88E-05 |
| Cpm | skyblue | -2.1195646 | 9.62E-08 | -1.4344354 | 0.00052581 |
| E2f7 | yellow | 1.80654173 | 1.30E-10 | 2.27278222 | 4.90E-15 |

| | | | | | |
|---|---|---|---|---|---|
| Phlda1 | orange | 1.02189254 | 5.96E-05 | 1.21459389 | 2.52E-06 |
| Jsrp1 | red | -1.187658 | 0.00016517 | -0.7141545 | 0.03479706 |
| Wif1 | skyblue | -4.0522417 | 8.80E-14 | -2.7457035 | 1.24E-07 |
| Tmbim4 | orange | 0.69911921 | 7.66E-07 | 0.59725447 | 4.16E-05 |
| Helb | orange | 0.93215301 | 6.46E-12 | 0.81991968 | 3.71E-09 |
| Prmt2 | red | -0.7889717 | 3.64E-11 | -0.4959403 | 6.85E-05 |
| Fzr1 | orange | 0.63035801 | 3.22E-06 | 0.66750181 | 1.31E-06 |
| Txnrd1 | orange | 0.74252923 | 2.26E-05 | 0.7830257 | 1.16E-05 |
| Glt8d2 | skyblue | -3.2657469 | 3.38E-16 | -2.2017098 | 1.08E-08 |
| Slc36a1 | orange | 0.70380742 | 0.00021446 | 0.54051622 | 0.00626408 |
| Adarb1 | red | -0.9401372 | 5.42E-06 | -0.9414296 | 8.09E-06 |
| Appl2 | red | -1.0348894 | 9.36E-07 | -0.7321418 | 0.00083339 |
| Slc36a2 | red | -0.9324381 | 0.00022996 | -0.9229062 | 0.00037703 |
| Hint1 | orange | 0.71877185 | 8.99E-07 | 0.46528889 | 0.00238699 |
| Lyrm7 | red | -0.3414895 | 0.30490242 | -0.8448362 | 0.0084991 |
| Stk10 | orange | 1.18127617 | 1.47E-07 | 1.1668721 | 3.51E-07 |
| Rel | orange | 1.281837 | 6.99E-10 | 1.16528266 | 4.20E-08 |
| 1700093K21I | yellow | 0.31876556 | 0.73581625 | 2.84077542 | 0.00429329 |
| Mpg | orange | 0.66565598 | 2.82E-05 | 0.45549617 | 0.0059974 |
| Nsg2 | orange | 0.43203658 | 0.47026587 | 1.8196399 | 0.00096939 |
| Cdc34 | orange | 0.70740765 | 6.08E-05 | 0.56927478 | 0.00183915 |
| Shc2 | skyblue | -2.2226064 | 2.25E-17 | -1.1692133 | 1.04E-05 |
| Sptbn1 | red | -0.8711198 | 1.04E-05 | -0.6574266 | 0.00134273 |
| Wdpcp | red | -0.9736794 | 1.37E-10 | -0.741988 | 1.82E-06 |
| Fstl3 | orange | 1.0817481 | 4.49E-06 | 1.25132073 | 1.43E-07 |
| Nudcd2 | orange | 0.80843273 | 4.39E-05 | 0.51536718 | 0.01344426 |
| Hmmr | yellow | 3.31678963 | 6.33E-28 | 3.51684776 | 1.85E-27 |
| Hcn2 | red | -0.906826 | 0.00807747 | -0.7546165 | 0.03945055 |
| Zfp354b | skyblue | -1.3429729 | 1.63E-05 | -1.3452072 | 2.62E-05 |
| Cyfip2 | orange | 1.82842261 | 6.40E-34 | 1.6071308 | 2.98E-26 |
| Hnrnpab | orange | 0.7522464 | 0.00126373 | 0.97687712 | 3.42E-05 |
| Gfpt2 | orange | 0.87874866 | 3.46E-05 | 0.97825895 | 5.75E-06 |
| Zfp354a | red | -0.9347752 | 0.00114708 | -0.8678365 | 0.00295649 |
| Gnb2l1 | orange | 0.58819568 | 1.29E-07 | 0.50019303 | 1.25E-05 |
| Il13 | black | 5.74821873 | 9.56E-05 | 2.3191206 | 0.0084676 |
| Jade2 | orange | 0.62167976 | 1.99E-08 | 0.31855748 | 0.00642155 |
| Cdkl3 | orange | 0.86246477 | 0.00021797 | 0.89389151 | 0.00016942 |
| Kremen1 | red | -0.7284294 | 3.37E-06 | -0.2425757 | 0.16138401 |
| Itk | yellow | 2.55967704 | 7.89E-19 | 2.405104 | 1.34E-16 |
| Nefh | yellow | 3.24710416 | 2.44E-13 | 3.04574131 | 1.80E-13 |
| Med7 | orange | 0.67169332 | 0.00071512 | 0.79998179 | 8.82E-05 |
| Havcr2 | yellow | 3.74926452 | 1.28E-22 | 3.16553186 | 2.77E-16 |
| Tnip1 | orange | 1.3756104 | 6.43E-13 | 1.22502106 | 2.96E-10 |

| | | | | | |
|---|---|---|---|---|---|
| Fam71b | yellow | 3.45491604 | 6.09E-07 | 3.92182063 | 1.42E-06 |
| Upp1 | yellow | 2.53604164 | 4.43E-21 | 2.59051298 | 8.72E-22 |
| Tns3 | red | -0.7459515 | 6.42E-11 | -0.4933731 | 3.17E-05 |
| Igfbp3 | skyblue | -1.3649469 | 2.32E-25 | -1.2187163 | 1.91E-20 |
| Pes1 | orange | 0.60381702 | 0.00058943 | 0.41755166 | 0.02385107 |
| Adcy1 | skyblue | -1.5191279 | 0.02351021 | -1.7655027 | 0.00957316 |
| Osbp2 | skyblue | -1.6963775 | 0.03002868 | -2.1416574 | 0.0066853 |
| Myo1g | yellow | 2.27600692 | 8.40E-32 | 2.15059232 | 2.43E-28 |
| Smtn | red | -0.653385 | 0.00012709 | -0.5284007 | 0.00277941 |
| Arf5 | orange | 0.74381217 | 5.29E-06 | 0.5726686 | 0.00074247 |
| Patz1 | red | -0.6707452 | 3.03E-05 | -0.3267495 | 0.05810767 |
| Ogdh | red | -0.5962631 | 2.88E-06 | -0.5198159 | 7.37E-05 |
| Rps27a | orange | 0.65668463 | 4.94E-07 | 0.50440445 | 0.00019338 |
| Pnpt1 | orange | 1.03226683 | 8.47E-14 | 1.01748916 | 2.98E-13 |
| Efemp1 | skyblue | -1.5441403 | 4.86E-06 | -1.0550518 | 0.00275197 |
| Myl7 | skyblue | -1.5803056 | 0.07308532 | -2.3945308 | 0.00635518 |
| Pold2 | orange | 0.68417799 | 6.11E-05 | 0.45472863 | 0.01112121 |
| Aebp1 | red | -0.8095185 | 5.27E-08 | -0.4232593 | 0.0069959 |
| Pgam2 | skyblue | -1.7908524 | 1.16E-08 | -1.3172919 | 5.24E-05 |
| Dbnl | orange | 0.95822863 | 6.39E-14 | 0.90843583 | 1.98E-12 |
| Dynll2 | red | -1.1025714 | 1.45E-17 | -0.7930075 | 2.42E-09 |
| Sept4 | red | -0.9370337 | 5.44E-06 | -0.7497649 | 0.00043808 |
| Prr11 | yellow | 3.00635604 | 3.50E-22 | 3.01983289 | 3.20E-21 |
| Tubd1 | red | -0.3694042 | 0.06837476 | -0.6558111 | 0.00102179 |
| Acaca | red | -0.7602519 | 0.00957567 | -1.0612126 | 0.00031792 |
| Shmt1 | orange | 0.71614439 | 0.0008586 | 0.80608617 | 0.00023935 |
| Tom1l1 | red | -0.7667675 | 0.00070582 | -0.2977125 | 0.23432108 |
| Myocd | red | -1.0578015 | 0.00451015 | -1.0136796 | 0.00699169 |
| Cox11 | orange | 0.33396594 | 0.27020529 | 0.83289076 | 0.00451562 |
| Stxbp4 | skyblue | -1.5018543 | 2.81E-09 | -1.2004136 | 2.16E-06 |
| Efcab10 | skyblue | -2.7908684 | 8.99E-06 | -1.6852148 | 0.00665942 |
| Atxn7l1 | orange | 0.74321712 | 9.64E-07 | 0.61225647 | 8.80E-05 |
| Atp6v1c2 | red | -1.5415859 | 6.36E-06 | -0.9774569 | 0.00626378 |
| Sypl | red | -0.572531 | 7.86E-07 | -0.5842273 | 7.53E-07 |
| Nampt | orange | 1.70296664 | 6.14E-35 | 1.25356609 | 3.16E-19 |
| Pik3cg | yellow | 2.75937433 | 1.76E-26 | 2.03292993 | 7.26E-15 |
| Tspan13 | skyblue | -1.3704554 | 2.88E-09 | -1.1824651 | 5.66E-07 |
| Agr2 | orange | 0.01891873 | 0.96882077 | 1.60188068 | 4.41E-05 |
| Lpin1 | red | -0.91106 | 0.00727305 | -1.0643055 | 0.00198861 |
| Rgs9 | skyblue | -1.8182835 | 1.19E-06 | -1.2322571 | 0.00145995 |
| Hs1bp3 | red | -0.7195343 | 8.83E-08 | -0.4435425 | 0.00157477 |
| Fam84a | skyblue | -1.7220744 | 2.22E-30 | -0.8839657 | 4.02E-09 |
| Gna13 | orange | 1.19482945 | 4.35E-13 | 1.13209247 | 1.16E-11 |

| | | | | | |
|---|---|---|---|---|---|
| Abca8b | red | -1.4369124 | 1.04E-09 | -0.985271 | 5.52E-05 |
| Map2k6 | red | -1.0481278 | 9.02E-05 | -1.2278701 | 8.15E-06 |
| Klhl29 | red | -0.5780553 | 0.04256182 | -0.7423708 | 0.00919383 |
| Adi1 | red | -1.0714082 | 2.68E-13 | -0.9580274 | 1.12E-10 |
| Fkbp1b | orange | 1.31429755 | 2.95E-06 | 1.04878317 | 0.00032302 |
| Cmpk2 | yellow | 2.6295269 | 1.85E-37 | 2.66780921 | 1.68E-38 |
| Rsad2 | black | 5.82808577 | 4.27E-64 | 5.45456007 | 2.44E-56 |
| Rnf144a | skyblue | -1.7611913 | 4.70E-17 | -0.9820607 | 7.47E-06 |
| Id2 | orange | 1.41984114 | 6.94E-18 | 1.03571383 | 9.84E-10 |
| Mboat2 | red | -0.6999059 | 0.00097324 | -0.7405773 | 0.0006704 |
| Rrm2 | yellow | 2.97737961 | 4.20E-27 | 3.41865805 | 3.50E-34 |
| Bcap29 | orange | 0.86442263 | 0.00039683 | 0.40673246 | 0.12679278 |
| Slc26a4 | yellow | 3.49058307 | 1.35E-27 | 3.93736538 | 4.73E-33 |
| Cenpo | orange | 0.76429922 | 5.29E-05 | 0.58781952 | 0.00303962 |
| Klf11 | red | -0.7511798 | 3.06E-06 | -0.4586702 | 0.00666795 |
| Adcy3 | red | -0.599408 | 0.00019366 | -0.5010685 | 0.00271117 |
| Efr3b | orange | 0.82611098 | 0.00426493 | 1.07278325 | 0.00024315 |
| Kif3c | red | -0.7961863 | 5.36E-12 | -0.8084308 | 5.53E-12 |
| Pxdn | red | -0.9701171 | 9.12E-14 | -0.8191947 | 6.62E-10 |
| Ccl11 | orange | 1.18838675 | 1.62E-06 | 1.58418222 | 1.59E-10 |
| Hnf1b | red | -0.8997325 | 3.27E-06 | -0.6376144 | 0.00132854 |
| Ace | red | -0.4905314 | 3.25E-05 | -0.6417503 | 6.38E-08 |
| Mmp28 | skyblue | -1.7428435 | 5.78E-09 | -1.3195289 | 1.88E-05 |
| Rasl10b | yellow | 2.48444133 | 6.25E-09 | 1.56498621 | 0.00041539 |
| Gas2l2 | skyblue | -1.6475379 | 2.50E-07 | -0.879502 | 0.0062221 |
| Itgb3 | orange | 0.93808193 | 7.98E-05 | 0.85870434 | 0.00045706 |
| Nle1 | orange | 0.85230756 | 0.00033701 | 0.90449353 | 0.00019601 |
| Mrc2 | red | -0.6692106 | 0.00820882 | -0.4931352 | 0.06409948 |
| Ccl1 | black | 8.48516845 | 3.85E-10 | 7.1445923 | 2.93E-07 |
| Asic2 | yellow | 3.31551012 | 6.66E-05 | 2.75486996 | 0.00160296 |
| Ftsj3 | orange | 0.81839458 | 2.02E-06 | 0.76015115 | 1.69E-05 |
| Psmc5 | orange | 0.61259802 | 7.42E-06 | 0.45719963 | 0.00129459 |
| Adap2 | orange | 1.18743809 | 7.93E-08 | 0.96341613 | 2.29E-05 |
| Psmd12 | orange | 0.76663652 | 6.62E-07 | 0.63216794 | 7.16E-05 |
| Cacng1 | red | -1.1056862 | 2.84E-05 | -0.6921711 | 0.01408352 |
| Cep112 | red | -0.7982737 | 6.05E-05 | -0.7646753 | 0.00018053 |
| Rab37 | yellow | 2.17630467 | 5.43E-07 | 2.01524325 | 6.04E-06 |
| Slc9a3r1 | orange | 0.95826615 | 4.73E-09 | 0.75163188 | 8.28E-06 |
| Nt5c | orange | 0.79558114 | 4.71E-08 | 0.68658235 | 4.64E-06 |
| Hn1 | orange | 1.22097622 | 6.72E-14 | 1.07427154 | 9.33E-11 |
| Mif4gd | orange | 0.88909783 | 5.57E-22 | 0.69043347 | 2.83E-13 |
| 2310067B10I | red | -0.8181011 | 4.02E-08 | -0.6270003 | 4.57E-05 |
| Itgb4 | red | -0.7711245 | 2.47E-10 | -0.3373895 | 0.00892422 |

| | | | | | |
|---|---|---|---|---|---|
| Aspa | skyblue | -1.5883007 | 7.84E-08 | -1.1799695 | 8.23E-05 |
| Fbf1 | red | -0.6295765 | 1.95E-05 | -0.5046128 | 0.00093085 |
| Acox1 | red | -0.5824988 | 0.00054858 | -0.3794163 | 0.03278858 |
| Camkk1 | red | -1.2222801 | 8.60E-26 | -0.5888821 | 1.01E-06 |
| Atp2a3 | skyblue | -1.5503895 | 5.02E-11 | -1.2926161 | 8.73E-08 |
| Ankfy1 | orange | 0.60141247 | 3.58E-07 | 0.62588999 | 1.87E-07 |
| Spns3 | orange | 1.61297738 | 0.00470516 | 1.08702839 | 0.08090044 |
| Tekt1 | red | -1.0481737 | 2.23E-09 | -0.9060601 | 3.44E-07 |
| Rhbdf2 | orange | 1.21229866 | 4.38E-07 | 1.15587082 | 2.42E-06 |
| Fam64a | yellow | 3.07961971 | 4.70E-17 | 3.29309813 | 1.12E-17 |
| Cygb | red | -0.8195906 | 1.30E-06 | -0.5669005 | 0.00131453 |
| Wscd1 | skyblue | -2.4047478 | 2.99E-17 | -1.9227514 | 3.20E-11 |
| 1810032O08 | orange | 0.95558221 | 3.48E-07 | 1.09516766 | 5.82E-09 |
| Mxra7 | red | -0.8476815 | 1.95E-06 | -0.6869721 | 0.00019256 |
| Nos2 | yellow | 2.07301706 | 3.99E-07 | 2.39469293 | 8.35E-09 |
| Slc46a1 | red | -0.6553702 | 0.00243979 | -0.9174988 | 3.42E-05 |
| Tmigd1 | skyblue | -1.7242998 | 0.00253096 | -0.7277195 | 0.25301268 |
| Rph3al | red | -1.4546338 | 1.28E-19 | -0.8723107 | 1.10E-07 |
| Doc2b | yellow | 3.45800304 | 1.60E-25 | 1.07516186 | 0.00128944 |
| Cacna1g | skyblue | -1.3749408 | 0.01026509 | -1.7531331 | 0.0013565 |
| Lrrc59 | orange | 1.12938831 | 3.15E-06 | 1.12301891 | 5.60E-06 |
| Slc35b1 | orange | 0.59135254 | 3.38E-06 | 0.54654904 | 2.82E-05 |
| Scrn2 | red | -0.6557557 | 1.61E-07 | -0.4014202 | 0.00252962 |
| Lrrc46 | red | -1.4481579 | 1.01E-08 | -0.7145894 | 0.00744123 |
| Cacnb1 | skyblue | -1.6620024 | 1.55E-14 | -1.0203041 | 5.76E-06 |
| Asgr1 | skyblue | -3.6928568 | 8.89E-14 | -3.4377982 | 9.01E-11 |
| Nr1d1 | skyblue | -2.3786861 | 1.00E-12 | -1.5671008 | 5.93E-06 |
| Tmem107 | red | -1.0098835 | 1.54E-09 | -0.8370862 | 9.55E-07 |
| Aurkb | yellow | 3.22753682 | 8.43E-24 | 3.70000994 | 1.85E-29 |
| Myh10 | red | -1.2705526 | 1.79E-09 | -0.8348718 | 0.00014478 |
| Pik3r5 | yellow | 2.42655688 | 6.79E-34 | 2.3140465 | 8.38E-31 |
| Cfap52 | skyblue | -1.5601668 | 6.29E-13 | -0.9273746 | 2.54E-05 |
| Krt19 | orange | 0.82052423 | 0.01054166 | 1.07297217 | 0.00090887 |
| Top2a | yellow | 3.15002794 | 1.11E-26 | 3.66567423 | 3.34E-35 |
| Lsm12 | orange | 0.81858898 | 5.79E-07 | 0.6582216 | 0.00010426 |
| Adam11 | orange | 0.62697642 | 0.00038431 | 0.14308555 | 0.48477605 |
| Higd1b | skyblue | -1.6186573 | 1.32E-05 | -1.5167234 | 6.77E-05 |
| Ccdc103 | skyblue | -1.5915892 | 5.66E-05 | -0.7393967 | 0.07421833 |
| Nmt1 | orange | 0.71331445 | 1.15E-09 | 0.52658247 | 1.41E-05 |
| Map3k14 | orange | 1.21085338 | 1.94E-19 | 1.18454467 | 1.48E-18 |
| Scfd1 | orange | 0.62251286 | 5.45E-07 | 0.52118978 | 4.69E-05 |
| Pole2 | orange | 1.5325934 | 2.04E-06 | 1.26573371 | 0.0001802 |
| L2hgdh | red | -0.8272035 | 4.00E-07 | -0.6069527 | 0.00031643 |

| | | | | | |
|---|---|---|---|---|---|
| Cdkl1 | skyblue | -1.9163027 | 0.00019504 | -1.5455282 | 0.00309114 |
| Ttc8 | red | -1.1473855 | 2.03E-09 | -0.8282304 | 2.02E-05 |
| Polr2h | orange | 0.63296131 | 8.88E-06 | 0.4233977 | 0.00538015 |
| Psma6 | orange | 0.82919011 | 2.22E-05 | 0.67266568 | 0.00088967 |
| Nfkbia | orange | 1.0447492 | 2.20E-10 | 0.60070062 | 0.00050097 |
| Ngb | skyblue | -1.8045055 | 1.22E-06 | -1.5904279 | 2.63E-05 |
| Gstz1 | red | -1.0832015 | 1.72E-05 | -0.8193643 | 0.00168264 |
| Nova1 | red | -1.0201232 | 4.02E-07 | -0.923945 | 1.69E-05 |
| Esr2 | skyblue | -2.0953256 | 0.00010462 | -1.9059067 | 0.000585 |
| Akap5 | red | -0.9650822 | 0.00056119 | -0.3463792 | 0.26667481 |
| Atl1 | red | -0.9928799 | 0.00043936 | -0.7485904 | 0.0106068 |
| Pygl | red | -0.7043453 | 0.00091056 | -0.6483268 | 0.00299072 |
| Timm9 | orange | 0.61833747 | 0.00314133 | 0.31155591 | 0.17402392 |
| Rtn1 | red | -1.0357044 | 0.00024458 | -0.9945858 | 0.00062201 |
| Serpina3n | yellow | 3.15244253 | 8.17E-28 | 3.04438788 | 6.15E-26 |
| Dhrs7 | red | -0.6179714 | 3.51E-08 | -0.6603458 | 6.43E-09 |
| Clmn | red | -0.9563685 | 1.54E-06 | -0.693648 | 0.00070122 |
| Hif1a | orange | 1.6498011 | 4.00E-17 | 1.46702886 | 1.44E-13 |
| Mpp5 | red | -1.0520161 | 4.40E-05 | -0.9185555 | 0.00053906 |
| Atp6v1d | orange | 0.59795465 | 6.92E-05 | 0.36905314 | 0.0201881 |
| Vrk1 | orange | 0.72838182 | 6.12E-11 | 0.62625993 | 4.75E-08 |
| Eif2s1 | orange | 1.03505803 | 1.36E-08 | 0.73043423 | 0.00011404 |
| Arg2 | yellow | 2.56482079 | 6.84E-13 | 1.50791827 | 1.74E-05 |
| Erh | orange | 0.86878037 | 4.47E-09 | 0.76746692 | 4.71E-07 |
| Susd6 | orange | 0.6050992 | 1.86E-07 | 0.4245088 | 0.00043216 |
| Smoc1 | skyblue | -1.6119313 | 6.42E-07 | -1.2202521 | 0.00022174 |
| Zmynd11 | red | -0.6456647 | 5.34E-06 | -0.5560937 | 0.00014173 |
| Cdca7l | orange | 0.88561979 | 3.83E-08 | 0.74196665 | 6.46E-06 |
| Psmc1 | orange | 0.7278851 | 5.13E-07 | 0.53643974 | 0.00036775 |
| Fbln5 | skyblue | -1.6553973 | 1.30E-08 | -1.1862616 | 8.44E-05 |
| Tc2n | orange | 0.63843586 | 0.00401362 | 0.60415322 | 0.00901599 |
| Lgmn | yellow | 2.4479671 | 3.84E-11 | 2.08937498 | 3.39E-08 |
| Pfkp | orange | 1.51234927 | 2.25E-18 | 1.31563903 | 5.97E-14 |
| Unc79 | skyblue | -3.5188505 | 7.69E-13 | -2.0428144 | 9.74E-06 |
| Asb2 | red | -0.7635579 | 0.00118116 | -0.6778195 | 0.00564527 |
| Otub2 | red | -0.7212815 | 0.0015545 | -0.6774828 | 0.00384655 |
| Ifi27l2b | yellow | 3.42580307 | 1.17E-06 | 2.76430767 | 0.00020143 |
| Ppp4r4 | red | -1.012483 | 8.40E-05 | -0.798895 | 0.00277669 |
| Akr1c13 | orange | 0.67806749 | 0.00810456 | -0.1166204 | 0.7084278 |
| Net1 | red | -0.825797 | 0.00010464 | -0.8470075 | 9.83E-05 |
| Rgs6 | skyblue | -1.4816588 | 6.45E-10 | -1.5695249 | 8.43E-11 |
| Fam161b | skyblue | -1.8938803 | 4.52E-07 | -1.1562011 | 0.00269492 |
| Entpd5 | red | -0.8418624 | 3.97E-08 | -0.7100093 | 6.35E-06 |

| | | | | | |
|---|---|---|---|---|---|
| Aldh6a1 | red | -1.447906 | 0.00015524 | -1.0634933 | 0.0076681 |
| Fcf1 | orange | 0.83383757 | 3.89E-06 | 0.82544454 | 8.75E-06 |
| Mlh3 | red | -0.836732 | 1.78E-12 | -0.8565376 | 1.84E-12 |
| Esrrb | skyblue | -0.9732317 | 0.22038364 | -2.4410904 | 0.00264499 |
| Vash1 | red | -0.7212801 | 8.12E-05 | -0.6123887 | 0.00120646 |
| Angel1 | red | -0.5814492 | 0.00018765 | -0.4062772 | 0.01351907 |
| Hhipl1 | orange | 1.08256697 | 0.00021783 | 0.90089226 | 0.00289367 |
| Evl | orange | 1.24277483 | 1.57E-11 | 1.29569237 | 3.39E-12 |
| Wars | yellow | 2.36925045 | 4.04E-66 | 2.3196848 | 2.32E-63 |
| Meg3 | skyblue | -1.7140899 | 2.91E-13 | -1.0844558 | 5.31E-06 |
| Traf3 | orange | 0.62309174 | 0.00023689 | 0.71224099 | 3.57E-05 |
| Amn | yellow | 3.40213366 | 4.22E-05 | 3.21674468 | 0.00022657 |
| Exoc3l4 | orange | 0.87899126 | 0.00014795 | 0.84514215 | 0.00036096 |
| Tnfaip2 | yellow | 2.40346466 | 4.21E-23 | 2.62582253 | 2.29E-27 |
| Zfyve21 | red | -1.0202581 | 0.00023809 | -0.7735459 | 0.00742792 |
| Kif26a | skyblue | -1.9293714 | 5.86E-14 | -1.418247 | 7.46E-08 |
| Gpr132 | yellow | 2.30569427 | 9.09E-31 | 2.26653665 | 1.43E-29 |
| Gng4 | skyblue | -2.0624442 | 7.48E-06 | -1.3653661 | 0.00421081 |
| Amph | red | -1.4634601 | 3.68E-07 | -0.9771333 | 0.0010696 |
| Sfrp4 | skyblue | -3.8248314 | 3.07E-08 | -2.0809554 | 0.00339118 |
| Aoah | yellow | 3.85330633 | 3.59E-15 | 3.83490122 | 1.54E-14 |
| Irf4 | orange | 1.27815061 | 3.86E-07 | 1.26629083 | 9.88E-07 |
| Mak | skyblue | -1.872215 | 3.69E-15 | -1.1245446 | 2.62E-06 |
| Edn1 | red | -0.2701281 | 0.08556922 | -0.7415766 | 9.03E-07 |
| Cap2 | skyblue | -1.4724747 | 1.24E-07 | -1.389119 | 1.15E-06 |
| Kif13a | red | -0.7518924 | 5.75E-05 | -0.6014828 | 0.00187458 |
| Dek | orange | 0.79488012 | 5.50E-07 | 0.62597361 | 0.00014407 |
| Id4 | red | -1.3062991 | 5.61E-07 | -1.141227 | 2.19E-05 |
| Susd3 | orange | 2.03442632 | 2.05E-10 | 1.81796865 | 3.31E-08 |
| Aspn | skyblue | -1.8170598 | 2.05E-11 | -1.7038808 | 8.27E-10 |
| Ogn | skyblue | -1.6415826 | 1.23E-06 | -1.3478397 | 0.00011254 |
| Serpinb9b | yellow | 3.36308927 | 1.86E-09 | 3.75265812 | 1.07E-10 |
| Ripk1 | orange | 1.15058305 | 6.95E-20 | 0.9506234 | 1.06E-13 |
| Ly86 | yellow | 3.04202226 | 1.21E-31 | 2.64277047 | 1.46E-23 |
| Riok1 | orange | 0.81827746 | 1.95E-09 | 0.69078405 | 8.74E-07 |
| Sema4d | yellow | 2.49827577 | 5.44E-30 | 2.39542473 | 1.83E-27 |
| Gadd45g | orange | 1.96502292 | 2.73E-24 | 1.79761918 | 2.09E-20 |
| Syk | orange | 1.12857676 | 1.16E-06 | 1.07761926 | 5.61E-06 |
| Fancc | red | -0.9924391 | 1.34E-12 | -0.6305339 | 1.82E-05 |
| Ptch1 | red | -0.9720768 | 1.35E-05 | -0.5198765 | 0.02855703 |
| Habp4 | red | -0.6545505 | 2.40E-08 | -0.4236633 | 0.00053776 |
| Aaed1 | skyblue | -1.2500001 | 2.86E-08 | -1.3721114 | 1.59E-09 |
| Mxd3 | yellow | 2.84303307 | 5.66E-18 | 3.27183482 | 8.23E-22 |

| | | | | | |
|---|---|---|---|---|---|
| Prelid1 | orange | 0.88750167 | 2.11E-06 | 0.63178811 | 0.00118365 |
| Pdlim7 | orange | 0.54785701 | 0.01840318 | 0.61409742 | 0.00939572 |
| Ddx41 | orange | 0.73211251 | 2.30E-09 | 0.54089299 | 1.87E-05 |
| Fam193b | red | -0.7084992 | 1.62E-06 | -0.6343562 | 2.87E-05 |
| Fastkd3 | orange | 0.60184909 | 0.00016439 | 0.33137914 | 0.05210296 |
| 1700001L19I | red | -1.4743155 | 5.80E-05 | -0.4735958 | 0.23772295 |
| Adcy2 | red | -0.774173 | 1.21E-07 | -0.8516993 | 9.49E-09 |
| Nkd2 | skyblue | -1.4244595 | 2.24E-09 | -1.3873375 | 8.95E-09 |
| Trip13 | yellow | 2.92095918 | 1.43E-27 | 2.86025272 | 5.54E-24 |
| Tppp | skyblue | -2.2451849 | 0.00011589 | -1.7871006 | 0.00305708 |
| Pdcd6 | orange | 0.58234323 | 1.57E-06 | 0.44501461 | 0.0004156 |
| Erap1 | orange | 0.8703294 | 2.80E-18 | 0.83986365 | 6.32E-17 |
| Rhobtb3 | red | -0.7249741 | 0.00085304 | -0.6544463 | 0.00340913 |
| Glrx | orange | 2.00503335 | 3.10E-12 | 1.71075557 | 5.56E-09 |
| Arsk | red | -0.9854825 | 2.55E-05 | -1.2419151 | 1.24E-07 |
| Srd5a1 | red | -0.7054896 | 0.03592985 | -1.0966631 | 0.00170943 |
| Nsun2 | orange | 0.67026066 | 0.00024591 | 0.48071934 | 0.01198581 |
| Lpcat1 | red | -0.8916868 | 9.16E-05 | -0.5685529 | 0.01781496 |
| Hapln1 | skyblue | -1.3461509 | 0.02261029 | -1.6975255 | 0.00490076 |
| Vcan | yellow | 2.26762506 | 2.01E-13 | 2.29611056 | 1.42E-13 |
| Ckmt2 | skyblue | -2.8024942 | 1.52E-09 | -2.0353778 | 2.22E-05 |
| Cd180 | yellow | 3.63136321 | 5.72E-10 | 3.25264294 | 5.41E-08 |
| Ocln | red | -0.7781958 | 9.88E-16 | -0.5744625 | 7.93E-09 |
| Smn1 | orange | 1.05401426 | 8.81E-09 | 0.69321568 | 0.00029304 |
| Mccc2 | red | -1.0048025 | 5.62E-06 | -0.9667578 | 2.10E-05 |
| Btf3 | orange | 0.68829723 | 1.46E-05 | 0.45942613 | 0.00573643 |
| Arhgef28 | red | -1.374576 | 2.93E-12 | -0.8064045 | 8.35E-05 |
| Hexb | orange | 0.9305982 | 6.53E-05 | 0.55256295 | 0.02535688 |
| Hmgcr | orange | 0.65837279 | 1.66E-08 | 0.72520602 | 7.16E-10 |
| F2rl2 | orange | 1.17274681 | 1.35E-05 | 0.76651007 | 0.01080797 |
| Iqgap2 | orange | 0.98204846 | 4.85E-11 | 0.93501133 | 7.18E-10 |
| Pde8b | skyblue | -1.7144006 | 2.65E-09 | -1.3563377 | 4.67E-06 |
| Jmy | red | -1.1550529 | 1.58E-08 | -0.8162803 | 0.00011896 |
| Depdc1b | yellow | 2.59848645 | 1.21E-07 | 3.37306594 | 2.25E-10 |
| Thbs4 | skyblue | -2.1679181 | 2.00E-05 | -1.0050062 | 0.06740693 |
| Serinc5 | red | -0.8877112 | 3.10E-06 | -0.5007286 | 0.0126766 |
| Dhfr | orange | 1.8613128 | 5.57E-11 | 1.4605111 | 7.04E-07 |
| Rasgrf2 | skyblue | -1.7675423 | 3.73E-07 | -1.2905856 | 0.00034724 |
| Cenpk | yellow | 3.0883551 | 2.35E-08 | 2.28804362 | 7.08E-05 |
| Cwc27 | orange | 0.64149818 | 0.00385747 | 0.73974818 | 0.00114513 |
| Rgs7bp | skyblue | -1.8444734 | 2.01E-05 | -1.5501652 | 0.00050088 |
| Rnf180 | orange | 1.8928958 | 4.74E-07 | 0.78988974 | 0.050507 |
| Parp8 | orange | 0.71945947 | 1.23E-08 | 0.49835859 | 0.00015162 |

| | | | | | |
|---|---|---|---|---|---|
| Hcn1 | skyblue | -1.5540875 | 1.25E-05 | -1.5536297 | 3.96E-05 |
| Fgf10 | red | -1.0493411 | 1.08E-07 | -0.8952102 | 1.19E-05 |
| Slc4a7 | orange | 0.80872476 | 2.83E-07 | 0.69533135 | 2.07E-05 |
| 4930452B06I | skyblue | -1.6570384 | 1.33E-09 | -1.6272453 | 9.32E-08 |
| Fam107a | orange | 0.92646095 | 0.00264794 | -0.3403305 | 0.32078262 |
| Acox2 | red | -1.2813349 | 2.22E-11 | -0.7940874 | 4.68E-05 |
| Gpx8 | red | -0.8775298 | 1.84E-14 | -0.5401721 | 6.25E-06 |
| BC067074 | red | -0.8121659 | 3.17E-07 | -0.3889908 | 0.02049839 |
| Fst | yellow | 3.28339787 | 1.05E-13 | 2.11665414 | 3.40E-06 |
| Kat6b | red | -0.6112611 | 0.00107421 | -0.3916544 | 0.04754225 |
| Nr1d2 | skyblue | -1.5629382 | 1.08E-11 | -1.3223052 | 1.81E-08 |
| Thrb | red | -1.3227371 | 5.41E-10 | -1.0337309 | 1.84E-06 |
| Sftpa1 | red | -1.0698106 | 0.00452703 | -0.6141471 | 0.12935999 |
| Dydc1 | skyblue | -1.3678221 | 0.02691306 | -1.5893011 | 0.00638978 |
| Dydc2 | skyblue | -2.2775564 | 0.00130476 | -0.7586577 | 0.27901795 |
| Fam213a | red | -1.1178325 | 2.96E-07 | -0.956245 | 2.03E-05 |
| Bmpr1a | red | -0.8426915 | 1.25E-08 | -0.6540982 | 1.89E-05 |
| Ldb3 | skyblue | -1.5888358 | 7.73E-06 | -1.1622613 | 0.00164872 |
| Nid2 | red | -1.0180367 | 7.75E-07 | -0.777644 | 0.00025219 |
| Dnajc9 | orange | 0.87521973 | 1.03E-10 | 0.60609792 | 1.81E-05 |
| Plau | yellow | 2.29908045 | 4.91E-14 | 2.35688288 | 1.70E-14 |
| Vcl | red | -0.629016 | 0.00274388 | -0.3065945 | 0.17980801 |
| Txndc16 | red | -1.039664 | 7.02E-09 | -0.9663312 | 1.32E-07 |
| Bmp4 | red | -0.7899316 | 6.15E-05 | -0.5070097 | 0.01448886 |
| Samd4 | red | -0.9633661 | 0.00010479 | -0.7376881 | 0.0042369 |
| Rnase4 | skyblue | -1.583519 | 2.26E-11 | -1.3506833 | 2.34E-08 |
| Dnah12 | skyblue | -1.8370718 | 2.03E-11 | -0.7702379 | 0.00669709 |
| Rnase6 | orange | 1.02387629 | 0.00017093 | 1.33353057 | 1.82E-06 |
| Gpr65 | yellow | 3.73446345 | 3.92E-32 | 3.12208697 | 3.19E-22 |
| Asb14 | red | -1.2629505 | 2.81E-06 | -1.0895886 | 9.36E-05 |
| Btd | red | -0.5700976 | 0.00059515 | -0.6262293 | 0.00022222 |
| Sema3g | skyblue | -1.323555 | 1.94E-19 | -1.2329983 | 7.30E-17 |
| Glt8d1 | red | -0.683734 | 4.73E-08 | -0.6498817 | 4.10E-07 |
| Kpna3 | orange | 0.58769394 | 0.00039731 | 0.32496912 | 0.06755748 |
| Rnaseh2b | orange | 1.00669056 | 1.36E-06 | 0.77208748 | 0.00039701 |
| Ctsb | orange | 1.48365795 | 2.98E-07 | 1.22052624 | 4.28E-05 |
| Gdf10 | skyblue | -1.8263364 | 3.25E-07 | -0.9717583 | 0.00994214 |
| Gata4 | skyblue | -1.0940134 | 0.11572865 | -1.8446454 | 0.00749588 |
| Prkcd | orange | 0.90286902 | 1.07E-08 | 0.77093169 | 1.94E-06 |
| Anxa8 | red | -0.7939498 | 0.00028253 | -0.5344051 | 0.01990462 |
| Pinx1 | orange | 0.87778255 | 0.00499725 | 0.62901599 | 0.06002996 |
| Ska3 | yellow | 2.61066455 | 3.97E-10 | 3.0901325 | 1.12E-12 |
| Mrpl57 | orange | 0.67174212 | 0.00016001 | 0.63488826 | 0.00056491 |

| | | | | | |
|---|---|---|---|---|---|
| Cdadc1 | red | -0.6465644 | 9.90E-05 | -0.5455776 | 0.00147649 |
| Amer2 | yellow | 2.26817325 | 0.00066998 | 1.5431864 | 0.02186601 |
| Spata13 | orange | 1.08892795 | 3.05E-11 | 0.88300355 | 1.47E-07 |
| Esd | orange | 0.95962216 | 1.95E-06 | 0.78641429 | 0.00016143 |
| Lcp1 | orange | 2.06436491 | 1.25E-16 | 1.65786104 | 7.24E-11 |
| Nufip1 | orange | 0.90723099 | 3.94E-08 | 0.87064014 | 3.38E-07 |
| Tsc22d1 | red | -0.945306 | 3.54E-09 | -0.9640656 | 2.67E-09 |
| Dnajc15 | orange | 0.68835782 | 4.05E-05 | 0.41175238 | 0.02163624 |
| Epsti1 | yellow | 3.52984546 | 1.80E-46 | 3.28190914 | 1.02E-39 |
| Tnfsf11 | yellow | 3.77538153 | 6.76E-12 | 2.38178862 | 2.68E-06 |
| Diap3 | orange | 1.66076908 | 1.64E-07 | 1.57691659 | 1.33E-06 |
| Scara5 | red | -1.2089614 | 7.44E-08 | -0.6668567 | 0.00466193 |
| Pbk | yellow | 2.21007129 | 1.44E-07 | 3.42443068 | 3.79E-14 |
| Esco2 | yellow | 2.74177761 | 3.78E-16 | 3.32285249 | 3.46E-20 |
| Ccdc25 | orange | 1.22990932 | 3.29E-17 | 1.14728381 | 8.02E-15 |
| Ephx2 | skyblue | -1.9424786 | 8.27E-07 | -1.7546409 | 1.54E-05 |
| Chrna2 | orange | -0.0682316 | 0.93190243 | 2.36706333 | 0.0016111 |
| Stmn4 | skyblue | -1.7381615 | 0.03156679 | -2.3673519 | 0.00960255 |
| Dpysl2 | red | -1.119044 | 9.19E-15 | -1.036768 | 1.32E-12 |
| Bnip3l | red | -0.7514927 | 2.22E-06 | -0.7466445 | 4.15E-06 |
| Ebf2 | skyblue | -2.346969 | 8.45E-07 | -1.5061869 | 0.00235244 |
| Adam7 | skyblue | -2.8819767 | 0.00031442 | -3.2820576 | 0.00013103 |
| Adamdec1 | skyblue | -1.901449 | 2.91E-11 | -0.9694609 | 0.00079054 |
| Bora | orange | 1.52857028 | 9.34E-11 | 1.46420213 | 1.21E-09 |
| Tnfrsf10b | orange | 0.25197664 | 0.30004628 | 0.63587293 | 0.00684822 |
| Bin3 | orange | 1.00220878 | 5.11E-10 | 0.76965272 | 3.78E-06 |
| Ppp3cc | orange | 0.93516818 | 1.66E-07 | 0.82647307 | 7.96E-06 |
| Slc39a14 | yellow | 2.37587872 | 7.99E-17 | 2.23150639 | 7.94E-15 |
| Hr | skyblue | -1.7040261 | 8.73E-13 | -1.123573 | 5.64E-06 |
| Sftpc | skyblue | -2.0050133 | 6.98E-05 | -1.2378531 | 0.01993711 |
| Dmtn | red | -1.3688185 | 2.31E-18 | -0.9245632 | 8.93E-09 |
| Dok2 | yellow | 2.25143081 | 2.80E-17 | 2.32383717 | 5.28E-18 |
| Gfra2 | red | -1.1390693 | 3.92E-05 | -1.1894626 | 2.85E-05 |
| Sucla2 | red | -0.6723564 | 5.58E-09 | -0.5244976 | 1.16E-05 |
| Uchl3 | orange | 0.8868092 | 0.00036705 | 0.68742065 | 0.00828655 |
| Ednrb | skyblue | -1.9077686 | 9.16E-08 | -1.5181056 | 3.71E-05 |
| Scel | red | -1.0840304 | 2.16E-18 | -0.9686134 | 1.11E-14 |
| Irg1 | black | 11.9299782 | 4.23E-18 | 9.59200186 | 2.49E-29 |
| Cldn10 | red | -1.2812045 | 4.96E-07 | -0.6252484 | 0.01999532 |
| Osmr | orange | 0.71921609 | 4.16E-06 | 0.8195583 | 2.16E-07 |
| Fyb | yellow | 3.44431529 | 4.39E-33 | 3.11959437 | 4.06E-27 |
| Dab2 | orange | 0.83911413 | 0.00110269 | 0.92787479 | 0.00040318 |
| Ttc33 | red | -0.5260279 | 0.01067507 | -0.6664574 | 0.00147814 |

| | | | | | |
|---|---|---|---|---|---|
| Rem2 | red | -0.7898483 | 0.0108876 | -1.0200811 | 0.00244522 |
| Haus4 | orange | 1.15413245 | 1.05E-17 | 0.60268319 | 2.39E-05 |
| Ajuba | red | -0.5821492 | 2.93E-09 | -0.3876337 | 0.00014561 |
| 4931414P19I | red | -0.6330367 | 3.40E-05 | -0.4643846 | 0.00406421 |
| Slc7a8 | yellow | 3.62747131 | 2.10E-19 | 3.41604343 | 3.71E-17 |
| C6 | red | -0.5932127 | 0.00376891 | -0.4925102 | 0.02075815 |
| Oxct1 | red | -0.8303979 | 9.29E-11 | -0.7585471 | 6.79E-09 |
| Psmb5 | orange | 0.70366067 | 3.19E-05 | 0.60877275 | 0.00050969 |
| Pdzd2 | red | -1.180141 | 0.00018123 | -0.7890542 | 0.01712267 |
| Slc22a17 | red | -1.341644 | 5.42E-09 | -1.1031044 | 2.91E-06 |
| Ngdn | orange | 0.67360886 | 4.69E-05 | 0.67799583 | 6.57E-05 |
| Sub1 | orange | 1.02922466 | 8.04E-07 | 0.69907059 | 0.00131726 |
| Npr3 | skyblue | -2.1054855 | 0.00643402 | -0.80452 | 0.35018466 |
| Fitm1 | red | -1.5298776 | 5.03E-05 | -1.0080293 | 0.01138333 |
| Psme1 | orange | 1.88973875 | 2.19E-45 | 1.70332928 | 6.23E-37 |
| Emc9 | red | -0.6180078 | 0.00288749 | -0.5646711 | 0.00846472 |
| Ripk3 | yellow | 2.23531182 | 1.87E-19 | 2.20482451 | 8.27E-19 |
| Zscan26 | red | -0.8479148 | 9.50E-06 | -0.717165 | 0.00027921 |
| Cmbl | red | -1.070293 | 6.90E-08 | -0.9889025 | 1.20E-06 |
| Ctnnd2 | red | -1.3298543 | 6.34E-07 | -0.9001007 | 0.00112673 |
| Tars | orange | 0.69185126 | 8.49E-09 | 0.63385834 | 2.51E-07 |
| Amacr | red | -0.8736788 | 0.00178714 | -0.7195673 | 0.01299873 |
| Brix1 | orange | 0.62363674 | 0.005692 | 0.51589241 | 0.02845156 |
| Nadk2 | red | -0.7554613 | 0.00021879 | -0.6833279 | 0.00124368 |
| Laptm4b | red | -0.8507026 | 7.13E-12 | -0.6895059 | 5.44E-08 |
| Sdc2 | skyblue | -1.6184191 | 9.61E-14 | -1.2830598 | 8.13E-09 |
| Dnah5 | skyblue | -3.3511212 | 8.52E-15 | -2.0521148 | 3.85E-06 |
| Ank | red | -0.9742563 | 6.46E-08 | -1.008361 | 3.72E-08 |
| Pabpc1 | orange | 0.60088319 | 6.54E-08 | 0.37868114 | 0.00111885 |
| Grhl2 | red | -1.2199746 | 1.17E-10 | -0.632427 | 0.00133451 |
| Rrm2b | red | -0.8955664 | 4.25E-07 | -0.8605056 | 2.11E-06 |
| Fzd6 | red | -1.1105462 | 1.23E-10 | -1.0519617 | 2.33E-09 |
| Dcaf13 | orange | 0.88060568 | 1.60E-08 | 0.68269196 | 2.39E-05 |
| Zfpm2 | red | -1.0517444 | 4.68E-06 | -0.7793091 | 0.00093507 |
| Oxr1 | red | -0.7604342 | 6.02E-08 | -0.5542571 | 0.0001381 |
| Angpt1 | skyblue | -1.3380705 | 7.69E-06 | -1.3276362 | 1.41E-05 |
| Shcbp1 | yellow | 3.98682478 | 1.76E-25 | 3.76994708 | 8.22E-22 |
| Hrsp12 | red | -1.064908 | 6.29E-07 | -0.6881415 | 0.00226759 |
| Matn2 | red | -0.9422219 | 2.03E-09 | -0.8466423 | 1.78E-07 |
| Pop1 | orange | 0.71815285 | 8.21E-05 | 0.94868086 | 3.32E-07 |
| Sybu | red | -0.8722042 | 0.00478279 | -0.8198587 | 0.00981674 |
| Myc | yellow | 2.48935079 | 2.41E-15 | 2.68404982 | 1.67E-17 |
| Mtss1 | red | -0.7516598 | 1.92E-13 | -0.5289994 | 5.81E-07 |

| | | | | | |
|---|---|---|---|---|---|
| Klhl38 | skyblue | -1.528464 | 0.01263824 | -1.8018802 | 0.00771923 |
| Atad2 | orange | 0.87551729 | 7.27E-07 | 0.82043862 | 6.87E-06 |
| Has2 | orange | 1.52587328 | 0.02477263 | 2.46636187 | 0.00055894 |
| Mtbp | orange | 1.42165595 | 1.14E-14 | 1.12356262 | 3.17E-09 |
| Mrpl13 | orange | 0.73318996 | 0.00402711 | 0.50952018 | 0.05931965 |
| Col14a1 | skyblue | -1.9981735 | 1.61E-21 | -1.2382465 | 1.04E-08 |
| Sla | yellow | 2.60621381 | 7.37E-24 | 2.5134241 | 3.98E-22 |
| Lrrc6 | skyblue | -2.5389482 | 8.41E-10 | -1.431156 | 0.00040558 |
| Adcy8 | red | -1.3245729 | 7.02E-05 | -0.7954707 | 0.02373064 |
| Fam49b | yellow | 2.19122891 | 1.21E-20 | 1.83681853 | 1.33E-14 |
| Ppara | skyblue | -1.6110043 | 3.00E-05 | -1.1320461 | 0.00446616 |
| Gtse1 | yellow | 3.00401636 | 2.34E-27 | 2.65026555 | 2.25E-21 |
| Tef | skyblue | -1.5496759 | 5.17E-05 | -1.2565197 | 0.00151231 |
| Fam83f | orange | 1.07746214 | 0.00162032 | 0.72456692 | 0.04584592 |
| Deptor | red | -1.0244126 | 5.06E-05 | -0.8926662 | 0.00061767 |
| Dnal4 | red | -0.664441 | 1.71E-14 | -0.4440027 | 7.49E-07 |
| Nptxr | red | -0.7139928 | 0.00047079 | -0.4425911 | 0.04128306 |
| Dscc1 | yellow | 2.60185393 | 5.03E-07 | 2.60067748 | 1.27E-06 |
| Enpp2 | skyblue | -1.6318772 | 2.80E-08 | -1.2243254 | 5.54E-05 |
| Sh3bp1 | orange | 1.3254862 | 4.34E-12 | 0.8687746 | 1.50E-05 |
| Parvg | yellow | 2.58145932 | 9.60E-25 | 2.49845305 | 5.30E-23 |
| C1qtnf6 | orange | 0.88985526 | 0.00010422 | 0.77199371 | 0.00112936 |
| Efcab6 | skyblue | -2.1597244 | 1.42E-08 | -1.2770208 | 0.00100776 |
| Ttll1 | red | -0.5941791 | 6.33E-05 | -0.7951688 | 2.09E-07 |
| Myh9 | orange | 0.61061699 | 0.00305522 | 0.70956958 | 0.00070867 |
| Adamts20 | skyblue | -2.1202025 | 1.38E-05 | -0.7991775 | 0.13341746 |
| Desi1 | orange | 0.60881399 | 0.00043267 | 0.66066374 | 0.00018548 |
| Aco2 | red | -0.6979137 | 3.32E-09 | -0.6058765 | 5.35E-07 |
| Vdr | orange | 0.69846004 | 0.0002143 | 0.13797781 | 0.53320068 |
| Col2a1 | skyblue | -3.8175945 | 0.00070529 | -2.7367511 | 0.01827306 |
| Nckap1l | yellow | 2.40492885 | 1.44E-18 | 2.32027554 | 3.41E-17 |
| Pde1b | yellow | 2.0278518 | 6.13E-09 | 2.10219881 | 2.49E-09 |
| Litaf | orange | 2.0143322 | 8.60E-17 | 1.94104111 | 1.62E-15 |
| Prm1 | black | 4.68718221 | 0.00019339 | 5.65708094 | 8.76E-05 |
| Nubp1 | orange | 1.07678263 | 3.67E-08 | 1.12219508 | 1.56E-08 |
| Ciita | yellow | 1.89136676 | 1.70E-22 | 2.0700855 | 1.00E-26 |
| Emp2 | red | -0.7439188 | 2.12E-06 | -0.4423864 | 0.00735002 |
| Bcl6 | orange | -0.1366113 | 0.3415519 | 0.63758222 | 1.65E-06 |
| Cldn1 | red | -1.2451169 | 4.83E-06 | -0.9404988 | 0.00086388 |
| 4930562C15 | red | -1.297508 | 8.20E-07 | -0.5190866 | 0.05939231 |
| Srl | skyblue | -1.5134079 | 3.39E-06 | -1.1906908 | 0.00042109 |
| Fgf12 | skyblue | -1.5082009 | 0.13092093 | -2.642464 | 0.00756147 |
| Hrasls | skyblue | -2.3556193 | 5.22E-08 | -2.1540462 | 2.52E-06 |

| | | | | | |
|---|---|---|---|---|---|
| Hes1 | red | -0.5942906 | 0.00192251 | -0.3558076 | 0.08201961 |
| Mefv | black | 4.89327972 | 8.68E-29 | 6.06842196 | 2.93E-35 |
| Tmem44 | skyblue | -1.3761909 | 2.47E-08 | -1.1926531 | 2.42E-06 |
| Lsg1 | orange | 0.69388528 | 1.97E-06 | 0.58763264 | 9.54E-05 |
| 4930451G09 | skyblue | -1.9155551 | 4.62E-06 | -0.7318126 | 0.08877305 |
| Gpt | skyblue | -1.6726282 | 8.59E-07 | -1.2315044 | 0.00047988 |
| Apod | yellow | 2.38213375 | 1.74E-37 | 2.40976234 | 3.28E-38 |
| Sharpin | orange | 0.74896425 | 5.15E-11 | 0.66591915 | 1.01E-08 |
| Hgh1 | orange | 0.39067058 | 0.09814119 | 0.77661928 | 0.00083356 |
| Bop1 | orange | 0.70585893 | 0.00045799 | 0.79391812 | 0.00010983 |
| Slc52a2 | orange | 0.66714839 | 4.51E-05 | 0.40352642 | 0.02006499 |
| Oplah | red | -0.7897095 | 4.69E-05 | -0.6417985 | 0.00137202 |
| Grina | orange | 1.0492035 | 1.86E-20 | 0.90119999 | 3.34E-15 |
| Tsta3 | orange | 0.62412015 | 1.28E-06 | 0.41213664 | 0.00219345 |
| Gsdmd | orange | 1.5414346 | 3.15E-33 | 1.28793387 | 2.25E-23 |
| Gpihbp1 | red | -0.7359135 | 0.00256302 | -0.7634434 | 0.0022249 |
| Ly6c2 | black | 5.64537496 | 5.03E-61 | 5.38729598 | 2.58E-55 |
| Ly6i | black | 5.48203406 | 1.98E-62 | 4.60788894 | 1.57E-45 |
| Ly6e | orange | 1.24373303 | 2.17E-29 | 1.07294893 | 5.77E-22 |
| Lynx1 | red | -1.433952 | 2.12E-12 | -0.8539655 | 6.15E-05 |
| Lypd2 | skyblue | -1.8332717 | 7.73E-07 | -1.3896274 | 0.0002953 |
| Psca | yellow | 2.61280025 | 0.00048717 | 2.27957318 | 0.0058691 |
| Arc | orange | 1.69393716 | 6.92E-07 | 1.04415324 | 0.00291651 |
| Ptk2 | red | -0.6856994 | 2.38E-13 | -0.4290851 | 1.06E-05 |
| Mapk12 | red | -0.9086602 | 1.39E-09 | -0.5836914 | 0.00018905 |
| Tymp | skyblue | -2.8190755 | 7.81E-15 | -1.8273763 | 9.16E-07 |
| Rabl2 | skyblue | -1.5080745 | 1.31E-11 | -0.8915202 | 0.00011365 |
| Acr | orange | 1.45313936 | 9.28E-06 | 1.13623805 | 0.00070039 |
| Kif21a | red | -1.3689115 | 8.41E-10 | -1.0025958 | 1.38E-05 |
| Alcam | red | -0.9541777 | 5.40E-07 | -0.6197107 | 0.00181834 |
| Cblb | orange | 1.13201618 | 2.14E-11 | 0.61561658 | 0.00052865 |
| Dubr | red | -0.7519308 | 6.51E-05 | -0.7285927 | 0.00017601 |
| Retnlb | black | 6.10274154 | 9.91E-05 | 4.45870685 | 0.00279007 |
| Retnlg | yellow | 3.22767566 | 6.99E-11 | 2.69030485 | 5.31E-08 |
| Morc1 | orange | 0.51954581 | 0.41502942 | 1.94453906 | 0.00144523 |
| Pvrl3 | red | -1.3038402 | 4.44E-09 | -1.0803025 | 2.23E-06 |
| Cd96 | yellow | 2.65412724 | 4.23E-13 | 2.74119712 | 3.02E-13 |
| Ccdc80 | red | -0.4163734 | 0.04264056 | -0.6151685 | 0.00273085 |
| Cd200r1 | orange | 1.67549012 | 0.00045394 | 1.80228574 | 0.00023283 |
| Mcm4 | orange | 1.11923318 | 1.20E-16 | 0.73781405 | 1.46E-07 |
| Snai2 | red | -0.3922794 | 0.1024344 | -1.0822206 | 3.64E-06 |
| Mpv17l | red | -1.4671345 | 1.39E-07 | -0.9814612 | 0.0007185 |
| B3gnt5 | orange | 0.93666893 | 9.62E-06 | 0.92578593 | 1.85E-05 |

| | | | | | |
|---|---|---|---|---|---|
| Sidt1 | yellow | 1.86169214 | 6.73E-17 | 2.24527857 | 1.96E-23 |
| 2610015P09I | red | -1.0725252 | 7.47E-07 | -0.7005959 | 0.00204054 |
| Mrpl40 | orange | 0.61393796 | 0.00035694 | 0.37052663 | 0.04316663 |
| Gbe1 | red | -0.6888035 | 3.98E-05 | -0.616804 | 0.00035466 |
| Zbtb20 | red | -0.8337556 | 1.01E-05 | -0.9168468 | 1.79E-06 |
| Pmm2 | orange | 0.67962128 | 1.80E-09 | 0.5786756 | 6.11E-07 |
| St3gal6 | red | -1.1464169 | 1.86E-12 | -0.886748 | 1.18E-07 |
| Adgrg7 | black | 6.7836695 | 1.11E-05 | 3.09122928 | 0.0229785 |
| Slc7a4 | skyblue | -2.0204689 | 9.20E-09 | -1.3891608 | 0.00013253 |
| P2rx6 | skyblue | -2.8347775 | 1.84E-14 | -1.8012105 | 1.81E-06 |
| Lrrc74b | skyblue | -2.6568773 | 2.80E-09 | -1.4066332 | 0.00224185 |
| Aifm3 | red | -1.1224154 | 0.00099425 | -0.9305134 | 0.00884865 |
| Sdf2l1 | orange | 1.40191377 | 5.75E-08 | 1.37247878 | 2.02E-07 |
| Spag6 | skyblue | -2.4698531 | 4.08E-19 | -1.5494644 | 1.28E-08 |
| B4galt4 | red | -1.1572726 | 8.85E-09 | -1.1716845 | 9.14E-09 |
| Maats1 | skyblue | -1.5924026 | 2.48E-10 | -0.858971 | 0.00095638 |
| Nr1i2 | skyblue | -2.6255201 | 3.99E-06 | -1.9271495 | 8.54E-05 |
| Umps | orange | 0.94816364 | 1.62E-07 | 0.79420652 | 2.07E-05 |
| Muc13 | yellow | 1.85460934 | 0.00111379 | 2.05240282 | 0.0006244 |
| Rabl3 | orange | 0.48228364 | 0.02231504 | 0.79674633 | 0.00015621 |
| Hcls1 | yellow | 3.04414216 | 1.03E-23 | 2.86493745 | 5.68E-21 |
| Mylk | red | -0.7239023 | 5.44E-07 | -0.5982325 | 5.91E-05 |
| Adcy5 | skyblue | -1.6722857 | 2.93E-07 | -1.0732482 | 0.00158517 |
| Clcn2 | red | -0.8183064 | 2.57E-05 | -0.5961694 | 0.00304926 |
| Pdia5 | red | -0.7954371 | 1.91E-06 | -0.7211969 | 3.06E-05 |
| Dirc2 | red | -1.039713 | 7.03E-06 | -0.9597084 | 5.32E-05 |
| Dgkg | skyblue | -1.6206759 | 4.63E-05 | -0.7070558 | 0.0889971 |
| D16Ertd472e | orange | 0.78508037 | 2.32E-11 | 0.59507221 | 9.83E-07 |
| Cxadr | red | -1.0956749 | 4.94E-20 | -0.6422128 | 2.37E-07 |
| Usp25 | orange | 0.72386983 | 3.40E-14 | 0.57993243 | 2.97E-09 |
| Fetub | skyblue | -1.9788692 | 5.87E-06 | -1.1389284 | 0.01312875 |
| Samsn1 | yellow | 2.86962834 | 4.73E-31 | 2.4977848 | 1.74E-23 |
| Adipoq | skyblue | -2.8702884 | 0.00041214 | -1.7578817 | 0.04078991 |
| Rfc4 | orange | 1.35755666 | 7.49E-09 | 1.32176812 | 4.97E-08 |
| Robo1 | red | -0.9212905 | 5.07E-05 | -0.9180354 | 7.59E-05 |
| St6gal1 | red | -0.6172828 | 2.37E-09 | -0.3131955 | 0.00410127 |
| Adamts5 | red | -1.5486896 | 9.87E-09 | -1.0040519 | 0.0003459 |
| Cd86 | yellow | 2.76196345 | 6.32E-15 | 2.41030635 | 2.16E-11 |
| Parp9 | yellow | 2.54673485 | 5.74E-78 | 2.35310213 | 1.16E-66 |
| Pigp | red | -0.782442 | 1.22E-05 | -0.7652902 | 3.15E-05 |
| Ripply3 | red | -1.1219659 | 1.49E-06 | -0.8375265 | 0.00053986 |
| Chaf1b | yellow | 3.04138091 | 6.25E-29 | 2.25532228 | 8.05E-17 |
| Cbr3 | red | -0.908083 | 4.16E-06 | -0.747583 | 0.00023834 |

| | | | | | |
|---|---|---|---|---|---|
| Clic6 | skyblue | -2.8159993 | 1.72E-34 | -1.5647125 | 1.79E-11 |
| Rcan1 | orange | 0.47470052 | 0.00269891 | 0.61041803 | 0.0001394 |
| Runx1 | orange | 0.6368799 | 0.0001081 | 0.51274317 | 0.00263527 |
| Itsn1 | red | -0.723339 | 1.34E-12 | -0.4004555 | 0.0001901 |
| Donson | orange | 0.6106862 | 8.19E-06 | 0.35724551 | 0.01503108 |
| Tmem50b | red | -1.050985 | 6.35E-09 | -0.895784 | 1.34E-06 |
| Mis18a | orange | 0.9421183 | 5.07E-08 | 0.75488522 | 2.75E-05 |
| Adcy6 | red | -0.7826924 | 4.58E-06 | -0.6032185 | 0.00065367 |
| Enah | red | -0.8917351 | 1.41E-08 | -0.3927187 | 0.01905571 |
| Wnt10b | skyblue | -2.4928809 | 1.19E-11 | -1.6903959 | 7.79E-06 |
| Tuba1b | orange | 0.81017673 | 1.47E-08 | 0.66182844 | 7.01E-06 |
| Prpf40b | red | -0.664335 | 7.38E-05 | -0.6494947 | 0.00015394 |
| Faim2 | skyblue | -3.1936677 | 1.16E-07 | -2.22811 | 0.00036073 |
| Racgap1 | yellow | 2.84757332 | 5.57E-53 | 2.73808554 | 4.80E-48 |
| Asic1 | red | -0.6139943 | 0.00129487 | -0.4260496 | 0.03672948 |
| Gpd1 | skyblue | -2.5802999 | 4.36E-05 | -1.8953287 | 0.00388485 |
| Cela1 | yellow | 2.21129029 | 1.91E-05 | 1.73751956 | 0.00126509 |
| Krt7 | orange | 0.90395957 | 5.28E-05 | 1.09752313 | 1.20E-06 |
| Krt6b | skyblue | -2.3457402 | 0.0040136 | -1.2278751 | 0.16695183 |
| Krt18 | orange | 1.97405063 | 1.05E-11 | 1.81536495 | 7.37E-10 |
| Csad | red | -1.0678114 | 4.00E-08 | -0.8533969 | 2.07E-05 |
| Soat2 | black | 4.31312925 | 9.01E-37 | 4.82865854 | 4.32E-44 |
| Igfbp6 | skyblue | -1.8845916 | 0.00072961 | -1.3644472 | 0.01948362 |
| Prr13 | orange | 1.02852582 | 5.13E-09 | 0.8516071 | 2.58E-06 |
| Map3k12 | red | -0.6613866 | 2.32E-05 | -0.385633 | 0.01812528 |
| Calcoco1 | red | -1.328262 | 1.25E-05 | -1.0606792 | 0.00074643 |
| Sncg | red | -0.5266814 | 0.01303527 | -0.7372584 | 0.00042894 |
| Cdkn1a | yellow | 2.65926043 | 7.37E-58 | 2.2225687 | 1.69E-40 |
| Cxcl13 | yellow | 3.11378354 | 2.92E-06 | 3.25384392 | 1.53E-06 |
| Lrrc71 | red | -1.3786842 | 1.70E-07 | -0.5585856 | 0.04703664 |
| Noct | yellow | 2.25344633 | 2.92E-11 | 1.88800269 | 4.87E-08 |
| Fhl1 | skyblue | -1.5068859 | 1.13E-09 | -1.103569 | 1.61E-05 |
| Msrb2 | red | -1.4665975 | 2.45E-11 | -1.0483231 | 2.90E-06 |
| Rfc2 | orange | 0.63872866 | 7.83E-07 | 0.42070565 | 0.00183353 |
| Denr | orange | 0.59780398 | 0.00051672 | 0.41147527 | 0.0233534 |
| Prmt5 | orange | 0.58796664 | 8.13E-05 | 0.43442391 | 0.00523467 |
| Gzma | yellow | 4.23448179 | 1.14E-55 | 3.82118602 | 4.93E-45 |
| Wrb | red | -0.8774686 | 3.76E-05 | -0.8032708 | 0.00023368 |
| Ivns1abp | red | -0.6721866 | 0.00050117 | -0.6420986 | 0.00121064 |
| Slc38a1 | orange | 0.83702525 | 9.94E-08 | 0.81182732 | 3.82E-07 |
| Vwa5a | orange | 0.70571157 | 1.06E-14 | 0.62408605 | 1.78E-11 |
| P3h3 | red | -0.8543696 | 1.56E-10 | -0.6152792 | 8.51E-06 |
| Il15ra | orange | 1.49766355 | 5.43E-13 | 1.3253632 | 3.52E-10 |

| | | | | | |
|---|---|---|---|---|---|
| Serping1 | orange | 1.03549099 | 1.12E-14 | 0.89603263 | 4.85E-11 |
| Kcnk5 | orange | 0.57560807 | 0.00026213 | 0.84951975 | 9.42E-08 |
| Parp3 | orange | 1.1389542 | 1.09E-11 | 1.25937235 | 6.45E-14 |
| Frs3 | red | -0.4593102 | 0.03380915 | -0.7309285 | 0.00082163 |
| Cd4 | yellow | 2.06879966 | 1.91E-06 | 3.26071601 | 4.19E-14 |
| Twf2 | orange | 0.74573877 | 8.95E-05 | 0.39638974 | 0.05196715 |
| Ube2j2 | orange | 0.81941162 | 7.16E-07 | 0.72465518 | 2.06E-05 |
| March5 | orange | 0.74263998 | 2.47E-08 | 0.70584021 | 2.01E-07 |
| Ache | red | -1.5526839 | 0.00012279 | -0.524617 | 0.24483723 |
| Wfdc1 | skyblue | -2.8052143 | 1.21E-21 | -1.8527321 | 6.24E-10 |
| Mx2 | yellow | 4.03589189 | 9.89E-51 | 3.73047103 | 5.22E-44 |
| Poc1a | orange | 1.30547953 | 1.60E-08 | 1.04159812 | 1.39E-05 |
| Clec4n | orange | 1.80118371 | 3.14E-05 | 1.4620219 | 0.0010895 |
| Slc17a9 | orange | 0.63012134 | 0.00012834 | 0.61116907 | 0.00031153 |
| Pisd | orange | 0.6935434 | 2.70E-05 | 0.71070859 | 2.55E-05 |
| Celsr3 | yellow | 2.98346362 | 2.58E-05 | 1.70784645 | 0.00922741 |
| Cdca3 | yellow | 2.94929951 | 3.01E-19 | 3.16629718 | 2.47E-21 |
| Fam132a | red | -0.529878 | 0.0104078 | -0.6763239 | 0.00130184 |
| Mrps23 | orange | 0.68709347 | 0.00067242 | 0.45990657 | 0.03158392 |
| Pigx | orange | 0.87004153 | 1.03E-05 | 0.56257333 | 0.00677079 |
| Rsph3b | red | -0.7626499 | 2.51E-06 | -0.733179 | 1.36E-05 |
| Park2 | red | -0.7557113 | 0.00747588 | -0.6008842 | 0.0399142 |
| Slc22a3 | skyblue | -1.184899 | 0.00025909 | -1.363816 | 4.30E-05 |
| Acat2 | red | -0.5471504 | 0.01538104 | -0.6274224 | 0.00618195 |
| Lnpep | orange | 0.62124948 | 1.26E-05 | 0.43038697 | 0.00378691 |
| Pde10a | orange | 1.55855925 | 0.00078286 | 1.46720595 | 0.00198098 |
| Smoc2 | red | -0.6785074 | 2.11E-05 | -0.2909365 | 0.09216983 |
| Mmp25 | yellow | 3.36367474 | 1.56E-14 | 2.84975241 | 5.54E-11 |
| Hcfc1r1 | red | -0.5864993 | 8.89E-06 | -0.5195728 | 0.00015394 |
| Tnfrsf12a | yellow | 2.54968292 | 4.10E-13 | 2.7623898 | 4.64E-15 |
| Pkmyt1 | orange | 1.25971594 | 2.10E-08 | 1.20892551 | 1.51E-07 |
| Slc25a27 | red | -1.2265236 | 0.00032765 | -1.1017198 | 0.00172844 |
| Pla2g7 | yellow | 3.82216061 | 1.96E-19 | 3.59164926 | 3.93E-17 |
| Cenpq | orange | 1.38205521 | 1.36E-06 | 1.08107175 | 0.00030601 |
| Mut | red | -0.9062704 | 4.62E-05 | -0.6876849 | 0.00291295 |
| Satb1 | orange | 0.12738112 | 0.87132917 | 1.87981194 | 0.00525535 |
| Sgol1 | yellow | 3.16755172 | 6.60E-16 | 3.7055058 | 5.48E-19 |
| Slc29a1 | red | -0.8564042 | 6.50E-13 | -0.857583 | 8.54E-13 |
| Nfkbie | yellow | 2.25691331 | 1.83E-19 | 2.18477474 | 3.79E-18 |
| Tcte1 | skyblue | -2.7462822 | 3.36E-15 | -1.3341902 | 0.00012057 |
| Vegfa | red | -1.1115389 | 1.22E-09 | -0.7156787 | 0.00017263 |
| Gtpbp2 | orange | 0.69296062 | 8.03E-09 | 0.72067714 | 3.02E-09 |
| Enpp5 | red | -1.2460665 | 1.19E-06 | -0.9538758 | 0.00032887 |

| | | | | | |
|---|---|---|---|---|---|
| Cyp39a1 | skyblue | -1.9101361 | 1.08E-06 | -1.7074819 | 2.10E-05 |
| Rsph9 | red | -0.9266356 | 1.25E-05 | -0.6746019 | 0.00212163 |
| Crip3 | skyblue | -2.0441336 | 0.00014492 | -0.6431994 | 0.27388445 |
| Guca1a | yellow | 2.19687787 | 0.00284771 | 2.11157015 | 0.00590307 |
| Bysl | orange | 0.71654618 | 0.00022702 | 0.70982282 | 0.00037536 |
| Trem2 | yellow | 4.05376491 | 2.00E-15 | 3.22008859 | 6.34E-10 |
| Kif6 | skyblue | -1.855221 | 4.39E-13 | -0.9943887 | 2.44E-05 |
| Ppil1 | orange | 0.84908657 | 1.46E-08 | 0.66813252 | 1.71E-05 |
| Pi16 | skyblue | -1.5591768 | 8.80E-19 | -0.7815172 | 2.28E-05 |
| Fgd2 | orange | 1.66730204 | 1.24E-12 | 1.65993347 | 2.46E-12 |
| Pim1 | yellow | 2.36666466 | 6.63E-46 | 2.39564391 | 5.41E-47 |
| Glo1 | red | -0.6312045 | 3.02E-06 | -0.4756646 | 0.00071618 |
| Glp1r | skyblue | -2.3063579 | 1.63E-11 | -1.820264 | 2.12E-07 |
| Rsph1 | skyblue | -1.8270041 | 2.93E-14 | -1.2613459 | 3.38E-07 |
| Tmprss3 | black | 5.19385602 | 0.00049886 | 3.23219885 | 0.011448 |
| Cbs | skyblue | -1.7344979 | 0.00038752 | -0.9057877 | 0.07848839 |
| Cryaa | skyblue | -1.3806175 | 0.01623502 | -1.5224562 | 0.00984258 |
| Epb41l3 | red | -1.1013619 | 8.90E-10 | -0.9444437 | 3.47E-07 |
| Myom1 | red | -0.6803685 | 0.00131962 | -0.5792142 | 0.00834943 |
| Emilin2 | orange | 1.13064798 | 0.00263134 | 1.23265968 | 0.00130537 |
| Smchd1 | orange | 0.74633113 | 1.77E-07 | 0.68168293 | 3.41E-06 |
| Ndc80 | yellow | 3.07660915 | 6.19E-22 | 2.74745366 | 5.59E-17 |
| Clip4 | orange | 0.58661252 | 0.00080747 | 0.55083737 | 0.0022494 |
| Lbh | red | -0.7073183 | 1.47E-08 | -0.607374 | 2.13E-06 |
| Xdh | orange | 1.77129058 | 3.70E-27 | 1.41406977 | 1.70E-17 |
| Crim1 | red | -0.9943608 | 3.73E-08 | -0.6141618 | 0.00114513 |
| Vit | skyblue | -1.2707916 | 0.00080969 | -1.56601 | 7.97E-05 |
| Eif2ak2 | yellow | 2.16723741 | 2.40E-61 | 1.95349547 | 1.06E-49 |
| Qpct | red | -0.6822338 | 0.00044354 | -0.5862392 | 0.00398773 |
| Man2a1 | orange | 0.82092412 | 1.10E-13 | 0.78793716 | 1.68E-12 |
| Twsg1 | red | -1.0787094 | 7.38E-11 | -0.8533658 | 5.34E-07 |
| Nrxn1 | red | -1.3235394 | 0.0007653 | -1.0375861 | 0.01289517 |
| Atp6v0c | orange | 0.73126076 | 5.97E-05 | 0.6780523 | 0.00029914 |
| Prss30 | orange | 1.69601021 | 0.0020602 | 1.58999745 | 0.00393318 |
| Prepl | red | -0.6514393 | 3.04E-06 | -0.6619944 | 3.12E-06 |
| Abca3 | red | -0.8275915 | 0.00338054 | -0.4247662 | 0.16516266 |
| Eci1 | red | -0.835036 | 2.68E-09 | -0.7046706 | 9.76E-07 |
| Epas1 | red | -0.9592363 | 0.00032685 | -0.7452671 | 0.00724547 |
| Spsb3 | orange | 0.45627774 | 0.00421312 | 0.58277505 | 0.00032778 |
| C3 | orange | 1.33365881 | 1.41E-10 | 1.03214527 | 1.41E-06 |
| Hn1l | orange | 0.99826 | 0.00037649 | 1.08682662 | 0.00014749 |
| Tmem204 | red | -1.3314558 | 1.17E-05 | -1.1181718 | 0.00035956 |
| Tekt4 | red | -1.1771247 | 1.63E-07 | -0.6435108 | 0.00555548 |

| | | | | | |
|---|---|---|---|---|---|
| Sox8 | skyblue | -2.1631303 | 0.00019057 | -0.7656933 | 0.17255184 |
| Rgs11 | skyblue | -1.9484871 | 1.32E-08 | -1.1660914 | 0.00104977 |
| Itfg3 | red | -0.6252554 | 2.90E-06 | -0.4644847 | 0.00081745 |
| Bnip1 | orange | 0.61824936 | 0.00010242 | 0.62264177 | 0.00015865 |
| Rfx2 | red | -1.0042495 | 2.86E-12 | -0.6968342 | 1.93E-06 |
| Snrpc | orange | 0.73570254 | 7.16E-08 | 0.72647924 | 2.04E-07 |
| Anks1 | red | -0.6092356 | 2.30E-09 | -0.298273 | 0.00565549 |
| Fkbp5 | orange | 2.11318748 | 2.77E-34 | 0.7793772 | 2.19E-05 |
| Nudt12 | red | -1.3269955 | 3.35E-07 | -0.8929119 | 0.00082163 |
| Bambi | red | -0.9168111 | 1.47E-08 | -0.4870555 | 0.00379168 |
| Map3k8 | orange | 1.42088488 | 2.95E-16 | 1.24109833 | 1.61E-12 |
| Tmem178 | red | -1.137499 | 3.87E-11 | -0.8908966 | 5.52E-07 |
| Pkdcc | red | -0.6112491 | 0.00013295 | -0.3795489 | 0.02545113 |
| Dync2li1 | red | -1.2293357 | 3.36E-15 | -0.9093649 | 1.77E-08 |
| Zfp871 | orange | 0.65135353 | 0.00339873 | 0.95567775 | 2.14E-05 |
| Adamts10 | red | -1.1558388 | 6.88E-09 | -0.8177899 | 7.70E-05 |
| Myo1f | yellow | 2.46726015 | 3.29E-18 | 2.28784221 | 1.28E-15 |
| Kifc5b | orange | 1.6113146 | 1.05E-14 | 1.97722691 | 5.52E-20 |
| Dtna | red | -0.8244619 | 0.00352942 | -0.6362129 | 0.03106623 |
| Cdh2 | skyblue | -1.727063 | 7.39E-05 | -2.0176642 | 5.48E-06 |
| Tapbp | orange | 1.99665507 | 5.05E-25 | 1.82030916 | 7.76E-21 |
| Wdr46 | orange | 0.68307396 | 0.00040792 | 0.65307293 | 0.00097615 |
| Dsc2 | red | -0.7062699 | 0.0878756 | -1.1720062 | 0.00417826 |
| Psmb8 | yellow | 3.01856519 | 1.68E-54 | 2.75484889 | 2.02E-45 |
| Tap2 | yellow | 2.08308984 | 4.88E-75 | 1.97125336 | 3.94E-67 |
| Btnl2 | yellow | 1.88928076 | 0.00088049 | 2.08605286 | 0.00069676 |
| Tmem173 | yellow | 2.89255986 | 2.10E-35 | 2.77555401 | 1.56E-32 |
| Slc23a1 | skyblue | -1.622066 | 3.73E-06 | -0.8994844 | 0.01412653 |
| Etf1 | orange | 0.67646623 | 1.72E-06 | 0.49483613 | 0.00076104 |
| Gfra3 | skyblue | -2.283517 | 4.33E-05 | -1.2917564 | 0.01957193 |
| C2 | orange | 2.02505212 | 1.37E-21 | 1.80820036 | 5.65E-17 |
| Epb41l4a | red | -0.9863455 | 3.31E-09 | -0.694332 | 6.05E-05 |
| Tslp | skyblue | -1.1445495 | 0.00080318 | -1.6057872 | 8.93E-06 |
| Proc | skyblue | -3.5638041 | 0.0001771 | -2.2268493 | 0.01212049 |
| Csnk2b | orange | 0.73061185 | 2.26E-11 | 0.46606411 | 4.33E-05 |
| Lims2 | red | -0.9020336 | 0.00024354 | -0.9154277 | 0.00027264 |
| Aif1 | black | 4.8784305 | 9.26E-42 | 4.58583382 | 4.83E-36 |
| Ltb | orange | 1.66048059 | 1.17E-10 | 1.54148598 | 5.12E-09 |
| Tnf | black | 5.17196877 | 3.69E-30 | 3.20393407 | 1.32E-14 |
| Riok3 | orange | 0.6458367 | 2.09E-09 | 0.45634969 | 4.41E-05 |
| Psors1c2 | skyblue | -1.8980118 | 0.00116824 | -1.0520174 | 0.09369313 |
| Aqp4 | red | -1.4676966 | 2.05E-06 | -0.9086295 | 0.00443407 |
| Mrpl27 | orange | 0.59152821 | 9.72E-05 | 0.61198987 | 9.32E-05 |

| | | | | | |
|---|---|---|---|---|---|
| Lama3 | skyblue | -1.6676137 | 3.50E-07 | -0.9588506 | 0.00533826 |
| Ttc39c | orange | 1.63143703 | 4.36E-17 | 1.35333328 | 7.13E-12 |
| Spry4 | orange | 0.90182578 | 0.00013573 | 0.87067746 | 0.00033419 |
| Cabyr | orange | 2.07949765 | 2.07E-05 | 0.69161244 | 0.19660381 |
| Nr3c1 | red | -0.7137958 | 9.62E-08 | -0.451628 | 0.0012424 |
| Mrps18b | orange | 0.75199421 | 0.00128006 | 0.71303942 | 0.0028161 |
| Pcdh12 | skyblue | -2.5822005 | 1.01E-22 | -1.6276192 | 1.81E-09 |
| 0610009O20 | red | -0.6613397 | 3.07E-05 | -0.5119259 | 0.0018487 |
| Arap3 | red | -1.0195247 | 3.61E-08 | -0.7919093 | 3.37E-05 |
| Trim26 | orange | 0.8516767 | 2.01E-13 | 0.68999388 | 5.84E-09 |
| H2-M5 | skyblue | -1.7724663 | 0.00014104 | -0.9615139 | 0.04477413 |
| Myot | red | -1.0848355 | 0.00086083 | -0.6540122 | 0.06019612 |
| Dcp2 | orange | 0.92765855 | 2.98E-16 | 0.86870263 | 3.18E-14 |
| Mal2 | red | -1.0516509 | 2.41E-06 | -0.5489853 | 0.02049839 |
| Ap3s1 | orange | 0.62343485 | 0.0004463 | 0.57428454 | 0.00180783 |
| Hbegf | orange | 1.14520531 | 0.00019393 | 1.30860499 | 2.79E-05 |
| Tcerg1 | orange | 0.90390256 | 3.04E-07 | 0.60631992 | 0.00099874 |
| Ppp2r2b | skyblue | -1.8682335 | 3.04E-06 | -1.2832002 | 0.00135555 |
| Hsd17b4 | red | -0.6656249 | 1.27E-06 | -0.5334613 | 0.00017272 |
| Rab27b | red | -1.0158587 | 2.09E-06 | -0.7381858 | 0.00090468 |
| Mbd2 | orange | 0.80200182 | 2.69E-07 | 0.7131043 | 8.21E-06 |
| Sec11c | orange | 0.69222312 | 6.15E-16 | 0.61692625 | 1.30E-12 |
| Grp | red | -1.3053054 | 0.00042859 | -0.7130922 | 0.07022881 |
| Pmaip1 | orange | 1.42471646 | 4.64E-14 | 0.68955599 | 0.00051781 |
| Impa2 | orange | 1.20209906 | 4.21E-05 | 1.21275686 | 5.66E-05 |
| Cidea | skyblue | -3.1430831 | 0.00043235 | -2.3814627 | 0.01053995 |
| Lox | orange | 2.06109172 | 5.07E-22 | 1.82227507 | 2.59E-17 |
| Sncaip | skyblue | -1.7205482 | 8.51E-05 | -1.8314948 | 4.43E-05 |
| Ptpn2 | orange | 1.35563981 | 4.16E-19 | 1.26855251 | 1.28E-16 |
| Cep192 | orange | 1.04568739 | 5.63E-15 | 0.88934316 | 7.05E-11 |
| Me2 | orange | 0.92686083 | 1.58E-14 | 0.6673304 | 8.65E-08 |
| Pcyox1l | red | -0.8742873 | 6.39E-12 | -0.7715266 | 2.81E-09 |
| Nars | orange | 0.59708069 | 2.22E-05 | 0.52142624 | 0.00032775 |
| Fech | red | -0.6159182 | 8.49E-08 | -0.4164825 | 0.00050672 |
| Lmnb1 | yellow | 2.08877091 | 2.69E-35 | 2.08365947 | 9.31E-35 |
| C330018D20 | red | -0.2223351 | 0.3313705 | -0.7378507 | 0.00067446 |
| Fbn2 | skyblue | -2.2661357 | 0.00135711 | -0.9261238 | 0.23723295 |
| Cd74 | orange | 1.26209044 | 3.44E-14 | 1.03327037 | 1.22E-09 |
| Tcof1 | orange | 1.12765344 | 1.15E-11 | 1.06104684 | 3.21E-10 |
| Camk2a | skyblue | -2.256547 | 5.46E-09 | -1.7306168 | 1.55E-05 |
| Pdgfrb | red | -0.5853218 | 0.00211361 | -0.3739231 | 0.06415961 |
| Csf1r | orange | 0.70299414 | 0.00609414 | 0.41920087 | 0.12714474 |
| Psat1 | orange | 1.89543197 | 1.68E-26 | 1.67302856 | 9.83E-21 |

| | | | | | |
|---|---|---|---|---|---|
| Cndp2 | orange | 1.67197334 | 2.21E-13 | 1.42464472 | 8.66E-10 |
| Cyb5a | red | -0.9980745 | 1.81E-07 | -0.8457205 | 1.72E-05 |
| Scgb1a1 | skyblue | -2.1567231 | 1.70E-08 | -1.1431356 | 0.00458865 |
| Anxa1 | orange | 0.8595003 | 1.78E-05 | 0.81086548 | 7.88E-05 |
| Incenp | yellow | 2.09062243 | 3.50E-23 | 2.17408263 | 9.20E-25 |
| Rab3il1 | orange | 1.98746982 | 9.70E-16 | 1.8610749 | 9.10E-14 |
| Fads2 | red | -0.584923 | 0.00101907 | -0.5157363 | 0.00498627 |
| Cd5 | yellow | 2.67345559 | 5.91E-25 | 2.96530438 | 4.96E-30 |
| Cd6 | yellow | 3.9278196 | 1.01E-44 | 3.95886855 | 3.92E-45 |
| Ms4a7 | black | 4.68462702 | 9.44E-13 | 4.35025386 | 7.55E-11 |
| Ms4a4c | black | 5.99696033 | 6.10E-48 | 5.829883 | 2.39E-44 |
| Ms4a6b | yellow | 4.00905498 | 4.06E-45 | 3.70261893 | 1.26E-38 |
| Ms4a6d | black | 5.17471271 | 2.13E-19 | 4.69469714 | 5.02E-16 |
| Mrpl16 | orange | 0.78638739 | 2.00E-05 | 0.42510625 | 0.03102756 |
| Fam111a | orange | 1.66101333 | 3.28E-15 | 1.36596577 | 2.64E-10 |
| Lpxn | yellow | 3.64544097 | 2.25E-23 | 3.42578153 | 1.75E-20 |
| Ostf1 | orange | 0.64498621 | 9.49E-10 | 0.45255388 | 3.53E-05 |
| Trpm6 | red | -0.619189 | 0.0022015 | -0.2329624 | 0.29937973 |
| Ms4a8a | yellow | 2.76636767 | 1.07E-13 | 2.47204158 | 5.23E-11 |
| Ccdc86 | orange | 1.4739752 | 1.55E-07 | 1.40183118 | 1.06E-06 |
| Slc15a3 | yellow | 3.05757683 | 3.82E-16 | 2.87768603 | 2.92E-14 |
| Fen1 | orange | 1.27854803 | 0.00049719 | 0.86828917 | 0.02585589 |
| Syt7 | red | -1.1400379 | 2.72E-06 | -0.7542262 | 0.00293358 |
| Aldh1a7 | skyblue | -1.6658607 | 3.15E-15 | -1.1035108 | 4.24E-07 |
| Lipo1 | red | -0.4512849 | 0.00053897 | -0.6216193 | 2.10E-06 |
| Otub1 | orange | 0.62933345 | 3.38E-05 | 0.45854627 | 0.00374937 |
| Cdc42bpg | red | -1.3837489 | 1.25E-06 | -0.9911521 | 0.00083405 |
| Fas | orange | 0.5932394 | 0.00018702 | 0.22512779 | 0.19937565 |
| Ak3 | red | -0.8340729 | 9.43E-06 | -0.8709673 | 5.61E-06 |
| 1700123I01R | yellow | 2.93033701 | 2.01E-13 | 4.43468522 | 1.46E-21 |
| Jak2 | orange | 1.12372335 | 7.58E-15 | 1.11660318 | 1.64E-14 |
| Cdca5 | yellow | 3.12290191 | 8.52E-11 | 3.6910428 | 1.15E-13 |
| Tnfrsf25 | red | -1.2419683 | 2.43E-07 | -1.0152967 | 3.89E-05 |
| Kif20b | yellow | 2.3627007 | 1.62E-21 | 2.65371112 | 2.18E-24 |
| Vps51 | orange | 0.62837891 | 0.00755766 | 0.54647106 | 0.02237566 |
| Htr7 | yellow | 3.70896994 | 3.94E-10 | 4.66110881 | 1.61E-13 |
| Rpp30 | orange | 0.77652043 | 7.36E-06 | 0.56448958 | 0.00192002 |
| Ankrd1 | orange | 1.69072301 | 2.05E-06 | 1.14622685 | 0.0020177 |
| Pcgf5 | orange | 0.82963144 | 5.27E-10 | 0.61020457 | 1.06E-05 |
| Mlana | skyblue | -0.7538376 | 0.25896592 | -1.8801001 | 0.0034814 |
| Gldc | orange | 1.88700775 | 5.11E-07 | 0.62909177 | 0.12193749 |
| Mrpl21 | orange | 0.60067319 | 0.0004677 | 0.40326909 | 0.02622545 |
| Pola2 | orange | 1.10036144 | 7.58E-13 | 1.01299945 | 8.02E-11 |

| | | | | | |
|---|---|---|---|---|---|
| Coro1b | orange | 0.73270152 | 9.03E-07 | 0.51073426 | 0.00100953 |
| Cabp4 | yellow | 3.7493299 | 1.35E-06 | 3.07923899 | 3.49E-05 |
| Banf1 | orange | 0.68882862 | 0.00063902 | 0.60581167 | 0.00370052 |
| Pitpnm1 | orange | 1.15528651 | 9.32E-10 | 0.90597429 | 3.07E-06 |
| Pold4 | orange | 0.65430776 | 3.12E-06 | 0.27774521 | 0.07396877 |
| Adrbk1 | orange | 0.87622 | 1.36E-14 | 0.75923778 | 5.35E-11 |
| Acy3 | red | -0.9346284 | 0.00179183 | -0.757743 | 0.01476843 |
| Doc2g | skyblue | -0.5351204 | 0.30567337 | -2.2496585 | 4.75E-06 |
| Rin1 | orange | 1.26026003 | 1.36E-06 | 1.53797504 | 4.37E-09 |
| Pcx | red | -1.42332 | 5.74E-07 | -1.001498 | 0.00070725 |
| Papss2 | red | -0.6633453 | 0.00183384 | -0.660987 | 0.00240146 |
| Peli3 | skyblue | -1.9384891 | 1.01E-08 | -1.5717952 | 5.46E-06 |
| Ctsw | yellow | 4.03752749 | 2.50E-44 | 3.44515101 | 2.03E-32 |
| Smarca2 | skyblue | -1.5118119 | 1.05E-13 | -1.3183743 | 1.87E-10 |
| Vldlr | red | -0.8068849 | 0.00076608 | -0.826993 | 0.00075888 |
| Slc1a1 | red | -1.0442042 | 9.82E-08 | -1.0765767 | 8.27E-08 |
| Ehbp1l1 | orange | 0.50683702 | 0.0021429 | 0.65972777 | 7.72E-05 |
| Ltbp3 | red | -1.1123346 | 2.03E-09 | -0.7612263 | 7.75E-05 |
| Prdx5 | orange | 1.07972266 | 3.67E-06 | 0.89550341 | 0.00020025 |
| Vegfb | red | -1.1428922 | 1.16E-08 | -1.0779036 | 1.39E-07 |
| Fermt3 | yellow | 2.54290387 | 4.32E-23 | 2.3665363 | 5.60E-20 |
| Rcor2 | red | -1.091668 | 0.00081389 | -0.4763488 | 0.17398597 |
| Lgals12 | skyblue | -2.7110847 | 4.77E-06 | -2.0236821 | 0.00086309 |
| Gpam | red | -0.7768091 | 8.00E-05 | -0.2227218 | 0.31560671 |
| Cep55 | yellow | 3.57569098 | 1.01E-35 | 4.03125298 | 2.75E-37 |
| Rbp4 | skyblue | -2.4187972 | 1.87E-10 | -1.5637385 | 6.25E-05 |
| Plce1 | red | -1.3314327 | 7.68E-08 | -0.901951 | 0.00047101 |
| Hells | yellow | 2.61528751 | 3.29E-18 | 2.28958646 | 1.17E-13 |
| Aldh18a1 | orange | 1.52991979 | 4.48E-12 | 1.51850351 | 1.01E-11 |
| Tll2 | skyblue | -4.4731844 | 7.02E-05 | -1.8192171 | 0.08294389 |
| Pik3ap1 | yellow | 2.66819662 | 8.49E-13 | 2.44972446 | 9.56E-11 |
| Lcor | orange | 0.601381 | 0.2162954 | 1.33579459 | 0.00527732 |
| Mxi1 | red | -1.0819374 | 7.11E-06 | -1.1158809 | 5.50E-06 |
| Add3 | red | -0.5373454 | 0.00055715 | -0.5916274 | 0.00019313 |
| Xpnpep1 | orange | 0.73443313 | 4.35E-17 | 0.54694108 | 9.56E-10 |
| Arl3 | red | -0.7866665 | 2.57E-08 | -0.7091046 | 1.05E-06 |
| Maoa | red | -1.2024507 | 1.28E-16 | -1.0538987 | 1.03E-12 |
| Nt5c2 | orange | 0.59838172 | 0.00015701 | 0.45676902 | 0.00551077 |
| Msr1 | black | 5.97567641 | 2.20E-18 | 5.37127985 | 4.41E-15 |
| 5430427O19 | yellow | 3.3208908 | 1.04E-14 | 2.8108081 | 8.11E-11 |
| Gk | orange | 1.00829138 | 0.00045621 | 0.80318305 | 0.0073869 |
| Col17a1 | red | -1.0930893 | 2.98E-05 | -1.1893583 | 8.37E-06 |
| Casp7 | orange | 0.77662225 | 2.11E-07 | 0.93984632 | 4.19E-10 |

| | | | | | |
|---|---|---|---|---|---|
| Nhlrc2 | red | -0.7952308 | 3.61E-11 | -0.6167805 | 5.62E-07 |
| Gfra1 | skyblue | -2.1612364 | 2.27E-10 | -1.1977205 | 0.00063307 |
| Hspa12a | red | -1.0648587 | 3.54E-09 | -0.7372873 | 7.36E-05 |
| Slc18a2 | red | -0.4417467 | 0.22666873 | -1.1783014 | 0.0009477 |
| Gcgr | skyblue | -2.1490186 | 0.00835587 | -0.7370154 | 0.37619671 |
| Arhgdia | orange | 0.62953722 | 0.00038611 | 0.55660702 | 0.00234005 |
| Alyref | orange | 1.01792532 | 5.86E-09 | 0.83480608 | 3.54E-06 |
| Lrrc45 | red | -0.8197553 | 6.53E-06 | -0.6202964 | 0.0009865 |
| Cbr2 | skyblue | -2.5228025 | 2.03E-06 | -1.6353263 | 0.00323872 |
| Fasn | skyblue | -1.4262264 | 0.00010436 | -1.4653612 | 9.56E-05 |
| Arhgap19 | orange | 1.44426206 | 1.48E-10 | 1.410987 | 7.94E-10 |
| Slc16a3 | orange | 0.86521259 | 0.00021681 | 1.01871642 | 1.92E-05 |
| Cd7 | yellow | 2.34135521 | 3.80E-19 | 2.39138387 | 4.10E-19 |
| Sectm1a | black | 7.18358726 | 3.53E-21 | 9.15129249 | 1.92E-12 |
| Rab40b | red | -1.3992077 | 4.30E-06 | -0.5096285 | 0.11413624 |
| Ubtd1 | orange | 0.68742434 | 0.00011914 | 0.76883499 | 2.35E-05 |
| Fn3k | skyblue | -3.4071157 | 1.96E-07 | -1.789384 | 0.00694634 |
| Hoga1 | orange | 0.70994838 | 6.66E-05 | 0.76481939 | 2.63E-05 |
| Loxl4 | orange | 0.7084516 | 0.01802563 | 1.07904809 | 0.00035416 |
| Hps1 | orange | 0.71541758 | 2.20E-05 | 0.70828191 | 4.18E-05 |
| Entpd7 | orange | 0.77428058 | 0.00039561 | 0.92042867 | 3.51E-05 |
| Dnmbp | red | -0.6014467 | 1.34E-07 | -0.3821874 | 0.00129302 |
| Cpn1 | skyblue | -1.9783985 | 4.15E-11 | -1.1212972 | 0.00026425 |
| Cyp2c44 | skyblue | -3.2919495 | 0.0003309 | -2.7214186 | 0.00098479 |
| Scd2 | red | -1.1739436 | 0.00142471 | -0.9173911 | 0.01672506 |
| Peo1 | red | -0.7454619 | 1.60E-09 | -0.4227087 | 0.00103874 |
| Kazald1 | skyblue | -1.6309947 | 2.21E-09 | -1.3064429 | 3.02E-06 |
| Nfkb2 | orange | 1.41191491 | 2.34E-12 | 1.35331674 | 3.11E-11 |
| Fam120c | red | -1.1607846 | 4.38E-07 | -0.6894299 | 0.00393374 |
| Apex2 | orange | 0.59942529 | 0.00019841 | 0.30297261 | 0.0799365 |
| Tro | skyblue | -1.8752689 | 2.06E-07 | -1.5976968 | 1.17E-05 |
| Flnb | orange | 1.22214686 | 1.17E-06 | 1.19900984 | 2.97E-06 |
| Dnase1l3 | yellow | 2.87936978 | 3.25E-11 | 2.35871683 | 1.05E-07 |
| Sat1 | orange | 0.95973647 | 8.98E-11 | 0.6874253 | 7.07E-06 |
| Acot9 | orange | 0.77127194 | 3.99E-12 | 0.69747733 | 7.60E-10 |
| Ptprj | orange | 0.70280023 | 0.00017168 | 0.65835543 | 0.00062591 |
| Banp | red | -0.7788183 | 2.68E-05 | -0.6849594 | 0.00034896 |
| Atp10a | orange | 1.1928809 | 3.85E-11 | 1.04842777 | 1.05E-08 |
| Padi4 | orange | 0.61589925 | 0.09021588 | 1.35512361 | 0.00014287 |
| Rdh5 | red | -1.1638652 | 4.35E-05 | -0.8483288 | 0.00397459 |
| Cd63 | orange | 0.84591815 | 0.00056594 | 0.83047069 | 0.00097026 |
| Mmp19 | orange | 1.09102352 | 0.00171323 | 0.81433944 | 0.02526513 |
| Rps26 | orange | 0.70987342 | 5.49E-13 | 0.589841 | 4.64E-09 |

| | | | | | |
|---|---|---|---|---|---|
| Pa2g4 | orange | 0.69285205 | 1.00E-05 | 0.55262949 | 0.00066726 |
| Esyt1 | orange | 1.03603606 | 5.61E-08 | 0.84515205 | 1.70E-05 |
| Nabp2 | orange | 0.594568 | 0.00019107 | 0.3367656 | 0.0479603 |
| Aatk | red | -0.6393457 | 1.45E-05 | -0.3251335 | 0.03899809 |
| Il23a | yellow | 3.42116677 | 0.00024605 | 3.27442352 | 0.00021492 |
| Prim1 | orange | 1.83082079 | 1.40E-21 | 1.2465045 | 2.41E-10 |
| Nab2 | orange | 0.64895266 | 0.00010816 | 0.69213928 | 5.48E-05 |
| Shmt2 | orange | 0.89362259 | 2.99E-11 | 0.75718596 | 3.87E-08 |
| Gli1 | red | -1.3733257 | 3.65E-06 | -0.7269699 | 0.02015234 |
| Katnal2 | red | -1.0194119 | 0.00348474 | -0.3068408 | 0.42944803 |
| Agap2 | yellow | 2.85731659 | 1.06E-32 | 2.73764491 | 7.81E-30 |
| Pstpip2 | orange | 0.70073322 | 0.02927222 | 1.13786771 | 0.00037839 |
| Crisp1 | red | -1.5971823 | 0.00144872 | -0.6461752 | 0.23432108 |
| Avil | red | -0.6594388 | 0.00695188 | -0.8049385 | 0.00120855 |
| Paox | orange | 0.60443634 | 0.00053149 | 0.42528512 | 0.02056676 |
| Adam8 | yellow | 2.3305673 | 5.14E-06 | 1.89193498 | 0.00034151 |
| Inpp5a | red | -0.6114923 | 4.52E-06 | -0.6159124 | 6.02E-06 |
| Psmd13 | orange | 0.83201192 | 6.00E-11 | 0.60481883 | 4.31E-06 |
| Cox8b | skyblue | -2.1427324 | 2.08E-13 | -1.907707 | 2.52E-10 |
| Ifitm5 | orange | 0.95566809 | 0.00920152 | 0.22954166 | 0.59052005 |
| Ifitm1 | orange | 1.26307875 | 5.20E-12 | 0.84267689 | 9.07E-06 |
| Ifitm3 | yellow | 2.14252192 | 1.95E-35 | 2.0909679 | 9.97E-34 |
| Drd4 | yellow | 3.60831636 | 0.00011975 | 4.47426418 | 0.00011496 |
| Irf7 | black | 5.32734858 | 1.88E-98 | 5.0690693 | 3.24E-89 |
| Pidd1 | yellow | 1.90822049 | 1.35E-17 | 2.14293092 | 1.29E-21 |
| Rplp2 | orange | 0.66753382 | 1.56E-07 | 0.54082617 | 3.81E-05 |
| Tmem192 | orange | 1.18121756 | 1.05E-07 | 0.86436872 | 0.0001939 |
| Asl | orange | 0.60946349 | 4.91E-06 | 0.51721842 | 0.00018073 |
| Gusb | orange | 0.73472942 | 0.00056215 | 0.44467273 | 0.04975381 |
| Phkg1 | skyblue | -2.0219738 | 6.78E-05 | -2.1820177 | 2.60E-05 |
| Clybl | red | -0.8667321 | 6.53E-09 | -0.756746 | 1.34E-06 |
| Tk1 | yellow | 2.09514147 | 3.80E-25 | 1.95139297 | 1.34E-21 |
| Rbfox3 | red | -0.2449971 | 0.60625381 | -1.3331037 | 0.00181078 |
| Cbx2 | red | -0.7954643 | 0.0002847 | -0.8870476 | 6.01E-05 |
| Cbx8 | red | -0.7261558 | 0.00011203 | -0.7255707 | 0.00020023 |
| Gaa | red | -0.7458441 | 1.83E-08 | -0.5988407 | 1.16E-05 |
| Nptx1 | skyblue | -3.5238886 | 5.17E-49 | -2.4432773 | 5.47E-25 |
| Cpeb1 | red | -0.6162721 | 0.00021437 | -0.6029109 | 0.00039165 |
| Tma16 | orange | 0.80087167 | 0.00194604 | 0.96128621 | 0.00026009 |
| Map3k7cl | skyblue | -1.679039 | 2.68E-10 | -0.8148578 | 0.00326568 |
| Hprt | orange | 0.6874734 | 3.55E-05 | 0.41084077 | 0.01958429 |
| Shisa5 | orange | 1.26431729 | 1.96E-18 | 1.17243288 | 7.89E-16 |
| Col7a1 | red | -0.9970846 | 5.35E-05 | -0.4891725 | 0.06525863 |

| | | | | | |
|---|---|---|---|---|---|
| Arhgef9 | red | -1.2471431 | 2.04E-06 | -0.9149122 | 0.00064337 |
| Cnksr2 | skyblue | -1.7830518 | 0.00363062 | -0.6836736 | 0.27951202 |
| Rps6ka6 | skyblue | -0.9489117 | 0.08815324 | -1.8264137 | 0.00119507 |
| Tmem47 | red | -1.0551702 | 5.72E-09 | -0.7310726 | 1.00E-04 |
| March8 | red | -0.6297631 | 3.52E-11 | -0.4414352 | 7.30E-06 |
| 0610011F06I | red | -0.8214444 | 1.63E-05 | -0.6426324 | 0.00119278 |
| Fam195a | red | -0.799202 | 9.38E-05 | -0.9558348 | 7.35E-06 |
| Jmjd8 | red | -1.1181285 | 4.48E-11 | -0.7830238 | 7.71E-06 |
| Fbxl16 | red | -1.1695655 | 1.57E-05 | -0.6698622 | 0.01681535 |
| Prps2 | red | -0.6391837 | 0.0031096 | -0.7225972 | 0.00103315 |
| Sdc3 | orange | 0.78731901 | 0.00030159 | 0.60774026 | 0.0072998 |
| Il6 | black | 5.14651871 | 5.87E-15 | 7.85106891 | 2.46E-16 |
| Tyms | orange | 1.56543491 | 7.73E-11 | 1.45618832 | 3.32E-09 |
| Hspa4l | red | -1.4521266 | 5.24E-15 | -1.0097167 | 1.07E-07 |
| Plk4 | orange | 1.28490498 | 1.64E-07 | 1.54829093 | 7.64E-10 |
| Jade1 | red | -0.862817 | 1.12E-06 | -0.7433475 | 4.56E-05 |
| D3Ertd751e | red | -1.1147924 | 5.69E-05 | -1.1023993 | 0.00010058 |
| Itih5 | red | -1.1453355 | 0.00012131 | -0.7922228 | 0.01101929 |
| Clec3b | skyblue | -2.3834738 | 6.87E-19 | -1.5484355 | 2.32E-08 |
| St8sia2 | skyblue | -1.9866759 | 1.88E-06 | -1.2980551 | 0.00271131 |
| Slc25a10 | red | -0.6219736 | 4.38E-05 | -0.5132599 | 0.00110065 |
| Rassf3 | red | -1.2226318 | 8.94E-12 | -0.9274823 | 4.95E-07 |
| Ccr1 | yellow | 4.05551869 | 1.85E-24 | 4.26585541 | 9.57E-27 |
| Nrp1 | red | -1.0701272 | 6.34E-11 | -0.8670343 | 2.44E-07 |
| Homer2 | red | -1.48746 | 7.56E-07 | -0.7071711 | 0.02868853 |
| Dhtkd1 | skyblue | -1.5779654 | 0.00385837 | -1.1951891 | 0.03305597 |
| Pdgfa | red | -1.0624068 | 5.04E-11 | -0.8734262 | 1.49E-07 |
| Cplx2 | red | -0.8641203 | 2.39E-06 | -0.5068064 | 0.00821226 |
| Nop16 | orange | 0.56861196 | 0.00257934 | 0.63641983 | 0.0009939 |
| Tspan17 | red | -0.6521435 | 1.69E-08 | -0.6123627 | 2.36E-07 |
| Hk3 | black | 5.29987359 | 7.24E-52 | 5.39745103 | 2.78E-53 |
| Smad7 | red | -0.6530414 | 1.06E-08 | -0.5676283 | 1.15E-06 |
| Myo5b | red | -0.7592208 | 0.00322209 | -0.3897799 | 0.16242991 |
| Casp12 | orange | 0.79178514 | 1.18E-10 | 0.8218485 | 2.91E-11 |
| Casp1 | yellow | 2.30347932 | 2.44E-15 | 2.21885419 | 4.93E-14 |
| Kbtbd3 | red | -1.3994513 | 1.60E-05 | -0.7309871 | 0.03616911 |
| Rp1 | skyblue | -2.9528495 | 2.43E-12 | -2.0158346 | 1.97E-06 |
| Sox17 | red | -0.7080303 | 3.06E-07 | -0.7976252 | 1.20E-08 |
| Adhfe1 | red | -0.9686714 | 0.00031801 | -0.8222181 | 0.00304561 |
| Ppp1r42 | skyblue | -2.3096684 | 0.00068642 | -1.8662993 | 0.0062426 |
| Stau2 | red | -0.8438966 | 1.37E-05 | -0.7474462 | 0.00018159 |
| Rdh10 | red | -0.8101501 | 3.51E-11 | -0.3247939 | 0.01248189 |
| Paqr8 | red | -0.7795479 | 1.85E-07 | -0.3082373 | 0.05795071 |

| | | | | | |
|---|---|---|---|---|---|
| Eya1 | skyblue | -1.5447085 | 5.63E-08 | -0.8604119 | 0.00508614 |
| Tmem14a | skyblue | -1.4544303 | 3.57E-08 | -1.5694594 | 6.85E-09 |
| Gsta3 | skyblue | -1.4814738 | 0.00031136 | -1.2333831 | 0.00369034 |
| Gm4956 | skyblue | -1.6453443 | 0.00014152 | -1.4244167 | 0.00138858 |
| Slco5a1 | skyblue | -1.5937219 | 5.71E-08 | -1.3020011 | 3.78E-05 |
| Idh1 | red | -0.8740345 | 1.07E-09 | -0.5524938 | 0.00021152 |
| Mdh1b | skyblue | -2.4752431 | 6.89E-08 | -1.3600507 | 0.00348952 |
| Eef1b2 | orange | 0.65066685 | 1.01E-06 | 0.44351876 | 0.00139985 |
| Rftn2 | red | -1.3069251 | 3.21E-05 | -1.0994337 | 0.00068976 |
| Wdr75 | orange | 0.58218544 | 0.00011412 | 0.38551147 | 0.01522452 |
| Ikzf2 | orange | 1.31250301 | 4.42E-07 | 0.91760926 | 0.00066411 |
| Lancl1 | red | -0.7014246 | 4.45E-07 | -0.4449002 | 0.00219887 |
| Icos | black | 5.61394466 | 3.47E-39 | 5.12221125 | 1.76E-34 |
| Ctla4 | black | 7.18181704 | 2.45E-52 | 6.78987048 | 6.07E-53 |
| Cd28 | yellow | 3.57918367 | 2.72E-10 | 3.11495247 | 4.49E-08 |
| Wdr12 | orange | 0.6754628 | 0.00040887 | 0.7057073 | 0.00033135 |
| Nop58 | orange | 1.32105864 | 2.02E-11 | 1.10500007 | 4.69E-08 |
| Stradb | red | -0.7130409 | 0.00486065 | -0.4564838 | 0.09036791 |
| Casp8 | orange | 1.48927519 | 3.39E-14 | 1.28301867 | 1.42E-10 |
| Cflar | orange | 0.62621064 | 8.11E-08 | 0.56344349 | 2.43E-06 |
| Sgol2a | yellow | 2.39210644 | 4.52E-10 | 2.6267161 | 4.97E-10 |
| Col5a2 | orange | 0.72082847 | 0.02470606 | 0.8735143 | 0.00715021 |
| Col3a1 | orange | 0.61430051 | 0.06933169 | 1.06641572 | 0.00139667 |
| Tex30 | orange | 1.23626026 | 2.11E-09 | 0.91914024 | 3.00E-05 |
| Slc9a2 | skyblue | -1.708381 | 5.45E-05 | -0.9113445 | 0.03917126 |
| Il18rap | yellow | 3.36987945 | 1.38E-42 | 3.37395558 | 2.36E-42 |
| Il1rl1 | orange | 0.85765175 | 4.03E-06 | 0.81144459 | 2.63E-05 |
| Il18r1 | orange | 0.96040245 | 1.10E-07 | 1.03732171 | 1.45E-08 |
| Il1r2 | yellow | 2.29171409 | 1.89E-08 | 2.55842546 | 5.67E-10 |
| Chst10 | orange | 1.06545343 | 0.00583843 | 1.79548898 | 5.43E-06 |
| Mrpl30 | orange | 0.63161319 | 0.00016032 | 0.52298236 | 0.00264051 |
| Mitd1 | orange | 1.90508092 | 7.05E-30 | 1.6136289 | 1.60E-21 |
| 2010300C02 | red | -1.1897817 | 5.28E-09 | -0.7852273 | 0.00020085 |
| Stk17b | orange | 1.56459702 | 4.47E-20 | 1.34954037 | 6.57E-15 |
| Pms1 | orange | 0.96143389 | 0.00583216 | 0.94650754 | 0.0083065 |
| Inpp1 | orange | 1.00426287 | 1.06E-22 | 0.88720263 | 8.68E-18 |
| Stat1 | yellow | 2.32873497 | 4.33E-39 | 2.40935317 | 1.12E-41 |
| Nabp1 | orange | 0.81749064 | 1.68E-10 | 0.80881323 | 5.40E-10 |
| Tmeff2 | skyblue | -2.0440867 | 1.39E-18 | -1.5916234 | 2.65E-11 |
| Mgat4a | orange | 1.09831693 | 7.51E-10 | 0.88277356 | 1.44E-06 |
| Zap70 | yellow | 3.02392723 | 2.17E-66 | 2.80664237 | 6.30E-57 |
| Sema4c | orange | 0.89498818 | 5.50E-07 | 0.83704798 | 4.61E-06 |
| Ptpn18 | orange | 0.66048375 | 1.06E-05 | 0.52588527 | 0.00078284 |

| | | | | | |
|---|---|---|---|---|---|
| Prim2 | orange | 1.38020337 | 1.61E-15 | 1.12869962 | 2.45E-10 |
| Ogfrl1 | orange | 0.937649 | 3.48E-06 | 0.52634882 | 0.01399719 |
| Ccl20 | black | 6.18444999 | 1.90E-11 | 5.87201685 | 4.83E-10 |
| Cyp27a1 | red | -0.7402943 | 0.00181734 | -0.6706964 | 0.00620267 |
| Plcd4 | red | -1.4808337 | 3.92E-06 | -0.7035208 | 0.04665444 |
| Rqcd1 | orange | 0.76789376 | 2.43E-06 | 0.56794296 | 0.00079801 |
| Slc11a1 | yellow | 3.29930458 | 4.90E-10 | 3.16175883 | 4.26E-09 |
| Pnkd | red | -1.1881747 | 7.11E-07 | -1.1129667 | 5.77E-06 |
| Cxcr2 | yellow | 2.61261386 | 5.74E-12 | 2.29293377 | 1.22E-09 |
| Igfbp5 | skyblue | -1.5133566 | 1.11E-17 | -1.2320793 | 7.55E-12 |
| Bard1 | yellow | 2.56381041 | 1.37E-11 | 2.21818638 | 9.05E-09 |
| Ankzf1 | red | -0.4909014 | 0.00248536 | -0.6278775 | 0.00012892 |
| Dnajb2 | red | -0.7936104 | 4.87E-09 | -0.6002222 | 1.87E-05 |
| Speg | red | -0.9743217 | 4.49E-09 | -0.7520702 | 1.18E-05 |
| Des | red | -0.934743 | 4.66E-06 | -0.5387731 | 0.01238949 |
| Obsl1 | red | -0.699788 | 0.00043913 | -0.6831504 | 0.0008265 |
| Sp100 | orange | 1.87182585 | 4.11E-79 | 1.81315972 | 5.16E-74 |
| 2810459M11 | skyblue | -2.8067755 | 0.00609757 | -1.7946641 | 0.0786753 |
| Ncl | orange | 1.00168203 | 1.35E-09 | 0.80875274 | 1.92E-06 |
| Nmur1 | skyblue | -1.9661086 | 0.00014626 | -1.9820088 | 0.00026723 |
| Eif4e2 | orange | 0.60161175 | 4.66E-05 | 0.51685738 | 0.0007177 |
| Ngef | skyblue | -1.7421997 | 1.86E-11 | -1.067346 | 7.20E-05 |
| Gpr35 | yellow | 3.11133302 | 2.24E-21 | 2.97017473 | 1.88E-19 |
| Pask | orange | 1.58216173 | 2.36E-06 | 1.4561459 | 2.04E-05 |
| Bok | red | -0.6311034 | 0.00222414 | -0.5560771 | 0.00901121 |
| Dtymk | orange | 0.58256219 | 6.38E-05 | 0.37367637 | 0.01534389 |
| Pdcd1 | black | 7.7919514 | 1.81E-66 | 6.52463904 | 6.92E-55 |
| Inpp5d | orange | 1.20629827 | 7.40E-14 | 0.93195177 | 1.73E-08 |
| Iqca | skyblue | -2.8365373 | 1.00E-07 | -2.0325614 | 0.00019311 |
| Rab17 | skyblue | -1.6395037 | 4.33E-05 | -1.5109591 | 0.00039259 |
| Lrrfip1 | orange | 0.53755422 | 0.00113006 | 0.65540125 | 9.08E-05 |
| Scly | orange | 0.80837273 | 1.05E-13 | 0.62407858 | 2.23E-08 |
| Klhl30 | skyblue | -1.3865203 | 6.85E-06 | -1.3796003 | 2.40E-05 |
| Tnfrsf11a | yellow | 2.24665001 | 1.54E-07 | 1.86091378 | 2.43E-05 |
| Pam | skyblue | -1.2943523 | 0.00055278 | -1.6165926 | 2.03E-05 |
| Actr3 | orange | 1.43961013 | 5.61E-15 | 1.23519889 | 4.36E-11 |
| Tmem163 | red | -0.7483763 | 0.01360208 | -1.0095418 | 0.0009654 |
| Mcm6 | orange | 1.62826153 | 4.80E-27 | 1.37812571 | 1.85E-19 |
| Rgs1 | black | 5.30560631 | 5.21E-28 | 4.17088832 | 1.77E-19 |
| Nifk | orange | 0.7352035 | 6.65E-05 | 0.68850407 | 0.00029907 |
| Epb41l5 | red | -1.0479853 | 3.77E-09 | -0.6875393 | 0.00020551 |
| Ptpn4 | red | -0.7479781 | 0.000161 | -0.7179821 | 0.000474 |
| Steap3 | skyblue | -1.5504105 | 5.73E-32 | -1.3743694 | 7.05E-25 |

| | | | | | |
|---|---|---|---|---|---|
| Ptprc | yellow | 2.47692632 | 6.67E-54 | 2.50500815 | 7.15E-55 |
| Cd55 | red | -1.4643144 | 4.59E-17 | -1.0172232 | 1.29E-08 |
| Pigr | orange | -0.3751016 | 0.10855447 | 0.70480807 | 0.00194957 |
| Il24 | black | 6.76803021 | 6.33E-05 | 5.75051364 | 0.00102342 |
| Gpr37l1 | skyblue | -2.0504378 | 7.25E-05 | -1.225091 | 0.0267181 |
| Ube2t | orange | 1.6148251 | 1.91E-10 | 1.72415428 | 5.78E-11 |
| Rassf5 | orange | 1.21680704 | 6.17E-13 | 1.09902311 | 1.62E-10 |
| Rab29 | orange | 0.79579542 | 5.85E-08 | 0.58957981 | 0.00012202 |
| Lrrn2 | red | -1.2974065 | 0.00030491 | -0.9251776 | 0.01483816 |
| Pik3c2b | red | -0.9462238 | 8.24E-11 | -0.5043921 | 0.00100761 |
| Ppfia4 | orange | 2.0443587 | 2.81E-09 | 1.6645321 | 2.75E-06 |
| Myog | skyblue | -2.8510376 | 8.55E-05 | -2.4386996 | 0.0032624 |
| Tor1aip1 | orange | 0.86473268 | 3.23E-14 | 0.7023157 | 1.67E-09 |
| Mr1 | red | -0.5994919 | 7.97E-06 | -0.4954256 | 0.00031733 |
| Rgs16 | yellow | 3.93351057 | 8.99E-38 | 4.03773943 | 5.76E-38 |
| Ncf2 | yellow | 2.09406691 | 2.79E-13 | 1.81508386 | 4.97E-10 |
| Adck3 | red | -1.3410772 | 1.40E-06 | -1.1379529 | 7.32E-05 |
| Cdc42bpa | red | -0.8424203 | 2.22E-06 | -0.71604 | 9.31E-05 |
| Tfb2m | red | -0.7249191 | 4.77E-07 | -0.7570916 | 2.64E-07 |
| Kif26b | skyblue | -1.9791958 | 1.25E-12 | -1.6856321 | 3.17E-09 |
| Efcab2 | red | -1.1696101 | 1.83E-05 | -1.1882314 | 2.19E-05 |
| Trp53bp2 | red | -1.0796359 | 1.14E-09 | -0.8245944 | 6.53E-06 |
| Srp9 | orange | 0.64930833 | 1.06E-07 | 0.41067823 | 0.00136079 |
| Rgs7 | skyblue | -3.0217304 | 4.48E-05 | -2.6577289 | 0.0008621 |
| Spta1 | skyblue | -0.9076004 | 0.1194254 | -2.4195927 | 0.00015851 |
| Mnda | black | 5.61382305 | 1.03E-45 | 5.2648848 | 3.65E-40 |
| Cfap45 | red | -0.7714071 | 3.93E-05 | -0.3100802 | 0.12846313 |
| Slamf9 | yellow | 3.41767715 | 2.69E-09 | 2.83731484 | 1.51E-06 |
| Vangl2 | red | -0.8227055 | 2.18E-06 | -0.6095122 | 0.0007329 |
| Uck2 | orange | 0.71033526 | 0.00018333 | 0.78335622 | 5.18E-05 |
| Dusp27 | skyblue | -1.3769567 | 1.46E-08 | -1.1497931 | 1.10E-05 |
| Mpc2 | red | -0.5466377 | 0.00012744 | -0.593804 | 5.02E-05 |
| Xcl1 | black | 4.55110773 | 1.64E-15 | 4.5739231 | 3.33E-12 |
| Dpt | red | -1.1303013 | 7.50E-12 | -0.8628518 | 3.83E-07 |
| Atp1b1 | red | -0.6066488 | 0.00045799 | -0.2456016 | 0.19697203 |
| Ccdc181 | red | -0.6925333 | 0.00014273 | -0.5899278 | 0.0016643 |
| Selp | yellow | 2.5947715 | 1.46E-22 | 3.02212729 | 6.23E-30 |
| Sell | yellow | 2.10996344 | 1.17E-38 | 2.37623214 | 2.86E-48 |
| Sele | orange | 1.99826512 | 0.00055678 | 1.74481284 | 0.00288814 |
| Scyl3 | orange | 0.66343882 | 5.80E-06 | 0.57122406 | 0.00013499 |
| Kifap3 | red | -0.9293088 | 3.14E-08 | -0.9079071 | 1.14E-07 |
| Prrx1 | red | -0.8908372 | 2.52E-07 | -0.5729234 | 0.00159921 |
| Sec16b | red | -0.7778467 | 1.15E-05 | -0.6234548 | 0.00075317 |

| | | | | | |
|---|---|---|---|---|---|
| Abl2 | orange | 1.08313838 | 4.46E-08 | 0.87915717 | 1.65E-05 |
| Cenpf | yellow | 2.67631248 | 1.21E-20 | 3.28665498 | 8.60E-28 |
| Esrrg | skyblue | -2.0118556 | 4.19E-05 | -2.3243479 | 4.17E-06 |
| Spata17 | skyblue | -1.48003 | 0.00011257 | -1.1898875 | 0.00198098 |
| Cr2 | skyblue | -1.7924005 | 0.00333565 | -1.5642635 | 0.01605409 |
| Nek2 | orange | 1.27255046 | 1.54E-05 | 1.38369751 | 3.61E-06 |
| Tmem206 | red | -0.5683534 | 0.00063448 | -0.697171 | 5.26E-05 |
| Atf3 | yellow | 3.25820199 | 4.42E-85 | 2.4898596 | 5.38E-52 |
| Batf3 | yellow | 2.82267153 | 4.57E-22 | 2.71445074 | 1.08E-19 |
| Irf6 | red | -0.8940074 | 8.39E-06 | -0.5100234 | 0.01611685 |
| Plxna2 | orange | 0.79759206 | 3.26E-07 | 0.58503095 | 0.00029611 |
| Suv39h2 | orange | 1.51904189 | 0.00018444 | 1.32128585 | 0.00157488 |
| Dclre1c | orange | 1.42770999 | 7.02E-15 | 1.18064013 | 2.24E-10 |
| Cfap126 | skyblue | -1.6986369 | 1.02E-13 | -1.1015356 | 2.55E-06 |
| Meig1 | skyblue | -2.1030198 | 9.95E-11 | -0.9914455 | 0.00097495 |
| Fam107b | orange | 0.84026107 | 1.89E-09 | 0.87782756 | 5.76E-10 |
| Fcgr2b | yellow | 3.16646103 | 8.33E-27 | 3.10827507 | 1.00E-25 |
| Frmd4a | red | -0.4803047 | 0.00081305 | -0.5893313 | 5.02E-05 |
| Mcm10 | yellow | 2.93317826 | 5.53E-25 | 3.36039899 | 8.65E-30 |
| Ddr2 | red | -0.7820304 | 5.22E-14 | -0.4652647 | 1.81E-05 |
| Hsd17b7 | orange | 0.96374508 | 0.00066086 | 0.9528712 | 0.00102349 |
| Ccdc3 | skyblue | -2.6552639 | 5.31E-09 | -1.4028946 | 0.00163211 |
| Rgs5 | skyblue | -1.5936439 | 1.73E-08 | -1.5401776 | 9.84E-08 |
| Enkur | skyblue | -1.4273852 | 6.93E-06 | -1.507766 | 3.31E-06 |
| Nuf2 | yellow | 3.26258501 | 8.73E-25 | 3.60072001 | 6.45E-27 |
| Fmo3 | skyblue | -1.8234105 | 1.93E-05 | -0.9065679 | 0.04674546 |
| Myoc | skyblue | -1.8385897 | 0.00017216 | -2.3781194 | 2.33E-06 |
| Prdx6 | red | -0.6417354 | 0.00468063 | -0.5350455 | 0.02323841 |
| Cenpl | orange | 1.77346022 | 2.91E-22 | 1.41048272 | 2.63E-14 |
| Tnn | skyblue | -2.3701346 | 1.22E-12 | -1.5356453 | 5.12E-06 |
| 4930562F07I | yellow | 2.37306878 | 0.00378877 | 1.60326307 | 0.01854755 |
| Pip4k2a | orange | 0.91649844 | 9.88E-08 | 0.74814316 | 2.57E-05 |
| Plxdc2 | skyblue | -1.3723887 | 5.29E-22 | -1.1530881 | 1.19E-15 |
| Nek6 | orange | 1.2222891 | 8.16E-07 | 1.30536162 | 2.11E-07 |
| Psmb7 | orange | 0.78743192 | 4.58E-05 | 0.68775242 | 0.00056215 |
| Lypd6b | red | -0.8744081 | 0.00127437 | -0.5043473 | 0.07734984 |
| Fam188a | orange | 0.61254491 | 7.05E-08 | 0.54107747 | 3.67E-06 |
| Itga8 | skyblue | -2.1339515 | 6.22E-09 | -1.4730836 | 0.00011115 |
| Il2ra | yellow | 4.36457139 | 2.42E-64 | 4.21532681 | 3.17E-59 |
| Pfkfb3 | orange | 1.30389468 | 4.25E-18 | 1.13467515 | 9.17E-14 |
| Prkcq | orange | 0.97800393 | 7.58E-08 | 1.08821953 | 3.56E-09 |
| Mastl | yellow | 2.52553286 | 1.45E-07 | 2.77153768 | 4.65E-08 |
| Pdss1 | orange | 0.92606447 | 0.00058386 | 0.93493246 | 0.00075317 |

| | | | | | |
|---|---|---|---|---|---|
| Pkn3 | red | -0.6562704 | 4.46E-05 | -0.4671242 | 0.00540602 |
| Apbb1ip | orange | 2.00650728 | 6.29E-17 | 1.78911152 | 1.90E-13 |
| Lrsam1 | red | -0.7922173 | 4.95E-09 | -0.541182 | 0.00011694 |
| Ddx31 | orange | 0.67707951 | 8.11E-06 | 0.49288588 | 0.00189613 |
| Ak8 | skyblue | -1.9232243 | 1.15E-06 | -1.1312878 | 0.00507573 |
| 1700026L06F | skyblue | -1.7416928 | 3.39E-14 | -1.3180951 | 8.28E-09 |
| St6galnac6 | red | -0.7694265 | 4.19E-11 | -0.7654926 | 1.16E-10 |
| Eng | red | -0.6012699 | 3.09E-07 | -0.5280559 | 1.19E-05 |
| Ak1 | red | -1.319115 | 7.83E-17 | -1.0103786 | 5.31E-10 |
| Slc25a25 | orange | 1.00549229 | 2.26E-06 | 0.96706453 | 8.64E-06 |
| Lcn2 | yellow | 3.65905501 | 3.15E-68 | 3.17019467 | 2.53E-51 |
| Dnm1 | red | -0.8283729 | 2.69E-06 | -0.7941341 | 1.16E-05 |
| Galnt5 | skyblue | -2.1059823 | 7.44E-05 | -0.7972734 | 0.14721002 |
| Ermn | black | 6.04321859 | 2.37E-05 | 5.48439769 | 0.00019125 |
| 1700007K13I | skyblue | -2.1294201 | 5.03E-13 | -1.5042772 | 6.57E-07 |
| Cytip | orange | 1.50269343 | 4.21E-32 | 1.23566423 | 7.48E-22 |
| Acvr1c | skyblue | -2.5943883 | 0.00278352 | -1.0951938 | 0.23505792 |
| Upp2 | skyblue | -1.6522781 | 4.48E-09 | -1.4127576 | 9.84E-07 |
| Crat | red | -0.8971858 | 3.48E-10 | -0.7247893 | 8.08E-07 |
| Ntmt1 | orange | 0.53487184 | 0.00571894 | 0.59761332 | 0.00251806 |
| Fam73b | red | -0.8011165 | 3.22E-06 | -0.5282909 | 0.00328117 |
| Kynu | orange | 1.7366438 | 2.34E-05 | 1.39107696 | 0.00085839 |
| Cutal | skyblue | -2.1402377 | 3.39E-11 | -1.4514886 | 1.18E-05 |
| Phf19 | orange | 0.7553749 | 3.39E-05 | 0.7730224 | 3.87E-05 |
| Hc | skyblue | -2.2697397 | 0.00798853 | -1.3212073 | 0.15046723 |
| Gsn | red | -1.2529339 | 1.64E-05 | -0.9521612 | 0.00159685 |
| Dab2ip | red | -0.962565 | 3.20E-08 | -0.5963462 | 0.00103923 |
| Ttll11 | orange | 0.30604702 | 0.31207267 | 0.99126469 | 0.00059813 |
| Grb14 | red | -1.1247838 | 4.01E-05 | -0.5459841 | 0.06367628 |
| Lhx6 | orange | 1.04318169 | 4.82E-08 | 1.32808082 | 1.45E-11 |
| Ifih1 | yellow | 1.99664093 | 1.82E-20 | 1.9962778 | 2.31E-20 |
| Slc4a10 | skyblue | -3.7939623 | 0.00274288 | -0.7818859 | 0.50811705 |
| Psmd14 | orange | 1.1803096 | 2.02E-10 | 0.9357642 | 9.87E-07 |
| Strbp | red | -0.7251425 | 1.09E-07 | -0.5164226 | 0.00024731 |
| Brd3 | red | -0.61983 | 0.00010605 | -0.5327709 | 0.00125079 |
| Lcn4 | yellow | 3.19963517 | 1.62E-05 | 3.24945915 | 0.00021638 |
| Agpat2 | red | -0.6571265 | 0.00180635 | -0.560407 | 0.01010329 |
| Card9 | orange | 0.94906736 | 4.08E-08 | 0.87953301 | 5.97E-07 |
| Gpsm1 | red | -0.6533491 | 0.00180313 | -0.6761782 | 0.00155851 |
| Tmem141 | red | -0.6611896 | 0.00571717 | -0.4378471 | 0.08544177 |
| Traf2 | orange | 0.66119645 | 1.17E-11 | 0.61248697 | 7.67E-10 |
| Nmi | orange | 2.02184037 | 1.19E-36 | 1.8377808 | 2.97E-30 |
| Neb | red | -0.9746073 | 0.0032317 | -0.5396036 | 0.13032403 |

| | | | | | |
|---|---|---|---|---|---|
| Sapcd2 | yellow | 2.25668346 | 3.02E-12 | 2.7296409 | 2.24E-16 |
| Lrrc26 | red | -1.2517954 | 4.81E-07 | -0.6653385 | 0.01115202 |
| Rbms1 | orange | 0.64349473 | 5.28E-06 | 0.48075977 | 0.00105791 |
| Itgb6 | orange | 1.41414117 | 2.65E-06 | 1.62516963 | 9.97E-08 |
| Zmynd19 | orange | 0.76830484 | 8.10E-05 | 0.84256737 | 2.31E-05 |
| Psd4 | yellow | 2.21100081 | 9.08E-27 | 2.05239577 | 4.69E-23 |
| Ly75 | orange | 1.21082266 | 4.52E-10 | 1.42947335 | 3.08E-13 |
| Il1rn | black | 4.5362517 | 7.66E-30 | 4.53418203 | 1.50E-29 |
| Il1f5 | red | -1.1819998 | 0.00944699 | -0.6847258 | 0.15911466 |
| Il1f6 | skyblue | -1.8035572 | 9.22E-05 | -0.9170043 | 0.06135728 |
| Hnmt | red | -1.0117047 | 0.00089167 | -1.037336 | 0.00096017 |
| Wdsub1 | red | -0.9330898 | 3.56E-08 | -0.7174835 | 3.98E-05 |
| Dusp19 | red | -0.5893647 | 0.00066148 | -0.6054286 | 0.00079041 |
| Frzb | skyblue | -1.595957 | 0.00031248 | -0.8753056 | 0.06344341 |
| Itga4 | yellow | 1.94476345 | 7.25E-26 | 2.04887702 | 2.82E-28 |
| Cybrd1 | skyblue | -2.1142513 | 2.77E-12 | -2.1117183 | 4.82E-12 |
| Zfp385b | red | -1.3248345 | 0.00474208 | -1.0892627 | 0.02575576 |
| Hat1 | orange | 1.41957337 | 4.96E-10 | 1.06754692 | 6.07E-06 |
| Xirp2 | skyblue | -1.8908941 | 9.77E-07 | -1.3314879 | 0.00093952 |
| Stk39 | orange | 1.13856633 | 2.31E-07 | 0.7021307 | 0.00234654 |
| Cers6 | orange | 1.24843189 | 1.30E-10 | 1.37977899 | 1.44E-12 |
| Timm10 | orange | 0.71075594 | 0.00363769 | 0.69787777 | 0.00586624 |
| Smtnl1 | red | -1.2041125 | 0.00277446 | -0.6358823 | 0.14531556 |
| Ube2l6 | yellow | 2.24248144 | 8.06E-32 | 2.02487774 | 5.30E-26 |
| Clp1 | orange | 0.67707893 | 1.22E-06 | 0.4408802 | 0.00255173 |
| Tfpi | red | -0.5974701 | 3.35E-05 | -0.5831219 | 7.82E-05 |
| Hoxd8 | red | -1.1796596 | 0.00023242 | -0.8508856 | 0.01573086 |
| Chrna1 | red | -0.9623802 | 8.10E-05 | -0.4288344 | 0.13850723 |
| Ola1 | orange | 0.78891918 | 0.00010749 | 0.6061438 | 0.0043131 |
| Kif18a | orange | 1.38892554 | 0.00012537 | 1.30703074 | 0.000474 |
| Lpcat4 | orange | 0.60411323 | 1.48E-06 | 0.38120765 | 0.00383903 |
| Ccdc34 | orange | 1.02658183 | 1.44E-05 | 0.50603958 | 0.04786208 |
| B230118H07 | red | -0.7260319 | 0.00053483 | -0.6249504 | 0.00403687 |
| Depdc7 | red | -0.9881848 | 0.0019641 | -0.8099521 | 0.02268116 |
| Elf5 | skyblue | -1.4895116 | 1.33E-06 | -1.1079713 | 0.00052411 |
| Cat | skyblue | -1.5393353 | 4.53E-07 | -1.2195191 | 0.00010917 |
| Pamr1 | skyblue | -2.3334534 | 3.42E-42 | -1.8223541 | 6.27E-26 |
| Gatm | yellow | 3.57173147 | 1.20E-17 | 3.46161759 | 2.60E-16 |
| Dut | orange | 1.54729606 | 7.19E-30 | 1.39204415 | 4.79E-23 |
| Fbn1 | orange | 1.13786734 | 0.00016104 | 1.43828481 | 2.32E-06 |
| Fgf7 | red | -0.953474 | 7.33E-07 | -0.773973 | 9.64E-05 |
| Meis2 | red | -0.7096721 | 1.30E-05 | -0.4488948 | 0.00850731 |
| Tspan18 | skyblue | -1.8639797 | 4.44E-13 | -1.5530431 | 3.32E-09 |

| | | | | | |
|---|---|---|---|---|---|
| Slc28a2 | yellow | 3.28417913 | 2.67E-20 | 3.3523582 | 8.00E-21 |
| Syt13 | yellow | -1.5351684 | 0.18553218 | 3.32845227 | 0.00337089 |
| Chst1 | red | -0.8738108 | 3.91E-07 | -0.5709205 | 0.00148888 |
| Mapk8ip1 | red | -1.3777559 | 6.16E-10 | -0.8657852 | 0.00017476 |
| Sord | skyblue | -2.1970714 | 3.65E-11 | -1.7165632 | 4.66E-07 |
| Creb3l1 | orange | 1.52375151 | 4.79E-10 | 1.54309743 | 4.69E-10 |
| Frmd5 | skyblue | -1.4628532 | 0.00350695 | -1.2004382 | 0.02069082 |
| Wdr76 | orange | 1.12510663 | 4.71E-11 | 1.03889934 | 3.62E-09 |
| Map1a | red | -1.2108422 | 9.71E-14 | -0.3578464 | 0.0438651 |
| Pacsin3 | red | -0.8051267 | 3.07E-06 | -0.5454864 | 0.00256139 |
| Mkks | skyblue | -1.5013394 | 2.89E-10 | -1.5686946 | 1.28E-10 |
| Jag1 | red | -0.7709761 | 3.72E-05 | -0.4561981 | 0.02049839 |
| Slx4ip | orange | 0.47556569 | 0.04816868 | 0.69482109 | 0.00388331 |
| Ehd4 | red | -0.7519104 | 4.70E-09 | -0.7450818 | 1.05E-08 |
| Itpka | red | -0.7927714 | 1.28E-06 | -0.3791122 | 0.0262349 |
| Ltk | red | -0.7977193 | 0.00910101 | -0.3597309 | 0.28296486 |
| Tyro3 | red | -0.9525972 | 1.33E-13 | -0.3897251 | 0.00351618 |
| Nusap1 | yellow | 2.45866276 | 1.53E-18 | 2.54408953 | 5.82E-19 |
| 4930402H24 | red | -0.7871811 | 5.72E-19 | -0.6077806 | 1.90E-11 |
| Atrn | red | -0.6129502 | 7.96E-05 | -0.4299809 | 0.00805848 |
| Dll4 | skyblue | -1.7692534 | 9.35E-18 | -1.335679 | 2.26E-10 |
| Gfra4 | skyblue | -1.3492403 | 0.0073938 | -1.6867486 | 0.00151227 |
| Ppp1r14d | orange | 1.21558726 | 0.03296245 | 1.78744085 | 0.00239615 |
| Adam33 | skyblue | -1.5574695 | 4.59E-12 | -1.1010978 | 2.09E-06 |
| Siglec1 | yellow | 3.39517186 | 1.29E-07 | 3.09231855 | 2.57E-06 |
| Rad51 | yellow | 3.17697201 | 2.02E-24 | 2.92791691 | 2.82E-20 |
| Casc5 | yellow | 3.59663215 | 1.32E-17 | 3.82898707 | 4.27E-18 |
| Spef1 | skyblue | -1.5951839 | 4.29E-14 | -1.2236978 | 1.58E-08 |
| Knstrn | yellow | 2.54370282 | 2.44E-20 | 2.67424007 | 2.43E-21 |
| Ivd | red | -1.1001146 | 1.04E-05 | -0.8740909 | 0.000712 |
| Smox | orange | 0.99775129 | 0.00047947 | 1.09976862 | 0.00015847 |
| Adra1d | skyblue | -1.4682776 | 8.15E-05 | -1.1450739 | 0.00275612 |
| Rassf2 | orange | 1.37599776 | 6.01E-11 | 1.18090303 | 4.04E-08 |
| Slc23a2 | red | -0.847978 | 1.43E-09 | -0.569351 | 9.02E-05 |
| Pcna | orange | 1.30863308 | 2.11E-16 | 1.04954283 | 1.12E-10 |
| Fsip1 | red | -1.3285758 | 0.00023766 | -1.0607468 | 0.00420746 |
| Rasgrp1 | orange | 1.31127663 | 1.20E-08 | 1.8438599 | 1.18E-15 |
| Fermt1 | red | -0.776034 | 0.00362322 | -0.4270995 | 0.13574805 |
| Slc27a2 | skyblue | -2.1288147 | 1.81E-07 | -1.1440389 | 0.00767897 |
| Hdc | orange | 0.69822131 | 2.55E-07 | 0.78603522 | 9.92E-09 |
| Sppl2a | orange | 0.69639935 | 4.73E-11 | 0.67208117 | 3.85E-10 |
| Dusp2 | yellow | 4.11609364 | 6.39E-42 | 3.93549964 | 2.79E-37 |
| Mall | red | -0.9685441 | 2.17E-06 | -0.627838 | 0.00338924 |

| | | | | | |
|---|---|---|---|---|---|
| Bub1 | yellow | 2.51562672 | 1.59E-21 | 2.81903266 | 2.03E-23 |
| Acoxl | skyblue | -2.3406099 | 9.13E-08 | -1.8100635 | 6.39E-05 |
| Bcl2l11 | orange | 0.68286785 | 3.04E-08 | 0.36927338 | 0.00448162 |
| Fbln7 | skyblue | -1.8793955 | 0.00034954 | -1.617248 | 0.00472462 |
| Ttl | red | -0.973383 | 6.12E-13 | -0.7747619 | 2.16E-08 |
| Slc20a1 | orange | 1.04609963 | 1.38E-09 | 0.90803526 | 2.71E-07 |
| Il1b | yellow | 3.16729043 | 1.00E-24 | 2.6192458 | 7.36E-19 |
| Il1a | orange | 0.17705233 | 0.62172165 | 1.50227289 | 1.86E-06 |
| Nop56 | orange | 0.81569445 | 2.36E-07 | 0.7073838 | 1.29E-05 |
| Cpxm1 | orange | 0.3501494 | 0.2288674 | 0.74842046 | 0.00818788 |
| Vps16 | orange | 0.61228716 | 6.06E-15 | 0.47177446 | 4.50E-09 |
| Rrbp1 | orange | 0.9816198 | 5.02E-06 | 0.92335498 | 2.80E-05 |
| Snx5 | orange | 1.45513579 | 3.19E-12 | 1.10622641 | 2.63E-07 |
| Mgme1 | red | -0.3994836 | 0.02343065 | -0.7396396 | 2.96E-05 |
| Rbbp9 | red | -0.8102744 | 9.49E-07 | -0.7389861 | 1.31E-05 |
| Sec23b | orange | 0.61781659 | 1.61E-08 | 0.58507226 | 1.54E-07 |
| Xrn2 | orange | 0.8000451 | 5.06E-15 | 0.43876868 | 4.34E-05 |
| Cst8 | skyblue | -2.6586 | 4.96E-07 | -2.473142 | 2.24E-06 |
| 9230104L09F | skyblue | -1.4430732 | 0.00018024 | -1.2499471 | 0.00166922 |
| Acss1 | red | -1.3138232 | 4.65E-08 | -0.9012118 | 0.00031376 |
| Gins1 | yellow | 2.5476872 | 5.89E-29 | 1.95518216 | 2.35E-17 |
| Sdcbp2 | orange | 1.46593705 | 2.64E-11 | 1.29864897 | 5.21E-09 |
| Tpx2 | yellow | 3.15184228 | 2.73E-37 | 3.49727021 | 1.48E-43 |
| Kif3b | orange | 0.58131324 | 5.31E-06 | 0.53661997 | 4.18E-05 |
| Bpifa1 | skyblue | -4.1591807 | 4.30E-30 | -3.1776704 | 8.31E-18 |
| Bpifb1 | skyblue | -2.8953319 | 3.14E-08 | -1.2974441 | 0.02009403 |
| E2f1 | orange | 0.77837563 | 0.00078942 | 0.76120995 | 0.00163274 |
| Aurka | yellow | 2.6328452 | 1.25E-39 | 2.62200949 | 3.10E-37 |
| Pkia | red | -1.3294802 | 2.72E-11 | -1.030893 | 5.52E-07 |
| Stmn2 | skyblue | -2.5524991 | 3.12E-20 | -2.135738 | 2.36E-14 |
| Rtfdc1 | orange | 0.64267033 | 1.39E-09 | 0.44132338 | 6.26E-05 |
| Tpd52 | orange | 0.81594895 | 0.00159061 | 0.78258836 | 0.00321913 |
| Pck1 | skyblue | -2.8970678 | 0.00049938 | -1.9568094 | 0.0252842 |
| Zbp1 | black | 5.68035632 | 3.19E-121 | 5.67629748 | 5.06E-121 |
| Edn3 | skyblue | -1.7276328 | 0.00011811 | -1.4666673 | 0.0015614 |
| Ptpn1 | orange | 1.25333908 | 1.20E-12 | 1.07706124 | 2.09E-09 |
| Nfatc2 | orange | 1.43671045 | 4.64E-13 | 1.11441521 | 5.61E-08 |
| Atp9a | red | -0.7916293 | 9.87E-10 | -0.5010475 | 0.00020551 |
| Car13 | orange | 1.0691209 | 0.00010032 | 0.59479351 | 0.04214749 |
| Car3 | skyblue | -2.7019647 | 0.00011085 | -1.8095731 | 0.01360025 |
| Psma7 | orange | 1.14989412 | 2.90E-13 | 0.87924134 | 5.64E-08 |
| Col9a3 | orange | 1.1255738 | 0.00045687 | 0.60047572 | 0.08115039 |
| Nkain4 | skyblue | -1.5145302 | 5.94E-08 | -1.4930934 | 1.73E-07 |

| | | | | | |
|---|---|---|---|---|---|
| Srms | red | -1.1759262 | 6.52E-05 | -1.1848529 | 6.06E-05 |
| Helz2 | orange | 1.49789223 | 2.25E-22 | 1.50001517 | 2.34E-22 |
| Zbtb46 | red | -0.6667721 | 2.43E-05 | -0.4637033 | 0.00480059 |
| Pcmtd2 | red | -0.973674 | 8.00E-05 | -0.8847813 | 0.00050277 |
| Ahcy | orange | 1.19982996 | 7.66E-14 | 0.9611218 | 4.75E-09 |
| Map1lc3a | red | -0.9904057 | 1.66E-08 | -0.743444 | 4.20E-05 |
| Acss2 | red | -1.3286996 | 1.48E-06 | -1.0260844 | 0.00032605 |
| Procr | orange | 1.30831168 | 2.06E-06 | 1.27611281 | 5.63E-06 |
| Mmp24 | yellow | 1.64363118 | 0.0785576 | 3.15957942 | 0.00177083 |
| Eif6 | orange | 1.16344369 | 1.65E-08 | 1.02554469 | 1.19E-06 |
| Dsn1 | orange | 1.5747656 | 1.55E-14 | 1.136092 | 6.39E-08 |
| Sla2 | yellow | 2.7744775 | 1.22E-15 | 3.41531387 | 1.72E-20 |
| Samhd1 | yellow | 2.41887731 | 1.16E-101 | 2.24425411 | 1.83E-87 |
| Rbl1 | orange | 1.54280919 | 3.70E-15 | 1.41347606 | 1.14E-12 |
| Ctnnbl1 | orange | 0.61752371 | 6.13E-05 | 0.61657358 | 9.37E-05 |
| Fam83d | orange | 1.52639173 | 1.78E-06 | 0.67456824 | 0.04904642 |
| Wisp2 | orange | 0.77033925 | 0.00686865 | 0.91339784 | 0.00159781 |
| Ocstamp | black | 6.51065088 | 4.20E-13 | 5.9820777 | 1.12E-11 |
| Ccdc39 | red | -1.1468388 | 8.11E-08 | -0.7472449 | 0.00062175 |
| Mecom | red | -0.8754674 | 4.35E-15 | -0.6160988 | 8.74E-08 |
| Tnik | orange | 0.64961576 | 0.00187566 | 0.7189737 | 0.00073947 |
| Ect2 | yellow | 2.15102236 | 1.08E-16 | 2.50418929 | 8.08E-21 |
| Lrrc34 | skyblue | -2.4410811 | 2.66E-07 | -1.394651 | 0.00311598 |
| Mccc1 | red | -0.8974706 | 4.38E-08 | -0.698972 | 3.64E-05 |
| Exosc9 | orange | 0.76179694 | 3.40E-08 | 0.53393401 | 0.00024168 |
| Ccna2 | yellow | 3.06108674 | 6.54E-26 | 3.20110437 | 1.66E-27 |
| Trpc3 | skyblue | -1.8348851 | 2.05E-06 | -0.7628089 | 0.06321614 |
| Il21 | black | 7.28566282 | 2.24E-08 | 7.09119416 | 9.31E-08 |
| Spata5 | orange | 0.7724285 | 0.00010154 | 0.66493702 | 0.0012962 |
| Slc7a11 | orange | 1.68426267 | 1.66E-07 | 2.3878042 | 2.82E-13 |
| Postn | red | -1.0249378 | 3.02E-09 | -0.6304465 | 0.00048068 |
| Exosc8 | orange | 1.30042096 | 4.03E-16 | 0.92770671 | 1.94E-08 |
| P2ry1 | skyblue | -1.2423024 | 8.04E-06 | -1.656505 | 1.40E-08 |
| Il12a | skyblue | -1.6167191 | 2.82E-05 | -1.3624702 | 0.00120212 |
| Ift80 | red | -0.7935146 | 0.00013857 | -0.549157 | 0.01155879 |
| Ppm1l | red | -1.0162977 | 1.21E-05 | -0.6059164 | 0.01318132 |
| Nmd3 | orange | 0.97172406 | 1.02E-10 | 0.72659479 | 2.99E-06 |
| Bche | skyblue | -1.3187571 | 0.00116887 | -1.2267185 | 0.00338851 |
| Ccna1 | orange | 1.44021285 | 0.00577891 | 2.26217017 | 2.87E-05 |
| Smad9 | skyblue | -2.4660886 | 2.80E-09 | -1.6591941 | 9.05E-05 |
| Dclk1 | orange | 1.06214467 | 0.00454728 | 0.95728863 | 0.01369654 |
| Nbea | red | -0.857202 | 2.50E-11 | -0.7729862 | 2.49E-09 |
| Wwtr1 | red | -0.5937282 | 1.39E-05 | -0.3422994 | 0.01769469 |

| | | | | | |
|---|---|---|---|---|---|
| Pfn2 | red | -0.8831288 | 1.40E-05 | -0.5017857 | 0.01985224 |
| 4930579G24 | orange | 1.40636051 | 3.24E-05 | 0.66907467 | 0.07073885 |
| Mme | skyblue | -1.5448333 | 4.66E-05 | -1.2949526 | 0.00094887 |
| Ptx3 | black | 3.67995015 | 1.72E-15 | 5.13342934 | 3.46E-26 |
| Shox2 | skyblue | -1.6582856 | 0.0061025 | -1.7513961 | 0.00451551 |
| Pdcd10 | orange | 0.71163018 | 4.55E-07 | 0.47589887 | 0.00127407 |
| Ptpn22 | yellow | 3.83089522 | 1.59E-44 | 3.41980799 | 2.41E-35 |
| Dclre1b | orange | 0.74256668 | 8.31E-06 | 0.55979314 | 0.00140218 |
| Tspan2 | red | -1.3091754 | 5.09E-05 | -0.9891741 | 0.00321222 |
| Vangl1 | red | -1.0315198 | 3.37E-12 | -0.6423418 | 2.74E-05 |
| Casq2 | skyblue | -1.7093481 | 0.01200861 | -2.2402145 | 0.00111124 |
| Cd2 | yellow | 2.71677011 | 1.13E-52 | 2.75813975 | 1.02E-52 |
| Ptgfrn | red | -0.820951 | 4.52E-13 | -0.7245642 | 3.31E-10 |
| Spag17 | skyblue | -1.6700772 | 1.13E-07 | -0.6717793 | 0.04181325 |
| Hmgcs2 | red | -1.1061863 | 0.00136175 | -0.6826179 | 0.06199639 |
| 1700013F07I | skyblue | -2.1038509 | 1.05E-08 | -1.1216686 | 0.00180435 |
| Sypl2 | skyblue | -2.124887 | 5.72E-11 | -1.2711926 | 0.00020071 |
| Slc6a17 | skyblue | -2.5421893 | 1.68E-26 | -1.8942711 | 9.98E-17 |
| Kcnc4 | red | -1.3589955 | 0.00164166 | -0.5595206 | 0.24059728 |
| Dennd2d | orange | 0.77402017 | 1.54E-06 | 0.95408258 | 3.95E-09 |
| S100a11 | orange | 0.83620974 | 5.14E-05 | 0.84876012 | 5.82E-05 |
| Crct1 | red | -0.866586 | 0.00262904 | -0.5397202 | 0.07868237 |
| Lce1g | skyblue | -1.8989518 | 4.21E-06 | -0.9292571 | 0.03337837 |
| Lce1b | red | -1.2440742 | 9.04E-05 | -0.6036145 | 0.07881701 |
| Npr1 | red | -1.2545866 | 1.99E-11 | -0.947954 | 8.56E-07 |
| Slc27a3 | red | -0.4997874 | 0.00839633 | -0.5801661 | 0.00275612 |
| Creb3l4 | red | -1.0312626 | 0.00227028 | -0.2574848 | 0.49747867 |
| Tpm3 | orange | 1.07943824 | 1.97E-09 | 0.93614114 | 3.64E-07 |
| Chrnb2 | orange | 1.18940523 | 2.29E-06 | 0.8718494 | 0.00067516 |
| Adar | orange | 1.59872781 | 8.52E-35 | 1.51300148 | 3.06E-31 |
| Pmvk | orange | 0.87674166 | 0.00110024 | 0.70899979 | 0.01097136 |
| Efna1 | skyblue | -1.5753139 | 1.08E-14 | -1.3178895 | 2.16E-10 |
| Fam198b | red | -1.1199154 | 4.11E-09 | -0.7775674 | 8.52E-05 |
| Tmem144 | red | -1.1301624 | 0.00045911 | -0.5002668 | 0.14929956 |
| Sass6 | orange | 1.36633326 | 2.76E-06 | 0.96529068 | 0.0014042 |
| Lrrc39 | skyblue | -1.3404923 | 5.42E-05 | -1.2034545 | 0.00068816 |
| Vcam1 | orange | 1.17434116 | 9.72E-07 | 1.19363785 | 1.03E-06 |
| Larp7 | orange | 0.69479139 | 1.67E-06 | 0.58262869 | 0.00012402 |
| Cyp2u1 | red | -0.9473364 | 0.00526273 | -0.8493946 | 0.01665082 |
| Hadh | red | -0.9540894 | 8.87E-08 | -0.8353024 | 5.09E-06 |
| Ccdc109b | orange | 1.36889959 | 1.37E-10 | 1.04594628 | 2.55E-06 |
| Tlr2 | yellow | 2.29759126 | 7.06E-12 | 1.84068461 | 8.34E-08 |
| Sfrp2 | skyblue | -1.6382995 | 5.50E-09 | -2.0010005 | 2.18E-12 |

| | | | | | |
|---|---|---|---|---|---|
| Fga | black | 5.03812752 | 1.40E-06 | 3.39298003 | 0.00023535 |
| Lrat | skyblue | -2.8025844 | 7.28E-11 | -2.4761674 | 1.67E-08 |
| Gucy1b3 | red | -1.3149027 | 5.84E-08 | -1.0034608 | 6.17E-05 |
| Gar1 | orange | 0.8040405 | 0.00072375 | 0.83231191 | 0.00066261 |
| Egf | red | -0.9821002 | 0.0014973 | -0.4366296 | 0.20410636 |
| Pitx2 | red | -1.4258249 | 0.00022072 | -0.5262737 | 0.23207412 |
| Enpep | skyblue | -1.5410399 | 1.96E-07 | -1.0882158 | 0.0004034 |
| Alpk1 | orange | 0.74218814 | 0.0005392 | 0.74106966 | 0.00074399 |
| Dkk2 | skyblue | -2.7957819 | 4.75E-10 | -1.688101 | 0.00011224 |
| Papss1 | red | -0.5843046 | 1.14E-08 | -0.3722605 | 0.00052127 |
| Kcnq5 | red | -0.6607323 | 0.00841973 | -0.8558679 | 0.000779 |
| Ptgfr | skyblue | -2.9050548 | 2.22E-05 | -2.4495745 | 0.0005258 |
| Ifi44 | yellow | 3.66771846 | 1.10E-50 | 3.90565927 | 1.56E-56 |
| Adam15 | orange | 0.82278861 | 1.41E-07 | 0.67501242 | 2.77E-05 |
| Cks1b | yellow | 2.3715481 | 3.13E-25 | 1.99636558 | 1.89E-17 |
| Thbs3 | skyblue | -2.4298186 | 3.97E-17 | -1.5979421 | 8.38E-08 |
| Gba | orange | 0.80554495 | 7.08E-06 | 0.68388534 | 0.00022199 |
| Sema4a | orange | 1.56428038 | 1.83E-17 | 1.38014621 | 1.35E-13 |
| Pmf1 | orange | 1.46875708 | 7.60E-15 | 1.16988109 | 1.66E-09 |
| Iqgap3 | yellow | 2.04844712 | 1.94E-08 | 2.68172137 | 5.24E-13 |
| Gpatch4 | orange | 0.75762387 | 2.16E-05 | 0.78784496 | 1.56E-05 |
| Sh2d2a | yellow | 4.31842731 | 1.98E-76 | 4.17482948 | 9.44E-71 |
| Dclk2 | orange | 1.01942581 | 5.05E-06 | 1.04387003 | 3.97E-06 |
| Sh3d19 | red | -1.2260686 | 1.21E-10 | -0.8043471 | 4.75E-05 |
| Fmo5 | skyblue | -2.0196551 | 4.57E-06 | -1.6904109 | 0.00019783 |
| Acp6 | red | -0.6011019 | 7.53E-05 | -0.5085617 | 0.00119731 |
| Rnf115 | orange | 0.80192463 | 8.43E-07 | 0.70564866 | 2.56E-05 |
| Polr3c | orange | 0.80820647 | 2.53E-07 | 0.67266633 | 3.14E-05 |
| Bnipl | red | -1.0405034 | 1.24E-05 | -0.7921586 | 0.00140521 |
| Myoz2 | skyblue | -1.4094051 | 0.00054101 | -2.0215734 | 8.64E-07 |
| Abcd3 | red | -0.7974374 | 7.80E-05 | -0.5347629 | 0.01145708 |
| F3 | orange | 2.0047208 | 9.13E-11 | 1.77274073 | 1.90E-08 |
| Tmem56 | red | -0.7288339 | 0.00029732 | -0.2335345 | 0.29264392 |
| Riiad1 | skyblue | -1.5884464 | 3.61E-11 | -1.3137635 | 7.61E-08 |
| Them5 | red | -1.0928169 | 0.00584067 | -0.9148296 | 0.02681502 |
| Mttp | red | -0.9789825 | 9.67E-05 | -0.6185278 | 0.01641561 |
| Dapp1 | orange | 1.17987811 | 2.63E-14 | 1.14734316 | 3.38E-13 |
| Wls | red | -0.7603371 | 7.41E-10 | -0.5859968 | 3.99E-06 |
| Depdc1a | yellow | 3.7791719 | 7.07E-17 | 4.17934855 | 4.80E-15 |
| Uox | skyblue | -2.4942856 | 0.00547869 | -1.5105738 | 0.07793209 |
| Bcl10 | orange | 0.77588837 | 5.05E-08 | 0.51975126 | 0.00046545 |
| Cyr61 | orange | 1.03561192 | 1.42E-05 | 1.21798228 | 4.38E-07 |
| Cryz | red | -0.6964635 | 0.00045687 | -0.522753 | 0.01253056 |

| | | | | | |
|---|---|---|---|---|---|
| Asph | red | -0.7402269 | 6.80E-07 | -0.5432416 | 0.00043729 |
| Ccne2 | orange | 0.74413902 | 0.00527658 | 0.37645991 | 0.20991613 |
| Gem | orange | 0.52485163 | 2.61E-07 | 0.63057062 | 7.46E-10 |
| Cdh17 | black | 5.18140655 | 1.77E-08 | 3.32659774 | 9.97E-08 |
| Decr1 | red | -0.7214225 | 7.33E-06 | -0.7415067 | 6.59E-06 |
| Rmdn1 | red | -1.0532525 | 4.16E-05 | -0.9743635 | 0.00022389 |
| Rps20 | orange | 0.74286277 | 2.28E-15 | 0.62997035 | 4.17E-11 |
| Faxc | red | -1.3716083 | 0.00058822 | -0.6899032 | 0.08554221 |
| Sdcbp | orange | 0.87863491 | 7.11E-06 | 0.68405102 | 0.00074247 |
| Tstd3 | orange | 0.81342767 | 0.00151116 | 0.20351126 | 0.49759145 |
| Clca3a2 | yellow | 2.42914617 | 0.00849033 | 2.67652362 | 0.00085408 |
| Lmo4 | orange | 0.87739404 | 3.32E-07 | 0.6600042 | 0.00021314 |
| Gbp3 | yellow | 3.44887892 | 8.64E-89 | 3.35806653 | 2.21E-84 |
| Gbp2 | yellow | 4.3244958 | 6.93E-193 | 4.13672949 | 7.06E-177 |
| Rragd | red | -1.3598405 | 4.10E-09 | -1.1006594 | 4.18E-06 |
| Casp8ap2 | orange | 0.54882055 | 3.39E-06 | 0.60421025 | 4.81E-07 |
| Slc35a1 | red | -0.6544555 | 2.50E-11 | -0.3513037 | 0.00072977 |
| 1700003M02 | skyblue | -2.2809868 | 2.40E-12 | -1.4458632 | 1.41E-05 |
| Smc2 | orange | 2.02062335 | 8.66E-20 | 1.6011434 | 1.81E-12 |
| Polr1e | orange | 0.82437018 | 3.89E-07 | 0.72468469 | 1.41E-05 |
| Exosc3 | orange | 0.82906301 | 1.79E-07 | 0.65583132 | 6.82E-05 |
| Stra6l | black | 4.3593537 | 9.22E-13 | 4.60986195 | 4.39E-13 |
| Tmod1 | red | -1.0633783 | 0.00742779 | -0.7708506 | 0.06512161 |
| Anp32b | orange | 0.87854113 | 7.92E-08 | 0.69202897 | 4.32E-05 |
| Nans | orange | 0.97047627 | 1.11E-05 | 0.66688274 | 0.00382305 |
| Coro2a | orange | 1.96330114 | 1.30E-18 | 1.4312962 | 3.61E-10 |
| Col15a1 | skyblue | -1.5736867 | 4.52E-10 | -0.8035033 | 0.00254451 |
| Invs | red | -0.9455134 | 8.48E-09 | -0.8515935 | 2.92E-07 |
| Tex10 | orange | 0.74258849 | 9.55E-07 | 0.51919452 | 0.00111028 |
| Tmeff1 | skyblue | -1.2573205 | 8.64E-13 | -1.278248 | 5.29E-13 |
| Murc | red | -1.1827427 | 1.83E-07 | -0.6313686 | 0.00961384 |
| Fmn2 | skyblue | -1.7235736 | 0.00401765 | -1.9867586 | 0.00166547 |
| Zfp618 | orange | 1.20282901 | 1.07E-05 | 1.04599954 | 0.00014405 |
| Tnfsf8 | yellow | 3.21455767 | 1.74E-16 | 2.18605384 | 6.12E-09 |
| Tnc | yellow | 1.96368746 | 0.000762 | 2.45457729 | 3.21E-05 |
| Txn1 | orange | 1.08905788 | 3.20E-19 | 0.99250353 | 6.09E-16 |
| Pappa | orange | 0.90706332 | 0.00404814 | 0.81380906 | 0.01012041 |
| Astn2 | red | -0.9119418 | 1.48E-06 | -0.7237769 | 0.00021615 |
| Ptgr1 | red | -0.9883298 | 6.54E-07 | -0.6744516 | 0.0011287 |
| Ugcg | orange | 0.85977126 | 1.41E-12 | 0.63329395 | 4.57E-07 |
| Ptbp3 | orange | 0.78711177 | 1.37E-08 | 0.60075751 | 2.78E-05 |
| Snx30 | orange | 0.50441863 | 0.00395431 | 0.60625053 | 0.00066322 |
| Wdr31 | red | -1.1061586 | 1.49E-05 | -0.4029911 | 0.14279171 |

| | | | | | |
|---|---|---|---|---|---|
| 2310002L09F | skyblue | -2.3941282 | 1.03E-05 | -1.3324127 | 0.02102977 |
| Ptprd | red | -0.7203967 | 3.63E-05 | -0.7269764 | 5.03E-05 |
| Mpdz | red | -0.8634131 | 6.16E-10 | -0.7478801 | 1.72E-07 |
| Zdhhc21 | orange | 0.65413532 | 2.00E-05 | 0.77424025 | 7.45E-07 |
| B4galt1 | orange | 0.98979995 | 5.49E-08 | 0.66302046 | 0.00047456 |
| Fktn | red | -0.8285963 | 3.79E-07 | -0.5575491 | 0.00106087 |
| Aqp7 | skyblue | -3.1470368 | 2.39E-08 | -2.1041535 | 0.00032626 |
| Epb41l4b | skyblue | -1.565367 | 4.84E-08 | -1.4036665 | 1.69E-06 |
| Kif24 | orange | 1.00310855 | 0.00048668 | 0.93363182 | 0.00096254 |
| 1110017D15 | skyblue | -1.8660387 | 2.12E-17 | -1.3717415 | 5.76E-10 |
| Cntfr | red | -1.0914406 | 0.00018621 | -1.0626474 | 0.00036967 |
| Unc13b | red | -0.7018474 | 5.33E-06 | -0.4719762 | 0.00341132 |
| Cd72 | yellow | 4.33624166 | 5.04E-16 | 4.23870198 | 3.98E-15 |
| Sit1 | yellow | 2.3385861 | 0.00015948 | 3.21426488 | 6.18E-07 |
| Tpm2 | red | -1.1048329 | 1.79E-05 | -0.5580457 | 0.04290622 |
| Creb3 | orange | 0.62813378 | 3.99E-05 | 0.51269267 | 0.00120574 |
| Npr2 | red | -1.2639384 | 2.21E-06 | -0.9069885 | 0.00108436 |
| Reck | skyblue | -1.5831401 | 3.28E-07 | -1.1844066 | 0.00022396 |
| Clta | orange | 0.66549474 | 2.04E-09 | 0.45718269 | 7.54E-05 |
| Glipr2 | yellow | 2.88290405 | 4.98E-35 | 2.73690078 | 1.71E-31 |
| Sh3gl2 | skyblue | -2.2573875 | 2.42E-11 | -1.3301681 | 5.21E-05 |
| Plin2 | orange | 1.87321764 | 1.75E-08 | 1.79175023 | 1.20E-07 |
| Rps6 | orange | 0.59015277 | 9.22E-08 | 0.47108242 | 3.59E-05 |
| Ppap2b | red | -0.9077488 | 3.78E-09 | -0.6257288 | 9.05E-05 |
| Prkaa2 | red | -0.997509 | 0.00024207 | -0.7635771 | 0.0069683 |
| Sgip1 | red | -0.9833913 | 1.91E-05 | -0.4929494 | 0.04435374 |
| Ak4 | red | -0.875964 | 0.00659256 | -0.6955805 | 0.0397134 |
| Dnajc6 | red | -1.3687174 | 0.00231753 | -0.287851 | 0.58249852 |
| Cachd1 | red | -0.6691933 | 1.63E-06 | -0.6576436 | 3.81E-06 |
| 2610528J11F | red | -0.8186639 | 0.00032899 | -0.7186145 | 0.00222948 |
| Dph2 | orange | 0.91013809 | 3.56E-05 | 0.71937922 | 0.00157797 |
| Ttc39a | skyblue | -1.5498641 | 1.85E-15 | -0.9496385 | 1.68E-06 |
| Usp1 | orange | 0.58531262 | 8.41E-05 | 0.57609237 | 0.00016678 |
| Nfia | red | -0.8054853 | 9.72E-08 | -0.9560155 | 3.25E-10 |
| Fggy | red | -0.7030441 | 0.00033608 | -0.921799 | 5.50E-06 |
| Plaa | orange | 0.85204075 | 6.07E-10 | 0.68790426 | 1.18E-06 |
| Laptm5 | yellow | 2.28683185 | 1.12E-22 | 2.07436242 | 1.12E-18 |
| Orc1 | orange | 1.54741331 | 4.08E-05 | 1.54706358 | 6.07E-05 |
| Tnfrsf1b | yellow | 2.33733181 | 4.40E-19 | 2.10604017 | 1.61E-15 |
| Podn | red | -0.5083113 | 0.07508426 | -0.758515 | 0.0076681 |
| Echdc2 | red | -0.9762285 | 4.72E-08 | -0.8358854 | 5.37E-06 |
| Tnfrsf8 | black | 3.91413192 | 9.91E-22 | 5.10014354 | 2.42E-26 |
| Cpt2 | red | -0.6144166 | 0.00043169 | -0.5701169 | 0.00148903 |

| | | | | | |
|---|---|---|---|---|---|
| Magoh | orange | 0.75919518 | 5.44E-07 | 0.58158005 | 0.00023114 |
| Lrp8 | yellow | 3.01135896 | 3.98E-15 | 3.13234363 | 5.35E-16 |
| Tmem59 | red | -0.6590589 | 2.97E-07 | -0.6407923 | 1.05E-06 |
| Cyb5rl | red | -0.7604114 | 2.54E-05 | -0.9069889 | 7.42E-07 |
| Col9a2 | skyblue | -2.2308404 | 0.00218165 | -0.9854805 | 0.19614548 |
| Dyrk2 | red | -0.5978111 | 6.62E-09 | -0.4569824 | 1.83E-05 |
| Kcnq4 | red | -0.9981201 | 0.00019085 | -0.9841892 | 0.00033633 |
| Ctps | orange | 1.56337813 | 1.17E-16 | 1.5775065 | 8.22E-17 |
| Hivep3 | orange | 0.99984689 | 0.00020884 | 0.90937594 | 0.00106138 |
| Ccdc30 | skyblue | -1.5809817 | 2.84E-11 | -0.78703 | 0.00129566 |
| Ccdc23 | orange | 0.82301005 | 3.67E-06 | 0.55182252 | 0.00315263 |
| Slc2a1 | orange | 1.37147957 | 1.30E-10 | 1.14792252 | 1.50E-07 |
| Macf1 | red | -0.697256 | 7.81E-05 | -0.5155279 | 0.00503309 |
| Ppie | orange | 0.64228227 | 0.00066407 | 0.33089066 | 0.10587659 |
| Mycl | red | -0.9788098 | 0.00132218 | -0.7433602 | 0.01911201 |
| Cap1 | orange | 0.89083473 | 5.63E-07 | 0.76034235 | 3.31E-05 |
| Lypla2 | orange | 0.6687576 | 4.32E-06 | 0.46206695 | 0.00237775 |
| Gale | orange | 1.04770629 | 1.27E-08 | 1.07515913 | 8.92E-09 |
| Kif2c | yellow | 3.44056655 | 6.79E-30 | 4.56416104 | 8.40E-41 |
| Plk3 | orange | 2.09036667 | 2.67E-15 | 1.72942048 | 1.25E-10 |
| Mmachc | red | -0.7581528 | 5.71E-05 | -0.6497159 | 0.00093804 |
| Nasp | orange | 1.11443177 | 5.41E-14 | 1.0261063 | 1.12E-11 |
| Pik3r3 | red | -0.5895186 | 0.0008203 | -0.654709 | 0.00027955 |
| Tspan1 | red | -1.2988845 | 7.54E-11 | -0.6321539 | 0.00228536 |
| Lurap1 | red | -1.177116 | 3.86E-05 | -0.8196037 | 0.0057546 |
| Rad54l | yellow | 2.78785465 | 4.94E-15 | 2.38899791 | 3.58E-11 |
| Cyp4b1 | red | -0.9515869 | 0.00187032 | -0.7336886 | 0.02156498 |
| Pdzk1ip1 | red | -1.1317322 | 3.80E-05 | -0.8435256 | 0.00317351 |
| Tal1 | red | -0.8366554 | 3.80E-07 | -0.7642468 | 5.77E-06 |
| Stil | yellow | 3.42185406 | 5.09E-19 | 3.2465678 | 2.97E-17 |
| Cmpk1 | orange | 0.74570076 | 5.17E-07 | 0.70734209 | 3.38E-06 |
| Ebna1bp2 | orange | 0.65751077 | 0.000228 | 0.46918363 | 0.01199588 |
| Cfap57 | skyblue | -2.2690217 | 5.11E-11 | -1.4230477 | 5.76E-05 |
| Aldh4a1 | red | -1.0825155 | 1.39E-06 | -0.8797919 | 0.00014383 |
| Mrto4 | orange | 1.05713411 | 5.64E-06 | 0.92038012 | 0.00012689 |
| Akr7a5 | red | -1.0437141 | 1.01E-12 | -0.6121387 | 6.08E-05 |
| Pqlc2 | orange | 0.82088649 | 1.30E-05 | 0.5546356 | 0.0047989 |
| Capzb | orange | 0.65250445 | 0.00034245 | 0.50380735 | 0.0078765 |
| Pla2g2f | skyblue | -1.4008146 | 0.00153901 | -1.2338065 | 0.00966508 |
| Pla2g2c | skyblue | -1.2573104 | 0.15278508 | -2.6271015 | 0.00435309 |
| Pink1 | red | -0.9590349 | 0.00017045 | -0.9069912 | 0.00054149 |
| Hspg2 | red | -0.7109395 | 0.00011678 | -0.1708877 | 0.42052201 |
| Alpl | orange | 1.52431296 | 1.20E-05 | 2.06588845 | 3.03E-09 |

| | | | | | |
|---|---|---|---|---|---|
| Fabp3 | skyblue | -1.6437394 | 0.0001979 | -1.8402524 | 4.37E-05 |
| Tinagl1 | orange | 1.55785551 | 6.53E-17 | 1.50711395 | 9.49E-16 |
| Hcrtr1 | yellow | 0.85870967 | 0.40178155 | 2.83127537 | 0.00260405 |
| Sema3c | red | -1.3821134 | 5.67E-19 | -1.0217168 | 1.31E-10 |
| Hpca | yellow | 2.81297955 | 1.19E-05 | 3.68249088 | 8.26E-07 |
| Azin2 | red | -0.6222107 | 0.00094266 | -0.2462328 | 0.23757871 |
| Ak2 | orange | 1.21502075 | 2.33E-09 | 1.08005084 | 2.09E-07 |
| Rnf19b | yellow | 2.89270122 | 4.34E-34 | 2.79615396 | 8.46E-32 |
| Phc2 | orange | 0.79325188 | 2.68E-14 | 0.67219434 | 2.34E-10 |
| Eif3i | orange | 0.60016975 | 3.22E-08 | 0.51157097 | 4.58E-06 |
| Stpg1 | skyblue | -2.3101718 | 2.13E-05 | -1.5774544 | 0.00338924 |
| Nipal3 | red | -0.7870506 | 9.52E-05 | -0.6763688 | 0.0011434 |
| Yars | orange | 0.79687447 | 1.49E-06 | 0.70322036 | 3.58E-05 |
| Trim63 | red | -0.7000708 | 0.01940251 | -1.2698072 | 2.65E-05 |
| Slc30a2 | orange | 1.0474029 | 0.02171513 | 1.26055888 | 0.00728937 |
| Psmb2 | orange | 1.2893377 | 1.99E-16 | 0.96788906 | 1.79E-09 |
| Extl1 | red | -1.1756756 | 8.47E-08 | -0.6503672 | 0.005004 |
| Cnksr1 | orange | 0.32887796 | 0.0644357 | 0.72094012 | 3.24E-05 |
| Sh3bgrl3 | orange | 1.60725748 | 5.52E-24 | 1.05816553 | 1.08E-10 |
| Tekt2 | red | -1.3936668 | 2.11E-05 | -0.9377653 | 0.00600034 |
| Gpn2 | orange | 0.60153095 | 2.21E-05 | 0.42482765 | 0.0042765 |
| Gpatch3 | orange | 0.84863835 | 0.00027046 | 0.74390875 | 0.00228536 |
| Slc9a1 | orange | 0.61029467 | 7.88E-05 | 0.55029261 | 0.00054428 |
| Csf3r | yellow | 2.84168277 | 5.36E-18 | 2.59500217 | 3.84E-15 |
| Map3k6 | orange | 0.94805293 | 1.40E-07 | 0.34861173 | 0.0749479 |
| Cdca8 | yellow | 2.95040101 | 1.86E-42 | 2.9920538 | 2.04E-40 |
| Fgr | yellow | 2.96388927 | 8.80E-20 | 2.81133661 | 8.33E-18 |
| Sema3a | red | -0.9428426 | 1.81E-09 | -0.6324955 | 0.00010306 |
| Rpa2 | orange | 1.18994733 | 2.98E-16 | 0.87449966 | 6.44E-09 |
| Smpdl3b | orange | 1.76645725 | 3.86E-08 | 1.58878339 | 1.31E-06 |
| Yrdc | orange | 0.92028481 | 7.16E-08 | 0.65591694 | 0.00022122 |
| Sesn2 | orange | 0.71235206 | 1.21E-06 | 0.39752071 | 0.00976916 |
| Rcc1 | orange | 1.28043066 | 5.18E-11 | 1.42523272 | 4.95E-13 |
| Sf3a3 | orange | 0.6049432 | 1.56E-06 | 0.59192205 | 4.45E-06 |
| Ptpru | skyblue | -1.530632 | 2.06E-07 | -1.287027 | 2.13E-05 |
| Arhgef19 | skyblue | -1.6843322 | 4.49E-16 | -1.1617231 | 4.58E-08 |
| Necap2 | orange | 0.66322716 | 5.34E-09 | 0.57014497 | 1.02E-06 |
| Cdk14 | red | -0.5423444 | 1.32E-05 | -0.6370524 | 4.21E-07 |
| Padi2 | yellow | 2.80107022 | 3.19E-29 | 2.50163104 | 4.21E-23 |
| Kcnab2 | yellow | 2.43041789 | 3.34E-38 | 2.28026459 | 1.29E-33 |
| Psmc2 | orange | 0.74614982 | 9.49E-08 | 0.55049361 | 0.00014697 |
| Hes2 | skyblue | -3.2307578 | 0.00014524 | -2.8490861 | 0.0011567 |
| Espn | red | -1.0023594 | 4.35E-06 | -0.8992525 | 6.70E-05 |

| | | | | | |
|---|---|---|---|---|---|
| Prkag2 | red | -0.6349621 | 0.00032404 | -0.83292 | 3.10E-06 |
| Smarcd3 | red | -0.6794246 | 0.00019365 | -0.3939238 | 0.04349012 |
| Tas1r1 | red | -1.1035712 | 0.0098522 | -0.6083892 | 0.16535684 |
| Per3 | skyblue | -2.0424355 | 3.05E-07 | -1.582278 | 0.00012362 |
| Pgd | orange | 0.61452417 | 0.00146383 | 0.52443336 | 0.00870464 |
| Tnfrsf9 | black | 4.48940356 | 8.08E-46 | 5.5165712 | 3.17E-54 |
| Errfi1 | orange | 0.75700922 | 0.00017572 | 0.40942288 | 0.05763078 |
| Abcb1b | orange | 1.32644361 | 3.29E-08 | 1.08982007 | 1.01E-05 |
| Casz1 | red | -0.6668761 | 2.88E-05 | -0.3553303 | 0.03628659 |
| Slc25a33 | red | -0.98276 | 7.96E-06 | -1.0776123 | 1.66E-06 |
| Klhl7 | red | -0.7630533 | 9.19E-12 | -0.5322138 | 3.78E-06 |
| Angptl7 | red | -0.2724503 | 0.51839664 | -1.3541574 | 0.00058425 |
| Rbp7 | skyblue | -3.0287623 | 0.00035937 | -4.0113711 | 1.42E-06 |
| Mad2l2 | orange | 1.29929645 | 4.40E-15 | 1.07444804 | 2.09E-10 |
| Draxin | orange | 2.15110427 | 0.0001671 | 0.16839377 | 0.80917766 |
| Mthfr | orange | 0.81642295 | 4.64E-05 | 0.73181379 | 0.00038776 |
| Dnajc2 | orange | 1.16078665 | 1.27E-08 | 0.93939544 | 8.62E-06 |
| Miip | orange | 0.61140992 | 2.66E-07 | 0.38156535 | 0.0020675 |
| Trp73 | skyblue | -1.976975 | 2.69E-14 | -1.2859906 | 9.41E-07 |
| Tprgl | red | -0.7429689 | 1.15E-10 | -0.5593714 | 2.50E-06 |
| Pex10 | red | -1.0204617 | 7.80E-11 | -0.8131233 | 5.23E-07 |
| Morn1 | red | -1.1828993 | 7.12E-07 | -0.9197988 | 0.00016175 |
| Prkcz | red | -1.5573695 | 7.88E-09 | -1.0152133 | 0.00029905 |
| Plch2 | red | -1.3176183 | 3.34E-06 | -0.728044 | 0.01471564 |
| Fam213b | red | -0.7520422 | 0.00014081 | -0.4892787 | 0.01959683 |
| Mib2 | red | -1.1248001 | 1.35E-09 | -1.0205302 | 6.88E-08 |
| Mmp23 | red | -0.6310796 | 0.00186238 | -0.3810715 | 0.07587952 |
| Mxra8 | red | -0.965194 | 1.65E-07 | -0.7195997 | 0.00016503 |
| Ttll10 | red | -1.1295416 | 0.00400829 | -0.216012 | 0.63972609 |
| Tnfrsf4 | yellow | 3.29539712 | 1.27E-70 | 2.53925996 | 1.44E-47 |
| Bst1 | orange | 1.9803397 | 2.85E-09 | 1.70242118 | 6.17E-07 |
| Prom1 | skyblue | -1.8516593 | 2.98E-12 | -1.1056415 | 5.36E-05 |
| Adgra2 | red | -0.7094695 | 0.00836011 | -0.4390386 | 0.12616601 |
| D5Ertd615e | yellow | 2.24841904 | 0.05926686 | 4.02845689 | 0.00957667 |
| Ablim2 | red | -1.1816258 | 3.19E-12 | -0.8433096 | 1.68E-06 |
| Htra3 | red | -0.995062 | 6.39E-06 | -0.9559393 | 2.30E-05 |
| Htt | orange | 0.9332273 | 1.19E-10 | 0.91363269 | 4.80E-10 |
| Pcdh7 | red | -0.7865189 | 0.00063809 | -0.8147362 | 0.00055362 |
| Ppp2r2c | skyblue | -1.0943295 | 0.01804987 | -2.4160463 | 8.50E-08 |
| Crmp1 | yellow | 2.83458985 | 4.47E-20 | 2.02123207 | 8.53E-11 |
| Evc | red | -1.3653645 | 4.95E-13 | -1.0781637 | 2.38E-08 |
| Stk32b | skyblue | 0.47083555 | 0.69507448 | -4.2756063 | 0.0041133 |
| Stx18 | orange | 0.59660079 | 6.24E-05 | 0.34855329 | 0.02726685 |

| | | | | | |
|---|---|---|---|---|---|
| Rnf32 | red | -0.9530935 | 0.00665815 | -1.1884831 | 0.0009629 |
| Fosl2 | orange | 0.89925842 | 6.59E-10 | 0.80235457 | 6.50E-08 |
| Rbks | orange | 0.98671381 | 0.00012336 | 0.57924654 | 0.03489359 |
| Ppm1g | orange | 0.64235181 | 2.87E-08 | 0.52465237 | 1.08E-05 |
| Ociad2 | skyblue | -1.6855322 | 5.84E-13 | -1.0369463 | 2.52E-05 |
| Spata18 | skyblue | -2.3885086 | 1.42E-14 | -0.9631432 | 0.00229323 |
| Yipf7 | skyblue | -1.2477174 | 0.06838568 | -2.2177452 | 0.00155275 |
| Emilin1 | orange | 0.77086872 | 0.00890243 | 0.8470685 | 0.00477786 |
| Mapre3 | red | -0.8675276 | 3.75E-17 | -0.5804909 | 5.04E-08 |
| Ppargc1a | red | -1.1337185 | 0.00057026 | -0.8562226 | 0.01278962 |
| Dpysl5 | red | -0.867685 | 0.04295395 | -1.1550279 | 0.00721704 |
| Tbc1d1 | orange | 0.61741025 | 0.0005138 | 0.40826619 | 0.02916142 |
| Slc35f6 | orange | 0.74174904 | 0.00013925 | 0.66162211 | 0.00098223 |
| Cenpa | yellow | 2.21450938 | 2.15E-37 | 1.98964082 | 6.44E-30 |
| 1700001C02 | skyblue | -2.0440568 | 1.85E-09 | -1.5124563 | 1.12E-05 |
| Pi4k2b | red | -0.6002794 | 9.95E-05 | -0.4676166 | 0.00354299 |
| Slc34a2 | red | -1.2722221 | 0.00035411 | -0.7185849 | 0.05869862 |
| Sel1l3 | red | -1.2361348 | 2.56E-07 | -0.1786965 | 0.53164944 |
| Tbc1d14 | orange | 0.68461059 | 0.00043209 | 0.58953248 | 0.00335275 |
| Cckar | skyblue | -2.1228837 | 4.02E-16 | -1.1487705 | 2.38E-05 |
| Klb | skyblue | -1.6474993 | 0.0024469 | -1.5350131 | 0.01169121 |
| Rhoh | yellow | 2.11035044 | 4.70E-12 | 2.86351387 | 1.04E-20 |
| Tec | orange | 1.24760008 | 9.97E-09 | 0.96220423 | 1.98E-05 |
| Lnx1 | red | -0.985466 | 0.00015617 | -0.5582932 | 0.04030695 |
| Chic2 | orange | 0.65325057 | 5.53E-07 | 0.55072375 | 4.33E-05 |
| Thegl | skyblue | -1.8821256 | 0.00760501 | -1.1069341 | 0.11472406 |
| Cenpc1 | orange | 0.66447547 | 0.00028247 | 0.3583239 | 0.06834015 |
| Stap1 | orange | 0.98994486 | 0.00022501 | 1.38128875 | 7.59E-07 |
| Pigg | red | -0.7097775 | 1.70E-05 | -0.47018 | 0.00633347 |
| Sult1d1 | skyblue | -2.2846078 | 7.84E-08 | -1.909575 | 1.09E-05 |
| Gfi1 | yellow | 2.88649373 | 7.74E-11 | 4.10148754 | 2.39E-18 |
| Glmn | orange | 1.29400107 | 7.04E-05 | 0.62750058 | 0.07154206 |
| Cdc7 | orange | 1.42123482 | 7.26E-09 | 1.07763151 | 2.04E-05 |
| Tgfbr3 | red | -0.7147809 | 0.0041586 | -0.6753288 | 0.00848929 |
| Rufy3 | orange | 0.70225576 | 0.00177386 | 0.77340393 | 0.00076758 |
| Gbp9 | yellow | 2.23113011 | 4.60E-47 | 2.31488438 | 1.37E-50 |
| Abcg3 | red | -1.122505 | 0.00302188 | -0.343773 | 0.42326176 |
| Spp1 | orange | 2.09524979 | 5.61E-08 | 1.52135431 | 0.00014052 |
| Hsd17b11 | red | -0.5312612 | 2.07E-05 | -0.6184136 | 1.02E-06 |
| Klhl8 | red | -1.0585135 | 2.17E-06 | -0.73849 | 0.00137683 |
| Agpat9 | red | -0.7358525 | 4.63E-05 | -0.2097197 | 0.30490797 |
| Plac8 | yellow | 3.75298729 | 3.65E-44 | 3.80261857 | 3.64E-45 |
| Bmp3 | skyblue | -1.7318463 | 1.76E-09 | -1.1299164 | 0.00016107 |

| | | | | | |
|---|---|---|---|---|---|
| Antxr2 | red | -1.129878 | 3.83E-13 | -0.9645241 | 1.16E-09 |
| Srrd | orange | 0.40870236 | 0.06716095 | 0.66849857 | 0.00315361 |
| Crybb3 | skyblue | -1.7331174 | 9.40E-08 | -1.1499251 | 0.00052507 |
| Rfc5 | orange | 1.6286037 | 1.90E-19 | 1.43268537 | 6.57E-15 |
| Dck | orange | 1.81224928 | 1.51E-11 | 1.96087024 | 4.66E-13 |
| Cxcl5 | yellow | 4.39491616 | 5.43E-15 | 4.19329305 | 1.34E-19 |
| Ppbp | red | 0.04260219 | 0.88747658 | -1.019683 | 5.99E-05 |
| Ereg | orange | 1.73982006 | 0.00405517 | 1.51369188 | 0.01530564 |
| Areg | yellow | 2.12263936 | 1.45E-05 | 2.58552703 | 1.80E-07 |
| Cxcl3 | black | 5.40938834 | 3.18E-06 | 7.76851641 | 1.65E-07 |
| Cxcl1 | yellow | 2.9206574 | 1.92E-21 | 3.81787166 | 1.25E-33 |
| Tctn2 | red | -0.711747 | 3.63E-05 | -0.3900668 | 0.0320844 |
| Rilpl1 | red | -0.6717476 | 9.65E-05 | -0.5781568 | 0.00116141 |
| Rilpl2 | orange | 1.11497884 | 2.19E-12 | 0.99611206 | 6.60E-10 |
| Cdkl2 | red | -1.069707 | 1.11E-05 | -0.8699231 | 0.00055059 |
| G3bp2 | orange | 0.74742495 | 3.64E-06 | 0.62299445 | 0.00018852 |
| Naaa | orange | 0.85495411 | 3.96E-07 | 0.75104165 | 1.51E-05 |
| Kntc1 | yellow | 2.87639995 | 5.84E-21 | 2.8323872 | 4.24E-19 |
| Sdad1 | orange | 1.15275338 | 5.56E-07 | 0.98142253 | 3.44E-05 |
| Cxcl9 | black | 8.52371505 | 3.32E-96 | 7.23535103 | 3.14E-81 |
| Rimbp2 | skyblue | -2.3215186 | 0.00079437 | -0.415504 | 0.54149282 |
| Ran | orange | 1.17163114 | 3.85E-10 | 1.07063844 | 2.00E-08 |
| Gbas | red | -0.702916 | 7.37E-06 | -0.7049017 | 1.09E-05 |
| Bcl7a | red | -1.398817 | 1.67E-06 | -0.9372523 | 0.00206908 |
| Wdr66 | skyblue | -1.8647158 | 5.67E-14 | -1.0425489 | 4.95E-05 |
| Psph | orange | 0.7320747 | 0.00061747 | 0.41963986 | 0.06774657 |
| Cct6a | orange | 0.58893466 | 0.00023033 | 0.39240101 | 0.01972635 |
| Rhof | yellow | 2.07326777 | 9.10E-24 | 2.05166292 | 5.10E-23 |
| Aldh2 | red | -0.8055961 | 0.00081065 | -0.6507138 | 0.00917888 |
| Acad10 | red | -1.0869877 | 5.12E-10 | -0.9811829 | 3.14E-08 |
| Arpc3 | orange | 0.78315496 | 1.02E-10 | 0.71674213 | 6.44E-09 |
| Ift81 | red | -1.281067 | 1.92E-13 | -0.9388803 | 1.51E-07 |
| Rnf34 | orange | 0.61546419 | 0.00681594 | 0.38268535 | 0.11527882 |
| Morn3 | skyblue | -1.4919262 | 9.66E-08 | -1.4627563 | 3.41E-07 |
| Anxa3 | orange | 0.66258922 | 3.32E-06 | 0.73403426 | 3.73E-07 |
| Mfsd7a | orange | 0.86537037 | 4.85E-05 | 0.7996787 | 0.00029142 |
| Pxmp2 | red | -1.110927 | 0.00107841 | -1.0184492 | 0.00378844 |
| Ep400 | orange | 0.59066444 | 5.40E-07 | 0.45752079 | 0.00017677 |
| Pus1 | orange | 0.74253962 | 0.00234374 | 0.81140273 | 0.00114737 |
| Ulk1 | red | -0.6045476 | 0.00215214 | -0.3821362 | 0.06748333 |
| Cit | orange | 0.97626971 | 4.64E-09 | 0.69816809 | 5.12E-05 |
| Pla2g1b | skyblue | -1.8277961 | 0.0005801 | -1.2158691 | 0.02815754 |
| Sirt4 | red | -0.7519682 | 2.60E-05 | -0.605933 | 0.00106959 |

| | | | | | |
|---|---|---|---|---|---|
| Ccr9 | yellow | -0.4670977 | 0.73236893 | 3.46769016 | 0.00444269 |
| Srsf9 | orange | 0.67742986 | 3.92E-06 | 0.50148278 | 0.00107332 |
| Tes | orange | 1.17709859 | 2.00E-13 | 0.91818666 | 2.32E-08 |
| Tfec | orange | 2.07698228 | 2.24E-09 | 1.51181539 | 2.30E-05 |
| Oasl2 | yellow | 3.46265124 | 1.03E-93 | 3.45947281 | 1.90E-93 |
| Foxp2 | skyblue | -1.2202087 | 2.30E-07 | -1.3852805 | 1.02E-08 |
| Lfng | orange | 0.77985918 | 3.01E-10 | 0.44758919 | 0.00057679 |
| Mmab | red | -0.8439513 | 3.68E-06 | -0.4063028 | 0.0369049 |
| Radil | red | -1.1729967 | 3.18E-05 | -0.5708847 | 0.05984284 |
| Actb | orange | 1.32325685 | 3.43E-14 | 1.08025871 | 1.36E-09 |
| Fscn1 | orange | 1.54924773 | 4.93E-11 | 1.4117147 | 4.20E-09 |
| Zfp12 | red | -0.5980013 | 0.00051249 | -0.5708696 | 0.0012167 |
| Ung | orange | 1.21058673 | 0.00035704 | 1.05033686 | 0.0029951 |
| Rita1 | red | -0.9089466 | 1.43E-06 | -0.5724435 | 0.0039453 |
| Iqcd | skyblue | -2.5752971 | 4.01E-11 | -1.4064813 | 0.00020958 |
| Oas1b | yellow | 2.04718971 | 1.44E-40 | 2.14323843 | 8.09E-45 |
| Rpl6 | orange | 0.72241059 | 4.47E-08 | 0.46513235 | 0.00074247 |
| Erp29 | orange | 0.90553361 | 1.24E-14 | 0.64620454 | 1.04E-07 |
| Ccz1 | orange | 0.73279339 | 5.35E-10 | 0.47832584 | 0.00010506 |
| Arpc1b | orange | 1.55833019 | 5.35E-13 | 1.30816838 | 2.96E-09 |
| Cpsf4 | orange | 0.8785612 | 1.90E-12 | 0.69287769 | 5.94E-08 |
| Wasf3 | skyblue | -1.6561524 | 2.21E-09 | -1.3439922 | 2.27E-06 |
| Pomp | orange | 1.07398216 | 1.21E-08 | 0.8508463 | 1.22E-05 |
| Slc46a3 | red | -0.6973694 | 1.99E-06 | -0.4883349 | 0.00125124 |
| Mtus2 | skyblue | -1.6356316 | 0.0007436 | -2.0661364 | 2.58E-05 |
| Hsph1 | red | -1.060138 | 5.92E-07 | -1.1117724 | 2.63E-07 |
| Tex26 | skyblue | -1.2250093 | 0.00488549 | -1.3340266 | 0.00161749 |
| Tfpi2 | orange | 1.56634253 | 0.00327734 | 1.35488396 | 0.01405996 |
| Tspan12 | skyblue | -1.9587273 | 1.74E-26 | -1.6046898 | 5.50E-18 |
| Auts2 | red | -1.0357689 | 0.00339964 | -0.0051596 | 0.99114126 |
| Limk1 | orange | 0.72414075 | 0.00010251 | 0.36749865 | 0.06589018 |
| Eln | orange | 2.03677434 | 0.00106152 | 1.83945551 | 0.00412022 |
| Lmod2 | red | -0.7257163 | 0.02189539 | -0.8613284 | 0.00742969 |
| Asb15 | skyblue | -2.3354091 | 0.0001983 | -2.0412697 | 0.00154797 |
| Ezh2 | orange | 1.50208739 | 6.09E-16 | 1.42263867 | 4.00E-14 |
| Aass | skyblue | -2.6937917 | 7.68E-05 | -1.3151655 | 0.06782833 |
| Ssc4d | orange | 1.20005376 | 0.00054055 | 0.82854234 | 0.02049127 |
| Ephb4 | red | -1.0416505 | 1.39E-11 | -0.8779478 | 2.47E-08 |
| Pop7 | orange | 0.81676631 | 1.69E-05 | 0.85391554 | 1.48E-05 |
| Pcolce | red | -0.6459248 | 4.30E-05 | -0.3097862 | 0.0685469 |
| Agfg2 | orange | 0.62171791 | 8.06E-06 | 0.49654917 | 0.0005688 |
| Tsc22d4 | orange | 0.58376545 | 5.50E-05 | 0.5090701 | 0.00065572 |
| Mcm7 | orange | 1.34612703 | 5.64E-26 | 1.03475387 | 1.61E-15 |

| | | | | | |
|---|---|---|---|---|---|
| Tpk1 | red | -0.7390269 | 4.71E-05 | -0.5365647 | 0.00367343 |
| Asns | orange | 0.79314003 | 7.49E-08 | 0.88281374 | 3.24E-09 |
| Pon3 | red | -0.7030507 | 2.03E-05 | -0.6191967 | 0.00027033 |
| Akr1b8 | orange | 0.77950703 | 0.00597062 | 0.84904941 | 0.00328913 |
| Ccdc136 | red | -1.0384477 | 2.30E-05 | -0.7005989 | 0.00788897 |
| Irf5 | yellow | 2.57261272 | 1.84E-14 | 2.73363534 | 5.34E-16 |
| Gars | orange | 1.05815091 | 2.00E-09 | 0.9584644 | 1.01E-07 |
| Adcyap1r1 | skyblue | -1.6408067 | 8.71E-10 | -0.7181945 | 0.00888122 |
| Nt5c3 | orange | 1.14854757 | 1.89E-15 | 0.9519892 | 1.35E-10 |
| Fkbp9 | red | -0.6910305 | 3.20E-12 | -0.4943849 | 1.41E-06 |
| Sspo | skyblue | -2.1607375 | 4.33E-07 | -1.5102153 | 0.0005291 |
| Herc6 | yellow | 2.77275405 | 2.75E-92 | 2.84064924 | 5.52E-96 |
| Herc3 | orange | 1.03415776 | 8.04E-14 | 0.78267582 | 3.49E-08 |
| Igf2bp3 | orange | 1.25364268 | 3.01E-05 | 0.9079699 | 0.00291651 |
| Malsu1 | orange | 0.77459782 | 1.88E-05 | 0.63840443 | 0.00063934 |
| Gpnmb | orange | 1.6532211 | 3.24E-06 | 1.44818826 | 7.31E-05 |
| Dfna5 | orange | 1.01180581 | 0.00187347 | 0.243506 | 0.52381811 |
| Osbpl3 | orange | 0.82478465 | 3.65E-06 | 0.40462904 | 0.03363885 |
| Zc3hav1 | orange | 1.31746292 | 3.09E-23 | 1.1627484 | 3.79E-18 |
| Tmem213 | yellow | 3.40297014 | 0.00081614 | 0.96705523 | 0.23077053 |
| Nfe2l3 | red | -0.9262184 | 9.53E-05 | -0.3868728 | 0.1193793 |
| Ptn | red | -1.2540837 | 4.60E-06 | -0.8926068 | 0.00186044 |
| Tcaf2 | red | -0.7096781 | 0.00015204 | -0.5360483 | 0.00633554 |
| Epha1 | skyblue | -1.3653616 | 1.90E-13 | -1.1669292 | 6.10E-10 |
| Fam131b | skyblue | -1.146721 | 0.01203838 | -1.6280178 | 0.00033516 |
| Clcn1 | skyblue | -1.3245402 | 5.13E-07 | -1.2708627 | 4.73E-06 |
| Gstk1 | red | -1.2205254 | 5.19E-09 | -0.8262503 | 0.00013879 |
| Ccdc184 | orange | 1.03753329 | 0.00510717 | 1.33396329 | 0.00037182 |
| Mad2l1 | orange | 1.90913921 | 3.95E-24 | 1.82769355 | 1.30E-21 |
| Clec5a | yellow | 3.71587427 | 7.50E-12 | 3.24848886 | 3.87E-09 |
| Hpgds | orange | 2.06345845 | 1.64E-07 | 1.53623007 | 0.00017702 |
| Rab19 | orange | 1.4433583 | 8.45E-16 | 1.2982441 | 1.21E-12 |
| Tbxas1 | yellow | 2.16929198 | 1.84E-09 | 1.7678739 | 1.91E-06 |
| Pcyox1 | red | -0.9346421 | 8.76E-09 | -0.8825596 | 9.58E-08 |
| Add2 | skyblue | -0.9173144 | 0.32264548 | -2.7803121 | 0.00950464 |
| Nat8 | skyblue | -2.1426549 | 0.00066998 | -1.0900625 | 0.07642089 |
| Pradc1 | red | -0.6468328 | 7.69E-06 | -0.5541356 | 0.00025379 |
| Prickle2 | red | -0.6886722 | 1.25E-06 | -0.4215459 | 0.00467592 |
| Adamts9 | orange | 1.36653111 | 1.05E-06 | 1.38521661 | 1.19E-06 |
| Lrig1 | red | -1.0103402 | 2.63E-09 | -0.5206566 | 0.00349558 |
| 1700003E16l | skyblue | -1.5879454 | 6.89E-10 | -1.0574412 | 5.45E-05 |
| Wdr54 | red | -0.8399622 | 0.00171808 | -1.0744691 | 9.81E-05 |
| Wbp1 | red | -0.6368855 | 5.86E-07 | -0.6278351 | 1.51E-06 |

| | | | | | |
|---|---|---|---|---|---|
| M1ap | orange | 0.38285766 | 0.42727138 | 1.51680648 | 0.00111233 |
| Tacr1 | skyblue | -4.5859018 | 1.36E-06 | -2.2381826 | 0.00364222 |
| Mrpl19 | orange | 0.81589381 | 3.63E-08 | 0.57363163 | 0.00021372 |
| Bmp10 | orange | 9.82627151 | 0.00121101 | -10.742795 | 0.00048251 |
| Arhgap25 | orange | 1.27983118 | 2.88E-15 | 1.13978446 | 4.68E-12 |
| Gkn3 | skyblue | -3.9098689 | 7.35E-11 | -7.4370597 | 2.22E-08 |
| Rab43 | orange | 1.71107005 | 2.33E-08 | 1.6727655 | 9.61E-08 |
| Hmces | orange | 0.58054546 | 4.06E-06 | 0.35990805 | 0.00671882 |
| Foxp1 | red | -0.7229771 | 8.93E-07 | -0.6893183 | 4.62E-06 |
| Gxylt2 | skyblue | -1.6224271 | 1.41E-10 | -0.9079192 | 0.00079837 |
| Chl1 | orange | 1.20971742 | 0.00061846 | 0.54417818 | 0.15871249 |
| Plxna1 | red | -0.7651453 | 1.65E-06 | -0.3789263 | 0.02586575 |
| Chchd6 | red | -0.654028 | 0.00014762 | -0.3817841 | 0.03733686 |
| Nup210 | orange | 1.2649421 | 2.62E-22 | 1.25811032 | 7.79E-22 |
| Cntn6 | skyblue | -1.8530184 | 0.0885198 | -3.4200678 | 0.0095846 |
| Xpc | red | -0.9242107 | 9.49E-06 | -0.6059754 | 0.00544605 |
| Grip2 | skyblue | -1.6779047 | 1.15E-09 | -0.8612214 | 0.00111246 |
| Bhlhe40 | orange | 0.51841391 | 0.00517748 | 0.58638129 | 0.00188191 |
| Edem1 | orange | 1.23003585 | 9.85E-22 | 1.09737756 | 2.66E-17 |
| Usp18 | yellow | 3.03309456 | 8.25E-59 | 3.00089597 | 2.57E-57 |
| Slc6a12 | yellow | 2.44798666 | 0.00629008 | 3.36190867 | 0.0005893 |
| Ret | red | -0.9969143 | 0.00013108 | -0.7551985 | 0.00553924 |
| A2m | yellow | 3.40058846 | 0.00024976 | 2.75460429 | 0.00400887 |
| Klrg1 | orange | 1.77652581 | 3.47E-06 | 1.30211463 | 0.00123253 |
| Mfap5 | red | -0.6427945 | 0.00092249 | -0.3080674 | 0.14421519 |
| Gdf3 | black | 5.432136 | 0.00063448 | 3.31155009 | 0.00677079 |
| Plxnd1 | red | -0.5855988 | 0.00172804 | -0.3320741 | 0.097391 |
| Lag3 | black | 4.73500328 | 2.52E-64 | 4.50369151 | 4.65E-57 |
| Lrrc23 | skyblue | -1.648105 | 1.71E-10 | -1.1623756 | 1.31E-05 |
| Rasgef1a | skyblue | -1.7173165 | 2.40E-20 | -1.3813238 | 2.53E-13 |
| Clec4e | black | 5.52238957 | 9.74E-23 | 4.4059342 | 4.64E-17 |
| Clec4d | yellow | 4.00745755 | 7.56E-16 | 3.51895888 | 8.22E-13 |
| Zfp248 | red | -1.0134222 | 0.00010477 | -1.022748 | 0.00013813 |
| Clec4a2 | yellow | 3.48013377 | 3.82E-16 | 2.69461023 | 6.71E-10 |
| Klrk1 | yellow | 3.8774696 | 1.90E-48 | 3.53257781 | 1.04E-39 |
| Klrb1f | orange | 1.78022302 | 2.67E-05 | 1.78567446 | 0.0001106 |
| Cd69 | yellow | 3.03477559 | 1.11E-17 | 3.29036105 | 1.90E-19 |
| Clec12b | red | -1.0858261 | 0.00115013 | -1.2820783 | 0.00016175 |
| Clec1b | red | -0.2295159 | 0.35323928 | -1.0879266 | 2.69E-06 |
| Gabarapl1 | red | -0.74104 | 6.97E-06 | -0.5857295 | 0.00059635 |
| Klrd1 | yellow | 3.14350741 | 2.13E-44 | 2.74158128 | 1.86E-32 |
| Klrc1 | black | 5.04939511 | 1.11E-31 | 4.7918127 | 5.11E-27 |
| Klra13-ps | orange | 1.28427564 | 3.00E-05 | 1.99956186 | 3.41E-09 |

| | | | | | |
|---|---|---|---|---|---|
| Klra2 | yellow | 3.64688861 | 1.77E-18 | 2.94996224 | 2.41E-12 |
| Magohb | orange | 1.74912677 | 2.67E-14 | 1.41037571 | 4.23E-09 |
| Etv6 | orange | 1.32911412 | 6.47E-13 | 1.2365671 | 4.01E-11 |
| Bcl2l14 | orange | 0.89682487 | 0.00055555 | 0.86099803 | 0.00114502 |
| Lrp6 | red | -0.9192465 | 7.43E-09 | -0.71225 | 1.43E-05 |
| Emp1 | orange | 0.76056816 | 0.00945643 | 0.8535115 | 0.00419239 |
| Plbd1 | orange | 1.26816443 | 2.46E-09 | 1.03106689 | 2.52E-06 |
| Art4 | skyblue | -1.4707708 | 8.38E-12 | -1.2825113 | 4.83E-09 |
| Mgp | orange | 0.56555091 | 0.02812293 | 0.7199717 | 0.0055993 |
| Arhgdib | yellow | 2.30395588 | 3.43E-27 | 2.31216667 | 3.47E-27 |
| Rerg | red | -0.8934456 | 6.63E-05 | -0.7792076 | 0.00074331 |
| Ptpro | orange | 1.74895157 | 9.68E-06 | 1.29800143 | 0.00159849 |
| Dera | orange | 1.60836886 | 7.76E-13 | 1.38469277 | 1.48E-09 |
| Aebp2 | orange | 0.71851177 | 7.22E-09 | 0.59232505 | 3.57E-06 |
| Ldhb | skyblue | -1.6593725 | 5.40E-07 | -1.1189703 | 0.0011811 |
| Abcc9 | red | -1.4117483 | 1.72E-08 | -1.0934454 | 2.55E-05 |
| Rad18 | orange | 1.62902984 | 6.79E-10 | 1.74289207 | 7.50E-11 |
| Sspn | skyblue | -1.7174248 | 1.96E-07 | -1.6589427 | 8.46E-07 |
| Bhlhe41 | skyblue | -2.3250326 | 3.48E-19 | -1.8199108 | 5.95E-12 |
| Srgap3 | orange | 0.70981437 | 2.75E-05 | 0.33994279 | 0.06077967 |
| Rassf8 | red | -1.183507 | 1.02E-09 | -0.921521 | 3.82E-06 |
| Lrmp | yellow | 2.04344378 | 9.24E-21 | 2.38606928 | 2.32E-27 |
| Bcat1 | yellow | 2.40049017 | 3.22E-08 | 2.58064961 | 6.52E-09 |
| Ttll3 | red | -0.8502716 | 7.80E-05 | -0.5986066 | 0.00705883 |
| Cidec | skyblue | -2.3523216 | 0.00348584 | -1.3561717 | 0.11603848 |
| St8sia1 | orange | 0.94214502 | 0.0006625 | 1.55097528 | 5.25E-08 |
| Creld1 | red | -0.6667175 | 3.24E-05 | -0.6318126 | 0.0001279 |
| Ccdc91 | red | -0.9373402 | 4.62E-08 | -0.758992 | 1.86E-05 |
| Atp2b2 | skyblue | -3.6169843 | 8.65E-18 | -1.8789065 | 6.05E-06 |
| Caprin2 | red | -0.8503119 | 0.00470179 | -0.8603995 | 0.00485234 |
| Tamm41 | orange | 0.62817899 | 9.51E-05 | 0.58941431 | 0.00042659 |
| Timp4 | red | -0.8651618 | 0.00315351 | -0.7083943 | 0.01811731 |
| Cand2 | red | -1.0140845 | 9.32E-05 | -0.5638679 | 0.04181228 |
| Efcab12 | red | -1.211849 | 1.24E-06 | -0.8013646 | 0.00103539 |
| Ift122 | red | -0.990645 | 3.40E-17 | -0.6900733 | 8.67E-09 |
| Klrb1c | orange | 1.14168187 | 0.00415011 | 1.76995456 | 1.82E-05 |
| Pianp | red | -1.3318963 | 0.00047921 | -0.6062061 | 0.11736503 |
| Cd27 | yellow | 2.43488522 | 1.93E-14 | 2.99669662 | 1.62E-20 |
| Tnfrsf1a | orange | 0.68829969 | 1.12E-07 | 0.72395371 | 3.63E-08 |
| Rad51ap1 | yellow | 2.58525098 | 9.92E-14 | 2.80101264 | 6.28E-15 |
| Fkbp4 | red | -0.980978 | 4.65E-16 | -0.813708 | 4.00E-11 |
| Klrb1a | yellow | 1.10445229 | 0.18047469 | 3.58031817 | 0.00076565 |
| Clec2i | yellow | 1.8629233 | 3.08E-19 | 2.27011841 | 2.36E-25 |

| | | | | | |
|---|---|---|---|---|---|
| Slc8a2 | red | -1.3649121 | 1.02E-05 | -0.7348395 | 0.02022941 |
| Zfp606 | red | -0.6718606 | 5.56E-05 | -0.6220075 | 0.00031141 |
| Ckm | red | -1.1667208 | 0.00012385 | -0.8351933 | 0.00851912 |
| Rtn2 | red | -1.1569314 | 1.53E-05 | -1.1317295 | 3.99E-05 |
| Vasp | orange | 1.13493125 | 8.28E-09 | 1.0084634 | 5.59E-07 |
| Dmpk | red | -1.4679612 | 1.50E-15 | -1.0561695 | 2.46E-08 |
| Pglyrp1 | orange | 1.46957846 | 2.69E-10 | 1.25478953 | 1.46E-07 |
| Pdcd5 | orange | 0.58070169 | 0.00010154 | 0.51952869 | 0.00077229 |
| Pop4 | orange | 0.64183089 | 4.45E-07 | 0.47800687 | 0.00031695 |
| Lilra6 | orange | 1.53687818 | 0.00032282 | 1.2643772 | 0.00353344 |
| Rpl28 | orange | 0.67180984 | 2.57E-05 | 0.53008848 | 0.00135235 |
| Sbk2 | skyblue | -1.854689 | 7.35E-08 | -1.5742019 | 9.27E-06 |
| Zfp583 | red | -0.9118339 | 0.0034664 | -0.7374192 | 0.02122449 |
| Zfp273 | red | -0.8196734 | 0.00418404 | -0.8214352 | 0.00665188 |
| Nipa2 | orange | 0.86375554 | 2.77E-06 | 0.78607896 | 3.37E-05 |
| Psd3 | red | -1.0626422 | 1.00E-09 | -0.951179 | 8.37E-08 |
| Zfp719 | red | -0.6510426 | 0.00401514 | -0.4102861 | 0.09017366 |
| Csrp3 | skyblue | -2.0615436 | 2.33E-08 | -1.7634328 | 3.33E-06 |
| Zdhhc13 | orange | 0.60036065 | 6.98E-06 | 0.59832813 | 1.36E-05 |
| Siglece | orange | 1.31971926 | 0.00029756 | 1.13546177 | 0.00256066 |
| Cyp2b10 | skyblue | -1.4827135 | 0.00885927 | -1.2759437 | 0.02988239 |
| Zfp108 | skyblue | -2.0986252 | 5.63E-08 | -0.9573542 | 0.01534389 |
| Faap24 | orange | 0.98137718 | 1.36E-08 | 0.66318248 | 0.00026836 |
| Slc7a10 | skyblue | -2.7175573 | 3.99E-12 | -2.2882686 | 1.04E-08 |
| Kctd15 | red | -0.6821914 | 1.02E-08 | -0.5753511 | 2.87E-06 |
| Snrpa1 | orange | 1.12506864 | 2.27E-10 | 0.80886637 | 1.29E-05 |
| Pcsk6 | skyblue | -1.8621045 | 2.10E-15 | -1.2468238 | 2.39E-07 |
| Tarsl2 | red | -0.879671 | 7.73E-06 | -0.7624522 | 0.00021159 |
| Tjp1 | red | -0.7443029 | 1.55E-05 | -0.5165387 | 0.00401979 |
| Apba2 | skyblue | -1.6571659 | 0.00173766 | -0.6396192 | 0.23509991 |
| Mphosph10 | orange | 0.74150348 | 1.60E-05 | 0.37843996 | 0.04027914 |
| Chrna7 | yellow | 1.92366612 | 0.00058605 | 2.88425857 | 6.21E-07 |
| Blm | orange | 2.00224755 | 2.29E-13 | 1.6789603 | 1.87E-09 |
| Furin | orange | 1.02274974 | 9.43E-07 | 0.84297776 | 8.88E-05 |
| Hddc3 | red | -1.4086911 | 0.00050483 | -0.9888597 | 0.01783903 |
| Cib1 | orange | 0.60606554 | 0.00016381 | 0.42791569 | 0.01133761 |
| Idh2 | red | -0.9371183 | 4.00E-08 | -0.8105617 | 3.65E-06 |
| Mesp2 | skyblue | -1.6747995 | 9.54E-05 | -1.3262247 | 0.00300698 |
| Mesp1 | skyblue | -3.6224556 | 1.76E-08 | -1.4045243 | 0.02111241 |
| Plin1 | skyblue | -3.0274686 | 7.07E-05 | -2.0031319 | 0.0120582 |
| Nr2f2 | red | -0.8097704 | 3.39E-11 | -0.6455531 | 2.62E-07 |
| Pgpep1l | skyblue | -1.903134 | 5.40E-08 | -1.5074689 | 0.00024059 |
| Synm | skyblue | -2.2731173 | 2.77E-09 | -1.6735397 | 2.32E-05 |

| | | | | | |
|---|---|---|---|---|---|
| Ttc23 | red | -0.8022472 | 7.10E-06 | -0.4563635 | 0.0151002 |
| Rab38 | red | -0.891071 | 7.39E-05 | -0.85816 | 0.00019828 |
| Ctsc | orange | 1.4320098 | 1.03E-05 | 1.26574736 | 0.00015109 |
| Nox4 | red | -1.1265059 | 0.00048079 | -1.0256491 | 0.00199972 |
| Tyrobp | yellow | 2.75623947 | 1.25E-17 | 2.48909016 | 2.18E-14 |
| 2200002D01 | orange | 0.99536074 | 1.04E-05 | 0.79485627 | 0.00072047 |
| Psmd8 | orange | 0.86419384 | 8.71E-08 | 0.75714264 | 4.85E-06 |
| Ryr1 | red | -0.7231578 | 0.00674464 | -0.3059664 | 0.30003162 |
| Nfkbib | orange | 1.21352856 | 8.76E-15 | 1.01857966 | 1.80E-10 |
| Psmc4 | orange | 0.88505651 | 8.26E-09 | 0.74310396 | 2.50E-06 |
| Mfge8 | red | -0.7769267 | 8.80E-12 | -0.666346 | 9.66E-09 |
| Aen | orange | 0.83833744 | 5.05E-07 | 0.75907589 | 9.61E-06 |
| Mrps11 | orange | 0.63053575 | 0.00220337 | 0.19884753 | 0.39268573 |
| Ccdc90b | red | -0.5986258 | 0.00085125 | -0.6209784 | 0.00075241 |
| Eed | orange | 0.70691731 | 1.61E-07 | 0.52369015 | 0.00019544 |
| Me3 | red | -1.22719 | 0.00484566 | -0.8213806 | 0.07514179 |
| Ddias | yellow | 2.74059337 | 4.84E-08 | 1.5228549 | 0.00283622 |
| Anapc15 | orange | 0.84977443 | 7.64E-06 | 1.01430685 | 2.06E-07 |
| Tmc5 | red | -1.3145099 | 0.00583603 | -0.4621644 | 0.37452102 |
| Xylt1 | orange | 1.42303394 | 3.23E-08 | 1.31626659 | 7.09E-07 |
| Calca | orange | 1.50875554 | 7.91E-05 | 1.26299794 | 0.00055603 |
| Mylpf | red | -1.452933 | 0.00010376 | -1.0241106 | 0.00889071 |
| Qprt | orange | 1.67989521 | 4.64E-05 | 1.71851172 | 8.30E-05 |
| Kif22 | yellow | 2.7132151 | 3.41E-31 | 2.99102754 | 4.71E-36 |
| Mvp | orange | 1.23158083 | 7.38E-14 | 1.15692431 | 3.67E-12 |
| Stard10 | red | -1.10143 | 1.55E-05 | -1.0204683 | 9.52E-05 |
| Ppp4c | orange | 0.75921203 | 3.38E-07 | 0.59664847 | 0.00010806 |
| Tbx6 | red | -1.3666544 | 3.35E-06 | -0.7973679 | 0.00843595 |
| Plekhb1 | skyblue | -1.9213134 | 8.63E-09 | -1.2334793 | 0.00038567 |
| Coro1a | yellow | 2.82304986 | 2.66E-44 | 2.74345689 | 9.21E-42 |
| Klk7 | skyblue | -1.8174651 | 0.00298746 | -1.2582319 | 0.05254871 |
| Nupr1 | orange | 0.79363102 | 1.62E-07 | 0.74486576 | 1.54E-06 |
| Cln3 | orange | 1.01215898 | 1.13E-08 | 0.97723175 | 6.38E-08 |
| Pold3 | orange | 0.89535485 | 1.06E-09 | 0.55784445 | 0.00027483 |
| Pgm2l1 | skyblue | -1.3294562 | 5.35E-05 | -1.2620498 | 0.00018811 |
| Syt3 | skyblue | -1.2857409 | 0.00321185 | -1.4852818 | 0.00086195 |
| Slco2b1 | red | -1.4520502 | 0.00019582 | -0.7937464 | 0.05592166 |
| Myh14 | red | -1.2422954 | 4.04E-06 | -0.7972822 | 0.00472948 |
| Lat | yellow | 2.67247289 | 8.17E-13 | 3.42834641 | 3.89E-20 |
| Il21r | yellow | 2.94239359 | 1.31E-17 | 3.58509302 | 4.29E-25 |
| Dgat2 | skyblue | -1.9071743 | 8.66E-10 | -1.7642625 | 2.48E-08 |
| Il4ra | orange | 1.77075085 | 1.63E-24 | 1.65828051 | 1.55E-21 |
| Nsmce1 | orange | 0.75580844 | 5.09E-13 | 0.66005242 | 6.75E-10 |

| | | | | | |
|---|---|---|---|---|---|
| Psma1 | orange | 0.9223171 | 6.83E-07 | 0.66296053 | 0.00060166 |
| Parva | red | -0.7077399 | 8.87E-08 | -0.5056353 | 0.00023041 |
| Dkk3 | red | -0.1725131 | 0.48450253 | -0.6339541 | 0.0056278 |
| Pak1 | orange | 0.606749 | 0.00374562 | 0.36170802 | 0.10795072 |
| Cox6a2 | skyblue | -1.173406 | 0.00288071 | -1.4316366 | 0.00034807 |
| Itgam | yellow | 3.45628272 | 2.68E-08 | 3.61967836 | 9.68E-09 |
| Lyve1 | orange | 1.00037951 | 4.59E-08 | 0.99194648 | 9.66E-08 |
| Rnf141 | red | -0.6411716 | 0.00011121 | -0.5272903 | 0.00211193 |
| Itgax | yellow | 2.23198302 | 9.44E-09 | 1.82166323 | 5.23E-06 |
| Adm | orange | 1.49933526 | 3.98E-09 | 1.45411113 | 1.22E-08 |
| Pycard | orange | 1.81297018 | 2.80E-23 | 1.48566072 | 2.35E-15 |
| Tead2 | red | -1.292317 | 5.83E-15 | -1.1956104 | 9.26E-13 |
| Cd37 | orange | 1.19686059 | 9.78E-08 | 1.20273378 | 1.45E-07 |
| Prss8 | red | -1.4287264 | 1.52E-06 | -0.9879301 | 0.00139985 |
| Itgal | yellow | 2.32730446 | 2.46E-26 | 2.23847868 | 2.52E-24 |
| Abcc6 | skyblue | -3.2333684 | 0.00012244 | -0.8263654 | 0.28318488 |
| Sergef | orange | 0.78903418 | 5.73E-05 | 0.65757736 | 0.00120855 |
| Rgs10 | orange | 0.98796618 | 6.34E-06 | 0.78939166 | 0.00054985 |
| Fgfr2 | red | -1.4333452 | 2.33E-06 | -0.7627288 | 0.01778265 |
| Cpxm2 | red | -1.2366901 | 6.95E-05 | -0.7097073 | 0.03026601 |
| Chp2 | skyblue | -2.2289654 | 4.04E-05 | -1.3929077 | 0.01234153 |
| Plk1 | yellow | 2.60109993 | 1.10E-26 | 2.64685092 | 5.90E-27 |
| Scnn1b | red | -0.9817028 | 0.00029701 | -0.5918982 | 0.03959877 |
| Vwa3a | red | -1.4403119 | 9.09E-07 | -0.9103303 | 0.00272695 |
| Hpx | orange | 0.76726668 | 0.01090471 | 1.00712396 | 0.00090151 |
| Tmem159 | red | -0.7998761 | 5.87E-06 | -0.6293069 | 0.00057036 |
| Trim30a | yellow | 2.78122713 | 9.33E-60 | 2.72232028 | 4.35E-57 |
| Lyrm1 | orange | 0.78258411 | 0.00677473 | 0.71755159 | 0.01679439 |
| 2610020H08 | orange | 0.89123469 | 0.00113728 | 0.57998551 | 0.04941739 |
| Acsm3 | skyblue | -1.4508873 | 0.0137203 | -1.6091172 | 0.00756147 |
| Trim21 | orange | 1.5403653 | 4.20E-40 | 1.65852959 | 1.76E-46 |
| Rrm1 | orange | 1.46930869 | 1.63E-25 | 1.19684668 | 5.46E-17 |
| Uros | red | -0.7757079 | 4.90E-07 | -0.6970997 | 1.39E-05 |
| Art1 | red | -1.26338 | 0.00323908 | -1.0031605 | 0.02536736 |
| Mki67 | yellow | 3.22656009 | 7.95E-44 | 3.66543711 | 3.78E-55 |
| Wee1 | red | -0.2835942 | 0.13404963 | -0.7323154 | 5.97E-05 |
| BC051019 | red | -1.1942614 | 8.87E-07 | -0.6776363 | 0.00742792 |
| St5 | red | -0.8138986 | 1.43E-05 | -0.5666098 | 0.00373984 |
| Stk33 | skyblue | -2.0790266 | 7.45E-11 | -1.5268526 | 2.20E-06 |
| Mrgprf | skyblue | -2.0902841 | 5.71E-17 | -1.0215482 | 7.15E-05 |
| Ano1 | red | -1.1251635 | 1.07E-05 | -0.654962 | 0.01512419 |
| Dock11 | orange | 1.37496964 | 6.61E-26 | 1.33783348 | 6.39E-24 |
| Tnni2 | red | -1.3228721 | 7.91E-05 | -0.863371 | 0.01428355 |

| | | | | | |
|---|---|---|---|---|---|
| Syt8 | orange | 1.23386206 | 0.00759078 | 1.51737427 | 0.00122065 |
| Smarca1 | skyblue | -1.51988 | 0.00267476 | -1.113198 | 0.03441917 |
| Sash3 | yellow | 2.09282611 | 6.96E-33 | 2.44817312 | 6.48E-44 |
| Elf4 | orange | 1.55818727 | 3.16E-13 | 1.47688257 | 8.62E-12 |
| Rbmx2 | orange | 1.02009477 | 2.84E-05 | 1.0022323 | 8.39E-05 |
| Igsf1 | skyblue | -1.9175551 | 0.00019595 | -1.8103812 | 0.00072153 |
| Stk26 | orange | 1.32152558 | 2.94E-10 | 0.79034763 | 0.00031269 |
| Cd40lg | yellow | 2.40159608 | 0.0003975 | 3.2936925 | 5.92E-06 |
| Syp | red | -1.1131508 | 4.91E-05 | -1.164957 | 3.87E-05 |
| Magix | skyblue | -3.3854487 | 2.95E-19 | -2.2487393 | 2.17E-09 |
| Pim2 | orange | 0.63579836 | 0.00427975 | 0.59607362 | 0.01021087 |
| Was | yellow | 2.62763893 | 2.21E-17 | 2.47241732 | 2.84E-15 |
| Rbm3 | orange | 1.9071651 | 3.56E-21 | 1.62253928 | 2.03E-15 |
| Porcn | red | -0.6032081 | 3.02E-09 | -0.6083792 | 3.00E-09 |
| Slc38a5 | skyblue | -2.2601748 | 4.84E-24 | -1.6050193 | 2.19E-12 |
| Rpgr | red | -0.8995619 | 0.00024632 | -0.7599938 | 0.00270343 |
| Dynlt3 | red | -0.7973188 | 4.81E-16 | -0.6399819 | 1.77E-10 |
| Heph | red | -1.1360628 | 5.23E-05 | -0.9972321 | 0.00055724 |
| Stard8 | red | -1.0091763 | 0.00026725 | -0.6415977 | 0.02802595 |
| Pbdc1 | orange | 0.78835376 | 0.00459827 | 0.58239293 | 0.04680769 |
| Magee1 | red | -0.9415124 | 9.32E-09 | -0.8044975 | 2.20E-06 |
| Fgf16 | skyblue | -1.7644531 | 0.00337777 | 0.0986804 | 0.89105193 |
| Srpx2 | red | -1.2627915 | 2.83E-07 | -0.9445347 | 0.00019478 |
| Sytl4 | red | -0.956414 | 2.77E-07 | -0.5565796 | 0.00424558 |
| Cenpi | yellow | 2.32456685 | 5.21E-09 | 2.6276115 | 8.36E-11 |
| Btk | orange | 1.59076388 | 1.92E-09 | 1.61078757 | 2.81E-09 |
| Gla | orange | 1.54925305 | 3.23E-08 | 1.41550415 | 7.95E-07 |
| Col4a6 | skyblue | -2.1807847 | 2.89E-16 | -1.0596083 | 0.00012492 |
| Col4a5 | red | -1.0515147 | 0.00015292 | -0.6374098 | 0.02989418 |
| Chrdl1 | skyblue | -2.0875533 | 4.00E-06 | -1.5161764 | 0.00123362 |
| Glt28d2 | red | -0.6798211 | 0.01789624 | -1.1154887 | 0.00011895 |
| Lrch2 | red | -1.3483082 | 7.52E-05 | -0.9616603 | 0.00519135 |
| Cdkl5 | skyblue | -1.9577097 | 3.05E-08 | -1.3619292 | 0.00020487 |
| D630029K05 | skyblue | -2.1407875 | 4.10E-07 | -1.6854674 | 0.00013693 |
| Phka2 | orange | 0.74267929 | 3.25E-11 | 0.51665286 | 7.87E-06 |
| Pdha1 | red | -0.6387063 | 0.00040675 | -1.405222 | 1.06E-15 |
| Il2rg | orange | 1.91084166 | 1.87E-20 | 1.63634344 | 4.80E-15 |
| Itgb1bp2 | red | -1.0449127 | 7.91E-06 | -0.8170545 | 0.00081829 |
| Gabre | red | -0.907691 | 0.00049424 | -0.344826 | 0.22519106 |
| Gabra3 | skyblue | -1.6872482 | 2.24E-16 | -0.6045066 | 0.00384251 |
| Gabrq | red | -1.7509221 | 0.00038054 | -0.553643 | 0.28440814 |
| Xlr4c | yellow | 2.9183785 | 3.03E-12 | 2.45558143 | 2.15E-09 |
| Haus7 | orange | 0.55879077 | 0.00048033 | 0.66878197 | 4.97E-05 |

| | | | | | |
|---|---|---|---|---|---|
| Bmx | skyblue | -1.9211784 | 1.48E-09 | -1.4319655 | 8.52E-06 |
| Pir | red | -1.2271144 | 3.81E-08 | -1.1746959 | 2.43E-07 |
| Asb11 | red | -0.7087071 | 0.0310905 | -0.9633904 | 0.00487556 |
| Arhgap4 | orange | 1.35983864 | 1.66E-17 | 1.67042826 | 1.18E-25 |
| Avpr2 | yellow | 2.23537208 | 2.85E-07 | 2.36568536 | 1.02E-07 |
| Plxna3 | red | -0.9187669 | 0.00014478 | -0.738999 | 0.00320631 |
| G6pdx | orange | 0.63088771 | 5.58E-05 | 0.54486181 | 0.00072664 |
| Dkc1 | orange | 0.61493549 | 0.00208876 | 0.31849132 | 0.14272206 |
| Plp1 | red | -1.1008132 | 2.51E-05 | -0.7078341 | 0.01031583 |
| Zcchc18 | orange | 0.98709014 | 0.00121389 | 0.85931431 | 0.00755825 |
| Psmd10 | orange | 0.71808422 | 0.00304984 | 0.32729247 | 0.22052191 |
| Morc4 | red | -1.4582941 | 9.49E-11 | -0.8886538 | 9.28E-05 |
| Mcf2l | skyblue | -1.2889566 | 5.85E-05 | -1.4881219 | 4.70E-06 |
| F7 | orange | 1.09001734 | 0.00017387 | 0.73397223 | 0.01586236 |
| F10 | yellow | 3.45394044 | 1.45E-17 | 2.89071866 | 1.32E-12 |
| Gas6 | red | -1.0130854 | 7.44E-06 | -0.5759525 | 0.01593696 |
| Rasa3 | orange | 0.55096125 | 2.21E-06 | 0.58358817 | 8.08E-07 |
| Myom2 | red | -0.968078 | 0.00212476 | -1.1330191 | 0.00040777 |
| Thsd1 | red | -0.9122229 | 8.63E-08 | -0.6763469 | 0.00012921 |
| Adrb3 | skyblue | -3.1779797 | 6.93E-06 | -2.2573705 | 0.00205646 |
| Cd209a | skyblue | -3.2661037 | 4.28E-13 | -2.7834739 | 1.26E-09 |
| Col4a1 | orange | 0.97828681 | 6.35E-07 | 1.10413756 | 2.66E-08 |
| Col4a2 | orange | 0.82058527 | 1.77E-05 | 0.9554523 | 7.91E-07 |
| Rab20 | yellow | 2.18635698 | 1.76E-19 | 1.97051916 | 8.29E-16 |
| Ptpn7 | yellow | 2.72097398 | 1.77E-25 | 3.01887067 | 1.19E-30 |
| Gpm6a | skyblue | -1.5768898 | 5.62E-09 | -1.3803542 | 6.50E-07 |
| Asb5 | red | -1.1561818 | 0.00147159 | -1.0139568 | 0.00839866 |
| Vegfc | red | -0.6267286 | 0.00661394 | -0.4918411 | 0.04112175 |
| Dlc1 | red | -0.9132913 | 6.25E-12 | -0.6625494 | 1.34E-06 |
| Eri1 | orange | 0.66391935 | 5.15E-05 | 0.68041665 | 5.10E-05 |
| Plat | orange | 1.55429128 | 6.52E-08 | 1.72308474 | 2.88E-09 |
| Ank1 | skyblue | -1.128475 | 0.00308466 | -1.6452378 | 2.03E-05 |
| Sfrp1 | yellow | 2.32075186 | 1.62E-15 | 1.84703525 | 5.59E-10 |
| Ido2 | yellow | 2.55400157 | 4.17E-06 | 2.15781256 | 9.17E-05 |
| Ido1 | black | 5.54105876 | 3.81E-124 | 5.95923996 | 2.70E-127 |
| Adam3 | skyblue | -1.3916658 | 0.00083033 | -1.6128222 | 0.00011894 |
| Adam5 | skyblue | -2.1056016 | 4.45E-05 | -2.2007477 | 4.42E-05 |
| Slit2 | red | -0.6714342 | 0.00057811 | -0.5276663 | 0.00936017 |
| Tenm3 | red | -1.1761093 | 2.08E-05 | -0.8170013 | 0.00581608 |
| Dctd | yellow | 2.11849346 | 8.92E-06 | 2.21716219 | 5.46E-06 |
| Mak16 | orange | 0.85670428 | 4.23E-06 | 0.65193818 | 0.00070867 |
| Wrn | orange | 0.59294728 | 0.00015928 | 0.44125593 | 0.0070598 |
| Gsr | orange | 0.84559436 | 4.84E-05 | 0.93912484 | 8.99E-06 |

| | | | | | |
|---|---|---|---|---|---|
| Fgl1 | yellow | 3.68068538 | 9.84E-06 | 1.95440127 | 0.00188057 |
| Galnt7 | orange | 0.9532752 | 1.34E-06 | 0.55474293 | 0.00771822 |
| Sap30 | orange | 1.46899408 | 3.05E-07 | 1.14998484 | 0.0001173 |
| Nr3c2 | red | -1.3106629 | 7.90E-06 | -1.0476164 | 0.00054834 |
| Irf2 | orange | 0.72214961 | 8.12E-12 | 0.52840759 | 1.23E-06 |
| Casp3 | orange | 1.93642801 | 1.60E-14 | 1.91046966 | 6.25E-14 |
| Cenpu | orange | 1.2986388 | 6.50E-05 | 1.41144062 | 1.83E-05 |
| Pdlim3 | red | -0.8939488 | 0.00025348 | -0.663358 | 0.00946768 |
| Tlr3 | orange | 1.02616668 | 4.75E-08 | 0.96661851 | 4.53E-07 |
| Nek1 | red | -0.9363282 | 6.54E-10 | -0.8792617 | 1.30E-08 |
| Mfap3l | red | -1.3836034 | 0.00024884 | -0.839157 | 0.03542594 |
| N4bp1 | orange | 1.27446652 | 1.63E-14 | 1.09022856 | 1.08E-10 |
| Adcy7 | orange | 0.5765469 | 0.00203345 | 0.61895981 | 0.00117852 |
| Snx20 | yellow | 2.75444599 | 3.94E-29 | 2.52406616 | 1.84E-24 |
| Aktip | skyblue | -1.4509383 | 1.25E-10 | -1.2117499 | 1.53E-07 |
| Gins3 | orange | 0.59913002 | 0.02398348 | 0.86889065 | 0.00148336 |
| Orc6 | orange | 0.592503 | 2.02E-05 | 0.37244306 | 0.01115821 |
| Itfg1 | red | -0.6654934 | 6.52E-15 | -0.6301653 | 4.85E-13 |
| Gab1 | red | -0.7228706 | 0.00050535 | -0.573264 | 0.00790424 |
| Hp | red | -0.7500038 | 0.00016325 | -0.1840984 | 0.42042746 |
| Ces1f | skyblue | -3.1565829 | 1.05E-09 | -2.1120321 | 6.86E-05 |
| Irx3 | red | -1.1850039 | 1.54E-07 | -0.6802871 | 0.00413949 |
| Crnde | skyblue | -1.5185253 | 1.56E-05 | -1.1018571 | 0.00235481 |
| Irx5 | skyblue | -1.5103381 | 6.99E-12 | -0.8952726 | 8.16E-05 |
| Il34 | red | -1.1411641 | 1.41E-06 | -0.9926438 | 4.51E-05 |
| Bbs2 | red | -1.1928103 | 4.02E-07 | -0.9813708 | 5.12E-05 |
| Cenpn | yellow | 2.23327941 | 2.16E-21 | 2.27910497 | 3.50E-21 |
| Mt4 | red | -1.1770014 | 0.00058445 | -0.8277723 | 0.02133633 |
| Cdyl2 | orange | 0.90881708 | 7.08E-05 | 1.14295241 | 1.30E-06 |
| Mt2 | yellow | 3.6175676 | 2.87E-20 | 3.21718201 | 1.25E-16 |
| Mt1 | yellow | 2.51496187 | 2.44E-44 | 2.13903694 | 2.40E-32 |
| Nudt7 | red | -0.9601881 | 5.23E-05 | -0.8127052 | 0.0008996 |
| Herpud1 | red | -0.9218837 | 0.00063226 | -0.8923571 | 0.0012624 |
| Pllp | red | -1.1541832 | 4.01E-05 | -1.0116725 | 0.0004375 |
| Ciapin1 | orange | 0.86821309 | 2.08E-08 | 0.71372001 | 7.72E-06 |
| Coq9 | red | -0.8818781 | 3.69E-11 | -0.7462528 | 4.66E-08 |
| Drc7 | skyblue | -2.0331614 | 8.80E-24 | -0.8315061 | 5.64E-05 |
| Katnb1 | orange | 0.68938214 | 3.61E-09 | 0.67281928 | 1.20E-08 |
| Cngb1 | skyblue | -1.7701323 | 0.00052362 | -1.6072885 | 0.00255518 |
| Mmp15 | skyblue | -1.8920782 | 4.02E-11 | -1.2126124 | 4.73E-05 |
| Tmem38a | red | -1.260192 | 2.04E-11 | -1.0950819 | 1.31E-08 |
| Usb1 | orange | 1.11096611 | 7.29E-09 | 1.06163666 | 5.34E-08 |
| B3gnt3 | orange | 0.86120316 | 2.26E-06 | 0.76645465 | 4.17E-05 |

| | | | | | |
|---|---|---|---|---|---|
| Jak3 | orange | 0.9794156 | 2.21E-12 | 0.774091 | 6.35E-08 |
| Slc27a1 | red | -0.5838657 | 0.00716466 | -0.7105102 | 0.00122749 |
| Mvb12a | orange | 0.73236879 | 6.52E-10 | 0.51159914 | 3.26E-05 |
| Gins2 | yellow | 2.3998873 | 1.03E-11 | 2.2555799 | 3.16E-10 |
| 6430548M08 | red | -0.8112944 | 3.23E-05 | -0.738407 | 0.00023193 |
| Crispld2 | orange | 0.666943 | 0.00093143 | 0.40248346 | 0.06042649 |
| Cotl1 | yellow | 2.79839074 | 3.36E-22 | 2.41256518 | 1.39E-16 |
| Dnaaf1 | skyblue | -2.101146 | 5.42E-18 | -1.172141 | 1.29E-06 |
| Pik3r2 | red | -0.6096796 | 1.52E-06 | -0.6466533 | 5.34E-07 |
| Ifi30 | yellow | 2.15413602 | 7.97E-41 | 1.66186882 | 1.52E-24 |
| Rab3a | skyblue | -1.484961 | 2.02E-08 | -1.219751 | 6.89E-06 |
| Cdh13 | skyblue | -1.652095 | 3.02E-07 | -1.0373386 | 0.00205857 |
| Mphosph6 | orange | 0.71765734 | 0.00018537 | 0.5941485 | 0.00296149 |
| 1700030J22F | red | -0.8723041 | 0.00604647 | -0.7614949 | 0.01589888 |
| Lsm4 | orange | 0.85480423 | 4.81E-06 | 0.7882298 | 4.18E-05 |
| Comp | skyblue | -1.2300033 | 0.0094267 | -1.9178388 | 7.11E-05 |
| Ntpcr | orange | 1.09510895 | 2.36E-05 | 0.79993758 | 0.00311105 |
| BC021891 | skyblue | -1.7074306 | 0.0003193 | -1.1425662 | 0.01661621 |
| Atp13a1 | orange | 0.75577943 | 7.91E-08 | 0.75396786 | 1.36E-07 |
| Pgr | skyblue | -1.3328766 | 0.02004688 | -1.5505035 | 0.00850312 |
| Cdh5 | red | -0.6149081 | 7.64E-05 | -0.4551237 | 0.00491752 |
| Bean1 | yellow | 2.19065411 | 1.79E-05 | 1.75992859 | 0.0006221 |
| Ces2g | skyblue | -1.6384592 | 3.02E-07 | -1.1197321 | 0.00076295 |
| Cdh16 | skyblue | -2.7625922 | 3.06E-06 | -1.5720275 | 0.010878 |
| Cbfb | orange | 0.80962881 | 8.96E-06 | 0.63697875 | 0.00075488 |
| Ces2e | skyblue | -2.4758385 | 2.13E-06 | -2.8915624 | 1.65E-07 |
| Tradd | orange | 0.76443337 | 1.09E-05 | 0.56955166 | 0.00170338 |
| D230025D16 | red | -0.980706 | 1.96E-05 | -0.6993072 | 0.00341653 |
| Hsd11b2 | orange | 1.73940096 | 0.00397175 | 0.02454015 | 0.97477362 |
| Tsnaxip1 | skyblue | -2.0242513 | 9.84E-12 | -1.1676427 | 0.00011264 |
| Ctrl | orange | 1.0123714 | 0.01188873 | 1.38009952 | 0.00087538 |
| Psmb10 | yellow | 2.23710606 | 5.13E-75 | 2.04840341 | 8.02E-63 |
| Dpep3 | yellow | 2.46912937 | 0.00013465 | 1.4846336 | 0.04379761 |
| Slc7a6 | orange | 0.59906091 | 5.27E-08 | 0.66025473 | 2.58E-09 |
| Smpd3 | red | -0.6341268 | 0.00890691 | -0.2388636 | 0.37414889 |
| Zfp90 | orange | 0.45027171 | 0.10487137 | 0.79906837 | 0.00370789 |
| Has3 | red | -1.4440686 | 6.02E-08 | -1.0174158 | 0.00016435 |
| Nip7 | orange | 0.76368702 | 0.00015283 | 0.68340063 | 0.00103577 |
| Mre11a | orange | 0.65245066 | 0.00012493 | 0.43823789 | 0.01461758 |
| Ankrd49 | orange | 0.58168429 | 0.00140733 | 0.40426702 | 0.03611018 |
| Izumo1r | yellow | 3.22747684 | 7.16E-16 | 2.94907489 | 4.99E-13 |
| Panx1 | orange | 1.52454692 | 2.16E-15 | 1.52679425 | 5.03E-15 |
| Vstm5 | orange | 1.00383657 | 0.00485213 | 0.41296265 | 0.28423471 |

| | | | | | |
|---|---|---|---|---|---|
| 4931406C07 | red | -1.0578704 | 2.53E-12 | -0.8083946 | 1.81E-07 |
| Taf1d | orange | 1.19297152 | 1.25E-12 | 1.04399324 | 1.25E-09 |
| Kars | orange | 1.04335053 | 1.21E-09 | 0.92153928 | 1.48E-07 |
| Tmem231 | skyblue | -1.7254597 | 1.17E-19 | -1.2231922 | 1.01E-10 |
| Ldhd | red | -0.8008078 | 0.00498938 | -0.6685144 | 0.02387961 |
| Aars | orange | 0.92470009 | 8.41E-10 | 0.93957478 | 7.04E-10 |
| Glb1l3 | skyblue | -3.0444105 | 3.59E-06 | -1.8706291 | 0.00460951 |
| Afg3l1 | orange | 0.58464111 | 0.00015884 | 0.50870421 | 0.00147099 |
| Acta1 | red | -1.1307921 | 0.00153315 | -0.6921598 | 0.06836149 |
| Urb2 | orange | 0.70142495 | 5.61E-05 | 0.96402878 | 5.05E-08 |
| Gnpat | red | -0.6661061 | 3.20E-08 | -0.4411841 | 0.00044436 |
| Egln1 | red | -0.6400563 | 2.63E-07 | -0.6132921 | 1.39E-06 |
| Jam3 | red | -0.9387214 | 7.21E-08 | -0.862511 | 1.69E-06 |
| Adamts8 | red | -1.0900059 | 8.62E-06 | -0.6977577 | 0.00647695 |
| Aplp2 | red | -1.1879659 | 8.88E-13 | -0.8840022 | 2.40E-07 |
| Trpc6 | red | -1.0275217 | 5.53E-14 | -0.904623 | 6.54E-11 |
| Birc3 | orange | 1.31123189 | 8.27E-27 | 1.14789907 | 9.97E-21 |
| Pdgfd | skyblue | -1.1857647 | 2.96E-13 | -1.2556964 | 1.50E-14 |
| Sesn3 | orange | 0.41349265 | 0.00507601 | 0.60196256 | 5.32E-05 |
| Usp2 | red | -1.0705383 | 5.03E-05 | -1.2450403 | 3.39E-06 |
| Thy1 | yellow | 2.66608224 | 3.21E-16 | 3.05153257 | 8.76E-21 |
| Ubash3b | yellow | 2.24669607 | 2.02E-21 | 1.8741648 | 5.46E-15 |
| Crtam | yellow | 4.54198276 | 1.33E-20 | 4.06980517 | 3.12E-17 |
| 4931429I11R | red | -0.9752041 | 0.00485892 | -0.5052124 | 0.14719645 |
| Nxpe2 | skyblue | -1.4594445 | 7.92E-06 | -1.2937395 | 0.00011465 |
| Barx2 | red | -0.9680851 | 0.00209699 | -0.7714101 | 0.01948904 |
| Kirrel3 | red | -1.4762985 | 0.00022857 | -0.1595447 | 0.73338212 |
| Abhd12 | orange | 0.7381072 | 0.00014188 | 0.61163399 | 0.0023379 |
| 4833427G06 | skyblue | -1.6025106 | 3.01E-08 | -1.0963059 | 0.00018071 |
| Cryab | red | -0.7785107 | 2.80E-06 | -0.5675276 | 0.00101965 |
| 2310030G06 | red | -0.6756747 | 0.00026367 | -0.3313729 | 0.09441347 |
| Dixdc1 | skyblue | -2.6377922 | 1.08E-10 | -1.6087174 | 0.00014449 |
| Plet1 | red | -1.0352646 | 4.68E-05 | -0.5781187 | 0.03117299 |
| Cadm1 | red | -1.4711483 | 4.46E-11 | -1.0607575 | 4.29E-06 |
| Bace1 | red | -0.7335607 | 4.56E-06 | -0.6321488 | 0.0001263 |
| Il10ra | yellow | 3.40486514 | 2.83E-35 | 2.91014657 | 6.51E-26 |
| Cd3e | yellow | 3.0336001 | 2.09E-24 | 3.16545561 | 7.57E-26 |
| Cd3d | yellow | 3.38090879 | 1.43E-24 | 3.71255432 | 1.83E-28 |
| Nlrx1 | orange | 0.68659813 | 3.75E-06 | 0.44679298 | 0.0040822 |
| Chek1 | orange | 1.89372488 | 1.73E-07 | 1.87113278 | 6.31E-07 |
| Fez1 | skyblue | -1.926269 | 2.20E-07 | -0.7825037 | 0.04623936 |
| Tmem218 | red | -0.7145238 | 8.70E-06 | -0.5549437 | 0.00089528 |
| Slc37a2 | yellow | 2.33146002 | 5.54E-10 | 1.99119612 | 2.21E-07 |

| | | | | | |
|---|---|---|---|---|---|
| Olfm2 | skyblue | -1.7200251 | 0.00038954 | -2.4190743 | 6.39E-07 |
| Icam5 | orange | 0.66767371 | 0.17240801 | 1.54860332 | 0.0014106 |
| Pde4a | red | -0.7266458 | 1.88E-05 | -0.7941633 | 4.16E-06 |
| Bmp5 | red | -1.0855314 | 1.07E-10 | -0.746736 | 1.95E-05 |
| Scg3 | skyblue | -1.4231912 | 0.00715687 | -1.3413811 | 0.01774919 |
| Gnb5 | red | -0.9486697 | 3.58E-06 | -0.7063245 | 0.00083177 |
| Ldlr | orange | 0.55858474 | 0.01045965 | 0.62862262 | 0.00457663 |
| Kank2 | red | -0.7887041 | 1.10E-05 | -0.6489005 | 0.00046612 |
| Dock6 | red | -1.0376633 | 8.14E-08 | -0.8400701 | 2.49E-05 |
| Aqp9 | orange | 0.98184377 | 0.01902973 | 1.54644262 | 0.00045838 |
| Nedd4 | red | -0.9121798 | 4.66E-12 | -0.7501193 | 2.75E-08 |
| Ccnb2 | yellow | 3.53827404 | 1.08E-57 | 3.82187596 | 4.54E-62 |
| Fam81a | skyblue | -2.3609717 | 9.36E-19 | -1.4583404 | 2.31E-08 |
| Anxa2 | orange | 0.65453919 | 0.00013574 | 0.6099261 | 0.00054599 |
| Cgnl1 | red | -0.8273988 | 2.44E-09 | -0.3775537 | 0.01046814 |
| Rora | red | -0.5906169 | 0.00022557 | -0.6973684 | 1.81E-05 |
| Itga11 | red | -0.8918707 | 0.00105568 | -0.8590605 | 0.00222899 |
| Calml4 | red | -1.5125995 | 8.73E-08 | -1.0697282 | 0.00022652 |
| Irak1bp1 | red | -0.7744803 | 0.00312808 | -0.2410603 | 0.41691565 |
| Kif23 | yellow | 1.88696375 | 1.92E-19 | 2.10990251 | 5.43E-23 |
| Lca5 | red | -1.2169324 | 1.45E-14 | -0.5319348 | 0.00078183 |
| Fam46a | orange | 1.48064958 | 1.40E-13 | 1.10648308 | 8.24E-08 |
| Tmprss5 | skyblue | -1.768821 | 2.65E-05 | -0.9797562 | 0.03154098 |
| Paqr5 | red | -0.9991589 | 0.00518763 | -1.0209354 | 0.00581608 |
| Tle3 | orange | 0.66720399 | 1.70E-07 | 0.88175329 | 4.33E-12 |
| Dnaja4 | skyblue | -1.3965756 | 1.52E-14 | -1.1757635 | 2.17E-10 |
| Thsd4 | red | -0.9257729 | 3.54E-06 | -0.5108483 | 0.01505384 |
| Pkm | orange | 1.25646211 | 8.14E-10 | 1.06889817 | 3.33E-07 |
| Commd4 | orange | 0.62332348 | 5.59E-09 | 0.58339521 | 1.02E-07 |
| 1700017B05I | orange | 1.96685196 | 1.22E-14 | 1.51149584 | 7.34E-09 |
| Psma4 | orange | 1.13305214 | 2.11E-12 | 0.90545716 | 4.64E-08 |
| Csk | orange | 0.61377057 | 2.69E-06 | 0.60375295 | 6.23E-06 |
| Cyp1a1 | skyblue | -4.5267367 | 1.70E-10 | -3.5212443 | 1.07E-06 |
| Rcn2 | red | -0.5850002 | 1.61E-08 | -0.4286731 | 6.78E-05 |
| Pstpip1 | yellow | 3.47880003 | 6.92E-43 | 3.46417261 | 3.11E-42 |
| Hmg20a | red | -0.5936537 | 0.00026617 | -0.457321 | 0.00696476 |
| Col12a1 | red | -1.1992567 | 9.83E-07 | -0.4866911 | 0.06683918 |
| Loxl1 | orange | 0.72139756 | 0.07212407 | 1.15495815 | 0.00367502 |
| Hcn4 | skyblue | -3.320111 | 9.13E-09 | -1.752245 | 0.00215364 |
| Mb21d1 | orange | 2.17342022 | 2.11E-16 | 1.34699638 | 6.39E-07 |
| Gsta4 | red | -0.5427126 | 0.00997901 | -0.6191567 | 0.00381496 |
| Elovl5 | red | -0.5900725 | 7.04E-05 | -0.5259254 | 0.00058521 |
| Lrrc1 | red | -0.9613273 | 1.32E-08 | -0.7287949 | 3.16E-05 |

| | | | | | |
|---|---|---|---|---|---|
| Mlip | red | -1.0916356 | 0.001685 | -1.1007521 | 0.00217891 |
| Tinag | skyblue | -2.2242875 | 8.66E-17 | -1.7321659 | 3.22E-10 |
| Ctsh | orange | 1.67369269 | 3.43E-11 | 1.40518558 | 5.36E-08 |
| Adamts7 | red | -0.6396894 | 0.00659256 | -0.8721921 | 0.00024236 |
| Tpm1 | red | -0.8267051 | 2.09E-08 | -0.5568651 | 0.00028888 |
| Lactb | orange | 0.70847456 | 7.40E-06 | 0.49266033 | 0.00313918 |
| Plod2 | red | -0.9387689 | 3.11E-07 | -0.9392375 | 5.12E-07 |
| Aph1b | red | -1.0612478 | 2.53E-07 | -0.9915159 | 2.16E-06 |
| Plscr4 | skyblue | -1.5021777 | 3.12E-13 | -1.3774305 | 4.37E-11 |
| Fam96a | orange | 0.87671856 | 6.77E-06 | 0.68776291 | 0.00069004 |
| Rbpms2 | red | -1.1556874 | 7.45E-10 | -0.8097827 | 2.70E-05 |
| Spg21 | orange | 0.66182529 | 4.82E-06 | 0.44035345 | 0.00368169 |
| Tipin | orange | 0.87888009 | 5.75E-06 | 0.77519681 | 0.00011167 |
| Zwilch | yellow | 2.3765427 | 4.50E-12 | 2.11979988 | 1.40E-09 |
| Lctl | skyblue | -3.1561161 | 0.00155038 | -2.9699365 | 0.0014556 |
| Me1 | red | -1.0760571 | 3.43E-07 | -1.1906702 | 2.35E-08 |
| Nt5e | orange | 0.71492618 | 0.0025666 | 0.7645877 | 0.00157538 |
| Snx14 | red | -0.6787387 | 0.00011165 | -0.5533306 | 0.00234014 |
| Zfp949 | red | -0.631915 | 0.00524655 | -0.391758 | 0.10527907 |
| Cmtm7 | orange | 1.7319061 | 2.48E-19 | 1.4284928 | 5.38E-13 |
| Stt3b | orange | 0.8182978 | 1.74E-13 | 0.40315269 | 0.00057499 |
| Eomes | yellow | 2.6279439 | 1.90E-20 | 2.08687484 | 3.70E-13 |
| Slc25a36 | red | -0.8138309 | 1.21E-07 | -0.5239428 | 0.00108997 |
| Clstn2 | skyblue | -1.5164355 | 1.23E-06 | -1.3608091 | 2.14E-05 |
| Mrps22 | orange | 0.70284875 | 5.86E-05 | 0.55591865 | 0.00214478 |
| Faim | orange | 0.96227732 | 1.19E-06 | 0.90957249 | 7.98E-06 |
| Dbr1 | orange | 0.77263558 | 1.46E-09 | 0.49461263 | 0.00020551 |
| Cldn18 | red | -0.9252993 | 9.81E-08 | -0.6180227 | 0.0006365 |
| Cdc25a | orange | 0.58783479 | 7.15E-05 | 0.74350313 | 7.73E-07 |
| Cspg5 | red | -1.4495384 | 0.00839235 | 0.17337116 | 0.80917766 |
| Ngp | yellow | 2.25956392 | 0.00608516 | 3.56924817 | 0.00081895 |
| Ptgs2 | orange | 0.6490943 | 0.00098867 | 0.80032409 | 3.70E-05 |
| Kif9 | red | -1.1933377 | 4.07E-09 | -0.9817305 | 1.87E-06 |
| Tdgf1 | orange | 1.41584236 | 0.00028178 | 0.95299118 | 0.01710863 |
| Lrrc2 | skyblue | -1.342158 | 1.96E-06 | -1.2842438 | 1.23E-05 |
| Dclk3 | skyblue | -1.7004452 | 4.85E-05 | -1.7273849 | 6.19E-05 |
| Trib1 | orange | 0.73894918 | 0.00026389 | 0.79689797 | 0.00011438 |
| Stac | skyblue | -2.7265641 | 2.54E-05 | -0.6848064 | 0.24943829 |
| Myd88 | orange | 2.02456214 | 1.16E-36 | 1.79678644 | 5.57E-29 |
| Scn5a | skyblue | -2.3280783 | 0.00140008 | -1.7957654 | 0.01787464 |
| Ttc21a | skyblue | -1.952902 | 2.26E-16 | -0.8440619 | 0.0007034 |
| Csrnp1 | yellow | 2.55000307 | 4.22E-30 | 2.00912715 | 6.19E-19 |
| Rpsa | orange | 1.02007767 | 1.64E-10 | 0.74948306 | 5.46E-06 |

| | | | | | |
|---|---|---|---|---|---|
| Hhatl | skyblue | -1.1383993 | 0.00216436 | -1.6763315 | 7.93E-06 |
| Nktr | orange | 0.68573133 | 2.62E-07 | 0.67164631 | 7.45E-07 |
| Pccb | red | -0.937737 | 3.11E-07 | -0.7727992 | 4.37E-05 |
| Vipr1 | red | -1.292996 | 0.00070486 | -0.9381801 | 0.01851602 |
| Ryk | red | -0.680327 | 8.31E-07 | -0.654124 | 3.66E-06 |
| Rab6b | skyblue | -1.6938094 | 2.72E-09 | -1.3721702 | 2.73E-06 |
| Trf | red | -0.9012384 | 0.00288705 | -0.6465827 | 0.04464584 |
| Topbp1 | orange | 0.93271367 | 1.83E-12 | 0.84366418 | 3.92E-10 |
| Dnajc13 | orange | 0.66278804 | 3.63E-07 | 0.648201 | 1.09E-06 |
| Acpp | red | -0.8653275 | 0.00187585 | -0.081048 | 0.81019753 |
| Aste1 | orange | 0.93638475 | 0.00014843 | 0.58993456 | 0.02299474 |
| Manf | orange | 0.931134 | 2.12E-07 | 0.63041023 | 0.00076168 |
| Mapkapk3 | orange | 1.1738594 | 4.13E-08 | 1.0347658 | 2.43E-06 |
| Cish | orange | 0.81750664 | 5.94E-08 | 0.49424585 | 0.0017023 |
| Traip | yellow | 2.28802275 | 1.69E-13 | 1.85507117 | 2.59E-09 |
| Bsn | orange | 1.25818868 | 0.00017652 | 1.09451564 | 0.00151286 |
| Mst1 | red | -0.9797815 | 0.0019968 | -1.0908549 | 0.00062469 |
| Cdhr4 | red | -1.349028 | 1.89E-08 | -0.5518431 | 0.02867905 |
| Uba7 | yellow | 2.14919849 | 2.43E-61 | 2.12771802 | 3.80E-60 |
| Nicn1 | red | -1.0455921 | 5.15E-14 | -1.0622222 | 3.05E-14 |
| Amt | red | -0.9402238 | 7.56E-06 | -0.9947852 | 3.68E-06 |
| Klhdc8b | skyblue | -1.6259075 | 1.08E-11 | -1.3045468 | 1.06E-07 |
| Pygm | red | -1.0730563 | 5.26E-05 | -0.840232 | 0.00227319 |
| Colgalt2 | skyblue | -2.8243163 | 4.94E-13 | -2.0848896 | 1.87E-07 |
| Crebl2 | red | -1.2976534 | 1.05E-05 | -0.9654835 | 0.00157335 |
| Oas3 | black | 6.53452424 | 1.38E-34 | 6.17445452 | 5.47E-31 |
| 1700025G04 | red | -0.7028376 | 0.00014574 | -0.603985 | 0.00159921 |
| Cd59a | skyblue | -1.5243768 | 8.70E-09 | -1.3912662 | 2.77E-07 |
| 6820408C15 | red | -0.8965704 | 6.36E-05 | -0.9274982 | 5.02E-05 |
| Malt1 | orange | 1.70215753 | 3.55E-23 | 1.39904863 | 9.04E-16 |
| Oas2 | yellow | 3.7181752 | 7.87E-39 | 3.82395303 | 6.53E-41 |
| Nlrp3 | yellow | 2.89768173 | 4.48E-14 | 2.78630425 | 5.92E-13 |
| Kank1 | red | -0.8346273 | 3.14E-06 | -0.575444 | 0.0020177 |
| Exd2 | red | -0.8251732 | 4.70E-17 | -0.7881001 | 1.04E-15 |
| 2810474O19 | orange | 0.64257647 | 0.00121257 | 0.59041915 | 0.00390452 |
| Trib3 | orange | 0.91064588 | 5.14E-05 | 0.82302897 | 0.00034986 |
| Mansc1 | red | -0.688093 | 3.87E-05 | -0.4397162 | 0.01168347 |
| Sbspon | red | -1.1639729 | 0.00442212 | -1.0153824 | 0.01061428 |
| Abtb2 | orange | 1.67016039 | 6.14E-14 | 1.56753651 | 3.14E-12 |
| Snx33 | red | -0.6306255 | 0.00019951 | -0.4952059 | 0.00484565 |
| Ablim3 | red | -0.9375992 | 3.44E-06 | -0.5561439 | 0.00888342 |
| Tpcn1 | red | -1.1982281 | 1.64E-09 | -1.0668529 | 1.43E-07 |
| Chrm1 | skyblue | -3.2886243 | 9.06E-10 | -2.6471305 | 4.66E-07 |

| | | | | | |
|---|---|---|---|---|---|
| Mctp2 | orange | 1.97973908 | 8.55E-10 | 1.69844666 | 2.30E-07 |
| Troap | yellow | 3.59814003 | 2.93E-35 | 4.2639286 | 5.78E-40 |
| Alas1 | red | -0.7223023 | 0.00075633 | -0.7116084 | 0.00120782 |
| Pdxk | red | -0.5620827 | 0.00041408 | -0.7562421 | 2.14E-06 |
| Lama1 | yellow | 2.17702001 | 4.64E-06 | 2.73601238 | 6.17E-08 |
| Srxn1 | orange | 0.69505156 | 0.02858374 | 0.95088889 | 0.00285183 |
| Fanca | orange | 1.30152658 | 2.39E-12 | 0.87068297 | 4.57E-06 |
| Igdcc4 | red | -1.3651654 | 3.68E-14 | -0.9743191 | 4.24E-07 |
| Ank2 | red | -0.9316028 | 0.00036478 | -0.7361329 | 0.00723169 |
| Ppp1r9a | red | -1.1245667 | 0.00024332 | -0.7148105 | 0.02707912 |
| Trpc1 | red | -1.0117557 | 6.90E-05 | -0.6765278 | 0.01087342 |
| Abcc4 | orange | 0.62534712 | 0.00205966 | 0.74019778 | 0.00034083 |
| Ugt8a | skyblue | -3.4643538 | 2.57E-07 | -1.384363 | 0.01173294 |
| Pkd1 | red | -0.6516685 | 0.00081149 | -0.550373 | 0.00629147 |
| P2ry2 | orange | 0.58594293 | 0.06005977 | 0.93468356 | 0.00251067 |
| Smap2 | orange | 0.92047157 | 3.37E-12 | 0.69348491 | 3.63E-07 |
| Arhgef17 | red | -0.8360736 | 3.15E-08 | -0.6127403 | 8.94E-05 |
| Ccdc85a | skyblue | -1.2322734 | 3.43E-07 | -1.2378681 | 4.64E-07 |
| Acsl3 | red | -1.1993665 | 3.56E-07 | -0.8558411 | 0.00046837 |
| Fbxo21 | red | -0.8899206 | 1.06E-07 | -0.6845323 | 7.50E-05 |
| Styk1 | yellow | 2.05531576 | 8.60E-10 | 2.1246276 | 5.30E-10 |
| Cspg4 | red | -0.8172552 | 1.80E-07 | -0.5786327 | 0.00036865 |
| Nup93 | orange | 1.00659559 | 5.35E-11 | 0.78447077 | 7.09E-07 |
| Rasgrp2 | red | -0.6310519 | 0.00859821 | -0.7280718 | 0.00284567 |
| Sox10 | skyblue | -1.5564999 | 5.40E-07 | -1.083703 | 0.00074131 |
| Polr2f | orange | 0.61030766 | 0.00089221 | 0.26497875 | 0.19010798 |
| Klrc3 | yellow | 2.5946192 | 0.00165314 | 0.85600518 | 0.30469946 |
| C330027C09 | yellow | 2.63651938 | 1.01E-25 | 2.42752881 | 3.30E-21 |
| Afap1l1 | red | -1.1409667 | 9.70E-10 | -0.8365615 | 1.44E-05 |
| Calhm2 | red | -0.5530833 | 0.00074592 | -0.7693616 | 3.89E-06 |
| Dhrs7c | skyblue | -2.2458543 | 3.37E-09 | -2.4888632 | 1.34E-10 |
| 1700028P14I | skyblue | -2.0049512 | 5.96E-09 | -1.7173285 | 3.21E-07 |
| Pygb | red | -0.6261453 | 0.00024334 | -0.6880788 | 7.58E-05 |
| Lmo7 | red | -0.5836273 | 5.07E-06 | -0.3694891 | 0.00588384 |
| Pfkm | skyblue | -1.4165196 | 3.38E-07 | -1.2432438 | 1.32E-05 |
| Clec1a | red | -1.1228104 | 8.53E-05 | -0.8947424 | 0.00251129 |
| Tbc1d4 | red | -0.3206532 | 0.14704545 | -0.6570519 | 0.00232903 |
| Triobp | orange | 0.68458601 | 1.06E-06 | 0.56926909 | 8.33E-05 |
| Nol12 | orange | 1.41319839 | 2.53E-09 | 1.14638627 | 2.74E-06 |
| Ybey | red | -0.6128455 | 0.00406246 | -0.8260191 | 0.00019223 |
| Phldb2 | red | -0.6611926 | 1.65E-07 | -0.3096409 | 0.02166238 |
| Podxl2 | skyblue | -1.9333539 | 2.22E-09 | -1.0512846 | 0.00202925 |
| Dis3 | orange | 0.66078009 | 0.00025592 | 0.54014216 | 0.00407392 |

| | | | | | |
|---|---|---|---|---|---|
| Card10 | red | -0.9541043 | 6.46E-09 | -0.7993728 | 2.10E-06 |
| Mgll | red | -0.9217689 | 1.60E-13 | -1.0726124 | 8.34E-18 |
| Tie1 | red | -1.1431519 | 6.41E-19 | -1.058415 | 3.39E-16 |
| Lpcat2 | orange | 1.6343695 | 1.12E-08 | 1.322676 | 7.72E-06 |
| Myh2 | skyblue | -3.9414964 | 0.0070713 | -2.9157449 | 0.05818557 |
| Mamdc2 | skyblue | -2.3681919 | 3.31E-16 | -2.0270773 | 5.77E-12 |
| S100b | skyblue | -2.227575 | 2.69E-07 | -1.8063338 | 0.00012869 |
| Ttc28 | red | -0.9123866 | 2.80E-05 | -0.6564848 | 0.00377402 |
| AA467197 | black | 8.22770642 | 2.14E-29 | 7.07526833 | 1.43E-23 |
| Eefsec | orange | 0.58826646 | 2.38E-05 | 0.25396485 | 0.09238624 |
| Rac2 | yellow | 2.84439823 | 1.62E-34 | 2.68453612 | 1.18E-30 |
| Ttf2 | orange | 1.03570517 | 7.54E-09 | 0.90218941 | 1.01E-06 |
| Trim45 | red | -0.9510279 | 1.27E-06 | -0.6645119 | 0.00129566 |
| Hsf4 | red | -1.0218287 | 5.99E-08 | -0.6645164 | 0.00067477 |
| Duox1 | red | -0.8036078 | 0.06198696 | -1.5435488 | 0.00032775 |
| Stk36 | red | -1.1983429 | 1.54E-05 | -0.9848586 | 0.00052257 |
| Ptprm | red | -1.2402788 | 8.21E-16 | -0.9743127 | 6.08E-10 |
| Rpgrip1l | red | -1.0297713 | 4.11E-08 | -0.4852709 | 0.01395711 |
| Noc4l | orange | 1.76371387 | 1.61E-23 | 1.57473469 | 8.20E-19 |
| Ptprf | red | -0.8836137 | 3.62E-05 | -0.4201351 | 0.06804784 |
| Mif | orange | 1.03868683 | 3.49E-12 | 0.73570596 | 1.92E-06 |
| Dpyd | red | -0.9182277 | 0.0006942 | -0.8909548 | 0.00136295 |
| Ctdp1 | orange | 0.7730023 | 1.80E-06 | 0.66109288 | 7.22E-05 |
| Tnxb | skyblue | -2.3099293 | 6.12E-23 | -1.7148633 | 7.15E-13 |
| Chst2 | red | -0.8287092 | 1.57E-07 | -0.545216 | 0.0008971 |
| Rtp4 | yellow | 3.09744982 | 5.23E-59 | 2.84429772 | 7.02E-50 |
| Pus7l | orange | 0.96028696 | 2.56E-05 | 0.81450794 | 0.00058755 |
| Palmd | red | -1.1764537 | 9.29E-07 | -0.80896 | 0.00121382 |
| Frrs1 | orange | 0.87610711 | 0.00021474 | 0.72404714 | 0.00324877 |
| Arhgap44 | red | -0.6238711 | 0.00063046 | -0.3069726 | 0.12008155 |
| Agl | red | -0.6465133 | 5.50E-06 | -0.4517735 | 0.00242445 |
| Snap91 | skyblue | -2.2955811 | 0.00204123 | -2.1986674 | 0.00379966 |
| Antxr1 | red | -1.2436205 | 9.11E-12 | -0.9764053 | 1.84E-07 |
| Mcee | red | -1.2658325 | 6.04E-07 | -1.0249446 | 9.45E-05 |
| Trmt13 | orange | 0.67891116 | 0.0025392 | 0.68115806 | 0.0028259 |
| Specc1l | red | -0.9697521 | 7.24E-09 | -0.6843306 | 8.25E-05 |
| Lpar6 | orange | 0.19505306 | 0.24875927 | 0.58517011 | 0.00032743 |
| Tagap | yellow | 2.53998338 | 1.79E-23 | 1.99658185 | 1.04E-14 |
| Adamts15 | orange | 1.76484952 | 2.56E-12 | 1.26899697 | 1.06E-06 |
| Armcx1 | red | -0.865953 | 1.28E-06 | -0.6254915 | 0.00075482 |
| Crlf2 | orange | 1.94784624 | 1.63E-16 | 1.7471271 | 2.85E-13 |
| Cysltr2 | orange | 2.05545666 | 6.56E-11 | 1.645854 | 3.82E-07 |
| Fndc3a | orange | 0.60435645 | 0.00020592 | 0.55625174 | 0.00089509 |

| | | | | | |
|---|---|---|---|---|---|
| BC026585 | red | -0.4720748 | 0.01349654 | -0.6132819 | 0.00154327 |
| Prss35 | skyblue | -1.8001118 | 2.62E-05 | -1.3844251 | 0.00175348 |
| Cdc14a | red | -0.7686671 | 2.96E-10 | -0.7044917 | 1.30E-08 |
| Asprv1 | red | -0.8492766 | 0.00746778 | -0.6280244 | 0.05992819 |
| Ppip5k1 | red | -0.6914868 | 2.06E-07 | -0.4687339 | 0.00072153 |
| Acsm1 | red | -1.3606664 | 5.57E-09 | -0.0613298 | 0.83151616 |
| Casp4 | yellow | 2.66544729 | 6.43E-32 | 2.50936731 | 2.87E-28 |
| Angptl1 | skyblue | -2.2431146 | 3.01E-05 | -2.5538275 | 1.70E-05 |
| Znrf1 | orange | 0.66860096 | 7.67E-08 | 0.82541289 | 3.65E-11 |
| Dph5 | orange | 0.98047387 | 0.00110631 | 1.22569431 | 5.91E-05 |
| Rbfox2 | red | -0.6298367 | 9.03E-17 | -0.4074429 | 2.26E-07 |
| Apol6 | yellow | 3.23047961 | 2.01E-32 | 3.43387359 | 2.53E-36 |
| Myo6 | red | -1.0899739 | 1.37E-06 | -0.8692183 | 0.00019537 |
| Tmem35 | skyblue | -2.9369421 | 1.92E-10 | -2.7945623 | 4.03E-09 |
| Igf2bp2 | orange | 1.0797884 | 6.08E-08 | 0.78328925 | 0.0001267 |
| Ndn | red | -1.3018856 | 6.20E-15 | -1.1207998 | 4.16E-11 |
| Myo5c | red | -1.3641824 | 6.55E-08 | -0.7439182 | 0.00511835 |
| Lgi3 | skyblue | -2.2612536 | 0.00074754 | -1.4926382 | 0.03480823 |
| Rfwd3 | orange | 0.65322236 | 1.08E-06 | 0.39515517 | 0.00501138 |
| Caskin1 | skyblue | -1.0449008 | 0.00634231 | -1.3227383 | 0.00071414 |
| Pank1 | skyblue | -1.9614877 | 2.86E-11 | -1.4673081 | 1.48E-06 |
| Cplx1 | skyblue | -0.9586148 | 0.13046927 | -1.8103522 | 0.0029951 |
| Map3k13 | red | -0.9722082 | 0.00088427 | -0.6977599 | 0.02149035 |
| Hacd3 | red | -0.5279137 | 3.86E-05 | -0.5884113 | 6.78E-06 |
| Piwil2 | yellow | 2.81255071 | 3.05E-07 | 2.74455803 | 8.17E-07 |
| Qsox1 | orange | 0.74216437 | 6.80E-09 | 0.43857932 | 0.00106716 |
| Ucp2 | orange | 1.3451984 | 1.46E-23 | 0.91405648 | 3.58E-11 |
| Stard9 | red | -0.824094 | 1.20E-05 | -0.6328685 | 0.00118085 |
| Adra2a | skyblue | -2.1268936 | 4.16E-06 | -0.500315 | 0.2953029 |
| Egr3 | orange | 1.32137773 | 2.94E-10 | 1.16479998 | 6.41E-08 |
| 3300002A11I | skyblue | -2.7345194 | 7.33E-09 | -1.4201091 | 0.00179464 |
| Recql4 | orange | 1.92739236 | 6.50E-15 | 1.9686942 | 1.61E-15 |
| Mtss1l | skyblue | -1.7497609 | 6.69E-18 | -1.2082132 | 7.44E-09 |
| Calm4 | red | -1.1052097 | 0.00640934 | -0.5761351 | 0.19026681 |
| Nrxn2 | red | -1.0180315 | 0.00378717 | -1.0426513 | 0.00380994 |
| Tlr13 | yellow | 4.11528315 | 4.16E-20 | 3.53687879 | 4.96E-15 |
| Dysf | orange | 0.69741644 | 0.00072501 | 0.72986335 | 0.0005499 |
| Atp7a | orange | 0.11425381 | 0.60252149 | 0.60271026 | 0.00225214 |
| Fam208b | orange | 0.54877224 | 0.00146145 | 0.85994906 | 5.95E-07 |
| Ephx4 | skyblue | -2.5843545 | 1.39E-05 | -1.0883511 | 0.06417172 |
| Pla2g4c | yellow | 1.9186633 | 0.0001503 | 3.67332888 | 2.72E-10 |
| B3galt2 | skyblue | -3.2639272 | 2.57E-06 | -1.3803141 | 0.05520079 |
| Fgg | yellow | 2.52277777 | 2.10E-05 | 2.04644504 | 0.00061409 |

| | | | | | |
|---|---|---|---|---|---|
| Klf9 | red | -0.6785916 | 0.00161655 | -0.9514241 | 1.09E-05 |
| Lgals3bp | yellow | 2.74146671 | 1.95E-27 | 2.60784771 | 7.76E-25 |
| Map9 | red | -1.1811342 | 1.26E-08 | -0.6329988 | 0.00294806 |
| Zdhhc15 | orange | 1.10124189 | 2.49E-05 | 0.85166185 | 0.00159407 |
| Gucy1a3 | skyblue | -1.4525622 | 4.39E-06 | -1.167396 | 0.00035714 |
| Gde1 | red | -0.7765379 | 7.21E-06 | -0.5724762 | 0.00143734 |
| Parl | orange | 0.69745578 | 0.00012156 | 0.54550603 | 0.00385057 |
| Rbm34 | orange | 0.55203761 | 0.01278652 | 0.61025727 | 0.00725959 |
| Zswim5 | red | -1.0890047 | 0.00030202 | -0.6688969 | 0.03457806 |
| Aspm | yellow | 2.90160433 | 1.43E-13 | 3.37704891 | 1.68E-16 |
| 9430020K01I | red | -0.5890914 | 5.68E-07 | -0.7732649 | 5.04E-11 |
| Slc16a2 | red | -1.1897048 | 5.33E-18 | -1.0175124 | 2.64E-13 |
| Cdkl4 | skyblue | -2.0850843 | 7.56E-09 | -1.1560993 | 0.00184297 |
| Rfc3 | orange | 1.61044779 | 1.66E-17 | 0.97863627 | 7.09E-07 |
| Dnah17 | orange | 1.44516939 | 8.77E-10 | 1.37859631 | 6.13E-09 |
| Fancd2 | yellow | 2.15623017 | 1.36E-10 | 1.80322085 | 2.34E-07 |
| Cct2 | orange | 0.67020725 | 4.41E-09 | 0.52261342 | 9.15E-06 |
| Cd226 | yellow | 2.42297366 | 5.39E-14 | 1.70716179 | 2.75E-07 |
| Ccdc17 | skyblue | -1.6071784 | 3.19E-07 | -1.0712226 | 0.00104037 |
| Fgd5 | red | -0.656434 | 9.90E-07 | -0.6300303 | 4.22E-06 |
| Wbscr17 | red | -1.3762756 | 6.03E-06 | -0.4706778 | 0.16216365 |
| Phka1 | red | -0.9573353 | 6.62E-07 | -0.5646976 | 0.00542364 |
| Farp2 | red | -1.2862902 | 3.83E-05 | -0.9669207 | 0.00260405 |
| Golim4 | red | -0.7675073 | 2.70E-17 | -0.5477092 | 4.30E-09 |
| Kctd7 | red | -1.149111 | 2.56E-12 | -0.6743905 | 9.05E-05 |
| Atp2c2 | red | -1.2956242 | 3.42E-11 | -0.6558064 | 0.00110477 |
| Vav1 | yellow | 3.25474405 | 6.33E-46 | 3.19254797 | 6.69E-44 |
| Bzrap1 | orange | 0.61836915 | 0.00056369 | 0.33603796 | 0.07902611 |
| Emid1 | red | -1.1567613 | 2.42E-05 | -1.0360391 | 0.00025288 |
| Faah | red | -0.6023909 | 2.59E-05 | -0.3211313 | 0.03508142 |
| Zbed5 | red | -0.0480676 | 0.88738247 | -0.9387176 | 0.00127994 |
| Rnf43 | red | -1.3668856 | 1.95E-07 | -0.7233431 | 0.00887716 |
| Lsm3 | orange | 0.70579904 | 9.64E-05 | 0.57036896 | 0.00256139 |
| Polq | yellow | 2.39460698 | 4.38E-20 | 2.54832845 | 1.72E-18 |
| Rasl10a | red | -0.7818193 | 0.02495469 | -0.9567578 | 0.00666795 |
| Ankmy1 | red | -1.3706605 | 0.00241428 | -0.5953486 | 0.21105521 |
| Rhov | orange | 1.62490137 | 6.12E-05 | 1.59368993 | 0.00013506 |
| Foxj1 | red | -1.222248 | 2.63E-07 | -0.4151788 | 0.11065705 |
| Hdac11 | skyblue | -1.9164432 | 1.95E-11 | -1.6225303 | 2.39E-08 |
| Mfsd7c | red | -0.5275349 | 0.02099675 | -0.6725433 | 0.00350401 |
| Vwa9 | orange | 0.61644089 | 2.77E-05 | 0.4354807 | 0.00455426 |
| Zdhhc14 | skyblue | -1.7450674 | 3.10E-16 | -0.8660464 | 0.00011115 |
| Batf | yellow | 3.23293197 | 9.88E-32 | 2.96834104 | 1.22E-26 |

| | | | | | |
|---|---|---|---|---|---|
| Dnajc17 | orange | 0.83853054 | 5.36E-06 | 0.67348485 | 0.00040283 |
| Traf3ip1 | red | -1.1417017 | 5.38E-12 | -0.6623302 | 0.00012361 |
| Fhod3 | skyblue | -1.9992836 | 5.07E-06 | -2.1169592 | 2.24E-06 |
| Sdr42e1 | red | -0.6967447 | 0.00855809 | -0.5305983 | 0.0564671 |
| Kif4 | yellow | 2.28061477 | 8.89E-14 | 2.93389589 | 1.17E-20 |
| Iqsec1 | red | -0.588065 | 9.12E-05 | -0.5748248 | 0.00018786 |
| Trim59 | orange | 1.62623011 | 1.67E-10 | 1.62589778 | 1.07E-09 |
| Brip1 | yellow | 2.20261054 | 5.68E-08 | 2.02830655 | 1.31E-06 |
| Plcg2 | orange | 1.4173063 | 3.35E-08 | 1.18574116 | 7.03E-06 |
| Zbed4 | orange | 0.61245534 | 0.00033508 | 0.59588332 | 0.00066323 |
| Fam151b | red | -0.7230646 | 0.0024448 | -0.6097824 | 0.01398177 |
| Ube2f | orange | 0.58836994 | 1.13E-07 | 0.32398591 | 0.00579221 |
| Smc4 | orange | 0.89319955 | 1.71E-07 | 0.64167318 | 0.00031414 |
| Ramp1 | red | -1.3677228 | 0.0001075 | -0.7183586 | 0.05782609 |
| Cpne2 | orange | 1.89901376 | 1.99E-10 | 1.72623925 | 1.40E-08 |
| AI661453 | red | -0.7706076 | 1.23E-05 | -0.3745058 | 0.04715928 |
| Ssu2 | skyblue | -3.8332086 | 0.00031801 | -3.066229 | 0.00080643 |
| Lif | yellow | 2.34997227 | 9.42E-12 | 3.09934471 | 1.34E-19 |
| Neurl1b | red | -0.6734756 | 3.24E-07 | -0.6159086 | 5.94E-06 |
| Pkd1l2 | orange | 1.88999276 | 0.00626105 | 1.99716751 | 0.00386196 |
| Parp14 | yellow | 2.34376412 | 3.47E-143 | 2.38037313 | 1.55E-147 |
| Tmem30b | red | -0.7540663 | 7.64E-05 | -0.2046625 | 0.34120337 |
| Gbp8 | yellow | 3.66239242 | 6.08E-63 | 3.88690214 | 9.31E-71 |
| Ifit1 | yellow | 4.0171973 | 9.82E-107 | 4.01138642 | 2.75E-106 |
| Six4 | skyblue | -2.3915804 | 2.30E-08 | -1.5834851 | 0.0003401 |
| Pkd2 | red | -0.6556907 | 4.78E-07 | -0.5589991 | 2.96E-05 |
| Scara3 | red | -1.1310401 | 8.39E-08 | -0.3130425 | 0.17846891 |
| Dynlrb2 | skyblue | -1.93857 | 1.34E-20 | -1.3027636 | 8.28E-10 |
| Rasd2 | skyblue | -1.6935971 | 1.24E-05 | -1.0644048 | 0.00831528 |
| Snx2 | orange | 0.58120173 | 1.02E-05 | 0.45659424 | 0.00082954 |
| Kdelc2 | red | -0.9113445 | 1.00E-10 | -0.7873131 | 4.93E-08 |
| Edil3 | skyblue | -1.8913957 | 9.50E-12 | -1.061856 | 0.00022498 |
| Pcnxl4 | red | -0.7189244 | 1.23E-09 | -0.3637689 | 0.00327445 |
| Mad2l1bp | orange | 0.62377567 | 0.00017168 | 0.52862703 | 0.00221382 |
| Gjc1 | red | -0.6217434 | 3.66E-08 | -0.5897548 | 2.97E-07 |
| Zfp395 | red | -1.0466744 | 2.78E-06 | -0.845933 | 0.00024828 |
| Zfyve9 | red | -0.5929784 | 7.77E-07 | -0.5980518 | 1.04E-06 |
| Inpp5j | orange | 1.84327408 | 7.57E-05 | 1.33176676 | 0.00597391 |
| Papd7 | orange | 0.61638349 | 1.91E-07 | 0.56168877 | 3.43E-06 |
| Exph5 | red | -1.0983751 | 0.00918975 | -0.5671997 | 0.21515782 |
| Hid1 | red | -0.7052653 | 3.71E-05 | -0.2900525 | 0.11961776 |
| Slc41a2 | orange | 0.66173101 | 0.00638183 | 0.55410843 | 0.02969249 |
| Myo5a | orange | 1.03489954 | 1.85E-09 | 0.69354668 | 0.00011534 |

| | | | | | |
|---|---|---|---|---|---|
| Ppp1r18 | orange | 1.61008537 | 5.18E-16 | 1.43182427 | 1.09E-12 |
| Ppm1h | orange | 0.60970414 | 8.50E-05 | 0.57187799 | 0.00034581 |
| Setmar | red | -0.3254702 | 0.18997683 | -0.753047 | 0.00166074 |
| Cd300ld | orange | 1.53792007 | 0.00153329 | 1.55765884 | 0.00172833 |
| Ankrd12 | red | -0.4994504 | 0.01329138 | -0.5975787 | 0.00350645 |
| Lrrn1 | red | -1.2043715 | 8.58E-05 | -0.8696113 | 0.01173637 |
| Cd300a | yellow | 2.65926006 | 3.31E-13 | 2.3437214 | 2.29E-10 |
| Ythdc2 | orange | 0.38372108 | 0.02376042 | 0.58843523 | 0.00055017 |
| Cacna1a | red | -1.4432099 | 3.32E-06 | -0.9716885 | 0.0028956 |
| Tmem109 | red | -0.7257082 | 1.06E-11 | -0.6543555 | 1.73E-09 |
| Bmp2k | orange | 0.85233285 | 4.65E-08 | 0.77144033 | 1.43E-06 |
| Itga2b | orange | 0.77585204 | 4.65E-06 | 0.18505512 | 0.33893427 |
| Tdg | orange | 0.60973077 | 6.87E-06 | 0.6737723 | 1.11E-06 |
| Dbn1 | orange | 0.67684588 | 0.05101389 | 0.97525197 | 0.00493796 |
| Rnps1 | orange | 0.76897374 | 8.92E-06 | 0.6861092 | 0.00011884 |
| Fam171a2 | skyblue | -1.6851356 | 1.63E-12 | -1.4984213 | 8.28E-10 |
| Prr7 | orange | 1.4694751 | 2.10E-05 | 1.48792384 | 4.88E-05 |
| Fras1 | skyblue | -1.2582112 | 0.00012051 | -1.6396928 | 1.68E-06 |
| Dnaic2 | skyblue | -1.8259991 | 3.07E-16 | -1.1360274 | 6.02E-07 |
| Gns | orange | 0.8140009 | 9.91E-05 | 0.4999459 | 0.02354956 |
| Grn | orange | 1.74171721 | 1.00E-11 | 1.37669703 | 1.60E-07 |
| Adgrb1 | orange | 1.13245916 | 0.01884764 | 1.32962468 | 0.00566145 |
| Tle6 | red | -0.8805994 | 4.69E-05 | -0.9455233 | 2.11E-05 |
| Map4k5 | red | -0.6277734 | 7.38E-07 | -0.3742544 | 0.0047989 |
| Dusp5 | yellow | 1.95058904 | 6.73E-21 | 2.05539475 | 7.81E-23 |
| Asb16 | red | -1.4455616 | 2.09E-05 | -1.0339725 | 0.00467293 |
| Tle2 | skyblue | -1.6377313 | 4.00E-08 | -1.1321257 | 0.00025616 |
| BC030867 | yellow | 3.3576637 | 1.84E-21 | 3.61368565 | 4.69E-20 |
| B3galt1 | red | -1.0038166 | 0.00299828 | -0.3847266 | 0.29745209 |
| Gpsm3 | orange | 1.44001528 | 1.71E-11 | 1.31766772 | 1.61E-09 |
| Gna15 | orange | 1.54312303 | 1.90E-15 | 1.39399597 | 2.41E-12 |
| Cpne7 | orange | 1.3444046 | 0.00144411 | 0.68887131 | 0.13605879 |
| Unc13a | red | -0.988006 | 8.17E-07 | -0.8062804 | 0.0001016 |
| Scn7a | skyblue | -2.1499921 | 7.66E-15 | -1.8140229 | 1.19E-10 |
| Celf5 | skyblue | -1.0128787 | 0.09764513 | -1.6885541 | 0.00827518 |
| Nup54 | orange | 0.68361417 | 7.64E-06 | 0.47053508 | 0.00320683 |
| Tespa1 | orange | 1.17934581 | 0.00160615 | 1.80789599 | 2.84E-06 |
| Larp6 | red | -1.1025255 | 1.96E-05 | -0.2125906 | 0.48282081 |
| Mfsd12 | orange | 0.67025101 | 0.00401449 | 0.6072424 | 0.0115224 |
| Cxcl10 | black | 8.02361014 | 7.63E-119 | 8.03714332 | 1.25E-119 |
| Fam214a | red | -0.7306172 | 0.00418136 | -0.7656364 | 0.00333181 |
| Gipc3 | red | -0.78849 | 4.78E-07 | -0.7829898 | 1.03E-06 |
| Nudt19 | orange | 0.65829429 | 4.56E-06 | 0.32306858 | 0.0380084 |

| | | | | | |
|---|---|---|---|---|---|
| Tbxa2r | red | -1.3910243 | 1.16E-05 | -0.7335935 | 0.02993191 |
| Lrr1 | yellow | 3.16602259 | 1.89E-08 | 4.2980019 | 6.24E-08 |
| Cobll1 | red | -0.6225862 | 0.00271382 | -0.5699569 | 0.00772179 |
| Ncaph | orange | 1.79605102 | 8.72E-15 | 1.88840206 | 8.95E-16 |
| Pygo1 | skyblue | -1.3915196 | 5.50E-10 | -1.2688237 | 2.99E-08 |
| Ushbp1 | skyblue | -1.7725901 | 7.86E-16 | -1.4446113 | 1.19E-10 |
| Rtkn | red | -0.9191562 | 2.50E-08 | -0.6322649 | 0.00020885 |
| Mrpl54 | orange | 1.01889541 | 1.43E-08 | 0.73064069 | 9.71E-05 |
| Tmem106a | yellow | 2.28425501 | 9.34E-18 | 2.04611007 | 2.89E-14 |
| Zfr2 | red | -1.1961573 | 3.07E-10 | -0.8147319 | 2.54E-05 |
| Cebpa | red | -0.6588205 | 6.40E-09 | -0.434397 | 0.00024129 |
| Atcay | orange | 0.49472517 | 0.18801588 | 1.04477729 | 0.00533378 |
| 5031414D18 | yellow | 2.01242413 | 1.39E-08 | 2.06529845 | 1.28E-08 |
| Lbx2 | red | -1.0832463 | 0.0076482 | -0.4943964 | 0.23695094 |
| Dapk3 | orange | 0.61106251 | 1.65E-05 | 0.6057117 | 3.15E-05 |
| Parm1 | red | -0.9712402 | 2.45E-05 | -0.7476572 | 0.0016512 |
| Hrh2 | yellow | 3.69871017 | 5.01E-24 | 3.96660387 | 3.33E-27 |
| Dpp4 | red | -1.2292395 | 2.23E-11 | -1.0195681 | 5.92E-08 |
| Igsf6 | yellow | 2.94704009 | 5.78E-11 | 2.73066296 | 2.24E-09 |
| Baz1a | orange | 1.66811739 | 9.80E-27 | 1.4238283 | 1.60E-19 |
| Nek11 | skyblue | -2.0455361 | 2.94E-11 | -1.0366111 | 0.00037936 |
| Ccl5 | yellow | 3.29421841 | 8.21E-38 | 3.20334408 | 3.41E-35 |
| Kri1 | orange | 0.66106395 | 0.0012031 | 0.63617877 | 0.00245725 |
| Rrp12 | orange | 1.13280204 | 2.27E-08 | 1.31190061 | 1.31E-10 |
| Eef2k | red | -0.9298249 | 3.47E-08 | -0.7516368 | 1.48E-05 |
| Xkr6 | red | -0.9754337 | 1.95E-05 | -0.4986242 | 0.03474257 |
| 1700020L24F | skyblue | -1.1871385 | 0.00080514 | -1.2668782 | 0.00047536 |
| Dcbld2 | red | -0.8003774 | 1.84E-06 | -0.5510186 | 0.00160274 |
| Wnk4 | red | -0.6350693 | 0.00100506 | -0.4965815 | 0.01353065 |
| Wdr78 | skyblue | -1.8858652 | 1.26E-11 | -1.1120251 | 9.88E-05 |
| Gpr33 | yellow | 3.00819293 | 0.00376113 | 1.18797124 | 0.25193381 |
| Kcne3 | red | -1.4784365 | 0.00316798 | -0.6939726 | 0.19631309 |
| Ppp1r32 | red | -1.5332081 | 4.19E-08 | -0.737312 | 0.00924933 |
| Ubd | black | 5.23404096 | 1.36E-36 | 5.6865258 | 3.94E-36 |
| Nkx6-1 | red | -1.2351237 | 0.00220246 | -0.6262335 | 0.16142154 |
| Ano4 | skyblue | -2.092211 | 0.02743899 | -3.3964536 | 0.00142 |
| Slfn8 | black | 5.03744774 | 6.55E-30 | 4.32552851 | 9.29E-25 |
| Lsm7 | orange | 0.64670562 | 2.39E-05 | 0.30866583 | 0.06358522 |
| Rims4 | skyblue | -2.7346204 | 0.00515371 | -1.7041132 | 0.06579209 |
| Pdk3 | orange | 0.5957197 | 7.78E-07 | 0.6003876 | 1.18E-06 |
| Eogt | red | -0.5901089 | 3.98E-05 | -0.4853134 | 0.00106457 |
| Abi3bp | skyblue | -2.5059799 | 1.79E-12 | -1.6478724 | 7.62E-06 |
| Pkig | orange | 0.61082086 | 0.00048637 | 0.34142324 | 0.06838406 |

| | | | | | |
|---|---|---|---|---|---|
| Hpse | orange | 1.54289041 | 1.94E-09 | 1.30182872 | 8.29E-07 |
| Tpbg | red | -1.0828875 | 0.00622263 | -0.380594 | 0.38758988 |
| Raver2 | red | -1.0365919 | 0.00012759 | -0.4954243 | 0.08909016 |
| Plekhj1 | orange | 0.60796727 | 0.00015292 | 0.49186606 | 0.00328481 |
| Adrb1 | skyblue | -1.5494588 | 1.29E-16 | -0.8745703 | 7.47E-06 |
| Nat14 | red | -0.9239141 | 1.62E-07 | -0.8831566 | 2.04E-06 |
| Sgcg | skyblue | -1.3117527 | 7.72E-05 | -1.3888822 | 4.60E-05 |
| Mid1 | orange | 0.93918697 | 4.33E-07 | 1.34550251 | 1.45E-13 |
| Gnptab | orange | 0.85008782 | 1.26E-05 | 0.64996011 | 0.00128388 |
| Nup37 | orange | 0.92251297 | 2.55E-05 | 0.81340692 | 0.00037841 |
| Ccl12 | black | 5.91632869 | 2.91E-15 | 5.44722767 | 5.53E-13 |
| Kcnh4 | orange | 2.05701442 | 0.00011534 | 1.73286574 | 0.00160909 |
| Nfkbiz | orange | 1.41328322 | 1.33E-17 | 1.326793 | 1.55E-15 |
| Parpbp | yellow | 3.86205775 | 5.45E-13 | 4.10028015 | 5.21E-12 |
| Ccl7 | black | 7.52306965 | 5.43E-67 | 7.69567528 | 1.22E-64 |
| Ccl2 | black | 7.09275848 | 3.39E-31 | 6.80610205 | 3.80E-29 |
| Dennd1a | orange | 0.61759515 | 0.00013954 | 0.39561454 | 0.02053066 |
| Cfap53 | red | -0.7819815 | 0.00239717 | -0.310218 | 0.27758012 |
| Kank4 | skyblue | -2.6847471 | 7.21E-06 | -1.6860402 | 0.00708105 |
| Tmem98 | red | -0.616312 | 0.0001874 | -0.6154209 | 0.000276 |
| Ccr3 | black | 3.70617384 | 0.00604517 | 5.58368807 | 0.00018125 |
| Foxa1 | red | -0.9921894 | 2.55E-12 | -0.3823393 | 0.01083305 |
| Fignl1 | yellow | 2.74680273 | 1.23E-12 | 3.14930171 | 2.25E-15 |
| Prdm8 | red | -1.5033987 | 2.20E-08 | -0.9578721 | 0.00062314 |
| Tnni3 | skyblue | -1.3049753 | 0.05651485 | -1.9295676 | 0.00468641 |
| Galm | red | -0.6033775 | 0.00697519 | -0.6077508 | 0.00825433 |
| Plk5 | skyblue | -3.0850237 | 4.52E-13 | -1.9405657 | 1.65E-06 |
| Tgfbi | yellow | 3.73624135 | 9.23E-22 | 3.26047923 | 1.19E-16 |
| Reep6 | skyblue | -1.6529534 | 1.78E-08 | -1.1872167 | 7.77E-05 |
| Cox18 | orange | 1.13602426 | 1.35E-10 | 0.82385675 | 5.77E-06 |
| Tdrd7 | orange | 0.85365594 | 1.03E-10 | 0.69251417 | 3.31E-07 |
| Ccdc180 | skyblue | -1.9337699 | 2.40E-19 | -0.9906531 | 6.88E-06 |
| Krt17 | yellow | 2.08718509 | 8.57E-08 | 2.33532586 | 3.48E-09 |
| Aldh1b1 | orange | 1.20944695 | 0.00189309 | 0.37169301 | 0.3992404 |
| Iqcg | skyblue | -1.7813452 | 2.98E-12 | -1.1671833 | 6.76E-06 |
| Ky | skyblue | -3.0022915 | 1.36E-12 | -2.0871111 | 1.62E-06 |
| Frmpd1 | red | -1.4654081 | 1.96E-06 | -1.0751858 | 0.00066834 |
| Grhpr | red | -0.6729185 | 3.42E-05 | -0.6783828 | 4.17E-05 |
| Cbarp | red | -0.6153284 | 0.00275976 | -0.3307983 | 0.1337142 |
| Aamdc | red | -0.805053 | 0.00012297 | -0.8155422 | 0.00016289 |
| Zswim4 | orange | 0.47295871 | 0.00048133 | 0.62135567 | 4.75E-06 |
| Sbno2 | yellow | 2.02200391 | 3.64E-24 | 2.0591496 | 5.70E-25 |
| Tnfsf9 | yellow | 2.13013639 | 1.22E-11 | 1.7761657 | 4.56E-08 |

| | | | | | |
|---|---|---|---|---|---|
| Melk | yellow | 2.7227919 | 7.42E-25 | 2.37040898 | 8.96E-19 |
| Thrsp | skyblue | -3.5154841 | 3.20E-05 | -2.7046538 | 0.00199196 |
| Isg15 | black | 5.35382118 | 1.44E-119 | 5.21394305 | 3.01E-113 |
| Caps2 | red | -1.0498005 | 0.02951245 | -1.2722815 | 0.00904504 |
| Hmha1 | orange | 1.17157167 | 1.33E-31 | 1.33537077 | 1.12E-40 |
| Alg8 | orange | 1.03508881 | 1.78E-08 | 0.89858989 | 1.71E-06 |
| Dok3 | yellow | 2.30263519 | 4.28E-13 | 2.05983413 | 1.81E-10 |
| Dagla | red | -0.9262005 | 6.02E-11 | -0.6621486 | 5.79E-06 |
| Bbs10 | red | -0.8639612 | 0.00491901 | -0.536032 | 0.09607135 |
| Krt20 | yellow | 2.66263816 | 0.01633908 | 4.07152039 | 0.00037353 |
| Cd99l2 | red | -0.9337425 | 2.62E-12 | -0.7288543 | 1.10E-07 |
| Acta2 | red | -0.9135144 | 3.54E-06 | -0.1601057 | 0.48976545 |
| Mlc1 | red | -0.9074926 | 0.00324382 | -0.5722223 | 0.08106271 |
| Plekhs1 | yellow | 2.90498851 | 2.16E-05 | 3.09899186 | 6.38E-06 |
| Pim3 | orange | 0.60482825 | 0.00169656 | 0.40187313 | 0.04843531 |
| Ppp1r26 | red | -1.0183427 | 1.49E-07 | -0.9548402 | 1.31E-06 |
| Polr3g | red | -0.4823545 | 0.05813849 | -0.9554467 | 0.00014422 |
| Ddx11 | orange | 0.85115441 | 8.46E-07 | 0.6995439 | 9.85E-05 |
| Ids | red | -1.0723326 | 5.94E-09 | -0.7990053 | 2.70E-05 |
| Misp | yellow | 2.5720257 | 7.10E-34 | 2.38885437 | 8.68E-30 |
| Cdhr3 | skyblue | -2.7767469 | 2.16E-16 | -1.6213954 | 3.36E-06 |
| Syt1 | skyblue | -2.3746655 | 0.0017679 | -1.7728237 | 0.02129217 |
| 3110052M02 | red | -0.8071075 | 0.00601373 | -0.5331031 | 0.08101599 |
| Zhx3 | red | -0.6156973 | 0.0008615 | -0.6730899 | 0.00035239 |
| Hykk | skyblue | -1.6124293 | 0.0001033 | -1.2945679 | 0.00250068 |
| Rnf126 | orange | 0.64467988 | 6.43E-05 | 0.57784308 | 0.00050839 |
| Cerk | orange | 0.8140182 | 6.63E-08 | 0.91473282 | 1.73E-09 |
| Gramd4 | orange | 0.64349255 | 0.00092726 | 0.67567576 | 0.00066824 |
| Dcdc2a | skyblue | -2.4920395 | 1.23E-07 | -1.7294564 | 0.0003128 |
| Cd276 | orange | 1.34313915 | 0.00014156 | 1.2927967 | 0.0003607 |
| Bbs9 | red | -0.7493154 | 0.00138167 | -0.7163085 | 0.00284816 |
| Myf6 | skyblue | -1.7332153 | 1.29E-06 | -0.7124337 | 0.08118936 |
| H2-Q4 | yellow | 2.43344273 | 1.76E-47 | 2.24343903 | 2.17E-40 |
| Chst4 | skyblue | -2.0381549 | 5.39E-06 | -2.1154128 | 4.95E-06 |
| Pknox2 | red | -0.8203398 | 9.91E-05 | -0.8622455 | 5.81E-05 |
| Aldh5a1 | red | -1.2499761 | 1.92E-08 | -0.98653 | 1.43E-05 |
| Ttc38 | red | -0.8168544 | 0.00151012 | -0.4825885 | 0.07909184 |
| Apex1 | orange | 0.69342509 | 0.00868917 | 0.74713209 | 0.00600527 |
| Tmem59l | red | -1.0019288 | 0.00024726 | -0.7321149 | 0.00931787 |
| Rusc2 | red | -0.7820074 | 4.10E-11 | -0.6481217 | 9.11E-08 |
| Nme5 | skyblue | -1.6919426 | 5.83E-13 | -1.0178991 | 1.74E-05 |
| Fam65b | orange | 1.12821839 | 2.51E-20 | 1.29165681 | 6.45E-26 |
| Tmtc2 | red | -0.9519768 | 2.82E-07 | -1.2249034 | 4.34E-11 |

| | | | | | |
|---|---|---|---|---|---|
| Tmem63b | red | -0.6700633 | 1.15E-06 | -0.4271887 | 0.00301476 |
| Fmnl2 | orange | 1.1059807 | 7.47E-20 | 0.75654262 | 1.13E-09 |
| Phf24 | red | -1.173694 | 0.00861053 | -0.6921915 | 0.14657881 |
| Slc2a6 | yellow | 2.72628972 | 2.33E-17 | 2.41019718 | 1.40E-13 |
| Zranb3 | orange | 0.95305351 | 0.0022946 | 0.28224844 | 0.43087446 |
| Dgkb | skyblue | -2.5591928 | 6.72E-09 | -1.7599138 | 5.81E-05 |
| Colec12 | red | -0.6048597 | 4.59E-05 | -0.471474 | 0.00219887 |
| Prr5 | orange | 0.6125033 | 0.00150512 | 0.13956481 | 0.53418626 |
| Mbnl3 | orange | 1.72371412 | 1.69E-09 | 1.42350754 | 1.41E-06 |
| Meox2 | red | -1.4133305 | 4.71E-21 | -0.8144091 | 1.36E-07 |
| Tm6sf2 | red | -0.8747026 | 0.00942842 | -0.6707578 | 0.05411585 |
| Prickle1 | skyblue | -1.7441749 | 2.45E-18 | -1.4317339 | 1.68E-12 |
| Gatad2a | orange | 0.58244635 | 2.03E-05 | 0.45211136 | 0.00139968 |
| Sh3bp4 | red | -0.7116458 | 6.51E-15 | -0.4268343 | 7.36E-06 |
| BC027231 | orange | 1.01195653 | 7.31E-06 | 0.68423617 | 0.00350998 |
| Ska1 | black | 3.39677503 | 7.92E-07 | 4.91466516 | 5.73E-10 |
| Agr3 | skyblue | -2.0909099 | 1.64E-06 | -0.9661808 | 0.02366325 |
| 3632451O06 | skyblue | -1.8612851 | 3.03E-09 | -1.5091985 | 2.32E-06 |
| Gmip | orange | 1.59003534 | 7.02E-18 | 1.4237705 | 2.39E-14 |
| Rbm43 | orange | 0.68732753 | 7.73E-07 | 0.65449934 | 4.48E-06 |
| Pnpla8 | red | -0.6158162 | 8.64E-05 | -0.670358 | 2.78E-05 |
| Macrod1 | red | -1.257115 | 5.01E-10 | -0.9970781 | 1.88E-06 |
| Gramd1c | red | -0.9564227 | 2.83E-05 | -0.557515 | 0.01994266 |
| Lrrn3 | skyblue | -2.1964698 | 6.22E-08 | -1.6328796 | 9.13E-05 |
| Slc2a13 | red | -0.8147126 | 0.00056285 | -0.8743396 | 0.00034368 |
| Lzts1 | red | -0.604423 | 0.00100842 | -0.3583907 | 0.07275897 |
| Skp1a | red | -0.6002839 | 1.56E-06 | -0.5540191 | 1.58E-05 |
| Slc18a1 | skyblue | -2.0014865 | 6.04E-05 | -0.2137263 | 0.708635 |
| Igsf10 | skyblue | -2.0813683 | 1.57E-12 | -1.2309564 | 6.31E-05 |
| P2ry12 | orange | 1.7153047 | 0.00012005 | 0.77795952 | 0.10288344 |
| P2ry13 | yellow | 2.2225734 | 6.39E-09 | 2.05588648 | 1.77E-07 |
| Rmdn2 | red | -0.8365966 | 0.00081044 | -0.4230151 | 0.11429364 |
| Serbp1 | orange | 0.59928532 | 6.86E-05 | 0.4210652 | 0.00745023 |
| C530008M17 | skyblue | -0.9159996 | 0.10737662 | -1.8858102 | 0.0008215 |
| P2ry14 | orange | 0.89180084 | 6.27E-05 | 0.54679635 | 0.02049127 |
| Glb1l2 | skyblue | -1.598222 | 0.00019582 | -1.3668838 | 0.00199489 |
| Arsi | yellow | 0.42507815 | 0.21739489 | 2.457075 | 3.33E-14 |
| Tbcc | orange | 0.6637662 | 0.00035588 | 0.60139327 | 0.00178994 |
| Exoc1 | orange | 0.64834267 | 4.62E-06 | 0.64104487 | 9.24E-06 |
| Npy1r | skyblue | -2.1201299 | 2.54E-06 | -1.9631988 | 1.75E-05 |
| Lum | skyblue | -2.3914507 | 2.04E-15 | -1.8361741 | 2.83E-09 |
| Elf1 | orange | 0.77518299 | 3.67E-16 | 0.66705679 | 4.19E-12 |
| March1 | yellow | 2.56954614 | 2.12E-10 | 2.16771056 | 1.64E-07 |

| | | | | | |
|---|---|---|---|---|---|
| Btg1 | orange | 1.21434904 | 4.61E-11 | 1.13295814 | 1.52E-09 |
| Eea1 | orange | 0.58396451 | 3.59E-08 | 0.54045998 | 6.35E-07 |
| Card11 | orange | 1.84183572 | 4.12E-10 | 1.83036029 | 8.96E-10 |
| Ppfibp2 | red | -1.0048893 | 4.18E-07 | -0.5439799 | 0.00945603 |
| Akap14 | skyblue | -2.3642744 | 1.09E-05 | -1.6237063 | 0.0028889 |
| Lgi4 | red | -1.2560001 | 4.10E-08 | -0.5234942 | 0.03334002 |
| Ndrg4 | skyblue | -1.1895361 | 0.00578156 | -1.641974 | 0.00015678 |
| Gltscr1l | red | -0.6290471 | 0.00039931 | -0.5056519 | 0.00603018 |
| Fxyd1 | skyblue | -1.3097256 | 9.89E-08 | -1.2104902 | 1.48E-06 |
| Fxyd7 | red | -0.8062572 | 0.00437823 | -1.0518081 | 0.00033132 |
| Fgf1 | skyblue | -1.8551878 | 3.32E-07 | -1.3211997 | 0.0004681 |
| Fut7 | yellow | 2.77039562 | 1.69E-11 | 2.55759157 | 1.92E-09 |
| H2-Aa | orange | 1.34692036 | 2.26E-19 | 1.05576594 | 4.67E-12 |
| Ccdc113 | skyblue | -1.6019734 | 5.15E-13 | -1.0516531 | 3.46E-06 |
| Tssc1 | orange | 1.00938484 | 1.91E-12 | 0.79080344 | 8.64E-08 |
| Etl4 | red | -1.2954229 | 4.27E-13 | -1.0234995 | 2.30E-08 |
| Alg5 | orange | 0.79420069 | 1.81E-05 | 0.50365811 | 0.00976916 |
| Clcn7 | orange | 0.79852604 | 1.07E-05 | 0.66368574 | 0.00039418 |
| Nudt1 | orange | 0.80015866 | 0.0009878 | 0.57801455 | 0.02435682 |
| Ccdc148 | red | -1.1961605 | 0.00398033 | -1.0278911 | 0.01344824 |
| Tcaf1 | red | -0.855644 | 2.96E-10 | -0.7613616 | 3.89E-08 |
| Cenpt | orange | 0.66287514 | 2.47E-05 | 0.64789415 | 5.49E-05 |
| Aaas | orange | 0.65453998 | 9.19E-06 | 0.51221016 | 0.00084488 |
| Tmem184a | red | -0.6447642 | 0.00258592 | -0.4156644 | 0.06548634 |
| Nop14 | orange | 0.69022794 | 4.91E-07 | 0.56717749 | 6.41E-05 |
| Cyld | orange | 0.59880866 | 1.05E-06 | 0.6309357 | 4.50E-07 |
| Dner | skyblue | -1.4436605 | 0.00231129 | -1.5970668 | 0.00170943 |
| Kif15 | yellow | 2.84299927 | 5.97E-37 | 2.89513573 | 3.43E-34 |
| Anln | orange | 1.21955489 | 1.85E-07 | 1.14792327 | 1.54E-06 |
| Rps27l | orange | 0.61343411 | 0.00058373 | 0.60435499 | 0.00098325 |
| Klhl13 | skyblue | -1.5338163 | 3.02E-12 | -1.078738 | 1.82E-06 |
| Amdhd2 | orange | 0.86984309 | 9.65E-09 | 0.63747555 | 5.04E-05 |
| Lpar3 | skyblue | -2.2962 | 5.03E-11 | -1.5221031 | 2.44E-05 |
| Pnpla7 | orange | 0.88719897 | 1.01E-09 | 0.65383861 | 1.31E-05 |
| Mcoln3 | skyblue | -1.7785463 | 0.00389519 | -1.0011117 | 0.12295362 |
| Hspb6 | skyblue | -1.6477142 | 3.87E-09 | -0.9636075 | 0.00103056 |
| Wnt4 | red | -0.9911104 | 7.35E-07 | -0.9037638 | 1.23E-05 |
| Dchs1 | red | -1.0025299 | 1.69E-06 | -0.6128919 | 0.00522399 |
| Syde2 | red | -1.3783093 | 1.46E-07 | -1.0798 | 6.06E-05 |
| Proser3 | red | -0.8487646 | 0.00010793 | -0.5050538 | 0.02916031 |
| Smad6 | red | -1.3534675 | 4.21E-15 | -1.1290047 | 1.26E-10 |
| Dna2 | orange | 1.46924562 | 1.00E-12 | 1.17118123 | 2.98E-08 |
| Phkb | red | -0.8793147 | 6.16E-10 | -0.7968733 | 4.49E-08 |

| | | | | | |
|---|---|---|---|---|---|
| Acaa2 | red | -0.7972356 | 1.72E-05 | -0.7630463 | 5.95E-05 |
| C1qa | yellow | 3.66522925 | 9.95E-21 | 2.8996783 | 3.93E-13 |
| Rap2b | orange | 1.34002271 | 6.60E-05 | 1.23590463 | 0.0003676 |
| C1qc | yellow | 3.36842899 | 1.70E-18 | 2.87095967 | 1.64E-13 |
| Fzd8 | red | -0.8534247 | 5.49E-05 | -0.7171909 | 0.00098305 |
| C1qb | yellow | 3.82427099 | 3.39E-31 | 3.3151297 | 1.57E-23 |
| Unc93b1 | orange | 1.66611034 | 4.11E-14 | 1.37741482 | 9.31E-10 |
| Piwil4 | orange | 0.84245984 | 0.00027895 | 0.73491744 | 0.00212541 |
| Zfp280c | orange | 0.61646618 | 0.00066061 | 0.51788663 | 0.00538987 |
| Stox1 | red | -1.3786702 | 6.71E-07 | -0.8900453 | 0.00158576 |
| Stag3 | yellow | 2.55385165 | 0.00011278 | 2.26601303 | 0.0006849 |
| Nfkbid | yellow | 2.42928262 | 1.06E-23 | 2.3582267 | 2.02E-22 |
| Rab8b | orange | 1.26944312 | 1.00E-11 | 0.94071252 | 1.02E-06 |
| Tmem71 | orange | 0.58774034 | 6.61E-07 | 0.48962308 | 7.15E-05 |
| BC037034 | orange | 0.83492841 | 5.67E-07 | 0.85048434 | 6.00E-07 |
| Kif1bp | red | -0.5485994 | 2.97E-06 | -0.6305234 | 1.12E-07 |
| Bcorl1 | red | -0.8216842 | 6.37E-06 | -0.6885371 | 0.00024545 |
| Cfap221 | skyblue | -2.0817119 | 7.40E-11 | -1.575284 | 8.74E-08 |
| Trim17 | red | -1.0491223 | 0.00939378 | -0.6147763 | 0.14731655 |
| Pml | orange | 1.85122303 | 4.33E-39 | 1.78248265 | 2.93E-36 |
| Nxt1 | orange | 0.75851585 | 0.0002698 | 0.57486432 | 0.0086579 |
| Asap3 | orange | 0.73592398 | 4.82E-06 | 0.93653252 | 6.73E-09 |
| Zfp39 | red | -0.8519854 | 0.00057746 | -0.4205131 | 0.11616105 |
| Tns2 | red | -0.8209677 | 0.00095336 | -0.7010271 | 0.0063878 |
| Zfp113 | red | -0.7875021 | 3.19E-07 | -0.3848732 | 0.01960104 |
| Apln | red | -1.463988 | 1.78E-07 | -0.7985529 | 0.00693929 |
| Hk1 | orange | 0.82175116 | 4.89E-06 | 0.70476785 | 0.00014256 |
| Sstr4 | red | -0.9823463 | 0.00125591 | -0.9471504 | 0.00226644 |
| Frem2 | skyblue | -2.5471107 | 8.03E-09 | -1.6782562 | 0.00019783 |
| Wdr62 | orange | 1.15712634 | 1.11E-09 | 0.88583869 | 5.61E-06 |
| Mmaa | red | -1.0330052 | 1.06E-07 | -0.9068436 | 5.51E-06 |
| Foxa2 | red | -0.8155473 | 3.31E-07 | -0.3014384 | 0.0829628 |
| Apbb1 | red | -1.3045798 | 2.88E-09 | -1.1109867 | 7.97E-07 |
| Clca3b | yellow | 2.1105162 | 0.01579943 | 2.83500709 | 0.00118505 |
| Inhbb | orange | 0.96877427 | 0.01034508 | 1.27762552 | 0.00080194 |
| Paip2 | red | -0.8355525 | 1.08E-06 | -0.8030773 | 4.54E-06 |
| Prkcdbp | orange | 0.79244818 | 0.00105443 | 0.73514948 | 0.00323349 |
| Rbmxl1 | orange | 0.85954447 | 1.07E-05 | 0.73077113 | 0.00029981 |
| Scd1 | skyblue | -1.7710485 | 7.71E-06 | -1.7128985 | 2.37E-05 |
| Prr32 | skyblue | -2.8116072 | 0.00496506 | -1.8076294 | 0.08720812 |
| Lrg1 | orange | 1.54099206 | 1.74E-21 | 1.09961886 | 3.23E-11 |
| Rab11fip3 | red | -0.9610535 | 3.35E-10 | -0.6804309 | 1.76E-05 |
| Ttc29 | skyblue | -3.4376679 | 3.09E-11 | -2.0105624 | 0.0001332 |

| | | | | | |
|---|---|---|---|---|---|
| Dcaf15 | orange | 1.41613628 | 1.00E-28 | 1.22160089 | 2.03E-21 |
| Psd | orange | 0.76413976 | 1.66E-05 | 0.62085564 | 0.00076498 |
| Tmprss13 | red | -1.1331543 | 0.00123982 | -0.7777658 | 0.03485583 |
| Myom3 | skyblue | -2.5639988 | 1.31E-05 | -1.2865381 | 0.04084059 |
| Cfap61 | skyblue | -2.1456321 | 5.90E-06 | -1.6184786 | 0.0008011 |
| Lrrc20 | red | -0.7213927 | 4.54E-10 | -0.5831827 | 1.35E-06 |
| Ppp1r14a | red | -0.6838807 | 0.00027028 | -0.5230879 | 0.00747818 |
| Mycn | skyblue | -1.7318358 | 1.82E-07 | -1.1038176 | 0.00154658 |
| Cyb561d2 | orange | 0.68580131 | 4.03E-07 | 0.50146877 | 0.0004528 |
| Pacrg | skyblue | -1.3813824 | 6.61E-05 | -1.2552141 | 0.0004094 |
| Prf1 | yellow | 4.13059328 | 4.76E-38 | 4.12231929 | 1.12E-37 |
| Islr | red | -1.0459942 | 9.80E-15 | -0.5250764 | 0.00021573 |
| Spry1 | red | -0.7272354 | 0.00101617 | -0.321078 | 0.18512681 |
| Spred3 | orange | 0.93977319 | 0.00012977 | 0.91650278 | 0.00027264 |
| Clic4 | orange | 0.77782812 | 1.02E-06 | 0.65361841 | 6.66E-05 |
| Itih2 | red | -1.2767254 | 3.33E-06 | -0.5336046 | 0.06615458 |
| Dzank1 | skyblue | -2.3066946 | 6.41E-14 | -1.1453598 | 0.00032701 |
| Galnt6 | yellow | 2.16447188 | 9.81E-18 | 1.94813982 | 2.20E-14 |
| Tbcel | red | -0.785204 | 5.16E-05 | -0.560188 | 0.00565993 |
| Tacc3 | yellow | 2.76726554 | 2.29E-41 | 2.52248336 | 8.04E-34 |
| Traf3ip3 | yellow | 1.81260125 | 7.42E-38 | 2.24921935 | 1.26E-53 |
| Tap1 | yellow | 3.34477665 | 1.16E-104 | 3.15172065 | 5.61E-93 |
| Bbs7 | red | -1.0792017 | 4.31E-11 | -0.5818143 | 0.00057484 |
| Larp1 | orange | 0.87006454 | 8.28E-12 | 0.72763879 | 2.31E-08 |
| Map4k1 | yellow | 2.23395206 | 8.61E-32 | 2.2198963 | 5.21E-31 |
| Slc12a9 | orange | 0.91182162 | 1.25E-09 | 0.79969421 | 2.02E-07 |
| Chst7 | red | -1.1407885 | 0.00015162 | -0.9392147 | 0.0025685 |
| Paqr7 | red | -1.0409965 | 3.04E-06 | -0.864894 | 0.00017154 |
| 4930578C19 | red | -1.2346576 | 4.57E-11 | -0.721782 | 0.00032919 |
| Sf3b6 | orange | 0.630137 | 1.58E-06 | 0.48712491 | 0.00038115 |
| Nov | skyblue | -1.6106253 | 2.57E-08 | -0.7174652 | 0.01874194 |
| Letm2 | red | -0.7222899 | 4.12E-05 | -0.6841846 | 0.00016538 |
| Pafah2 | red | -0.9686249 | 3.89E-07 | -0.6922224 | 0.00047456 |
| Trmt6 | orange | 1.239832 | 2.46E-10 | 0.82418489 | 5.12E-05 |
| Icam1 | orange | 0.87197771 | 1.33E-06 | 0.931227 | 3.63E-07 |
| Serpine1 | yellow | 1.99737419 | 2.02E-09 | 2.00414388 | 2.73E-09 |
| Fer1l5 | red | -1.0595519 | 1.51E-06 | -0.9714154 | 1.82E-05 |
| Vnn1 | red | -0.9667847 | 0.00379923 | -0.2905842 | 0.44335274 |
| Cep85 | orange | 2.01234948 | 1.82E-25 | 1.33270141 | 1.81E-11 |
| Arid5a | yellow | 2.35397328 | 3.76E-27 | 2.50127795 | 1.97E-30 |
| Slc18b1 | red | -1.0793481 | 1.15E-11 | -0.7708359 | 2.31E-06 |
| 4930427A07I | yellow | 2.19052518 | 2.31E-22 | 2.26760627 | 3.45E-23 |
| Papl | skyblue | -1.7036934 | 0.00521443 | -0.8919812 | 0.16557242 |

| | | | | | |
|---|---|---|---|---|---|
| Dtl | yellow | 2.54143247 | 5.90E-14 | 2.80021624 | 1.30E-15 |
| Slc2a12 | skyblue | -2.0708465 | 9.50E-12 | -1.288071 | 3.11E-05 |
| Zmat4 | skyblue | -2.6624315 | 1.11E-05 | -1.5409072 | 0.01634503 |
| Cib2 | skyblue | -1.638215 | 2.74E-10 | -1.0228141 | 0.0001605 |
| Arhgef4 | red | -0.7260993 | 0.0092285 | -0.2515919 | 0.41249627 |
| Mavs | red | -0.5855781 | 9.69E-06 | -0.4873838 | 0.00036796 |
| Atg14 | red | -0.6600044 | 0.00177008 | -0.4178327 | 0.06183987 |
| Rapgef6 | orange | 0.60231335 | 7.30E-06 | 0.47920448 | 0.00057349 |
| Shank2 | skyblue | -2.3005242 | 3.63E-19 | -0.9905822 | 0.00015025 |
| Dlgap5 | yellow | 3.67737046 | 2.68E-51 | 4.14544062 | 1.38E-55 |
| Rps16 | orange | 0.59400704 | 4.20E-05 | 0.41153524 | 0.00662766 |
| Vash2 | skyblue | -1.83205 | 1.24E-05 | -1.6924323 | 0.00013072 |
| Wdhd1 | yellow | 2.30384678 | 1.19E-25 | 1.79117484 | 1.29E-15 |
| Tob1 | red | -1.2472463 | 1.97E-10 | -0.9278888 | 4.30E-06 |
| Kcnh3 | red | -1.2864647 | 8.86E-05 | -0.5966182 | 0.08636439 |
| Gch1 | orange | 1.4372693 | 9.04E-11 | 1.22385928 | 6.39E-08 |
| BC022687 | red | -0.797044 | 0.00019125 | -0.8120091 | 0.00027144 |
| Nme1 | orange | 1.27689791 | 4.96E-09 | 0.79912234 | 0.00051166 |
| Osbpl5 | red | -0.7158592 | 1.25E-05 | -0.5067892 | 0.00297024 |
| Kcnmb2 | skyblue | -2.5184036 | 2.25E-11 | -1.8563864 | 8.14E-07 |
| Tnfrsf23 | orange | 1.37758855 | 0.00058165 | 1.32516696 | 0.00131096 |
| Wdtc1 | red | -0.6586282 | 5.29E-08 | -0.6089203 | 8.64E-07 |
| Kcnk2 | skyblue | -3.0898259 | 4.88E-13 | -1.8004437 | 4.68E-05 |
| Cldn11 | skyblue | -3.4253522 | 0.00070044 | -2.6758615 | 0.00291295 |
| Rgs22 | skyblue | -2.255154 | 3.33E-09 | -1.5427554 | 6.09E-05 |
| Cdkn3 | yellow | 3.27269514 | 7.06E-21 | 3.24298809 | 1.98E-17 |
| Slc25a43 | orange | 1.23943632 | 7.66E-05 | 0.93539118 | 0.00502783 |
| Zfp60 | red | -0.9453937 | 5.52E-07 | -0.6222428 | 0.00165476 |
| Prkci | red | -0.7225033 | 1.16E-10 | -0.4517056 | 0.0001122 |
| H2-DMa | orange | 1.60404384 | 1.04E-13 | 1.31644668 | 2.45E-09 |
| Kctd8 | skyblue | -2.7505287 | 1.67E-07 | -1.6261203 | 0.00165054 |
| Gpr160 | orange | 0.91858752 | 2.82E-07 | 1.18282751 | 1.31E-10 |
| Cdkn1c | red | -0.7661179 | 0.00141908 | -1.0064333 | 3.23E-05 |
| Esyt3 | red | -1.2317732 | 0.00012391 | -0.9101924 | 0.00617297 |
| Armc3 | skyblue | -1.8309447 | 9.04E-14 | -1.5860325 | 6.77E-11 |
| Cd81 | red | -0.6502518 | 3.10E-06 | -0.4463356 | 0.00214478 |
| Fam13a | skyblue | -2.2156065 | 6.64E-08 | -1.6956572 | 6.13E-05 |
| Cisd1 | red | -0.5917894 | 0.00015161 | -0.4919571 | 0.00249923 |
| Ccdc33 | skyblue | -2.1095311 | 2.77E-11 | -1.183459 | 0.00015741 |
| Ckap2 | yellow | 3.35923397 | 8.22E-25 | 2.75867987 | 3.27E-17 |
| Themis2 | yellow | 3.09792854 | 2.52E-24 | 3.34349549 | 4.97E-28 |
| Limch1 | red | -1.0983776 | 3.96E-05 | -0.855757 | 0.00201122 |
| Nek5 | red | -1.3503632 | 9.93E-09 | -0.8070801 | 0.00084848 |

| | | | | | |
|---|---|---|---|---|---|
| Ptger2 | orange | 1.03720081 | 0.00034776 | 0.80486546 | 0.00798054 |
| Slc16a9 | red | -1.1052962 | 0.00047346 | -0.9629747 | 0.00315308 |
| Dzip1l | red | -0.7960076 | 1.40E-06 | -0.4626527 | 0.00728115 |
| Iqch | skyblue | -3.3042394 | 0.00023638 | -1.8421217 | 0.03178823 |
| Rpl10a | orange | 0.63617257 | 3.25E-07 | 0.5086241 | 7.66E-05 |
| D630003M21 | skyblue | -1.35057 | 1.25E-05 | -1.4784489 | 4.59E-06 |
| Fbxw17 | orange | 1.36931284 | 3.75E-19 | 1.22892888 | 1.52E-15 |
| Tgm2 | orange | 1.80527087 | 4.44E-14 | 1.73075255 | 7.48E-13 |
| Smim14 | red | -0.6249173 | 1.71E-05 | -0.6183564 | 3.19E-05 |
| Rtkn2 | skyblue | -2.7963074 | 0.00087201 | -2.1442828 | 0.01436256 |
| Nmrk1 | orange | 0.62010058 | 0.00020972 | 0.48288689 | 0.00718103 |
| LOC1026395 | yellow | 4.23219923 | 8.50E-44 | 3.9956066 | 5.40E-39 |
| Cpe | skyblue | -1.3467285 | 5.29E-15 | -1.1931359 | 9.13E-12 |
| Zfp365 | red | -1.2240121 | 6.02E-13 | -0.8479149 | 1.51E-06 |
| Aim2 | yellow | 2.82701324 | 1.48E-29 | 2.70208724 | 2.30E-26 |
| Egr2 | yellow | 2.4330051 | 0.00152843 | 2.7820941 | 0.00055612 |
| Ackr1 | yellow | 2.2499875 | 5.58E-10 | 2.44048824 | 3.65E-10 |
| Wdr19 | red | -0.8422298 | 3.16E-05 | -0.4283423 | 0.04648501 |
| Pcdh18 | red | -0.9305976 | 0.00019422 | -0.7462483 | 0.00389022 |
| H2afz | orange | 1.11349967 | 1.28E-09 | 1.03607511 | 2.97E-08 |
| Sirpa | orange | 1.1032679 | 2.32E-05 | 0.95903689 | 0.00035917 |
| Bri3bp | orange | 0.93456998 | 9.19E-10 | 0.83565553 | 1.14E-07 |
| Tmem156 | orange | 1.87671589 | 2.05E-07 | 1.72471759 | 3.31E-06 |
| Ddx60 | yellow | 2.6702349 | 1.15E-33 | 2.71954418 | 1.12E-34 |
| Inpp4b | orange | 1.1581915 | 2.30E-06 | 0.63794803 | 0.01429936 |
| Ccr7 | orange | 1.80778357 | 7.42E-22 | 1.8438003 | 2.95E-22 |
| Fgd3 | yellow | 2.08392038 | 9.55E-11 | 2.15241141 | 3.71E-11 |
| Ccdc55 | orange | 0.7136818 | 0.00031148 | 0.40739054 | 0.05499579 |
| 1110007C09 | orange | 0.62091269 | 0.00349664 | 0.57697505 | 0.00852762 |
| Fam101a | red | -1.2896858 | 3.20E-05 | -0.8797986 | 0.00673619 |
| Zc3h7a | orange | 0.92983384 | 1.36E-23 | 0.88611353 | 2.42E-21 |
| Ninj1 | orange | 1.12261117 | 1.49E-06 | 0.9471919 | 8.12E-05 |
| 1110032A03I | red | -0.8582973 | 6.05E-09 | -0.6897796 | 5.17E-06 |
| Ccdc129 | skyblue | -2.8615344 | 3.46E-07 | -1.835969 | 0.00130927 |
| Ccdc92 | red | -0.8501806 | 4.12E-06 | -0.7919181 | 2.83E-05 |
| Wnk2 | red | -0.9907036 | 0.0039972 | -1.103705 | 0.00167956 |
| Rmi2 | yellow | 2.2150252 | 0.00159835 | 4.14054804 | 5.48E-08 |
| Slc24a2 | skyblue | -1.879941 | 0.00520399 | -1.7557078 | 0.01375874 |
| Parp11 | orange | 1.81363 | 1.58E-42 | 1.8208729 | 1.51E-42 |
| 2700029M09 | orange | 1.01651068 | 1.52E-08 | 0.5906073 | 0.00179153 |
| Acer2 | red | -0.643236 | 0.00805386 | -0.7879319 | 0.00133636 |
| Dnah10 | skyblue | -1.7729769 | 8.37E-08 | -1.1716948 | 0.00063236 |
| Fam188b | red | -0.7203649 | 0.000108 | -0.7368779 | 9.05E-05 |

| | | | | | |
|---|---|---|---|---|---|
| Socs1 | black | 4.6214377 | 4.27E-82 | 5.26110486 | 2.58E-98 |
| Ptpdc1 | red | -1.2393881 | 7.13E-18 | -0.7094132 | 1.60E-06 |
| Sult6b1 | yellow | 2.11895851 | 0.0033239 | 2.19328681 | 0.00977605 |
| Haus6 | orange | 0.67426892 | 3.15E-07 | 0.55749652 | 4.49E-05 |
| Nod1 | orange | 0.62285812 | 1.03E-18 | 0.65917929 | 9.02E-21 |
| Smim3 | orange | 0.97979706 | 9.44E-05 | 0.78638944 | 0.0025408 |
| Dlec1 | skyblue | -2.5144649 | 6.58E-10 | -1.1014013 | 0.00921886 |
| Mturn | skyblue | -2.2432185 | 1.39E-07 | -1.942826 | 8.66E-06 |
| Csf3 | yellow | 3.99739971 | 6.53E-07 | 3.48345594 | 4.79E-07 |
| Cntln | red | -0.8382158 | 9.81E-07 | -0.5279117 | 0.0029946 |
| Kcna6 | skyblue | -2.9975408 | 1.25E-12 | -2.0543756 | 3.58E-07 |
| Cnbd2 | orange | 1.42620056 | 1.14E-09 | 1.08153678 | 1.44E-05 |
| Atp13a4 | skyblue | -1.3327946 | 0.00049836 | -1.3635178 | 0.00059589 |
| Ano2 | skyblue | -0.5627212 | 0.50523529 | -2.411775 | 0.00926687 |
| Cdon | red | -0.7681229 | 2.93E-06 | -0.5585951 | 0.00111086 |
| Fam210a | red | -0.6768376 | 1.36E-09 | -0.6235223 | 6.30E-08 |
| Tbc1d32 | red | -1.1325466 | 8.63E-07 | -0.8335397 | 0.00043744 |
| Camk4 | yellow | 1.76948623 | 8.95E-06 | 2.6312618 | 7.47E-11 |
| Rbm24 | skyblue | -1.2824257 | 1.07E-05 | -1.391357 | 2.79E-06 |
| Stox2 | red | -0.7647565 | 1.01E-05 | -0.7038561 | 7.71E-05 |
| Notch3 | skyblue | -1.3831464 | 8.54E-08 | -1.1615701 | 1.21E-05 |
| Cd84 | yellow | 2.440806 | 2.22E-22 | 2.15852619 | 1.70E-17 |
| Ormdl3 | red | -0.7465681 | 2.94E-09 | -0.5819463 | 7.74E-06 |
| Gstp2 | skyblue | -1.4894296 | 5.00E-07 | -1.2577679 | 4.15E-05 |
| P3h2 | red | -1.518693 | 6.92E-07 | -0.9638767 | 0.00240604 |
| Ttc39b | orange | 2.05204344 | 1.43E-23 | 1.82680281 | 1.01E-18 |
| Enpp6 | skyblue | -0.5472098 | 0.47577982 | -2.9701775 | 3.88E-05 |
| Fam126b | red | -0.6657079 | 0.00051516 | -0.4488777 | 0.02590412 |
| Slamf7 | yellow | 3.35296406 | 6.48E-21 | 2.97168541 | 2.22E-16 |
| Chpf2 | orange | 0.78602278 | 1.67E-07 | 0.74076328 | 1.32E-06 |
| Rilp | red | -1.0395174 | 1.14E-05 | -0.6806368 | 0.00608691 |
| Kcna7 | skyblue | -1.8779519 | 1.20E-05 | -0.7137563 | 0.12971707 |
| Asb10 | skyblue | -1.7082659 | 9.72E-06 | -2.0488575 | 2.98E-07 |
| Tapbpl | orange | 1.63282148 | 1.37E-38 | 1.64336891 | 6.11E-39 |
| Tlcd2 | red | -0.5513074 | 0.02967973 | -0.707576 | 0.00685817 |
| Fam198a | skyblue | -1.7384266 | 2.74E-09 | -0.8341175 | 0.00626537 |
| Hrc | skyblue | -1.8914027 | 1.79E-10 | -1.9564216 | 7.55E-11 |
| Sobp | red | -1.1259869 | 0.00142709 | -0.691612 | 0.06836149 |
| Ncapd2 | yellow | 2.02839765 | 4.55E-28 | 2.18158804 | 6.53E-32 |
| Hoxa5 | red | -1.1124539 | 4.58E-09 | -0.9616792 | 7.63E-07 |
| Trpm4 | red | -1.1482231 | 1.35E-10 | -0.7979928 | 1.61E-05 |
| Sema7a | orange | 2.0933365 | 8.39E-25 | 1.37710715 | 4.60E-11 |
| Slc22a23 | red | -0.9962604 | 2.44E-06 | -0.6073933 | 0.00619218 |

| | | | | | |
|---|---|---|---|---|---|
| Fau | orange | 0.85128812 | 8.89E-09 | 0.53607818 | 0.00053118 |
| Nop2 | orange | 0.85983035 | 2.04E-07 | 0.75272822 | 9.44E-06 |
| Ostm1 | orange | 0.79517457 | 2.46E-05 | 0.63419064 | 0.00114737 |
| Bphl | skyblue | -1.5727387 | 1.35E-18 | -1.1133876 | 1.12E-09 |
| Snx25 | red | -1.1767633 | 4.38E-06 | -0.8984425 | 0.0007214 |
| Ccdc155 | skyblue | -2.3495727 | 6.55E-06 | -1.241532 | 0.02123631 |
| Snx10 | yellow | 2.27331588 | 3.48E-18 | 2.0014559 | 3.37E-14 |
| Lace1 | red | -1.2572653 | 5.02E-10 | -1.1417689 | 2.96E-08 |
| Cd160 | yellow | 3.43252339 | 3.89E-11 | 2.82867713 | 7.98E-08 |
| Kcnh2 | skyblue | -1.397774 | 9.17E-05 | -1.4761292 | 5.02E-05 |
| Tsr1 | orange | 0.61411085 | 0.00156387 | 0.80683011 | 3.98E-05 |
| Txlng | red | -0.6951584 | 0.00110156 | -0.5600507 | 0.01154568 |
| Tcte2 | red | -0.8575562 | 0.0066359 | -0.6097636 | 0.06698444 |
| Plcl1 | red | -0.8629215 | 3.51E-05 | -0.8040935 | 0.00017476 |
| Arl5c | orange | 1.14752467 | 0.00378425 | 1.59454014 | 6.66E-05 |
| Lasp1 | orange | 0.69586094 | 0.00056301 | 0.73780657 | 0.00034533 |
| Pcp4l1 | skyblue | -2.5823158 | 1.04E-19 | -2.2331806 | 6.95E-15 |
| Gmds | orange | 0.84455282 | 1.07E-07 | 0.57706434 | 0.00047917 |
| Trp53inp2 | red | -1.2042169 | 1.05E-12 | -0.9726299 | 1.92E-08 |
| Ttk | yellow | 2.56778831 | 3.83E-14 | 2.9281863 | 1.67E-15 |
| Mpp6 | orange | 0.71209047 | 0.00198822 | 0.58854481 | 0.01422301 |
| Pmepa1 | orange | 0.83055003 | 0.00036811 | 0.7416644 | 0.00204486 |
| Foxf2 | red | -1.3352865 | 4.53E-15 | -0.9164346 | 1.85E-07 |
| Hfe2 | skyblue | -1.591042 | 2.81E-07 | -1.3450908 | 2.74E-05 |
| Egr1 | orange | 1.13576583 | 4.07E-06 | 0.88505558 | 0.00051175 |
| Srcin1 | red | -1.3428805 | 3.38E-07 | -0.6222993 | 0.02604227 |
| Dennd2a | red | -1.1365971 | 7.44E-16 | -0.95348 | 2.63E-11 |
| Tmem255b | skyblue | -1.5115372 | 0.0002135 | -1.9865425 | 4.33E-07 |
| Chmp4b | orange | 1.360056 | 5.96E-17 | 1.27405385 | 7.90E-15 |
| Cdk19 | red | -0.3919842 | 0.01991003 | -0.6050554 | 0.0003278 |
| Tfdp1 | orange | 0.57243755 | 3.52E-05 | 0.59178565 | 2.89E-05 |
| Tmco3 | red | -0.7482433 | 2.13E-09 | -0.6564523 | 2.73E-07 |
| Ptov1 | red | -0.7225363 | 4.13E-08 | -0.5685197 | 2.89E-05 |
| Dcun1d2 | red | -0.7699883 | 6.22E-06 | -0.7104697 | 5.01E-05 |
| Parp12 | orange | 1.78785564 | 1.80E-42 | 1.8292908 | 2.14E-44 |
| Gdf15 | yellow | 3.29040865 | 4.92E-11 | 3.36872524 | 2.86E-11 |
| Rpf2 | orange | 1.18551613 | 2.09E-07 | 1.07303436 | 5.12E-06 |
| Grtp1 | red | -1.0082839 | 0.00480873 | -0.5174763 | 0.18068813 |
| AI317395 | red | -1.1110918 | 0.00033143 | -0.6962929 | 0.03370497 |
| Car14 | skyblue | -1.4333014 | 5.61E-07 | -1.6598783 | 9.68E-09 |
| Rgs4 | red | -0.791608 | 4.94E-05 | -0.7307159 | 0.00027668 |
| Atf5 | red | -0.6619139 | 1.60E-05 | -0.4373068 | 0.00650243 |
| BC028528 | red | -1.2201994 | 3.01E-07 | -1.0731428 | 1.27E-05 |

| | | | | | |
|---|---|---|---|---|---|
| Ciart | skyblue | -1.4640894 | 4.33E-08 | -1.26233 | 3.93E-06 |
| Ift172 | red | -0.934568 | 1.15E-10 | -0.7852855 | 1.20E-07 |
| Saxo2 | red | -1.5004818 | 2.97E-07 | -0.999846 | 0.00085415 |
| Susd4 | red | -0.8856559 | 1.30E-07 | -0.2803244 | 0.12788774 |
| Susd1 | orange | 0.76832328 | 0.00281885 | 0.74997858 | 0.00387934 |
| Pln | skyblue | -1.5219389 | 0.13619182 | -3.0916386 | 0.00204202 |
| Akap12 | orange | 0.74198097 | 0.03011862 | 0.97938659 | 0.00444563 |
| Colec10 | skyblue | -6.3500378 | 3.46E-05 | -4.0103658 | 0.00614707 |
| Slc35f1 | skyblue | -2.5745978 | 0.00112293 | -1.9112663 | 0.02426082 |
| Dock10 | orange | 1.26258663 | 5.36E-11 | 1.08135723 | 4.05E-08 |
| Mcl1 | orange | 0.59365346 | 1.83E-06 | 0.50346076 | 8.60E-05 |
| Rassf7 | red | -0.8195917 | 1.20E-05 | -0.4499819 | 0.02346764 |
| Med30 | orange | 0.40083458 | 0.0521969 | 0.62486402 | 0.00261047 |
| Tm6sf1 | orange | 0.74202667 | 0.00097993 | 0.84942019 | 0.00021159 |
| Lrrc56 | red | -0.9720464 | 1.08E-07 | -0.6731531 | 0.00032743 |
| Ctss | yellow | 3.82191139 | 1.08E-27 | 3.32726763 | 4.44E-21 |
| Creb3l2 | red | -0.6861085 | 2.05E-08 | -0.53493 | 2.26E-05 |
| Rnh1 | orange | 0.86274857 | 1.50E-08 | 0.75909509 | 1.13E-06 |
| Fsd2 | skyblue | -1.5077658 | 0.00054146 | -1.8921858 | 2.07E-05 |
| Mybpc2 | red | -0.9302816 | 0.00173411 | -0.4426101 | 0.1713231 |
| Scube3 | red | -1.5446307 | 6.73E-16 | -0.5911456 | 0.00231773 |
| Trps1 | orange | 0.67199344 | 2.34E-05 | 0.43827468 | 0.00895643 |
| Hoxb4 | red | -0.5931554 | 6.28E-08 | -0.4279368 | 0.00016208 |
| Taf5l | orange | 0.64162945 | 0.00023606 | 0.53178508 | 0.00321091 |
| Hoxb5 | red | -0.8314081 | 0.00016964 | -0.6719315 | 0.00332939 |
| Dsel | red | -0.7844374 | 0.0001096 | -0.4948283 | 0.0204091 |
| Bud31 | orange | 0.68339799 | 2.68E-05 | 0.45126718 | 0.00848437 |
| Pkhd1l1 | red | -1.190005 | 0.00070049 | -1.1924171 | 0.00097439 |
| Mvb12b | red | -0.8032425 | 3.20E-10 | -0.4207884 | 0.00168207 |
| Ttll6 | skyblue | -1.6016989 | 3.81E-05 | -0.8580164 | 0.036435 |
| Nup205 | orange | 0.87682011 | 1.44E-08 | 0.79700265 | 4.66E-07 |
| Alpk3 | skyblue | -1.4698231 | 1.66E-07 | -1.2769602 | 1.07E-05 |
| Ptpn3 | red | -0.9423821 | 2.48E-08 | -0.6074859 | 0.00059262 |
| Ascc3 | orange | 0.83753323 | 9.92E-10 | 0.93996349 | 7.69E-12 |
| Vill | orange | 0.87370045 | 2.66E-05 | 0.69147537 | 0.00131164 |
| Ephx1 | skyblue | -2.3576591 | 1.22E-10 | -1.8349212 | 1.11E-06 |
| Sema6c | skyblue | -2.2900177 | 1.80E-09 | -1.3747572 | 0.00052464 |
| Scgb3a2 | skyblue | -2.9590729 | 1.05E-14 | -2.0431108 | 2.31E-07 |
| Lefty1 | red | -0.9550447 | 0.00415584 | -1.3081311 | 0.00011264 |
| Zscan2 | red | -0.7029819 | 4.17E-05 | -0.3524198 | 0.06239963 |
| Rap1gap2 | orange | 1.16315206 | 5.99E-05 | 0.84462277 | 0.00518513 |
| Gngt2 | orange | 1.11121074 | 1.21E-07 | 1.0553153 | 9.39E-07 |
| Ctnnal1 | red | -0.8268477 | 0.00010605 | -0.6301513 | 0.00454639 |

| | | | | | |
|---|---|---|---|---|---|
| Agbl3 | red | -0.6935455 | 0.00405237 | -0.6786087 | 0.00594386 |
| Gcnt1 | orange | 1.7173562 | 2.40E-09 | 1.75271248 | 2.02E-09 |
| Baiap2l1 | red | -0.7905202 | 2.36E-05 | -0.5473887 | 0.00498189 |
| Zcchc2 | orange | 1.22474516 | 1.53E-16 | 0.94208482 | 5.15E-10 |
| Bpgm | red | -0.7075383 | 0.00088325 | -0.7304711 | 0.00080595 |
| Nipal2 | red | -0.7755304 | 2.71E-07 | -0.5616597 | 0.00030106 |
| Mrps34 | orange | 0.38770625 | 0.11698487 | 0.73353449 | 0.00262235 |
| A230050P20I | orange | 0.52092104 | 0.00013938 | 0.62716227 | 5.70E-06 |
| Fam117a | red | -0.6684225 | 8.47E-05 | -0.4388053 | 0.01446123 |
| Rpl12 | orange | 0.80524236 | 9.05E-09 | 0.67612294 | 2.60E-06 |
| Plcl2 | orange | 1.52509062 | 6.06E-14 | 1.49567507 | 3.94E-13 |
| Sccpdh | red | -0.5652089 | 5.65E-06 | -0.6617598 | 2.06E-07 |
| Prc1 | yellow | 2.4488486 | 8.93E-21 | 2.34493119 | 6.87E-19 |
| Cnst | red | -0.6715053 | 0.00039106 | -0.5856843 | 0.00270157 |
| Slco4a1 | yellow | 1.71375215 | 7.76E-09 | 2.27478638 | 4.74E-14 |
| Pdk2 | skyblue | -1.5487013 | 2.33E-05 | -1.4095311 | 0.00018073 |
| Cldnd2 | yellow | 2.66009973 | 6.78E-05 | 2.26977994 | 0.00158395 |
| Rabggtb | orange | 0.58972496 | 7.92E-07 | 0.56410869 | 4.22E-06 |
| Rbbp8nl | skyblue | -0.8977459 | 0.22337414 | -2.0716483 | 0.00365966 |
| Bloc1s5 | red | -0.6596953 | 3.59E-05 | -0.4093324 | 0.0159898 |
| Tspyl5 | skyblue | -1.2307477 | 0.00112664 | -1.7105958 | 1.17E-05 |
| 1700019L03F | red | -1.41598 | 2.25E-11 | -0.6032548 | 0.00617734 |
| Bmp6 | skyblue | -1.6185499 | 2.38E-12 | -1.3307521 | 1.74E-08 |
| Tlr4 | orange | 0.71245586 | 1.06E-05 | 0.83302928 | 3.41E-07 |
| Ttc16 | red | -1.4779157 | 3.35E-10 | -0.9629933 | 7.21E-05 |
| Adrm1 | orange | 0.82617079 | 4.10E-09 | 0.67703266 | 2.76E-06 |
| Eme1 | yellow | 3.04773381 | 3.60E-16 | 3.69612447 | 1.01E-18 |
| Myo16 | skyblue | -2.149621 | 8.04E-05 | -1.8241308 | 0.00098373 |
| Ak5 | red | -1.4511241 | 0.00214632 | -0.8896676 | 0.08359975 |
| Echdc3 | skyblue | -1.5655813 | 1.41E-06 | -0.8117744 | 0.01665082 |
| Zfp503 | red | -0.8943979 | 1.30E-05 | -0.4287571 | 0.05154671 |
| Chad | skyblue | -2.7180332 | 3.07E-10 | -1.8789406 | 2.46E-05 |
| Sptlc3 | red | -1.1546623 | 0.00058227 | -1.2701862 | 0.00024477 |
| Wdr93 | skyblue | -2.2513339 | 2.88E-09 | -1.3490811 | 0.00035605 |
| Nexn | red | -0.6438149 | 0.00067915 | -0.6119603 | 0.00180634 |
| F13a1 | orange | 0.61606653 | 0.032723 | 0.82449017 | 0.00447075 |
| Mycbpap | skyblue | -2.162218 | 5.43E-13 | -1.4469764 | 2.47E-06 |
| Nrn1 | skyblue | -1.9006694 | 3.97E-10 | -1.1334743 | 0.00030929 |
| Adgrg6 | red | -1.1706 | 6.33E-14 | -1.0052788 | 2.55E-10 |
| Gipc2 | red | -0.5000935 | 0.00361232 | -0.6029234 | 0.00057655 |
| Whrn | red | -0.7311535 | 0.00317547 | -0.3077215 | 0.25321974 |
| Camk1d | orange | 0.60005163 | 0.00329463 | 0.59700639 | 0.00464828 |
| Runx2 | yellow | 2.76247613 | 1.21E-21 | 2.43455546 | 6.44E-17 |

| | | | | | |
|---|---|---|---|---|---|
| Cdh26 | skyblue | -4.3567068 | 3.30E-20 | -4.5016055 | 1.40E-18 |
| Akna | orange | 1.43407899 | 3.52E-14 | 1.18251431 | 9.04E-10 |
| Akap7 | red | -0.9507835 | 7.75E-10 | -0.6258598 | 9.32E-05 |
| Tbc1d19 | red | -0.8659712 | 4.10E-13 | -0.4935683 | 7.83E-05 |
| AW209491 | red | -0.6822883 | 0.00181527 | -0.635838 | 0.00480865 |
| Nubp2 | orange | 1.08135392 | 6.04E-09 | 0.96729411 | 3.60E-07 |
| Fanci | yellow | 3.09288651 | 1.57E-11 | 4.35404613 | 1.29E-17 |
| Rbpj | orange | 0.59063975 | 0.00028247 | 0.45766326 | 0.00691874 |
| Nlrc4 | orange | 1.52605726 | 6.86E-05 | 1.31220896 | 0.00107914 |
| 1110008P14I | orange | 0.69444237 | 0.00016713 | 0.49091906 | 0.01146246 |
| Orm1 | orange | 1.30770836 | 0.0098192 | 0.17422292 | 0.77633674 |
| Adk | red | -0.8587018 | 2.74E-12 | -0.7218467 | 9.11E-09 |
| Zdhhc1 | red | -1.2063011 | 5.59E-10 | -0.79649 | 8.04E-05 |
| Metrnl | orange | 1.34276677 | 1.92E-06 | 1.52907214 | 9.19E-08 |
| Suv39h1 | orange | 0.96250655 | 2.28E-08 | 0.77007601 | 1.55E-05 |
| Stx11 | orange | 1.63638575 | 7.72E-12 | 1.68710206 | 2.57E-12 |
| Isg20 | yellow | 2.213804 | 1.65E-29 | 1.91513257 | 2.89E-22 |
| Lyplal1 | red | -1.2379516 | 0.00430892 | -0.6631935 | 0.14835159 |
| Lgi2 | red | -0.9590069 | 6.78E-08 | -0.4919729 | 0.00878117 |
| Fn3krp | red | -0.7473123 | 0.00223939 | -0.6717719 | 0.00794639 |
| Pomt1 | red | -0.6721857 | 4.55E-07 | -0.4369513 | 0.00149583 |
| Vstm2b | skyblue | -3.0949337 | 2.70E-08 | -2.1765181 | 6.15E-05 |
| Gimap3 | yellow | 2.15545884 | 1.03E-23 | 2.21261371 | 9.72E-25 |
| Megf9 | red | -0.8132511 | 0.00014567 | -0.6560722 | 0.00310386 |
| Pcsk1n | skyblue | -1.7504649 | 0.00122415 | -0.8174669 | 0.1681277 |
| Azi2 | orange | 0.76767239 | 1.20E-13 | 0.59613904 | 1.90E-08 |
| BC017643 | orange | 0.82973308 | 3.73E-11 | 0.66449807 | 2.40E-07 |
| Ndst2 | orange | 0.63470293 | 1.37E-05 | 0.62332408 | 2.86E-05 |
| Clnk | orange | 1.39613209 | 0.0167842 | 2.00371428 | 0.00385095 |
| Rftn1 | orange | 1.00436388 | 1.98E-12 | 0.96301935 | 2.94E-11 |
| Rab3gap2 | orange | 0.48885528 | 0.00308336 | 0.62275791 | 0.00019552 |
| Atp6v0e2 | red | -1.0872661 | 1.19E-06 | -0.8403698 | 0.00028481 |
| C130074G19 | red | -0.9975963 | 2.42E-20 | -0.7941989 | 5.21E-13 |
| Exosc2 | orange | 0.69102273 | 2.14E-06 | 0.45513042 | 0.0029769 |
| Picalm | orange | 1.06929163 | 2.59E-09 | 0.90160582 | 9.71E-07 |
| Sectm1b | black | 4.12369642 | 2.63E-09 | 5.20976747 | 5.43E-14 |
| March4 | skyblue | -2.3678359 | 2.12E-18 | -1.1659329 | 8.20E-06 |
| Synpo2l | skyblue | -1.5148469 | 0.00027502 | -1.10986 | 0.0107862 |
| Wdr45 | red | -0.7879073 | 0.00011469 | -0.7628494 | 0.00027508 |
| Dusp10 | orange | 0.88184824 | 3.28E-07 | 0.96390491 | 5.91E-08 |
| Cdh6 | red | -1.2976437 | 3.28E-06 | -0.8864512 | 0.00212121 |
| Ccdc81 | skyblue | -2.2921709 | 1.19E-12 | -1.6225528 | 3.65E-07 |
| Neil3 | yellow | 3.20261026 | 4.55E-18 | 4.31746082 | 1.38E-24 |

| | | | | | |
|---|---|---|---|---|---|
| Prdm16 | red | -1.2090519 | 1.59E-15 | -0.6373138 | 5.29E-05 |
| Dcxr | red | -1.1554521 | 0.00011414 | -0.7627683 | 0.01520513 |
| Snx22 | red | -1.1462846 | 2.47E-05 | -0.502023 | 0.08730022 |
| Morc3 | orange | 1.10643713 | 3.80E-11 | 0.89086414 | 2.16E-07 |
| Col10a1 | skyblue | -2.6115049 | 0.01863648 | -3.8842553 | 0.00104912 |
| Prrx2 | orange | 0.41627638 | 0.50319832 | 1.88889419 | 0.00172041 |
| Micu3 | red | -0.4374371 | 0.02077554 | -0.6165349 | 0.00119956 |
| Tspyl4 | red | -1.1430728 | 5.34E-06 | -0.7516367 | 0.00413651 |
| Cdnf | red | -1.3452139 | 2.35E-05 | -1.0570193 | 0.00173958 |
| Znfx1 | orange | 1.57771329 | 3.03E-30 | 1.59890208 | 5.47E-31 |
| Cyp7b1 | yellow | 2.1517242 | 5.50E-11 | 1.94285301 | 7.42E-09 |
| Foxp3 | yellow | 3.914145 | 2.18E-23 | 3.54594339 | 9.41E-21 |
| Atp8b1 | orange | 0.75160321 | 3.81E-05 | 0.55353042 | 0.00360689 |
| Zufsp | orange | 1.45381349 | 2.16E-40 | 0.98584797 | 1.12E-18 |
| Ncam1 | red | -0.7675069 | 7.76E-06 | -0.8447815 | 4.04E-06 |
| Cfap70 | skyblue | -2.4074148 | 8.26E-16 | -1.3850864 | 3.82E-06 |
| Ajap1 | red | -0.9060587 | 0.03806461 | -1.1564309 | 0.00844907 |
| Rsph4a | skyblue | -2.7214859 | 1.32E-19 | -1.4528827 | 1.19E-06 |
| Ccser1 | red | -0.9706898 | 2.63E-06 | -0.482664 | 0.02490086 |
| Rcan2 | skyblue | -1.5571066 | 0.00028547 | -1.5865931 | 0.00031026 |
| Rbms3 | red | -1.0878459 | 8.08E-12 | -0.9130983 | 2.07E-08 |
| Tmem246 | red | -1.0776516 | 1.10E-09 | -0.8144563 | 8.04E-06 |
| 6430573F11I | red | -1.3744863 | 1.14E-05 | -0.862346 | 0.00661722 |
| Prex1 | orange | 1.79870457 | 2.73E-17 | 1.66536908 | 8.67E-15 |
| Hs3st6 | red | -1.4410537 | 0.00056667 | -1.0604865 | 0.01512422 |
| Hnrnpu | orange | 0.5818189 | 8.17E-05 | 0.54856112 | 0.00030322 |
| Ccdc151 | red | -1.3417859 | 4.41E-10 | -0.9214497 | 2.91E-05 |
| Lonrf1 | red | -0.7897959 | 1.38E-06 | -0.907211 | 5.12E-08 |
| Zfp189 | red | -0.7043777 | 0.00607258 | -0.4248762 | 0.12170264 |
| Coro7 | orange | 1.16951657 | 5.87E-15 | 1.01122775 | 3.21E-11 |
| Ccbl1 | red | -1.0765192 | 1.45E-05 | -0.8140329 | 0.00155127 |
| Cpeb3 | red | -0.776636 | 1.25E-05 | -0.6709926 | 0.00027264 |
| Dusp26 | skyblue | -1.952643 | 0.00018152 | -1.7308224 | 0.00089966 |
| Kcne2 | skyblue | -2.0574264 | 3.40E-06 | -1.0138253 | 0.03044067 |
| Capsl | skyblue | -1.6007931 | 2.10E-09 | -1.0140941 | 0.00024731 |
| Lap3 | orange | 1.67821637 | 3.68E-24 | 1.59089081 | 1.02E-21 |
| Ncoa7 | orange | 0.66316862 | 9.85E-05 | 0.78983211 | 4.90E-06 |
| Batf2 | yellow | 4.02520941 | 2.73E-75 | 4.11272569 | 9.44E-79 |
| Ldb2 | red | -0.8224672 | 1.58E-06 | -0.6396728 | 0.0003138 |
| Plekhg5 | red | -1.1057443 | 8.27E-11 | -0.9458066 | 5.44E-08 |
| Wdr34 | red | -0.9468567 | 2.91E-12 | -0.881527 | 1.45E-10 |
| Slx4 | orange | 0.86498296 | 4.51E-05 | 0.81254365 | 0.00018617 |
| Bahcc1 | red | -1.1289389 | 5.20E-06 | -0.8489638 | 0.000938 |

| | | | | | |
|---|---|---|---|---|---|
| Orai2 | orange | 1.0776144 | 9.87E-14 | 1.09099951 | 5.66E-14 |
| Exo1 | yellow | 3.22143289 | 6.44E-13 | 2.31340936 | 1.03E-07 |
| Alkbh4 | orange | 0.50489331 | 0.00011414 | 0.6093587 | 5.73E-06 |
| Dnajc28 | skyblue | -1.9942291 | 0.00053881 | -1.9833992 | 0.00078964 |
| Cc2d2a | red | -1.1914336 | 1.39E-09 | -0.7756954 | 0.00013993 |
| Galnt12 | orange | 1.05297746 | 0.00012602 | 1.1047214 | 9.63E-05 |
| Cep131 | red | -0.6527293 | 7.30E-06 | -0.4229734 | 0.00548159 |
| Kmo | orange | 0.71705062 | 0.0148362 | 1.49945734 | 3.41E-07 |
| Tbc1d2 | orange | 0.7740339 | 0.00169425 | 1.12099433 | 5.64E-06 |
| Xkr5 | yellow | 3.52380196 | 1.20E-06 | 3.23069936 | 8.96E-06 |
| Nhsl1 | red | -1.0122812 | 1.93E-08 | -0.3729895 | 0.05640137 |
| Mcph1 | orange | 0.7638606 | 0.00032885 | 0.77827023 | 0.00040283 |
| Endov | red | -0.718085 | 3.03E-06 | -0.6273886 | 6.93E-05 |
| Rere | red | -0.7950968 | 7.50E-06 | -0.5943043 | 0.00125425 |
| Trim14 | orange | 1.13826342 | 1.43E-07 | 0.92416393 | 3.29E-05 |
| Slc44a3 | skyblue | -1.6152131 | 3.15E-10 | -1.3266253 | 4.42E-07 |
| Neurl2 | red | -0.924689 | 1.46E-05 | -1.0017167 | 7.99E-06 |
| Lrrc17 | skyblue | -2.991089 | 1.53E-32 | -2.3799556 | 2.06E-22 |
| Alg14 | red | -0.8996346 | 1.55E-08 | -0.7846374 | 1.40E-06 |
| Txlnb | skyblue | -1.0338666 | 0.00057 | -1.3725881 | 6.14E-06 |
| Fgl2 | yellow | 2.23251646 | 4.35E-19 | 1.89346853 | 8.06E-14 |
| Gpr37 | skyblue | -2.9222265 | 0.00722594 | -3.5027515 | 0.00219308 |
| Slc26a11 | red | -0.7784408 | 6.68E-09 | -0.5774262 | 3.24E-05 |
| Spsb1 | orange | 0.69500887 | 8.26E-07 | 0.36782739 | 0.01345297 |
| Coq10a | red | -0.747492 | 5.12E-08 | -0.7402506 | 1.37E-07 |
| Urb1 | orange | 0.69157199 | 7.09E-06 | 0.60705338 | 0.00012682 |
| Pik3cd | orange | 2.0084118 | 5.73E-24 | 1.86739282 | 9.41E-21 |
| Ptger4 | yellow | 2.19050978 | 1.96E-19 | 2.07216139 | 3.03E-17 |
| Dag1 | red | -0.614692 | 0.00080168 | -0.3418459 | 0.08182587 |
| Clstn1 | red | -1.211619 | 6.90E-09 | -0.8648078 | 6.59E-05 |
| Stk32a | red | -1.0325545 | 0.00521796 | -0.4486214 | 0.26487625 |
| Mettl20 | red | -1.1265811 | 1.18E-08 | -0.9344544 | 4.13E-06 |
| Rhou | orange | 0.75456346 | 0.00027417 | 0.62136636 | 0.00386717 |
| Ccdc40 | skyblue | -1.5597109 | 1.03E-10 | -0.8636278 | 0.0005727 |
| Tbc1d16 | red | -1.2156837 | 4.34E-08 | -0.9903305 | 1.44E-05 |
| Zc3h12d | yellow | 3.25515687 | 3.86E-28 | 3.28829242 | 1.79E-28 |
| Dtx4 | orange | 0.74572888 | 0.00265136 | 0.90449713 | 0.00032401 |
| Phtf2 | red | -1.2024061 | 2.47E-08 | -0.6557669 | 0.00406744 |
| Timeless | orange | 1.2171242 | 3.11E-11 | 1.06290685 | 1.30E-08 |
| Ifi203 | yellow | 2.30067562 | 9.82E-35 | 2.07180765 | 3.68E-28 |
| Magi2 | skyblue | -1.3510001 | 3.39E-07 | -1.3370397 | 1.12E-06 |
| Gnaz | red | -1.2269226 | 8.96E-07 | -0.6074894 | 0.02456797 |
| Slc7a5 | yellow | 2.24731415 | 5.36E-25 | 1.96605456 | 2.74E-19 |

| | | | | | |
|---|---|---|---|---|---|
| Ptger3 | skyblue | -1.5057221 | 4.47E-05 | -1.1687332 | 0.00214336 |
| Saa3 | black | 8.62825322 | 2.38E-192 | 8.054942 | 4.19E-173 |
| Stat2 | yellow | 2.2683768 | 4.91E-43 | 2.19912052 | 1.57E-40 |
| Nup43 | orange | 1.02011422 | 0.00083326 | 0.71121185 | 0.02795735 |
| Negr1 | skyblue | -2.8267826 | 1.58E-07 | -2.0146434 | 0.00026451 |
| Ift88 | red | -0.9553746 | 3.66E-08 | -0.6756364 | 0.00014729 |
| Gjb6 | red | -1.0405565 | 0.00081261 | -0.7563478 | 0.01989989 |
| Bub1b | yellow | 2.49196379 | 5.40E-27 | 2.47512638 | 3.16E-26 |
| Tnni3k | skyblue | -1.7379946 | 0.08416401 | -2.6295825 | 0.00923128 |
| Bmf | red | -0.9181501 | 2.60E-05 | -0.7089293 | 0.00173207 |
| Klhl42 | red | -0.7048993 | 1.13E-08 | -0.5475631 | 1.56E-05 |
| Mettl11b | skyblue | -2.4458784 | 0.00307405 | -1.1974563 | 0.20211957 |
| Cacna2d1 | red | -1.038095 | 3.35E-07 | -0.8644655 | 3.98E-05 |
| Gpr176 | yellow | 1.82512214 | 0.00090866 | 2.05820497 | 0.00034675 |
| Abcc8 | skyblue | -1.4816525 | 0.17678111 | -2.7770117 | 0.00992267 |
| 9430038I01R | orange | 0.90224361 | 3.67E-06 | 0.7836659 | 0.00011462 |
| Rgl3 | red | -0.946591 | 5.41E-09 | -0.5738539 | 0.00067076 |
| Maob | skyblue | -1.7953253 | 1.40E-10 | -1.3834569 | 1.55E-06 |
| Hs2st1 | red | -1.0174466 | 3.42E-27 | -0.7745447 | 6.37E-16 |
| Thbs1 | yellow | 1.99165898 | 0.00013194 | 2.36111449 | 7.74E-06 |
| Cd209c | skyblue | -5.0270131 | 1.63E-08 | -1.4632144 | 0.01013972 |
| Fmo2 | red | -1.1005244 | 0.0025615 | -1.1902583 | 0.00138198 |
| Alkbh3 | orange | 0.5954988 | 6.57E-07 | 0.52267393 | 2.35E-05 |
| Fmo1 | skyblue | -2.0421631 | 1.38E-07 | -1.6822958 | 2.50E-05 |
| Ankrd6 | red | -1.1368793 | 3.86E-05 | -0.6639783 | 0.01959683 |
| Scamp2 | orange | 0.78298446 | 1.02E-06 | 0.57891832 | 0.00051781 |
| Ccdc114 | red | -1.1202132 | 7.98E-14 | -0.5271569 | 0.00073214 |
| Tmem194 | orange | 0.66839594 | 1.48E-05 | 0.48589841 | 0.00260586 |
| 2810417H13 | yellow | 3.41269032 | 2.03E-09 | 3.73921952 | 9.55E-11 |
| Zfp704 | red | -1.3161964 | 2.19E-12 | -1.0575934 | 3.59E-08 |
| Emp3 | orange | 1.37079059 | 2.81E-07 | 1.00910797 | 0.00027433 |
| Ttc12 | red | -0.9290334 | 4.59E-08 | -0.8305925 | 1.61E-06 |
| Tm7sf3 | red | -0.8043816 | 1.14E-08 | -0.6630216 | 4.66E-06 |
| Tbc1d10c | yellow | 2.17870252 | 8.62E-40 | 2.27349575 | 6.41E-42 |
| Lrp1 | orange | 0.6329984 | 0.00060824 | 0.56464278 | 0.00302247 |
| Gbp7 | yellow | 2.77432941 | 3.62E-59 | 2.57297902 | 5.46E-51 |
| Sema3d | red | -0.7817038 | 0.00014724 | -0.5293837 | 0.01431699 |
| Daam2 | red | -0.964222 | 0.00030814 | -0.7797401 | 0.00486138 |
| Klhdc4 | orange | 0.76546484 | 1.92E-07 | 0.61912439 | 4.65E-05 |
| Gbp2b | black | 4.46208756 | 1.19E-20 | 4.77572561 | 4.85E-19 |
| Dnm3 | red | -0.6971648 | 0.00035963 | -0.4687385 | 0.0225338 |
| Pacsin1 | yellow | 2.24413265 | 2.17E-11 | 3.02051865 | 2.20E-18 |
| Ndufa4l2 | skyblue | -1.2515657 | 7.68E-06 | -1.2334984 | 1.66E-05 |

| | | | | | |
|---|---|---|---|---|---|
| BC052040 | orange | 0.43103732 | 0.09130012 | 0.75595036 | 0.0034709 |
| Btnl9 | skyblue | -2.0818507 | 3.46E-05 | -1.7256596 | 0.00110893 |
| Stac3 | red | -1.4090916 | 1.52E-06 | -0.8230706 | 0.00838504 |
| Hey1 | skyblue | -2.7539914 | 3.22E-16 | -2.3105488 | 1.53E-11 |
| Ddx58 | orange | 1.60102936 | 7.57E-29 | 1.52007155 | 5.44E-26 |
| Btbd16 | yellow | 1.10295792 | 0.20540674 | 3.28271889 | 0.0009792 |
| Cchcr1 | orange | 0.50070726 | 0.00240008 | 0.6133954 | 0.00025595 |
| Olfr56 | black | 4.91275807 | 1.02E-78 | 4.61386871 | 5.16E-74 |
| Arhgap9 | yellow | 2.40849241 | 5.43E-33 | 2.19929939 | 1.94E-27 |
| Trim7 | red | -0.7485767 | 0.00674957 | -0.2922888 | 0.35438601 |
| Mars | orange | 0.52412777 | 8.20E-05 | 0.6407037 | 1.93E-06 |
| Lyrm5 | red | -1.1731919 | 1.19E-11 | -1.2141401 | 5.91E-12 |
| Klhl32 | red | -1.5339884 | 0.0010417 | -0.3055601 | 0.54670268 |
| 5330417C22 | skyblue | -2.1018663 | 1.81E-09 | -1.2519782 | 0.00060223 |
| Slc25a28 | orange | 0.63993738 | 1.87E-05 | 0.64447279 | 2.47E-05 |
| Plekha4 | orange | 0.81853351 | 0.0001761 | 1.15353402 | 1.38E-07 |
| Pitpnc1 | red | -0.7231455 | 5.09E-08 | -0.7335009 | 5.48E-08 |
| Slc26a10 | red | -0.6396854 | 0.0071876 | -1.324502 | 7.74E-08 |
| Spns2 | red | -0.9943052 | 3.97E-22 | -0.6530502 | 7.23E-10 |
| Mybbp1a | orange | 0.87905035 | 2.46E-06 | 0.7382568 | 0.00012448 |
| Blvrb | orange | 0.68102757 | 0.0003366 | 0.52176827 | 0.00840168 |
| Ggt6 | red | -0.9715585 | 0.00276872 | -0.9652557 | 0.00364019 |
| Cfap69 | red | -0.8664227 | 0.00049652 | -0.6982231 | 0.00655836 |
| Xaf1 | yellow | 2.74259972 | 1.07E-58 | 2.72889412 | 5.24E-58 |
| Ltbp4 | skyblue | -1.9492635 | 4.57E-09 | -1.3350469 | 0.00011108 |
| Chrm4 | yellow | 2.97546635 | 0.00023622 | 1.89101012 | 0.01359585 |
| Pvr | orange | 1.5558691 | 7.77E-09 | 1.48397164 | 6.30E-08 |
| Tlr8 | yellow | 2.75194516 | 1.89E-06 | 2.0585181 | 0.00052424 |
| Milr1 | yellow | 3.41105696 | 1.64E-18 | 2.84871163 | 4.65E-13 |
| Adam22 | skyblue | -1.9130149 | 2.61E-10 | -1.3793355 | 7.47E-06 |
| Pitpnm3 | red | -1.2554803 | 8.24E-05 | -0.735788 | 0.02926869 |
| C3ar1 | yellow | 4.43027159 | 1.29E-12 | 3.75473846 | 3.67E-09 |
| Wbscr27 | red | -0.9332904 | 0.00939025 | -0.6417261 | 0.09188018 |
| Gstm2 | red | -0.7634644 | 0.00020689 | -0.4929867 | 0.02289136 |
| Lppr2 | red | -0.8643932 | 2.73E-08 | -0.8182461 | 1.72E-07 |
| Apoc1 | red | -1.0525612 | 0.00588886 | -0.7765178 | 0.05348277 |
| Rundc3b | orange | 1.69642684 | 3.66E-05 | 1.49878096 | 0.00033929 |
| Abcb1a | red | -0.720323 | 4.57E-09 | -0.6614578 | 1.21E-07 |
| 1110051M20 | red | -0.9390723 | 4.56E-07 | -0.6073389 | 0.00161691 |
| Bace2 | red | -0.8286394 | 3.10E-06 | -0.7979466 | 1.42E-05 |
| Apobec1 | yellow | 2.74722857 | 3.24E-18 | 2.30590363 | 5.81E-13 |
| Tmem51 | orange | 1.18317621 | 2.34E-05 | 0.73493417 | 0.01268095 |
| Aicda | yellow | 5.14521619 | 0.00114055 | 0.77870785 | 0.5268967 |

| | | | | | |
|---|---|---|---|---|---|
| Oaz2 | red | -1.1845815 | 6.32E-18 | -0.8942702 | 2.08E-10 |
| Ppp1r14c | skyblue | -2.1902343 | 2.51E-10 | -1.5092292 | 2.60E-05 |
| Efhd2 | orange | 1.83400115 | 5.21E-12 | 1.59275173 | 4.16E-09 |
| Sh3bgr | skyblue | -2.3926557 | 4.04E-16 | -1.7032595 | 1.67E-08 |
| Mthfd1l | orange | 1.22665968 | 2.53E-09 | 0.81556594 | 0.00013656 |
| Kremen2 | orange | 1.65750656 | 9.71E-07 | 1.0741463 | 0.00126509 |
| Tbl3 | orange | 0.71736966 | 7.64E-06 | 0.77151443 | 2.16E-06 |
| Col16a1 | red | -0.7329452 | 1.42E-07 | -0.4683999 | 0.0012845 |
| Slco4c1 | skyblue | -1.5816269 | 2.22E-06 | -0.8202283 | 0.02066076 |
| Apobec2 | skyblue | -2.4339855 | 1.76E-08 | -1.4475375 | 0.00139247 |
| Limd2 | orange | 0.93357235 | 6.74E-14 | 0.85457378 | 1.51E-11 |
| Ap1g2 | orange | 0.76332342 | 2.16E-10 | 0.61784531 | 6.29E-07 |
| Cyp2s1 | skyblue | -2.0711566 | 1.41E-21 | -1.3866125 | 4.99E-10 |
| A930016O22 | skyblue | -1.1684159 | 0.03683756 | -1.5358008 | 0.00849792 |
| Sh3pxd2b | orange | 1.76776139 | 2.40E-07 | 1.43283017 | 4.86E-05 |
| Il17rd | red | -1.1603213 | 1.14E-05 | -0.8153939 | 0.00306084 |
| Zfhx2 | red | -0.6720212 | 0.00037621 | -0.1991544 | 0.34888193 |
| Kcna2 | red | -1.0248684 | 0.00030733 | -0.917261 | 0.00171657 |
| Cep126 | skyblue | -2.0226396 | 3.15E-15 | -1.1082199 | 2.20E-05 |
| Rnf167 | red | -0.8092253 | 0.00021152 | -0.6099588 | 0.00728853 |
| Cd53 | yellow | 2.88043622 | 2.79E-28 | 2.85846207 | 1.18E-27 |
| Lat2 | yellow | 2.24216584 | 1.61E-10 | 2.24531885 | 2.70E-10 |
| Cmtm5 | skyblue | -2.2863031 | 0.00011561 | -2.0261973 | 0.00087998 |
| Snrnp25 | orange | 0.76353374 | 3.48E-06 | 0.41771481 | 0.01769114 |
| Ttc3 | red | -0.7920046 | 1.98E-08 | -0.6945499 | 1.67E-06 |
| S100g | skyblue | -2.0331265 | 2.77E-05 | -1.5082617 | 0.00275131 |
| Chil3 | orange | 1.54059738 | 2.53E-05 | 1.03766106 | 0.00667395 |
| Agbl2 | skyblue | -1.6204288 | 3.35E-07 | -1.0092212 | 0.00194617 |
| Snrpd2 | orange | 0.62275427 | 0.00169729 | 0.45547501 | 0.02879462 |
| Zmynd15 | orange | 2.04741561 | 1.17E-12 | 1.62286916 | 3.96E-08 |
| Gpr161 | red | -0.5877994 | 0.00180137 | -0.0965008 | 0.66403357 |
| Six5 | skyblue | -1.8009515 | 1.21E-21 | -1.4286988 | 8.05E-14 |
| Plekhh2 | skyblue | -1.5532294 | 2.56E-07 | -0.8594902 | 0.0066853 |
| Reps2 | skyblue | -1.8141344 | 2.66E-08 | -1.5014575 | 7.62E-06 |
| Crocc | red | -0.8462804 | 1.30E-08 | -0.5646666 | 0.00026115 |
| Osbpl10 | red | -0.6139805 | 0.0048728 | -0.4266658 | 0.06227677 |
| Tmem205 | red | -0.9172364 | 1.68E-07 | -0.861392 | 1.54E-06 |
| Kcnd3 | red | -1.3190983 | 0.00056826 | -0.4010061 | 0.33114872 |
| Atp1a3 | yellow | 2.27453902 | 0.00011473 | 1.91176794 | 0.00169985 |
| Ulk4 | skyblue | -1.758092 | 6.95E-11 | -1.0854079 | 7.84E-05 |
| Slc16a11 | red | -1.4706786 | 1.43E-07 | -0.973517 | 0.00081858 |
| Tet2 | orange | 0.60014635 | 1.35E-06 | 0.5884606 | 4.00E-06 |
| Mgl2 | skyblue | -2.2136844 | 3.72E-08 | -1.356838 | 0.00118467 |

| | | | | | |
|---|---|---|---|---|---|
| Cables1 | red | -1.1181582 | 4.51E-10 | -0.82276 | 8.81E-06 |
| Arhgef38 | skyblue | -1.965468 | 1.14E-08 | -1.2020144 | 0.00077239 |
| Gm11992 | skyblue | -1.8707051 | 1.56E-06 | -1.5013125 | 0.00012464 |
| Mill2 | red | -1.1325736 | 1.86E-06 | -0.9232047 | 0.00016726 |
| Sh3kbp1 | orange | 1.12441443 | 4.71E-09 | 0.97004246 | 8.62E-07 |
| Abhd4 | red | -0.761392 | 3.21E-05 | -0.6156805 | 0.00113917 |
| Npnt | red | -1.5020795 | 1.29E-07 | -1.0233897 | 0.00055059 |
| Cmtm8 | red | -1.3124614 | 2.35E-14 | -0.9827835 | 2.66E-08 |
| Iffo2 | orange | 0.60499975 | 0.01850126 | 0.74010867 | 0.004382 |
| Fam117b | red | -0.0767584 | 0.70193805 | -0.5805902 | 0.00106069 |
| Ramp3 | orange | 0.9255248 | 6.60E-05 | 0.72399813 | 0.00263113 |
| Wdr43 | orange | 0.81519741 | 1.71E-07 | 0.75359958 | 2.41E-06 |
| Wwp1 | red | -0.8777518 | 3.46E-05 | -0.4996452 | 0.02603649 |
| Mslnl | yellow | 2.5584806 | 0.00169479 | 3.35972077 | 0.00010002 |
| Pif1 | black | 3.49334886 | 2.71E-21 | 5.11012889 | 2.79E-28 |
| Nacad | red | -1.0040573 | 0.00017045 | -0.5463491 | 0.05054142 |
| Fzd7 | red | -1.3010408 | 6.93E-12 | -1.0263084 | 1.36E-07 |
| Rwdd2b | orange | 0.94035988 | 4.00E-07 | 0.99062645 | 2.50E-07 |
| Ostc | orange | 0.66676972 | 0.00072569 | 0.54478288 | 0.00795784 |
| Ccdc8 | red | -1.0486953 | 1.17E-15 | -0.9414453 | 1.44E-12 |
| Nbl1 | red | -0.9800886 | 1.58E-11 | -0.6896939 | 4.58E-06 |
| Cyyr1 | red | -0.7225879 | 3.59E-06 | -0.6141563 | 0.00013281 |
| Ripk2 | orange | 0.81730681 | 1.75E-10 | 0.73376491 | 3.00E-08 |
| Tmco4 | orange | 1.12165666 | 8.35E-08 | 1.22049818 | 8.66E-09 |
| Dnah7b | red | -1.4287559 | 3.75E-08 | -0.5166322 | 0.05814212 |
| Brca2 | orange | 0.91422911 | 0.00218258 | 0.82616622 | 0.00793612 |
| Pla2g5 | skyblue | -0.5453569 | 0.2929509 | -3.1022589 | 1.04E-07 |
| 2310036O22 | orange | 0.63482289 | 4.04E-07 | 0.5639929 | 1.20E-05 |
| Arhgap11a | yellow | 2.46929768 | 1.11E-25 | 2.40526737 | 5.07E-24 |
| Elovl6 | red | -0.9137916 | 0.00764667 | -0.9802825 | 0.00498585 |
| Chd7 | orange | 1.61444126 | 1.04E-16 | 1.53309238 | 5.44E-15 |
| Rbbp8 | orange | 1.13013392 | 1.44E-13 | 0.591153 | 0.00023247 |
| Lamp3 | red | -0.9096243 | 0.0001911 | -0.6837891 | 0.00709476 |
| Car8 | skyblue | -1.9776577 | 4.93E-15 | -1.3932935 | 8.37E-08 |
| Tox | yellow | 2.2120317 | 1.97E-29 | 2.07690727 | 5.53E-24 |
| Katnal1 | red | -0.7309089 | 1.36E-06 | -0.5181952 | 0.00095638 |
| Slc7a1 | orange | 1.45829852 | 1.85E-09 | 1.39580127 | 1.48E-08 |
| Thoc6 | orange | 1.1012044 | 4.79E-06 | 1.30171295 | 9.65E-08 |
| Ak7 | skyblue | -2.5291161 | 6.84E-15 | -1.5665001 | 2.83E-06 |
| Inhba | orange | 1.25112586 | 0.0015716 | 1.39375402 | 0.00059356 |
| Atp1b2 | skyblue | -1.458674 | 5.47E-13 | -1.2679024 | 9.67E-10 |
| Ankrd42 | red | -1.0974228 | 2.69E-11 | -0.682174 | 5.49E-05 |
| Wrap53 | orange | 0.61871516 | 1.53E-05 | 0.47821762 | 0.00121424 |

| | | | | | |
|---|---|---|---|---|---|
| Bdkrb1 | orange | 1.33212613 | 3.02E-05 | 1.44163394 | 1.17E-05 |
| Ssr2 | orange | 0.62979448 | 0.00047436 | 0.57130222 | 0.00210403 |
| Myzap | red | -1.1945491 | 0.00010991 | -0.9565439 | 0.00278619 |
| BC055324 | orange | 1.82468284 | 4.79E-07 | 1.45325953 | 9.43E-05 |
| Paqr6 | skyblue | -1.2981711 | 6.32E-06 | -1.3673624 | 3.76E-06 |
| Hibch | red | -0.8441047 | 2.17E-07 | -0.6999718 | 3.44E-05 |
| Nthl1 | red | -0.7597312 | 1.08E-06 | -0.5888631 | 0.00027228 |
| Ccnb1 | yellow | 3.48102365 | 3.05E-40 | 3.80900097 | 1.01E-44 |
| Cirh1a | orange | 0.75476563 | 9.42E-05 | 0.68639014 | 0.00057818 |
| Mfsd6 | red | -0.6608748 | 1.68E-08 | -0.4848338 | 6.28E-05 |
| Arhgap32 | red | -0.764034 | 0.00011523 | -0.5145808 | 0.01306643 |
| Mmrn2 | red | -0.7016293 | 7.38E-07 | -0.6044473 | 3.29E-05 |
| Serpina3h | yellow | 3.66397567 | 1.89E-18 | 3.39128305 | 5.60E-16 |
| Cacna2d4 | skyblue | -1.8928689 | 4.37E-06 | -1.0457659 | 0.01520513 |
| Smpx | skyblue | -1.9008906 | 1.20E-05 | -1.7172258 | 0.00013058 |
| Serpina3g | black | 5.13266768 | 2.39E-64 | 4.23650002 | 1.48E-44 |
| Piezo2 | red | -0.8328795 | 0.00012214 | -0.6596808 | 0.00335764 |
| Zfp281 | orange | 0.95099592 | 1.11E-07 | 1.00931535 | 2.75E-08 |
| Kif14 | yellow | 3.33652619 | 5.75E-25 | 4.0069147 | 7.79E-30 |
| Rrp9 | orange | 0.56514755 | 0.00089739 | 0.58175142 | 0.00085121 |
| Irf8 | yellow | 2.36035004 | 3.47E-39 | 2.20259036 | 3.90E-34 |
| Upk2 | skyblue | -2.1786311 | 0.00242996 | -1.6602286 | 0.0255686 |
| Ago1 | red | -0.8148844 | 7.25E-09 | -0.6460194 | 8.54E-06 |
| H2-Ob | red | -0.6648722 | 0.00131658 | -0.176518 | 0.46208699 |
| Fmod | skyblue | -1.8215031 | 7.91E-05 | -1.4972188 | 0.00166002 |
| Sepw1 | orange | 0.71950511 | 0.00257337 | 0.55510835 | 0.02598613 |
| Prelp | skyblue | -1.8077953 | 1.51E-07 | -1.3367298 | 0.00017879 |
| Sdk2 | orange | 0.75424993 | 1.96E-05 | 0.85292475 | 2.03E-06 |
| Cdc42ep4 | orange | 0.84271986 | 4.57E-07 | 0.60893272 | 0.00045024 |
| 5730559C18 | orange | 0.7333982 | 0.01162718 | 0.82185727 | 0.00565212 |
| Entpd3 | orange | -0.00609 | 0.98388608 | 0.93905786 | 0.00012689 |
| Ccdc74a | red | -0.8477757 | 0.00046377 | -0.313239 | 0.23738202 |
| Kctd12b | red | -1.4796499 | 1.35E-06 | -0.9870997 | 0.00191748 |
| Kif21b | yellow | 1.94261852 | 4.84E-32 | 2.10648691 | 2.85E-37 |
| Slc5a12 | skyblue | -3.9314971 | 0.00138414 | -3.5416472 | 0.00482379 |
| Ddx24 | orange | 0.64142912 | 1.73E-13 | 0.50512867 | 1.49E-08 |
| Pcca | red | -0.6217989 | 0.00028957 | -0.5633745 | 0.00144874 |
| Pnpla3 | skyblue | -2.3962698 | 7.50E-07 | -2.3409452 | 2.12E-06 |
| Bbox1 | skyblue | -2.5787129 | 0.00080013 | -2.2053711 | 0.00392089 |
| Prima1 | skyblue | -0.8735071 | 0.16276005 | -2.3654316 | 0.00108359 |
| Lrrc18 | red | -1.488588 | 3.90E-05 | -0.519314 | 0.15189185 |
| Fcho2 | red | -0.5909195 | 0.00141886 | -0.5103569 | 0.00769887 |
| Amot | red | -0.9861762 | 8.14E-06 | -0.5142081 | 0.0304049 |

| | | | | | |
|---|---|---|---|---|---|
| Rasl12 | red | -0.9307903 | 1.71E-06 | -0.7172061 | 0.00037272 |
| Lhfpl1 | skyblue | -1.5282218 | 0.02412405 | -2.7044234 | 0.00083942 |
| Alg13 | orange | 0.68260593 | 0.00187099 | 0.63695772 | 0.00479434 |
| Coro2b | red | -0.6616428 | 0.00193284 | -0.56252 | 0.01089993 |
| Pgm5 | skyblue | -1.6381494 | 5.27E-13 | -1.4196182 | 8.28E-10 |
| Tspo | orange | 1.50517378 | 1.19E-10 | 1.3098496 | 4.11E-08 |
| Tmem45b | skyblue | -4.028869 | 3.14E-06 | -2.0054651 | 0.02567331 |
| Trem3 | yellow | 3.65901806 | 4.81E-11 | 2.16624176 | 2.23E-05 |
| Plekha6 | skyblue | -1.6682458 | 4.35E-13 | -0.9919914 | 3.66E-05 |
| Gpr155 | skyblue | -1.6577961 | 3.88E-07 | -1.7057042 | 2.65E-07 |
| Ppp2r2d | orange | 0.6759113 | 4.86E-06 | 0.45543971 | 0.0032073 |
| Enc1 | orange | 1.70031364 | 1.44E-08 | 1.7360282 | 1.10E-08 |
| Ydjc | orange | 0.77581011 | 0.02399433 | 1.1386251 | 0.0009609 |
| Mapk1ip1 | red | -0.8225058 | 6.49E-05 | -0.7501472 | 0.00037093 |
| Cpsf2 | orange | 0.86960537 | 6.91E-08 | 0.62741098 | 0.00018236 |
| 2700046A07I | skyblue | -4.51352 | 0.00032701 | -4.9156344 | 0.00010578 |
| Abca9 | skyblue | -1.5853134 | 1.39E-05 | -1.5527104 | 3.18E-05 |
| Gck | red | -1.4736752 | 4.56E-06 | -0.4802 | 0.14505033 |
| Efhc1 | skyblue | -1.8210402 | 2.88E-08 | -1.3124045 | 0.00010005 |
| Oasl1 | black | 5.37644034 | 9.18E-62 | 5.44623564 | 1.61E-62 |
| Abca8a | skyblue | -1.9398234 | 1.52E-10 | -1.5423637 | 7.21E-07 |
| Sytl3 | yellow | 3.27310583 | 5.20E-18 | 3.60862494 | 1.14E-20 |
| Ptpre | orange | 0.55843147 | 0.0042821 | 0.72534841 | 0.00025893 |
| Fhdc1 | skyblue | -1.7997563 | 1.75E-10 | -1.1819354 | 5.12E-05 |
| Mcm3 | orange | 1.75352205 | 7.99E-19 | 1.54202466 | 1.50E-14 |
| Slc16a6 | orange | 1.48520318 | 3.37E-08 | 1.51753737 | 2.91E-08 |
| Rnpep | orange | 0.68780758 | 6.35E-07 | 0.62360695 | 1.13E-05 |
| Fam222a | skyblue | -1.9671906 | 1.60E-08 | -1.3668991 | 0.00012099 |
| Tnfrsf18 | yellow | 2.61654242 | 1.82E-33 | 2.0348614 | 1.74E-22 |
| Pkp2 | red | -0.6301592 | 0.00127217 | -0.522042 | 0.01025288 |
| S100a10 | orange | 1.14803865 | 3.90E-11 | 0.95101812 | 9.29E-08 |
| Znrf3 | red | -0.6093104 | 0.0001465 | -0.2259256 | 0.20412044 |
| Mettl8 | red | -0.9445608 | 8.70E-06 | -0.5239883 | 0.01949919 |
| Elmod1 | skyblue | -1.9236858 | 8.08E-07 | -1.1527837 | 0.00364824 |
| Foxn4 | orange | 1.72220071 | 0.00018116 | 1.6339587 | 0.00021544 |
| Acacb | skyblue | -1.3282737 | 4.91E-06 | -1.3299726 | 7.42E-06 |
| Ncapg2 | yellow | 2.79065277 | 1.61E-23 | 2.48983105 | 1.38E-18 |
| Lce3b | red | -0.9402408 | 0.0047591 | -0.6576249 | 0.06252158 |
| Igsf3 | red | -0.7300223 | 1.82E-05 | -0.3612899 | 0.04913914 |
| Sln | red | -0.611215 | 0.1298588 | -1.4097715 | 0.0003155 |
| Myo3b | orange | 1.82107076 | 0.00039106 | 0.82292459 | 0.06755222 |
| Abhd14b | skyblue | -1.4030875 | 7.49E-10 | -1.2378384 | 8.73E-08 |
| Hnrnpf | orange | 0.58398767 | 3.41E-05 | 0.41824823 | 0.0044326 |

| | | | | | |
|---|---|---|---|---|---|
| Arsb | orange | 1.0055214 | 1.04E-05 | 0.58766746 | 0.01480186 |
| 4933440N22 | orange | 1.67989165 | 1.78E-05 | 1.46016514 | 0.0003044 |
| Lce1c | red | -1.1229565 | 0.00053486 | -0.8515846 | 0.01163667 |
| Kank3 | red | -0.9607795 | 5.87E-05 | -0.8743555 | 0.00037751 |
| Uggt2 | red | -0.6305264 | 0.00498818 | -0.2272022 | 0.36453992 |
| Klhdc8a | orange | 1.79168466 | 5.82E-07 | 1.79238216 | 9.32E-07 |
| Vwa1 | red | -0.9543735 | 3.66E-08 | -0.4420948 | 0.01672506 |
| Lce1f | skyblue | -2.1900285 | 2.92E-08 | -1.1679667 | 0.00446923 |
| Rassf4 | orange | 0.59838275 | 0.00098266 | 0.34832573 | 0.07317261 |
| Klhl23 | red | -0.986448 | 5.01E-06 | -0.7334916 | 0.00106706 |
| Dzip1 | red | -1.2234345 | 2.81E-05 | -0.7156715 | 0.01933886 |
| Papd4 | orange | 0.70642859 | 7.80E-05 | 0.50563841 | 0.00677757 |
| Tekt3 | skyblue | -1.9206196 | 0.00115542 | -1.3319072 | 0.02297122 |
| Slc35f2 | yellow | 2.08198751 | 9.11E-10 | 1.85553278 | 6.09E-08 |
| Chchd7 | red | -0.5111744 | 0.00011853 | -0.5964315 | 1.14E-05 |
| Abhd14a | red | -0.7024448 | 2.00E-05 | -0.6617667 | 9.51E-05 |
| Zfand4 | orange | 0.78125223 | 0.00131581 | 0.59474098 | 0.01605276 |
| Bag2 | red | -0.786432 | 2.33E-07 | -0.5176975 | 0.00090838 |
| Sgsm1 | red | -1.330397 | 2.09E-16 | -0.5189597 | 0.00185448 |
| Ammecr1 | orange | 0.62531919 | 0.00533573 | 0.35916837 | 0.13829776 |
| Lyn | orange | 0.64618726 | 0.00154894 | 0.50376732 | 0.01801915 |
| Pm20d1 | skyblue | -1.9338839 | 5.66E-06 | -1.3637419 | 0.00174723 |
| Cilp | skyblue | -2.7471217 | 1.53E-18 | -1.0653708 | 0.00133636 |
| Ccr8 | black | 5.12638906 | 5.13E-24 | 2.82658223 | 4.19E-10 |
| Trem1 | yellow | 4.21479088 | 3.65E-08 | 4.36597265 | 2.52E-09 |
| Fam92b | skyblue | -1.8356634 | 6.03E-05 | -1.231626 | 0.00986802 |
| Nxt2 | red | -0.7518577 | 5.37E-07 | -0.7576798 | 6.82E-07 |
| Ehbp1 | red | -0.6958875 | 0.00022316 | -0.7852289 | 4.28E-05 |
| Tnfrsf14 | orange | 1.16325605 | 1.11E-08 | 1.16669893 | 1.44E-08 |
| Ctf1 | red | -0.871154 | 0.00020364 | -0.5412291 | 0.0311547 |
| Ubash3a | yellow | 4.42596379 | 1.04E-31 | 4.2908811 | 5.80E-29 |
| Arl15 | skyblue | -1.2696501 | 3.39E-18 | -1.3699593 | 1.29E-20 |
| Ikbke | yellow | 2.13626144 | 4.71E-21 | 1.96158077 | 7.97E-18 |
| Arel1 | orange | 0.66281244 | 3.21E-06 | 0.63707482 | 1.18E-05 |
| Grap2 | yellow | 2.23265548 | 8.73E-31 | 2.23422084 | 1.61E-30 |
| Gnl3 | orange | 1.13908758 | 1.59E-07 | 1.14352183 | 2.32E-07 |
| Osbpl6 | skyblue | -2.2331156 | 5.31E-09 | -1.4728335 | 0.00021138 |
| Esm1 | skyblue | -2.0290977 | 5.55E-14 | -1.7874229 | 6.23E-11 |
| Gzmk | black | 8.36891779 | 6.69E-28 | 8.11521894 | 7.86E-22 |
| Atf4 | orange | 0.84198612 | 1.36E-08 | 0.80757674 | 8.90E-08 |
| Zmym6 | red | -0.5745089 | 0.00436597 | -0.5911814 | 0.00400597 |
| Ccno | orange | 0.65714125 | 0.0212655 | 1.28066844 | 4.48E-06 |
| Mgat3 | skyblue | -2.0023675 | 2.30E-15 | -0.9675021 | 0.00025644 |

| | | | | | |
|---|---|---|---|---|---|
| Pih1h3b | skyblue | -3.3250173 | 0.00018254 | -2.3560706 | 0.00631895 |
| Zfp532 | red | -1.1486269 | 1.56E-13 | -0.8065793 | 4.60E-07 |
| Fam63b | red | -0.6348006 | 0.00049927 | -0.4895906 | 0.01001651 |
| Mybph | skyblue | -3.2516335 | 8.55E-08 | -1.2628022 | 0.05153961 |
| Reln | red | -1.0580146 | 3.93E-05 | -0.38806 | 0.16782533 |
| Dctpp1 | orange | 1.30587392 | 1.23E-06 | 0.8439603 | 0.00301023 |
| Fcmr | red | -0.8921554 | 0.00206739 | -0.0328112 | 0.9323009 |
| Abcb4 | skyblue | -1.4825084 | 0.00016529 | -1.1582598 | 0.0062958 |
| Mustn1 | red | -1.2250389 | 2.10E-17 | -0.8483163 | 1.16E-08 |
| Leo1 | orange | 1.07129469 | 3.83E-06 | 1.31621026 | 2.30E-08 |
| Clspn | yellow | 3.65779805 | 6.30E-38 | 4.22536127 | 4.93E-45 |
| Ago4 | red | -1.3038457 | 2.06E-10 | -0.9167138 | 1.60E-05 |
| Mum1l1 | skyblue | -1.5620001 | 1.79E-06 | -1.0538055 | 0.00152008 |
| Dnal1 | red | -0.7641934 | 0.00017985 | -0.5183149 | 0.01525317 |
| Kcnj12 | red | -0.7832491 | 0.0036241 | -0.8514705 | 0.00194063 |
| Shfm1 | orange | 0.63090903 | 6.52E-09 | 0.43882087 | 0.00011128 |
| Fam227a | red | -1.1880884 | 0.00022264 | -0.6200643 | 0.06967166 |
| Nek10 | skyblue | -1.8289337 | 8.00E-05 | -1.2404436 | 0.00507378 |
| Cux2 | skyblue | -1.6331233 | 0.00013047 | -0.9867548 | 0.02756991 |
| Kdm7a | orange | 0.6120652 | 3.86E-06 | 0.26950554 | 0.05958685 |
| Asb4 | yellow | 2.57348175 | 1.33E-05 | 3.032079 | 2.20E-06 |
| Oscp1 | red | -1.1356404 | 1.77E-08 | -0.6589877 | 0.00167647 |
| Maff | orange | 1.11971831 | 2.80E-05 | 1.17510215 | 1.62E-05 |
| Acad12 | skyblue | -1.2786812 | 1.81E-08 | -1.3295595 | 7.49E-09 |
| Arrdc4 | orange | 0.48951044 | 0.0102157 | 0.65913008 | 0.00059381 |
| Rgs8 | black | 4.71084253 | 3.37E-07 | 5.39769321 | 3.32E-06 |
| Ypel3 | red | -0.9065511 | 2.17E-07 | -0.73556 | 4.59E-05 |
| Zc3h12a | orange | 1.80132519 | 3.07E-24 | 1.58077146 | 7.74E-19 |
| Jph1 | red | -0.9579678 | 0.00046848 | -1.0479725 | 0.00021398 |
| Mapk6 | orange | 0.81038813 | 3.41E-05 | 0.94408348 | 1.92E-06 |
| Dnali1 | skyblue | -2.3074154 | 1.06E-19 | -1.6947215 | 3.61E-11 |
| Wbp5 | orange | 0.74168423 | 2.88E-06 | 0.55683917 | 0.00072557 |
| Ppp1r3a | red | -1.1864168 | 0.00051273 | -0.9293369 | 0.00934734 |
| Naa25 | orange | 1.32593354 | 1.39E-09 | 1.33709007 | 1.50E-09 |
| Trafd1 | yellow | 2.24041725 | 9.87E-43 | 2.16416493 | 7.47E-40 |
| Sgtb | red | -0.6270918 | 0.0047923 | -0.3689121 | 0.11717971 |
| Bex2 | skyblue | -3.1851529 | 1.38E-09 | -1.6749443 | 0.00212076 |
| Nmnat2 | red | -0.8551374 | 2.55E-07 | -0.3770548 | 0.03844976 |
| Apobr | yellow | 2.38837385 | 3.01E-14 | 2.24734916 | 1.50E-12 |
| Hebp1 | red | -0.8856323 | 6.24E-07 | -0.6506278 | 0.00044012 |
| Muc1 | red | -1.0373827 | 4.01E-06 | -0.6840847 | 0.00361443 |
| Fam166b | skyblue | -1.9333089 | 2.26E-14 | -1.6254862 | 8.01E-11 |
| Lgr6 | skyblue | -1.8146202 | 4.31E-11 | -1.4839155 | 1.15E-07 |

| | | | | | |
|---|---|---|---|---|---|
| Flt3 | orange | 1.02301824 | 0.00053462 | 0.86365946 | 0.00489531 |
| Fgf11 | skyblue | -1.6096474 | 7.12E-16 | -1.0397551 | 4.08E-07 |
| Trim72 | red | -1.1018683 | 2.63E-05 | -0.7583371 | 0.00591569 |
| Nrep | red | -0.9890667 | 7.23E-07 | -1.0127932 | 6.64E-07 |
| Lhfpl4 | red | -0.9242824 | 0.00269898 | -1.3189657 | 2.85E-05 |
| Abra | red | -1.2998883 | 0.00022414 | -0.6758262 | 0.07771448 |
| Aida | orange | 0.87439079 | 1.25E-15 | 0.71758108 | 1.15E-10 |
| Foxo4 | red | -1.1488404 | 1.77E-08 | -0.9612663 | 4.43E-06 |
| Egflam | red | -1.1204018 | 6.95E-07 | -0.9774651 | 2.54E-05 |
| Upk3b | skyblue | -1.9626179 | 1.80E-10 | -1.7606996 | 2.09E-08 |
| Notum | red | -1.0554499 | 5.71E-05 | -1.1478336 | 2.52E-05 |
| Rasef | skyblue | -1.7815861 | 5.06E-15 | -1.0124048 | 1.47E-05 |
| Gng2 | orange | 0.73262899 | 0.00127569 | 0.23878679 | 0.35137033 |
| Klhl6 | yellow | 2.39106268 | 8.45E-21 | 1.99933216 | 1.25E-14 |
| Wdr63 | skyblue | -2.5937361 | 4.87E-13 | -1.3483086 | 0.00026282 |
| Trpv3 | red | -0.9880601 | 0.00637344 | -0.8910628 | 0.03106623 |
| Ccdc63 | orange | 2.25458685 | 0.00978529 | 1.27943893 | 0.18621469 |
| Zfp513 | orange | 0.77729823 | 2.05E-05 | 0.78990487 | 2.35E-05 |
| Spice1 | red | -1.2780256 | 3.12E-10 | -1.0095707 | 1.15E-06 |
| Tmem82 | orange | 1.01766772 | 0.01347178 | 1.25368552 | 0.00222948 |
| Il17re | red | -0.9507244 | 1.36E-05 | -0.4757481 | 0.04155561 |
| Tuba1c | yellow | 2.35908779 | 1.42E-21 | 2.05229405 | 2.32E-16 |
| Qrfp | yellow | 2.87641505 | 6.03E-12 | 1.6662136 | 9.52E-05 |
| A530016L24I | skyblue | -3.0568754 | 5.96E-06 | -2.3965861 | 0.00032775 |
| Ispd | red | -0.9188653 | 0.00211361 | -0.7905096 | 0.01125055 |
| Ppp2r3a | red | -0.9893455 | 3.25E-06 | -0.9254623 | 2.23E-05 |
| Arl11 | yellow | 2.51414761 | 2.34E-09 | 1.95904109 | 6.05E-06 |
| Pyurf | red | -1.0821807 | 5.05E-10 | -0.3582803 | 0.0578672 |
| Tmem212 | skyblue | -2.0360945 | 3.22E-16 | -1.1536634 | 6.23E-06 |
| Lor | red | -0.6635672 | 0.00728793 | -0.1927368 | 0.49752832 |
| Fam129c | red | -0.9077138 | 0.00131282 | 0.02397341 | 0.950103 |
| Exoc3l | red | -0.8316321 | 5.29E-06 | -0.4923006 | 0.01061247 |
| Tmem64 | red | -0.9314682 | 8.41E-11 | -0.7850415 | 8.61E-08 |
| Fam13c | red | -1.1316073 | 0.00036431 | -1.23102 | 0.00014583 |
| Pyhin1 | yellow | 3.72358667 | 7.86E-36 | 3.59115498 | 4.51E-33 |
| Trim56 | orange | 0.46212942 | 8.38E-05 | 0.61757844 | 1.72E-07 |
| Pcdhb19 | red | -1.2226621 | 0.00899358 | -0.2157546 | 0.68771617 |
| 6530409C15 | orange | 0.58471278 | 0.32651735 | 1.59324825 | 0.0062221 |
| Hoxd9 | red | -1.3325489 | 0.00784478 | -0.7503733 | 0.17336977 |
| Olfr1342 | skyblue | -1.965411 | 0.00183384 | -1.1206529 | 0.08189391 |
| Gprasp1 | skyblue | -1.4810041 | 1.58E-16 | -1.1909889 | 7.84E-11 |
| 2510009E07I | red | -0.7659294 | 1.13E-14 | -0.5233686 | 3.33E-07 |
| Otud1 | red | -1.1254718 | 9.86E-10 | -0.8774651 | 3.84E-06 |

| | | | | | |
|---|---|---|---|---|---|
| Hilpda | red | -0.6666077 | 6.88E-06 | -0.4299543 | 0.00571834 |
| Gjc2 | skyblue | -1.368561 | 3.73E-05 | -1.1804447 | 0.00087928 |
| Lce3d | skyblue | -1.8528788 | 1.50E-05 | -0.8069379 | 0.08264014 |
| Acot6 | red | -0.9227501 | 0.00263688 | -0.3551033 | 0.29618097 |
| Tril | skyblue | -3.1976728 | 1.14E-06 | -2.4121046 | 0.00038702 |
| Lgals2 | red | -1.2274127 | 0.00111951 | -0.5894085 | 0.14829133 |
| Gimap5 | orange | 0.79564739 | 3.98E-10 | 0.94547751 | 1.77E-13 |
| Casc1 | red | -1.3483357 | 5.44E-09 | -0.9592443 | 4.51E-05 |
| Zc2hc1a | red | -0.9425884 | 2.47E-09 | -0.6492735 | 7.01E-05 |
| Fbxl7 | red | -1.1147515 | 4.66E-08 | -0.7239389 | 0.00056798 |
| Pxylp1 | red | -0.6649143 | 4.28E-05 | -0.7202512 | 1.33E-05 |
| Mmp3 | orange | 0.28712691 | 0.29518256 | 1.05698553 | 4.06E-05 |
| Ubxn10 | skyblue | -2.5747542 | 5.65E-20 | -1.6381289 | 8.79E-09 |
| Ecm2 | skyblue | -2.023011 | 4.76E-07 | -1.576146 | 0.00014528 |
| Adamts3 | orange | 1.50809489 | 5.17E-06 | 0.82803073 | 0.0178705 |
| Rbm20 | skyblue | -1.3111847 | 0.0675586 | -2.3317075 | 0.00101936 |
| Tmem221 | red | -0.8350142 | 0.00012489 | -0.8109664 | 0.00032456 |
| Tox3 | skyblue | -1.7558323 | 1.99E-05 | -0.9190176 | 0.03423572 |
| Kcns3 | skyblue | -1.6973088 | 1.19E-11 | -1.0004631 | 0.00010971 |
| Fam25c | red | -1.1056632 | 0.00060552 | -0.6302813 | 0.06789439 |
| Capn13 | skyblue | -3.1353259 | 0.00038967 | -0.9744118 | 0.18894506 |
| Col6a6 | skyblue | -3.3865189 | 2.23E-11 | -2.0671724 | 1.39E-05 |
| B430306N03 | black | 5.19649422 | 2.34E-10 | 3.76295065 | 7.93E-07 |
| Ly6g5b | orange | 1.29990073 | 0.00818509 | 1.035759 | 0.04085119 |
| Adamtsl5 | red | -1.2243241 | 1.49E-10 | -0.8923501 | 5.94E-06 |
| Clec4a3 | yellow | 2.21420221 | 1.82E-11 | 2.30324584 | 6.64E-12 |
| Taf10 | orange | 0.80693748 | 2.25E-05 | 0.641526 | 0.00116941 |
| Ccdc60 | skyblue | -2.0493003 | 5.30E-11 | -0.8676427 | 0.00435633 |
| Gimap7 | yellow | 3.23690933 | 1.70E-20 | 2.56001363 | 3.43E-13 |
| Klri2 | orange | 1.35404147 | 0.00701231 | 1.86225435 | 0.00029685 |
| Wdfy3 | red | -0.6083201 | 7.97E-05 | -0.4681774 | 0.00343306 |
| Naalad2 | skyblue | -1.2119793 | 7.16E-07 | -1.4084895 | 1.97E-08 |
| 2900052L18F | red | -1.2761674 | 1.89E-05 | -0.5716557 | 0.07350939 |
| Mgat2 | orange | 0.6730175 | 0.00013792 | 0.63584146 | 0.00045706 |
| Cilp2 | skyblue | -2.5832001 | 0.0516669 | -4.7318991 | 0.00035769 |
| Slc35g1 | orange | 1.06425699 | 4.88E-06 | 0.55598354 | 0.02296852 |
| Ccdc141 | red | -0.9038341 | 0.00688713 | -0.7365537 | 0.0342582 |
| Als2cl | red | -0.6994191 | 2.38E-06 | -0.3167268 | 0.04672432 |
| Pcdhb14 | red | -0.8010149 | 0.00169729 | -0.5999523 | 0.02290554 |
| Ccr10 | skyblue | -1.7454382 | 0.00148411 | -1.415958 | 0.01202275 |
| Cep68 | red | -0.6608847 | 8.24E-08 | -0.591909 | 2.90E-06 |
| Zrsr1 | red | -1.3040836 | 3.44E-07 | -1.156286 | 1.01E-05 |
| Efcab8 | yellow | 2.35707911 | 0.00066201 | 2.61161525 | 0.00036126 |

| | | | | | |
|---|---|---|---|---|---|
| Lmod3 | skyblue | -1.9910355 | 1.70E-07 | -1.8423627 | 3.34E-06 |
| C130050O18 | orange | 1.56457893 | 6.33E-05 | 1.67651985 | 2.45E-05 |
| Proca1 | red | -1.5094775 | 1.03E-06 | -0.9882965 | 0.0020355 |
| Fam109a | red | -0.6641533 | 0.00156988 | -0.8358639 | 8.99E-05 |
| Arf6 | orange | 0.98242285 | 3.60E-10 | 0.79969025 | 6.78E-07 |
| Hepacam2 | skyblue | -3.6732129 | 1.38E-09 | -2.5025763 | 1.65E-05 |
| Tnip3 | black | 4.22605718 | 4.92E-29 | 4.57896174 | 2.40E-31 |
| Bcl2l15 | yellow | 4.49967959 | 0.00011854 | 2.30563346 | 0.01282693 |
| Cdc25c | yellow | 4.26977561 | 8.70E-21 | 3.85014834 | 4.72E-17 |
| Nxpe4 | skyblue | -1.5481353 | 3.80E-19 | -1.6395406 | 5.18E-21 |
| Il20rb | orange | 0.73724629 | 0.00030896 | 0.6796446 | 0.00130395 |
| Osbpl1a | red | -0.6004082 | 1.46E-05 | -0.6123232 | 1.51E-05 |
| Zfp879 | skyblue | -2.3239996 | 3.06E-05 | -1.6138602 | 0.00300662 |
| Mab21l3 | orange | 1.5669643 | 0.00443762 | 1.76036269 | 0.00162326 |
| Ackr3 | orange | 1.29190652 | 3.80E-08 | 1.3099545 | 4.07E-08 |
| Aplnr | skyblue | -4.6186406 | 1.83E-33 | -3.5127477 | 7.83E-20 |
| Phlpp1 | red | -0.9656345 | 2.61E-09 | -0.7901964 | 2.07E-06 |
| Marveld1 | red | -1.0981266 | 4.27E-13 | -1.0186074 | 3.20E-11 |
| Snhg11 | red | -1.4346085 | 0.00068734 | -0.9500378 | 0.03289882 |
| Lacc1 | yellow | 2.76225059 | 5.35E-19 | 2.01968721 | 1.87E-10 |
| Sowaha | skyblue | -2.8354106 | 1.72E-05 | -1.0122453 | 0.11174706 |
| Cxxc4 | skyblue | -2.639874 | 1.93E-06 | -1.6746267 | 0.00217445 |
| Dsg2 | red | -1.1306641 | 0.00015771 | -0.900209 | 0.00348415 |
| Adig | skyblue | -1.9998329 | 0.00068709 | -1.8223643 | 0.00264183 |
| Sptssa | red | -0.4926847 | 1.51E-05 | -0.6363413 | 3.30E-08 |
| Dock5 | orange | 0.648582 | 4.93E-06 | 0.43914333 | 0.00299959 |
| Rin3 | orange | 0.62584481 | 1.43E-05 | 0.28795179 | 0.06348288 |
| Shisa2 | skyblue | -2.144078 | 8.99E-08 | -1.4589191 | 0.00049755 |
| Tnfaip8l1 | orange | 1.3413003 | 4.55E-09 | 0.92100478 | 0.00013613 |
| A730020M07 | orange | 1.90702833 | 0.00552401 | 0.79041696 | 0.29149149 |
| Tram1l1 | skyblue | -1.4878193 | 0.0001679 | -1.3758951 | 0.0005281 |
| Rps2 | orange | 0.88648295 | 4.18E-05 | 0.85852597 | 0.00010802 |
| Tceal3 | skyblue | -2.4065427 | 1.52E-06 | -0.9820144 | 0.05814212 |
| Tlr7 | yellow | 3.01926413 | 3.54E-11 | 2.46186212 | 1.42E-07 |
| Rnf208 | red | -1.0497853 | 4.51E-05 | -0.4508273 | 0.10761431 |
| Cnrip1 | red | -0.6304399 | 0.00028397 | -0.4040273 | 0.03167462 |
| B530045E10I | skyblue | -2.4089726 | 8.26E-05 | -1.670265 | 0.00656321 |
| Zfp612 | skyblue | -1.8914011 | 1.72E-10 | -1.4143275 | 2.82E-06 |
| 2010007H06 | skyblue | -1.5831758 | 1.29E-08 | -0.8205819 | 0.00482379 |
| Il27 | black | 4.78427496 | 3.88E-20 | 4.47602005 | 5.73E-18 |
| Palb2 | orange | 1.33058654 | 1.09E-05 | 1.49268706 | 2.33E-06 |
| Phf11a | black | 6.1011627 | 1.73E-39 | 5.1349743 | 1.84E-32 |
| Serpinb1a | red | -0.7969773 | 9.17E-06 | -0.4855087 | 0.01077919 |

| | | | | | |
|---|---|---|---|---|---|
| Scml4 | orange | 1.05422263 | 8.88E-11 | 1.45167337 | 3.84E-18 |
| Sntn | skyblue | -2.1822633 | 2.70E-06 | -1.9460427 | 3.96E-05 |
| Zfp36 | orange | 1.35231116 | 1.77E-08 | 1.11749878 | 5.70E-06 |
| Fads6 | red | -0.8306481 | 0.00128083 | -0.5767783 | 0.03274161 |
| Isca1 | red | -0.5784032 | 4.31E-07 | -0.6889839 | 2.47E-09 |
| Zfp354c | red | -0.6896601 | 0.004825 | -0.8550407 | 0.00057655 |
| AF251705 | yellow | 2.96725926 | 1.21E-18 | 2.49425176 | 2.90E-13 |
| Tlr1 | yellow | 2.90880902 | 2.30E-20 | 2.77673709 | 1.88E-18 |
| Ankrd45 | skyblue | -1.9617417 | 2.03E-11 | -1.3409732 | 2.76E-06 |
| 1700056E22l | orange | 1.34195641 | 0.00415769 | 0.92003916 | 0.10777131 |
| Ankrd50 | red | -0.6292787 | 0.0004937 | -0.218135 | 0.2793846 |
| Gimap1os | orange | 1.47309981 | 4.18E-07 | 0.93295412 | 0.00230716 |
| Rassf9 | red | -1.4818395 | 1.23E-12 | -0.5687932 | 0.01013972 |
| Ttc41 | red | -0.5375501 | 0.03899741 | -0.7051142 | 0.00673193 |
| Klhl31 | skyblue | -1.8103776 | 2.91E-08 | -1.5617313 | 4.22E-06 |
| Mylk4 | skyblue | -2.22035 | 1.13E-09 | -1.6947112 | 6.60E-06 |
| Kctd21 | skyblue | -1.5849458 | 2.51E-08 | -1.2153418 | 3.07E-05 |
| Fbxo48 | orange | 2.16333596 | 0.0004893 | 1.32265435 | 0.03668595 |
| Wdr72 | skyblue | -2.148272 | 2.52E-09 | -1.7976192 | 1.36E-07 |
| Tst | red | -0.9477632 | 3.41E-08 | -0.7671035 | 1.46E-05 |
| 1110034G24 | red | -0.4822606 | 0.02123653 | -1.0779316 | 2.76E-07 |
| Prss22 | yellow | 3.72394779 | 8.41E-11 | 3.73369741 | 3.70E-10 |
| Cetn4 | red | -0.6907523 | 0.01850126 | -0.9623533 | 0.00132903 |
| Tmem232 | skyblue | -2.8652854 | 3.17E-10 | -2.1096485 | 2.54E-06 |
| Prkce | red | -0.7931571 | 4.61E-09 | -0.7842069 | 1.12E-08 |
| Megf8 | red | -0.7019481 | 3.70E-11 | -0.495949 | 6.20E-06 |
| Rpsa-ps2 | orange | 2.51725097 | 0.00257915 | -0.1090415 | 0.92182732 |
| Zc2hc1c | red | -0.963955 | 0.00010094 | -1.034184 | 3.89E-05 |
| 9930022D16 | black | 3.29056431 | 0.00077511 | 5.51856401 | 8.24E-05 |
| S1pr1 | orange | 0.58311664 | 0.00011638 | 0.41837679 | 0.00807681 |
| Arhgef37 | orange | 1.96282973 | 5.68E-15 | 1.85431417 | 3.80E-14 |
| Magi1 | red | -0.7456393 | 3.90E-05 | -0.5680662 | 0.00255236 |
| Slc25a26 | red | -0.599648 | 0.00013031 | -0.4915119 | 0.00276397 |
| Dmd | red | -0.704963 | 0.00153816 | -0.6536234 | 0.00434795 |
| Rpl18a | orange | 0.59844 | 5.19E-06 | 0.4247113 | 0.00189631 |
| Tubb2b | orange | 0.41532748 | 0.08109941 | 1.03524536 | 7.74E-06 |
| AI467606 | orange | 0.57945387 | 7.84E-08 | 0.60263878 | 5.31E-08 |
| Lrrc3b | skyblue | -2.7678531 | 0.00223714 | -3.8306828 | 0.00021455 |
| Kcng4 | skyblue | -2.0997703 | 0.00951843 | -1.6278126 | 0.05220313 |
| Morn2 | red | -1.2505446 | 3.00E-05 | -0.5801948 | 0.07211015 |
| Cenph | yellow | 2.46290269 | 5.00E-07 | 2.33995247 | 5.27E-06 |
| Lca5l | red | -1.1131057 | 9.55E-06 | -0.0931878 | 0.75925746 |
| Dcaf12l1 | skyblue | -1.9353803 | 0.00565799 | -2.0545711 | 0.0079273 |

| | | | | | |
|---|---|---|---|---|---|
| Rtn4rl1 | red | -0.9077803 | 3.33E-06 | -0.6076775 | 0.00271131 |
| Insig1 | red | -0.7948404 | 3.06E-05 | -0.4421113 | 0.0288559 |
| Preb | orange | 0.65865608 | 1.48E-12 | 0.50473995 | 1.32E-07 |
| Fahd1 | red | -0.684213 | 8.65E-06 | -0.7971293 | 5.23E-07 |
| Tlr9 | yellow | 3.35660067 | 1.57E-08 | 3.1459156 | 2.08E-07 |
| Cenpe | yellow | 3.43978037 | 7.19E-29 | 3.62640503 | 7.80E-30 |
| Sh2d5 | yellow | 2.70225583 | 3.76E-08 | 2.71827182 | 3.70E-08 |
| Tnfrsf26 | yellow | 1.80319405 | 2.65E-06 | 2.23800157 | 5.59E-09 |
| Olfr1033 | red | -0.6630956 | 0.00258185 | -0.0987738 | 0.71045773 |
| Kcnk13 | orange | 1.72861288 | 1.02E-05 | 1.14638044 | 0.00512564 |
| 1190002N15 | skyblue | -1.7892917 | 3.54E-12 | -1.3110411 | 7.64E-07 |
| 2810002D19 | red | -0.6897982 | 0.00647209 | -0.8435102 | 0.00124232 |
| Ttll13 | red | -1.2594766 | 0.00115174 | -0.9337194 | 0.01521098 |
| Lce3c | red | -0.94556 | 0.0074065 | -0.666559 | 0.07420993 |
| Pcdhb3 | skyblue | -1.6118669 | 0.0026197 | -1.4543232 | 0.0078712 |
| Hcar2 | orange | 1.65258402 | 1.88E-08 | 1.57626874 | 1.02E-07 |
| Sys1 | orange | 0.71356679 | 2.74E-08 | 0.49737577 | 0.0002308 |
| Kcna5 | skyblue | -1.4959836 | 0.00010729 | -2.0643739 | 1.72E-07 |
| Ddx28 | orange | 0.88307637 | 1.96E-08 | 0.61989361 | 0.00018491 |
| Krt14 | red | -0.8880228 | 0.00151933 | -0.8362223 | 0.00379966 |
| Mettl24 | skyblue | -1.9598254 | 5.06E-15 | -1.5388781 | 1.71E-09 |
| Penk | red | -0.7433363 | 0.00625766 | -0.6697648 | 0.01879095 |
| Olig3 | skyblue | -2.3886735 | 0.00015087 | -0.9970336 | 0.09331292 |
| Chrm2 | red | -0.5714987 | 0.06048987 | -0.9699192 | 0.00125396 |
| Odf3l1 | skyblue | -3.0080279 | 0.00625795 | -2.5440635 | 0.00978171 |
| Esf1 | orange | 0.6262064 | 0.00013317 | 0.189476 | 0.30757687 |
| Sh3tc2 | red | -1.1615255 | 9.65E-08 | -0.9870747 | 1.03E-05 |
| Mtus1 | red | -1.1095954 | 4.27E-08 | -0.8013099 | 0.00013408 |
| Fam216b | skyblue | -2.4618453 | 7.85E-10 | -1.4622381 | 0.00032384 |
| Pid1 | red | -0.414836 | 0.01007947 | -0.6297109 | 9.86E-05 |
| Col27a1 | skyblue | -1.3952392 | 8.45E-14 | -1.2670721 | 3.00E-11 |
| Tcf21 | skyblue | -1.2739325 | 3.26E-07 | -1.292326 | 3.50E-07 |
| Pcdhb4 | skyblue | -1.6578436 | 1.32E-05 | -0.9197756 | 0.02022007 |
| Thtpa | red | -0.6129564 | 0.015786 | -0.7872296 | 0.00180309 |
| Smkr-ps | skyblue | -1.5853009 | 5.19E-05 | -0.8985163 | 0.02392329 |
| Mms22l | yellow | 2.27217595 | 9.85E-22 | 2.36424136 | 3.48E-22 |
| Zfp764 | red | -0.4925958 | 0.00299652 | -0.6084959 | 0.00029672 |
| Fam179a | red | -1.2509597 | 7.85E-10 | -0.6586737 | 0.00183101 |
| Basp1 | yellow | 2.41657557 | 2.19E-09 | 2.51223666 | 7.80E-10 |
| Slc16a5 | skyblue | -3.0004332 | 0.00056002 | 1.41467123 | 0.12105763 |
| Lrtm1 | red | -0.8213686 | 0.00280628 | -0.5004517 | 0.0951453 |
| Ccdc149 | red | -0.6357118 | 0.00124229 | -0.4755002 | 0.02177577 |
| Whamm | orange | 0.68458264 | 1.32E-12 | 0.72581498 | 4.66E-14 |

| | | | | | |
|---|---|---|---|---|---|
| Zfp36l2 | orange | 0.69795607 | 0.00358874 | 0.64788831 | 0.00868644 |
| Ptprcap | yellow | 2.60096738 | 2.87E-25 | 2.72172216 | 3.85E-26 |
| Serpinb9 | orange | 1.0349298 | 1.26E-10 | 0.98162186 | 1.85E-09 |
| Gvin1 | yellow | 2.56940512 | 5.84E-20 | 2.37491237 | 3.18E-17 |
| Slitrk6 | skyblue | -1.4595309 | 7.39E-06 | -1.6736626 | 6.83E-07 |
| Adra1a | red | -0.9496626 | 0.00631572 | -0.5659801 | 0.12858455 |
| Ccdc42 | skyblue | -2.1322881 | 1.99E-08 | -0.9504703 | 0.01605319 |
| 6330416G13 | orange | 0.97794679 | 1.13E-05 | 0.9610469 | 2.44E-05 |
| Clec14a | red | -1.3609564 | 7.02E-08 | -1.0544335 | 5.24E-05 |
| Ifit2 | yellow | 4.15147551 | 3.47E-39 | 4.16654465 | 2.00E-39 |
| Sdpr | red | -1.3586705 | 1.09E-05 | -1.1053187 | 0.0005399 |
| Slc25a51 | red | -0.6725596 | 8.89E-08 | -0.5059762 | 0.00010243 |
| C2cd2 | red | -0.892224 | 3.60E-10 | -0.7832092 | 7.30E-08 |
| Tmem104 | orange | 0.96034019 | 5.47E-05 | 0.57677069 | 0.02177577 |
| Gapt | orange | 0.6611528 | 0.30092181 | 2.20980532 | 0.00073108 |
| Fam26f | black | 5.95310499 | 1.01E-79 | 5.9214366 | 2.65E-74 |
| Otulin | orange | 0.90420269 | 6.71E-12 | 0.69536842 | 2.70E-07 |
| Ppp1r15b | orange | 0.67032005 | 4.13E-09 | 0.54229452 | 3.81E-06 |
| Igfals | skyblue | -1.5987724 | 0.01362763 | -1.7580415 | 0.00465738 |
| Clec9a | red | -1.2591631 | 2.70E-06 | -1.0209557 | 0.00022853 |
| Hpcal4 | skyblue | -3.4386724 | 1.35E-19 | -2.1497475 | 4.30E-09 |
| Krt32 | skyblue | -1.7204638 | 0.00230497 | -1.0227164 | 0.08832356 |
| Cep295 | orange | 0.7533473 | 3.50E-07 | 0.7024683 | 3.32E-06 |
| Fut10 | red | -1.0911786 | 1.13E-07 | -0.903704 | 1.68E-05 |
| Tmem229b | orange | 0.96603718 | 3.04E-08 | 0.85984786 | 1.52E-06 |
| Chrm3 | skyblue | -1.6920271 | 0.00066044 | -2.099056 | 4.34E-05 |
| Kcnrg | skyblue | -1.9344127 | 2.30E-12 | -0.9699109 | 0.00048409 |
| Adamts6 | orange | 1.6332276 | 1.43E-23 | 1.4298065 | 2.26E-18 |
| E2f8 | yellow | 2.85486332 | 8.93E-31 | 2.92310081 | 7.95E-31 |
| Gsg1l | skyblue | -2.892675 | 2.79E-06 | -3.6982163 | 1.29E-08 |
| Cd109 | red | -0.8405852 | 0.00018708 | -0.7640606 | 0.00121025 |
| Pcdhb20 | skyblue | -1.3908202 | 1.83E-08 | -1.2880019 | 2.42E-07 |
| Iqub | skyblue | -2.2010229 | 2.12E-13 | -1.2508577 | 1.91E-05 |
| Rprml | skyblue | -2.1412053 | 6.71E-12 | -0.8747307 | 0.00684294 |
| Plaur | orange | 1.81584584 | 9.40E-09 | 1.75914796 | 4.59E-08 |
| Scand1 | orange | 1.174877 | 0.00019858 | 1.4258943 | 1.18E-05 |
| Hepacam | skyblue | -1.8611357 | 0.00021122 | -0.6831349 | 0.2071387 |
| 1600029I14R | skyblue | -2.4314125 | 1.56E-13 | -1.6550725 | 9.03E-07 |
| Pilra | orange | 2.06656837 | 9.50E-05 | 1.76376818 | 0.00122749 |
| Tusc5 | red | -1.2725907 | 3.66E-05 | -1.1745365 | 0.00020025 |
| She | red | -1.127515 | 3.76E-17 | -0.8687885 | 2.09E-10 |
| Ankle1 | yellow | 1.99006179 | 1.02E-13 | 3.31579827 | 8.88E-28 |
| AI464131 | red | -1.1318233 | 8.37E-05 | -0.9941762 | 0.00081652 |

| | | | | | |
|---|---|---|---|---|---|
| Stxbp6 | red | -1.0390724 | 0.00031106 | -0.5664578 | 0.06585494 |
| Slc25a23 | skyblue | -1.723698 | 3.65E-11 | -1.0974499 | 4.79E-05 |
| Fam178b | skyblue | -2.0476301 | 1.27E-06 | -0.6425367 | 0.12403751 |
| Smco1 | skyblue | -1.3251881 | 0.12890946 | -3.2586249 | 0.00359195 |
| Asphd1 | yellow | 3.35591106 | 0.00180632 | -0.0666134 | 0.9534411 |
| Rbp1 | red | -0.9978033 | 3.17E-12 | -0.873061 | 2.43E-09 |
| Irx3os | red | -0.9531899 | 0.00061118 | -0.3642409 | 0.22945895 |
| Lrrc75a | red | -0.9354804 | 5.73E-05 | -0.7124988 | 0.0033641 |
| Hnrnpa1 | orange | 0.62237221 | 5.47E-09 | 0.63702997 | 3.73E-09 |
| Ppm1e | red | -1.562884 | 0.00111461 | -0.6140025 | 0.2343724 |
| Camk2n1 | red | -1.1124003 | 2.18E-17 | -0.8639618 | 1.15E-10 |
| Sox18 | skyblue | -1.6688763 | 2.10E-13 | -1.3416782 | 7.81E-09 |
| Scn4b | skyblue | -1.8663301 | 0.00028412 | -0.8244228 | 0.14190572 |
| C1qtnf2 | red | -1.2502904 | 1.06E-06 | -0.9634619 | 0.0003044 |
| Ar | red | -1.3380506 | 9.05E-06 | -0.9370934 | 0.00300698 |
| Fam43a | red | -0.808872 | 1.55E-13 | -0.681285 | 1.24E-09 |
| Zfp319 | red | -0.584011 | 0.00022565 | -0.4330563 | 0.00863311 |
| 4930430F08I | red | -0.5626931 | 0.027109 | -0.6797314 | 0.00975094 |
| Cfap58 | skyblue | -2.1928438 | 2.05E-10 | -1.4035223 | 6.06E-05 |
| Lrrc8e | red | -0.7002948 | 2.77E-05 | -0.5542294 | 0.00153616 |
| Ticrr | yellow | 3.5858467 | 1.82E-23 | 4.15805107 | 1.16E-26 |
| Olfml2a | skyblue | -2.5977209 | 1.36E-19 | -1.7440468 | 2.69E-09 |
| Ttc34 | skyblue | -2.5761219 | 1.58E-08 | -0.9814921 | 0.03594934 |
| Zfp316 | red | -0.8147206 | 1.44E-07 | -0.5410016 | 0.00081202 |
| Cxxc5 | red | -0.9266662 | 1.28E-10 | -0.4517485 | 0.00302696 |
| Tmem251 | orange | 1.0800935 | 1.69E-05 | 0.75416077 | 0.00422169 |
| Lce1l | skyblue | -2.0655258 | 1.18E-06 | -2.064277 | 3.75E-06 |
| C87436 | red | -0.5918109 | 0.00151679 | -0.3789688 | 0.05258304 |
| Gm5424 | orange | 2.06908801 | 3.39E-05 | 0.44838403 | 0.4185153 |
| Tifa | yellow | 2.11669626 | 1.33E-29 | 2.10273658 | 1.20E-28 |
| Mapk10 | skyblue | -1.7196422 | 0.00074635 | -1.778684 | 0.0007033 |
| Hmga1 | orange | 2.05164855 | 1.44E-12 | 1.9403606 | 3.68E-11 |
| Foxc2 | red | -1.4235476 | 1.81E-05 | -0.8759666 | 0.01201456 |
| Bst2 | yellow | 2.53459333 | 4.74E-43 | 2.30153635 | 1.49E-35 |
| Nxph3 | red | -0.8530133 | 7.11E-05 | -0.991502 | 1.26E-05 |
| Cdc42se1 | orange | 0.61952784 | 1.22E-07 | 0.47069414 | 0.00010496 |
| Fat4 | red | -1.2253907 | 1.98E-18 | -1.0169497 | 9.39E-13 |
| Epm2aip1 | red | -0.5873845 | 0.00113841 | -0.3169909 | 0.10174106 |
| Phgr1 | yellow | 3.04402274 | 0.00067256 | 1.28841919 | 0.10559857 |
| Mpeg1 | yellow | 2.88654841 | 1.96E-13 | 2.63601413 | 3.58E-11 |
| Lrrc75b | red | -1.1848647 | 0.00519243 | -0.7378493 | 0.09737018 |
| Ddit4l | skyblue | -1.8068175 | 3.37E-08 | -1.1521734 | 0.00082251 |
| Ckap4 | orange | 0.69118339 | 0.00223964 | 0.58541721 | 0.01256485 |

| | | | | | |
|---|---|---|---|---|---|
| Vat1l | skyblue | -2.3780313 | 1.44E-06 | -1.4398053 | 0.00382646 |
| Fbl | orange | 1.22219333 | 2.42E-07 | 1.10510337 | 5.30E-06 |
| Irgm1 | yellow | 3.8854562 | 2.77E-117 | 3.57868399 | 3.28E-100 |
| Zfp474 | red | -1.1549447 | 2.14E-09 | -0.4852523 | 0.01448522 |
| Ltb4r1 | yellow | 2.27454054 | 1.45E-11 | 2.35579153 | 4.80E-12 |
| Myct1 | skyblue | -1.8016145 | 8.93E-32 | -1.4091581 | 2.38E-19 |
| Wdr27 | skyblue | -1.5715661 | 2.00E-05 | -0.6962988 | 0.07216343 |
| Gm5637 | yellow | 2.25635374 | 0.00025131 | 1.7467706 | 0.00506391 |
| 1110032F04I | orange | 2.27681195 | 0.00017013 | 1.32078057 | 0.04863519 |
| Ccdc108 | red | -1.4936249 | 2.06E-06 | -0.74289 | 0.02599963 |
| Nipa1 | red | -1.1153587 | 1.69E-06 | -0.7022488 | 0.00412588 |
| Dusp28 | orange | 0.77442645 | 1.95E-09 | 0.70262104 | 1.28E-07 |
| Lrrc4b | skyblue | -1.3346677 | 0.00090628 | -1.4271846 | 0.00065019 |
| Rnf31 | orange | 0.92892954 | 4.86E-23 | 0.93645676 | 2.09E-23 |
| Fam221a | red | -1.1472301 | 1.74E-05 | -0.8537549 | 0.00204766 |
| Tet1 | skyblue | -2.1527667 | 9.99E-07 | -0.9566514 | 0.04094549 |
| Neurl3 | yellow | 2.0374723 | 1.23E-16 | 2.22452408 | 1.77E-19 |
| Irs3 | red | -1.4950144 | 0.00172043 | -0.6719412 | 0.18859746 |
| Dync2h1 | red | -0.9354251 | 2.59E-06 | -0.9748524 | 1.33E-06 |
| Gjd3 | yellow | 3.98944776 | 0.00205534 | 0.99237156 | 0.38422982 |
| Dusp18 | skyblue | -1.5971743 | 1.10E-18 | -1.0723337 | 5.56E-09 |
| Cdh19 | skyblue | -1.1166313 | 0.00691538 | -1.3882465 | 0.00122587 |
| Gm527 | red | -0.6268171 | 0.05894414 | -0.9223468 | 0.00749147 |
| BC048546 | skyblue | -2.8103087 | 6.95E-24 | -1.9908718 | 3.06E-13 |
| Cldn2 | orange | 1.58055582 | 0.00152534 | 2.27122232 | 9.47E-07 |
| Mageh1 | red | -0.7379297 | 8.99E-06 | -0.4186639 | 0.01699661 |
| Hist1h2be | skyblue | -2.3043713 | 9.92E-06 | -1.4622744 | 0.00594364 |
| Ptgs1 | orange | 0.67699604 | 1.66E-07 | 0.51083699 | 0.00013103 |
| Gap43 | skyblue | -2.1387724 | 4.55E-12 | -2.0964345 | 2.59E-11 |
| Sfn | orange | 0.72956191 | 0.00872639 | 0.75367286 | 0.00813306 |
| Gpr15 | yellow | 4.12357841 | 1.69E-10 | 2.6644091 | 3.01E-07 |
| Kcne4 | orange | 0.68687862 | 0.04068867 | 1.0814678 | 0.00114448 |
| Lancl3 | orange | 2.18978377 | 0.00088798 | 1.1645418 | 0.09769153 |
| Gm973 | skyblue | -2.3862728 | 6.34E-17 | -1.6338009 | 1.49E-08 |
| Dnah14 | orange | 0.50072547 | 0.22563135 | 1.0790159 | 0.00817445 |
| B4gat1 | red | -0.7467393 | 1.63E-08 | -0.6532939 | 1.51E-06 |
| Odf3b | skyblue | -1.9524914 | 1.18E-08 | -1.2916875 | 0.00026451 |
| Tgif1 | orange | 1.50607142 | 2.90E-07 | 1.38089212 | 4.40E-06 |
| Ctdspl | red | -1.1414965 | 0.00010316 | -0.8736668 | 0.00421081 |
| Gpr68 | yellow | 3.48053126 | 3.56E-21 | 3.0631056 | 3.45E-16 |
| Cmya5 | red | -0.85576 | 0.00110412 | -1.0492576 | 8.25E-05 |
| Fam180a | skyblue | -0.9491807 | 0.04270654 | -2.5555104 | 1.37E-06 |
| Zfp30 | red | -1.5268511 | 4.71E-08 | -1.016864 | 0.00032748 |

| | | | | | |
|---|---|---|---|---|---|
| Klhl34 | skyblue | -2.1967489 | 0.00310235 | -1.3507494 | 0.08106451 |
| Adamts12 | orange | 1.079689 | 0.00516066 | 1.31100656 | 0.00094042 |
| Cldn4 | orange | 1.4181113 | 3.48E-06 | 1.43711052 | 3.99E-06 |
| Mis18bp1 | yellow | 3.58792986 | 4.96E-18 | 2.87689249 | 4.78E-12 |
| Cltb | orange | 0.61513504 | 0.00028532 | 0.56706414 | 0.00115254 |
| Mafa | red | -1.3234947 | 0.00353421 | -0.3924891 | 0.45243506 |
| Nxpe5 | yellow | 2.72055206 | 5.14E-07 | 3.07401497 | 4.16E-08 |
| Zfp235 | red | -0.6457567 | 0.00930167 | -0.583746 | 0.02220696 |
| Apof | black | 5.49066552 | 0.00061568 | 2.75548334 | 0.05640654 |
| 2810006K23I | red | -0.7031307 | 0.00603436 | -0.8178391 | 0.00183353 |
| Krt83 | orange | 2.04036076 | 2.44E-06 | 1.75998019 | 4.61E-05 |
| D930020B18 | skyblue | -3.2183401 | 2.25E-06 | -0.940223 | 0.16801699 |
| Fbxo30 | red | -0.9463243 | 8.07E-08 | -0.8158331 | 6.26E-06 |
| Trpt1 | red | -0.66227 | 0.000181 | -0.5263653 | 0.00371869 |
| Gal3st3 | skyblue | -1.4324686 | 0.01265361 | -1.5593689 | 0.00626177 |
| Tctex1d4 | red | -1.2260619 | 3.20E-06 | -0.7300468 | 0.00726805 |
| Rpsa-ps10 | orange | 1.11731772 | 3.19E-08 | 0.62248057 | 0.00508614 |
| Bola2 | orange | 0.79423212 | 1.67E-05 | 0.64628766 | 0.00081273 |
| Samd9l | orange | 1.64397797 | 2.26E-30 | 1.59419142 | 1.69E-28 |
| Fbxo40 | skyblue | -1.5132189 | 1.54E-06 | -1.1465284 | 0.00046545 |
| Rnf150 | red | -0.9187061 | 1.22E-05 | -0.7077941 | 0.00117642 |
| Zc3hav1l | red | -0.9114313 | 5.93E-06 | -0.7791051 | 0.00021709 |
| Utf1 | yellow | 4.91441962 | 0.0001125 | 2.15129113 | 0.02381139 |
| Lix1 | skyblue | -1.9085459 | 0.00049214 | -1.4464067 | 0.01263326 |
| Sned1 | red | -1.0079376 | 0.00028773 | -0.6524271 | 0.02535315 |
| Gjb1 | skyblue | -1.705749 | 0.00758204 | -1.3262687 | 0.04919595 |
| Cd300lf | black | 4.61908725 | 2.29E-32 | 4.44202754 | 5.41E-30 |
| Ccdc88b | yellow | 2.90019864 | 2.64E-23 | 3.01537824 | 5.47E-25 |
| Cdca4 | orange | 0.96384666 | 8.49E-10 | 0.86892309 | 7.25E-08 |
| Diras2 | skyblue | -2.7262076 | 2.77E-16 | -1.4991711 | 1.28E-05 |
| Bex4 | red | -1.0370684 | 0.0043305 | -0.8198635 | 0.03059368 |
| Stx19 | red | -0.7536577 | 0.00888401 | -0.7160077 | 0.01802762 |
| Ccr4 | orange | 1.07147251 | 0.11549192 | 2.44724389 | 0.00071436 |
| Marcksl1 | orange | 1.57239031 | 1.18E-14 | 1.71577941 | 5.50E-17 |
| Kcna3 | yellow | 1.74377819 | 8.32E-06 | 2.48125125 | 7.63E-10 |
| Stbd1 | red | -1.0977135 | 1.69E-09 | -0.6357669 | 0.00090364 |
| Kcna1 | red | -1.0759941 | 0.00033119 | -0.1126979 | 0.76218079 |
| Ino80c | orange | 0.70075854 | 3.22E-05 | 0.59246939 | 0.00066941 |
| Prrg1 | skyblue | -1.452131 | 9.55E-07 | -1.2802058 | 2.69E-05 |
| Timm8a1 | orange | 1.09010808 | 0.00026389 | 0.76983872 | 0.01447743 |
| Zfp473 | orange | 2.03968292 | 9.39E-05 | 1.83655659 | 0.00076036 |
| Rgmb | red | -0.8989998 | 4.03E-08 | -0.6826552 | 5.51E-05 |
| Eno4 | skyblue | -2.5289295 | 2.56E-15 | -1.2165131 | 0.00018884 |

| | | | | | |
|---|---|---|---|---|---|
| Fcrl5 | yellow | 2.7983237 | 3.76E-06 | 2.98321897 | 5.74E-06 |
| 4932418E24I | skyblue | -1.7303109 | 2.30E-10 | -1.1070666 | 9.16E-05 |
| Arxes2 | skyblue | -1.440559 | 0.00100506 | -1.3386002 | 0.00304265 |
| Ldlrad3 | red | -0.6236859 | 3.51E-05 | -0.6125909 | 7.42E-05 |
| Cyb5r2 | red | -1.4386091 | 0.00548566 | -0.9155698 | 0.10770939 |
| Lmod1 | skyblue | -1.557061 | 5.25E-15 | -1.0476547 | 3.39E-07 |
| Entpd1 | orange | 0.60035353 | 1.75E-05 | 0.41292018 | 0.00465794 |
| Nat8l | skyblue | -2.3906609 | 1.91E-07 | -2.2363177 | 1.75E-06 |
| Selplg | yellow | 2.82158401 | 1.84E-39 | 2.57260736 | 8.33E-33 |
| Bend7 | red | -0.7615099 | 3.06E-05 | -0.639016 | 0.00068023 |
| Cracr2b | red | -1.3087571 | 3.81E-07 | -1.0933038 | 3.75E-05 |
| A630023P12I | orange | 0.68461416 | 0.14942775 | 1.2470738 | 0.00947555 |
| Gpr85 | orange | 1.05829597 | 0.11463023 | 2.17350791 | 0.00146004 |
| Amigo2 | red | -0.6444991 | 0.00042252 | -0.5963539 | 0.00151757 |
| Fbxo10 | red | -0.9782613 | 2.08E-07 | -0.7524649 | 0.00010256 |
| Rnf149 | orange | 2.07342396 | 5.86E-11 | 1.77333861 | 4.32E-08 |
| Gng7 | red | -1.0551202 | 1.38E-05 | -1.2304113 | 3.95E-07 |
| Bcl11b | orange | 1.26342532 | 0.03249164 | 2.32534436 | 7.07E-05 |
| Syngr2 | orange | 0.76005828 | 1.39E-05 | 0.57928356 | 0.00141634 |
| Frmd6 | red | -0.781361 | 6.75E-05 | -0.6038768 | 0.00297639 |
| Ckap2l | yellow | 2.64382219 | 2.07E-18 | 2.72628903 | 8.89E-19 |
| Ric3 | red | -1.4344434 | 7.17E-05 | -0.9278759 | 0.01117103 |
| Lhfp | red | -0.6627485 | 1.03E-06 | -0.4469317 | 0.00157477 |
| Omd | red | -0.6032633 | 0.25475363 | -1.6397681 | 0.00119713 |
| Pdp2 | red | -0.7163323 | 0.00113016 | -0.8187056 | 0.00028708 |
| Scrt1 | yellow | 1.67316701 | 0.08576287 | 4.48911088 | 0.0008215 |
| Osr1 | orange | 0.57561182 | 0.0008966 | 0.89946715 | 2.19E-07 |
| Fam171b | red | -0.8786039 | 0.00191592 | -0.198213 | 0.56492847 |
| Gli2 | red | -0.5817251 | 0.01168153 | -0.9820351 | 2.42E-05 |
| Mlf1 | skyblue | -2.0504711 | 4.22E-14 | -1.5327821 | 4.00E-08 |
| Smim5 | skyblue | -1.5449991 | 2.98E-08 | -1.0654689 | 0.00020654 |
| Sprr1b | skyblue | -2.3355198 | 0.00153315 | -1.1605718 | 0.14541102 |
| Fam212b | red | -1.0726741 | 2.53E-12 | -0.5512004 | 0.00061876 |
| Mypop | red | -0.6511083 | 0.00767669 | -0.8005214 | 0.00109097 |
| Bdnf | red | -0.9475133 | 0.0001962 | -0.5211815 | 0.05444982 |
| Fitm2 | red | -0.716798 | 1.14E-05 | -0.7401693 | 8.76E-06 |
| Fbxl13 | skyblue | -2.7670612 | 4.97E-13 | -1.1620231 | 0.00125303 |
| Cxcr6 | yellow | 4.20160403 | 2.15E-64 | 3.47754566 | 9.30E-44 |
| Amica1 | yellow | 3.81266368 | 2.50E-31 | 2.66564384 | 7.92E-16 |
| Phldb1 | red | -0.7738871 | 1.56E-07 | -0.6186394 | 4.76E-05 |
| Tmem252 | orange | 0.71981714 | 0.00438405 | 0.25533088 | 0.36764959 |
| Gm6377 | yellow | 3.47274008 | 1.79E-12 | 1.97002224 | 2.58E-05 |
| Ctxn1 | skyblue | -2.0189844 | 1.33E-23 | -1.1942254 | 8.07E-09 |

| | | | | | |
|---|---|---|---|---|---|
| Tpcn2 | orange | 1.16757037 | 3.36E-08 | 1.17835057 | 4.08E-08 |
| Lurap1l | red | -1.3867985 | 3.02E-28 | -0.8564279 | 3.68E-11 |
| Tprn | orange | 0.81471422 | 4.67E-05 | 0.40041129 | 0.0622766 |
| Fndc9 | orange | 1.03387225 | 0.00059751 | 0.48233581 | 0.13829776 |
| Zfp454 | skyblue | -2.0583814 | 0.0021178 | -1.346794 | 0.04904493 |
| Ggnbp1 | red | -0.7985412 | 1.39E-06 | -0.9253204 | 5.87E-08 |
| Klhl11 | orange | 0.66571798 | 0.0445292 | 1.19170421 | 0.00033184 |
| Rpl29 | orange | 0.62984752 | 1.67E-08 | 0.56844176 | 6.35E-07 |
| Tmem53 | red | -0.6392322 | 0.00493842 | -0.3620547 | 0.1347641 |
| P2ry6 | yellow | 2.1163198 | 1.01E-08 | 1.67516218 | 1.09E-05 |
| Ccdc37 | skyblue | -1.8188396 | 1.76E-06 | -1.3547765 | 0.00054944 |
| Ifnb1 | black | 6.9757004 | 3.20E-06 | 6.92852553 | 6.19E-06 |
| Lonrf2 | skyblue | -2.7589536 | 7.42E-07 | -2.7758928 | 2.47E-07 |
| Dact2 | red | -1.1089693 | 0.00211361 | -0.6262507 | 0.10512577 |
| Pkd1l3 | orange | 0.93320932 | 0.0004633 | 0.16301289 | 0.60061945 |
| Vstm2a | skyblue | -2.9867295 | 0.00235915 | -1.5091637 | 0.0933323 |
| Gm12185 | yellow | 3.2029912 | 1.78E-48 | 3.10344617 | 8.01E-48 |
| Arhgap30 | yellow | 2.49553228 | 8.16E-26 | 2.57246983 | 3.03E-27 |
| Cdk5r1 | orange | 1.37211223 | 1.83E-06 | 0.9588087 | 0.00127013 |
| 4930539E08I | yellow | 2.75482077 | 1.82E-05 | 3.82576935 | 3.66E-07 |
| Cdca2 | yellow | 3.31190818 | 1.12E-36 | 3.01818606 | 7.00E-31 |
| Prex2 | red | -1.352691 | 1.49E-07 | -1.0957188 | 3.60E-05 |
| Yjefn3 | red | -1.1803163 | 5.70E-05 | -0.9376216 | 0.00162709 |
| Elfn1 | skyblue | -1.4585491 | 7.87E-08 | -1.1733318 | 2.67E-05 |
| Clec4a1 | yellow | 3.07635729 | 3.60E-18 | 2.65619018 | 1.37E-13 |
| Ccr2 | yellow | 2.42449115 | 9.86E-26 | 2.61396709 | 1.85E-29 |
| Ntf3 | skyblue | -1.8533587 | 4.71E-06 | -1.2363443 | 0.00357209 |
| Themis | yellow | 2.90101295 | 1.05E-09 | 4.6396881 | 3.92E-21 |
| Agtr1a | red | -1.0187663 | 0.00087243 | -0.7737471 | 0.01473273 |
| Gm8186 | orange | 0.6868353 | 9.50E-05 | 0.52898248 | 0.00364693 |
| Ivl | orange | 2.26523355 | 0.00347628 | 1.14318275 | 0.1370462 |
| C5ar1 | yellow | 2.68260783 | 1.88E-06 | 2.37305791 | 4.05E-05 |
| Nrap | red | -1.4246159 | 4.37E-08 | -0.8392497 | 0.0021429 |
| Fam183b | skyblue | -2.0138207 | 1.13E-12 | -1.1912849 | 3.93E-05 |
| Myoz3 | skyblue | -2.326843 | 2.92E-07 | -1.5632823 | 0.00116975 |
| Purg | red | -1.062671 | 5.31E-07 | -0.9169305 | 2.91E-05 |
| Hcar1 | skyblue | -3.2675162 | 1.16E-07 | -2.346315 | 0.00011927 |
| Kcnk3 | red | -1.0171304 | 2.32E-05 | -1.0118104 | 3.89E-05 |
| Scn3b | red | -0.8680773 | 3.00E-05 | -1.0734293 | 3.41E-07 |
| Syt12 | orange | 1.73800028 | 2.02E-07 | 1.82687684 | 5.71E-08 |
| Fut4 | orange | 1.53198199 | 1.43E-09 | 1.07717719 | 3.57E-05 |
| Zfp2 | skyblue | -1.5929218 | 0.00175657 | -1.2824219 | 0.01494929 |
| Krt8 | orange | 1.50752228 | 1.37E-06 | 1.58976581 | 5.37E-07 |

| | | | | | |
|---|---|---|---|---|---|
| Ogfr | orange | 1.94659293 | 5.62E-26 | 1.81633525 | 1.26E-22 |
| Zfp683 | yellow | 3.5927593 | 8.27E-13 | 3.55109901 | 6.70E-11 |
| Tmem200a | skyblue | -1.7258001 | 1.16E-19 | -1.3659779 | 9.65E-13 |
| Chchd10 | red | -1.3359249 | 2.27E-07 | -1.0030918 | 0.00017944 |
| Ctu2 | orange | 0.7740726 | 1.36E-06 | 0.75457364 | 4.27E-06 |
| Slc36a3 | black | 4.83118866 | 0.00209699 | 2.87738168 | 0.02483603 |
| Dtx3l | yellow | 2.26363423 | 3.00E-46 | 2.19659493 | 1.19E-43 |
| Htr1b | skyblue | -2.6838399 | 6.69E-12 | -1.6521828 | 3.80E-06 |
| Rps23 | orange | 0.58569735 | 3.28E-10 | 0.41760282 | 1.53E-05 |
| Cdc42ep1 | red | -0.9489085 | 4.65E-13 | -0.8169318 | 9.51E-10 |
| Sall2 | red | -1.2274369 | 2.19E-14 | -0.9474548 | 8.34E-09 |
| Tceal1 | skyblue | -1.5400434 | 3.90E-13 | -1.2159968 | 2.48E-08 |
| Tmie | red | -0.9522582 | 0.00840928 | -0.7420002 | 0.0492554 |
| Lingo1 | skyblue | -1.1010905 | 0.00425443 | -2.0249567 | 2.89E-07 |
| Ap5b1 | orange | 0.87696953 | 1.15E-05 | 0.83010911 | 5.34E-05 |
| Cfap46 | skyblue | -2.0867584 | 5.11E-17 | -1.1574633 | 4.31E-06 |
| Ccdc69 | orange | 0.71342553 | 7.82E-05 | 1.10069059 | 6.84E-09 |
| Lce1h | skyblue | -1.6823581 | 2.61E-05 | -0.6896141 | 0.11040243 |
| Gpr55 | yellow | 4.06901008 | 1.09E-22 | 3.68823733 | 4.22E-18 |
| Slc17a5 | red | -0.7470332 | 0.00079183 | -0.7946261 | 0.00047163 |
| Tifab | yellow | 2.26189324 | 4.83E-09 | 2.12873242 | 6.48E-08 |
| Zbtb14 | red | -0.8808251 | 8.93E-06 | -0.7124575 | 0.00048805 |
| Nckap5 | red | -0.9187556 | 1.33E-11 | -0.9150717 | 2.75E-11 |
| Nlrp10 | red | -0.9015037 | 0.00735249 | -0.8612587 | 0.01226435 |
| Prss46 | black | 6.35339685 | 1.58E-05 | 6.80841606 | 6.06E-06 |
| Gal3st1 | red | -1.1076182 | 0.00015527 | -0.7398162 | 0.01513619 |
| Mmp12 | orange | 1.60573047 | 0.00195791 | 1.06030965 | 0.05300583 |
| Zfp668 | orange | 0.84589956 | 1.92E-09 | 0.36581006 | 0.01467513 |
| Arhgap15 | orange | 1.98448698 | 5.09E-13 | 1.843129 | 4.81E-11 |
| Rpl36al | orange | 2.10187126 | 5.06E-08 | 1.07613705 | 0.00469432 |
| Tmsb4x | orange | 0.73514367 | 2.22E-10 | 0.44626719 | 0.00022907 |
| Armcx4 | orange | 0.63476724 | 0.00015913 | 0.72831566 | 2.09E-05 |
| Arhgap23 | red | -0.6110377 | 5.43E-08 | -0.4317498 | 0.00021398 |
| Fam161a | skyblue | -2.2605433 | 1.12E-18 | -1.5618507 | 2.15E-09 |
| Arl4c | orange | 0.81253123 | 5.74E-07 | 0.93927994 | 1.06E-08 |
| Nlrc3 | yellow | 2.66339854 | 2.72E-22 | 2.4921321 | 2.36E-19 |
| 2810025M15 | red | -0.5310483 | 0.0023952 | -0.7798521 | 9.76E-06 |
| Vcpkmt | orange | 1.01908094 | 1.87E-05 | 0.92482252 | 0.00017669 |
| Rasd1 | orange | 1.15598986 | 0.00030561 | 0.59992812 | 0.07836533 |
| Rasl11b | orange | 0.69481096 | 2.49E-05 | 0.70635683 | 2.78E-05 |
| 2610318N02 | yellow | 3.13703948 | 7.19E-06 | 4.48779472 | 2.06E-09 |
| H2afx | orange | 1.40085477 | 3.17E-11 | 1.21965445 | 1.81E-08 |
| Lrrc4 | yellow | 2.53441954 | 1.99E-06 | 2.62626144 | 2.93E-06 |

| | | | | | |
|---|---|---|---|---|---|
| Mrps16 | orange | 0.69619257 | 7.12E-06 | 0.46310402 | 0.00471852 |
| Ankrd55 | yellow | 2.45560522 | 6.92E-05 | 3.70742891 | 4.62E-07 |
| Lrrc25 | yellow | 4.38463558 | 5.06E-36 | 3.95970683 | 8.95E-30 |
| Idnk | orange | 0.94873062 | 6.18E-34 | 0.85227175 | 1.43E-27 |
| Apol10b | yellow | 3.72396786 | 9.02E-52 | 3.08982649 | 1.93E-36 |
| Rap2c | orange | 1.00474185 | 6.76E-09 | 0.87053093 | 9.70E-07 |
| Tdrp | red | -0.8982482 | 0.00105813 | -0.7102735 | 0.01276048 |
| Zfp697 | orange | 0.64712072 | 0.00142141 | 0.49551493 | 0.01911313 |
| Grem2 | skyblue | -2.3348963 | 4.85E-24 | -1.6261517 | 5.32E-12 |
| Gpr171 | yellow | 4.23463061 | 4.86E-80 | 4.28676641 | 1.51E-67 |
| Ces2b | skyblue | -5.0684993 | 8.15E-15 | -2.9624894 | 5.12E-09 |
| Agmo | skyblue | -1.6738169 | 6.60E-05 | -1.457259 | 0.00074802 |
| Grrp1 | orange | 0.67135711 | 5.05E-06 | 0.51138134 | 0.00092915 |
| Tmc8 | orange | 1.10696727 | 9.77E-08 | 1.22050483 | 6.87E-09 |
| Gsg2 | yellow | 2.58601816 | 1.24E-14 | 2.88181349 | 1.21E-16 |
| Vwa3b | skyblue | -1.7016276 | 1.59E-07 | -0.9661222 | 0.00364483 |
| Kcnk12 | skyblue | -2.1569815 | 3.31E-07 | -0.8799198 | 0.04249518 |
| F2rl3 | orange | 2.00312067 | 1.77E-06 | 1.43910471 | 0.00108436 |
| Gm867 | red | -1.0971802 | 1.03E-07 | -0.5886897 | 0.00588016 |
| Lgr4 | red | -0.6186474 | 0.00023558 | -0.7734352 | 5.67E-06 |
| Eva1b | red | -0.7108736 | 2.76E-06 | -0.5705948 | 0.0002831 |
| Il17d | skyblue | -1.7748309 | 6.47E-10 | -0.8738843 | 0.0042424 |
| Cxcr3 | yellow | 4.49193828 | 1.88E-49 | 3.93307487 | 9.77E-39 |
| Klre1 | orange | 1.6278577 | 6.63E-06 | 1.39146963 | 0.00024518 |
| Heatr1 | orange | 0.95172359 | 1.14E-08 | 0.81361188 | 2.03E-06 |
| Evc2 | red | -0.9934984 | 1.76E-13 | -0.6989778 | 4.25E-07 |
| Tmem220 | red | -1.0026908 | 0.00023606 | -0.9195704 | 0.00087688 |
| Fzd2 | red | -1.2759182 | 7.76E-12 | -0.7039041 | 0.00032749 |
| Gm9843 | orange | 1.15640511 | 0.000108 | 0.19352966 | 0.59452712 |
| Synpo2 | red | -1.1822186 | 1.99E-18 | -0.762666 | 4.81E-08 |
| Lgals3 | orange | 1.84474124 | 7.20E-10 | 1.69085937 | 2.97E-08 |
| Gpr18 | yellow | 2.39372847 | 6.95E-24 | 2.14681671 | 1.77E-19 |
| Rltpr | orange | 1.57477485 | 4.32E-06 | 2.30947534 | 1.61E-11 |
| Ch25h | black | 4.69067944 | 1.13E-44 | 4.38303783 | 4.77E-39 |
| Snx21 | red | -0.5899024 | 0.00408082 | -0.7848311 | 0.00015306 |
| Sept6 | orange | 1.43410674 | 5.64E-30 | 1.48028403 | 6.94E-31 |
| Tcf19 | orange | 1.62831293 | 1.59E-21 | 1.43687172 | 7.43E-17 |
| Enthd1 | black | 4.79771693 | 0.00150667 | 3.64349465 | 0.00370354 |
| Hamp | skyblue | -1.8417462 | 9.43E-06 | -1.4232885 | 0.00080239 |
| Lypd6 | skyblue | -2.2028757 | 0.0003482 | -1.0675733 | 0.09509652 |
| Krt78 | red | -1.0272448 | 0.00360034 | -0.7161237 | 0.05452806 |
| Slc35d3 | skyblue | -2.5424473 | 7.68E-05 | -0.9613297 | 0.14110948 |
| Fbxl22 | red | -1.3057936 | 1.43E-05 | -0.8193441 | 0.00956962 |

| | | | | | |
|---|---|---|---|---|---|
| Cldn8 | skyblue | -2.8357 | 4.84E-17 | -2.116452 | 3.13E-11 |
| Fam171a1 | red | -1.1160176 | 8.70E-09 | -0.9748645 | 9.12E-07 |
| 5730508B09I | orange | 0.64429503 | 0.0092285 | 0.65699353 | 0.0100529 |
| Hyls1 | orange | 1.13200467 | 1.77E-11 | 0.8858359 | 2.78E-07 |
| Kcnb1 | red | -0.8848447 | 8.83E-07 | -0.4434431 | 0.02109278 |
| Tor1aip2 | orange | 0.86909386 | 1.20E-08 | 0.84298109 | 5.57E-08 |
| Mmp13 | yellow | 3.66291551 | 8.80E-11 | 2.71238342 | 1.25E-06 |
| Lrrc4c | skyblue | -1.0936654 | 0.03803569 | -1.9814576 | 0.00016881 |
| Fam78a | orange | 1.58764769 | 1.53E-20 | 1.58008118 | 5.18E-20 |
| Zfp831 | orange | 1.40961441 | 1.37E-06 | 1.3912585 | 4.12E-06 |
| Ccdc121 | skyblue | -2.5082348 | 5.36E-16 | -1.2242698 | 3.07E-05 |
| Gpd1l | red | -0.758102 | 2.12E-08 | -0.647917 | 3.07E-06 |
| Tmem150c | red | -0.6831305 | 0.00228947 | -0.6234536 | 0.00714047 |
| Ccdc96 | skyblue | -1.6421815 | 2.58E-11 | -0.9859743 | 7.24E-05 |
| Ftl1 | orange | 0.72790576 | 0.0011147 | 0.43181784 | 0.06962969 |
| Plekho2 | orange | 1.73272423 | 5.51E-11 | 1.6545519 | 6.52E-10 |
| Arhgap42 | red | -1.0843746 | 2.30E-05 | -0.747397 | 0.00510901 |
| Ptges | orange | 1.30773698 | 9.80E-06 | 1.31488485 | 1.34E-05 |
| Trim15 | orange | 1.21059328 | 0.00605317 | 1.70963314 | 0.0001401 |
| Gm5084 | skyblue | -2.5165189 | 5.86E-05 | -0.844231 | 0.14929956 |
| Tmem37 | orange | 1.57381168 | 1.64E-18 | 1.42587798 | 3.13E-15 |
| Fam131a | red | -0.9537518 | 1.55E-07 | -0.8986898 | 1.35E-06 |
| Rtn4rl2 | yellow | 2.81800583 | 2.90E-15 | 2.59813802 | 6.17E-13 |
| Tvp23a | yellow | 2.14244618 | 3.31E-12 | 2.31300821 | 3.78E-13 |
| Cdr2l | red | -0.8648057 | 2.03E-06 | -0.4252627 | 0.02844283 |
| Ankrd37 | orange | 1.41242128 | 2.65E-07 | 0.50170182 | 0.10758068 |
| P2ry10 | yellow | 2.05167698 | 2.25E-13 | 2.56931279 | 1.89E-19 |
| Sgms2 | orange | 0.67303671 | 0.09070555 | 1.14919846 | 0.00384941 |
| Zfp438 | red | -0.7989115 | 0.00112043 | -0.4814359 | 0.06338336 |
| Amigo1 | red | -1.1759541 | 3.22E-11 | -0.9355339 | 2.99E-07 |
| Zfp169 | orange | 0.73462413 | 0.00224884 | 1.20272081 | 9.35E-07 |
| Insl6 | yellow | 2.56644032 | 2.06E-07 | 1.20740895 | 0.01461758 |
| Creg2 | yellow | 2.56795743 | 1.12E-05 | 2.78772528 | 3.61E-05 |
| Adgb | skyblue | -1.9202762 | 1.12E-09 | -1.0760717 | 0.00087628 |
| Hs3st1 | red | -1.3283363 | 5.65E-05 | -0.7571683 | 0.02999816 |
| Zfp11 | red | -0.7196611 | 0.00218847 | -0.4690746 | 0.05779656 |
| Olfml1 | red | -0.7654514 | 1.80E-05 | -0.6361408 | 0.00056966 |
| Gprc5c | red | -1.2562549 | 6.29E-12 | -0.5429603 | 0.00453905 |
| P4ha3 | yellow | 1.87991776 | 0.00084658 | 2.15353953 | 0.00019387 |
| Lingo3 | skyblue | -2.5337424 | 0.00428409 | -2.9516826 | 0.01396406 |
| Vtcn1 | red | -1.114102 | 0.00679186 | -0.1280242 | 0.79229502 |
| Mblac2 | red | -1.0026006 | 6.05E-06 | -1.0116343 | 1.29E-05 |
| Sv2c | orange | 1.68007508 | 3.73E-06 | 1.05260059 | 0.0065167 |

| | | | | | |
|---|---|---|---|---|---|
| Gimap9 | orange | 1.30490175 | 2.21E-18 | 1.32859327 | 4.08E-18 |
| Camk2n2 | orange | 0.89688257 | 0.00397082 | 0.5373572 | 0.1116838 |
| Nat2 | orange | 0.66834101 | 0.00012365 | 0.29228833 | 0.12894527 |
| Adnp | orange | 0.62984341 | 0.0700545 | 0.95259039 | 0.00646008 |
| Plcb1 | red | -0.8127408 | 0.00035963 | -0.9659983 | 3.15E-05 |
| Gpr183 | orange | 1.51512964 | 1.95E-07 | 1.17868332 | 0.00010145 |
| Ercc6l | orange | 2.04654334 | 1.26E-08 | 1.91177303 | 3.66E-07 |
| Gen1 | yellow | 2.61282397 | 1.52E-11 | 2.16554711 | 9.34E-08 |
| Cml3 | skyblue | -2.1831127 | 7.63E-08 | -1.8385598 | 9.17E-06 |
| Zgrf1 | orange | 1.35755152 | 3.94E-06 | 1.49919827 | 8.74E-07 |
| Ffar2 | yellow | 1.97279571 | 2.48E-09 | 2.3515914 | 1.39E-12 |
| 2900026A02I | red | -0.6925137 | 3.59E-13 | -0.54476 | 2.42E-08 |
| Zfp52 | orange | 0.78823791 | 0.00016975 | 1.08560983 | 4.29E-07 |
| Samd3 | yellow | 2.07495952 | 7.47E-07 | 2.40709313 | 9.72E-08 |
| Ncald | red | -0.7913833 | 5.83E-06 | -0.5812478 | 0.00138981 |
| Six1 | red | -1.3426445 | 2.82E-05 | -0.6036668 | 0.08020208 |
| Ppapdc3 | red | -1.3696572 | 7.08E-10 | -0.9175405 | 0.00014864 |
| Pcdh1 | red | -1.1459358 | 1.45E-14 | -0.8870532 | 6.12E-09 |
| Kif18b | yellow | 3.37624757 | 6.14E-29 | 3.85836778 | 6.25E-34 |
| Gm45902 | orange | 1.21871847 | 2.96E-06 | 0.70482503 | 0.01164619 |
| Tacstd2 | orange | 1.12884287 | 7.87E-12 | 1.25818982 | 2.62E-14 |
| Plagl2 | orange | 0.61412823 | 0.00032478 | 0.56073719 | 0.00139081 |
| Ccdc157 | red | -0.9358713 | 1.09E-08 | -0.6358493 | 0.00017787 |
| Fhad1 | red | -1.4496291 | 1.19E-15 | -0.8836824 | 1.84E-06 |
| Cd14 | orange | 1.00750456 | 6.25E-05 | 1.27383182 | 5.22E-07 |
| Spn | yellow | 2.86175075 | 4.02E-59 | 2.62820553 | 1.22E-49 |
| Cbr1 | red | -1.0799952 | 4.83E-08 | -0.7512825 | 0.00025768 |
| Tlr6 | yellow | 2.46989049 | 1.30E-09 | 2.00191213 | 1.36E-06 |
| Siglech | orange | 0.49119086 | 0.3608591 | 1.52937626 | 0.002425 |
| Wdfy4 | orange | 0.8323446 | 0.00070863 | 0.77825928 | 0.00210403 |
| Fam181b | skyblue | -1.9883379 | 0.00077136 | -1.4755027 | 0.01175941 |
| Rps19bp1 | orange | 0.81738112 | 0.00010276 | 0.6540217 | 0.00310896 |
| Zfp579 | red | -0.5501697 | 0.00045626 | -0.6298348 | 8.20E-05 |
| Map3k19 | skyblue | -2.1715202 | 1.92E-11 | -1.1710104 | 0.00030525 |
| Otop1 | skyblue | -4.2051852 | 0.00176225 | -3.1415839 | 0.02497844 |
| 2010001K21I | skyblue | -2.1250978 | 0.00017442 | -1.8920479 | 0.00097718 |
| Rap2a | red | -0.7071452 | 1.49E-07 | -0.4783222 | 0.00066834 |
| Hist1h1e | red | -1.6745889 | 0.00318185 | -0.4133813 | 0.52896906 |
| B3gnt2 | orange | 0.78743419 | 0.02053634 | 0.92801658 | 0.00776292 |
| AU021092 | red | -1.2686161 | 1.10E-07 | -1.0634946 | 1.47E-05 |
| Dcun1d4 | red | -0.7790664 | 5.10E-05 | -0.765015 | 9.47E-05 |
| Trim32 | red | -0.711842 | 2.46E-09 | -0.4938627 | 6.69E-05 |
| Apon | yellow | 2.33694848 | 0.00267178 | 2.56402159 | 0.00292525 |

| | | | | | |
|---|---|---|---|---|---|
| Rinl | orange | 2.01229794 | 9.87E-67 | 1.95646134 | 1.71E-61 |
| Ttn | red | -1.1071645 | 0.00023457 | -1.0259226 | 0.00092409 |
| Nlgn2 | orange | 0.65112355 | 0.00112776 | 0.55756996 | 0.00707791 |
| Cox6b2 | red | -1.0290755 | 7.76E-09 | -0.8798192 | 1.85E-06 |
| Mest | red | -0.7722808 | 0.00042859 | -0.8214694 | 0.00027593 |
| Klhdc1 | red | -1.1142461 | 0.00017709 | -1.061741 | 0.00052673 |
| Cd209f | skyblue | -3.2356413 | 1.05E-09 | -2.20296 | 5.54E-05 |
| Sox6 | red | -1.2408948 | 5.00E-09 | -0.9150943 | 3.56E-05 |
| Sec31b | red | -0.835492 | 0.00168292 | -0.587092 | 0.03489359 |
| Lax1 | yellow | 2.95525219 | 5.05E-32 | 2.6256569 | 5.46E-25 |
| Btla | orange | 1.43822911 | 6.47E-15 | 1.50932078 | 8.30E-16 |
| Slc6a7 | skyblue | -2.0557878 | 0.00056196 | -1.892869 | 0.00537694 |
| Pfdn4 | orange | 0.61274938 | 0.00441766 | 0.37107044 | 0.11371487 |
| Dock8 | orange | 0.60675772 | 4.85E-05 | 0.48954076 | 0.00157139 |
| Rgs14 | yellow | 1.82857451 | 1.89E-13 | 2.09579222 | 6.12E-17 |
| Gnpda1 | orange | 0.88560271 | 7.11E-11 | 0.81748197 | 4.65E-09 |
| Mtcl1 | red | -1.4895911 | 5.21E-08 | -1.0136364 | 0.00036733 |
| D630039A03 | red | -1.5041963 | 2.93E-07 | -0.6947195 | 0.024587 |
| F730043M19 | orange | 1.75312687 | 5.05E-13 | 1.85135785 | 3.64E-14 |
| Akr1b7 | skyblue | -1.4020273 | 0.07489506 | -2.4797932 | 0.00256066 |
| Foxo6 | skyblue | -1.508955 | 1.80E-06 | -0.9001015 | 0.00714922 |
| Bre | orange | 0.61297831 | 2.00E-05 | 0.45890551 | 0.00223711 |
| Rasal3 | yellow | 2.23822346 | 1.87E-32 | 2.37166774 | 5.80E-36 |
| Gm9869 | orange | 1.48130595 | 0.00106169 | 1.84740996 | 0.0001387 |
| Pld4 | yellow | 2.70271858 | 2.14E-14 | 2.57359463 | 6.04E-13 |
| Cd177 | orange | 1.79437451 | 2.54E-12 | 1.82009969 | 6.54E-13 |
| Ppp1r36 | skyblue | -2.1166694 | 7.58E-09 | -1.1245333 | 0.00293805 |
| Zeb2os | orange | 1.65943463 | 0.00735596 | 1.40852796 | 0.03368535 |
| Fpr2 | yellow | 2.63928936 | 1.04E-07 | 2.16782142 | 2.12E-05 |
| Dnah3 | skyblue | -1.8097987 | 1.33E-06 | -0.9261794 | 0.01867899 |
| Tbc1d30 | skyblue | -1.9988784 | 1.31E-11 | -1.5337546 | 4.15E-07 |
| Hbb-bs | red | -0.4203381 | 0.28026506 | -1.0410674 | 0.00518063 |
| Lrrc15 | skyblue | -2.4037431 | 0.00273387 | -2.2505541 | 0.00776963 |
| Cx3cr1 | orange | 0.90258699 | 0.00840595 | 0.67466935 | 0.06376742 |
| Cemip | orange | 0.34333821 | 0.58014076 | 1.99128155 | 0.00045706 |
| Tmem106c | red | -0.69127 | 1.30E-06 | -0.5049474 | 0.00073285 |
| Actn2 | skyblue | -1.7294333 | 1.08E-06 | -1.4281042 | 9.52E-05 |
| Nrros | orange | 1.45367981 | 3.12E-12 | 1.57749097 | 4.94E-14 |
| Ccdc171 | red | -0.8664713 | 0.00404172 | -0.475654 | 0.14187557 |
| Tchh | red | -0.8808918 | 0.0008857 | -0.5159686 | 0.06406441 |
| Bmpr1b | skyblue | -2.009215 | 4.75E-09 | -0.8974353 | 0.01164445 |
| Cebpe | black | 2.60736898 | 0.00189317 | 5.0252932 | 4.63E-05 |
| C130026I21F | orange | 1.56503287 | 0.0175577 | 1.97900813 | 0.00217517 |

| | | | | | |
|---|---|---|---|---|---|
| Tmem171 | yellow | 3.20166611 | 1.17E-13 | 3.23909586 | 2.17E-13 |
| Nav2 | red | -0.60433 | 0.00050169 | -0.3583064 | 0.05234431 |
| Robo2 | skyblue | -1.6227072 | 7.28E-11 | -1.4247454 | 2.20E-08 |
| Pbx1 | red | -0.597702 | 7.76E-13 | -0.5305078 | 4.27E-10 |
| Magi3 | red | -1.0169068 | 2.16E-06 | -0.5427113 | 0.01707498 |
| D930048N14 | red | -0.7836045 | 2.49E-07 | -0.6560799 | 2.59E-05 |
| Zmat1 | red | -0.7324896 | 0.00395061 | -0.9873541 | 0.00012353 |
| Rap1b | orange | 0.73375695 | 4.91E-06 | 0.63838943 | 0.00011335 |
| Rab7b | orange | 1.53627366 | 4.72E-08 | 1.17659337 | 5.34E-05 |
| Tln2 | red | -1.2701764 | 1.43E-08 | -1.1189363 | 1.07E-06 |
| Zfp608 | red | -0.5966655 | 0.00027505 | -0.2732108 | 0.12448022 |
| Gm9888 | yellow | 1.72287964 | 0.01324872 | 2.21819386 | 0.00907409 |
| Map1b | red | -0.8913376 | 2.87E-10 | -0.6007155 | 4.18E-05 |
| Klrc2 | yellow | 2.47742609 | 5.78E-13 | 1.76533867 | 4.09E-07 |
| Swt1 | red | -0.3875794 | 0.03388336 | -0.6040454 | 0.00097882 |
| Trim30b | yellow | 3.0091351 | 6.27E-19 | 3.67572105 | 4.92E-27 |
| Gpr25 | black | 4.57650898 | 0.00042203 | 5.4678027 | 0.00024669 |
| A630001G21 | orange | 1.96168295 | 3.42E-16 | 1.75944861 | 6.88E-13 |
| Gm9889 | orange | 1.91883478 | 9.87E-05 | 1.46419111 | 0.00280232 |
| Oas1a | yellow | 4.37674319 | 1.47E-56 | 3.91980046 | 1.13E-45 |
| Grk4 | red | -0.7579674 | 0.00333894 | -0.5470616 | 0.04506024 |
| 1700030K09I | red | -0.648123 | 0.00161839 | -0.4747901 | 0.02700274 |
| Nup107 | orange | 0.70822278 | 2.54E-09 | 0.61287307 | 5.88E-07 |
| Cysltr1 | orange | 0.90681204 | 0.00474148 | 0.0594542 | 0.88556426 |
| Gm9892 | orange | 1.93766804 | 0.00138115 | 1.24568589 | 0.06792232 |
| Junb | orange | 1.73683658 | 4.61E-27 | 1.62197424 | 1.06E-23 |
| Dnah6 | skyblue | -2.616004 | 6.41E-13 | -1.6524593 | 1.07E-05 |
| Prkcb | yellow | 1.95857899 | 2.58E-20 | 2.10727015 | 2.34E-22 |
| Lamb2 | red | -1.0155005 | 1.27E-10 | -0.762154 | 2.85E-06 |
| Cyp2j6 | red | -0.7565559 | 3.61E-11 | -0.4341732 | 0.00032919 |
| Prkg1 | red | -1.01334 | 0.00458736 | -0.4759503 | 0.22102697 |
| Arhgef15 | red | -1.1402274 | 1.82E-11 | -0.9904556 | 1.04E-08 |
| Rnaseh2a | orange | 0.60536172 | 1.38E-05 | 0.41262104 | 0.0049842 |
| Glis3 | red | -0.8217035 | 5.90E-06 | -0.6359439 | 0.00056393 |
| Rnf157 | orange | 0.81654375 | 0.00018206 | 0.89433003 | 5.56E-05 |
| C130021I20F | skyblue | -3.1774043 | 0.00159377 | -1.7186037 | 0.05251035 |
| Gas1 | skyblue | -1.2791752 | 4.38E-07 | -1.2368503 | 1.94E-06 |
| Cyp2f2 | skyblue | -2.7154508 | 2.21E-08 | -1.7222624 | 0.00067943 |
| Creb5 | orange | 1.42258276 | 5.78E-08 | 1.14553537 | 2.36E-05 |
| Gm6180 | orange | 1.27345433 | 7.92E-05 | 0.87047598 | 0.01145892 |
| Cd8b1 | yellow | 3.0664243 | 2.31E-06 | 4.07262997 | 4.99E-10 |
| Gm15413 | yellow | 4.56718757 | 0.00286332 | 1.60039775 | 0.23826564 |
| Clec12a | yellow | 3.35246752 | 1.01E-14 | 2.7712292 | 4.07E-10 |

| | | | | | |
|---|---|---|---|---|---|
| 9230110C19 | red | -1.1843369 | 5.23E-05 | -0.5600758 | 0.06895051 |
| 2700081O15 | red | -1.2768523 | 1.16E-13 | -0.9132832 | 2.63E-07 |
| Myh7 | skyblue | -2.2797102 | 2.50E-07 | -1.5558538 | 0.00071058 |
| Gpr141 | yellow | 3.85283327 | 1.42E-18 | 3.44116984 | 3.56E-15 |
| Yap1 | red | -0.6954008 | 1.33E-09 | -0.4620165 | 0.00010662 |
| Fank1 | red | -1.3752883 | 0.00152509 | -0.6864366 | 0.14447516 |
| Socs3 | yellow | 2.95872179 | 1.11E-42 | 3.12391357 | 1.79E-47 |
| Spag16 | skyblue | -3.3032545 | 3.24E-18 | -1.7692245 | 3.98E-06 |
| Fes | orange | 1.4793281 | 1.47E-10 | 1.35056889 | 8.95E-09 |
| Daw1 | skyblue | -1.9529148 | 1.07E-07 | -1.2955531 | 0.00034103 |
| Gpr21 | skyblue | -2.8318498 | 0.00022162 | -2.1412515 | 0.00490485 |
| Cdh22 | skyblue | -1.035835 | 0.00648859 | -1.5508351 | 0.0001065 |
| Bcl3 | yellow | 2.2213804 | 3.86E-37 | 2.35135793 | 2.56E-41 |
| Arhgap20 | red | -1.0066519 | 0.00111198 | -0.6509976 | 0.04656733 |
| Gm9899 | skyblue | -1.8158505 | 0.00150535 | -1.9521556 | 0.00080752 |
| Dand5 | red | -0.7216501 | 0.00150535 | -0.6815928 | 0.00379678 |
| Aldh1a1 | skyblue | -1.9119987 | 4.00E-08 | -1.4753989 | 4.07E-05 |
| Rab4b | orange | 0.67162986 | 0.00831255 | 0.63355898 | 0.01350566 |
| AI854703 | red | -1.3002527 | 0.0092285 | -0.5577656 | 0.2980491 |
| Sec61b | orange | 0.76649107 | 2.52E-07 | 0.65052487 | 2.30E-05 |
| Slamf8 | black | 4.76183887 | 2.59E-40 | 4.12691662 | 5.27E-31 |
| D10Jhu81e | red | -0.8283241 | 1.15E-05 | -0.6430308 | 0.00103415 |
| Gas5 | orange | 0.65509537 | 8.42E-12 | 0.65195385 | 2.06E-11 |
| Tarm1 | black | 7.59837197 | 2.68E-31 | 7.37036926 | 2.03E-27 |
| Phgdh | orange | 1.60942322 | 9.86E-26 | 1.38530784 | 4.40E-19 |
| Cbx7 | red | -1.0043436 | 8.68E-06 | -0.7359965 | 0.00169349 |
| Hunk | red | -0.9466119 | 2.22E-05 | -0.7267856 | 0.00163557 |
| Ggcx | red | -1.1726673 | 6.81E-08 | -0.9204615 | 3.96E-05 |
| Tg | yellow | 2.28710824 | 4.17E-10 | 1.54898125 | 2.02E-05 |
| Lgals7 | red | -1.1328781 | 3.43E-05 | -0.6679824 | 0.02095695 |
| Shisa7 | red | -1.4102975 | 0.00047731 | -0.8600494 | 0.03943999 |
| Ebf4 | red | -0.9127064 | 0.00377115 | -0.8027258 | 0.01342924 |
| 3110082I17R | orange | 0.76721083 | 0.00132543 | 0.59436014 | 0.0171444 |
| Zfand2a | orange | 0.57940157 | 0.00123757 | 0.64118821 | 0.00045706 |
| Gm9913 | yellow | 1.11238503 | 0.25408558 | 5.58535916 | 0.00036095 |
| Sh3pxd2a | red | -0.8882178 | 3.61E-09 | -0.5615973 | 0.0003479 |
| Tll1 | red | -0.7373825 | 5.07E-05 | -0.9294373 | 1.40E-06 |
| Dennd4a | orange | 1.28715919 | 1.24E-17 | 0.99985678 | 8.12E-11 |
| Aldh7a1 | red | -0.7739877 | 1.93E-10 | -0.5457772 | 1.53E-05 |
| Plxnb1 | red | -0.8795639 | 8.00E-06 | -0.5410289 | 0.00890491 |
| Dpep2 | yellow | 3.07399093 | 3.26E-09 | 1.36272987 | 0.0081782 |
| Nebl | red | -0.9845488 | 0.00070435 | -0.574602 | 0.06358522 |
| Ptrh1 | orange | 0.6136728 | 0.00784263 | 0.61576338 | 0.00934421 |

| | | | | | |
|---|---|---|---|---|---|
| 1700016K19l | red | -0.9402278 | 0.00075503 | -0.8284324 | 0.00405753 |
| Grwd1 | orange | 0.9781743 | 0.00015185 | 0.9008554 | 0.00071927 |
| Camk2d | orange | 0.67987456 | 8.99E-07 | 0.66366348 | 2.68E-06 |
| H2-T24 | orange | 1.64014452 | 9.01E-30 | 1.57350044 | 2.30E-27 |
| Sh2d4a | red | -1.240328 | 1.54E-13 | -0.7099962 | 5.13E-05 |
| Cyhr1 | red | -0.5845103 | 2.12E-08 | -0.4529514 | 2.60E-05 |
| 6330403A02l | red | -1.420103 | 6.16E-06 | -0.8320707 | 0.00979358 |
| Pde5a | skyblue | -1.9800375 | 4.67E-17 | -1.5578041 | 9.82E-11 |
| Cd8a | yellow | 3.12862238 | 2.55E-05 | 4.57682599 | 7.55E-10 |
| Zfp14 | red | -1.3046948 | 0.00073263 | -1.1307664 | 0.00467031 |
| Tusc1 | orange | 0.84112529 | 0.00056369 | 0.56430968 | 0.03265677 |
| Ndst1 | red | -1.0089642 | 4.61E-09 | -0.5645847 | 0.00178816 |
| Tmem179 | skyblue | -2.4724835 | 0.00056911 | -1.9484368 | 0.00554033 |
| Sirt5 | red | -1.0060186 | 2.81E-09 | -0.8506414 | 1.33E-06 |
| A930004D18 | red | -0.8177099 | 1.58E-06 | -0.732524 | 4.36E-05 |
| Iigp1 | black | 4.65996271 | 1.43E-103 | 4.38089156 | 1.07E-91 |
| Skida1 | skyblue | -1.6770846 | 1.48E-06 | -1.6207197 | 5.69E-06 |
| Utp18 | orange | 0.68261115 | 4.40E-05 | 0.61799878 | 0.00035399 |
| Syne3 | orange | 1.49599823 | 2.89E-12 | 1.71169982 | 1.98E-15 |
| Fam83e | red | -1.4904602 | 1.15E-06 | -0.6300941 | 0.05154671 |
| Spock3 | skyblue | -2.7094679 | 0.00207047 | -0.7497698 | 0.40790775 |
| Cthrc1 | skyblue | -1.6697651 | 0.00522518 | -1.6373211 | 0.00867396 |
| Ifi205 | black | 4.97757774 | 1.44E-48 | 4.3151539 | 2.99E-37 |
| Fgfr3 | red | -0.7179799 | 0.0021677 | -0.7152744 | 0.00287927 |
| Lifr | red | -0.9162339 | 1.05E-06 | -0.787987 | 4.44E-05 |
| A630033H20 | orange | 2.25114812 | 2.95E-06 | 1.0564822 | 0.03841051 |
| Lrrc36 | skyblue | -2.0368506 | 2.83E-14 | -1.1539851 | 2.26E-05 |
| Taf4b | orange | 1.16423469 | 5.44E-05 | 1.0077208 | 0.0006446 |
| Lce3a | red | -1.1330183 | 0.00044539 | -0.5231953 | 0.13654275 |
| Kcnn4 | yellow | 2.91942023 | 5.20E-19 | 2.76524601 | 6.58E-17 |
| Pnma1 | red | -1.1682642 | 0.00016125 | -1.0734515 | 0.00086617 |
| Cklf | orange | 1.11345083 | 4.29E-07 | 1.05927621 | 2.75E-06 |
| Slfn5 | orange | 1.1952114 | 4.11E-17 | 1.04109735 | 5.02E-13 |
| Fabp1 | skyblue | -4.9923064 | 4.66E-08 | -4.0239282 | 7.00E-07 |
| Tmem120b | red | -0.8434118 | 0.00042228 | -0.4491135 | 0.07882358 |
| Gimap4 | orange | 1.19260911 | 2.42E-10 | 1.04751219 | 5.34E-08 |
| Vsnl1 | skyblue | -1.7203913 | 3.07E-07 | -1.4597649 | 2.41E-05 |
| Kcnn2 | skyblue | -2.8895638 | 2.49E-08 | -2.2114893 | 1.59E-05 |
| Trim65 | red | -0.6961576 | 1.25E-07 | -0.6103912 | 6.19E-06 |
| Zfp867 | red | -0.5079568 | 0.00851878 | -0.6006916 | 0.00195449 |
| Sh3bp2 | yellow | 2.52170509 | 1.35E-20 | 2.36706407 | 3.65E-18 |
| Pla2r1 | red | -1.2268784 | 6.59E-09 | -0.8887847 | 4.69E-05 |
| Pabpc1l | yellow | 2.11180353 | 9.30E-05 | 1.88141603 | 0.00071314 |

| | | | | | |
|---|---|---|---|---|---|
| 9130230L23F | skyblue | -2.1193677 | 5.69E-08 | -1.4744105 | 0.00021492 |
| Mettl7a1 | skyblue | -2.5851041 | 7.41E-06 | -1.9729568 | 0.00097127 |
| Mmrn1 | red | -0.6024774 | 0.00524376 | -0.1861994 | 0.44635124 |
| Pm20d2 | red | -1.3798745 | 0.00286902 | -1.0155078 | 0.03361479 |
| 5830411N06 | orange | 1.11132848 | 0.13940399 | 2.02878219 | 0.0063878 |
| 1600014C10 | orange | 1.53528239 | 2.54E-10 | 1.43956055 | 5.47E-09 |
| Emcn | red | -0.6504428 | 1.82E-05 | -0.7824853 | 3.90E-07 |
| Ap1s3 | orange | 0.88112679 | 5.97E-07 | 0.59455702 | 0.00137116 |
| Ankrd24 | red | -1.006596 | 3.54E-09 | -0.6987224 | 6.93E-05 |
| Zfp771 | orange | 0.32379189 | 0.1187489 | 0.68973001 | 0.00073827 |
| Hmgb2 | yellow | 2.25675716 | 6.66E-34 | 1.87986928 | 1.43E-23 |
| Vmac | red | -0.8448577 | 3.87E-07 | -0.8777246 | 2.11E-07 |
| Phactr1 | red | -0.7674259 | 6.34E-05 | -0.6705907 | 0.00069676 |
| Msra | red | -0.6697243 | 0.0001787 | -0.5354666 | 0.00386162 |
| Fsd1l | skyblue | -1.4925726 | 3.92E-09 | -0.8952629 | 0.0006849 |
| Set | orange | 0.70169024 | 0.0003193 | 0.62845001 | 0.00176618 |
| Cadm4 | red | -1.1644704 | 7.16E-17 | -0.765456 | 1.19E-07 |
| Usp46 | red | -1.0678965 | 2.21E-09 | -0.9054197 | 7.58E-07 |
| Whsc1l1 | orange | 0.6196177 | 2.77E-07 | 0.56532632 | 4.63E-06 |
| Atrnl1 | red | -0.6433339 | 2.82E-06 | -0.4916332 | 0.00055383 |
| Smim10l2a | skyblue | -1.8202634 | 0.00010625 | -0.8010153 | 0.08624365 |
| Rnd1 | orange | 1.41802602 | 6.63E-08 | 1.57117868 | 3.13E-09 |
| Fam19a5 | red | -1.4078193 | 0.00104494 | -0.988768 | 0.02849423 |
| Tmem158 | red | -0.7085854 | 0.00329521 | -0.2523968 | 0.35815108 |
| Dsp | red | -0.7522523 | 0.0036011 | -0.7470848 | 0.00482563 |
| Txk | yellow | 2.10900454 | 1.19E-10 | 2.70094645 | 1.02E-15 |
| Zfp667 | red | -1.0546028 | 0.00239717 | -0.9245874 | 0.01044713 |
| Atp5s | red | -0.6533468 | 0.00128878 | -0.5831471 | 0.0071974 |
| Zkscan4 | skyblue | -1.3720769 | 1.00E-05 | -1.5404156 | 1.49E-06 |
| Afp | yellow | 2.99051289 | 4.61E-06 | 2.22686945 | 0.00054188 |
| Zfp174 | red | -0.8492593 | 0.00280757 | -0.7857573 | 0.00779394 |
| Ifltd1 | skyblue | -1.8149203 | 0.00285356 | -2.4246219 | 8.54E-05 |
| Zfp647 | red | -0.8337514 | 0.0027308 | -0.848671 | 0.00284833 |
| Kbtbd13 | skyblue | -2.1212909 | 4.92E-08 | -1.4756719 | 0.00035693 |
| Sec14l3 | skyblue | -2.1282271 | 2.27E-07 | -1.3934635 | 0.00115959 |
| Agap1 | red | -0.890389 | 3.02E-08 | -0.8371058 | 3.23E-07 |
| Cntn1 | skyblue | -2.0566022 | 0.00064088 | -1.9561557 | 0.00125378 |
| Smyd1 | skyblue | -1.5330489 | 3.43E-09 | -1.4541674 | 4.15E-08 |
| Nfic | red | -0.6351491 | 2.98E-16 | -0.5651473 | 6.66E-13 |
| A630012P03I | yellow | 4.23537484 | 0.00079246 | 1.86373177 | 0.11144424 |
| Arntl | orange | 1.67915759 | 1.10E-08 | 2.4028926 | 5.98E-16 |
| Cgrrf1 | red | -0.5963993 | 0.00016692 | -0.5622909 | 0.00055477 |
| Sugct | red | -1.0664244 | 0.00435798 | -0.2475605 | 0.56829091 |

| | | | | | |
|---|---|---|---|---|---|
| Ifng | black | 7.54189677 | 7.57E-38 | 5.48750246 | 5.97E-50 |
| C1ra | orange | 1.22388096 | 4.21E-26 | 1.14546461 | 8.00E-23 |
| Sertad3 | orange | 0.67472894 | 8.38E-05 | 0.44292547 | 0.01416842 |
| Zfp811 | skyblue | -1.08157 | 0.04472366 | -1.603489 | 0.00856179 |
| Ankrd17 | orange | 0.63408052 | 4.81E-10 | 0.47979247 | 5.05E-06 |
| Ntrk2 | red | -0.0391632 | 0.90801939 | -0.8155746 | 0.00454644 |
| Tmem245 | red | -0.8848743 | 8.07E-08 | -0.5779071 | 0.00078131 |
| Zfp93 | red | -1.2282641 | 3.90E-08 | -0.7975291 | 0.00053719 |
| Pgbd1 | skyblue | -1.4652672 | 0.00246743 | -1.239815 | 0.0126856 |
| Gm9967 | red | -1.2926763 | 1.99E-05 | -1.1153644 | 0.00068158 |
| Fat2 | red | -1.0399689 | 8.69E-06 | -0.719082 | 0.00321952 |
| Slc6a2 | red | -0.9757784 | 1.39E-05 | -0.7306899 | 0.00170394 |
| Fut9 | skyblue | -3.8745905 | 0.00121733 | -1.1450553 | 0.26558578 |
| Fbxo6 | orange | 0.68003287 | 4.85E-08 | 0.62269636 | 1.06E-06 |
| Nell1 | skyblue | -3.5399203 | 0.00141305 | -2.6942464 | 0.00631746 |
| H2-Q5 | orange | 2.01978825 | 9.51E-22 | 1.92591447 | 9.09E-20 |
| Nap1l5 | skyblue | -2.053949 | 0.00062149 | -1.7525832 | 0.00415976 |
| Maf | orange | 0.80559552 | 9.51E-06 | 0.54960973 | 0.00385057 |
| Cd47 | orange | 0.77759222 | 1.88E-09 | 0.47655283 | 0.00042882 |
| Ano5 | red | -1.1393086 | 0.00358925 | -0.9901663 | 0.01710316 |
| Pprc1 | orange | 1.12948126 | 1.39E-06 | 1.18389636 | 6.45E-07 |
| Epm2a | skyblue | -1.5805855 | 0.00010515 | -1.3913427 | 0.0009187 |
| Lair1 | yellow | 3.93748802 | 5.98E-27 | 3.75907556 | 1.64E-24 |
| B4galnt4 | black | 5.17885595 | 2.54E-41 | 4.96742931 | 2.78E-37 |
| Hmcn2 | skyblue | -1.9113282 | 1.68E-14 | -1.1076006 | 2.19E-05 |
| Dach1 | red | -0.730523 | 0.00049966 | -0.547335 | 0.01239773 |
| Gpc3 | red | -1.2338968 | 1.14E-10 | -0.9406447 | 1.86E-06 |
| Kbtbd11 | orange | 1.16577522 | 3.93E-09 | 0.75393465 | 0.0002565 |
| Gm5069 | skyblue | -1.6676344 | 0.00336907 | -1.3365928 | 0.02097285 |
| Klhl26 | red | -0.6971169 | 2.35E-07 | -0.4434026 | 0.00152213 |
| Slain1 | orange | 1.20336539 | 9.40E-08 | 1.18140648 | 3.70E-07 |
| Nap1l3 | skyblue | -2.9061344 | 1.90E-09 | -1.8325472 | 0.00011548 |
| Ghr | red | -0.4791668 | 0.00054093 | -0.5944465 | 2.28E-05 |
| Ldoc1l | red | -0.6875427 | 2.58E-06 | -0.447758 | 0.00366493 |
| Gemin6 | orange | 0.85742036 | 6.00E-05 | 0.70779105 | 0.00175987 |
| Abcd2 | red | -0.7529987 | 8.60E-06 | -0.6823833 | 8.36E-05 |
| Tcf7l1 | red | -0.6063894 | 4.18E-10 | -0.2158552 | 0.03769348 |
| Fmnl1 | yellow | 2.19261354 | 2.43E-24 | 2.1474274 | 2.85E-23 |
| Mta3 | orange | 0.88267894 | 7.96E-09 | 0.70044642 | 9.07E-06 |
| Zfp1 | orange | 0.87030439 | 6.65E-08 | 0.53950439 | 0.00144965 |
| Tceb2 | orange | 0.7368553 | 9.09E-05 | 0.51740003 | 0.00865467 |
| Izumo4 | red | -0.9491838 | 8.04E-05 | -0.6527564 | 0.00967845 |
| Fam19a3 | orange | 0.39342624 | 0.26628784 | 1.05096247 | 0.00378968 |

| | | | | | |
|---|---|---|---|---|---|
| Per2 | red | -0.8294616 | 0.00035707 | -0.8888616 | 0.00017527 |
| Fancm | orange | 0.93515874 | 1.40E-07 | 0.5103102 | 0.00605938 |
| Tmem150a | red | -0.7315952 | 0.00014992 | -0.6848454 | 0.00055362 |
| Aasdh | red | -0.6024483 | 3.19E-05 | -0.4969687 | 0.00102773 |
| Fto | red | -0.6473403 | 3.96E-12 | -0.4127099 | 2.13E-05 |
| Prr18 | skyblue | -2.2097609 | 1.60E-14 | -1.1526843 | 0.00011081 |
| Cldn23 | red | -1.033919 | 8.68E-05 | -0.7548612 | 0.00697197 |
| Irs1 | skyblue | -1.9309843 | 4.42E-05 | -1.0693347 | 0.0324293 |
| Nod2 | orange | 1.42019105 | 3.25E-07 | 1.59354356 | 1.62E-08 |
| A430033K04I | red | -0.8283271 | 0.00011233 | -0.517429 | 0.02130096 |
| Clca3a1 | black | 4.67285332 | 1.67E-80 | 3.64001212 | 6.02E-51 |
| S100a8 | yellow | 2.7145452 | 2.88E-09 | 2.06507843 | 8.61E-06 |
| Fam105a | orange | 1.42059765 | 6.58E-24 | 1.01415876 | 1.79E-12 |
| S100a9 | yellow | 2.44496454 | 9.21E-09 | 2.08423116 | 1.35E-06 |
| Grik2 | skyblue | -1.3826581 | 0.00400022 | -1.9527167 | 0.0002723 |
| Eif3b | orange | 0.53882742 | 0.00023991 | 0.6029191 | 5.41E-05 |
| H2-T22 | yellow | 2.73674849 | 1.51E-79 | 2.43467698 | 4.48E-63 |
| Trim34a | orange | 1.15556555 | 5.52E-17 | 1.19691595 | 6.51E-18 |
| AI504432 | orange | 1.27632203 | 1.27E-17 | 1.55247782 | 7.58E-25 |
| Col8a2 | skyblue | -2.2292335 | 2.11E-13 | -1.1962591 | 0.00013283 |
| Cfl1 | orange | 1.33224586 | 2.31E-09 | 1.22378241 | 7.30E-08 |
| Pgpep1 | red | -0.7502375 | 3.04E-13 | -0.6391939 | 1.21E-09 |
| Npm3 | orange | 0.95976539 | 4.90E-09 | 0.80632536 | 2.00E-06 |
| Pard6g | red | -0.5835074 | 3.04E-05 | -0.2977646 | 0.04610472 |
| Lrguk | skyblue | -2.2603059 | 1.06E-13 | -1.317793 | 1.23E-05 |
| Cep70 | red | -0.6162926 | 0.00188125 | -0.4962899 | 0.0155742 |
| Dennd1b | orange | 1.00317451 | 1.36E-11 | 0.92925416 | 7.37E-10 |
| Ms4a4b | yellow | 4.33613292 | 1.52E-49 | 3.83525876 | 1.26E-38 |
| 1810011O10 | red | -0.9054883 | 1.03E-09 | -1.212255 | 2.04E-16 |
| Sftpb | red | -0.9650912 | 0.00194604 | -0.4902141 | 0.14613751 |
| Lig1 | orange | 1.93362534 | 1.12E-21 | 1.55994263 | 3.28E-14 |
| Adap1 | yellow | 2.26109177 | 2.18E-21 | 2.05139816 | 1.39E-17 |
| Hoxaas2 | red | -0.7376915 | 0.01470783 | -0.9429652 | 0.00228206 |
| Zbtb25 | orange | 0.70815969 | 0.00090397 | 0.50678329 | 0.02342727 |
| Med12l | orange | 0.95593391 | 0.00672326 | 1.52918386 | 1.82E-05 |
| Chn1 | red | -0.7174334 | 0.00018975 | -0.8292051 | 2.18E-05 |
| Mettl7a2 | skyblue | -2.5557775 | 0.00541145 | -1.7157881 | 0.06270857 |
| Cngb3 | black | 7.05226258 | 2.58E-07 | 3.86598959 | 1.80E-06 |
| Tmem154 | orange | 0.58261196 | 0.00280574 | 0.46621809 | 0.02163518 |
| Cebpb | orange | 1.21863035 | 4.20E-05 | 1.42999377 | 2.22E-06 |
| Ptafr | yellow | 2.7819547 | 3.92E-12 | 2.75566921 | 9.34E-12 |
| Ccdc18 | orange | 1.61617344 | 0.00661394 | 2.13917737 | 0.00057228 |
| Pign | orange | 0.89202673 | 4.18E-06 | 0.74817776 | 0.00018328 |

| | | | | | |
|---|---|---|---|---|---|
| Mpz | skyblue | -1.9749539 | 0.00096383 | -0.8733074 | 0.18194684 |
| Zfp658 | red | -1.1859219 | 2.51E-05 | -0.9453678 | 0.00102239 |
| Trnp1 | red | -0.775895 | 0.00417775 | -0.6578607 | 0.01942078 |
| Lrrc48 | skyblue | -1.9519317 | 2.76E-11 | -1.1116841 | 0.00026473 |
| Fry | red | -0.5076818 | 0.00038204 | -0.7241957 | 4.48E-07 |
| Ppp1r14b | orange | 0.56929912 | 0.00064997 | 0.7453543 | 1.05E-05 |
| Apol8 | black | 4.77574003 | 2.93E-06 | 4.32815184 | 5.12E-06 |
| Retsat | red | -0.6651878 | 0.00034213 | -0.4712159 | 0.01552295 |
| Ier5 | orange | 0.82450197 | 0.0004677 | 0.85823273 | 0.0003675 |
| Nbeal2 | orange | 0.88702325 | 4.72E-14 | 0.86432962 | 3.10E-13 |
| Capg | orange | 1.89963584 | 1.04E-10 | 1.67225904 | 2.51E-08 |
| Nfil3 | yellow | 3.04380501 | 3.66E-21 | 2.69468727 | 7.36E-17 |
| Dnah9 | skyblue | -2.6110799 | 6.89E-15 | -1.3689196 | 7.88E-05 |
| Hmga2 | yellow | 4.03456755 | 0.00897154 | 3.55464553 | 0.01933886 |
| Zfp663 | skyblue | -3.9090195 | 0.00010674 | -2.3677978 | 0.00715891 |
| Ttc26 | red | -1.2404398 | 1.02E-07 | -0.8465282 | 0.00047127 |
| Gulp1 | red | -1.2026897 | 8.08E-07 | -1.1885016 | 1.85E-06 |
| Glipr1 | orange | 1.82074791 | 2.90E-10 | 1.19256896 | 7.80E-05 |
| Immp2l | red | -1.0547361 | 0.00613438 | -0.7583777 | 0.06191501 |
| Usp13 | red | -1.4720208 | 7.98E-06 | -1.0390375 | 0.00256339 |
| Acbd4 | red | -0.8895817 | 1.76E-05 | -0.69342 | 0.00122317 |
| Tatdn2 | orange | 0.9211726 | 4.50E-09 | 0.90199475 | 1.54E-08 |
| Ces1d | skyblue | -1.9054646 | 4.86E-08 | -1.3420763 | 0.00021492 |
| Myh4 | skyblue | -2.0523573 | 0.00033411 | -0.6857838 | 0.28410633 |
| Gm7536 | orange | 0.60272504 | 0.00661479 | 0.42047752 | 0.08225667 |
| Skap1 | yellow | 2.81795119 | 1.79E-16 | 3.04001612 | 2.73E-18 |
| Slc35f3 | skyblue | -2.8944315 | 6.71E-07 | -1.4690907 | 0.00882692 |
| Fam47e | skyblue | -2.0686295 | 5.75E-12 | -1.4359846 | 3.13E-06 |
| Ces1g | skyblue | -2.9967491 | 1.39E-10 | -3.1883695 | 1.42E-11 |
| C030039L03I | red | -1.2083634 | 0.00085505 | -1.1410042 | 0.00213149 |
| Ebf1 | red | -0.8861635 | 7.18E-09 | -0.7069888 | 7.98E-06 |
| Cntrl | orange | 0.92444171 | 3.48E-11 | 1.01167824 | 5.30E-13 |
| Npm1 | orange | 0.93677111 | 1.76E-09 | 0.6486159 | 5.97E-05 |
| Gja5 | red | -1.2122782 | 1.46E-05 | -0.9956092 | 0.00058571 |
| Scimp | yellow | 4.17893633 | 3.67E-30 | 3.8839408 | 1.62E-25 |
| Tmem140 | orange | 1.09458153 | 4.71E-14 | 1.27061684 | 1.51E-18 |
| Trim12c | orange | 1.34521508 | 1.10E-37 | 1.38566989 | 1.52E-39 |
| Gm166 | red | -0.8336537 | 0.0002183 | -0.5513176 | 0.01815576 |
| Scn3a | skyblue | -2.4124754 | 5.88E-12 | -1.3273051 | 0.00028568 |
| AB124611 | yellow | 3.23846146 | 4.97E-30 | 3.12373297 | 1.17E-27 |
| Armc7 | orange | 1.25120961 | 3.88E-09 | 1.33056296 | 7.18E-10 |
| BC051142 | orange | 1.16563498 | 0.00038277 | 0.74595224 | 0.0233718 |
| Ccdc176 | skyblue | -1.7862888 | 3.16E-06 | -1.2947108 | 0.00102341 |

| | | | | | |
|---|---|---|---|---|---|
| St6galnac2 | red | -1.1227037 | 0.00039646 | -0.9662015 | 0.00313139 |
| Rpl38 | orange | 0.66123219 | 3.52E-06 | 0.6050389 | 3.64E-05 |
| Cep170 | orange | 0.61344733 | 6.64E-06 | 0.58889451 | 2.49E-05 |
| Chst3 | red | -1.122864 | 1.84E-05 | -0.6386365 | 0.02137784 |
| Apol9a | black | 5.26608437 | 1.06E-75 | 5.11790653 | 3.33E-74 |
| Uxs1 | orange | 0.62176852 | 0.0027478 | 0.52779896 | 0.01445085 |
| Birc2 | orange | 0.64573847 | 1.10E-08 | 0.40936694 | 0.00053239 |
| Ryr3 | skyblue | -1.3172185 | 0.00080978 | -1.2123613 | 0.00329615 |
| Zfp759 | red | -0.9475656 | 0.00135397 | -0.917721 | 0.00307281 |
| Whsc1 | orange | 1.42881031 | 8.09E-14 | 1.16699147 | 2.25E-09 |
| Zfp53 | orange | 0.61931254 | 0.00155527 | 0.50487122 | 0.01399719 |
| Mpp7 | red | -1.1116452 | 3.16E-10 | -0.6507029 | 0.00042637 |
| Phex | red | -1.178997 | 8.54E-07 | -1.1174087 | 6.84E-06 |
| Saa2 | black | 5.2057052 | 2.61E-05 | 3.90352845 | 1.07E-05 |
| Bloc1s2 | orange | 0.71666455 | 0.00038578 | 0.61201218 | 0.00349346 |
| Dtnbp1 | orange | 0.79364619 | 0.00047673 | 0.71104284 | 0.00245067 |
| Lgals8 | orange | 0.64915032 | 8.83E-07 | 0.66551066 | 7.50E-07 |
| Eif1a | orange | 0.9855768 | 2.32E-07 | 0.87140786 | 8.36E-06 |
| Trim30d | yellow | 3.07390053 | 7.24E-52 | 2.6828 | 7.47E-40 |
| Colq | skyblue | -3.3065538 | 3.21E-05 | -1.3891608 | 0.10737081 |
| Lce1a1 | red | -1.1541695 | 0.00021057 | -0.7846429 | 0.01663218 |
| Gnai1 | red | -0.6524575 | 4.23E-08 | -0.5786381 | 2.14E-06 |
| Gapdh | orange | 0.65723485 | 0.00012423 | 0.64493646 | 0.00024043 |
| Mex3b | red | -1.0303311 | 8.95E-06 | -0.8550328 | 0.00041205 |
| A830018L16R | skyblue | -3.1912603 | 1.81E-07 | -1.008242 | 0.07335411 |
| Tmem178b | skyblue | -2.4013683 | 0.0005509 | -1.7332996 | 0.01405519 |
| Sh3rf2 | skyblue | -1.4728744 | 1.46E-14 | -0.8393428 | 2.62E-05 |
| Lepr | red | -1.0678494 | 1.36E-05 | -0.9561989 | 0.00015057 |
| Prtn3 | orange | 0.42466421 | 0.39665542 | 1.3727244 | 0.00371368 |
| Megf6 | skyblue | -1.7167327 | 2.14E-08 | -1.3112737 | 3.43E-05 |
| Cyb5d2 | red | -0.6046347 | 0.00551045 | -0.8046685 | 0.0002946 |
| Ddx49 | orange | 0.64780016 | 2.50E-05 | 0.58948059 | 0.0001936 |
| Bak1 | orange | 1.89362321 | 8.99E-14 | 1.7436156 | 1.24E-11 |
| 1700007G11 | skyblue | -2.2380728 | 1.04E-12 | -1.6195318 | 1.45E-07 |
| Rpl32 | orange | 0.89797531 | 5.80E-12 | 0.71823885 | 8.04E-08 |
| Gm10032 | skyblue | -2.9036781 | 0.0015205 | -1.2661773 | 0.15066306 |
| Cacnb2 | skyblue | -1.343359 | 7.85E-05 | -1.2224576 | 0.00044919 |
| Gsta2 | skyblue | -1.6356046 | 0.00019289 | -0.8804325 | 0.05554745 |
| Unc13d | orange | 0.65749389 | 0.00019079 | 0.74837618 | 2.97E-05 |
| Fgf18 | red | -1.3976021 | 7.44E-05 | -0.6363309 | 0.0973092 |
| Sema3b | red | -0.9312172 | 1.23E-05 | -0.3783725 | 0.10273263 |
| Mdn1 | orange | 0.53690159 | 0.02264313 | 0.62358326 | 0.00921886 |
| Adtrp | red | -1.4485231 | 0.00300522 | -1.0119399 | 0.04996651 |

| | | | | | |
|---|---|---|---|---|---|
| Palld | red | -0.7577293 | 9.40E-06 | -0.6724586 | 0.00013399 |
| Mettl7a3 | skyblue | -2.2729888 | 0.00158401 | -1.8118484 | 0.01056405 |
| Eml1 | red | -1.2339013 | 1.04E-22 | -0.960127 | 5.66E-14 |
| Nfam1 | orange | 1.2424166 | 1.49E-05 | 1.24341479 | 2.27E-05 |
| Tpm3-rs7 | orange | 1.30831246 | 1.04E-07 | 1.04998674 | 5.29E-05 |
| Gstm1 | skyblue | -1.712035 | 1.59E-06 | -1.2537959 | 0.00071414 |
| Chsy3 | orange | 0.89105691 | 0.05285676 | 1.50472264 | 0.00093153 |
| T2 | skyblue | -2.1282291 | 2.88E-06 | -1.3888551 | 0.00153522 |
| Gm5431 | yellow | 4.02681504 | 2.08E-34 | 4.11678381 | 3.41E-35 |
| Gm5148 | skyblue | -1.5878096 | 0.00074678 | -1.187897 | 0.01169121 |
| Zfp846 | red | -0.8185226 | 0.00064111 | -0.6413154 | 0.0105918 |
| BC021614 | orange | 1.97399083 | 2.97E-14 | 1.66279567 | 1.07E-09 |
| Arhgap35 | red | -0.5803668 | 1.04E-05 | -0.3172462 | 0.02299474 |
| Tspan7 | red | -1.0833615 | 2.40E-11 | -0.9206765 | 2.78E-08 |
| Serpina9 | skyblue | -2.1465031 | 9.64E-06 | -1.8065185 | 0.00027618 |
| Espl1 | yellow | 2.7519599 | 4.24E-31 | 2.81801225 | 1.04E-31 |
| Spock2 | red | -0.6655329 | 0.00038935 | -0.2208099 | 0.29313145 |
| Mcm9 | orange | 0.79045572 | 1.58E-07 | 0.73582097 | 1.97E-06 |
| Rrp1b | orange | 0.68751353 | 3.38E-06 | 0.5037689 | 0.00105698 |
| Gpr182 | red | -0.6713607 | 0.00039351 | -0.4485799 | 0.02422335 |
| Zfp420 | red | -0.9510196 | 5.23E-05 | -0.618861 | 0.01125055 |
| Syt17 | skyblue | -1.9543803 | 1.81E-07 | -1.8369526 | 1.01E-06 |
| Cxcl2 | black | 6.8084163 | 1.70E-17 | 5.44733173 | 5.10E-25 |
| Gm8369 | yellow | 2.96750908 | 7.96E-27 | 2.69576296 | 8.14E-22 |
| Rnf207 | skyblue | -1.3982424 | 0.17110216 | -2.6871113 | 0.00737751 |
| Gpc6 | skyblue | -2.4717953 | 3.43E-09 | -1.2360933 | 0.00297323 |
| Gda | orange | 2.02190091 | 2.39E-10 | 1.91765676 | 3.26E-09 |
| Samd12 | skyblue | -1.792066 | 5.08E-20 | -1.7936017 | 8.39E-20 |
| Tubb2a | orange | 0.67653076 | 5.89E-05 | 0.71885061 | 2.88E-05 |
| Memo1 | orange | 0.68705083 | 3.06E-05 | 0.53630742 | 0.00176846 |
| Fcer1g | yellow | 3.87858795 | 2.81E-35 | 3.38526169 | 6.01E-27 |
| Lin9 | orange | 0.71768722 | 0.00246458 | 0.48494621 | 0.05250404 |
| Kcnt1 | black | 5.37634923 | 4.66E-21 | 3.75050544 | 8.93E-15 |
| Osm | yellow | 3.59207214 | 7.62E-07 | 2.80700845 | 0.00015011 |
| Thra | red | -0.8067878 | 6.17E-06 | -0.5056219 | 0.00694634 |
| Rnf169 | orange | 0.67762442 | 5.17E-06 | 0.52176728 | 0.00073045 |
| Hist1h1b | yellow | 2.12785542 | 0.06342852 | 4.74187068 | 0.0032624 |
| Nfe2 | orange | 0.65912658 | 0.0081526 | 0.59164084 | 0.0229209 |
| 0610039K10I | yellow | 3.50534238 | 1.28E-07 | 1.54968036 | 0.00404866 |
| Pirb | yellow | 3.28289702 | 4.27E-25 | 3.16703588 | 2.32E-23 |
| C1qtnf3 | skyblue | -3.2148915 | 3.10E-07 | -1.4442326 | 0.02181117 |
| Slc10a5 | skyblue | -3.3543606 | 8.60E-10 | -1.2192768 | 0.01505384 |
| Gm10052 | orange | 0.65401261 | 0.00823149 | 0.49225224 | 0.06187594 |

| | | | | | |
|---|---|---|---|---|---|
| Vwa8 | red | -0.6502763 | 1.03E-05 | -0.5949691 | 8.45E-05 |
| Hnrnpa3 | orange | 0.7729625 | 1.97E-06 | 0.6967237 | 2.98E-05 |
| Frem1 | red | -0.6293891 | 0.00270774 | -0.4810406 | 0.02943806 |
| Rpl18 | orange | 0.79724269 | 2.51E-06 | 0.55922243 | 0.00152349 |
| Fcgr4 | black | 6.16121938 | 1.24E-60 | 5.26579317 | 1.04E-45 |
| Ifitm6 | yellow | 2.26037381 | 1.36E-05 | 3.1189551 | 8.28E-09 |
| Ntrk3 | skyblue | -2.1453223 | 1.05E-11 | -1.0278415 | 0.00163456 |
| Mtfmt | orange | 0.6067103 | 1.27E-07 | 0.38060912 | 0.00156484 |
| Fam19a1 | skyblue | -2.3858803 | 8.10E-05 | -1.6789485 | 0.00685876 |
| Gm12715 | orange | 0.80903584 | 0.00955881 | 0.76396207 | 0.01769469 |
| Lep | skyblue | -1.8078355 | 0.04006734 | -2.6329559 | 0.00176231 |
| Sept9 | orange | 0.83111716 | 2.70E-05 | 0.62518954 | 0.00237671 |
| R74862 | red | -1.0604182 | 0.00962428 | -0.6401024 | 0.13502905 |
| Rpl11 | orange | 0.71825311 | 2.59E-08 | 0.48693387 | 0.00028871 |
| Tonsl | orange | 1.22564025 | 2.52E-09 | 1.11643668 | 1.26E-07 |
| Hopx | red | -0.7396052 | 0.00323696 | -0.4917737 | 0.06442714 |
| Csf2ra | yellow | 2.22858525 | 6.39E-11 | 2.14429617 | 5.59E-10 |
| Slc14a1 | orange | 1.59630252 | 0.0011563 | 1.58758698 | 0.00198253 |
| Fxn | orange | 0.52355659 | 0.01924618 | 0.74247572 | 0.00114513 |
| Mamld1 | red | -0.5910939 | 0.00224114 | -0.6150827 | 0.00163904 |
| Fxyd2 | skyblue | -2.1780728 | 0.00027146 | 0.71689553 | 0.22598838 |
| Actg2 | red | -1.059682 | 1.58E-06 | -0.0900532 | 0.74534483 |
| Bcas3 | red | -0.4503377 | 0.00417432 | -0.5825715 | 0.00025189 |
| Hadhb | red | -0.6094015 | 0.00013246 | -0.5905637 | 0.0003166 |
| Ptk2b | yellow | 2.1888967 | 4.41E-20 | 1.94251166 | 7.64E-16 |
| B3gnt8 | orange | 0.82693206 | 0.00751121 | 0.44615941 | 0.18464314 |
| Fcgr3 | yellow | 3.25014274 | 6.83E-21 | 2.69290152 | 1.60E-14 |
| Znhit1 | orange | 0.66015534 | 3.69E-05 | 0.21670667 | 0.22670195 |
| Ccdc154 | skyblue | -2.0853264 | 1.34E-11 | -2.0966891 | 2.17E-11 |
| Nsmce2 | orange | 0.64361287 | 1.04E-05 | 0.44557977 | 0.00343857 |
| Calcrl | red | -1.2284647 | 1.59E-07 | -1.0208317 | 2.32E-05 |
| Stfa2l1 | black | 5.55557938 | 5.57E-05 | 3.54578724 | 0.00043052 |
| Cdh24 | orange | 0.50802714 | 0.04076138 | 1.14244133 | 3.32E-06 |
| Zfp637 | red | -0.6440316 | 1.72E-06 | -0.3822457 | 0.00676611 |
| Myl3 | red | -1.2906703 | 0.00026905 | -0.8141222 | 0.02971329 |
| Slx1b | red | -0.7609134 | 1.45E-06 | -0.705804 | 1.39E-05 |
| Nrm | orange | 1.04719462 | 1.04E-09 | 0.84248955 | 2.03E-06 |
| Eif4a1 | orange | 0.92887419 | 7.84E-09 | 0.91326524 | 2.30E-08 |
| Rgs3 | red | -1.0119701 | 2.40E-22 | -0.9806206 | 6.81E-21 |
| Dbp | skyblue | -3.2953509 | 1.80E-17 | -2.4267468 | 9.62E-10 |
| Zfp874b | red | -0.9050095 | 7.65E-05 | -0.7360957 | 0.00167943 |
| Hydin | skyblue | -3.005779 | 2.68E-19 | -1.351143 | 6.82E-05 |
| Tnip2 | orange | 0.74971756 | 8.79E-08 | 0.60861253 | 2.48E-05 |

| | | | | | |
|---|---|---|---|---|---|
| Irak4 | orange | 0.84913377 | 1.18E-09 | 0.67538297 | 2.84E-06 |
| Ptp4a3 | red | -0.5828124 | 0.00192686 | -0.6267457 | 0.0010895 |
| Dsc3 | red | -0.8608399 | 0.09048628 | -1.3563409 | 0.00759392 |
| Adamts14 | orange | 1.73703726 | 6.94E-20 | 1.19133229 | 8.32E-10 |
| Grb2 | orange | 0.68167459 | 0.00015228 | 0.58837535 | 0.00154658 |
| Hspa2 | red | -0.8795763 | 8.67E-08 | -0.8136927 | 1.29E-06 |
| Chpt1 | red | -1.241715 | 2.32E-08 | -1.1268963 | 7.10E-07 |
| Alox5ap | orange | 1.84705118 | 1.04E-06 | 1.66824696 | 1.73E-05 |
| Psma3 | orange | 0.80836485 | 2.84E-08 | 0.63069876 | 2.75E-05 |
| Atp8b4 | yellow | 3.76334193 | 9.37E-28 | 2.69304265 | 1.08E-14 |
| Kif27 | skyblue | -2.1191592 | 1.63E-11 | -0.8609572 | 0.00844742 |
| Slc35e3 | red | -1.0937254 | 3.64E-10 | -0.8422738 | 2.65E-06 |
| Cxcl11 | black | 7.10711772 | 7.19E-36 | 11.0296358 | 5.08E-18 |
| Lrrc10 | skyblue | -2.4002192 | 0.0182309 | -3.6149181 | 0.00044798 |
| Arrb2 | yellow | 2.41707337 | 3.95E-18 | 2.03461128 | 6.22E-13 |
| Casc4 | red | -0.9331476 | 3.26E-07 | -0.7565162 | 6.11E-05 |
| Gtf2i | red | -0.9468754 | 4.62E-10 | -0.8381647 | 6.51E-08 |
| Gm1661 | skyblue | -2.2414032 | 5.81E-10 | -0.9227294 | 0.0078272 |
| Ppih | orange | 1.19784082 | 2.34E-08 | 1.05092683 | 2.11E-06 |
| Bckdha | red | -0.9651532 | 0.00179062 | -0.8212019 | 0.01030756 |
| Zfp128 | red | -0.597235 | 0.06099452 | -0.8244551 | 0.00939572 |
| Chst8 | skyblue | -2.4264358 | 2.47E-23 | -1.322739 | 6.25E-08 |
| Sntb1 | red | -0.9527311 | 0.00013493 | -0.6952805 | 0.00749588 |
| Trim5 | orange | 1.00404593 | 5.82E-08 | 0.88787409 | 3.56E-06 |
| Adgrg3 | orange | 1.11244196 | 3.73E-06 | 1.10495568 | 6.41E-06 |
| Irak2 | orange | 0.70146965 | 2.76E-05 | 0.76259917 | 7.57E-06 |
| Xcr1 | orange | 0.98746712 | 0.02001931 | 1.19352445 | 0.00765841 |
| Zfp266 | red | -0.844718 | 5.46E-06 | -0.6241302 | 0.0012289 |
| 0610040J01F | red | -0.6685806 | 0.00157199 | -0.0840337 | 0.74803037 |
| Tor3a | orange | 1.10527259 | 2.75E-19 | 0.88414059 | 1.57E-12 |
| Tmem219 | orange | 1.11521699 | 2.52E-07 | 0.90326984 | 5.38E-05 |
| Tnfrsf19 | red | -1.3189935 | 8.23E-16 | -1.1370785 | 8.60E-12 |
| H2-Q9 | yellow | 2.33152836 | 4.63E-35 | 2.02443009 | 1.51E-26 |
| Mfap2 | red | -0.6644024 | 0.00381938 | -0.5913049 | 0.0127308 |
| H2-Eb1 | orange | 1.4655747 | 2.90E-21 | 1.11223461 | 1.92E-12 |
| Ifitm2 | orange | 0.79222983 | 0.00123382 | 0.48853352 | 0.06102192 |
| Layn | red | -1.0187379 | 4.85E-08 | -1.0581655 | 2.85E-08 |
| Eno3 | skyblue | -1.5269684 | 2.50E-07 | -1.1606054 | 0.00015395 |
| Nkpd1 | red | -1.0572154 | 0.00769026 | -0.9199095 | 0.02508251 |
| Hist1h4i | orange | 1.9921715 | 2.39E-08 | 1.44829209 | 0.00010875 |
| Atp8b2 | orange | 0.74055636 | 2.60E-08 | 0.70343896 | 2.16E-07 |
| Pla2g16 | orange | 0.7962025 | 6.82E-05 | 0.55322732 | 0.0081782 |
| Slc9a6 | red | -0.8319887 | 2.21E-11 | -0.8353892 | 3.31E-11 |

| | | | | | |
|---|---|---|---|---|---|
| Plekhh1 | red | -0.9909489 | 2.98E-05 | -0.7436884 | 0.00236997 |
| Nsa2 | orange | 0.85379949 | 4.64E-07 | 0.58044999 | 0.00104115 |
| Tsga10 | red | -1.0570985 | 1.24E-05 | -1.2197184 | 6.47E-07 |
| Gmfg | orange | 1.24048129 | 2.14E-10 | 1.16806988 | 4.60E-09 |
| Intu | red | -1.0919233 | 0.0008979 | -0.1887283 | 0.59594054 |
| B2m | yellow | 2.1224356 | 6.03E-35 | 1.8171023 | 9.72E-26 |
| Ube2s | orange | 0.79695435 | 2.66E-09 | 0.836449 | 7.87E-10 |
| Kcnd2 | skyblue | -2.4724546 | 0.00754156 | -0.8610173 | 0.38931766 |
| Trim55 | skyblue | -1.5651085 | 0.00030947 | -2.024154 | 6.45E-06 |
| Acyp2 | red | -0.6361981 | 0.00761572 | -0.3212668 | 0.21942719 |
| Trmt61a | orange | 1.21863175 | 4.23E-05 | 0.99470456 | 0.00127863 |
| Slc4a4 | red | -1.2134127 | 6.56E-10 | -0.6098157 | 0.00300698 |
| Hist1h4h | skyblue | -1.6391151 | 0.00050181 | -1.3957316 | 0.00516824 |
| Rrs1 | orange | 0.77294515 | 0.00030947 | 0.84776326 | 0.00010854 |
| Olfr920 | orange | 1.4349365 | 1.75E-06 | 0.84712013 | 0.0050985 |
| Cdh3 | orange | 0.31101385 | 0.29405691 | 0.91343186 | 0.00166143 |
| Mcpt4 | skyblue | -3.0148682 | 1.04E-08 | -1.6245898 | 0.00077515 |
| Lsamp | skyblue | -1.9836504 | 2.36E-06 | -1.7207087 | 6.27E-05 |
| Retnla | yellow | 3.71497589 | 1.76E-47 | 3.98465882 | 1.85E-54 |
| Prpf40a | orange | 0.71963739 | 5.33E-09 | 0.40319216 | 0.0019182 |
| Slc38a11 | skyblue | -1.1376972 | 0.08431463 | -1.9397469 | 0.00368655 |
| Fnip2 | orange | 0.82676088 | 0.0001149 | 0.85506688 | 9.37E-05 |
| Stk19 | orange | 0.94589731 | 1.38E-19 | 0.85670879 | 4.76E-16 |
| H2-K1 | yellow | 2.15300382 | 1.73E-39 | 1.86684298 | 8.48E-30 |
| Mmgt1 | red | -0.6150325 | 2.53E-07 | -0.6591831 | 5.90E-08 |
| Dnaic1 | skyblue | -2.8129908 | 6.71E-14 | -1.7798085 | 2.71E-06 |
| Hipk2 | orange | 0.96506996 | 6.42E-09 | 0.91963267 | 6.22E-08 |
| Nol10 | orange | 0.93949018 | 2.78E-06 | 0.83670696 | 5.37E-05 |
| Obscn | red | -0.7182117 | 0.00280628 | -0.9099217 | 0.00018792 |
| Snrpa | orange | 0.58664606 | 5.65E-05 | 0.45070869 | 0.0029951 |
| Krt5 | red | -1.0441023 | 0.00031442 | -1.093508 | 0.0002239 |
| C1qtnf7 | skyblue | -1.606313 | 9.04E-16 | -1.300669 | 1.59E-10 |
| Orm2 | yellow | 3.18420178 | 0.00036803 | 2.4833366 | 0.00365429 |
| Zfp229 | red | -0.7776462 | 4.19E-05 | -0.7527039 | 9.34E-05 |
| Dpp6 | skyblue | -1.974505 | 0.0002884 | -2.3757932 | 9.91E-06 |
| Adgrg5 | yellow | 3.05144222 | 4.43E-21 | 2.58819012 | 2.38E-15 |
| Dot1l | orange | 0.70365415 | 9.51E-05 | 0.74201497 | 5.45E-05 |
| Pclo | skyblue | -2.524526 | 7.89E-09 | -1.2569626 | 0.00340429 |
| Akap6 | skyblue | -1.8796335 | 1.95E-05 | -1.6928186 | 0.00019847 |
| U2af1 | orange | 0.74172494 | 3.53E-12 | 0.57091628 | 2.04E-07 |
| Gdpd1 | red | -0.8728976 | 0.00012908 | -0.7881253 | 0.00076295 |
| Ppp1r1b | red | -1.1672898 | 0.00255177 | -0.6918277 | 0.09288096 |
| Tnnt3 | red | -1.3594241 | 0.00019508 | -0.8899226 | 0.02049839 |

| | | | | | |
|---|---|---|---|---|---|
| Cyp2d22 | red | -0.9052559 | 0.00176157 | -0.7807072 | 0.00905827 |
| Kalrn | orange | 0.36457296 | 0.18665087 | 0.82323034 | 0.00198628 |
| Cfd | skyblue | -2.6495823 | 0.00415584 | -0.9448003 | 0.36162571 |
| Armc4 | skyblue | -2.1136939 | 5.93E-10 | -1.178448 | 0.00053272 |
| Myl1 | skyblue | -2.1153784 | 0.00039439 | -1.2400853 | 0.05107575 |
| Inadl | red | -0.8890769 | 0.00142225 | -0.5525724 | 0.06177605 |
| Sphk1 | orange | 1.0242907 | 5.29E-09 | 0.74407747 | 4.14E-05 |
| Ccdc62 | red | -0.5005652 | 0.1136592 | -1.1185277 | 0.00027003 |
| Ces1e | skyblue | -2.6495752 | 0.00090125 | -1.5832877 | 0.06086648 |
| Olfr99 | orange | 1.766216 | 0.00447047 | 1.87364525 | 0.00615406 |
| Rpl34 | orange | 0.59226243 | 1.68E-06 | 0.43801415 | 0.00065804 |
| Hsh2d | yellow | 3.2966961 | 4.25E-27 | 2.94639149 | 7.28E-22 |
| Zfp13 | red | -0.6402083 | 4.68E-05 | -0.3931325 | 0.01625994 |
| Lmtk3 | orange | 1.48951768 | 0.00046289 | 1.7965186 | 3.98E-05 |
| Pgk1 | orange | 0.78026689 | 1.52E-06 | 0.61744472 | 0.00023764 |
| Lmnb2 | orange | 0.52335231 | 0.00031915 | 0.60525945 | 4.40E-05 |
| Trim54 | red | -1.1128556 | 8.13E-05 | -1.0597926 | 0.00028881 |
| Cd200r4 | yellow | 2.50278422 | 1.01E-07 | 1.82892187 | 0.00017085 |
| Btbd3 | red | -1.3216074 | 8.55E-07 | -0.9852926 | 0.00039429 |
| Ear6 | black | 5.42330653 | 0.00758429 | 4.03222902 | 0.06130152 |
| Hs6st2 | skyblue | -2.4677368 | 5.48E-17 | -1.7829975 | 8.10E-11 |
| Erbb4 | skyblue | -0.6721033 | 0.46629474 | -2.389736 | 0.00723924 |
| Tnfaip8 | orange | 0.89770828 | 5.73E-07 | 0.77763381 | 2.61E-05 |
| Cks2 | yellow | 2.23780546 | 4.53E-10 | 2.36917153 | 1.16E-10 |
| Opcml | skyblue | -2.15113 | 4.91E-05 | -2.0644915 | 0.00021455 |
| Pvrl2 | orange | 0.57155321 | 0.00206871 | 0.58156401 | 0.00220992 |
| Rpp25 | skyblue | -0.8694746 | 0.18135773 | -1.9922202 | 0.00265786 |
| Gm10115 | skyblue | -0.5907572 | 0.4747596 | -2.6935956 | 0.00231307 |
| Rpl17 | orange | 0.59772429 | 2.89E-06 | 0.38048879 | 0.00448878 |
| Cytl1 | skyblue | -1.64198 | 0.00011209 | -1.8019537 | 3.24E-05 |
| Itgb1bp1 | orange | 0.60400322 | 1.79E-05 | 0.22827892 | 0.13943543 |
| Tubb3 | yellow | 1.80632927 | 1.29E-16 | 2.03364366 | 7.49E-20 |
| Arf2 | orange | 0.62331925 | 0.00128506 | 0.56523634 | 0.00467293 |
| Fbxl12os | red | -1.5092706 | 0.00084952 | -0.6265255 | 0.20941628 |
| Ifit3b | yellow | 3.14871876 | 3.68E-70 | 3.21082101 | 1.06E-72 |
| Nsl1 | yellow | 2.00328061 | 2.16E-09 | 2.10182914 | 1.33E-09 |
| Fabp4 | skyblue | -2.3508309 | 0.00133051 | -1.4641832 | 0.05984284 |
| Ncr1 | orange | 0.8012035 | 0.03446374 | 1.83367291 | 1.57E-06 |
| Cys1 | skyblue | -2.2261053 | 2.48E-09 | -1.8270628 | 1.85E-06 |
| Tubb4a | skyblue | -1.6006588 | 6.88E-10 | -0.9519683 | 0.00043241 |
| Rpl7a | orange | 0.70342858 | 4.81E-07 | 0.59181259 | 3.91E-05 |
| Ncs1 | red | -0.9399448 | 3.79E-05 | -0.6219852 | 0.00904897 |
| Cav3 | red | -0.9507723 | 0.0005573 | -0.1564018 | 0.64140601 |

| | | | | | |
|---|---|---|---|---|---|
| Sim2 | red | -1.0080941 | 0.00061303 | -0.5289054 | 0.09623452 |
| AI413582 | orange | 1.02928103 | 3.55E-11 | 0.93519847 | 5.82E-09 |
| 1810041L15F | skyblue | -2.1218204 | 1.48E-07 | -1.2290095 | 0.00314116 |
| Chia1 | red | -0.8520013 | 0.01022134 | -0.9598653 | 0.00437002 |
| Csprs | black | 5.83772344 | 0.00018032 | 1.69304502 | 0.12732344 |
| Kcnc3 | red | -0.934094 | 0.00054948 | -0.4714874 | 0.10608427 |
| Zfp599 | orange | 0.72873798 | 0.07412258 | 1.12061976 | 0.00696018 |
| Actg1 | orange | 0.8973618 | 5.45E-05 | 0.91871893 | 5.23E-05 |
| Tcp11 | skyblue | -1.5969726 | 1.28E-14 | -1.079383 | 2.53E-07 |
| Phactr2 | red | -0.989843 | 7.12E-08 | -0.7123866 | 0.00018528 |
| Rpl31-ps11 | orange | 1.0529478 | 0.00713124 | -0.0234819 | 0.96273917 |
| Klhl24 | red | -1.2067429 | 5.07E-05 | -0.9932202 | 0.00125425 |
| Hdac10 | red | -0.8229181 | 1.74E-07 | -0.5002108 | 0.00237678 |
| Acadm | red | -1.1645201 | 9.68E-08 | -1.026644 | 4.61E-06 |
| Stat4 | yellow | 3.43732107 | 1.40E-38 | 3.39636002 | 6.91E-34 |
| Kdr | red | -1.4890549 | 1.87E-19 | -1.0612685 | 3.74E-10 |
| Cped1 | skyblue | -1.9188074 | 4.10E-07 | -1.299831 | 0.00098183 |
| Rpl35 | orange | 0.77957903 | 2.64E-05 | 0.47402192 | 0.01538985 |
| Msln | orange | 1.33648183 | 0.00011148 | 1.25465932 | 0.0004156 |
| Sox7 | skyblue | -1.5808535 | 3.98E-45 | -1.4050793 | 9.68E-36 |
| Mapk3 | red | -0.6574415 | 3.85E-12 | -0.6629054 | 4.09E-12 |
| Calml3 | red | -0.9618875 | 0.00225869 | -0.7918767 | 0.01562405 |
| Kcnma1 | red | -1.4146659 | 0.00097913 | -0.8780568 | 0.04485596 |
| Clip2 | orange | 0.64540521 | 0.00386635 | 0.61760071 | 0.00717486 |
| Hspb11 | red | -0.7707903 | 2.05E-05 | -0.1385919 | 0.51696141 |
| Cd300lb | orange | 1.18072337 | 0.00053085 | 0.97083338 | 0.00593783 |
| Ldha | orange | 0.70675315 | 2.75E-05 | 0.57482699 | 0.00097969 |
| Gpr84 | black | 8.31360228 | 2.63E-09 | 7.08710103 | 7.67E-07 |
| Grm4 | skyblue | -2.8968337 | 5.72E-05 | -1.9766093 | 0.00298476 |
| Zfp993 | yellow | 2.35530788 | 0.00022186 | 1.56616192 | 0.02049127 |
| Parp10 | yellow | 2.30860815 | 9.79E-106 | 2.24382142 | 2.88E-100 |
| Naa15 | orange | 0.74292991 | 6.26E-07 | 0.43254762 | 0.00598871 |
| Gm8995 | yellow | 2.81665576 | 1.75E-65 | 2.68427341 | 3.91E-59 |
| Tmem117 | red | -1.1600966 | 0.00205049 | -0.4754297 | 0.24777384 |
| Rpl27 | orange | 0.92032193 | 2.16E-09 | 0.71517665 | 6.40E-06 |
| Slc39a4 | red | -0.6834066 | 2.66E-05 | -0.4627631 | 0.00647964 |
| BC023105 | black | 5.0951919 | 7.04E-10 | 5.19892408 | 4.50E-10 |
| Tmed5 | orange | 0.77087941 | 2.08E-06 | 0.49612441 | 0.00362599 |
| Lrrc43 | skyblue | -1.7097027 | 1.28E-06 | -0.8017605 | 0.02912478 |
| Gm10131 | orange | 2.06371598 | 0.00944699 | 1.12031896 | 0.18286968 |
| Ddo | skyblue | -2.0629745 | 1.41E-09 | -1.6112602 | 3.96E-06 |
| Wscd2 | red | -0.7963098 | 1.69E-05 | -0.1720354 | 0.40885664 |
| B9d2 | orange | 0.63182161 | 0.00023254 | 0.63283082 | 0.00034625 |

| | | | | | |
|---|---|---|---|---|---|
| Nmral1 | orange | 1.67589102 | 1.06E-10 | 1.89633216 | 8.61E-13 |
| Syne2 | red | -1.160494 | 8.54E-08 | -0.8803331 | 8.54E-05 |
| D630045J12l | red | -1.084945 | 1.83E-05 | -0.9591973 | 0.00021758 |
| Zkscan7 | red | -0.9309931 | 0.00049695 | -0.8063474 | 0.00335989 |
| Eno1 | orange | 1.09222418 | 2.26E-07 | 0.82901375 | 0.00014713 |
| Stmnd1 | skyblue | -3.244975 | 5.20E-20 | -1.9992514 | 8.19E-09 |
| Sema3e | skyblue | -1.4936342 | 9.15E-08 | -0.8198236 | 0.0053692 |
| Aox1 | red | -1.1641473 | 8.46E-07 | -0.6394246 | 0.01035898 |
| C230062I16F | red | -1.3531536 | 8.05E-05 | -1.0763785 | 0.00278912 |
| Unc5d | skyblue | -1.2274542 | 0.06215305 | -2.5329808 | 0.00051003 |
| Sox11 | skyblue | -2.3920218 | 3.56E-10 | -1.4004416 | 0.00043611 |
| Jakmip1 | orange | 1.58546378 | 1.65E-07 | 2.29658516 | 6.67E-14 |
| Brwd3 | orange | 0.70619552 | 9.99E-05 | 0.34943859 | 0.07045369 |
| Pcdhb5 | red | -1.1436867 | 0.00481451 | -0.0777708 | 0.87967941 |
| Sfxn4 | red | -0.9798125 | 4.58E-06 | -1.2513822 | 9.29E-09 |
| Mapk15 | red | -0.829145 | 4.19E-06 | -0.4961651 | 0.00840888 |
| Chadl | red | -1.1481797 | 0.00053076 | -1.1366001 | 0.00091609 |
| Chil4 | yellow | 1.75219603 | 0.03206387 | 3.24957934 | 2.01E-05 |
| Utp14a | orange | 0.78915235 | 3.65E-07 | 0.62751384 | 0.00011277 |
| Prpf38a | orange | 0.8187525 | 1.08E-06 | 0.57803084 | 0.00096427 |
| Dupd1 | skyblue | -3.7126917 | 0.00022996 | -3.2701008 | 0.0046715 |
| Slc24a3 | red | -1.381711 | 9.95E-11 | -0.8096481 | 0.00026314 |
| Crem | orange | 0.88036604 | 2.79E-07 | 0.91144817 | 1.94E-07 |
| Dhrsx | orange | 0.72826419 | 2.51E-05 | 0.80991061 | 4.33E-06 |
| Dpp3 | orange | 0.65435471 | 3.43E-07 | 0.61258851 | 3.17E-06 |
| Slco1a5 | skyblue | -1.7947794 | 8.37E-05 | -0.6679244 | 0.15031901 |
| Wibg | orange | 1.11447795 | 8.15E-14 | 0.96056815 | 3.21E-10 |
| Trerf1 | orange | 0.60753248 | 0.00499014 | 0.27080035 | 0.25456719 |
| Scgb3a1 | skyblue | -3.4577809 | 5.12E-09 | -2.2256549 | 0.00030219 |
| Ipcef1 | yellow | 2.91352764 | 5.43E-26 | 2.96638922 | 2.21E-26 |
| Fbln2 | red | -1.398191 | 1.50E-19 | -0.7845837 | 1.04E-06 |
| Vrk2 | orange | 1.08546954 | 2.93E-05 | 0.84357502 | 0.00189268 |
| Cnnm2 | red | -0.7797587 | 1.14E-07 | -0.4468919 | 0.00378968 |
| Hcst | yellow | 2.37765577 | 2.28E-15 | 2.24143069 | 2.59E-13 |
| Cadm2 | skyblue | -1.8625537 | 6.29E-05 | -1.6977097 | 0.00027366 |
| Prr36 | red | -1.0855809 | 2.19E-05 | -1.0595913 | 3.18E-05 |
| Cenpj | orange | 0.47406458 | 0.00313996 | 0.63439425 | 9.74E-05 |
| Rab44 | red | -1.1141118 | 0.00021831 | -0.4816693 | 0.13871355 |
| Gm4735 | orange | 1.44601168 | 0.00792178 | 1.20670746 | 0.03908509 |
| 4430402I18R | skyblue | -1.4280587 | 3.84E-07 | -1.4140654 | 5.34E-07 |
| Ifi27 | red | -0.9795739 | 0.00013064 | -0.522375 | 0.05663301 |
| Paqr9 | skyblue | -1.8464539 | 0.00145322 | -2.5140748 | 2.05E-05 |
| Hvcn1 | orange | 1.37439349 | 6.67E-05 | 0.885608 | 0.01516447 |

| | | | | | |
|---|---|---|---|---|---|
| Ccdc146 | skyblue | -1.7694037 | 1.71E-12 | -1.0399189 | 3.61E-05 |
| Tank | orange | 0.84575111 | 1.62E-10 | 0.71212621 | 1.39E-07 |
| Cntn4 | skyblue | -2.2548833 | 0.00394647 | -1.678622 | 0.04059136 |
| Aox3 | skyblue | -2.4081503 | 0.00056119 | -1.152215 | 0.12806409 |
| Lrrc51 | skyblue | -1.6623321 | 1.20E-13 | -0.9690996 | 1.85E-05 |
| Hhip | red | -1.1401287 | 4.06E-12 | -0.9893778 | 3.73E-09 |
| mt-Rnr1 | red | -0.6581435 | 0.00978093 | -0.4592345 | 0.08833306 |
| mt-Rnr2 | red | -0.8078053 | 7.68E-05 | -0.6599851 | 0.00179358 |
| ND1 | red | -0.6609651 | 0.00036811 | -0.7661511 | 4.80E-05 |
| ND2 | skyblue | -1.388137 | 5.86E-08 | -1.2832541 | 9.29E-07 |
| COX1 | red | -0.6635491 | 2.71E-07 | -0.6247223 | 2.15E-06 |
| ND4 | red | -1.050643 | 7.78E-07 | -1.1354079 | 1.39E-07 |
| ND5 | red | -0.9771964 | 3.42E-06 | -0.8089293 | 0.00019577 |
| ND6 | red | -0.9342604 | 1.94E-05 | -0.9778291 | 1.17E-05 |
| CYTB | red | -1.1281528 | 5.66E-09 | -1.1315972 | 8.16E-09 |
| Sepp1 | red | -0.9169387 | 0.00053149 | -0.8035904 | 0.00326331 |
| Gm25131 | yellow | 2.66498475 | 0.00190625 | 0.91331691 | 0.42836019 |
| Snord104 | orange | 1.43361973 | 4.04E-05 | 1.01544964 | 0.00551077 |
| Snora31 | orange | 1.19718637 | 0.00134504 | 1.26838033 | 0.00103986 |
| Mir142b | yellow | 1.61180972 | 0.0003014 | 2.8203024 | 2.44E-07 |
| Gm24494 | orange | 0.82009113 | 0.08907373 | 2.18726173 | 1.65E-05 |
| Cd209b | skyblue | -3.8248569 | 0.00042959 | -3.820855 | 0.0007162 |
| 2610305D13 | red | -0.9183084 | 0.0066034 | -0.8259088 | 0.01757659 |
| Dhrs3 | red | -0.8676962 | 1.96E-07 | -0.6253887 | 0.00030916 |
| Gm1976 | orange | 1.13101053 | 0.00475999 | 0.98106421 | 0.01625415 |
| Muc5b | skyblue | -2.7380102 | 1.60E-06 | -0.7442486 | 0.24552592 |
| Adamtsl1 | red | -0.9431039 | 5.01E-05 | -0.4331849 | 0.08445006 |
| Kndc1 | skyblue | -2.3202389 | 4.26E-11 | -1.1947443 | 0.00102849 |
| Slc31a2 | orange | 0.97388311 | 1.94E-06 | 0.73550975 | 0.00053751 |
| E230001N04 | orange | 1.71812903 | 2.14E-06 | 0.74304977 | 0.05803515 |
| Anks6 | red | -0.6824058 | 0.00027028 | -0.2847799 | 0.15831998 |
| Spag8 | skyblue | -1.5776017 | 7.92E-08 | -0.788206 | 0.00717665 |
| Pomgnt2 | skyblue | -1.5630891 | 4.82E-10 | -1.255951 | 1.08E-06 |
| Gm10156 | yellow | 3.11861008 | 1.94E-05 | 2.02019148 | 0.00102887 |
| Trim12a | orange | 1.82214775 | 9.02E-94 | 1.74230096 | 6.74E-85 |
| Vps37b | orange | 0.63550924 | 0.00164355 | 0.67770986 | 0.00102768 |
| Rtp3 | red | -1.0409199 | 2.78E-06 | -1.1572891 | 4.24E-07 |
| Wdr6 | red | -0.8328715 | 2.02E-10 | -0.4821783 | 0.00043744 |
| Serpina3c | orange | 1.30468236 | 0.00150535 | 1.32630097 | 0.00153604 |
| Serpina3f | black | 6.53564448 | 9.32E-115 | 5.19437234 | 6.03E-79 |
| Plekhd1 | skyblue | -1.6206241 | 1.57E-06 | -1.0290245 | 0.00351064 |
| Mthfs | orange | 1.09742262 | 0.00178627 | 1.00833379 | 0.0048087 |
| Gm6254 | orange | 0.76151535 | 0.00929266 | 0.74491654 | 0.01765359 |

| | | | | | |
|---|---|---|---|---|---|
| Gm6969 | skyblue | -1.6886491 | 0.0121681 | -2.0867233 | 0.00243907 |
| Gm10167 | orange | 0.99711537 | 0.000475 | 1.01302301 | 0.00055877 |
| 6030419C18 | red | -0.7236471 | 0.00621779 | -0.3923091 | 0.16709286 |
| Wdr35 | red | -1.092698 | 2.66E-29 | -0.7726387 | 2.22E-15 |
| Gm5113 | red | -0.7957708 | 0.0008976 | -0.8128475 | 0.0009567 |
| Pilrb2 | orange | 1.45058354 | 0.0004201 | 1.77782582 | 3.06E-05 |
| Pilrb1 | orange | 0.83317555 | 0.00149919 | 1.82588667 | 1.93E-11 |
| Tnfsf18 | yellow | 1.58990455 | 0.24261809 | 4.17068699 | 0.00374167 |
| Zfp772 | red | -0.7051395 | 1.67E-05 | -0.2172549 | 0.24673888 |
| Hmcn1 | skyblue | -1.6366846 | 2.25E-23 | -1.4336488 | 5.18E-18 |
| Oas1g | black | 5.42017044 | 5.61E-84 | 5.06789564 | 1.37E-75 |
| Vsig10 | red | -0.6271522 | 1.86E-12 | -0.2958521 | 0.00172769 |
| Myo1h | red | -1.2373383 | 0.00018749 | -0.6179015 | 0.07085237 |
| Cryba4 | yellow | 2.29571581 | 0.00187032 | 2.68615228 | 0.00085121 |
| Serpinb5 | red | -0.6339476 | 0.08286202 | -1.008117 | 0.00598779 |
| Gm3608 | orange | 1.67738036 | 0.0001978 | 0.3368399 | 0.51810875 |
| Unc93a | yellow | 4.3746277 | 5.84E-05 | 3.12021429 | 0.00167768 |
| H2-K2 | orange | 0.63773702 | 1.13E-06 | 0.58452563 | 1.43E-05 |
| H2-T23 | yellow | 2.64742812 | 5.35E-73 | 2.36382614 | 3.18E-58 |
| Rplp0 | orange | 0.71162934 | 3.93E-11 | 0.57254522 | 2.23E-07 |
| Ppp1r3c | red | -0.590035 | 0.01486382 | -0.677455 | 0.00620016 |
| Ifit1bl2 | orange | 0.82449247 | 0.00929136 | 0.59789643 | 0.07367882 |
| Bmpr2 | red | -0.818828 | 5.60E-05 | -0.7495426 | 0.00033547 |
| Tmem128 | orange | 0.93579302 | 2.11E-16 | 0.71160297 | 1.22E-09 |
| Lyar | orange | 0.88360208 | 1.41E-06 | 0.67564353 | 0.00039464 |
| B3galt4 | orange | 0.68236938 | 0.00096788 | 0.32455623 | 0.14508816 |
| Tspan6 | red | -0.8037269 | 7.08E-05 | -0.5668866 | 0.00714566 |
| Zfp763 | red | -0.7906528 | 0.00236213 | -0.8347582 | 0.00170485 |
| A430093F15I | yellow | 4.15794811 | 1.68E-30 | 4.09339123 | 7.46E-29 |
| Klra3 | yellow | 1.87784841 | 0.00323834 | 2.41204795 | 0.00045547 |
| Klra7 | yellow | 2.03161268 | 9.94E-10 | 3.74270896 | 2.44E-18 |
| Klk11 | red | -0.8775615 | 0.0059854 | -0.4342166 | 0.21281835 |
| Syngap1 | red | -1.0836189 | 3.59E-10 | -0.6869625 | 0.00012057 |
| Lpar5 | orange | 1.19694733 | 2.54E-05 | 0.37266865 | 0.24854838 |
| Gm10222 | red | -0.6049433 | 0.23238353 | -1.674002 | 0.00114699 |
| Khdc1a | yellow | 2.07883829 | 0.05147081 | 3.94968486 | 0.0021429 |
| Xlr4b | yellow | 2.48396555 | 1.83E-10 | 2.09111401 | 2.13E-07 |
| Myl9 | red | -1.1289396 | 1.14E-10 | -0.7812586 | 1.66E-05 |
| Pex26 | red | -0.8519893 | 8.27E-07 | -0.6255473 | 0.00050277 |
| Ccdc87 | red | -1.1537737 | 0.00414391 | -0.3927027 | 0.3495062 |
| Gm13139 | yellow | 2.0563656 | 4.98E-07 | 2.1598976 | 1.23E-06 |
| Zfp760 | red | -1.1376048 | 5.14E-08 | -0.879218 | 4.28E-05 |
| Gtf2f2 | orange | 1.24289938 | 1.11E-07 | 0.81417744 | 0.00085223 |

| | | | | | |
|---|---|---|---|---|---|
| Tcf15 | red | -1.3763241 | 0.00431559 | -0.4487922 | 0.43203311 |
| 1500009C09 | yellow | 2.79694651 | 0.01654294 | 4.4424631 | 0.00035175 |
| Cenpm | yellow | 3.14974746 | 1.18E-12 | 3.3398767 | 7.30E-13 |
| Cst7 | yellow | 4.11234139 | 2.30E-54 | 3.56329393 | 1.81E-43 |
| Col8a1 | red | -1.0922651 | 1.23E-06 | -1.2433082 | 7.96E-08 |
| Macrod2 | red | -0.6433251 | 0.01333153 | -1.0540209 | 9.06E-05 |
| Lgals1 | orange | 1.83254533 | 3.68E-09 | 1.48873337 | 3.15E-06 |
| Il2rb | yellow | 4.47747276 | 1.27E-87 | 4.21464468 | 2.18E-77 |
| Gm11808 | orange | 0.75275385 | 0.00390314 | 0.25427464 | 0.39400475 |
| Phf11d | yellow | 2.9876 | 1.35E-37 | 2.95823042 | 9.19E-37 |
| Apol9b | black | 4.60960857 | 2.66E-41 | 4.42815731 | 2.20E-38 |
| Efcc1 | skyblue | -2.838174 | 1.40E-17 | -1.9934188 | 3.16E-09 |
| Shroom4 | red | -0.853059 | 5.74E-07 | -0.4606494 | 0.01068352 |
| Slc4a5 | skyblue | -2.5596867 | 1.06E-05 | -1.4623533 | 0.01699715 |
| Aup1 | orange | 0.60608866 | 5.13E-07 | 0.36418689 | 0.00400079 |
| Dok1 | orange | 1.47250799 | 3.50E-13 | 1.2938359 | 3.54E-10 |
| Cep152 | orange | 0.91124692 | 1.64E-09 | 0.82898869 | 8.88E-08 |
| Pnp2 | yellow | 2.54135158 | 3.55E-13 | 2.00657954 | 2.52E-08 |
| Gmnc | skyblue | -1.6405477 | 0.00193446 | -0.8150822 | 0.12429299 |
| Duox2 | skyblue | -2.2724338 | 0.00011233 | -0.8829114 | 0.15385019 |
| Aard | skyblue | -2.4876422 | 1.94E-11 | -1.9133186 | 4.64E-07 |
| Zfp467 | red | -0.6756218 | 9.87E-07 | -0.5850183 | 3.51E-05 |
| Gm4841 | black | 6.35044088 | 5.92E-62 | 7.38296937 | 4.87E-66 |
| Actc1 | skyblue | -2.4104755 | 5.10E-06 | -1.8594751 | 0.00068934 |
| Efcab1 | skyblue | -2.3454481 | 2.44E-12 | -1.9764794 | 3.89E-09 |
| Gpr88 | orange | 1.08061286 | 0.008897 | 1.92525327 | 4.20E-06 |
| Flnc | red | -1.2949491 | 7.16E-07 | -0.5465154 | 0.0528145 |
| Trp53i11 | orange | 1.55828074 | 1.61E-12 | 1.82998435 | 9.87E-17 |
| Sars | orange | 0.5915235 | 1.66E-06 | 0.43388087 | 0.00072368 |
| Celsr2 | red | -0.798175 | 5.71E-05 | -0.4038863 | 0.05723876 |
| Cry2 | red | -0.9375552 | 3.23E-05 | -0.8005722 | 0.00058017 |
| Psrc1 | orange | 2.03548551 | 1.49E-09 | 1.88044893 | 4.17E-08 |
| Mybphl | skyblue | -1.9622614 | 0.01442407 | -2.4484802 | 0.00253933 |
| Sort1 | skyblue | -1.6719625 | 1.11E-07 | -1.3568987 | 2.90E-05 |
| Ptprz1 | red | -1.101258 | 1.51E-06 | -0.5039149 | 0.03370497 |
| Psma5 | orange | 1.34239546 | 3.71E-14 | 1.20591131 | 2.13E-11 |
| Il3ra | orange | 1.50108034 | 4.51E-11 | 1.27890434 | 4.11E-08 |
| Gstm6 | red | -1.1904183 | 0.00383479 | -0.8737886 | 0.04630157 |
| Hist2h2be | red | -1.1723303 | 2.32E-09 | -0.8050718 | 7.59E-05 |
| Sf3b4 | orange | 0.65028127 | 9.82E-05 | 0.39175372 | 0.02684043 |
| Ssb | orange | 0.63633908 | 3.37E-08 | 0.37355179 | 0.00208483 |
| Lce3f | red | -0.9232622 | 0.00274053 | -0.5121615 | 0.12308523 |
| Lce1j | skyblue | -2.0355385 | 0.00033006 | -1.0027588 | 0.09745122 |

| | | | | | |
|---|---|---|---|---|---|
| Lce1i | skyblue | -1.5708163 | 0.00312366 | -1.2868494 | 0.01990274 |
| Lce1e | skyblue | -2.0277919 | 6.30E-06 | -1.3418246 | 0.00415688 |
| Lce1a2 | red | -1.2553487 | 0.00030451 | -0.7152147 | 0.05402131 |
| Msto1 | orange | 0.68157514 | 0.00081877 | 0.62172541 | 0.00313918 |
| Gm5641 | orange | 1.66363849 | 0.00922454 | 1.15562586 | 0.12412865 |
| Cdk7 | orange | 0.6302054 | 0.0034485 | 0.48027351 | 0.03351039 |
| Adgrv1 | skyblue | -2.1265977 | 0.00098155 | -1.4547046 | 0.03097478 |
| Nr2f1 | skyblue | -1.3207864 | 5.00E-07 | -1.393494 | 2.85E-07 |
| Zfp72 | red | -1.0889672 | 0.00066658 | -1.0351595 | 0.00212752 |
| Zfp874a | red | -0.5355757 | 0.04948761 | -0.869674 | 0.00153027 |
| Hist1h2bj | yellow | 2.601034 | 0.03092256 | 4.39135527 | 0.00498065 |
| Hist1h2ag | black | 5.89395413 | 0.0001641 | 3.47419515 | 0.00817008 |
| Hist1h3c | yellow | 3.24504661 | 0.0001786 | 4.45274244 | 0.00056198 |
| Ctxn3 | red | -1.148679 | 0.00304457 | -0.9134828 | 0.02507013 |
| Prdm6 | red | -1.1578021 | 1.46E-07 | -0.8884489 | 9.34E-05 |
| Lyz1 | skyblue | -2.1688946 | 4.89E-05 | -1.5616597 | 0.00500819 |
| Lyz2 | red | -1.4623657 | 5.26E-06 | -1.0223627 | 0.00224451 |
| Ank3 | red | -0.7416005 | 0.00057026 | -0.4521968 | 0.04808581 |
| Gm10275 | orange | 0.86480927 | 0.002099 | 0.4405298 | 0.15081111 |
| Gm5595 | red | -0.8879873 | 0.00052537 | -1.1349521 | 1.10E-05 |
| Psmb3 | orange | 1.08275814 | 9.87E-12 | 0.90065024 | 3.22E-08 |
| Tmem100 | skyblue | -1.9755868 | 1.55E-10 | -1.3406276 | 2.83E-05 |
| Msi2 | red | -0.6123168 | 0.00046648 | -0.5334301 | 0.003137 |
| Wfdc17 | yellow | 3.38693605 | 1.72E-16 | 3.49857036 | 3.13E-17 |
| Slfn9 | yellow | 3.74551791 | 1.39E-39 | 3.58010987 | 5.04E-36 |
| Cacng7 | skyblue | -1.6539016 | 9.53E-08 | -0.9161871 | 0.00304371 |
| Nlrp1a | yellow | 2.39489144 | 0.00267196 | 3.16540274 | 0.0005688 |
| Ahnak | red | -0.5909832 | 9.29E-07 | -0.477926 | 0.00012162 |
| Sco1 | orange | 1.12752941 | 6.27E-07 | 0.40261976 | 0.10307839 |
| Pabpn1l | skyblue | -2.9218999 | 0.01098233 | -4.1223689 | 0.00472952 |
| 4930438A08I | black | 5.80624979 | 9.71E-05 | 3.77697633 | 0.00017774 |
| Irgm2 | yellow | 2.27363339 | 3.90E-29 | 2.32478932 | 2.16E-30 |
| 9930111J21F | orange | 1.58462806 | 9.35E-18 | 1.34070842 | 9.60E-13 |
| 9930111J21F | yellow | 2.08397068 | 2.47E-08 | 2.20071377 | 6.87E-09 |
| Gm12166 | orange | 0.72139765 | 0.00144949 | 0.40078577 | 0.10710552 |
| Spdl1 | yellow | 2.95938535 | 9.09E-23 | 2.80167293 | 2.05E-20 |
| Hba-a2 | red | -0.5882068 | 0.13929435 | -1.1286427 | 0.00370932 |
| Hba-a1 | red | -0.5052548 | 0.16699056 | -1.1685124 | 0.00093947 |
| Fcho1 | yellow | 2.4059204 | 9.45E-35 | 2.56390419 | 1.06E-38 |
| Ssbp4 | orange | 0.77182307 | 0.00013188 | 0.75363706 | 0.00027592 |
| Sp140 | yellow | 2.29726179 | 8.63E-27 | 2.32812915 | 3.64E-27 |
| Sp110 | yellow | 2.50332942 | 5.64E-48 | 2.46047296 | 3.91E-46 |
| Fam149a | red | -0.9417498 | 3.57E-08 | -0.592255 | 0.00084197 |

| | | | | | |
|---|---|---|---|---|---|
| Fat1 | red | -0.666716 | 1.52E-06 | -0.315858 | 0.03321981 |
| Gmppb | orange | 1.07542659 | 2.99E-12 | 1.05397576 | 1.32E-11 |
| Mpzl3 | orange | 0.80506695 | 0.00077125 | 0.36034384 | 0.16611578 |
| Ccdc153 | skyblue | -2.246412 | 1.91E-18 | -1.4033525 | 1.07E-07 |
| Rnf213 | yellow | 2.76579754 | 2.33E-68 | 2.72512616 | 2.27E-66 |
| Prss36 | skyblue | -1.6443081 | 2.74E-10 | -1.1898664 | 9.98E-06 |
| Capza1 | orange | 1.05429522 | 6.33E-09 | 0.8698902 | 3.16E-06 |
| Ampd1 | skyblue | -1.8553858 | 1.78E-05 | -1.0155894 | 0.0282268 |
| Fbxo39 | yellow | 3.22161866 | 1.01E-10 | 2.32623438 | 2.34E-08 |
| Nlrp1b | yellow | 3.70465283 | 9.16E-32 | 2.72295625 | 9.77E-20 |
| Art5 | red | -1.4324979 | 0.00390137 | -0.8245626 | 0.12430572 |
| Il18bp | yellow | 4.20132378 | 1.77E-63 | 3.69300568 | 6.36E-50 |
| Adamtsl3 | red | -1.051871 | 4.48E-06 | -0.8678837 | 0.00024297 |
| BC094916 | yellow | 4.04911016 | 1.71E-20 | 3.66382735 | 5.33E-16 |
| Fcrl6 | black | 5.05399821 | 0.00082672 | 4.48934197 | 0.00451562 |
| Rgma | skyblue | -1.5072681 | 9.61E-09 | -1.1006126 | 4.95E-05 |
| Fcrlb | black | 5.43565449 | 7.26E-27 | 5.77555386 | 1.85E-25 |
| Wfdc10 | skyblue | -6.5319191 | 2.13E-05 | -0.2116878 | 0.86868079 |
| Wfdc16 | skyblue | -6.6323734 | 3.84E-05 | -6.0524766 | 0.00024365 |
| Rasal2 | red | -0.7009034 | 0.00550358 | -0.8851603 | 0.00052673 |
| Slc17a7 | skyblue | -1.7862901 | 0.12223247 | -2.9838995 | 0.00919326 |
| Mn1 | red | -1.0842243 | 5.24E-07 | -0.8505971 | 0.00014589 |
| Gm572 | skyblue | -1.7082872 | 0.00050022 | -0.7706627 | 0.15138163 |
| Fv1 | red | -0.9246796 | 0.00518769 | -0.6728994 | 0.05011165 |
| Gm13251 | orange | 2.06671254 | 0.00208473 | 1.48151162 | 0.03404559 |
| Sox13 | red | -0.9704746 | 4.34E-08 | -0.7705104 | 2.44E-05 |
| Rnf186 | skyblue | -2.0349359 | 7.24E-14 | -1.4628294 | 1.60E-07 |
| Runx3 | yellow | 3.96013265 | 1.90E-31 | 3.57914481 | 9.40E-26 |
| Sars2 | orange | 0.7741406 | 1.57E-05 | 0.69910829 | 0.00016364 |
| 1700049G17 | red | -0.7475532 | 0.00062003 | -0.7641349 | 0.00062058 |
| Gm10282 | orange | 0.66487352 | 7.09E-05 | 0.63327349 | 0.00020734 |
| Tmem200b | skyblue | -1.6834443 | 1.85E-12 | -1.2073968 | 8.47E-07 |
| Rmdn3 | orange | 0.79789339 | 2.49E-10 | 0.73523177 | 1.06E-08 |
| Gm12942 | red | -0.7199931 | 9.15E-05 | -0.7746358 | 4.56E-05 |
| Rbm47 | orange | 0.32509024 | 0.04092757 | 0.61125937 | 1.00E-04 |
| Pnmal2 | skyblue | -1.7760368 | 6.61E-12 | -1.8462026 | 1.66E-12 |
| Zmynd12 | skyblue | -1.7688746 | 3.24E-06 | -1.139108 | 0.00219308 |
| 6330408A02I | red | -0.9631453 | 3.06E-09 | -1.0270408 | 6.08E-10 |
| Trabd2b | red | -0.6922333 | 0.00295939 | -0.8024925 | 0.00072664 |
| Lilra5 | red | -1.4813183 | 0.00031635 | -0.8761161 | 0.04532625 |
| Ldlrad1 | skyblue | -2.667737 | 2.99E-14 | -1.5106995 | 1.75E-05 |
| Rabepk | orange | 0.89226397 | 0.00017054 | 0.76224211 | 0.00194449 |
| Rasgrp3 | red | -1.1941597 | 1.00E-12 | -0.9490135 | 3.74E-08 |

| | | | | | |
|---|---|---|---|---|---|
| Treml2 | yellow | 2.03927738 | 9.82E-10 | 2.0743069 | 3.25E-10 |
| Nr2c2ap | orange | 0.8779066 | 0.00971113 | 1.16061858 | 0.00167472 |
| 1700029J07F | skyblue | -1.6294652 | 5.79E-08 | -1.0699025 | 0.00052524 |
| Arhgef10 | red | -0.9641228 | 3.65E-10 | -0.6748684 | 2.24E-05 |
| Ccdc78 | red | -1.2418989 | 2.65E-07 | -0.6796722 | 0.00683194 |
| Naip5 | yellow | 2.11097486 | 6.17E-17 | 1.95953021 | 1.18E-14 |
| Npw | skyblue | -2.0863799 | 1.63E-10 | -1.1338256 | 0.00079414 |
| Gpr31b | black | 7.39554331 | 4.85E-07 | 8.06991734 | 6.21E-08 |
| Bves | skyblue | -1.7976705 | 1.43E-08 | -1.6562566 | 3.38E-07 |
| Armc2 | skyblue | -1.7770502 | 3.21E-09 | -1.3654649 | 7.16E-06 |
| G630090E17 | skyblue | -2.2377853 | 0.00837382 | -1.6666665 | 0.0578368 |
| C1qtnf9 | skyblue | -1.48675 | 0.00543481 | -1.4819083 | 0.00745583 |
| Setdb2 | yellow | 2.08615114 | 1.30E-35 | 1.96619103 | 5.10E-31 |
| Ect2l | skyblue | -2.1431724 | 1.46E-08 | -1.1382182 | 0.00251877 |
| 2410004P03I | skyblue | -1.8006709 | 3.39E-20 | -1.1112633 | 2.14E-08 |
| LOC1008624 | orange | 0.58869981 | 2.71E-05 | 0.43382631 | 0.00297323 |
| Psmg4 | orange | 1.41594036 | 1.47E-13 | 1.13082583 | 8.39E-09 |
| 1110002L01F | red | -0.7944777 | 0.0092433 | -0.5847192 | 0.06604376 |
| Tmem139 | red | -0.7034747 | 0.02411417 | -0.927124 | 0.00242336 |
| Klrg2 | red | -0.6416424 | 0.00547361 | -0.1673733 | 0.54315754 |
| 3425401B19I | skyblue | -1.9390607 | 8.70E-09 | -1.5031741 | 1.81E-05 |
| Cfap44 | red | -1.5156325 | 1.14E-06 | -0.6884128 | 0.03731244 |
| Akr1c19 | red | -1.2073754 | 4.39E-05 | -0.554642 | 0.08420677 |
| Tigit | black | 7.48778557 | 1.36E-62 | 5.78554522 | 4.10E-63 |
| Fam189a2 | red | -1.0669796 | 8.13E-05 | -0.7643174 | 0.00683346 |
| Cebpd | orange | 1.60314618 | 1.41E-11 | 1.03186801 | 2.82E-05 |
| Mta2 | orange | 0.58141971 | 5.71E-17 | 0.47322449 | 2.51E-11 |
| Rom1 | orange | 0.64933054 | 4.24E-05 | 0.6966291 | 2.42E-05 |
| Ints5 | orange | 0.63884197 | 3.58E-06 | 0.28128311 | 0.059463 |
| Lrrn4cl | red | -1.4438288 | 1.86E-06 | -1.0966228 | 0.00059081 |
| Ttc9c | orange | 0.72357392 | 7.26E-07 | 0.60768201 | 5.41E-05 |
| Mpst | red | -0.7540962 | 9.89E-06 | -0.4953547 | 0.00550025 |
| Csf2rb | yellow | 2.27081888 | 9.02E-14 | 2.32442754 | 3.13E-14 |
| Csf2rb2 | yellow | 2.71174983 | 9.04E-14 | 2.83316993 | 8.63E-15 |
| Ncf4 | yellow | 3.09207267 | 2.01E-23 | 3.07219128 | 6.12E-23 |
| Apol7e | yellow | 3.44462721 | 3.48E-23 | 3.82042227 | 9.72E-25 |
| Tmem28 | skyblue | -1.6993658 | 4.33E-09 | -0.9846994 | 0.00094847 |
| Smpd5 | orange | 1.21856783 | 0.00081828 | 1.12702211 | 0.00272189 |
| Zhx2 | red | -0.9512685 | 5.46E-10 | -0.8836271 | 1.47E-08 |
| Apcdd1 | red | -1.1940298 | 2.95E-15 | -1.0423194 | 6.75E-12 |
| Mcc | red | -1.1656665 | 1.24E-08 | -0.8720869 | 3.86E-05 |
| Gm94 | red | -0.9172433 | 0.00829747 | -0.7192051 | 0.04833562 |
| Fndc1 | skyblue | -2.0917991 | 4.33E-33 | -1.4361113 | 4.76E-16 |

| | | | | | |
|---|---|---|---|---|---|
| Ccnf | yellow | 2.7279135 | 2.99E-34 | 3.22392103 | 1.04E-42 |
| A530040E14I | yellow | 3.01805184 | 3.84E-10 | 3.37270339 | 1.57E-11 |
| Ang | skyblue | -1.2602346 | 4.85E-13 | -1.2760662 | 9.90E-13 |
| Trim6 | orange | 1.69242577 | 3.93E-08 | 1.64032507 | 1.14E-07 |
| Klf12 | skyblue | -1.2594642 | 9.47E-06 | -1.2322725 | 2.41E-05 |
| 1700024G13 | skyblue | -3.9713637 | 3.05E-12 | -2.4265127 | 1.21E-05 |
| Ppp1r3e | red | -0.5902657 | 0.00030296 | -0.4375579 | 0.01049771 |
| Fam84b | red | -1.4117871 | 3.88E-10 | -1.0727733 | 3.91E-06 |
| Slc39a2 | orange | 1.26167462 | 0.00267297 | 0.62495473 | 0.17111879 |
| Gm10369 | orange | 1.81898053 | 0.00417432 | 1.21096937 | 0.05492083 |
| 5930412G12 | skyblue | -2.5406749 | 4.42E-05 | -0.8392255 | 0.19106467 |
| Slfn2 | yellow | 3.83747826 | 4.90E-44 | 3.76319781 | 8.65E-42 |
| Slfn10-ps | black | 4.32856762 | 8.18E-06 | 7.39511299 | 5.70E-07 |
| Lyrm9 | red | -0.6621198 | 0.00100818 | -0.7884964 | 0.00013498 |
| Adam1a | red | -0.6694319 | 0.06145344 | -1.1271127 | 0.00150424 |
| Spef2 | red | -1.3486936 | 6.31E-06 | -0.7970464 | 0.00996804 |
| Smim10l1 | red | -0.8856757 | 2.10E-08 | -0.8444778 | 1.66E-07 |
| Myo18b | red | -0.6367824 | 0.00299205 | -0.5289578 | 0.01822136 |
| Klra14-ps | orange | 1.10314595 | 0.06450949 | 1.61235299 | 0.00968954 |
| Gm10419 | skyblue | -1.7863498 | 0.00186814 | -0.9186326 | 0.12923552 |
| Acrbp | red | -1.048371 | 1.55E-05 | -1.1442635 | 3.95E-06 |
| G530011O06 | yellow | 3.74973428 | 8.84E-06 | 3.46963213 | 4.83E-05 |
| Gpr27 | skyblue | -1.6791875 | 1.02E-06 | -1.194102 | 0.00097098 |
| 4933439C10 | red | -0.6356239 | 0.00366057 | -0.2584482 | 0.2838747 |
| Gm10435 | skyblue | -2.4545671 | 0.0058843 | -3.1546565 | 0.00064214 |
| Gm12258 | red | -0.8842665 | 0.00019132 | -0.3661057 | 0.15681121 |
| Sod3 | red | -0.6594869 | 1.09E-05 | -0.3705641 | 0.01959683 |
| Nup62cl | skyblue | -1.8510908 | 0.00318319 | -1.2430074 | 0.05079321 |
| Ptgr2 | red | -0.7500519 | 5.89E-06 | -0.4431449 | 0.01111574 |
| Acot1 | red | -1.1639691 | 4.28E-05 | -0.4321234 | 0.16633178 |
| Gprasp2 | skyblue | -3.0817491 | 2.76E-06 | -1.6193244 | 0.01688374 |
| Oip5 | yellow | 3.23799677 | 7.63E-08 | 1.74473927 | 0.00359644 |
| Vamp5 | red | -0.5883994 | 2.37E-05 | -0.4466869 | 0.0022436 |
| Gpr174 | yellow | 2.29068743 | 6.35E-14 | 2.41242137 | 8.17E-14 |
| Drc1 | skyblue | -1.9889167 | 4.79E-19 | -1.4460976 | 1.40E-10 |
| 4930599N23 | yellow | 1.86085574 | 7.85E-06 | 2.06614123 | 1.17E-06 |
| Gm12000 | yellow | 2.67675201 | 0.00399587 | 3.95361719 | 0.0035577 |
| 9330158H04 | red | -0.623139 | 0.02691901 | -0.9237615 | 0.00137116 |
| 2500004C02 | skyblue | -1.358532 | 0.00021492 | -1.2247842 | 0.00131792 |
| Nudt10 | skyblue | -1.5952226 | 0.00070804 | -1.2346672 | 0.00848137 |
| Nudt11 | skyblue | -1.8490462 | 1.44E-05 | -1.290136 | 0.00311294 |
| C920025E04 | yellow | 2.55246101 | 2.75E-23 | 2.47256734 | 4.29E-21 |
| Gm6034 | yellow | 3.44630846 | 1.11E-08 | 2.20816052 | 8.78E-05 |

| | | | | | |
|---|---|---|---|---|---|
| H2-Q6 | yellow | 2.19315427 | 3.72E-30 | 2.03863195 | 4.43E-26 |
| H2-D1 | orange | 1.75993329 | 9.54E-31 | 1.49637198 | 2.42E-22 |
| Lst1 | yellow | 3.11672731 | 8.99E-23 | 2.99737614 | 1.13E-19 |
| Ly6g6d | skyblue | -1.4321929 | 1.13E-06 | -1.3068118 | 3.05E-05 |
| C4b | orange | 1.06556931 | 7.48E-09 | 1.2138045 | 5.70E-11 |
| H2-Ab1 | orange | 1.71742306 | 5.95E-20 | 1.31662759 | 6.54E-12 |
| Cyp4f15 | skyblue | -2.7233558 | 3.76E-10 | -1.4331653 | 0.00129432 |
| Gm10505 | yellow | 1.6916666 | 0.05968684 | 2.72632101 | 0.00448869 |
| Arhgdig | red | -1.3584901 | 1.40E-08 | -0.826306 | 0.00089842 |
| Sft2d1 | orange | 0.70290704 | 0.00035869 | 0.59315226 | 0.00365169 |
| Rsph3a | red | -0.756887 | 2.04E-06 | -0.5399559 | 0.00124368 |
| Ifi204 | black | 6.02132166 | 1.45E-55 | 5.22197265 | 1.21E-42 |
| AI607873 | yellow | 4.1503327 | 1.59E-32 | 3.43542126 | 1.17E-22 |
| Pydc4 | yellow | 2.74457156 | 1.49E-39 | 2.52952413 | 1.31E-33 |
| Gm10521 | yellow | 2.28356659 | 3.44E-07 | 3.25042944 | 2.29E-12 |
| Gm4951 | yellow | 3.9963364 | 4.02E-30 | 3.75785195 | 9.71E-27 |
| Ppp1r12b | red | -0.8669168 | 9.46E-12 | -0.4972908 | 0.00018954 |
| 3222401L13R | skyblue | -2.0800887 | 1.16E-06 | -1.4561598 | 0.00092602 |
| Pcdhb22 | red | -0.8791334 | 6.34E-05 | -0.6572812 | 0.00498608 |
| 1700066B19l | yellow | 2.51486745 | 1.81E-05 | 1.59390882 | 0.00470344 |
| Prob1 | skyblue | -1.6904387 | 5.57E-11 | -1.4289783 | 6.04E-08 |
| D2hgdh | red | -0.6272871 | 0.0005251 | -0.6908553 | 0.00018957 |
| Fbxo36 | red | -0.9307225 | 0.00119455 | -0.6868308 | 0.02190939 |
| Catip | skyblue | -1.7758451 | 2.24E-17 | -0.9303246 | 1.43E-05 |
| Nbeal1 | red | -0.8374537 | 1.47E-05 | -0.5996441 | 0.00287227 |
| Klhl21 | red | -0.9170462 | 6.26E-10 | -0.7115522 | 3.15E-06 |
| Rpl31 | orange | 0.59553241 | 0.00021546 | 0.37705654 | 0.02645714 |
| Apitd1 | yellow | 2.41606851 | 1.44E-12 | 1.70522341 | 2.96E-06 |
| Gm13241 | yellow | 2.67835143 | 0.00703054 | 1.8084504 | 0.04249949 |
| 5730409E04l | red | -0.7763373 | 4.95E-06 | -0.6788943 | 0.00011834 |
| Btbd19 | orange | 1.29522524 | 5.68E-09 | 1.02313206 | 7.62E-06 |
| Gm10575 | red | -0.7104591 | 0.00094363 | -0.4954382 | 0.02420595 |
| Alg6 | red | -0.6618221 | 0.00552464 | -0.2237415 | 0.41356167 |
| Cdkn2b | orange | 0.65572026 | 2.08E-06 | 0.44008838 | 0.0023161 |
| Fam196a | red | -0.5305739 | 0.07915151 | -1.0993324 | 0.00052411 |
| Iqck | red | -0.7836396 | 0.0020085 | -0.4269162 | 0.11403528 |
| Ccl27a | red | -0.8737089 | 0.00011157 | -1.012329 | 1.34E-05 |
| Il11ra1 | red | -0.6780681 | 7.41E-06 | -0.652733 | 2.59E-05 |
| Gm1966 | yellow | 2.81292408 | 1.78E-21 | 3.08565882 | 2.61E-25 |
| Mob3b | red | -0.9419289 | 1.85E-10 | -0.4965737 | 0.00140937 |
| Hbb-bt | red | -0.8892137 | 0.03128833 | -1.1841994 | 0.0043499 |
| Olfr550 | skyblue | -1.1242767 | 0.03455225 | -1.5131716 | 0.00600624 |
| Gm9574 | yellow | 1.77578782 | 1.04E-07 | 2.44909942 | 1.28E-12 |

| | | | | | |
|---|---|---|---|---|---|
| Rhog | orange | 1.35514223 | 6.96E-11 | 1.1123978 | 1.79E-07 |
| Ttpa | red | -0.9191602 | 0.00521211 | -0.637396 | 0.0593869 |
| Klhl40 | skyblue | -2.1087588 | 4.30E-08 | -0.8173287 | 0.04991458 |
| Omp | orange | 2.04538764 | 0.00020864 | 1.70566724 | 0.00177073 |
| Fam169b | orange | 1.22252319 | 1.43E-07 | 1.97101928 | 4.72E-17 |
| Snrnp40 | orange | 0.88203931 | 1.57E-07 | 0.74031118 | 2.02E-05 |
| Svip | skyblue | -1.7918303 | 2.33E-08 | -1.2608138 | 0.00014346 |
| Saa1 | black | 9.15344908 | 4.32E-12 | 7.17760219 | 1.96E-09 |
| Rpl13a | orange | 0.58287815 | 0.00051627 | 0.50634379 | 0.0039954 |
| Nlrc5 | yellow | 3.29760062 | 2.10E-69 | 3.41489981 | 2.40E-74 |
| Ces1h | red | -1.3831654 | 0.00094064 | -0.9484252 | 0.03089536 |
| Zfp788 | red | -0.5776711 | 0.00259384 | -0.5901595 | 0.00250323 |
| Zfp791 | red | -0.9576227 | 0.00372483 | -0.552878 | 0.10997279 |
| Adh1 | skyblue | -1.2663579 | 1.14E-18 | -1.2538183 | 4.46E-18 |
| 2210011C24 | skyblue | -1.9937487 | 2.27E-05 | -1.3512234 | 0.00592155 |
| Cox7a1 | skyblue | -2.2259714 | 1.19E-06 | -1.9599816 | 3.57E-05 |
| Zfp382 | orange | 1.53178056 | 0.00084029 | 1.56583003 | 0.00085121 |
| Dda1 | orange | 0.58423095 | 5.16E-05 | 0.551819 | 0.00019895 |
| Gm10654 | yellow | 2.05903139 | 0.00729852 | 4.6332301 | 1.02E-06 |
| Cyp2a4 | skyblue | -3.6143163 | 0.00142849 | -2.5897283 | 0.01398194 |
| Amy1 | red | -1.4315539 | 3.86E-05 | -0.9986547 | 0.00559589 |
| Zfp94 | red | -0.8806196 | 0.00239388 | -0.8853778 | 0.00282437 |
| I830077J02R | yellow | 2.89233959 | 4.40E-19 | 2.76792523 | 1.85E-17 |
| C5ar2 | orange | 1.53583002 | 2.59E-05 | 1.44265784 | 0.00012445 |
| Ehd2 | red | -0.6671096 | 1.99E-08 | -0.433711 | 0.00046858 |
| AI429214 | red | -0.7509426 | 0.02343843 | -0.9741232 | 0.00372491 |
| Pira11 | yellow | 3.84151892 | 2.18E-08 | 2.67297545 | 5.93E-05 |
| Gm15448 | yellow | 3.74537829 | 2.37E-08 | 3.24325013 | 7.88E-07 |
| Zfp872 | orange | 1.67992857 | 0.00111951 | 1.00379556 | 0.06959608 |
| Spc24 | yellow | 2.15234009 | 3.04E-12 | 2.31369223 | 2.11E-13 |
| Mex3a | orange | 0.09281356 | 0.79942726 | 1.0944106 | 0.00053892 |
| Clec4g | skyblue | -2.7479324 | 0.00352942 | -2.8514837 | 0.00286501 |
| A430078G23 | orange | 1.28057335 | 5.78E-07 | 0.96338567 | 0.00027736 |
| Fat3 | skyblue | -1.504627 | 1.08E-12 | -1.5726953 | 3.85E-13 |
| Zfas1 | orange | 1.59526429 | 3.18E-20 | 1.46740219 | 5.72E-17 |
| 4931440P22I | orange | 0.83221561 | 0.00806886 | 0.79717731 | 0.01625456 |
| Ankub1 | skyblue | -1.7829129 | 0.00021957 | -1.0881857 | 0.01344824 |
| Mafb | yellow | 2.92601367 | 1.40E-18 | 2.64309135 | 3.60E-15 |
| Tldc2 | yellow | 1.33070443 | 0.0590225 | 3.17591111 | 0.00169637 |
| Gm7120 | red | -0.173013 | 0.73879024 | -1.5596849 | 0.00075336 |
| Sox2 | red | -1.2746551 | 3.52E-09 | -0.6975786 | 0.00189795 |
| Mcidas | yellow | 1.88005772 | 0.00538244 | 2.61717184 | 8.56E-05 |
| Eif2s2 | orange | 1.24392535 | 2.13E-07 | 0.97838326 | 8.03E-05 |

| | | | | | |
|---|---|---|---|---|---|
| Ttll9 | red | -1.052925 | 0.00295334 | -0.8411428 | 0.01889423 |
| Foxs1 | orange | 1.20657231 | 0.0199216 | 1.42188206 | 0.00756585 |
| LOC1000389 | yellow | 4.18837556 | 6.96E-09 | 4.20467398 | 1.06E-08 |
| Defb23 | skyblue | -3.9797472 | 0.00019508 | -0.693228 | 0.38149283 |
| Zfp950 | red | -0.6196007 | 0.00445169 | -0.6777856 | 0.0023558 |
| Thbd | skyblue | -1.4631616 | 4.08E-10 | -1.2290427 | 2.95E-07 |
| Kiz | red | -0.9596054 | 2.23E-06 | -0.760775 | 0.00027736 |
| Gm5535 | yellow | 3.00601569 | 0.00164216 | 3.54286844 | 0.0010191 |
| Ankef1 | skyblue | -2.426814 | 0.00166133 | -1.5031225 | 0.05815933 |
| Ube2n | orange | 0.61546945 | 1.10E-05 | 0.38527229 | 0.00885995 |
| 4833422C13 | red | -1.4050759 | 0.00023967 | -1.0692454 | 0.0064653 |
| Plxnc1 | orange | 0.59250999 | 0.00023086 | 0.48825272 | 0.00355846 |
| Arrdc3 | red | -1.2774733 | 8.75E-05 | -1.0275294 | 0.00231374 |
| Slc4a11 | orange | 1.72566056 | 2.52E-06 | 1.66870864 | 6.81E-06 |
| Itpa | orange | 0.54405729 | 0.00131806 | 0.59870305 | 0.00052729 |
| Gas2l3 | orange | 1.6054283 | 1.01E-05 | 1.87734309 | 6.58E-07 |
| Gm14005 | orange | 1.0665129 | 0.0002764 | 1.15112517 | 0.00012932 |
| Pdzd7 | red | -0.7142167 | 0.00194916 | -0.9249574 | 6.67E-05 |
| Hpse2 | skyblue | -2.1298826 | 0.00017917 | -1.4257296 | 0.01136891 |
| Ctla2b | orange | 0.74375828 | 0.04473289 | 1.02358731 | 0.00752773 |
| Grk6 | orange | 1.10088266 | 3.74E-12 | 1.05618403 | 4.53E-11 |
| Ifit3 | yellow | 3.6383125 | 4.44E-132 | 3.52236213 | 6.08E-124 |
| Gm14207 | skyblue | -1.4053854 | 2.61E-07 | -1.3433382 | 1.02E-06 |
| Pak6 | orange | 1.4747955 | 2.00E-28 | 1.0146678 | 2.38E-14 |
| Fibin | skyblue | -1.6931627 | 0.00140266 | -1.3168171 | 0.01707498 |
| Dcdc5 | skyblue | -3.0769743 | 0.00219145 | -0.9092416 | 0.39098227 |
| Qser1 | red | -0.6345465 | 0.00037085 | -0.4269303 | 0.02268116 |
| AW112010 | yellow | 3.39628903 | 8.93E-58 | 3.03482563 | 5.94E-46 |
| Fjx1 | orange | 0.45514419 | 0.01762743 | 0.63897708 | 0.00101554 |
| Yae1d1 | red | -0.9069528 | 1.30E-10 | -0.6244098 | 2.20E-05 |
| Cd80 | yellow | 3.18269967 | 1.03E-14 | 2.09169161 | 4.69E-07 |
| Ccdc162 | skyblue | -1.9853191 | 1.52E-14 | -0.9707578 | 0.00028402 |
| Heg1 | red | -0.8580136 | 2.77E-09 | -0.5433768 | 0.00030803 |
| Cenpw | yellow | 2.2059763 | 1.53E-07 | 2.81710583 | 6.21E-10 |
| Ttc30a1 | red | -0.538668 | 0.04228785 | -0.791697 | 0.00360677 |
| Ttc30b | red | -1.4581599 | 6.54E-09 | -1.0338452 | 6.27E-05 |
| Wipf1 | orange | 0.78747923 | 2.27E-06 | 0.56349487 | 0.0011509 |
| Aldh3b2 | red | -0.9408339 | 0.0030665 | -0.7710736 | 0.02003912 |
| Erich2 | skyblue | -1.5663355 | 2.99E-10 | -1.2970293 | 3.00E-07 |
| Sp5 | red | -1.1217348 | 0.00339616 | -0.6985613 | 0.0918659 |
| Klhl41 | red | -1.0166326 | 0.00013989 | -0.7942049 | 0.00429444 |
| Scn2a1 | skyblue | -1.5146192 | 0.00702696 | -0.8361675 | 0.13673299 |
| Fign | skyblue | -1.4755009 | 4.33E-09 | -1.3541239 | 1.03E-07 |

| | | | | | |
|---|---|---|---|---|---|
| Rprm | red | -1.1743681 | 0.00765642 | -0.9848899 | 0.03138556 |
| 2810410L24F | red | -1.3200439 | 1.60E-10 | -0.9323185 | 1.06E-05 |
| Hoxc4 | skyblue | -1.5052693 | 7.10E-05 | -1.1420624 | 0.00371064 |
| Smim6 | red | -1.090428 | 0.00016583 | -0.645143 | 0.03546331 |
| Gm13375 | red | -0.8591454 | 0.00112444 | -0.8494156 | 0.00178237 |
| Rsph10b | skyblue | -1.8459088 | 7.89E-09 | -0.9638194 | 0.00362316 |
| 6330403L08F | red | -1.0074078 | 1.48E-14 | -0.7940613 | 3.49E-09 |
| Hoxb2 | red | -0.8153532 | 0.00309785 | -0.2651865 | 0.38342417 |
| Nynrin | red | -0.940571 | 1.88E-08 | -0.760938 | 7.72E-06 |
| Zfp652 | red | -0.6369638 | 0.00014706 | -0.5351497 | 0.00203295 |
| Ly6a | yellow | 2.18972903 | 4.46E-49 | 1.93749953 | 1.64E-38 |
| Msrb1 | orange | 1.18017533 | 1.82E-07 | 0.97663789 | 2.78E-05 |
| Sox4 | red | -1.467146 | 1.96E-16 | -0.6435943 | 0.00070558 |
| Wfdc3 | skyblue | -1.567679 | 0.00124225 | -0.946216 | 0.05743205 |
| Acsf2 | red | -0.6350379 | 5.70E-05 | -0.4845429 | 0.00302034 |
| 2700094K13I | orange | 0.75019962 | 3.43E-08 | 0.54675917 | 0.00011404 |
| Ass1 | orange | 1.44406369 | 2.38E-13 | 0.95864758 | 2.75E-06 |
| Trbv1 | yellow | 2.95082263 | 1.35E-07 | 3.35204854 | 4.96E-09 |
| Trbv2 | yellow | 1.77357579 | 0.00055415 | 3.37916028 | 2.04E-08 |
| Trbv3 | yellow | 3.22276127 | 4.48E-08 | 3.71945976 | 1.50E-09 |
| Trbv13-1 | black | 2.66029025 | 9.12E-06 | 4.94175983 | 1.25E-11 |
| Trbv13-2 | yellow | 0.89134767 | 0.14973655 | 3.2215148 | 1.81E-07 |
| Trbv13-3 | yellow | 2.32298164 | 7.02E-05 | 3.98546174 | 2.26E-07 |
| Trbv14 | black | 4.41519891 | 2.42E-11 | 4.85356772 | 1.25E-11 |
| Trbv15 | yellow | 2.24509667 | 0.00673768 | 4.03898373 | 1.60E-05 |
| Trbv16 | yellow | 2.78347476 | 3.43E-08 | 4.07389388 | 3.34E-12 |
| Trbv17 | yellow | 3.15687167 | 0.00035394 | 3.88106294 | 4.32E-05 |
| Trbv19 | yellow | 3.83286729 | 4.17E-12 | 3.92520559 | 2.04E-10 |
| Trbv20 | yellow | 3.23253268 | 8.81E-09 | 3.97663414 | 1.12E-10 |
| Trbv26 | yellow | 2.73753078 | 0.00406969 | 1.85135791 | 0.06534277 |
| Trbv29 | black | 3.33483232 | 2.11E-07 | 5.45613747 | 7.30E-11 |
| Trbj1-3 | yellow | 1.37391249 | 0.02348951 | 3.37631336 | 1.21E-06 |
| Trbj1-4 | yellow | 1.8177358 | 0.00505536 | 2.41197722 | 0.00035175 |
| Trbj1-5 | orange | 1.28274169 | 0.019666 | 2.44055675 | 3.48E-05 |
| Trbj1-7 | yellow | 1.88548329 | 0.04005106 | 3.43858914 | 0.00352952 |
| Trbc1 | yellow | 1.93425298 | 0.00225598 | 3.42106913 | 6.33E-08 |
| Trbj2-1 | yellow | 0.9254043 | 0.19924751 | 3.56840425 | 3.18E-05 |
| Trbj2-2 | yellow | 1.76936345 | 0.02496277 | 3.86611822 | 0.00065939 |
| Trbj2-3 | yellow | 1.56371512 | 0.08556506 | 3.37833565 | 0.00216434 |
| Trbj2-5 | yellow | 2.30791277 | 0.02671942 | 3.36800117 | 0.0059693 |
| Trbj2-7 | yellow | 2.15822433 | 0.00011603 | 3.20191175 | 8.99E-08 |
| Trbc2 | yellow | 2.30037326 | 0.00172858 | 3.43302093 | 3.28E-06 |
| Trbv31 | yellow | 2.25559512 | 0.00106864 | 4.52041285 | 9.62E-07 |

| | | | | | |
|---|---|---|---|---|---|
| Igkv2-112 | skyblue | -2.5303225 | 0.00741257 | -1.2697705 | 0.28681287 |
| Igkv4-50 | yellow | 4.002866 | 0.00978195 | 2.80775256 | 0.0885085 |
| Igkv5-48 | yellow | 3.0894726 | 1.94E-06 | 3.61701961 | 7.47E-08 |
| Ighg2c | yellow | 4.29122673 | 5.50E-05 | 1.667971 | 0.14967572 |
| Ighg2b | yellow | 2.99729162 | 0.00108346 | 3.12142834 | 0.00090028 |
| Ighg3 | yellow | 3.59738183 | 0.00696646 | 2.07884256 | 0.14596563 |
| Ighj1 | yellow | 3.12972948 | 0.00453379 | 1.50379757 | 0.13785546 |
| Tcrg-C2 | yellow | 2.45190854 | 4.42E-08 | 2.12596883 | 5.90E-06 |
| Trav3-1 | yellow | 1.18894123 | 0.47043227 | 6.04740263 | 0.00151007 |
| Trav8d-2 | black | 6.55548958 | 0.00010515 | 2.07064503 | 0.1606116 |
| Trav16d-dv11 | black | 6.22521331 | 0.0035154 | 3.72961046 | 0.08989549 |
| Trav16n | black | 4.73462539 | 4.89E-07 | 5.71700072 | 1.20E-05 |
| Trav14n-3 | black | 3.95610859 | 0.00244704 | 5.05801671 | 0.00066918 |
| Trav13-1 | black | 3.63278159 | 0.00095582 | 5.391157 | 0.000684 |
| Trav16 | black | 5.0468954 | 1.06E-06 | 5.61121666 | 7.44E-05 |
| Traj16 | black | 4.37815172 | 0.00018512 | 5.92808593 | 0.00011181 |
| Trac | yellow | 3.56359511 | 2.42E-64 | 3.55053044 | 2.06E-61 |
| Snora17 | orange | 1.59385409 | 0.00030205 | 0.39039827 | 0.42866346 |
| Snord89 | orange | 1.91100936 | 0.00054101 | 1.89415804 | 0.00074131 |
| Snord83b | orange | 1.85164107 | 0.00283975 | 1.75086614 | 0.00808935 |
| Mir208b | skyblue | -3.2040951 | 0.00069322 | -3.105674 | 0.00096254 |
| F630028O10 | orange | 1.17273649 | 0.00084736 | 0.65653555 | 0.07837707 |
| Ankrd63 | skyblue | -4.2088014 | 2.47E-07 | -2.6289037 | 0.00193387 |
| Capns2 | red | -0.8801928 | 0.00449607 | -0.8140902 | 0.01145084 |
| Erich3 | skyblue | -2.0325936 | 5.88E-22 | -0.5801153 | 0.00794399 |
| Dnm3os | orange | 0.78792812 | 0.00622034 | 0.73074673 | 0.01490247 |
| Gm2000 | orange | 1.43237732 | 0.00177199 | 0.19873895 | 0.72574913 |
| Nrarp | red | -0.9509811 | 0.00030486 | -0.6173818 | 0.02473091 |
| Klhdc7a | skyblue | -1.5950533 | 9.95E-11 | -0.7103561 | 0.00597746 |
| Krtap17-1 | skyblue | -1.4334065 | 0.00072832 | -2.0764563 | 5.13E-06 |
| AI593442 | skyblue | -0.4031235 | 0.7535279 | -3.608852 | 0.00752909 |
| AW011738 | orange | 1.01262086 | 1.64E-07 | 1.12425638 | 7.51E-09 |
| Smim1 | red | -0.777641 | 0.00746508 | -0.4507367 | 0.14773482 |
| Gm4294 | orange | 1.71046871 | 0.00581619 | 0.67622216 | 0.34827974 |
| Ppil6 | skyblue | -2.1792473 | 1.82E-08 | -1.2023682 | 0.00234686 |
| Raet1a | yellow | 3.18175043 | 0.00520582 | 4.6218015 | 0.00512302 |
| Abracl | orange | 1.43341346 | 4.15E-18 | 1.04149666 | 9.76E-10 |
| Ankrd65 | skyblue | -2.2905327 | 0.00014787 | -1.7995849 | 0.00337839 |
| Cfap74 | red | -1.440904 | 6.75E-06 | -0.9191766 | 0.00551077 |
| Gm13152 | red | -1.446711 | 0.0023209 | -0.8261885 | 0.09307604 |
| Gm13212 | yellow | 2.45696324 | 4.83E-18 | 1.41114198 | 9.39E-07 |
| Aunip | yellow | 4.08270665 | 6.86E-16 | 2.93084272 | 7.99E-10 |
| Nkain1 | skyblue | -1.5730189 | 1.14E-05 | -1.4810016 | 6.53E-05 |

| | | | | | |
|---|---|---|---|---|---|
| Dcdc2b | skyblue | -1.8125551 | 1.04E-08 | -1.1421934 | 0.00041761 |
| Bnip3 | red | -0.3546753 | 0.12307606 | -0.6239523 | 0.00596699 |
| Cyp4a12b | skyblue | -5.6609396 | 1.15E-11 | -4.165506 | 3.24E-10 |
| Gm4070 | yellow | 2.46333622 | 2.40E-37 | 2.27325882 | 1.20E-32 |
| 1810010H24 | skyblue | -1.8954083 | 1.20E-08 | -1.1584015 | 0.00081627 |
| Trim30c | black | 4.33223604 | 9.64E-16 | 7.19299891 | 1.59E-13 |
| Gm12695 | skyblue | -2.9660305 | 5.33E-05 | -1.8925462 | 0.00856385 |
| Gm11627 | skyblue | -1.980429 | 2.37E-06 | -1.2315454 | 0.00674585 |
| Aoc2 | orange | 0.54305518 | 0.01727317 | 0.70076613 | 0.00233396 |
| Fam174b | skyblue | -1.470575 | 1.54E-13 | -1.1733619 | 8.84E-09 |
| Tmem8b | red | -1.268394 | 8.33E-08 | -0.9431394 | 0.00011694 |
| Slfn1 | black | 5.44564733 | 5.47E-21 | 5.4954827 | 7.69E-21 |
| Evi2a | yellow | 2.63416132 | 3.59E-16 | 2.35426075 | 7.69E-13 |
| Rad54b | yellow | 2.19204607 | 4.46E-12 | 2.83427195 | 4.52E-16 |
| 9530053A07I | skyblue | -1.8801769 | 1.26E-10 | -1.3582615 | 4.81E-06 |
| Gm5150 | yellow | 3.36292494 | 2.15E-08 | 2.63862801 | 6.89E-06 |
| Gm9733 | yellow | 2.9453899 | 1.62E-06 | 4.27864004 | 9.68E-10 |
| Dact3 | red | -0.7638837 | 0.00035013 | -0.3962735 | 0.08559359 |
| Eif5a | orange | 1.08270324 | 1.51E-07 | 0.98509149 | 3.05E-06 |
| Hist3h2a | skyblue | -1.3065461 | 1.73E-05 | -1.270758 | 5.13E-05 |
| Igtp | yellow | 4.08078597 | 2.48E-68 | 4.00681014 | 5.54E-66 |
| Zfp931 | red | -1.5124457 | 0.00012091 | -0.7623128 | 0.06801689 |
| Gm14322 | red | -0.8830291 | 0.00738014 | -1.0045675 | 0.00231722 |
| Gm14420 | red | -0.8221183 | 0.00052341 | -0.7228367 | 0.00318648 |
| Gm14418 | skyblue | -1.8902776 | 2.99E-05 | -1.1885686 | 0.00701185 |
| Gm14305 | red | -1.2142598 | 0.0011979 | -0.5207426 | 0.19566152 |
| Ifi47 | yellow | 3.85284026 | 2.41E-34 | 3.29617708 | 7.96E-26 |
| Tgtp2 | yellow | 3.32652498 | 4.28E-73 | 3.19128259 | 1.41E-73 |
| Tgtp1 | black | 5.07803318 | 2.55E-62 | 5.12003124 | 2.04E-63 |
| Cdc20b | yellow | 1.84421087 | 0.00010843 | 3.76018558 | 1.93E-11 |
| Naip6 | yellow | 2.1388459 | 1.15E-09 | 1.96100819 | 4.09E-08 |
| Naip2 | yellow | 2.2312572 | 6.15E-09 | 2.09733316 | 8.27E-08 |
| Arhgap8 | orange | 0.88229128 | 8.07E-10 | 1.02264434 | 1.27E-12 |
| Hsbp1l1 | skyblue | -0.5263829 | 0.41385285 | -1.9502872 | 0.00224125 |
| Ces1b | red | -1.1808238 | 0.00528786 | -1.0060506 | 0.02218714 |
| Sec61g | orange | 0.81882065 | 4.42E-05 | 0.81824557 | 6.70E-05 |
| Zfp429 | orange | 1.11347973 | 1.74E-06 | 0.69653207 | 0.00452887 |
| Zfp456 | orange | 1.42781461 | 0.00328776 | 0.76965824 | 0.15183825 |
| Serpina3m | black | 5.0615289 | 2.36E-10 | 4.84055759 | 2.07E-09 |
| Serpina3i | black | 4.95895008 | 8.00E-34 | 4.63393091 | 1.17E-32 |
| Ifi27l2a | yellow | 2.70162387 | 2.32E-40 | 2.81332593 | 1.63E-43 |
| Ly6c1 | yellow | 2.47788084 | 4.74E-81 | 2.11210822 | 4.88E-59 |
| Slc45a4 | red | -0.5952172 | 2.15E-05 | -0.6244817 | 1.15E-05 |

| | | | | | |
|---|---|---|---|---|---|
| Nrp | orange | 0.71627315 | 4.77E-05 | 0.61517325 | 0.00071645 |
| Gm14010 | black | 2.90358422 | 0.0003692 | 5.72188557 | 7.84E-05 |
| Slc8a3 | skyblue | -2.1246067 | 1.17E-06 | -1.7391039 | 0.00013659 |
| Capn3 | red | -1.1531409 | 0.00282901 | -0.2421315 | 0.59792707 |
| Atxn7l1os1 | yellow | 2.83429948 | 0.00078984 | 1.51410863 | 0.06959608 |
| Cd209g | skyblue | -2.9282437 | 0.00090834 | -3.1997099 | 0.00159781 |
| Zan | red | -1.3126658 | 0.00174315 | 0.14909297 | 0.79041872 |
| Gzmc | black | 8.92288893 | 3.42E-12 | 7.92758571 | 1.33E-09 |
| Psme2 | yellow | 2.21530149 | 7.57E-24 | 1.99364094 | 2.40E-19 |
| Zfp664 | red | -0.7091103 | 2.00E-08 | -0.5203749 | 6.93E-05 |
| Gm9531 | orange | 1.57200957 | 0.00879543 | 0.69095212 | 0.30542529 |
| Ccr5 | black | 5.09257404 | 2.59E-43 | 4.92371857 | 2.34E-40 |
| Ccdc13 | skyblue | -1.9567844 | 3.18E-11 | -0.8158601 | 0.00553648 |
| Xirp1 | red | -1.387131 | 4.11E-07 | -0.8241095 | 0.00422925 |
| Gm5930 | skyblue | -4.398783 | 0.00059627 | -2.6039532 | 0.01510943 |
| Tmem233 | skyblue | -2.9491867 | 1.39E-09 | -1.9394111 | 0.00014994 |
| Clec7a | yellow | 2.22503598 | 6.96E-08 | 1.98732236 | 2.52E-06 |
| Klrb1b | orange | 1.26567089 | 1.71E-05 | 0.79544499 | 0.0106863 |
| Gm20661 | red | -1.075104 | 0.00020629 | -0.0172045 | 0.96177047 |
| Ifit1bl1 | yellow | 3.88218993 | 3.99E-40 | 3.49267837 | 4.44E-33 |
| Ackr4 | skyblue | -3.8259719 | 3.61E-12 | -2.2657375 | 5.65E-05 |
| Gm43302 | black | 4.36957892 | 0.0004326 | 4.56663227 | 0.00031775 |
| Gbp4 | yellow | 3.53970084 | 2.47E-53 | 3.44409607 | 1.60E-50 |
| Gm11110 | orange | 1.31013617 | 0.0064974 | 1.57718523 | 0.00204486 |
| Ms4a6c | yellow | 4.30588994 | 2.97E-23 | 4.05810464 | 1.46E-20 |
| Arpc4 | orange | 0.9930092 | 4.72E-08 | 0.75465754 | 6.02E-05 |
| Neu2 | skyblue | -3.4214999 | 8.35E-14 | -1.9878606 | 2.10E-05 |
| Kcnj13 | skyblue | -2.1802132 | 0.00017283 | -1.3797583 | 0.01520635 |
| St6galnac4 | orange | 0.62993476 | 0.00041375 | 0.74047181 | 4.38E-05 |
| Cldn34c1 | skyblue | -1.3565953 | 0.00173129 | -1.2637969 | 0.004739 |
| Gm7609 | yellow | 2.52568609 | 0.00278801 | 2.49543267 | 0.00125425 |
| Pin4 | orange | 0.80577082 | 1.11E-05 | 0.35040935 | 0.07946515 |
| Phyhd1 | orange | 1.06044529 | 0.00066998 | 0.47973853 | 0.15033939 |
| H2-T10 | yellow | 2.40826981 | 3.63E-42 | 2.31452674 | 9.03E-39 |
| 1700101E01l | skyblue | -2.4098951 | 2.45E-20 | -1.3015475 | 3.60E-07 |
| Gm11131 | orange | 1.01290796 | 1.17E-07 | 0.5161756 | 0.01220442 |
| Apoo | red | -0.5998255 | 0.00218258 | -0.3747728 | 0.0754232 |
| Tmsb10 | orange | 1.85877902 | 2.24E-17 | 1.6767836 | 3.84E-14 |
| Gm5640 | skyblue | -2.6758111 | 0.00415584 | -3.1245451 | 0.00777011 |
| H2-DMb1 | orange | 1.7575314 | 4.81E-16 | 1.50322214 | 9.07E-12 |
| Kifc1 | yellow | 3.02025938 | 4.49E-26 | 3.37759205 | 6.05E-31 |
| Haus3 | orange | 0.92821571 | 1.26E-05 | 0.95030597 | 1.29E-05 |
| Gm684 | red | -1.1164376 | 0.0004885 | -0.3964662 | 0.26039204 |

| | | | | | |
|---|---|---|---|---|---|
| Pglyrp2 | yellow | 2.6658106 | 3.92E-24 | 2.65275944 | 4.70E-23 |
| Tmem182 | skyblue | -1.2137131 | 0.00027071 | -1.4631455 | 1.74E-05 |
| Gm5483 | yellow | 4.10996895 | 0.00013175 | 3.37476444 | 0.00106998 |
| Muc4 | orange | 0.63666848 | 0.04496183 | 1.30238021 | 2.65E-05 |
| Tm4sf19 | black | 4.88392485 | 0.00022124 | 5.342237 | 0.00032114 |
| Gm17193 | orange | 1.73196597 | 0.00275606 | 0.6393053 | 0.2854249 |
| Gm17396 | red | -0.7005851 | 0.02266184 | -0.8277486 | 0.00858253 |
| 2610203C22 | red | -1.5321276 | 1.90E-05 | -0.3904344 | 0.30702039 |
| Ulbp1 | orange | 1.62743516 | 0.00947739 | 1.74623717 | 0.01258099 |
| LOC1026380 | yellow | 1.41261622 | 0.04907082 | 2.92180494 | 0.00020686 |
| Xlr4a | yellow | 2.42033573 | 4.35E-10 | 2.52057078 | 3.90E-10 |
| Klra18 | yellow | 2.24727062 | 1.03E-08 | 2.60060768 | 2.41E-10 |
| Klra1 | yellow | 3.3779297 | 0.00053076 | 1.64128951 | 0.0679176 |
| Gm15846 | orange | 1.24791653 | 0.11299382 | 1.96926439 | 0.00959342 |
| Brms1 | orange | 0.79949727 | 1.10E-08 | 0.57622308 | 7.20E-05 |
| B230307C23 | orange | 0.86270967 | 0.0071463 | 1.266909 | 0.00014097 |
| C920021L13I | red | -1.2866181 | 1.21E-09 | -0.9396065 | 1.46E-05 |
| Rpl38-ps2 | orange | 0.95354033 | 0.00172819 | 0.71715198 | 0.02823287 |
| Zfp36l1-ps | orange | 2.66444598 | 0.00416551 | 0.08077954 | 0.93022563 |
| Hist2h3c2 | skyblue | -3.7111107 | 2.40E-06 | -1.0573387 | 0.15094647 |
| Gm11400 | orange | 1.24504505 | 0.0002721 | 1.29008826 | 0.00016175 |
| Bambi-ps1 | yellow | 2.23426543 | 4.11E-07 | 2.88387197 | 3.85E-09 |
| Gm14760 | yellow | 0.80010035 | 0.38760603 | 3.46332412 | 0.00367316 |
| LOC1052442 | orange | 1.25918691 | 6.39E-06 | 0.19372445 | 0.56379583 |
| Gm15821 | orange | 1.68421082 | 7.50E-09 | 1.74692258 | 1.07E-09 |
| Slc48a1 | red | -0.6783269 | 2.16E-10 | -0.5158788 | 2.83E-06 |
| Pira1 | yellow | 3.4375982 | 1.96E-11 | 4.29008596 | 3.53E-15 |
| Fzd10 | red | -0.3602581 | 0.35537382 | -1.1716589 | 0.00284006 |
| Pira6 | orange | 1.72995419 | 0.00124072 | 1.90999234 | 0.00051781 |
| Gm12216 | orange | 0.34576138 | 0.04394768 | 0.60384215 | 0.00039464 |
| Snrpert | orange | 1.28100053 | 1.72E-05 | 1.03394821 | 0.00084908 |
| Gm11585 | orange | 1.07832184 | 6.40E-05 | 0.79645842 | 0.00520319 |
| Dnmt3c | orange | 1.86192457 | 0.01212084 | 2.00859834 | 0.00937673 |
| Gm13655 | skyblue | -1.7717681 | 0.0097492 | -1.4133652 | 0.05030923 |
| Gm16464 | yellow | 2.40675867 | 0.0002377 | 3.18772651 | 6.45E-06 |
| Gm14276 | orange | 1.42225634 | 0.00373879 | 0.90419004 | 0.08420677 |
| Gm12250 | black | 5.31687783 | 1.34E-69 | 5.39716724 | 1.23E-71 |
| Gm8822 | orange | 1.44472521 | 9.53E-05 | 1.61725998 | 6.38E-05 |
| Gm11425 | black | 6.53988327 | 2.67E-05 | 5.62489892 | 0.00055125 |
| Gm13204 | orange | 0.68860972 | 0.14332411 | 1.36232279 | 0.00287927 |
| Gm15056 | black | 7.5893047 | 3.50E-08 | 6.48398053 | 4.64E-06 |
| Gm15382 | skyblue | -2.0927211 | 4.88E-05 | -1.1784208 | 0.02434402 |
| Gm11427 | yellow | 2.26088923 | 4.06E-09 | 2.73209048 | 1.28E-11 |

| | | | | | |
|---|---|---|---|---|---|
| Gm15575 | skyblue | -1.2894519 | 0.15006605 | -3.0929761 | 0.00542791 |
| Ctsf | skyblue | -1.5201427 | 1.21E-05 | -1.5577953 | 1.12E-05 |
| Gm13688 | skyblue | -3.4666248 | 0.00413776 | 0.37343302 | 0.74028924 |
| Gm16016 | red | -0.4781158 | 0.17464819 | -1.0935375 | 0.00151223 |
| Gm13340 | red | -0.6854112 | 0.00136761 | -0.5362828 | 0.01519941 |
| Gm11771 | skyblue | -2.2995078 | 0.00226069 | -1.190463 | 0.13502123 |
| Zfp133-ps | skyblue | -0.9592628 | 0.18800945 | -2.615513 | 0.00359195 |
| Gm13436 | orange | 0.88507235 | 0.00012291 | 0.54399946 | 0.0302225 |
| Ccnb2-ps | red | -1.3427415 | 0.00203644 | -0.7946404 | 0.08304693 |
| Hmgb1-ps5 | black | 4.71061114 | 1.17E-10 | 3.74253312 | 4.47E-08 |
| Gm15712 | skyblue | -2.4232599 | 0.00069666 | -2.1394877 | 0.00329615 |
| Gm15696 | orange | 2.3981115 | 0.00043279 | 1.25547823 | 0.06516309 |
| Gm11478 | orange | 1.15096297 | 0.00087454 | 0.61927329 | 0.10024249 |
| Rpl3-ps1 | orange | 0.61763966 | 0.00062523 | 0.23000535 | 0.25899394 |
| Znf41-ps | orange | 1.3230285 | 0.00127569 | 1.17576128 | 0.0067863 |
| Gm22107 | orange | 1.8558781 | 0.00675846 | 0.62620783 | 0.37511766 |
| Gm15472 | black | 5.71687578 | 0.00055933 | 5.69304759 | 0.00084908 |
| Gm2568 | skyblue | -0.5095887 | 0.48733441 | -2.0836247 | 0.00357935 |
| Mir142hg | yellow | 2.05553386 | 1.06E-21 | 2.89485403 | 3.41E-35 |
| Gm14321 | orange | 1.47664158 | 1.62E-05 | 0.63138997 | 0.06041801 |
| Gm15232 | yellow | 2.91243336 | 0.00014762 | 2.13765803 | 0.01827974 |
| AI847159 | yellow | 1.5522543 | 0.02320974 | 2.91273629 | 0.00074802 |
| Gm14097 | red | -1.7158235 | 0.0070713 | -0.3015791 | 0.70846165 |
| A730011C13 | orange | 1.14178682 | 0.00062588 | 0.88439683 | 0.0100529 |
| Ccdc85c | red | -0.6321533 | 4.81E-07 | -0.324602 | 0.0145661 |
| Gm281 | skyblue | -2.8981604 | 2.24E-11 | -1.3918455 | 0.00125378 |
| Foxo6os | skyblue | -4.6071346 | 1.48E-09 | -3.5378749 | 2.84E-07 |
| Gm830 | skyblue | -2.1551367 | 0.0002249 | -2.9913823 | 3.88E-06 |
| B430010I23F | skyblue | -2.8612398 | 1.60E-10 | -2.2542877 | 8.18E-07 |
| Gm15503 | yellow | 3.06189254 | 0.00119399 | 1.89474349 | 0.04920528 |
| Gm16267 | yellow | 2.0192045 | 0.01235483 | 3.93467585 | 0.00014448 |
| Gm14011 | orange | 1.92004951 | 0.00619737 | -1.2613127 | 0.11841076 |
| Gm13111 | skyblue | -3.1090905 | 5.66E-09 | -1.822524 | 0.0007893 |
| Rptoros | skyblue | -1.4109391 | 0.00027078 | -1.401953 | 0.0003913 |
| 4931413K12I | orange | 1.09977175 | 0.00073263 | 0.92974981 | 0.00798778 |
| Mir22hg | orange | 1.17284187 | 1.78E-15 | 0.94670299 | 3.06E-10 |
| 1110018N20 | red | -1.7326061 | 0.00588421 | -0.3905737 | 0.59206404 |
| Snhg15 | orange | 1.54263376 | 1.05E-06 | 1.58688471 | 1.02E-06 |
| Gm10785 | orange | 1.11719358 | 0.00753898 | 0.41389644 | 0.36866398 |
| D830026I12F | skyblue | -1.2304535 | 7.01E-05 | -1.4000802 | 9.00E-06 |
| AI838599 | skyblue | -1.7541343 | 8.50E-06 | -1.6076856 | 7.07E-05 |
| Gm12576 | skyblue | -1.9395206 | 0.00027252 | -0.6123689 | 0.28597331 |
| Gm16151 | skyblue | -2.6053405 | 1.85E-05 | -1.3361665 | 0.02257463 |

| | | | | | |
|---|---|---|---|---|---|
| Gm15663 | skyblue | -1.6084238 | 1.01E-06 | -1.0745517 | 0.00215956 |
| 4930430E12I | black | 7.77118346 | 4.78E-14 | 6.44338355 | 9.15E-17 |
| Gm15964 | orange | 1.71065756 | 0.00358449 | 0.44766261 | 0.4954307 |
| Gm16188 | skyblue | -3.8190577 | 0.00050313 | -1.3617078 | 0.07925375 |
| Snhg17 | orange | 0.59774069 | 0.00296139 | 0.60429274 | 0.00337699 |
| Gm16150 | skyblue | -1.2732713 | 0.01327516 | -1.4915995 | 0.00454262 |
| Gm13056 | red | -1.1533814 | 0.0033239 | -0.6715223 | 0.12497183 |
| Foxd2os | orange | 1.199822 | 3.93E-05 | 1.59577612 | 6.70E-08 |
| Gm14319 | yellow | 2.18718602 | 0.00810932 | 1.56614556 | 0.06130152 |
| Gm11734 | skyblue | -2.4107637 | 5.56E-05 | -1.1975927 | 0.03050578 |
| 1110046J04F | red | -0.9713561 | 0.00047811 | -0.8502055 | 0.00376368 |
| Gm15347 | orange | 1.62957536 | 0.00260161 | 1.61893222 | 0.00257944 |
| 4933423P22I | orange | 1.7454326 | 0.00789379 | 0.89771434 | 0.19035268 |
| Gm11772 | orange | 1.25115006 | 0.00025657 | 1.06820797 | 0.00314116 |
| Gm11738 | black | 2.19347397 | 0.08078437 | 5.3566846 | 0.00116075 |
| Bcas3os2 | skyblue | -2.797159 | 1.31E-05 | -1.7621853 | 0.00335032 |
| Gm12963 | red | -1.2272046 | 0.00376526 | -1.0141197 | 0.0229979 |
| Gm13166 | yellow | 3.18042521 | 0.00123761 | 0.86151025 | 0.4781091 |
| Slc36a3os | black | 4.72403803 | 8.37E-08 | 4.28677664 | 3.65E-06 |
| 4930412C18 | skyblue | -2.0231465 | 1.04E-05 | -1.65426 | 0.00037152 |
| Gm15721 | skyblue | -4.4483875 | 0.00420018 | -2.8447087 | 0.01706197 |
| A730017L22F | red | -0.7883604 | 0.00166387 | -0.3900891 | 0.14098687 |
| Dhx58os | yellow | 1.77948519 | 8.26E-05 | 4.37585634 | 2.09E-11 |
| Gm15410 | skyblue | -3.5946859 | 0.00128285 | -2.0888707 | 0.03143013 |
| 4930469K13I | yellow | 4.06464393 | 0.00372862 | 2.65406236 | 0.01664094 |
| Mecomos | skyblue | -0.7353561 | 0.23216186 | -1.792853 | 0.00355323 |
| Gm16046 | red | -0.9959343 | 0.00051144 | -0.6011684 | 0.05005661 |
| D630024D03 | skyblue | -1.6446434 | 4.52E-07 | -0.7938878 | 0.02001059 |
| Gm15987 | orange | 0.91227035 | 0.06233224 | 1.91847049 | 0.00015121 |
| Lin52 | red | -0.5804758 | 0.00560596 | -0.6943409 | 0.00112768 |
| Gm15912 | skyblue | -1.360045 | 0.0001164 | -1.2297197 | 0.00087947 |
| 2610507I01F | skyblue | -1.7355474 | 2.37E-25 | -1.4504667 | 7.70E-18 |
| Arhgap27os3 | yellow | 3.0839583 | 8.95E-23 | 3.3778626 | 2.88E-25 |
| Gm15832 | yellow | 3.00924281 | 1.42E-21 | 1.96713904 | 1.37E-10 |
| Gm15601 | orange | 1.20846633 | 5.34E-05 | 1.34014081 | 7.01E-06 |
| 2610307P16I | red | -1.5868514 | 4.06E-08 | -0.4244122 | 0.16757126 |
| Cd63-ps | orange | 0.77777236 | 0.02455327 | 1.11165544 | 0.00155714 |
| 2310014F06I | red | -1.3128599 | 0.00025377 | -1.1045166 | 0.00368398 |
| 4930481B07I | red | 0.04193767 | 0.95229299 | -1.6289722 | 0.00990294 |
| Syna | red | -1.0953685 | 0.00577922 | -0.1657865 | 0.72675792 |
| Gm15904 | skyblue | -0.7809691 | 0.35856729 | -2.4922226 | 0.00276692 |
| Gm15411 | red | -1.3922756 | 0.00046662 | -0.7281441 | 0.07396877 |
| Gm13816 | skyblue | -2.6431169 | 6.72E-05 | -2.2303066 | 0.00064558 |

| | | | | | |
|---|---|---|---|---|---|
| Gm5970 | black | 5.7576616 | 1.69E-38 | 6.78449009 | 3.78E-41 |
| Gm16731 | yellow | 3.68427121 | 0.00401995 | 1.76005882 | 0.11576336 |
| B230206L02I | skyblue | -2.0624104 | 0.00046901 | -0.0859462 | 0.90327775 |
| Gm13293 | orange | 1.85477033 | 1.18E-05 | 0.72000249 | 0.09990326 |
| Wipf3 | red | -0.3713062 | 0.06312143 | -0.6274287 | 0.00139968 |
| Gm12473 | red | -0.8525851 | 0.00109432 | -0.7224597 | 0.00775989 |
| Unc45bos | red | -1.102576 | 0.00098873 | -0.2981845 | 0.42587748 |
| Gm11636 | skyblue | -3.1044928 | 0.00074187 | -1.0793148 | 0.23481863 |
| Gm15326 | orange | 1.05843922 | 0.07947797 | 1.81241212 | 0.00227861 |
| Gm2415 | red | -1.291158 | 5.42E-08 | -0.7981488 | 0.00124285 |
| 9030622O22 | skyblue | -1.6990145 | 7.65E-07 | -1.0065724 | 0.00263627 |
| Bach2os | yellow | 2.88446702 | 0.00040781 | 2.57418406 | 0.00283329 |
| Ccpg1os | skyblue | -1.630807 | 2.73E-12 | -1.058208 | 9.80E-06 |
| A330040F15I | yellow | 2.38802773 | 7.10E-06 | 2.00127887 | 0.00017631 |
| Gm7846 | orange | 1.80222188 | 0.00743852 | 0.97940647 | 0.16833706 |
| Tmcc3os | skyblue | -2.6115673 | 0.00327965 | -0.6750335 | 0.49982382 |
| 2810433D01 | skyblue | -2.2381549 | 0.0001698 | -0.5443186 | 0.42090669 |
| Snhg12 | orange | 1.07241673 | 3.58E-08 | 0.87466089 | 1.30E-05 |
| Gm12840 | skyblue | -2.9491291 | 0.00188125 | -1.0735186 | 0.23991752 |
| Gm15998 | skyblue | -1.6648579 | 0.00799745 | -0.6643796 | 0.32723344 |
| Gm16144 | orange | 0.79370365 | 0.15229502 | 1.5174568 | 0.00674059 |
| Prkag2os1 | red | -0.6562008 | 0.11089864 | -1.1263528 | 0.00953426 |
| Gm11707 | yellow | 2.78346355 | 0.00042643 | 1.98206919 | 0.01519938 |
| Adap2os | orange | 1.2861541 | 0.00040656 | 1.05519329 | 0.00527732 |
| 9130208D14 | yellow | 3.24960877 | 9.46E-07 | 2.4201025 | 7.59E-05 |
| Erich2os | skyblue | -2.3144606 | 0.0046003 | -1.4251639 | 0.08823704 |
| Gm15856 | orange | 2.17788403 | 0.0058461 | 1.02875275 | 0.20892996 |
| Cd101 | orange | 1.57126972 | 0.00046055 | 1.71946885 | 0.00019522 |
| Gm2830 | red | -0.7109507 | 0.10471332 | -1.4777537 | 0.00061146 |
| Gm15500 | orange | 0.72530558 | 0.00023377 | 0.44265098 | 0.0351455 |
| Gm12002 | skyblue | -3.5273134 | 0.00520687 | -2.1057046 | 0.13092236 |
| Dnah2os | orange | 1.74406975 | 0.00166573 | 0.64167749 | 0.2944033 |
| Susd5 | skyblue | -1.8757796 | 7.50E-06 | -0.9284523 | 0.02800387 |
| Gm15609 | orange | 0.69781152 | 0.02110268 | 1.27177871 | 2.77E-05 |
| Gm14029 | yellow | 3.48446407 | 1.82E-11 | 4.5780951 | 1.95E-10 |
| Gm15222 | red | -0.8764985 | 0.00049873 | -0.8191309 | 0.00168849 |
| Gm11827 | orange | 0.96171632 | 0.00901244 | -0.6037916 | 0.10171977 |
| 5330413P13I | skyblue | -2.7385333 | 2.89E-07 | -1.2393175 | 0.01232653 |
| Gm15675 | red | -0.8637253 | 0.00047433 | -0.3633932 | 0.1820233 |
| 2410006H16 | orange | 1.02848976 | 5.34E-09 | 0.72382511 | 8.46E-05 |
| E030013I19R | skyblue | -2.9651089 | 1.79E-13 | -2.4631526 | 4.01E-10 |
| A430108G06 | skyblue | -3.6589124 | 2.10E-08 | -0.910953 | 0.0959083 |
| 4930512H18 | yellow | 3.29747283 | 0.00016974 | 2.05308481 | 0.00333115 |

| | | | | | |
|---|---|---|---|---|---|
| Gm15883 | skyblue | -1.9372662 | 5.83E-06 | -0.9560108 | 0.03349515 |
| A230072C01 | red | -1.2883335 | 4.17E-07 | -1.0247697 | 0.00010373 |
| 4930429F24l | red | -1.3167089 | 5.20E-07 | -0.746028 | 0.00769887 |
| Gm13835 | orange | 1.08321586 | 0.00154217 | 0.62321618 | 0.09449884 |
| 4930481A15l | skyblue | -1.5310286 | 2.30E-07 | -1.1813085 | 7.08E-05 |
| Gm12596 | yellow | 3.07534622 | 0.04125803 | 4.34022497 | 0.00909523 |
| 4933431E20l | skyblue | -1.8359459 | 0.00144293 | -1.0505176 | 0.08717408 |
| 4930443O20 | skyblue | -0.9541861 | 0.31779786 | -4.2202203 | 0.00162753 |
| Gm15475 | skyblue | -2.3442749 | 2.22E-08 | -1.480783 | 0.00029471 |
| Gm13889 | orange | 1.91223618 | 2.82E-09 | 1.69066395 | 2.70E-07 |
| 2610027K06l | red | -0.8799791 | 0.00022267 | -0.9937755 | 3.80E-05 |
| Gm13387 | orange | 0.62203914 | 0.07402882 | 1.45757671 | 2.09E-05 |
| Gm11611 | skyblue | -1.800262 | 3.81E-05 | -0.5950144 | 0.2071815 |
| Gm8093 | yellow | 2.90798235 | 0.0014231 | 3.11573513 | 0.00295861 |
| Eldr | orange | 2.01257572 | 3.86E-05 | 1.21502512 | 0.01944938 |
| AI662270 | yellow | 2.9904211 | 3.08E-14 | 2.75218214 | 3.80E-11 |
| Gm11714 | yellow | 2.75215198 | 0.000194 | 2.29428521 | 0.00490076 |
| Gm16316 | red | -0.4942163 | 0.44215165 | -1.4749065 | 0.00774184 |
| Plcxd2 | orange | 1.61254473 | 2.07E-15 | 1.34511405 | 8.96E-11 |
| A830082K12l | skyblue | -1.7596787 | 1.15E-06 | -1.8480639 | 2.52E-07 |
| 2010001A14l | red | -0.7299584 | 0.00891796 | -1.1558468 | 6.89E-05 |
| Gm15133 | yellow | 2.64252548 | 2.48E-07 | 3.3521009 | 4.57E-08 |
| D230022J07l | skyblue | -2.2100399 | 0.00111521 | -0.0318115 | 0.97213252 |
| A230056P14l | skyblue | -1.8851037 | 6.71E-06 | -1.2869036 | 0.00278619 |
| 2810408I11R | yellow | 3.04125699 | 0.00735867 | 2.19519174 | 0.01402834 |
| Gm11377 | skyblue | -1.0380155 | 0.00356081 | -1.4263663 | 0.00043567 |
| E230016M11 | skyblue | -1.9902888 | 0.00101436 | -0.6079319 | 0.36343285 |
| Gm15990 | yellow | 2.37593283 | 0.00025075 | 3.08592 | 6.46E-05 |
| 2610035D17 | red | -1.2796219 | 0.00013201 | -0.9274149 | 0.00789192 |
| Gm15726 | yellow | 1.60714564 | 0.01718787 | 2.67702759 | 3.86E-05 |
| Gm16015 | orange | 0.90860494 | 0.00820335 | 1.58189804 | 2.17E-05 |
| 4933424M12 | orange | 2.46542802 | 0.00372694 | 0.32872813 | 0.75594527 |
| Gm13556 | yellow | 2.78828494 | 0.00327624 | 1.79728517 | 0.06070727 |
| C430049B03 | skyblue | -1.6260547 | 3.36E-09 | -0.9849194 | 0.00084672 |
| BC065397 | red | -1.1333619 | 0.00665796 | -1.2583363 | 0.00353478 |
| Kantr | red | -0.6044049 | 0.00177329 | -0.7806672 | 7.46E-05 |
| Cers1 | skyblue | -1.4287312 | 0.00250198 | -1.6914246 | 0.00069667 |
| 2310065F04l | skyblue | -3.0251501 | 2.51E-05 | -1.6845686 | 0.02800387 |
| 4930577N17 | orange | 1.75167288 | 0.00379644 | -0.0691076 | 0.92171684 |
| Ppp1r18os | orange | 1.85972555 | 0.00019371 | 2.01911 | 8.37E-05 |
| Gm14286 | orange | 0.56262519 | 0.02967973 | 0.86845275 | 0.00095206 |
| Gm13822 | black | 5.64519459 | 5.03E-05 | 3.83685937 | 3.30E-05 |
| 2900089D17 | red | -1.0128393 | 0.00215234 | -0.8837255 | 0.00993985 |

| | | | | | |
|---|---|---|---|---|---|
| Gm16091 | orange | 1.41996917 | 7.11E-09 | 1.09315765 | 9.89E-06 |
| 9830144P21I | orange | 1.31177347 | 4.33E-07 | 0.9223887 | 0.00028589 |
| Zmiz1os1 | orange | 0.92912372 | 0.00771333 | 0.13239767 | 0.76820904 |
| Gm16576 | red | -0.6948549 | 0.0033358 | -0.6004568 | 0.01501586 |
| Hectd2os | skyblue | -3.109504 | 0.00189863 | -1.3925641 | 0.17198126 |
| Epb41l4aos | orange | 0.73570771 | 0.00214322 | 0.20038514 | 0.46994219 |
| Gm16174 | yellow | 2.34565724 | 4.47E-06 | 2.16889007 | 1.59E-05 |
| Zfp111 | red | -0.7016471 | 2.48E-05 | -0.4693384 | 0.00739246 |
| 4930458D05 | red | -0.5559448 | 0.39053593 | -1.5706156 | 0.00936527 |
| 1500009L16F | orange | 1.25754731 | 0.00021808 | 1.0415103 | 0.00322899 |
| Hotairm1 | red | -0.9745769 | 0.0030076 | -1.0455984 | 0.00178994 |
| 1200007C13 | yellow | 3.27955048 | 4.00E-05 | 3.39831991 | 1.36E-05 |
| Gm15283 | yellow | 1.82441352 | 0.04988321 | 2.95539561 | 0.00130839 |
| Gm26280 | orange | 0.1774961 | 0.80981358 | 1.57684062 | 0.00940046 |
| Gm22079 | yellow | 2.60139885 | 0.00049247 | 2.39465049 | 0.00062587 |
| 1700058P15I | orange | 0.84700815 | 0.01186374 | 1.22493352 | 0.00026475 |
| Tnfsf13 | red | -1.0186733 | 4.01E-05 | -0.7579228 | 0.00323536 |
| Gm4477 | skyblue | -1.6673878 | 6.23E-05 | -1.8027894 | 3.34E-05 |
| Cbx6 | red | -0.7233445 | 1.98E-06 | -0.7266813 | 2.83E-06 |
| Mir17hg | orange | 0.98041849 | 6.61E-07 | 0.6210083 | 0.00307834 |
| Klra8 | orange | 1.79736762 | 7.64E-06 | 1.56550306 | 0.00016462 |
| Tgfbr3l | red | -1.0620875 | 0.00012069 | -0.4012536 | 0.17424625 |
| Ier5l | orange | 0.98212841 | 0.00177008 | 0.85901362 | 0.00823899 |
| Slc5a3 | red | -1.2313266 | 1.87E-12 | -0.9683649 | 1.45E-07 |
| Gm15684 | yellow | 2.13168607 | 0.0069049 | 1.98409313 | 0.01490247 |
| Gm16136 | red | -0.88923 | 0.00067608 | -0.8103153 | 0.00265129 |
| Shkbp1 | orange | 1.05879363 | 1.68E-12 | 0.77263862 | 6.09E-07 |
| Zfp882 | skyblue | -1.6026645 | 1.57E-06 | -1.2803534 | 0.00019111 |
| 9230117E06I | skyblue | -4.9044018 | 4.13E-08 | -2.9155979 | 1.75E-05 |
| Etohd2 | orange | 0.8025171 | 1.43E-05 | 0.46731422 | 0.01670614 |
| Gm16184 | orange | 1.8125506 | 0.00141301 | 1.45396073 | 0.01096232 |
| Bcl2a1b | yellow | 3.56841476 | 1.81E-14 | 3.0012903 | 3.90E-10 |
| Gm8459 | black | 5.65624123 | 9.45E-05 | 5.03543057 | 0.00083838 |
| Pira2 | yellow | 2.50448421 | 0.00019112 | 2.89773131 | 2.55E-05 |
| Gm15446 | yellow | 0.00643434 | 0.98671399 | 5.80891792 | 1.21E-21 |
| Gm16578 | black | 3.30731876 | 0.0324857 | 5.04166163 | 0.00267716 |
| A430072P03I | yellow | 1.78178557 | 0.0453821 | 3.02528116 | 0.00363331 |
| Galnt6os | yellow | 2.26585158 | 0.01128229 | 3.07321275 | 0.00076635 |
| LOC1026343 | skyblue | -1.8635986 | 5.12E-06 | -1.2196555 | 0.0045069 |
| Srpx | red | -1.0604784 | 6.94E-08 | -0.5506616 | 0.00728294 |
| AI480526 | red | -0.8161793 | 1.97E-07 | -0.6256794 | 0.00010815 |
| Alms1-ps2 | skyblue | -3.0990125 | 0.00020996 | -2.192246 | 0.01672506 |
| Ttbk2 | red | -0.624286 | 0.00318259 | -0.6305342 | 0.00366278 |

| | | | | | |
|---|---|---|---|---|---|
| Nhlrc4 | skyblue | -2.5829074 | 1.82E-10 | -1.3003198 | 0.00122584 |
| Ugt1a7c | orange | 1.19752899 | 6.08E-05 | 0.7896647 | 0.01379254 |
| Pou3f1 | yellow | 2.0298679 | 0.001272 | 2.02340158 | 0.00206965 |
| Uba52 | orange | 1.24154495 | 0.00104497 | 1.78303706 | 3.69E-06 |
| Gm614 | yellow | 2.9579276 | 2.94E-05 | 1.96749625 | 0.00323895 |
| Ugt1a6b | skyblue | -1.1332607 | 0.13046927 | -2.1939451 | 0.00436579 |
| Acad11 | red | -1.3527059 | 4.05E-09 | -1.0732921 | 5.46E-06 |
| Fbxw10 | yellow | 1.80787808 | 1.28E-05 | 2.22544679 | 2.66E-07 |
| Gm16556 | black | 5.92796915 | 0.00021628 | 2.90435654 | 0.02491181 |
| Tepp | skyblue | -2.0675943 | 0.00026392 | -2.6276945 | 5.86E-06 |
| Itga10 | red | -0.8959111 | 0.00023821 | -0.8807356 | 0.00041434 |
| Tmem189 | orange | 0.86588838 | 9.97E-09 | 0.65870318 | 2.46E-05 |
| Trim34b | orange | 1.97120719 | 0.00048614 | 1.08943186 | 0.05994608 |
| Cfb | yellow | 2.79412326 | 2.53E-09 | 2.82901492 | 3.68E-09 |
| Car15 | skyblue | -1.5232212 | 0.00944699 | -1.2662098 | 0.02992078 |
| Gm17017 | yellow | 2.17924394 | 4.55E-09 | 2.24740828 | 8.22E-09 |
| Alms1-ps1 | skyblue | -3.1876925 | 5.28E-05 | -1.7161234 | 0.01972635 |
| Gm4524 | skyblue | -2.7419871 | 2.93E-05 | -2.3088509 | 0.00060538 |
| Mndal | orange | 1.93924921 | 8.72E-39 | 1.84620849 | 7.44E-35 |
| Gm17296 | red | -0.4465901 | 0.01493726 | -0.6336954 | 0.00057041 |
| Lrrc10b | skyblue | -2.1381429 | 5.12E-13 | -1.0821902 | 0.00040669 |
| 1700071M16 | yellow | 4.0430522 | 3.75E-06 | 3.86230829 | 2.75E-05 |
| 2310050C09 | skyblue | -2.746961 | 0.00051079 | -0.9680077 | 0.25564871 |
| 4932443I19R | skyblue | -2.2720932 | 1.23E-06 | -1.180444 | 0.01639787 |
| Gm8229 | skyblue | -4.3040544 | 0.00137125 | -2.4171954 | 0.02641095 |
| Gm38399 | orange | 1.10533418 | 0.0006805 | 1.17317872 | 0.00050004 |
| Sfta2 | red | -1.3257443 | 0.00010139 | -0.7896545 | 0.02828334 |
| Rps11-ps1 | orange | 0.41420879 | 0.58925082 | 2.0210186 | 0.00936307 |
| Snrpe | orange | 1.09056167 | 5.79E-12 | 0.89013299 | 5.54E-08 |
| Olfr753-ps1 | black | 4.97695272 | 0.00051875 | 4.30209127 | 0.00114443 |
| A430057M04 | yellow | 0.8649555 | 0.28263289 | 2.60706374 | 0.00245148 |
| Tex9 | orange | 0.41936888 | 0.05624976 | 0.67387541 | 0.00182277 |
| Olfr456 | skyblue | -2.4761347 | 0.00058896 | -2.3939731 | 0.00154658 |
| Zfp712 | red | -0.7347748 | 0.02807011 | -0.8840976 | 0.00977621 |
| Gm17182 | skyblue | -2.1753252 | 0.00213894 | -0.3292747 | 0.68630782 |
| Gad1-ps | orange | 2.290435 | 0.00218176 | 1.29741169 | 0.09735954 |
| Gm340 | orange | 2.091754 | 2.05E-10 | 1.4090071 | 2.44E-05 |
| Gm17082 | orange | 0.50016463 | 0.09996348 | 0.93667788 | 0.00396443 |
| Apold1 | orange | 1.3332619 | 3.28E-07 | 0.90133905 | 0.0007018 |
| Gm17173 | yellow | 3.17508611 | 0.0236428 | 4.19457744 | 0.00748543 |
| Gm17201 | orange | 0.59097503 | 0.00130936 | 0.67183982 | 0.00045134 |
| Klhl33 | skyblue | -2.1366729 | 9.41E-06 | -1.5732697 | 0.00176878 |
| Trbj2-6 | yellow | 1.58330474 | 0.01865955 | 3.38628677 | 6.49E-06 |

| | | | | | |
|---|---|---|---|---|---|
| Rps13 | orange | 0.75396214 | 1.07E-10 | 0.54061102 | 7.83E-06 |
| Gm17112 | red | -0.8013758 | 0.00718933 | -0.1745864 | 0.6158187 |
| Hspa1b | skyblue | -2.8052625 | 1.28E-06 | -1.8599668 | 0.00206965 |
| Gm6904 | yellow | 4.19126654 | 6.50E-09 | 3.25325743 | 3.67E-06 |
| Pabpc4l | skyblue | -1.5459221 | 0.00792178 | -2.8926878 | 5.30E-05 |
| F830016B08I | black | 4.57461353 | 3.08E-37 | 5.67009975 | 4.57E-39 |
| Ccdc71l | orange | 0.58809881 | 0.00247309 | 0.38233008 | 0.06444527 |
| Phf11c | yellow | 3.09241693 | 2.99E-34 | 2.8076818 | 8.12E-29 |
| Gm17334 | orange | 2.0730583 | 3.80E-15 | 1.72345806 | 2.44E-11 |
| Smim13 | red | -0.6261104 | 1.20E-05 | -0.3983287 | 0.00829384 |
| Col6a5 | skyblue | -2.160279 | 2.38E-05 | -1.1043287 | 0.03771612 |
| Ak9 | skyblue | -1.8914699 | 2.09E-07 | -1.4975733 | 5.81E-05 |
| Vmn2r97 | yellow | 2.76302568 | 5.89E-11 | 2.15690497 | 3.35E-07 |
| Mpc1-ps | red | -0.32626 | 0.18157118 | -0.7007148 | 0.00341687 |
| Zfp422-ps | red | -1.2408265 | 8.98E-05 | -1.1166767 | 0.00150002 |
| Tma7 | orange | 0.68719101 | 3.31E-07 | 0.58337362 | 2.62E-05 |
| Gm6548 | orange | 1.46080418 | 5.68E-23 | 1.44687843 | 1.88E-22 |
| 2010016I18R | yellow | 1.9759169 | 6.99E-10 | 2.04073663 | 3.00E-10 |
| Lsm5 | orange | 0.96511834 | 1.82E-06 | 0.62693042 | 0.00368655 |
| Phf11b | black | 4.52113562 | 2.42E-47 | 4.37502389 | 6.01E-44 |
| Apol11a | black | 3.48039755 | 0.01010945 | 6.06123796 | 0.00038787 |
| Vmn2r96 | yellow | 3.74424124 | 0.0029843 | 3.37271587 | 0.00610392 |
| Apol11b | orange | 2.20065737 | 2.40E-06 | 1.28768574 | 0.00857129 |
| H2-Q2 | orange | 0.36230294 | 0.29093112 | 1.39064774 | 1.35E-05 |
| Sec14l5 | skyblue | -2.43659 | 6.23E-07 | -1.8701399 | 0.00020385 |
| Gm17098 | skyblue | -2.2913971 | 0.00022412 | -2.5819998 | 3.00E-05 |
| Zfp964 | skyblue | -1.4339805 | 1.48E-06 | -1.9367644 | 2.32E-10 |
| Tnnc1 | skyblue | -0.9581105 | 0.18894541 | -1.8728697 | 0.0085729 |
| Gm9844 | orange | 1.8041248 | 9.50E-07 | 1.38680465 | 0.00042921 |
| Rps2-ps10 | orange | 1.31882085 | 7.60E-05 | 1.46491733 | 3.66E-05 |
| Hspa1a | skyblue | -2.3429103 | 6.42E-05 | -1.6801991 | 0.00590621 |
| BC049352 | yellow | 2.18457755 | 0.03754549 | 3.60179721 | 0.0041242 |
| Gbp11 | black | 5.2950623 | 6.53E-117 | 5.33194265 | 1.88E-121 |
| Dynlt1a | orange | 1.32724553 | 0.00022555 | 1.19542289 | 0.00094253 |
| Tnxa | skyblue | -2.4155091 | 0.00086543 | -1.2751833 | 0.09058412 |
| 1110038B12I | orange | 1.69009313 | 5.67E-14 | 1.50296047 | 3.67E-11 |
| Gm19345 | black | 4.6456369 | 3.24E-07 | 3.80362897 | 3.66E-07 |
| Gm20519 | yellow | 0.66328824 | 0.60629976 | 3.46845616 | 0.00736785 |
| Neat1 | red | -0.6173848 | 1.10E-05 | -0.3224738 | 0.0310342 |
| Gm19684 | orange | 2.02920347 | 3.39E-06 | 1.7003005 | 0.00015541 |
| BB365896 | skyblue | -2.3764963 | 0.00055555 | -2.4742269 | 0.0004034 |
| Platr17 | yellow | 2.44457665 | 0.00685358 | 2.73752434 | 0.00535967 |
| C130080G10 | skyblue | -2.8068054 | 0.00041713 | -1.8007221 | 0.02570487 |

| | | | | | |
|---|---|---|---|---|---|
| Vhl-ps1 | black | 5.44395709 | 0.00015864 | 5.26973666 | 0.000491 |
| 2610524H06 | orange | 1.68232274 | 1.53E-07 | 1.2836338 | 0.00012494 |
| Gm20394 | yellow | 2.26870977 | 2.56E-05 | 2.42292072 | 2.85E-05 |
| Art2a-ps | yellow | 2.80338587 | 0.00022348 | 3.10698892 | 0.00050854 |
| Nlrp1c-ps | yellow | 3.29079694 | 1.09E-07 | 4.10854456 | 8.96E-10 |
| Gm20496 | yellow | 3.14072519 | 1.08E-06 | 2.7815003 | 4.05E-07 |
| Btnl6 | black | 5.94486497 | 0.00030608 | 5.84589134 | 0.00057319 |
| Gm22858 | yellow | 0.97655874 | 0.27976058 | 2.64216439 | 0.00771822 |
| Mir5107 | orange | 0.96743735 | 0.00438902 | 1.56772988 | 6.23E-06 |
| 6330562C20 | orange | 1.67250045 | 0.00478633 | 1.0561527 | 0.09775694 |
| Lrch4 | orange | 1.56652512 | 5.41E-07 | 0.82713199 | 0.01142688 |
| Gm20703 | yellow | 2.33109805 | 0.02053594 | 3.94041748 | 0.00686211 |
| Rpl41 | orange | 0.83617053 | 6.03E-07 | 0.50978244 | 0.00371482 |
| Gm20717 | yellow | 3.16053043 | 2.17E-05 | 1.97039341 | 0.00228114 |
| Gm20667 | yellow | 2.66340863 | 0.00446509 | 2.74162477 | 0.00995652 |
| 4931403E22I | skyblue | -1.791349 | 2.37E-06 | -0.8942932 | 0.0203576 |
| Trav4-3 | yellow | 2.90083604 | 0.00233038 | 3.58822211 | 0.00108671 |
| Gm2237 | orange | 2.45256005 | 8.95E-07 | 0.51659799 | 0.30798247 |
| Ighv1-74 | black | 6.44857104 | 0.00327746 | 1.39232929 | 0.544538 |
| Hist1h2ao | yellow | 3.83189091 | 5.44E-13 | 3.73389314 | 1.45E-08 |
| Ccdc42b | skyblue | -2.507536 | 1.04E-05 | -1.2196358 | 0.0297479 |
| Gm3788 | yellow | 2.17420485 | 8.13E-05 | 1.8612529 | 0.0020003 |
| Zfp955a | red | -0.8325129 | 0.00064381 | -0.4401578 | 0.09177737 |
| Trbv12-2 | yellow | 2.84766172 | 2.36E-05 | 4.50997333 | 1.18E-08 |
| Trbd2 | yellow | 4.00652231 | 6.89E-06 | 3.30701544 | 4.49E-05 |
| Cecr6 | skyblue | -2.6008042 | 1.74E-07 | -1.1864361 | 0.0204239 |
| Gal3st2 | orange | 1.24588451 | 0.02053817 | 1.41729134 | 0.00708575 |
| Trav7-4 | black | 4.01898102 | 0.00376874 | 4.72421915 | 0.00084927 |
| Hist1h2ap | yellow | 4.03181066 | 2.77E-06 | 3.34465421 | 0.00025018 |
| Gm14403 | skyblue | -1.9347363 | 2.49E-07 | -1.2256843 | 0.00151286 |
| BC147527 | yellow | 4.02587904 | 4.94E-29 | 3.82497955 | 4.18E-26 |
| Trav3-3 | black | 4.58599377 | 1.65E-09 | 6.01182546 | 5.63E-07 |
| D430019H16 | skyblue | -1.3511816 | 1.41E-10 | -1.1572419 | 9.52E-08 |
| Trav9-2 | black | 3.06078863 | 0.00527813 | 5.36700931 | 0.00057921 |
| Sirpb1b | yellow | 3.14574868 | 8.04E-12 | 3.38248033 | 1.80E-12 |
| AC149090.1 | red | -0.8970999 | 9.57E-08 | -0.4826537 | 0.00677079 |
| Ccdc85b | red | -0.8951043 | 3.93E-05 | -0.7454043 | 0.00094093 |
| Mid1-ps1 | orange | 1.09826503 | 0.0069146 | 1.8540414 | 4.14E-06 |
| Zfp799 | red | -0.7329147 | 0.00464624 | -0.7293753 | 0.00601646 |
| Gm10130 | orange | 1.13422506 | 0.00564503 | 0.98036331 | 0.01885444 |
| Noc2l | orange | 0.87537023 | 1.86E-05 | 0.73955514 | 0.00046388 |
| Ighv5-17 | orange | 2.45079195 | 0.00937356 | -0.2393924 | 0.83795987 |
| Trbv12-1 | yellow | 1.90341545 | 0.01137755 | 3.25178693 | 0.00013253 |

| | | | | | |
|---|---|---|---|---|---|
| Fmo6 | skyblue | -1.9179565 | 0.00062113 | -0.5388411 | 0.35604832 |
| Trav5-4 | black | 4.67058491 | 0.00064268 | 3.01904948 | 0.02156507 |
| Gm21188 | black | 4.63803278 | 6.52E-18 | 4.37565308 | 1.10E-15 |
| 2010005H15 | orange | 1.91155439 | 0.00156153 | 0.60548389 | 0.39137822 |
| Gm8979 | yellow | 2.87077573 | 0.0021359 | 3.67999241 | 0.00042273 |
| Trbd1 | yellow | 3.72222938 | 1.95E-05 | 2.49196675 | 0.00074331 |
| Rnaset2a | yellow | 3.0821374 | 0.02230301 | 3.98533943 | 0.00983266 |
| Trav14-2 | yellow | 3.5758347 | 0.00752777 | 3.08106212 | 0.00326226 |
| Igkv4-53 | yellow | 4.69935478 | 0.00226435 | 1.12251032 | 0.52206968 |
| Sirpb1a | yellow | 4.10180039 | 4.02E-17 | 3.73329263 | 1.51E-14 |
| Nupr1l | skyblue | -1.8010653 | 8.01E-05 | -1.3789434 | 0.0032935 |
| Igkv6-17 | yellow | 3.24930356 | 9.35E-10 | 1.96088753 | 0.00029507 |
| Lce1k | red | -1.5225375 | 0.00010066 | -0.7090091 | 0.09702734 |
| Syne1 | red | -0.6262413 | 2.84E-05 | -0.6843745 | 6.74E-06 |
| Trav4-4-dv10 | yellow | 3.47848282 | 0.00063448 | 4.24265264 | 0.0002455 |
| Gm16505 | skyblue | -2.6584628 | 0.00015283 | -2.0966885 | 0.00201122 |
| Ankrd66 | red | -0.8273319 | 0.00057269 | -0.9202136 | 0.00013044 |
| Dnah7a | skyblue | -2.048143 | 2.67E-07 | -1.0340543 | 0.00831777 |
| Vkorc1 | red | -1.1219552 | 0.0006144 | -0.3025906 | 0.42393416 |
| Kcnj11 | skyblue | -1.1382934 | 0.01001815 | -1.7985192 | 5.37E-05 |
| Cmtm4 | red | -0.9376798 | 1.64E-07 | -0.6252356 | 0.00080168 |
| Smim22 | red | -0.7881758 | 0.00094166 | -0.6569803 | 0.00940042 |
| Ighv1-78 | black | 5.52568545 | 2.22E-07 | 1.71677947 | 0.08763667 |
| Samd11 | yellow | 2.12773313 | 0.01412345 | 4.45352754 | 0.00019317 |
| Trav17 | black | 6.10284181 | 0.00094566 | 6.01428946 | 0.00154658 |
| Igkv12-44 | orange | -0.0379774 | 0.96452357 | 1.93090703 | 0.0073327 |
| Cdkn2d | orange | 0.9526098 | 3.21E-09 | 0.87258734 | 1.10E-07 |
| Trav10 | black | 5.4626051 | 0.00037902 | 4.94637613 | 0.00194824 |
| Trav12-2 | black | 3.48068569 | 9.87E-05 | 7.11139772 | 1.20E-06 |
| Ighv1-63 | black | 8.78675358 | 0.00561858 | 1.81953827 | 0.59653978 |
| Trav9-4 | black | 5.24547215 | 0.00018674 | 4.3466221 | 0.0008286 |
| Psmb9 | yellow | 2.46383585 | 1.95E-71 | 2.28347054 | 2.37E-61 |
| Gm28373 | yellow | 2.76905423 | 2.81E-05 | 2.84653906 | 2.52E-05 |
| Shisa8 | orange | 1.96701589 | 1.38E-08 | 1.86966138 | 1.42E-07 |
| Trav9-1 | black | 5.40926836 | 1.71E-05 | 4.40896033 | 8.92E-05 |
| 2500002B13I | red | -0.2858466 | 0.47930732 | -1.5483355 | 8.80E-05 |
| Gm9530 | orange | 1.04119022 | 0.00654249 | 0.39916461 | 0.37653884 |
| Gdap10 | yellow | 2.66817243 | 2.20E-10 | 2.43541944 | 8.00E-09 |
| E230013L22I | orange | 0.80240717 | 0.00129055 | 0.94709795 | 0.00022498 |
| A230028O05 | black | 4.97062324 | 8.08E-09 | 5.49276272 | 4.17E-09 |
| 4930556M19 | red | -1.3206714 | 8.52E-10 | -0.766217 | 0.0006881 |
| Gm26880 | orange | 2.26603862 | 0.00953071 | 0.00063495 | 0.9993873 |
| Gm17435 | orange | 1.50896656 | 0.00997901 | 0.61809892 | 0.34795021 |

| | | | | | |
|---|---|---|---|---|---|
| 4731419I09F | orange | 1.89854281 | 0.00378717 | 1.12729401 | 0.07836533 |
| 9530082P21I | orange | 0.85476331 | 3.45E-08 | 0.91159043 | 5.92E-09 |
| Pvt1 | orange | 1.41617504 | 0.00012504 | 1.59402831 | 3.55E-05 |
| Gm4262 | red | -1.0642109 | 0.00330517 | -0.5000787 | 0.18729407 |
| Gm26760 | skyblue | -1.780946 | 0.03125441 | -2.3450564 | 0.00839052 |
| Cdiptos | red | -0.9963071 | 0.00011379 | -1.2456774 | 4.43E-06 |
| Gm10425 | orange | 0.47089323 | 0.29322657 | 1.17732491 | 0.00545456 |
| Foxl1 | skyblue | -1.9743282 | 4.37E-06 | -0.7902237 | 0.0768043 |
| Gm9917 | red | -0.8702664 | 1.32E-06 | -0.7467153 | 5.42E-05 |
| 1810034E14I | red | -0.7201852 | 0.00802666 | -0.1973748 | 0.52506375 |
| Gm2885 | skyblue | -1.6134004 | 0.00556512 | -0.6827724 | 0.26529393 |
| Gm26580 | skyblue | -1.8706932 | 0.00938508 | -0.8335962 | 0.2832304 |
| Gm19705 | yellow | 2.18745156 | 6.82E-09 | 1.92958034 | 1.74E-07 |
| B230354K17I | red | -0.5396904 | 0.00568015 | -0.7185321 | 0.0003155 |
| A530020G20 | red | -0.9349212 | 0.00755766 | -0.6806421 | 0.06406441 |
| D230017M19 | skyblue | -1.3365075 | 0.00041812 | -1.3868354 | 0.00044543 |
| Gm26628 | yellow | 1.87915962 | 0.02258834 | 2.43460211 | 0.00358074 |
| 1110002J07F | yellow | 2.58777248 | 0.00861337 | 1.66432263 | 0.11736503 |
| Gm26510 | orange | 1.27672426 | 6.32E-05 | 1.14707513 | 0.00048391 |
| 2310002F09I | skyblue | -1.4712696 | 0.00626803 | -1.8086094 | 0.00376777 |
| 2700038G22 | orange | 1.99392666 | 6.92E-08 | 1.65530258 | 1.04E-05 |
| 4632428C04 | red | -1.0100244 | 0.00273254 | -0.6724622 | 0.06130152 |
| 9330175E14I | yellow | 3.18049304 | 6.18E-85 | 3.34711948 | 1.37E-91 |
| Snhg5 | orange | 0.64951186 | 8.07E-06 | 0.64978688 | 1.43E-05 |
| 1010001N08 | red | -0.4500365 | 0.27578783 | -1.0985862 | 0.00594364 |
| Gm16907 | orange | 1.1373409 | 4.65E-06 | 0.68153382 | 0.00981071 |
| Gm26767 | red | -0.7032027 | 0.00697174 | -0.1669603 | 0.57184442 |
| Gm26684 | skyblue | -2.0557429 | 0.00418257 | -1.5789122 | 0.0322404 |
| 2010300F17I | yellow | 4.13242744 | 4.57E-08 | 3.23612315 | 8.04E-06 |
| Hmgb1-ps8 | yellow | 2.41907816 | 3.14E-08 | 1.57119642 | 0.0005533 |
| Gm17249 | red | -0.3741863 | 0.29573491 | -1.5650691 | 2.32E-05 |
| Gm10516 | red | -0.9567916 | 0.00063283 | -0.9695003 | 0.00090016 |
| 2700012I20F | skyblue | -2.4576129 | 0.00025347 | -0.6498122 | 0.27372082 |
| Fendrr | red | -1.0519929 | 5.94E-05 | -0.7164462 | 0.00894341 |
| Gm26671 | red | -1.0640163 | 0.00045759 | -0.2573041 | 0.4295167 |
| C920009B18 | yellow | 3.56307547 | 6.75E-13 | 2.715986 | 1.38E-08 |
| C030034L19I | skyblue | -1.5057689 | 2.86E-05 | -1.3534815 | 0.00025721 |
| Gm26876 | yellow | 3.82254395 | 0.00744994 | 3.34254931 | 0.01163122 |
| Gm26816 | orange | 1.27228394 | 0.00596062 | 0.88522913 | 0.07881701 |
| Gm26814 | skyblue | -2.2762642 | 0.00126373 | -1.7431258 | 0.01594547 |
| 9230116N13 | orange | 1.0917284 | 0.00770208 | 0.87784587 | 0.04123156 |
| AU020206 | orange | 1.21005845 | 1.50E-17 | 0.89006798 | 9.83E-10 |
| Mir155hg | yellow | 3.82704043 | 1.52E-13 | 3.95437897 | 3.70E-13 |

| | | | | | |
|---|---|---|---|---|---|
| Gm26522 | orange | 1.78144546 | 7.30E-11 | 1.57408227 | 2.42E-07 |
| Gm10544 | skyblue | -1.8352698 | 0.00899939 | -0.7738982 | 0.29257386 |
| Gm26781 | black | 4.60160595 | 1.99E-11 | 4.68431538 | 1.43E-10 |
| Gm16754 | orange | 0.6281902 | 0.03754443 | 0.85794074 | 0.00498625 |
| Rian | skyblue | -2.0277183 | 1.55E-11 | -1.3423953 | 1.62E-05 |
| 2310031A07I | yellow | 2.13553861 | 0.00057185 | 1.78889439 | 0.0070147 |
| 9530026P05I | red | -1.439703 | 0.00347465 | -1.0704155 | 0.03755352 |
| D430036J16I | skyblue | -2.305565 | 1.86E-09 | -1.9021577 | 5.11E-07 |
| 5830432E09I | orange | 1.38861455 | 7.44E-05 | 1.75433734 | 2.48E-06 |
| Gm26586 | orange | 1.33791774 | 0.00204385 | 1.30572928 | 0.00454729 |
| 9930014A18I | red | -1.021249 | 2.46E-05 | -0.5231523 | 0.03858894 |
| 4933406C10 | skyblue | -1.5847139 | 0.00220558 | -1.5990619 | 0.00253544 |
| 4930516B21 | skyblue | -1.8502972 | 6.78E-08 | -1.4571935 | 3.41E-05 |
| Gm16675 | orange | 1.66667704 | 6.81E-09 | 1.70945998 | 2.49E-09 |
| 2610037D02 | orange | 1.09899435 | 0.00477833 | 0.56844836 | 0.16827121 |
| 2610020C07 | orange | 0.84274581 | 0.12170409 | 1.65307912 | 0.00130175 |
| Gm26902 | yellow | 3.10156819 | 1.23E-07 | 2.06890797 | 0.00028975 |
| D430018E03 | yellow | 1.22480277 | 0.07286705 | 2.72437814 | 0.00020654 |
| Gm26637 | yellow | 2.40178944 | 1.45E-06 | 2.52947038 | 1.83E-06 |
| Gm26797 | orange | 0.97584347 | 0.0091564 | 0.8371326 | 0.02892434 |
| Gm26799 | yellow | 1.93120057 | 0.00021931 | 2.14731208 | 4.91E-05 |
| E230029C05 | yellow | 2.44702214 | 1.74E-13 | 2.6772309 | 1.18E-13 |
| Gm26660 | red | -1.1654167 | 4.21E-06 | -0.887025 | 0.0011287 |
| Gm17322 | black | 3.73237134 | 0.00085476 | 6.76417812 | 9.51E-06 |
| A930012L18F | skyblue | -2.1346394 | 0.00055307 | -2.0918823 | 0.00060172 |
| Gm17597 | skyblue | -5.0445948 | 0.00069806 | -1.8590297 | 0.04282471 |
| 1110019D14 | skyblue | -1.5174245 | 0.000154 | -1.2873913 | 0.0020939 |
| Mirt1 | yellow | 1.9793524 | 1.19E-15 | 2.43268787 | 7.12E-23 |
| A130051J06F | orange | 0.79845166 | 0.07464139 | 1.4901113 | 0.00093709 |
| Mhrt | skyblue | -2.4120992 | 0.06141783 | -3.6929856 | 0.00470638 |
| Gm26764 | orange | 1.56896175 | 0.00626279 | 1.09362783 | 0.07958948 |
| Gm26740 | orange | 1.72647865 | 8.79E-07 | 1.80714717 | 1.94E-06 |
| 2810429I04F | yellow | 1.73389344 | 0.02911464 | 3.66607877 | 0.00012942 |
| 9530036O11 | skyblue | -2.1763719 | 0.00026162 | -0.824518 | 0.19953934 |
| Gm26530 | orange | 0.56259336 | 0.03443613 | 1.15585791 | 1.33E-05 |
| E030044B06I | skyblue | -2.8121261 | 0.00133798 | -3.7391042 | 0.00036614 |
| 2010110K18I | black | 3.02400187 | 0.00012391 | 6.76854074 | 1.11E-06 |
| A730056A06I | skyblue | -4.9549543 | 0.0015894 | -2.397447 | 0.04811746 |
| Snhg4 | orange | 1.67432693 | 7.23E-07 | 1.61508794 | 2.71E-06 |
| 4833407H14 | yellow | 2.23856817 | 2.72E-12 | 2.43006513 | 1.06E-13 |
| 9330136K24I | orange | 1.47379985 | 1.30E-05 | 0.5617539 | 0.11253107 |
| Gm2115 | skyblue | -3.0782733 | 9.96E-09 | -1.9500561 | 0.00029191 |
| Gm17259 | red | -0.7256343 | 0.18546626 | -1.3929843 | 0.00915621 |

| | | | | | |
|---|---|---|---|---|---|
| Gm16685 | yellow | 2.95811972 | 4.65E-12 | 2.5693488 | 7.60E-09 |
| Panct2 | skyblue | -2.7250112 | 1.48E-08 | -1.4135854 | 0.00313842 |
| Gm26809 | orange | 0.87682421 | 0.00055395 | 0.84810241 | 0.00113239 |
| Gm26813 | orange | 1.74315456 | 0.00103367 | 0.95101209 | 0.07988939 |
| E530011L22F | red | -0.7916101 | 0.00178022 | -0.9587442 | 0.0003155 |
| Gm16701 | red | -0.4356474 | 0.17620909 | -1.1166894 | 0.00044375 |
| Gm807 | black | 6.22271954 | 0.00057269 | 4.21901272 | 0.0078887 |
| A930007I19F | orange | 1.79157272 | 3.14E-07 | 1.16058594 | 0.0016403 |
| Gm26603 | yellow | 2.13636971 | 0.00297617 | 2.31281983 | 0.00120538 |
| Gm26703 | orange | 1.73173968 | 0.00012728 | 1.61951035 | 0.00047536 |
| Gm26682 | orange | 1.9866879 | 0.00501423 | 1.07451233 | 0.13359605 |
| Gm26541 | orange | 1.26851922 | 0.00012131 | 1.49788877 | 1.49E-05 |
| 4930547M16 | skyblue | -1.9977466 | 0.00988938 | -0.8678772 | 0.25771685 |
| Gm26711 | orange | 1.52832994 | 0.04503573 | 2.4185947 | 0.00262399 |
| A730020E08I | skyblue | -1.4568942 | 0.00920152 | -1.1766438 | 0.04304778 |
| Gm26917 | orange | 1.69299741 | 3.47E-06 | 1.26667046 | 0.0008225 |
| Ptprv | orange | 1.55968941 | 1.41E-05 | 2.21816637 | 4.22E-09 |
| A930001A20I | skyblue | -3.6247028 | 0.00673768 | -1.8098277 | 0.11590336 |
| Gm26947 | yellow | 2.66562591 | 1.48E-06 | 2.28888673 | 3.87E-05 |
| Gm6981 | red | -1.4185821 | 0.00657305 | -0.3177506 | 0.57184442 |
| 2700099C18 | yellow | 1.99649821 | 7.99E-08 | 2.25993797 | 9.52E-09 |
| Bvht | red | -0.6512627 | 0.00010785 | -0.6062409 | 0.00046335 |
| Bin2 | yellow | 2.73307863 | 8.64E-25 | 2.61705936 | 1.05E-22 |
| Rassf10 | skyblue | -1.9149848 | 4.14E-11 | -1.4521757 | 9.94E-07 |
| Gm26973 | skyblue | -1.7365486 | 1.78E-08 | -0.9018229 | 0.00451932 |
| Gm27010 | orange | 1.12609781 | 4.06E-08 | 0.80839277 | 0.00021774 |
| Snhg6 | orange | 1.18030662 | 1.57E-06 | 1.37231606 | 3.72E-08 |
| Gm27514 | orange | 1.59509981 | 0.06063266 | 2.12260559 | 0.00939572 |
| Lockd | yellow | 2.0296336 | 1.51E-06 | 2.91597246 | 1.74E-08 |
| 5830468F06I | yellow | 1.95391144 | 0.05229819 | 2.90535574 | 0.00497059 |
| Gm28043 | orange | 0.2526991 | 0.34048253 | 0.6580147 | 0.00970662 |
| Mir6377 | skyblue | -1.4126252 | 0.00758429 | -1.5243801 | 0.01026243 |
| Gm27202 | skyblue | -3.0407682 | 0.00176022 | -2.3544466 | 0.01254695 |
| C1rb | orange | 1.7791979 | 0.00065662 | 1.92527142 | 0.00039645 |
| 1500015A07I | orange | 0.88166493 | 7.25E-06 | 0.64489784 | 0.00195526 |
| Gm27252 | orange | 1.50719082 | 2.29E-05 | 0.183353 | 0.65492889 |
| Gm28053 | yellow | 2.5321044 | 0.01176363 | 2.56772723 | 0.00640326 |
| Gm18853 | yellow | 3.43370825 | 4.21E-06 | 2.71762673 | 0.00019566 |
| Gm18852 | yellow | 3.48447123 | 9.35E-09 | 3.94130337 | 7.20E-10 |
| 2810030D12 | skyblue | -1.5333992 | 0.00036409 | -1.0602 | 0.01899367 |
| Ms4a14 | yellow | 3.80657692 | 3.99E-13 | 3.96197629 | 9.23E-13 |
| Gm17767 | orange | 1.19313974 | 0.06740411 | 1.72147412 | 0.00629144 |
| 1700097N02 | black | 3.05731292 | 0.0038024 | 4.93205967 | 0.00076689 |

| | | | | | |
|---|---|---|---|---|---|
| Gm29538 | skyblue | -2.0989112 | 5.98E-05 | -1.6053155 | 0.00209352 |
| Gm28729 | skyblue | -2.4354297 | 3.10E-18 | -1.4542712 | 3.24E-08 |
| Gm18301 | black | 2.46783008 | 0.02137824 | 5.38067966 | 0.00025916 |
| 2310034G01 | red | -0.9466745 | 0.00011802 | -0.69794 | 0.00567311 |
| Zfp383 | red | -0.9716798 | 7.51E-11 | -1.1421238 | 4.18E-14 |
| BE692007 | yellow | 2.96618897 | 5.57E-23 | 2.82055131 | 1.87E-20 |
| Gm28884 | yellow | 4.46538787 | 0.00224114 | 2.04792064 | 0.08583137 |
| Gm5960 | black | 7.22940411 | 2.09E-06 | 6.93794066 | 8.82E-06 |
| Gm7160 | orange | 0.89158237 | 1.17E-05 | 0.92783949 | 6.42E-06 |
| Gm6652 | skyblue | -2.9268562 | 1.72E-06 | -1.8284113 | 0.00080067 |
| Rbm3-ps | yellow | 2.82847281 | 2.44E-11 | 2.4897863 | 5.94E-09 |
| 2810013P06I | red | -0.5822856 | 0.00011605 | -0.43845 | 0.00550859 |
| Bcl2a1d | yellow | 3.287741 | 1.34E-17 | 2.62104736 | 2.75E-11 |
| B130024G19 | red | -0.9100946 | 2.37E-07 | -0.9100711 | 6.22E-07 |
| 1700025N23 | yellow | 3.72762258 | 5.80E-08 | 4.03331807 | 1.02E-07 |
| Gm28439 | red | -0.8771905 | 0.05657707 | -1.267518 | 0.00774529 |
| Gm19585 | yellow | 3.91003302 | 3.77E-18 | 4.06368916 | 1.28E-15 |
| 2610306M01 | red | -0.6709821 | 0.0038982 | -0.5027081 | 0.04483809 |
| 1810006J02F | yellow | 2.62525551 | 0.00584318 | 1.70854102 | 0.0316208 |
| Gm28557 | skyblue | -2.3115849 | 2.03E-06 | -0.460684 | 0.35985836 |
| 4930422M22 | skyblue | -1.792468 | 0.00695641 | -1.1976885 | 0.06327165 |
| 2310008N11 | skyblue | -2.5425674 | 1.51E-05 | -2.2775872 | 0.00023114 |
| Gm28935 | yellow | 2.08803569 | 0.06720707 | 4.98696307 | 0.00068918 |
| 2310020H05 | skyblue | -1.7963298 | 0.00241959 | -1.17163 | 0.09763628 |
| C530043A13 | skyblue | -3.3978588 | 1.62E-07 | -0.7070267 | 0.17718228 |
| Gm19261 | black | 3.80415779 | 0.00697519 | 5.52720948 | 0.00043023 |
| Gm29585 | yellow | 2.37960404 | 2.74E-11 | 1.97134121 | 2.96E-09 |
| Gm5511 | yellow | 3.72681075 | 0.00111807 | 1.08255777 | 0.30194464 |
| F730311O21 | orange | 0.83655334 | 0.05951345 | 1.39643912 | 0.00172214 |
| Gm28523 | skyblue | -1.0233979 | 0.00771528 | -1.499037 | 0.00019847 |
| Gm28874 | orange | 1.7673554 | 0.00908482 | 0.55153045 | 0.47830553 |
| Lhb | skyblue | -2.2131663 | 4.69E-17 | -1.0890099 | 2.38E-05 |
| Gm28437 | red | -0.435669 | 0.09285364 | -0.7563767 | 0.00381158 |
| Eif4a-ps4 | orange | 1.15070145 | 5.21E-05 | 0.82678548 | 0.00566942 |
| Gm28720 | skyblue | -2.7566501 | 4.14E-06 | -2.2273524 | 0.0002648 |
| 4930568A12I | yellow | 2.91410037 | 1.03E-06 | 2.57676716 | 3.52E-05 |
| Dnah7c | skyblue | -1.9832141 | 1.72E-11 | -1.8157541 | 8.49E-11 |
| Ms4a4a | yellow | 4.45081868 | 8.03E-35 | 3.77934853 | 5.72E-26 |
| Ace3 | red | -1.1490255 | 2.66E-05 | -1.1995393 | 2.10E-05 |
| Kcnq1ot1 | red | -0.5446583 | 0.02952908 | -0.7661976 | 0.00241651 |
| 2310040G24 | red | -1.1019251 | 0.00875554 | -0.5819078 | 0.22036921 |
| Gm19461 | red | -1.2446653 | 0.00351905 | -0.9543893 | 0.0303791 |
| Gm29237 | red | -1.2961568 | 0.00013676 | -0.8303021 | 0.02005918 |

| | | | | | |
|---|---|---|---|---|---|
| 9130401M01 | orange | 1.15826641 | 0.00024875 | 0.55749322 | 0.09883937 |
| Gm28438 | red | -0.905983 | 0.00579719 | -0.8387359 | 0.01348904 |
| Bcl2a1a | yellow | 2.81083339 | 1.49E-09 | 2.22782146 | 3.26E-06 |
| I830127L07R | black | 5.26794994 | 8.51E-15 | 5.12090897 | 1.64E-14 |
| Gm20257 | orange | 0.93739874 | 9.70E-07 | 0.57324826 | 0.00376744 |
| Gm37472 | orange | 0.82768865 | 0.00506705 | 0.47558229 | 0.14421519 |
| Gm37468 | black | 5.0340901 | 0.00063809 | 4.83259308 | 0.00038314 |
| Gm33280 | skyblue | -2.5686163 | 7.09E-05 | -1.4723656 | 0.02433686 |
| Gm37718 | yellow | 2.48642241 | 0.00640934 | 2.08036431 | 0.06488985 |
| Snrpn | skyblue | -1.1044539 | 5.19E-05 | -1.6095449 | 1.04E-08 |
| Gm7357 | skyblue | -2.0673733 | 0.00076723 | -1.8877632 | 0.00347849 |
| Gm37626 | skyblue | -0.8401043 | 0.19106327 | -2.5574043 | 0.00016329 |
| Gm19721 | orange | 0.57148386 | 0.3472391 | 2.03917111 | 0.00071118 |
| Sh2d1b1 | orange | 0.77481335 | 0.03938638 | 1.01695019 | 0.00873157 |
| Gm37464 | skyblue | -2.7073623 | 8.21E-07 | -1.2248362 | 0.02383018 |
| Pcdhga12 | red | -1.1052934 | 1.29E-06 | -0.6768151 | 0.00472462 |
| Pcdhga9 | red | -1.5378545 | 2.07E-07 | -0.2984359 | 0.33926395 |
| Gm38197 | skyblue | -1.8717307 | 3.97E-06 | -1.8083556 | 2.11E-05 |
| A930004J17F | orange | 1.48527822 | 4.41E-09 | 1.03227594 | 0.00011409 |
| Gm37289 | yellow | 2.74597351 | 0.00809747 | 0.97647824 | 0.33576029 |
| Pcdhac2 | skyblue | -2.4264737 | 1.13E-08 | -1.7901757 | 3.73E-05 |
| A630081D01 | yellow | 2.52543087 | 0.00149796 | 1.34615013 | 0.11318525 |
| Gm7694 | red | -0.6492668 | 0.00313457 | -0.6767079 | 0.00260761 |
| Naaladl2 | red | -1.1141506 | 0.00117256 | -1.2762983 | 0.00037344 |
| Gm38253 | yellow | 1.81391041 | 0.03655723 | 2.93662215 | 0.00065701 |
| Gm37795 | orange | 1.19726051 | 8.89E-07 | 1.10233246 | 9.16E-06 |
| Pcdhgc3 | red | -0.9970994 | 1.10E-07 | -0.6053641 | 0.00201002 |
| 9430034N14 | orange | 1.32277005 | 0.0069977 | 1.65943119 | 0.00138757 |
| Pcdhgb6 | red | -1.008495 | 0.00025532 | -0.7263396 | 0.01125906 |
| Gm32200 | skyblue | -1.61181 | 1.74E-06 | -1.3824399 | 7.68E-05 |
| Gm37387 | yellow | 2.34995713 | 5.67E-05 | 2.19373751 | 0.00022108 |
| Gm37303 | yellow | 0.6050093 | 0.50878569 | 2.92031585 | 0.00139347 |
| Gm37169 | yellow | 2.67269533 | 4.05E-05 | 2.59316863 | 6.02E-05 |
| Gm37090 | red | -0.6796081 | 0.00049566 | -0.5925512 | 0.00372595 |
| Gm37728 | skyblue | -1.4259908 | 0.00251074 | -1.3402771 | 0.00838495 |
| Lce1d | red | -1.293508 | 8.26E-05 | -0.8121514 | 0.01945891 |
| Pcdhac1 | skyblue | -3.6628274 | 0.00573158 | -3.636747 | 0.00052424 |
| BC037039 | red | -1.420517 | 0.00306091 | -0.259915 | 0.65344149 |
| Gm38157 | orange | 1.09574133 | 0.00056186 | 0.26422058 | 0.5028267 |
| Gm38158 | yellow | 2.59471773 | 8.53E-11 | 2.02724043 | 5.32E-07 |
| Gm37581 | orange | 0.4002334 | 0.32502602 | 1.10364403 | 0.0030223 |
| Lsmem2 | red | -1.254998 | 0.00256393 | -0.9627125 | 0.0341215 |
| Gm37666 | yellow | 2.44916965 | 5.77E-11 | 2.25944835 | 4.62E-09 |

| | | | | | |
|---|---|---|---|---|---|
| Pcdhga5 | red | -1.0312835 | 9.59E-05 | -0.3787337 | 0.16751091 |
| Gm37352 | orange | 0.57013815 | 0.06863092 | 1.09394849 | 0.00033397 |
| Gm37652 | skyblue | -3.6029524 | 0.0004517 | -0.9713923 | 0.36426265 |
| Gm37859 | skyblue | -1.4751334 | 0.0001516 | -1.3373351 | 0.00081747 |
| Gm36932 | orange | 1.63976488 | 0.00780243 | 1.56176072 | 0.0187285 |
| Gm20056 | orange | 2.16544616 | 0.00164639 | 1.22826883 | 0.08758316 |
| Gm37249 | skyblue | -2.5947037 | 5.59E-05 | -0.7234216 | 0.29251374 |
| Gm37010 | black | 3.40368986 | 0.00051741 | 4.908753 | 0.00138173 |
| Gm37499 | red | -1.4182678 | 0.00835008 | -0.3940497 | 0.51044903 |
| Pcdhgb7 | red | -1.3580991 | 1.83E-07 | -0.7997872 | 0.00303939 |
| Gm38275 | black | 6.57318819 | 2.99E-06 | 3.79852789 | 0.00445008 |
| Gm38103 | skyblue | -2.0114701 | 0.00713599 | -2.1842699 | 0.00998019 |
| Gm38102 | red | -0.8749609 | 0.00038137 | -0.5141364 | 0.04865699 |
| A130071D04 | yellow | 2.51645287 | 4.54E-08 | 2.70625674 | 4.35E-08 |
| Gm37689 | yellow | 4.13400184 | 0.00333518 | 2.82529241 | 0.00664505 |
| Gm37691 | orange | 1.07852889 | 0.00056499 | 0.9708793 | 0.00340972 |
| Gm38244 | orange | 1.21059598 | 0.01686171 | 1.38617885 | 0.00865688 |
| Gm37516 | red | -1.4525424 | 0.00318319 | -1.1115609 | 0.03008553 |
| Gm37033 | orange | 0.69081621 | 0.00124305 | 0.2665997 | 0.28295859 |
| Gm37829 | red | -0.5697886 | 0.00429387 | -0.8493312 | 3.81E-05 |
| A430027H14 | skyblue | -1.8620196 | 0.01660499 | -2.2088157 | 0.00788246 |
| Gm33994 | yellow | 3.84999904 | 0.00042388 | 1.5889245 | 0.1334288 |
| Gm4610 | yellow | 2.63726958 | 4.62E-06 | 3.24549764 | 4.60E-08 |
| Gbp6 | black | 4.6987312 | 1.87E-109 | 4.86287289 | 1.03E-116 |
| Gm43162 | orange | 1.25582076 | 0.00054146 | 0.54501807 | 0.1984893 |
| Gm42462 | yellow | 3.74997623 | 1.77E-06 | 3.55244605 | 6.82E-06 |
| Gm42984 | skyblue | -2.1621422 | 0.00549979 | -0.9857297 | 0.23940501 |
| Gm5547 | orange | 1.45379188 | 5.64E-05 | 1.23159566 | 0.00128075 |
| Trdc | orange | -0.0661768 | 0.93990005 | 2.42966333 | 0.00091315 |
| Gm6560 | orange | 1.63525577 | 3.46E-07 | 1.49381357 | 7.07E-06 |
| 4833413G10 | red | -0.9973828 | 0.00016614 | -1.0818687 | 4.56E-05 |
| Gm43437 | skyblue | -1.9836933 | 4.70E-07 | -1.6655487 | 2.08E-05 |
| Gm43423 | red | -0.3368652 | 0.49822769 | -1.5568449 | 0.00098955 |
| Gbp10 | black | 6.03886169 | 1.28E-72 | 6.26011372 | 3.54E-79 |
| Gm43667 | skyblue | -1.7513217 | 1.00E-06 | -1.5904494 | 2.55E-05 |
| Snord43 | yellow | 2.79017685 | 0.00360839 | 1.873651 | 0.03622773 |
| Gm43668 | red | -1.1490196 | 0.00018765 | -0.7163373 | 0.0338654 |
| Sox2ot | skyblue | -3.7955305 | 2.58E-06 | -0.7963362 | 0.21861804 |
| Gm43071 | orange | 1.1386897 | 0.00067256 | 0.59084018 | 0.0989326 |
| Gm43719 | skyblue | -1.5175351 | 3.15E-07 | -1.5282426 | 8.74E-07 |
| Gm42515 | orange | 1.98403145 | 0.00097826 | -0.0128001 | 0.98874703 |
| Gm43618 | skyblue | -3.4279349 | 7.31E-06 | -4.5364492 | 1.98E-06 |
| Gm43205 | red | -1.5954964 | 0.00527813 | -0.3711441 | 0.57001983 |

| | | | | | |
|---|---|---|---|---|---|
| Gbp5 | yellow | 4.51219819 | 2.46E-127 | 4.21302919 | 7.00E-112 |
| Gm43714 | orange | 1.85553465 | 0.00723641 | 1.56165565 | 0.0232083 |
| Igkv2-109 | orange | 0.03604912 | 0.97148153 | 2.14394989 | 0.00774526 |
| Gm43272 | yellow | 2.31577474 | 0.00059146 | 1.86076576 | 0.0052487 |
| Gm42659 | orange | 1.50502474 | 5.80E-05 | 1.16440003 | 0.00322277 |
| Gm43328 | orange | 0.77640415 | 0.00049897 | 0.96855396 | 2.07E-05 |
| Gm43305 | yellow | 2.57743276 | 7.93E-14 | 2.04497568 | 6.80E-09 |
| Gm42748 | orange | 1.022565 | 0.06915051 | 1.58521958 | 0.00418479 |
| Gm43595 | skyblue | -2.2105175 | 3.66E-07 | -1.5076821 | 0.00069945 |
| Gm43351 | black | 5.63843062 | 0.00019048 | 6.13160804 | 6.72E-05 |
| Gm43006 | orange | 0.91847427 | 0.00392937 | 0.41174052 | 0.28929003 |
| Gm42141 | yellow | 2.92947738 | 0.0010223 | 2.32071576 | 0.01262595 |
| Gm19439 | skyblue | -1.890992 | 0.00021524 | -1.3997137 | 0.00836336 |
| Gm43508 | red | -1.3592769 | 0.00021984 | -0.8675322 | 0.02368544 |
| Gm42517 | skyblue | -2.1775385 | 1.39E-05 | -0.9971751 | 0.05407283 |
| 4932422M17 | red | -0.9086298 | 0.00018273 | -0.5771612 | 0.02379969 |
| Gm42780 | skyblue | -2.812696 | 0.00362357 | -0.9459885 | 0.39268573 |
| AI506816 | orange | 1.75988537 | 3.86E-12 | 1.7634608 | 5.47E-12 |
| Gm8493 | orange | 1.47669053 | 0.02599525 | 2.05192581 | 0.00255831 |
| Gm4332 | orange | 1.74289696 | 0.00062115 | 0.02615654 | 0.96837975 |
| Gm42892 | skyblue | -2.2298609 | 6.42E-05 | -1.0940425 | 0.07085237 |
| Gm30648 | skyblue | -1.9384013 | 0.00058447 | -0.8270661 | 0.16587993 |
| 4930555A03I | skyblue | -1.9299658 | 0.00435306 | -1.0129913 | 0.20547553 |
| Gm42918 | orange | 1.62312058 | 0.00436669 | 1.08993255 | 0.06001162 |
| Gm43464 | yellow | 2.07838166 | 0.0341869 | 3.36418441 | 0.00198095 |
| Gm42658 | orange | 1.13415776 | 0.00044807 | 1.29451896 | 0.00014193 |
| Gm8539 | skyblue | -1.8928508 | 0.00051548 | -0.40958 | 0.4645502 |
| Gm20755 | skyblue | -3.3136013 | 0.00020529 | -2.7698308 | 0.00066261 |
| C130083M11 | red | -0.8531543 | 0.00769049 | -0.3329676 | 0.34082196 |
| Gm40348 | skyblue | -4.0111493 | 0.00807526 | -0.3347805 | 0.80390948 |
| Gm36551 | yellow | 4.53733673 | 0.00119455 | 3.7215462 | 0.00511558 |
| Gm9403 | skyblue | -2.4677834 | 0.0065878 | -0.5928653 | 0.54326182 |
| Gm43181 | yellow | 3.84744773 | 0.00027239 | 3.49587789 | 0.00013152 |
| Gm36266 | red | -0.7860956 | 0.00495398 | -0.4768721 | 0.11164513 |
| Trav7-5 | black | 4.28245978 | 0.00511882 | 6.31728127 | 3.23E-05 |
| Gm20559 | orange | 1.51371124 | 5.06E-31 | 1.50562879 | 1.60E-30 |
| Gm43050 | red | -0.3937177 | 0.07737312 | -0.6332041 | 0.00441273 |
| Gm10441 | skyblue | -4.4087806 | 3.49E-05 | -2.7203139 | 0.00246707 |
| Peg13 | skyblue | -1.3939806 | 1.11E-07 | -1.2134153 | 6.59E-06 |
| Gm43621 | orange | 1.06049286 | 0.00432193 | 0.62360272 | 0.10710292 |
| 5930430L01F | red | -1.0715103 | 0.00164019 | -0.8555822 | 0.01615963 |
| Gm43633 | yellow | 2.06071318 | 0.00201549 | 2.94783285 | 6.60E-05 |
| Gm45495 | skyblue | -1.3467004 | 6.33E-05 | -1.1737173 | 0.00091766 |

| | | | | | |
|---|---|---|---|---|---|
| Gm43068 | black | 6.66654841 | 3.73E-06 | 6.33190991 | 1.91E-05 |
| Gm42726 | orange | 1.57199064 | 0.0022542 | 0.67148035 | 0.23289652 |
| Gm15997 | skyblue | -3.538033 | 0.00454728 | -1.5787388 | 0.12345487 |
| Gm42528 | skyblue | -1.7307407 | 0.00088239 | -1.1922981 | 0.0287851 |
| Gm43496 | skyblue | -2.3580751 | 0.00190625 | -1.6525979 | 0.04161501 |
| Gm43197 | orange | 1.59737465 | 6.47E-05 | 1.60199433 | 9.97E-05 |
| Gm43769 | yellow | 1.55372327 | 0.10517331 | 2.96309447 | 0.00433244 |
| Gm10461 | red | -1.0387531 | 0.00973469 | -0.5725188 | 0.2022919 |
| Mpv17 | red | -0.7677262 | 0.00068017 | -1.0541083 | 3.86E-06 |
| Gm43775 | orange | 1.67548861 | 0.00148228 | 1.72009052 | 0.00197977 |
| Gm19719 | yellow | 2.19664875 | 0.00010757 | 2.07405411 | 0.0002903 |
| Gm42549 | orange | 0.93558375 | 0.12329318 | 1.7350552 | 0.00559896 |
| AI839979 | orange | 1.3333106 | 3.28E-05 | 1.03993288 | 0.00149767 |
| Gstm2-ps1 | skyblue | -1.7907033 | 0.00365024 | -1.5022097 | 0.01524725 |
| Gm43588 | skyblue | -2.2185721 | 0.0032514 | -1.0024319 | 0.24498004 |
| Gm43323 | orange | 1.48744873 | 6.48E-05 | 0.88920967 | 0.03055499 |
| Zfp862-ps | red | -1.1224608 | 4.62E-05 | -0.9869866 | 0.00048127 |
| Gm44165 | black | 6.52247733 | 7.35E-07 | 4.87756516 | 3.90E-08 |
| Gm20560 | red | -1.4353833 | 1.76E-05 | -0.7530148 | 0.02508863 |
| Gm38910 | red | -0.9724603 | 0.00029 | -1.044138 | 0.00015737 |
| Gm44321 | orange | 0.91668137 | 0.09322223 | 1.91728026 | 0.00152571 |
| Gm44013 | yellow | 3.03340393 | 0.00128765 | 0.67855422 | 0.51674024 |
| Gm44401 | orange | 2.25300416 | 0.00223398 | 1.31137971 | 0.06014035 |
| Gm43914 | skyblue | -1.3157558 | 0.00158822 | -1.2310549 | 0.00504859 |
| Gm44000 | skyblue | -2.1172499 | 4.16E-08 | -1.3658749 | 0.00010169 |
| Gm45769 | red | -0.832537 | 0.00882709 | -0.6664829 | 0.04387047 |
| Gm43936 | skyblue | -2.4629524 | 0.00564994 | -2.9371261 | 0.0096533 |
| Gm44250 | red | -1.3605401 | 5.76E-08 | -1.0768737 | 3.14E-05 |
| Gm10388 | yellow | 2.19740299 | 1.86E-07 | 2.11525927 | 1.23E-06 |
| Gm6375 | orange | 1.36673709 | 0.00319159 | 1.12560233 | 0.01948904 |
| Gm44080 | yellow | 3.9999946 | 0.00043061 | -0.1071035 | 0.94031162 |
| Gm38708 | red | -0.982983 | 3.58E-05 | -0.8523315 | 0.00039554 |
| B130021K23I | orange | 0.6617335 | 0.35469316 | 2.32668742 | 0.00631557 |
| Gm43890 | yellow | 3.81672531 | 1.10E-05 | 3.88979785 | 5.45E-05 |
| Gm44423 | orange | 1.4683161 | 3.35E-05 | 1.95001725 | 5.44E-08 |
| Gm43953 | yellow | 0.82848297 | 0.38215113 | 2.81854526 | 0.00783685 |
| Gm44210 | skyblue | -2.7869468 | 6.16E-06 | -2.997447 | 5.01E-06 |
| Gm6637 | yellow | 3.79706183 | 2.92E-09 | 2.86327066 | 3.50E-06 |
| Gm45738 | yellow | 3.04476696 | 0.00532174 | 0.64429235 | 0.54580245 |
| 9930120I10R | orange | 1.15016741 | 0.11293555 | 2.18624299 | 0.00310431 |
| 4930417O13 | yellow | 2.60515124 | 0.00010454 | 3.3353675 | 2.11E-05 |
| Gm44175 | yellow | 1.70124582 | 0.07555434 | 5.42110399 | 6.46E-06 |
| Gm44020 | skyblue | -0.8874021 | 0.1511174 | -2.0459983 | 0.00508591 |

| | | | | | |
|---|---|---|---|---|---|
| 1810059H22 | skyblue | -1.958297 | 0.00322693 | 0.11657226 | 0.88956611 |
| Gm43982 | yellow | 1.60450338 | 0.13942736 | 3.38926067 | 0.00934267 |
| Olfr1372-ps1 | orange | 0.86474306 | 5.61E-05 | 0.7131999 | 0.00136588 |
| Gm43984 | orange | 1.09982158 | 0.00857903 | 1.87453173 | 7.08E-06 |
| Gm44187 | orange | 2.07793456 | 0.0052167 | 1.4295056 | 0.05181368 |
| Gm36640 | skyblue | -3.1256651 | 0.00165608 | -1.0395458 | 0.28410633 |
| Gm44224 | skyblue | -3.9351975 | 0.00623599 | -1.9884424 | 0.07014422 |
| Gm44174 | yellow | 4.00013293 | 6.40E-09 | 3.47734992 | 3.87E-07 |
| Gm44292 | yellow | 2.43679987 | 1.36E-10 | 2.40039988 | 5.59E-10 |
| Gm44663 | orange | 2.03931225 | 0.00415584 | 1.24951973 | 0.12098249 |
| 5430431A17I | red | -0.8348965 | 0.00238374 | -0.6897642 | 0.01753805 |
| Gm42372 | yellow | 2.71504527 | 5.95E-09 | 1.88537024 | 2.85E-05 |
| Gm44509 | skyblue | -2.1511105 | 4.27E-06 | -1.7700251 | 0.00026077 |
| Gm32633 | orange | 0.76278173 | 0.00882694 | 1.18504761 | 5.52E-05 |
| Snhg1 | orange | 1.12618258 | 6.53E-12 | 1.04938052 | 3.37E-10 |
| Gm30928 | skyblue | -2.5649308 | 1.63E-09 | -1.8850896 | 1.18E-05 |
| Gm44851 | yellow | 2.62833916 | 0.00021681 | 1.35918444 | 0.06878107 |
| 4930413G21 | orange | 0.22795486 | 0.3453769 | 0.80121481 | 0.00058329 |
| 4732496C06 | red | -1.0805392 | 0.0001081 | -0.968234 | 0.00075368 |
| Gm33248 | skyblue | -2.0371929 | 0.00196223 | -1.1531256 | 0.09294098 |
| Gm39094 | skyblue | -4.1863806 | 0.0023209 | -0.6106727 | 0.66914185 |
| Gm33989 | red | -0.8866789 | 0.00026368 | -0.8902901 | 0.00034103 |
| Gm34121 | skyblue | -2.5550897 | 0.00789379 | -1.3093933 | 0.17816636 |
| Gm44758 | red | -1.2421156 | 0.00330517 | -0.8358284 | 0.06507124 |
| Gm45226 | skyblue | -1.9920317 | 0.00372021 | -0.8839551 | 0.17682465 |
| 2310043P16I | orange | 2.03753226 | 2.95E-10 | 1.23884716 | 0.0001901 |
| Gm36028 | skyblue | -2.3720535 | 0.00667811 | -0.893392 | 0.27799673 |
| Gm626 | skyblue | -2.6235702 | 1.93E-10 | -1.0749265 | 0.01140684 |
| Gm44861 | black | 2.48381647 | 0.00033699 | 5.69338467 | 7.74E-06 |
| 9130015G15 | skyblue | -2.2314383 | 1.07E-06 | -1.6978025 | 0.00015335 |
| Gm45221 | orange | 1.10457853 | 0.05651485 | 2.41957039 | 5.14E-05 |
| 2210406H18 | yellow | 2.29507143 | 0.04948761 | 3.82697498 | 0.00533133 |
| 4833411C07 | red | -0.5662446 | 0.3331839 | -1.6531097 | 0.00478162 |
| Gm29683 | skyblue | -3.9112275 | 7.22E-07 | -1.4634307 | 0.02701306 |
| Gm32031 | orange | 0.08579031 | 0.85239732 | 1.11928988 | 0.00655159 |
| Gm44829 | orange | 1.61626529 | 0.00335263 | 0.7067598 | 0.25160536 |
| Gm35339 | skyblue | -1.3232854 | 4.06E-05 | -1.2193283 | 0.00023091 |
| Gm44659 | black | 4.91359672 | 0.00075347 | 3.71477812 | 0.00111086 |
| Fam81b | skyblue | -2.5012716 | 8.21E-16 | -1.4431312 | 1.00E-06 |
| Gm44860 | yellow | 3.01564761 | 0.000762 | 3.57925437 | 0.00759381 |
| Gm45220 | yellow | 1.90863465 | 0.00223941 | 2.44997609 | 0.0002925 |
| Dcst2 | orange | 1.25971757 | 0.0063231 | 0.3433018 | 0.53059805 |
| 1810010D01 | red | -1.1432688 | 0.00345901 | -1.2679627 | 0.00160852 |

| | | | | | |
|---|---|---|---|---|---|
| A330076H08 | red | -0.9115895 | 0.05230055 | -1.4605503 | 0.00242398 |
| Prmt1 | orange | 0.86716287 | 3.81E-06 | 0.79066227 | 4.14E-05 |
| Gm49396 | orange | 0.6557073 | 0.00077467 | 0.34485075 | 0.1030173 |
| Gm5737 | skyblue | -2.2814653 | 0.00055501 | -2.1735604 | 0.00143926 |
| A230057D06 | red | -1.2158206 | 0.00080514 | -0.8804705 | 0.02049839 |
| A930037H05 | orange | 2.00989284 | 2.60E-07 | 1.86255797 | 2.09E-06 |
| Gm45223 | yellow | 2.24180999 | 2.20E-05 | 1.72051763 | 0.00125425 |
| Gm44710 | red | -0.7763367 | 0.00143756 | -1.1536177 | 4.66E-06 |
| Gm45222 | yellow | 1.83196864 | 4.76E-09 | 2.10247792 | 9.13E-12 |
| Nup62 | orange | 0.91609703 | 1.19E-12 | 0.76659752 | 6.19E-09 |
| Gm44667 | orange | 1.47771482 | 0.00810446 | 1.75043263 | 0.00288339 |
| Muc16 | skyblue | -1.078832 | 0.00258539 | -1.3149246 | 0.00032933 |
| Cfap99 | skyblue | -2.0201598 | 1.62E-05 | -1.0685609 | 0.02360609 |
| Gm31105 | skyblue | -1.370315 | 0.00213794 | -1.2355096 | 0.00571051 |
| Gm45470 | skyblue | -2.8508735 | 5.46E-06 | -2.5607946 | 7.13E-05 |
| Gm36869 | yellow | 2.57557495 | 0.00140705 | 4.01264261 | 0.0003428 |
| Gm45606 | orange | 1.29793461 | 1.44E-05 | 1.20962359 | 0.00011383 |
| Gm45418 | yellow | 2.8685285 | 0.00070667 | 2.33028992 | 0.00655162 |
| Gm45640 | orange | 1.58171452 | 0.00023257 | 1.27373784 | 0.00477111 |
| Gm45605 | skyblue | -1.4942959 | 1.53E-07 | -1.2366969 | 2.29E-05 |
| Eid3 | orange | 0.88382106 | 0.03011283 | 1.37939803 | 0.00085408 |
| Hspa14 | orange | 0.65847882 | 5.91E-11 | 0.52609645 | 4.26E-07 |
| Gm45819 | skyblue | -2.2663951 | 3.76E-09 | -1.595464 | 4.60E-05 |
| Gm45663 | orange | 1.6052821 | 0.00396248 | 0.22696201 | 0.74037112 |
| Gm20071 | red | -0.6539339 | 0.00268536 | -0.32224 | 0.15372868 |
| 5430430B14I | red | -0.7428141 | 0.01832994 | -1.0568388 | 0.00059779 |
| Gm45353 | skyblue | -2.2317914 | 0.01250806 | -3.0727194 | 0.0055993 |
| 5430437J10F | yellow | 2.72382645 | 1.27E-09 | 1.94768668 | 3.88E-05 |
| Gm45471 | black | 4.16155302 | 8.63E-05 | 4.89999214 | 0.00011715 |
| Gm45564 | orange | 0.64396921 | 0.31336196 | 2.20892397 | 0.00059237 |
| 5830408C22 | orange | 1.21959591 | 7.58E-05 | 0.30609378 | 0.36921797 |
| Igip | red | -1.166909 | 8.86E-11 | -1.0402046 | 1.57E-08 |
| Flt3l | red | -1.1597032 | 4.17E-06 | -0.8634347 | 0.00096801 |
| Gm20219 | red | -0.9154911 | 0.00012908 | -0.6000539 | 0.01628575 |
| Gm20100 | yellow | 3.51803025 | 1.94E-05 | 2.63510879 | 0.00134443 |
| E330018M18 | skyblue | -2.6399343 | 0.00267726 | -1.8606511 | 0.02070819 |
| Gm45552 | orange | 0.8151143 | 0.01061233 | 0.88311806 | 0.00633521 |
| 4930412F12I | orange | 2.21167287 | 0.00808725 | 1.12527493 | 0.1979188 |
| Gm19935 | red | -1.0157633 | 0.00010255 | -0.3763088 | 0.18998851 |
| Gm45716 | orange | 1.44063572 | 5.81E-07 | 0.90123104 | 0.00226055 |
| Gm33543 | skyblue | -2.3162263 | 0.04712835 | -3.2878886 | 0.00640911 |
| Gm45332 | skyblue | -1.9514929 | 0.0006537 | -1.2397057 | 0.04230157 |
| Gm42031 | yellow | 3.07047317 | 1.48E-14 | 3.08061097 | 1.74E-14 |

| | | | | | |
|---|---|---|---|---|---|
| Gm45869 | skyblue | -2.9092788 | 1.02E-05 | -1.3429482 | 0.02468113 |
| 4933406B17I | skyblue | -2.4904573 | 2.85E-07 | -1.4007149 | 0.00297141 |
| Gm45752 | yellow | 2.09255787 | 0.00079181 | 2.51522141 | 0.00012316 |
| Gm45712 | black | 4.93737733 | 0.00114109 | 3.89255341 | 0.00159019 |
| Gm4316 | yellow | 1.81597225 | 0.00012093 | 2.52961469 | 1.11E-06 |
| Gm45774 | yellow | 2.59649572 | 0.00068537 | 3.50088232 | 4.84E-06 |
| Gm45854 | yellow | 1.63650938 | 0.03593266 | 2.94959548 | 0.00074427 |
| Gm39822 | orange | 1.0017291 | 6.69E-05 | 0.75267774 | 0.00387027 |
| Gm42047 | yellow | 2.12761999 | 3.07E-07 | 3.49745703 | 1.68E-17 |
| A530010L16R | yellow | 2.09938166 | 1.62E-07 | 2.05594233 | 8.08E-07 |
| Gm31718 | yellow | 3.90808771 | 3.73E-15 | 2.39447166 | 1.27E-07 |
| Gm45767 | red | -0.9602192 | 0.00758845 | -0.8658616 | 0.01715237 |
| Gm33104 | yellow | 2.50736397 | 1.97E-06 | 1.12682672 | 0.03160294 |
| D030034A15 | skyblue | -2.158877 | 0.0039659 | -1.8203403 | 0.01487096 |
| Gm35028 | black | 2.38531517 | 0.0266447 | 6.05786012 | 0.00012544 |
| Gm20300 | red | -1.0990336 | 3.96E-07 | -0.8062899 | 0.00040719 |
| Gm48065 | yellow | 4.33210197 | 0.00086649 | 2.28176038 | 0.00967129 |
| Gm6545 | black | 5.41804889 | 1.62E-27 | 4.97184154 | 1.99E-23 |
| AC160562.1 | yellow | 1.8583942 | 0.00797633 | 2.27687933 | 0.00274953 |
| Gm39323 | yellow | 3.8574184 | 2.74E-13 | 4.52419455 | 5.06E-15 |
| Gm48717 | skyblue | -1.4632099 | 0.00794271 | -1.148171 | 0.05094527 |
| E030022I16R | orange | 1.58975066 | 0.00501983 | 1.79831889 | 0.0027491 |
| Gm47445 | orange | 1.84001462 | 0.00404751 | 0.50828946 | 0.5008398 |
| AC160637.1 | orange | 1.77455025 | 7.48E-12 | 1.60562199 | 1.04E-09 |
| Gm48478 | yellow | 0.8936144 | 0.32900856 | 3.11227607 | 0.00417494 |
| Gm47050 | yellow | 3.87567504 | 3.22E-11 | 2.25543045 | 3.67E-05 |
| 1700019L13R | black | 6.35937137 | 1.14E-09 | 5.41311271 | 1.21E-10 |
| Gm47640 | red | -0.9813011 | 0.0011283 | -0.6489341 | 0.03265177 |
| Gm30373 | yellow | 3.93877652 | 9.13E-06 | 3.7292023 | 1.82E-05 |
| Gm33858 | yellow | 2.75902675 | 0.00267196 | 3.48926497 | 0.00126591 |
| Gm47950 | orange | 1.18090166 | 0.00072494 | 1.08166328 | 0.00212371 |
| 1190001M18 | yellow | 2.4937375 | 0.00070729 | 2.69880601 | 0.00093941 |
| Gm6477 | orange | 0.90521537 | 0.00992148 | 0.81697494 | 0.03254139 |
| AC153498.1 | black | 7.3821923 | 3.51E-07 | 5.99895871 | 6.45E-05 |
| Gm36660 | skyblue | -1.31977 | 0.0005278 | -1.2863242 | 0.00113394 |
| Gm47218 | orange | 1.4917721 | 0.00040513 | 1.70763337 | 2.77E-05 |
| Lilr4b | black | 5.02689019 | 8.49E-56 | 5.12231838 | 7.20E-58 |
| A130077B15I | black | 4.93527651 | 0.00075305 | 3.17348792 | 0.00127092 |
| C230072F16 | skyblue | -2.0411058 | 9.35E-06 | -0.6086631 | 0.20767152 |
| Lilrb4a | yellow | 3.77196945 | 1.58E-27 | 4.04140139 | 2.13E-31 |
| Gm19990 | red | -0.9542786 | 0.00080904 | -0.4036258 | 0.1945056 |
| Ifngas1 | yellow | 2.82251778 | 0.00047537 | 4.1584999 | 3.25E-05 |
| 4930466K18I | skyblue | -1.5077149 | 9.86E-05 | -1.4442838 | 0.00041457 |

| | | | | | |
|---|---|---|---|---|---|
| Sfta3-ps | red | -1.4597417 | 6.12E-05 | -1.0026668 | 0.00845476 |
| Gm48768 | red | -0.8183413 | 0.00677807 | -1.061223 | 0.00047163 |
| Gm47720 | skyblue | -1.8083234 | 0.00055555 | -1.6277915 | 0.00272674 |
| Gm29684 | yellow | 2.36925025 | 0.0007684 | 2.97844872 | 0.00025874 |
| Gm33091 | yellow | 3.80164389 | 9.17E-08 | 2.92530808 | 3.30E-05 |
| Gm35696 | skyblue | -2.6565004 | 6.98E-05 | -0.7692597 | 0.23774188 |
| Gm36283 | skyblue | -1.3459341 | 3.25E-05 | -1.1690254 | 0.00056109 |
| Gm47461 | skyblue | -5.7982979 | 0.00016003 | -3.3852684 | 0.00461519 |
| Gm48882 | skyblue | -1.806205 | 0.00562701 | -1.6474023 | 0.01740642 |
| Gm48835 | skyblue | -2.9959546 | 1.55E-06 | -2.0666848 | 0.0004246 |
| 4933412E12I | orange | 1.03095246 | 1.02E-11 | 0.83808275 | 6.69E-08 |
| Gm4798 | skyblue | -3.47023 | 0.00579537 | -1.0178637 | 0.36074186 |
| Gm47644 | red | -0.5339171 | 0.36350721 | -1.7771639 | 0.00250795 |
| Gm36172 | yellow | 3.42052329 | 0.00757858 | 1.07747156 | 0.33259946 |
| Gm33677 | skyblue | -3.1369004 | 0.00202496 | -4.3406724 | 0.00137808 |
| Gm38407 | yellow | 2.10656581 | 0.00025178 | 1.78048807 | 0.00398995 |
| Gm4739 | yellow | 2.38897493 | 4.10E-05 | 2.15217106 | 0.00173123 |
| Gm46224 | black | 4.37080425 | 1.77E-14 | 4.67711466 | 1.14E-12 |
| Gm47567 | black | 5.84032927 | 0.00023725 | 2.26928366 | 0.08212207 |
| Gm47015 | orange | 1.37852112 | 2.46E-05 | 1.42657582 | 2.16E-05 |
| D430020J02I | yellow | 1.81606957 | 2.23E-06 | 2.10421227 | 8.93E-08 |
| Gm33424 | skyblue | -1.9204816 | 0.0096079 | -1.55796 | 0.06048236 |
| A930040O22 | skyblue | -2.8669748 | 0.00350096 | 0.04797765 | 0.96368171 |
| Gm47139 | skyblue | -1.9045343 | 0.00019285 | -0.538702 | 0.30947578 |
| Gm19951 | yellow | 4.28033155 | 5.39E-21 | 3.82479285 | 9.10E-17 |
| Gm7511 | skyblue | 0.3247071 | 0.71114819 | -3.6811837 | 0.00646963 |
| AC125351.1 | red | -1.090083 | 4.54E-05 | -0.7363885 | 0.00784559 |
| Gm46430 | red | -0.6778895 | 0.00181271 | -0.7822626 | 0.00037517 |
| Gm32219 | skyblue | -1.9053523 | 0.00038499 | -2.0825866 | 0.00059518 |
| Gm47260 | orange | 0.89699722 | 0.00263174 | 0.70456267 | 0.0220235 |
| Gm49329 | skyblue | -0.3360595 | 0.64156089 | -2.159474 | 0.00304524 |
| AC163354.1 | yellow | 2.17366777 | 5.67E-07 | 1.66612789 | 0.00022383 |
| 5033406O09 | orange | 1.35468058 | 0.00578915 | 0.4277492 | 0.43541492 |
| AC121965.1 | orange | 0.65757642 | 0.00019629 | 0.55775321 | 0.00235012 |
| Gm6566 | orange | 1.56414097 | 0.00036751 | 0.53105903 | 0.27072114 |
| Gm40909 | yellow | 1.48418763 | 0.0316604 | 2.75071043 | 5.63E-05 |
| Gm47155 | skyblue | -1.3555022 | 0.00407154 | -1.1854482 | 0.01870128 |
| Gm47705 | yellow | 3.71108629 | 7.69E-07 | 4.02015322 | 3.84E-06 |
| 4930549C15 | skyblue | -1.020442 | 0.18283021 | -2.9119023 | 0.00126509 |
| Gm48662 | skyblue | -2.5627526 | 2.72E-13 | -2.4397828 | 1.51E-11 |
| Gm47662 | yellow | 3.35353997 | 0.00125591 | 2.94629136 | 0.00493009 |
| Gm35279 | yellow | 3.32457579 | 0.0001426 | 3.23335127 | 0.00108436 |
| Gm38655 | skyblue | -3.4144914 | 2.09E-06 | -1.966137 | 0.00590871 |

| | | | | | |
|---|---|---|---|---|---|
| Gm48600 | red | -1.2141706 | 0.00613197 | -1.0155638 | 0.02962795 |
| Gm49027 | skyblue | -1.9294472 | 4.92E-06 | -0.3104483 | 0.49904645 |
| Gm47773 | orange | 1.9273813 | 0.00703369 | -0.5213523 | 0.52097398 |
| Gm31544 | black | 5.90382423 | 3.70E-05 | 4.63853153 | 0.00057046 |
| F630042J09F | skyblue | -1.0647579 | 0.00378483 | -1.3899179 | 0.00044855 |
| Gm48500 | skyblue | -2.0615672 | 0.00169244 | -1.3799356 | 0.03633584 |
| Gm47732 | skyblue | -2.7808424 | 0.00041578 | -1.2997689 | 0.10448706 |
| Gm36161 | yellow | 4.31334215 | 3.36E-15 | 4.12929118 | 4.81E-13 |
| Gm47242 | black | 6.05854811 | 7.43E-14 | 6.56854656 | 3.08E-11 |
| Gm47754 | yellow | 2.92352375 | 0.00075421 | 2.48125529 | 0.00365013 |
| Gm21370 | black | 3.61747483 | 0.00029434 | 5.24837875 | 0.00011108 |
| 4930579J19F | skyblue | -3.0216992 | 1.07E-05 | -2.6507871 | 5.52E-05 |
| Gm49284 | red | -1.2548104 | 1.37E-11 | -1.0055666 | 4.82E-08 |
| G930009F23 | orange | 0.02789552 | 0.95533971 | 1.37143381 | 0.00169162 |
| Gm49041 | orange | 1.93798453 | 5.84E-07 | 1.90316739 | 3.67E-06 |
| Gm34934 | red | -1.3715394 | 9.96E-07 | -1.038268 | 0.00030681 |
| Eef1akmt4 | orange | 1.71818152 | 1.87E-06 | 1.32364021 | 0.00030081 |
| AC174678.1 | orange | 0.83985996 | 0.00528908 | 0.29349637 | 0.38702305 |
| AU022793 | yellow | 2.70773917 | 1.19E-06 | 1.71631554 | 0.00179829 |
| 9930017N22 | orange | 0.9756187 | 0.00846337 | 0.12625005 | 0.78534677 |
| Gm32618 | skyblue | -1.6827222 | 1.04E-09 | -1.7462057 | 1.65E-09 |
| Gm35019 | yellow | 3.29753783 | 0.00031595 | 1.16467435 | 0.19648641 |
| Pnp | yellow | 2.13943279 | 7.24E-27 | 1.81194477 | 2.25E-19 |
| Eppk1 | orange | 0.55441606 | 0.01101541 | 0.61750816 | 0.00551244 |
| Gm19510 | yellow | 3.65235619 | 0.00620704 | 3.98658012 | 0.00997863 |
| Gm36107 | black | 4.59821256 | 0.00730069 | 2.7930739 | 0.06690972 |
| Gm32857 | skyblue | -3.3832442 | 0.00024425 | -3.0173992 | 0.00042109 |
| Gm2682 | orange | 1.62717473 | 7.72E-06 | 1.89801507 | 5.29E-07 |
| Gm48972 | black | 5.36338208 | 0.00034084 | 3.83740991 | 0.00719402 |
| AC160336.1 | skyblue | -3.6506971 | 1.33E-05 | -2.4358546 | 0.00476411 |
| Gm49092 | skyblue | -2.7267305 | 0.00892564 | -1.1737613 | 0.17492024 |
| AC113595.1 | yellow | 2.38102927 | 1.41E-15 | 2.45553026 | 5.32E-16 |
| AC110211.1 | yellow | 2.1868477 | 0.00599264 | 1.60656402 | 0.03715913 |
| AC119264.1 | yellow | 3.05984496 | 1.86E-09 | 3.38756999 | 1.00E-08 |
| AC102334.1 | skyblue | -2.1121876 | 0.00044043 | -1.9652785 | 0.00083408 |
| AL592187.2 | black | 5.95534189 | 4.53E-05 | 4.99499198 | 0.00098223 |
| AC157583.2 | red | -1.3608246 | 0.00106093 | -0.8588569 | 0.04838148 |
| AC132863.2 | orange | 1.90739016 | 0.00041111 | 0.96105432 | 0.11552136 |
| AC123724.2 | red | -1.2039947 | 0.0001516 | -1.0597591 | 0.00125356 |
| AC147041.1 | orange | 2.29485656 | 0.00010624 | 1.2706041 | 0.03105285 |
| AC154311.1 | skyblue | -2.1202439 | 0.00025621 | -1.165666 | 0.04497613 |
| AC139671.1 | orange | 1.30594081 | 5.76E-05 | 0.92333568 | 0.00509047 |
| AC163018.2 | orange | 2.53300986 | 0.00233617 | -0.0593201 | 0.95154216 |

| | | | | | |
|---|---|---|---|---|---|
| AL592187.7 | yellow | 3.83653883 | 0.00097221 | 4.00342217 | 0.00378464 |
| AC138177.4 | skyblue | -2.7995145 | 0.00911745 | -2.6738957 | 0.02318518 |
| AC123724.3 | yellow | 2.46795076 | 1.13E-13 | 1.9099381 | 2.50E-07 |
| Gm2808 | yellow | 3.76976661 | 4.10E-07 | 2.32952518 | 0.00025999 |
| AC126280.1 | skyblue | -1.1277784 | 0.06595928 | -1.7980013 | 0.004754 |
| AC129574.2 | yellow | 4.04171405 | 1.47E-05 | 1.62342673 | 0.04749579 |
| AC131339.2 | red | -0.8706361 | 0.03991216 | -1.5276986 | 0.00037207 |
| AC127341.3 | yellow | 2.40912697 | 5.41E-05 | 3.00149841 | 2.62E-06 |
| AC154478.1 | black | 3.02407417 | 0.01290094 | 5.77647434 | 0.00056179 |
| AC125141.3 | orange | 2.04806676 | 0.00819024 | 1.92079534 | 0.03903846 |
| AC161607.1 | skyblue | -1.5647635 | 1.87E-05 | -1.3256148 | 0.00033469 |
| AC117241.2 | black | 6.98196009 | 2.30E-05 | 6.40927253 | 0.0001693 |
| AC130815.3 | orange | 1.26724043 | 5.43E-07 | 0.73765151 | 0.00526903 |
